# Large library docking for novel SARS-CoV-2 main protease non-covalent and covalent inhibitors

**DOI:** 10.1101/2022.07.05.498881

**Authors:** Elissa A. Fink, Conner Bardine, Stefan Gahbauer, Isha Singh, Kris White, Shuo Gu, Xiaobo Wan, Beatrice Ary, Isabella Glenn, Joseph O’Connell, Henry O’Donnell, Pavla Fajtová, Jiankun Lyu, Seth Vigneron, Nicholas J. Young, Ivan S. Kondratov, Anthony J. O’Donoghue, Yurii Moroz, Jack Taunton, Adam R. Renslo, John J. Irwin, Adolfo García-Sastre, Brian K. Shoichet, Charles S. Craik

## Abstract

Antiviral therapeutics to treat SARS-CoV-2 are much desired for the on-going pandemic. A well-precedented viral enzyme is the main protease (MPro), which is now targeted by an approved drug and by several investigational drugs. With the inevitable liabilities of these new drugs, and facing viral resistance, there remains a call for new chemical scaffolds against MPro. We virtually docked 1.2 billion non-covalent and a new library of 6.5 million electrophilic molecules against the enzyme structure. From these, 29 non-covalent and 11 covalent inhibitors were identified in 37 series, the most potent having an IC_50_ of 29 μM and 20 μM, respectively. Several series were optimized, resulting in inhibitors active in the low micromolar range. Subsequent crystallography confirmed the docking predicted binding modes and may template further optimization. Together, these compounds reveal new chemotypes to aid in further discovery of MPro inhibitors for SARS-CoV-2 and other future coronaviruses.

## Introduction

SARS-CoV-2 encodes two cysteine proteases that have essential roles in hydrolyzing viral polyproteins into nonstructural proteins, enabling virus replication. The main protease (MPro, also known as 3CL protease) cleaves 11 different sites in viral polyproteins^1, 2^. While MPro is highly conserved across other coronaviruses such as SARS-CoV-1 and MERS, it has no close human homolog^3–5^. This makes it attractive for potential pan-coronavirus targeting, and for selective action.

The therapeutic potential of MPro inhibitors was substantiated by the approval of Paxlovid in December 2021. The treatment combines nirmatrelvir, which covalently inhibits MPro, with ritonavir, which slows nirmatrelvir’s metabolism^6^. Nirmatrelvir was optimized from PF-00835231, an inhibitor of the SARS-CoV-1 MPro developed in response to the 2002 SARS outbreak. Meanwhile, other potent MPro inhibitors are advancing through the drug development pipeline. Among them is the orally active MPro inhibitor S-217622^7^, which has entered clinical trials. Other inhibitors show much promise^4, 8–16^, including a non-covalent MPro inhibitor from the international Covid-19 Moonshot consortium that may be characterized as an advanced pre-clinical candidate^17–19^, and more experimental molecules that are relatively potent but have not proceeded far from hit to lead^20^.

Notwithstanding these successes, both the resistance that may be expected to emerge^21, 22^, and the inevitable liabilities of the early drugs support the discovery of new scaffolds. Accordingly, we targeted the structure of MPro for large library docking, seeking new starting points for lead discovery. Docking a library of over 1.2 billion “tangible” (make-on-demand) lead-like molecules and 6.5 million tangible electrophiles from Enamine *REAL* space (https://enamine.net/compound-collections/real-compounds) led to MPro inhibitors from 37 scaffolds, with affinities ranging from the low μM to 200 μM. Crystal structures for eight of the new inhibitors bound to MPro largely confirmed the docking predictions, while cell-based antiviral activity for two of the new inhibitors supports their further optimization (**Fig. 1**).

**Figure 1.**
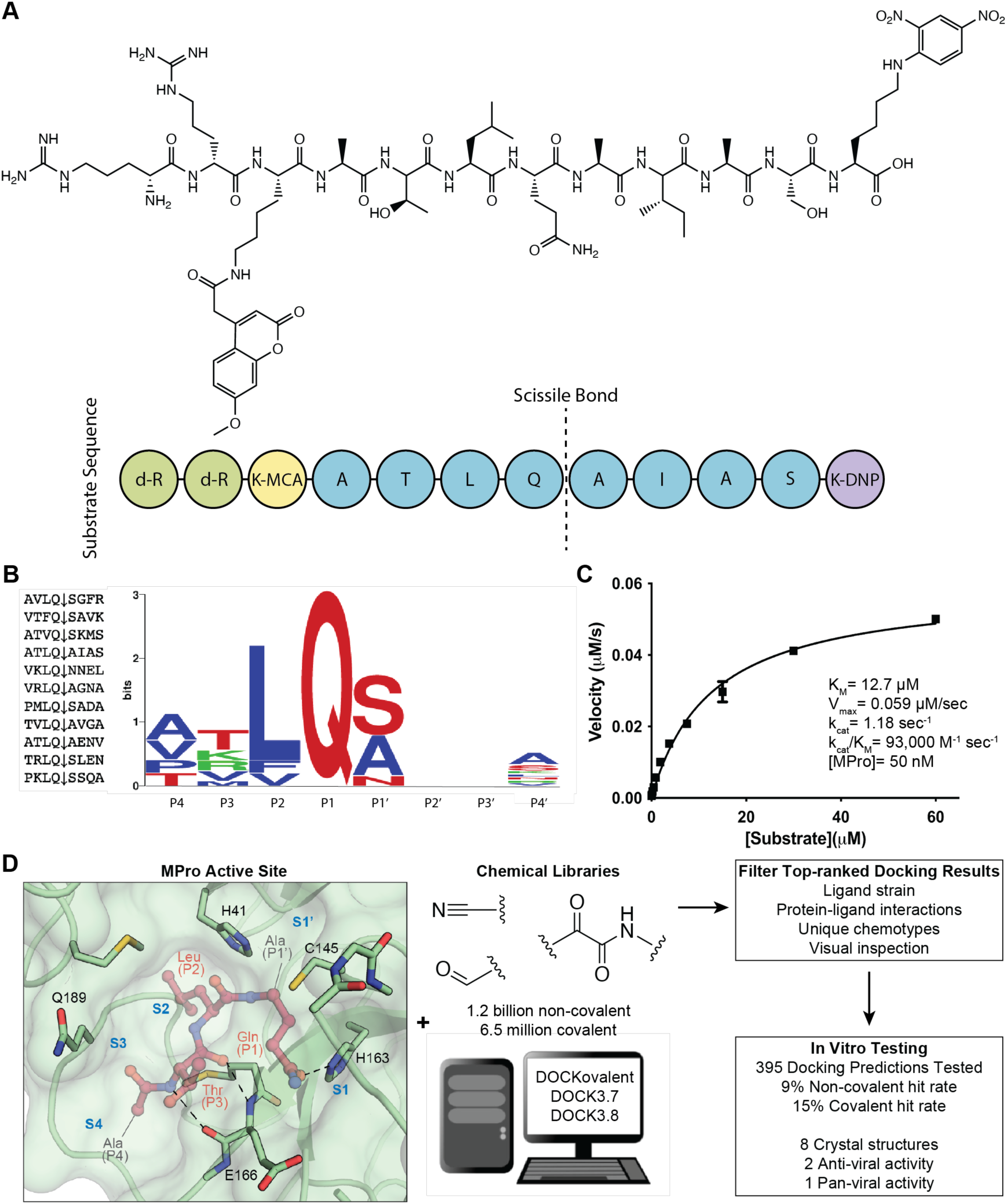
Substrate design and assay development allows structure-based inhibitor discovery. (**A**) The chemical structure of the optimized NSP7 substrate shown as a schematic (top) of the substrate sequence highlights the role of each residue (bottom). The substrate contains the P4-P4′ NSP7 extended substrate sequence (blue), the fluorophore (yellow), the fluorescent quencher (purple), and the residues for increasing solubility (green). (**B**) A list of the viral polypeptide NSP sequences (P4-P4′) that are cleaved by MPro (left). The sequenceLOGO highlighting the substrate specificity of MPro, yielding a P4-P4′ consensus sequence: ATLQ(S/A)XXA (right). (**C**) The Michaelis-Menten kinetics for the NSP7 substrate with MPro yield parameters indicative of an optimized, efficient substrate. (**D**) SARS-CoV-2 MPro active site (PDB 6Y2G)^26^ (green; sub-pockets S1′, S1, S2, S3, S4), shown here with substrate preferences (pink; P1′, P1, P2, P3, P4) (modeled after PDB 3SNE)^27^, was used to dock 1.2 billion non-covalent molecules and 6.5 million electrophile molecules. Top-ranked molecules were filtered and 395 were synthesized for *in vitro* testing. Some docking hits were prioritized for compound optimization, crystallography, pan-viral enzymatic activity, and cell-based antiviral activity. For C, experiments were performed in triplicate.

## Results

### Assay development and substrate design

MPro is the fifth nonstructural protein (Nsp5) encoded by SARS-CoV-2 and is a homodimeric cysteine protease with a catalytic diad comprised of Cys145 and His41. MPro has a P1 primary specificity determinant of glutamine and a preference for aliphatic residues in the P4 and P2 positions, while alanine and serine are preferred in the P1′ position^23^ (**Fig. 1D**). The catalytic cycle is typical of many cysteine proteases, with the catalytic Cys145 primed by proton transfer to His41 and formation of an acyl enzyme intermediate via nucleophilic attack of Cys145 at the scissile peptide carbonyl function. The thioester intermediate is then hydrolyzed by an attacking water to free the catalytic cysteine and initiate another catalytic cycle^24^.

Crucial to inhibitor testing was the design and synthesis of an optimal substrate, as was done previously for SARS CoV MPro^25^ (**Fig. 1**). The endogenous Nsp substrates of MPro were compiled and a consensus sequence was observed that closely matched the individual sequence of the Nsp7 cleavage site (ATLQAIAS) (**Fig. 1B**). This sequence was flanked with an N-terminal Lysine-MCA fluorophore and a C-terminal DNP-quencher. Noting the preference for nonpolar residues at multiple sites, we were concerned that this substrate would have low solubility. Accordingly, two D-Arginines were coupled to the N-terminal Lysine-MCA to increase solubility (**Fig. 1A**). This Nsp7-like substrate yielded a favorable K_m_ of 12 μM and a k_cat_/K_m_ of 93,000 M^-1^ s^-1^, 3.5-fold better than that of the commonly used commercial substrate (Nsp4: AVLQSGFR; k_cat_/K_m_ = 26,500 M^-1^ s^-1^)^2^; this substrate was used in all enzyme inhibition assays (**Fig. 1C**). The more efficient Nsp7-like substrate described here is readily synthesized and provides the field with an optimized MPro substrate.

In early proof-of-concept testing, we observed an intolerance of MPro activity to high concentrations of DMSO, which is introduced when evaluating inhibitors from (<10 mM) DMSO stocks (the sensitivity to DMSO may reflect oxidation of the catalytic cysteine). The increased solubility of the D-Arginine-modified substrate mitigated the DMSO effect by reducing the volume of DMSO needed in substrate alliquots. In addition, we found that ethanol and acetonitrile were better tolerated by the enzyme, though these solvents have issues with volatility (**Fig. S1A**). These observations highlight the importance of controlling and minimizing compound solvent concentrations for MPro activity assays and provide alternatives solvents when DMSO is not suitable for *in vitro* biochemical assays. We also found that small amounts of non-ionic detergent were crucial for retaining Mpro activity in our *in vitro* assays. Removing the 0.05% Tween-20 we used in our assays resulted in no observed substrate cleavage. Activity was then recovered by increasing addition of bovine serum albumin (BSA), highlighting the need of detergent or enzyme stabilizing additives **(Fig. S1B)**. We tested three previously reported compounds under our assay conditions. The covalent inhibitor nirmatrelvir had a similar IC_50_ as the reported potency^6^, whereas two non-covalent inhibitors (PET-UNK-29afea89-2 and VLA-UCB-1dbca3b4-15) had measured IC_50_ values 2 to 5-fold higher compared to reported values^17^, likely due to different substrate and substrate concentrations used in the published assays and those used here (**Table S1**). These rates provide a reference for comparing the different inhibitors.

### Non-covalent docking screen and compound optimization for MPro inhibitors

Seeking new inhibitors, we began with a SARS-CoV-2 MPro crystal structure in complex with an α-ketoamide covalent inhibitor (PDB 6Y2G)^26^. To define hot-spots for ligand docking in the active site, we modeled a complex of SARS-CoV-2 MPro bound to a non-covalent SARS-CoV MPro inhibitor (PubChem SID87915542)^28^ (non-covalent inhibitor complex crystal structures of the enzyme from SARS-CoV-2 were at that time unavailable). The crystal structure of the non-covalently ligated SARS-CoV MPro (PDB 3V3M)^28^ was structurally aligned onto the SARS-CoV-2 structure, the atomic coordinates of the α-ketoamide inhibitor were replaced with those of the non-covalent SARS-CoV MPro inhibitor SID87915542 (IC**_50_** = 5 µM)^28^ and the complex was energy-minimized (Methods). After calibration of the docking parameters^29^ (Methods), approximately 225 million neutral molecules, mainly from the lead-like subset of the ZINC15 library^30^ (molecular weight (MWT) ranging from 250-350 amu and clogP ≤4.5) were docked against MPro. Another 110 million molecules with 350 < MWT ≥ 500 were docked in a separate screen. Docked molecules were filtered for intramolecular strain^31^ and selected for their ability to hydrogen bond with Gly143, His163, or Glu166, and make favorable non-polar contacts with Met49 and Asp187. Ultimately, 220 molecules were prioritized, of which 194 (88%) were successfully synthesized by Enamine. In a primary screen, compounds were tested at a single concentration of 100 µM using the fluorescence-based substrate cleavage assay and 19 showed >30% inhibition of enzyme activity and were prioritized for full concentration-response curves. Overall, 12 molecules were defined as hits, with IC_50_ values ≤ 300 μM, for an overall hit rate of 6% (12 hits/194 molecules tested); potencies ranged from 97 to 291 µM (**Table 1, Table S1, Fig. S2.1, Fig. S2.2**).

**Table 1.**
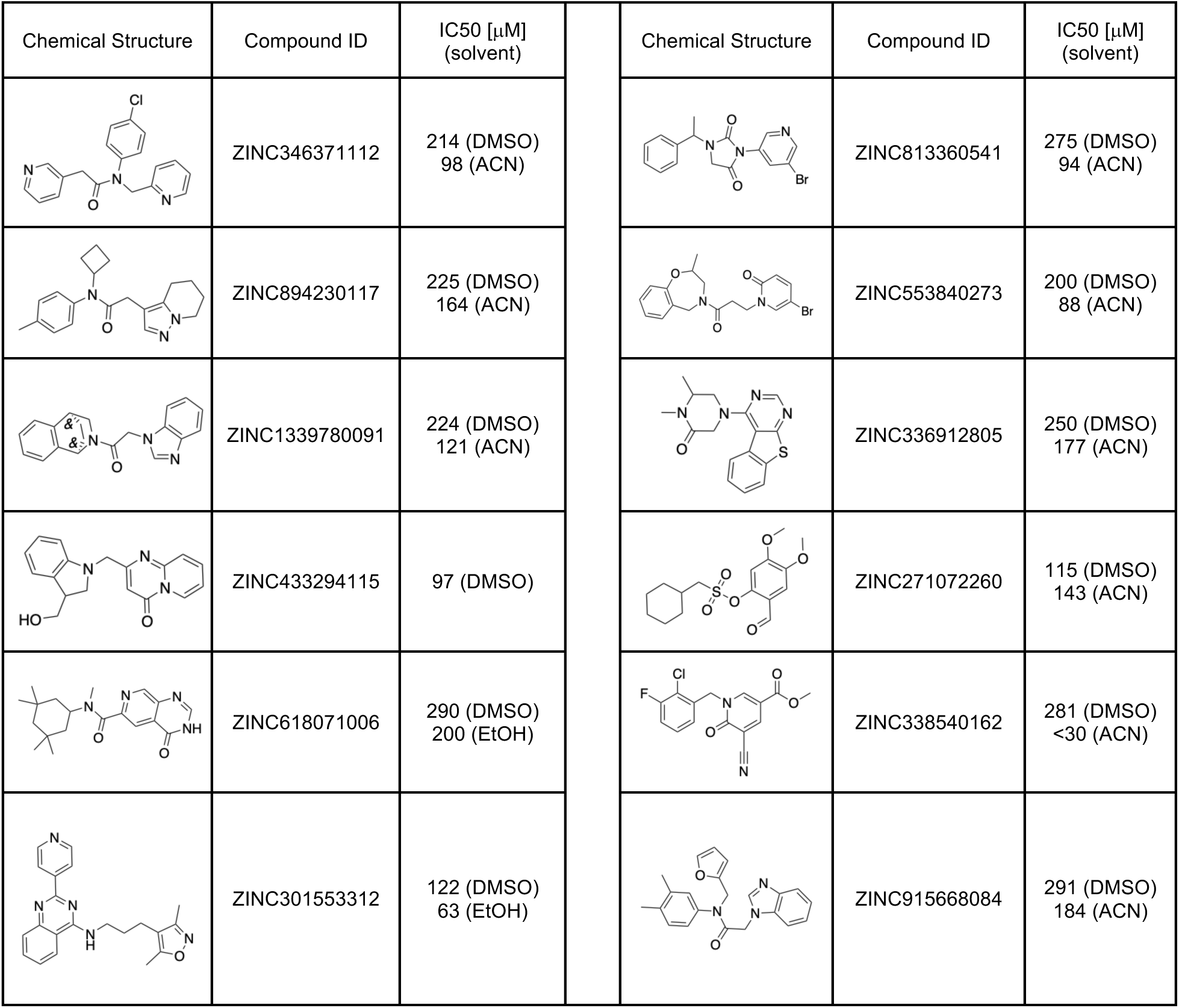
Hits from the first non-covalent docking screen.

| Chemical Structure | Compound ID | IC<sub>50</sub> [μM]<br>(solvent) | Chemical Structure | Compound ID | IC<sub>50</sub> [μM]<br>(solvent) |
| --- | --- | --- | --- | --- | --- |
|  | ZINC346371112 | 214 (DMSO)<br>98 (ACN) |  | ZINC813360541 | 275 (DMSO)<br>94 (ACN) |
|  | ZINC894230117 | 225 (DMSO)<br>164 (ACN) |  | ZINC553840273 | 200 (DMSO)<br>88 (ACN) |
|  | ZINC1339780091 | 224 (DMSO)<br>121 (ACN) |  | ZINC336912805 | 250 (DMSO)<br>177 (ACN) |
|  | ZINC433294115 | 97 (DMSO) |  | ZINC271072260 | 115 (DMSO)<br>143 (ACN) |
|  | ZINC618071006 | 290 (DMSO)<br>200 (EtOH) |  | ZINC338540162 | 281 (DMSO)<br><30 (ACN) |
|  | ZINC301553312 | 122 (DMSO)<br>63 (EtOH) |  | ZINC915668084 | 291 (DMSO)<br>184 (ACN) |

As mentioned above, DMSO was observed to lower enzyme activity, consequently the actives, initially tested from 10 mM DMSO stocks, were re-tested against MPro from 30 mM acetonitrile (ACN) or ethanol (EtOH) stocks. Eleven compounds showed clear dose-response with IC_50_ values ranging from 30 to 200 μM. Although covalent docking was not employed in this campaign, we noted three initial docking hits (**ZINC338540162**: IC_50_[ACN] = 30 μM, **ZINC271072260**: IC_50_[ACN] = 143 μM and **ZINC795258204**: IC_50_[DMSO] = 177 μM) could, in principle, inhibit MPro covalently as they contain warheads (nitrile, aldehyde) known to react with catalytic cysteines. Several initial docking hits were tested for colloidal aggregation using dynamic light scattering (DLS) and off-target counter screens against malate dehydrogenase (MDH) and AmpC β-lactamase^32, 33^ (**Fig. S3**). In DLS experiments, some scattering higher than 10^6^ is observed indicating potential aggregation. While a few compounds e.g., **‘3312** showed unspecific inhibition of MDH, off-target activities were reversed by addition of 0.01% Triton X-100. As the MPro enzymatic assay is run with 0.05% Tween-20, an even stronger disruptor of colloidal aggregation than 0.01% Triton-X 100, we deemed the weak aggregation of these compounds not relevant to their activity on Mpro.

We focused on four initial hits (**ZINC346371112**: IC_50_[ACN] = 98 μM, **ZINC301553312**: IC_50_[EtOH] = 63 μM, **ZINC813360541**: IC_50_[ACN] = 90 μM and **ZINC553840273**: IC_50_[ACN] = 88 μM) for structure-based optimization. We used the SmallWorld search engine (NextMove Software, Cambridge UK)^34^ to identify purchasable analogs of these inhibitors within a 12 billion compound version of the REAL library (https://enamine.net/compound-collections/real-compounds/real-space-navigator), docking each analog into the MPro structure to assess complementarity. Between 10-20 analogs of each of the four inhibitors were selected for testing in the initial round of optimization (**Fig. 2, Table S1**). For two initial hits, **‘0541** and **‘0273**, more potent analogs were identified in two to three rounds of this analog-by-catalog approach (**Table S1**). The **‘0273** analogs **Z4924562413** and **Z4946671001** had IC_50_ values of 13 µM and 5 µM, respectively (**Fig. 2A**). Analogs of the initial docking hit **‘0541**, such as **Z4929615577** and **Z4929616137**, reached similar potencies of 10 µM and 8 µM, respectively (**Fig. 2G**).

**Figure 2.**
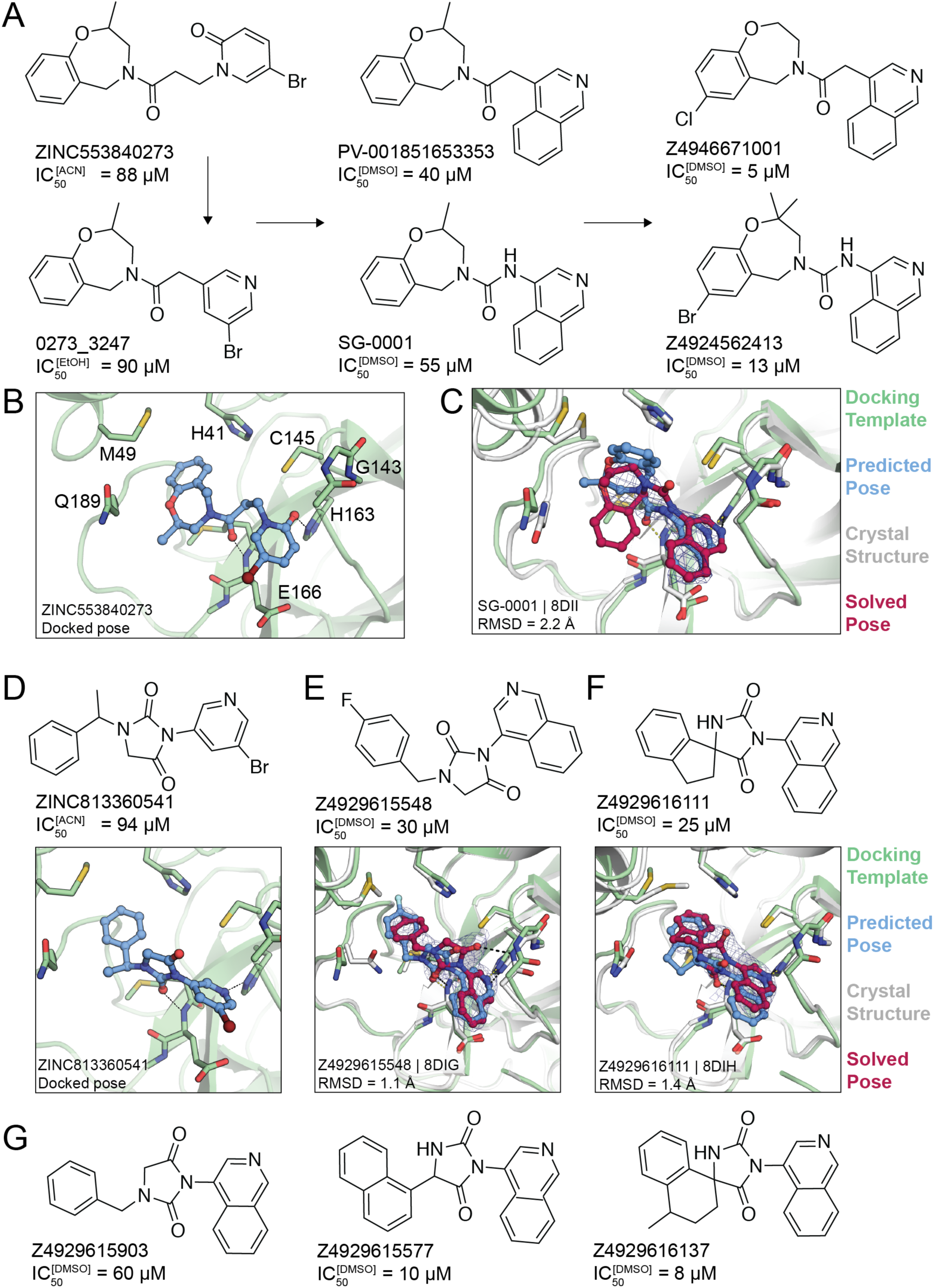
Non-covalent compound optimization to low-μM potencies. (**A**) Progression of the **‘0273** scaffold. (**B**) Predicted binding pose of **‘0273**. (**C**) Comparison of crystal structure (grey protein, red compound) and docked complex (green protein, blue compound) of **SG-0001** (PDB 8DII). (**D**) Predicted binding pose of **‘0541**. (**E**), (**F**) Comparison of crystal structures and docked complexes of **‘5548** (PDB 8DIG) and **‘6111** (PDB 8DIH), respectively. (**G**) Additional **‘0541** analogs with improved affinities. The 2fo-fc ligand density maps (blue contour) are shown at 1 α. Hungarian root mean square deviations (RMSD) were calculated with DOCK6.

### Crystal structures of the non-covalent inhibitors

To investigate how the docked poses of the new inhibitors corresponded to their true binding modes, and to inform further optimization, crystal structures of three of the optimized non-covalent inhibitors were determined with resolutions ranging from 2.12 Å to 2.59 Å. For the **‘0237** analog, **SG-0001** (IC_50_ = 55 µM, **Fig. 2A-C**), the crystal structure revealed only moderate density for the ligand. Still, the predicted binding pose compared well with the experimentally determined pose, with a Hungarian (symmetry corrected) root mean square deviation (RMSD) of 2.2 Å. The isoquinoline group of **SG-0001** is inserted in the S1 subpocket, hydrogen-bonding with His163; this was also predicted for the pyridone carbonyl in the parent molecule **‘0273** (**Fig. 2B,C, Fig. S4**). However, the tetrahydrobenzoxazepine ring, predicted to bind in the S2 subpocket in **‘0237**, appeared much less buried in the **SG-0001** experimental structure. The crystal structure of MPro in complex with the **‘0541** analog **‘5548** superimposed with high fidelity to the docking-predicted pose, with an RMSD of 1.1 Å (**Fig. 2E, Fig. S4**). Here, the compound’s hydantoin core hydrogen bonds with the backbone amine of Glu166 and Gly143. In addition, the crystal structure of MPro in complex with **‘6111** confirms the predicted biding pose (RMSD = 1.4 Å) with the isoquinoline placed in the S1 subpocket and the hydrophobic spirocyclic indane group occupying the S2 pocket (**Fig. 2F, Fig. S4**).

### A second docking screen for non-covalent inhibitors of MPro

Throughout the duration of this work, many groups have identified potent inhibitors with extensive structure-activity-relationship (SAR) data, with scaffolds resembling our own^20^. We therefore thought to perform a second docking campaign. Here, we tried to incorporate insights emerging from our own results and those from other studies (Methods) emphasizing the discovery of novel chemotypes.

The new docking screen targeted the SARS-CoV-2 MPro crystal structure in complex with MAT-POS-b3e365b9-1 (MPro-x11612.pdb)^17^, a non-covalent ligand reported by the COVID-19 Moonshot consortium. Compared to the previous docking template (PDB 6Y2G), the MAT-POS-b3e365b9-1-bound site is slightly smaller, with the 2-turn alpha helix between Thr45 and Leu50, and the loop between Arg188 and Ala191 shifted inwards by roughly 2 Å, constricting the shape of the P2 sub-pocket. After calibration of docking parameters, ensuring the model prioritizes 15 previously reported MPro inhibitors against different decoy sets^29, 35^, we used the ZINC-22 library (https://cartblanche22.docking.org/) to dock 862 million neutral compounds with 18-29 non-hydrogen atoms from the Enamine *REAL* database (Methods).

The high-ranking docked molecules were filtered for novelty by removing those with ECFP4-based Tanimoto coefficients (Tc) greater than 0.35 to 1,716 SARS-CoV-2 MPro inhibitors (Methods). Roughly 9,500 of these were graphically evaluated for favorable contacts, and 146 compounds were de novo synthesized by Enamine Ltd. Of these, 17 inhibited MPro with IC_50_ values ≤ 200 μM (**Table 2, Fig. S2.3, Fig. S2.4**) for a hit rate of 12% (17 hits/146 tested). To our knowledge, none the new actives fell into scaffolds that have been previously reported for MPro. Compared to the first docking screen, several initial hits from the second screen showed slightly higher activity, such as **Z3535317212**, with an IC_50_ value of 29 μM. For **‘7212**, the docked pose suggests hydrogen bonds between the compound’s dihydrouracil core and Glu166 as well as Gly143, in addition to hydrogen bonds between the compound’s pyridinol group (**Fig. S2.3**). Five docking hits (**Z5420225795**: IC_50_ = 40 μM, **Z1669286714**: IC_50_ = 110 μM, **Z1355254448**: IC_50_ = 110 μM, **ZINC5420738300**: IC_50_ = 160 μM, **Z2195811405**: IC_50_ ∼200 μM) share a common ketoamide functional group predicted to form one hydrogen bond to Glu166, however, we note that ketoamide might also inhibit MPro through covalent linkage to Cys145. As in the first docking campaign, hits were tested for colloidal aggregation. A few compounds (**‘7900, ‘8488, ‘1405, ‘8300**) had higher DLS scattering or caused >50% inhibition of MDH in the absence of detergent, which was reversed by 0.01% Triton X-100 (**Fig. S3**). We therefore conclude that the measured activities of those compounds at MPro, in presence of 0.05% Tween-20, originate from specific on-target actions, but care should be taken when using related scaffolds in detergent-free experiments.

**Table 2.**
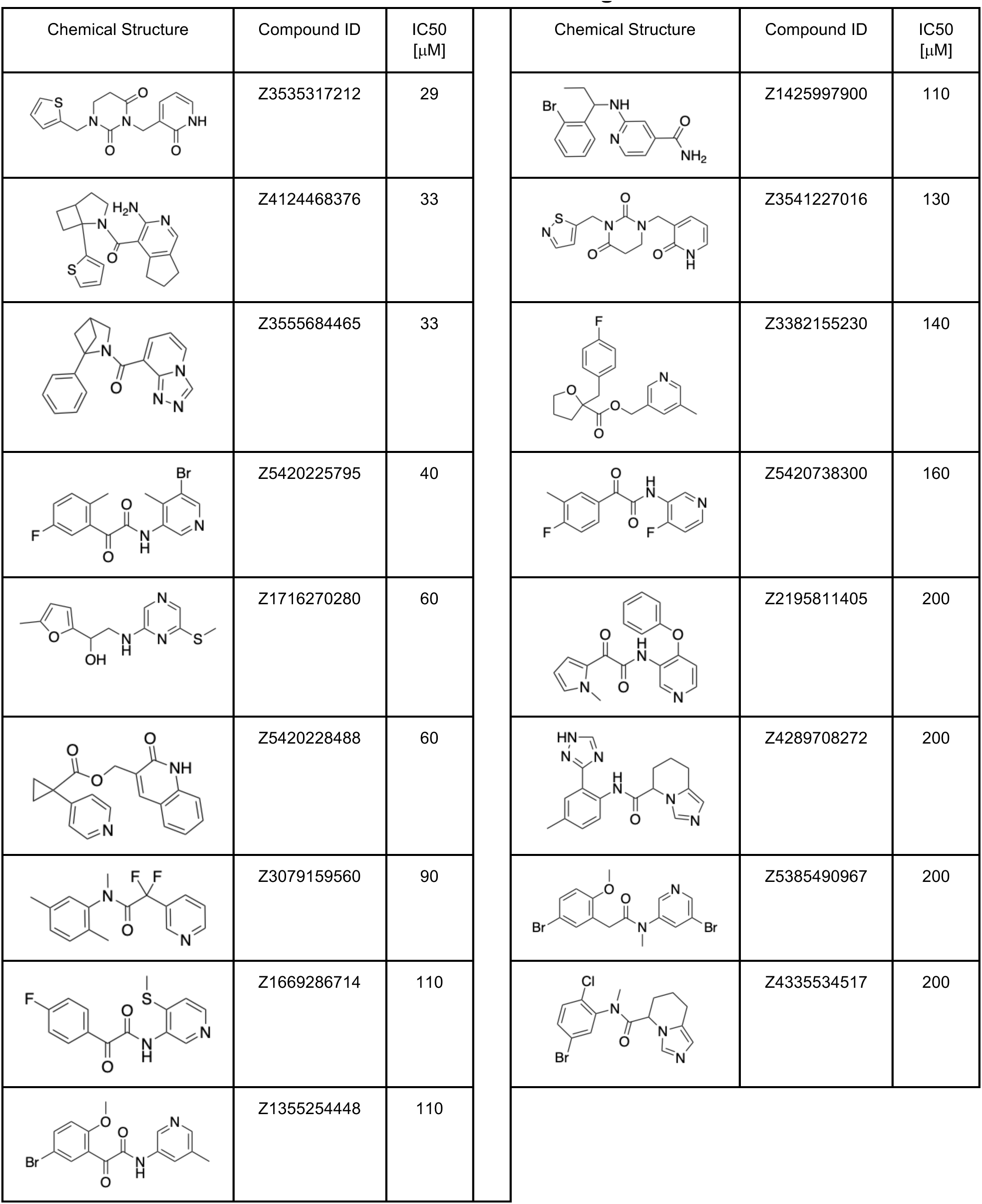
Hits from the second non-covalent docking screen.

| Chemical Structure | Compound ID | IC50<br>[μM] | Chemical Structure | Compound ID | IC50<br>[μM] |
| --- | --- | --- | --- | --- | --- |
|  | Z3535317212 | 29 |  | Z1425997900 | 110 |
|  | Z4124468376 | 33 |  | Z3541227016 | 130 |
|  | Z3555684465 | 33 |  | Z3382155230 | 140 |
|  | Z5420225795 | 40 |  | Z5420738300 | 160 |
|  | Z1716270280 | 60 |  | Z2195811405 | 200 |
|  | Z5420228488 | 60 |  | Z4289708272 | 200 |
|  | Z3079159560 | 90 |  | Z5385490967 | 200 |
|  | Z1669286714 | 110 |  | Z4335534517 | 200 |
|  | Z1355254448 | 110 |  |  |  |

Taken together, the actives from this campaign explored ten different scaffold classes with IC_50_ values better than 150 μM. These scaffolds represent new points of departure for MPro inhibitor discovery.

### A covalent docking screen targeting MPro Cys145

Seeking electrophiles that could covalently modify the catalytic Cys145, we searched the 1.4 billion molecules in the ZINC15/ZINC20^30, 34^ databases for three Cys-reactive covalent warheads, aldehydes, nitriles, and α-ketoamides. Dockable 3D molecules were built for covalent docking with DOCKovalent^36, 37^ (Methods). The molecules and their DOCKovalent files for the final 6.5 million molecules are available at http://covalent2022.docking.org.

We then virtually screened a total of 3.6 million nitriles, 1.5 million aldehydes, and 1.4 million α-ketoamides against MPro (PDB 6Y2G)^26^. The top-ranked molecules were filtered for torsional strain^31^, for favorable enzyme interactions, and clustered for chemical diversity using an ECFP4-based best first clustering algorithm (Methods). Remaining molecules were visually prioritized for favorable interactions with His41, Cys145, Gly143, Thr26, or Glu166. Ultimately, 35 aldehydes, 41 nitriles, and 21 α-ketoamides were selected for synthesis, of which 27, 31, 16, respectively, were successfully synthesized and tested for activity against MPro (**Table S1**). Those compounds with single-point percent inhibition >50% at 100 μM were prioritized for generation of full concentration-dose-response curves.

Defining actives as molecules with IC_50_ ≤ 150 μM, the hit rate for covalent docking was 15% (11 actives/74 compounds tested); the most potent had an IC_50_ of 20 μM (**Fig. 3, Fig. S5**). Eight others had IC_50_ values 25 to 100 μM. Initial nitriles and aldehyde docking hits had activities as low as 20 μM in compound ‘**5103**, and 55 μM in compound ‘**3620**, respectively. The α-ketoamide docking screens resulted in two compounds (**‘4072, ‘6634**) with inhibition ≥ 200 μM and were not considered actives (**Table S1**). Initial docking hits were evaluated for potential MPro inhibition through colloidal aggregation as previously described for non-covalent docking hits (**Fig. S3**). Some higher DLS scattering or non-specific inhibition is observed in the AmpC and MDH enzymatic assays. However, adding 0.01% Triton X-100 in the MDH inhibition assay largely recovered enzymatic activity and eliminated any non-specific inhibition thereby suggesting that the measured activities in the detergent-containing MPro enzymatic assays are not caused by aggregation (something also confirmed by subsequent crystallography, see below).

**Figure 3.**
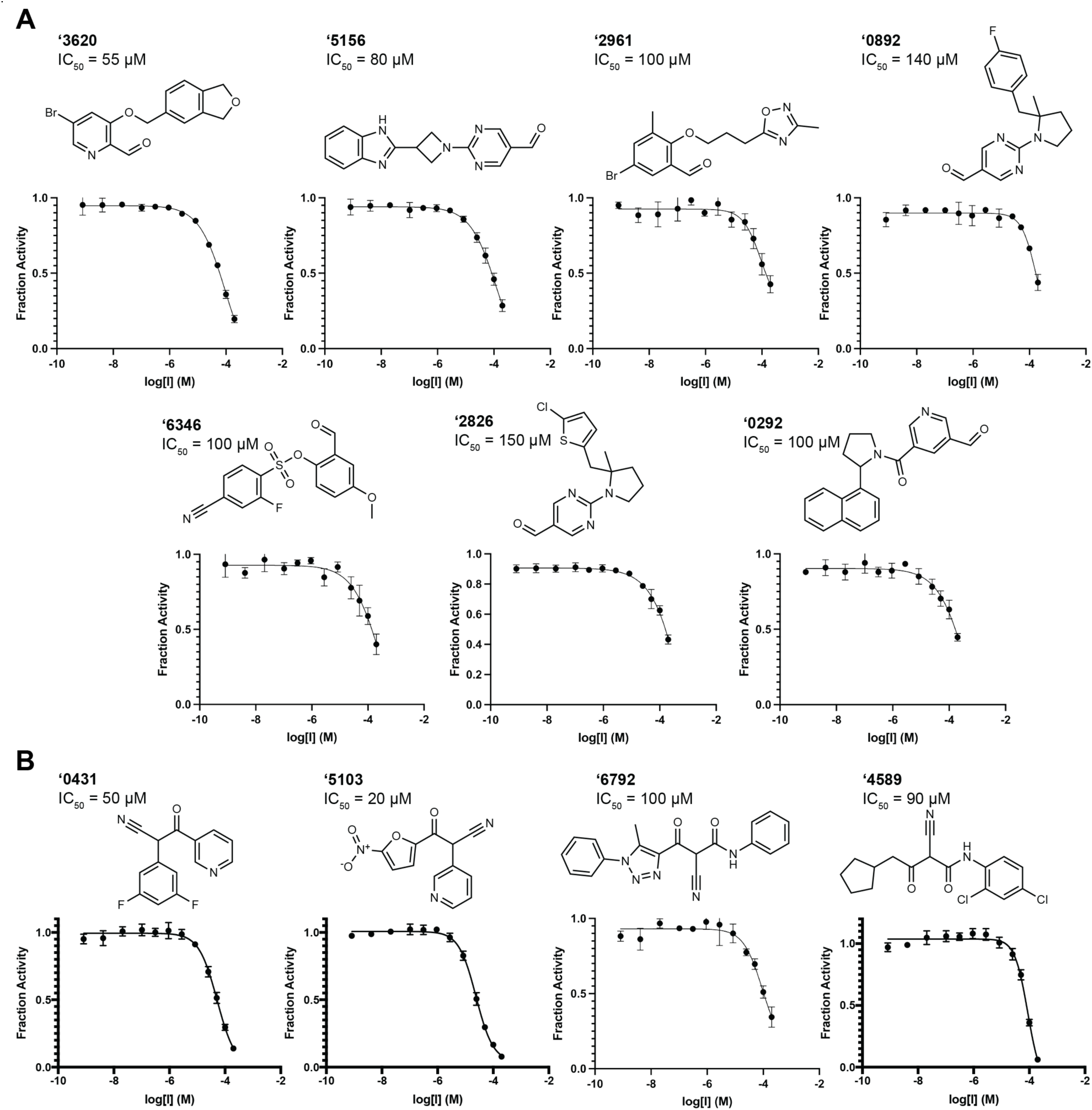
Covalent hits from 6.5 million virtual screen. Dose response curves for (**A**) aldehyde and (**B**) nitrile docking hits. IC_50_ values shown. All measurements done in triplicate.

The covalent inhibitors had diverse chemotype and their docked poses explored different enzyme sub-pockets (**Fig. 1, Fig. 3, Fig. S5**). In the S1′ pocket, hydrophobic interactions were made by compounds **‘3620**, **‘6345**, **‘6792** in their docked poses. Hydrogen bonding with His163 in the S1 pocket was made by **‘5103**, **‘0431**, **‘2961** in their docked poses. Several compounds, such as **‘0892** and **‘0292**, occupied the S2 and S3 pockets, making non-polar interactions with Met49 and Phe181. Other compounds appeared to span the binding site between the S1 and S2/S3 pockets, e.g. **‘5156** hydrogen-bonding with Glu166. Many compounds, such as **‘3620** and **‘6792**, formed hydrogen-bonds with the peptide backbone atoms of Cys145, Ser144 and Gly143.

We sought to optimize several of the new covalent inhibitors, focusing on the aldehyde **‘3620** with an IC_50_ of 55 μM (**Table S1**). These analogs were identified through multiple strategies, including simply seeking readily available “make-on-demand” congeners that fit in the enzyme site, using SmallWorld and Arthor (NextMove Software, Cambridge, UK)^34^, or testing perturbations to what seemed to be key interactions. From these studies emerged 39 analogs with IC_50_ values better than **‘3620**. The most potent analog **‘7021** had an IC_50_ of 1 μM and was observed to act as a reversible inhibitor (**Fig. S6**). Other analogs ranging from 2 to 48 μM had changes to different benzene substituents or bicyclic systems of **‘3620** (**Fig. 4**, **Table S2, Fig. S7**).

**Figure 4.**
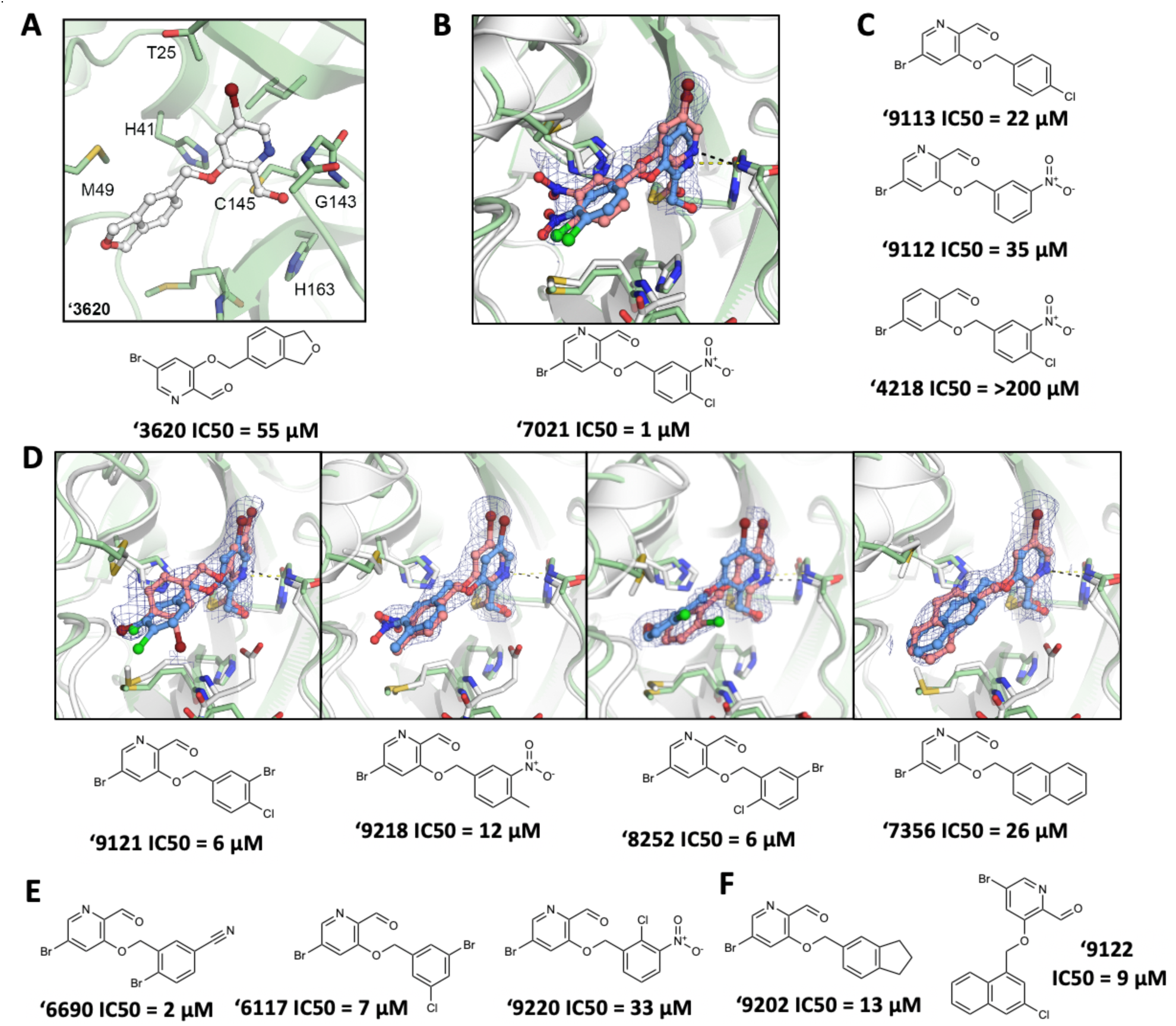
Compound optimization of aldehyde ‘3620. (**A**) Docked pose of docking hit **‘3620**. (**B**) Crystal structure (pink carbons) and docked pose (blue carbons) comparison for analog **‘7021** (RMSD 1.29 Å; PDB 8DIB). (**C**) Hypothesis testing analogs of **‘7021** included removing the nitro in **‘9113** and the chlorine in **‘9112**, both with weaker inhibition. Analog **‘4218** replaced the pyridine with a benzene eliminating inhibition. (**D**) Crystal structures of additional **‘3620** analogs comparing experimental (pink carbons) and docked (blue carbons) poses (RMSDs of 1.75 Å, 0.78 Å, 1.18 Å, and 0.84 Å, respectively; PDB 8DIC, 8DIE, 8DID, 8DIF, respectively). (**E**) Analogs with different benzene substituent orientations (**‘6690**, **‘6117**) inhibit MPro at similar potencies. Substituents oriented like **‘9220** were weaker inhibitors. (**F**) Examples of the most potent larger hydrophobic analogs of **‘3620**. For A-F, Mpro protein structure is PDB 6Y2G (green carbons) used in docking or from the solved structures (white carbons). Hydrogen bonds shown with dashed lines. The 2fo-fc ligand density maps (blue contour) are shown at 1 Λ. IC_50_ values are shown with concentration response curves in **Fig. S7**. All measurements done in triplicate.

In its docked pose, the pyridine nitrogen of **‘7021** hydrogen bonds to Gly143 (**Fig 4B**). To test the importance of this interaction, the phenyl analog of the pyridine, compound **‘4218**, was synthesized and tested. This molecule lost all measurable activity (IC_50_ > 200 μM), consistent with the importance of the pyridine hydrogen bonds (**Fig. 4C**). However, it is also likely that the more electro-deficient pyridine ring makes the aldehyde more reactive towards the catalytic Cys145. Meanwhile, removing non-polar groups from the distal phenyl ring of **‘7021,** as in analogs **‘9313** and **‘9112**, increased IC_50_ values to 22 μM and 35 μM, respectively, indicating more hydrophobic bulk was preferred in the shallow subsite in which this substituted phenyl was docked.

### Crystal structures of the covalent inhibitors

To investigate how the docked poses of the covalent inhibitors corresponded to true binding modes, and to aid further optimization, crystal structures of five aldehyde inhibitors complexed with MPro were determined: **‘7021** (IC_50_ = 1 μM), **‘9121** (IC_50_ = 6 μM), **‘8252** (IC_50_ = 6 μM), **‘9218** (IC_50_ = 12 μM), and **‘7356** (IC_50_ = 26 μM), with resolutions ranging from 1.90 Å to 2.17 Å (**Fig. 4B, Fig. 4D, Fig. S8**). The structures of these compounds recapitulated the docking predictions with high fidelity, with all-atom Hungarian RMSD values ranging from 0.78 Å to 1.75 Å (**Fig. 4B**). Consistent with the docking, and with the results of the analogs, the pyridine nitrogen in each inhibitor hydrogen bonds with Gly143 and the thioacetal adduct hydrogen bonds with the backbone of Cys145 in the oxyanion hole of the enzyme. The hydrophobic groups on the distal aryl ring interact with residues in the S2/S3 pockets, including Met49 and Phe181 (**Fig. 4B, Fig. 4D, Fig. S8**).

### Lead inhibitors are antiviral with pan-coronaviral MPro inhibition

With the progression of covalent and non-covalent inhibitor optimization, we tested several compounds in an RT-qPCR viral infectivity assay in HeLa-ACE2 cells. Compounds ‘**7021** and ‘**7356** had antiviral IC_50_ values of 6.2 **μ**M and 19.5 μM, respectively, directionally consistent with their *in vitro* IC_50_ values of 1 μM and 26 μM (**Fig. 5A, Table S3**). Meanwhile, no measurable antiviral activity was observed for the covalent aldehyde ‘**6690**, the covalent nitrile ‘**5103**, and the non-covalent compound ‘**6137**, with *in vitro* IC_50_ values of 2 μM, 20 μM, and 8 μM, respectively. What separates the antiviral actives from the inactives remains unclear. We also tested ‘**7021** for its ability to inhibit MPro of other coronaviruses, SARS-CoV-1 and MERS (**Fig. S9**). **‘7021** inhibited the SARS-CoV-1 MPro with an IC_50_ of 8 μΜ, similar to its SARS-CoV-2 MPro IC_50_ of 1 μM, however it was a weaker inhibitor for the MERS MPro with an IC_50_ of 50 μM (**Fig. 5B, Table S4**).

**Figure 5.**
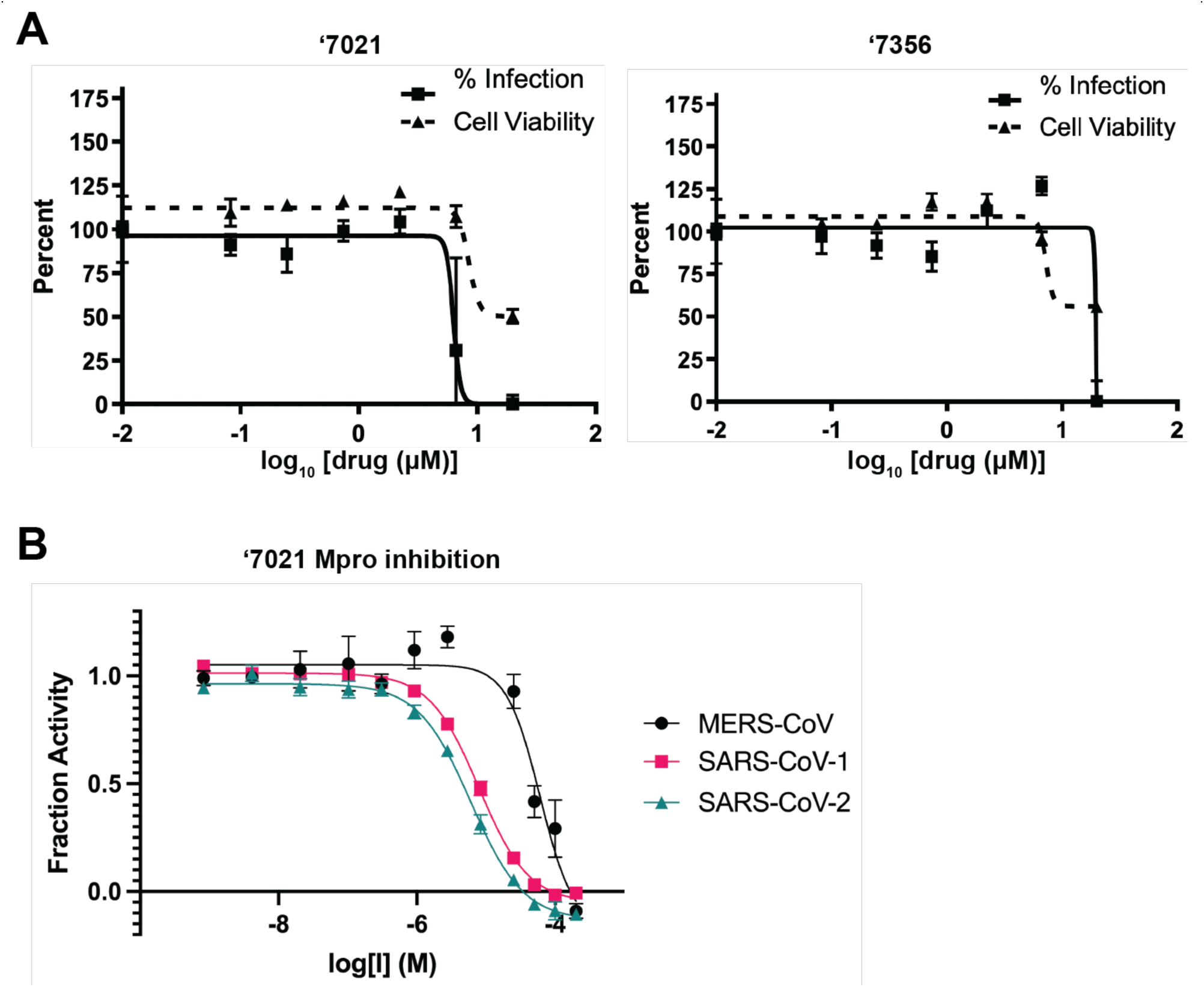
Antiviral activity and pan-coronaviral MPro inhibition by covalent analogs. (**A**) **‘7021** and **‘7356** inhibit SARS-CoV-2 infectivity with minimal impacts on cell viability. (**B**) **‘7021** also inhibits SARS-CoV-1 and MERS-CoV MPro. All measurements done in triplicate.

## Discussion

From this study emerged 132 MPro inhibitors with IC_50_ values less than 150 μM, covering 37 different scaffold classes (**Fig. 3, Table 1, Table 2**). Of these, 15 inhibitors in 3 scaffolds inhibited the enzyme with IC_50_ values less than 10 μM. The best covalent inhibitor, **’7021**, was confirmed to act reversibly (**Fig. S6**), likely reflecting the fast-on/fast-off kinetics characteristic of aldehyde covalent inhibitors. We also present an optimized MPro substrate for future inhibitor characterization (**Fig. 1**). To dock the electrophile library, we first had to create it, drawing on aldehydes, nitriles, and α-ketoamides in the expanding library of tangible molecules. This resulted in a library of over 6.5 million new electrophiles, which is openly available to the community at https://covalent2022.docking.org. Crystal structures of eight of the new inhibitors closely corresponded to the docking predictions (**Fig. 3, Fig. 4**). Two of the new aldehyde inhibitors had antiviral activities close to those of their enzymatic IC_50_ values, suggesting that further optimization of this class for on-enzyme potency may presage antiviral activity (**Fig. 5A**).

While the strengths of this study were the identification of multiple new MPro inhibitor scaffolds, with subsequent crystal structures supporting the docking predictions, the work also revealed liabilities of docking screens. In contrast to campaigns against G protein-coupled receptors^38–41^ and other integral membrane receptors^42, 43^, hit rates against MPro were in the 7 to 15% range, rather than the 25 to 60% range. Meanwhile, the activities of the better MPro docking hits were in the 20 to 100 μM range, not the low-to mid-nM range found against the integral membrane proteins. Here, the MPro docking campaigns resemble those against other soluble proteins such as β-lactamase^39, 44^ and the macrodomain of SARS-CoV-2^45, 46^, or even against allosteric sites in GPCRs^47, 48^. For physics-based scoring functions like that used in DOCK3.7, well-enclosed binding sites characterized by a key polar interaction are more amenable to ligand discovery than the more open, flatter sites such as those in MPro. This both reflects the physical characteristics of the site and the sorts of molecules in our docking libraries, where chemical space coverage is best in the fragment and lead-like ranges and erodes among larger and more hydrophobic molecules better suited to sites like MPro’s.

These caveats should not distract from the key observations of this study. Large library docking of both lead-like molecules and covalent electrophiles has revealed 11 scaffold families of MPro inhibitors (**Fig. 3, Table 1, Table 2**), the best of which act in the low μM range (**Fig. 2, Fig. 4**). Whereas neither hit rates nor affinities rose to levels seen against targets with well-defined binding sites, eight crystal structure of characteristic lead molecules confirmed the docking poses (**Fig. 2, Fig. 4**), suggesting that, notwithstanding the lower hit rates, when the docking was right it was right for the right reasons. These structures may template the further optimization of these new MPro inhibitors, several of which show initial antiviral activity against the virus.

## Methods

### Protein purification

E. coli strains and bacterial expression plasmids All bacterial expression plasmids were transformed into One Shot™ BL21(DE3)pLysS Chemically Competent E. coli (Thermo). We received four plasmids from Ursula Schulze-Gahmen at the Gladstone Institute encoding non-structural protein 5 (nsp5). Nsp5 codon optimized for E. coli was synthesized (Genscript). Nsp5 was cloned into pET His6 GST TEV LIC cloning vector (2G-T) (Addgene 29707), pET His6 TEV LIC cloning vector (2B-T) (Addgene 29666), pET His6 MBP TEV LIC cloning vector (2M-T) (Addgene 29708), pET His6 Sumo TEV LIC cloning vector (2S-T) (Addgene 29711). We received nsp5 cloned into pGEX6p-1with a N-terminal GST tag and Mpro cleavage-site SAVLQ↓SGFRK from Rolf Hilgenfeld. Nsp5-BA is Mpro cloned into pET28a(+) with a C-terminal TEV cleavage site and kanamycin resistance (TWIST).

### Expression and purification of MPro

The expression for MPro in E. coli was previously described^26^. In brief a transformed clone of BL21(DE3)pLysS E. coli was added to a 50 mL culture of 2xYT media supplemented with 2% glucose and ampicillin grown overnight at 37°C. 30 mL of overnight cultures were used to inoculate 1 L of 2xYT media ampicillin or appropriate antibiotic. A large 1 L culture was shaken at 225 rpm at 37°C. The 1 L culture was induced when culture OD600 reached 0.8 (after ∼3 hours) by adding 1 mL of 1 M IPTG. After 5 hours of expression at 37°C the culture was centrifuged at 9,000 rpm for 15 min. Supernatant was discarded and cell pellet stored at −80°C. The frozen cell pellet was thawed on ice in 30 mL of 20 mM TRIS 150 mM NaCl pH 7.4 buffer. The resuspended sample was sonicated for 5 mins or until lysis was complete. Sonicated cell lysate was centrifuged at 15000 rpm for 30 mins. 3 mL of Ni-NTA beads were incubated with 57 the supernatant for 1 hour at 4°C. Beads were centrifuged at 200 rpm for 2 mins. Supernatant was stored at 4°C. Ni-NTA beads were washed with ∼3 column volumes of wash buffer (20 mM TRIS 150 mM NaCl 20 mM imidazole). 6xHis tagged protein was eluted with 1 mL fractions of elution buffer (20 mM TRIS 150 mM NaCl 350 mM Imidazole). Sample was immediately buffer exchanged into 20% Glycerol 20 mM TRIS 150 mM NaCl pH 7.4 using Amicon concentrators. 3C protease was added in a 5:1 ratio of MPro to 3C protease and incubated overnight at 4°C. A 2 L of culture yielded 2.28 mg of MPro following 3C cleavage. 3C protease and 6xHis-tag were removed by incubation with Ni-NTA beads. Monomer was isolated with a MonoQ column. Buffer A: 20 mM Tris 1 mM DTT (fresh) pH 8. Buffer B: 1 M NaCl 20 mM Tris 1 mM DTT (fresh) pH 8. MonoQ column equilibrated with buffer A and eluted with a linear gradient of buffer B 0 mM to 500 mM NaCl over 20 column volumes.

### MPro Inhibition assay

A fluorescence-quenched substrate with the sequence H_2_N(d-Arg)(d-Arg)-K(MCA)-ATLQAIAS-K(DNP)-COOH was synthesized via the Fmoc solid-phase peptide synthesis as previously described^49^. Kinetic measurements were carried out in Corning black 384-well flat-bottom plates and read on a BioTek H4 multimode plate reader. The quenched fluorogenic peptide had a final concentration of K_M_ = 12.7 μM, and MPro had a final concentration of 50 nM. The reaction buffer was 20 mM Tris, 150 mM NaCl, 1 mM EDTA, 0.05% Tween-20 (v/v), and 1 mM DTT, pH 7.4. Compounds were incubated with protease prior to substrate addition at 37 °C for 1 h. After incubation, the substrate was added, and kinetic activity was monitored for 1 h at 37 °C. Initial velocities were calculated at 1 to 30 min in RFU/s. Velocities were corrected by subtracting the relative fluorescence of a substrate-only control, and fraction activity was calculated using a substrate-corrected no-inhibitor control where DMSO was added instead of a drug. Kinetics measurements were carried out in triplicate. SARS-CoV-1 and MERS major protease were both purchased from Bio-Techne (catalogue #: E-718-050 and E-719-050, respectively). K_M_ was derived with the NSP7 substrate for each protease (**Fig. S9**), which was the substrate concentration used for each protease for comparative dose-response curves. Enzyme concentration was 50 nM for SARS-CoV-1 and 100 nM for MERS. The same assay buffer described above was used for all kinetic assays with each protease.

### Non-covalent molecular docking

The protein template was modeled based on the crystal structure of the MPro dimer in complex with a covalent alpha-ketoamide inhibitor (PDB 6Y2G)^26^. All water molecules except for HOH 585 and HOH 602, which are located at the dimeric interface, were deleted. The binding pocket of the crystal structure’s chain A was selected for docking. The alpha-ketoamide inhibitor was replaced by the non-covalent SARS-CoV inhibitor SID87915542^28^. Here, the SID87915542-bound MPro crystal structure (PDB 3V3M) was aligned onto the SARS-CoV-2 MPro crystal structure in order to project SID87915542 into the SARS-CoV-2 MPro binding site. Next, the modeled protein-ligand complex and selected water molecules were prepared for docking with the protein prepwizard protocol of Maestro (Schrödinger v. 2019-3)^50^. Protons were added with Epik and protonation states were optimized with PropKa at pH 7. The C-terminus (Ser301) of each protein monomer structure was capped with N-methyl groups while the N-termini (Ser1) were positively charged. Subsequently, the modeled complex was energetically minimized using the OPLS3e force field. To better accommodate the modeled non-covalent ligand SID87915542, the CE atom of Met49 was displaced by 1.7Å from its initial position in the covalently ligated crystal structure (PDB 6Y2G).

Computational docking was performed using DOCK3.7^51^. Precomputed scoring grids for efficient quantification of van der Waals interaction between MPro and docked molecules were generated with CHEMGRID^52^. Using the AMBER united-atom partial charges^53^, electrostatic potentials within the binding pocket were computed following the numerical solution of the Poisson-Boltzmann equation with QNIFFT^54^. The partial charges of the hydrogen at the epsilon nitrogen of His163, as well as the hydrogen atoms of the backbone amines of Gly143 and Glu166 were increased by 0.4 elementary charge units (e). In turn, the partial charges of oxygen atoms of the corresponding backbone carbonyl groups were decreased by 0.4e to maintain the initial net charge of each residue^29^. The low dielectric protein environment was extended by 1.2 Å from the protein surface, as previously described^55^. Similarly, the low dielectric boundary was extended by 0.7 Å from the protein surface for the calculation of ligand desolvation scoring grids with SOLVMAP^56^. The atomic coordinates of SID87915542 (PDB 3V3M)^28^, the alpha-ketamide inhibitor of the initial crystal structure (PDB 6Y2G)^26^, BDBM512845 (PDB 4MDS)^57^, as well as fragment hits MAT-POS-7dfc56d9-1 (MPro-x0161)^17^ and AAR-POS-d2a4d1df-5 (MPro-x0305)^17^ obtained from the Covid-19 Moonshot screening efforts, were used to generate 80 matching spheres^51^ for ligand placement in the docking calculations.

The obtained docking parameters were evaluated based on their ability to prioritize 34 previously reported ligands of SARS-CoV MPro obtained from the Chembl database^58^, against a background of 1,805 property matched decoys generated with the DUDE-Z approach^35^. In addition, an ‘Extrema’ set^29, 40^ of 194,921 molecules, including compounds with net-charges ranging from −2 to +2, was screened against the docking model in order to assess the parameters’ ability to prioritize neutral molecules.

Using the ZINC15 database^30^, 225,327,212 neutral molecules mainly from the lead-like chemical space, i.e. molecular weight (MWT) between 250 and 350 amu and calculated (c)logP ≤ 4.5, from the make-on-demand compound libraries from Enamine Ltd. and WuXi Appetec. (Shanghai, China), were screened. Thereby, 219,305,079 molecules were successfully scored with each molecule sampling on average 3,588 orientations and 425 conformations which resulted in the evaluation of approximately 148 trillion complexes in roughly 70 hours on a 1,000-core computer cluster. In addition, 110,898,461 molecules with 350 < MWT ≤ 500 and clogP ≤ 4.5 from ZINC15 were screened in a separate docking campaign. 107,486,710 compounds were successfully scored, each exploring on average 4,175 orientations and 540 conformations within the binding pocket. Nearly 90 trillion complexes were scored in roughly 45 hours using a 1,000-core cluster.

From each docking screen, the predicted binding poses of the 500,000 top-ranked molecules were analyzed for internal molecular strain^31^. Molecules that passed the strain criteria (total strain <6.5 TEU; maximum single torsion <1.8 TEU), were judged by their ability to form hydrogen bonds with Gly143, His163 (S1 subpocket) or Glu166 and proximity to residues forming the S2 subpocket such as Met49 or Asp187. Finally, 120 compounds, selected from the lead-like docking screen, were ordered from Enamine Ltd., of which 105 were successfully synthesized (87.5%) in addition to 100 molecules of larger MWT that were ordered from the second docking screen, 89 of which were successfully synthesized by Enamine Ltd.

A second docking campaign for non-covalent inhibitors was performed against the crystal structure of MPro in complex with MAT-POS-b3e365b9-1 (MPro-x11612)^17^ from the Covid-19 Moonshot consortium. All water molecules except HOH6 and HOH300 were removed and the protein-ligand complex structure was prepared for docking following the protein prepwizard protocol of Maestro (Schrödinger v. 2019-3) as described above.

As described above in the previous docking campaign, the partial charges of the hydrogen atoms at the epsilon nitrogen of His163 and the backbone amine of Glu166 were increased by 0.4e, whereas the partial charges of corresponding backbone carbonyl oxygen atoms were decreased by 0.4e to maintain the net charge of each residue. For calculating electrostatic scoring grids, the low-dielectric volume of the protein was extended by 1.9Å from the protein surface (based on surface mapping spheres generated by Sphgen). In addition, the low dielectric boundary was extended by 1.0Å from the protein surface for calculating ligand desolvation scoring grids with SOLVMAP. The atomic coordinates of MAT-POS-b3e365b9-1 were used to generate 45 matching spheres for ligand placement with DOCK3.8. The performance of the obtained docking grids was evaluated by their ability to enrich 15 previously reported SARS-CoV-2 MPro inhibitors over 650 property-matched decoys or an Extrema set containing 153,256 molecules with net charges ranging from −2 to +2, molecular weight between 300 and 500 amu. Finally, 862,382,088 neutral compounds with 18-29 heavy atoms from the Enamine REAL chemical library were screened using the ZINC22 database (http://files.docking.org/zinc22/). Molecules with strained conformations (total strain > 8 TEU, maximum single strain > 3 TEU), were excluded by the docking program. 778,517,250 molecules were successfully scored, each sampled in approximately 836 conformations and 3,439 orientations, leading to the evaluation of roughly 905.8 trillion complexes within 481h on a 1000-core computer cluster.

21,284,498 compounds scored lower than −35 kcal/mol and the poses of top scoring 5,004,192 compounds were extracted. 214,580 compounds formed favorable interactions with key residues such as His163, Glu166 and the P2 subpocket, 181,866 of which obtained ECFP4-based TC coefficients of less than 0.35 to the 1,716 known SARS-CoV and SARS-CoV-2 MPro inhibitors reported in the literature^2–4, 8, 9, 11, 12, 14, 17, 20, 26, 28, 59–75^. Finally, roughly 9,000 top-ranking compounds were visually inspected, and 167 molecules were ordered from Enamine Ltd., 146 of which (87.4%) were successfully synthesized.

### Covalent molecular docking

Cysteine-reactive warheads of aldehydes, nitriles, and alpha-ketoamides were searched in the ZINC20/Enamine *REAL* databases of 1.4 billion molecules using their respective SMARTS patterns (ketoamides O=[CR0]([#6])[CR0](=O)N[#6]; aldehydes [CX3H1](=O)[#6]; nitriles [CX4]-C#N). This returned 25.7 million nitriles, 2.5 million aldehydes, and 1.5 million ketoamides. Molecules were filtered to have at least one ring, and to be fragment to lead-like molecular weights (<350). Three-dimensional “dockable” conformations were generated with molecules in their transition-state form and a dummy atom in place for the covalent docking algorithm to indicate which atom should be modeled covalently bound to the Cysteine sulfur^36, 37^. Overall, 6.5 million molecules were docked – 3.6 million nitriles, 1.4 million ketoamides, and 1.5 million aldehydes.

The receptor was prepared in DOCK3.7^51^. Pose reproduction of the truncated covalent molecule of PDB 6Y2G^26^ (smiles of dockable ligand: O=C1NCC[C@H]1CC[C@](O)([SiH3])C(=O)NCc2ccccc2) was checked for the docking setup. Default generated grids were used for electrostatic (radius size 1.9Å) and VDW scoring, and no matching spheres were used in docking calculations as they are not used by the covalent docking DOCKovalent^36, 37^ algorithm. For covalent docking, the Cys145 SH group was indicated as the anchor for molecules screened. The distance was slightly relaxed from the C-C bond distance to 1.85Å. For His41 protonation, aldehydes, nitriles, and neutral ketoamides used HID, while negative ketoamides used HIP. Each warhead was docked separately with a total 6.5 million molecules screened. Accordingly, each warhead was also processed separately.

For the aldehydes, the top 300,000 ranked molecules were evaluated for torsional strain^31^, and those with a total torsional strain greater than 9.8 (around 3.7 incurred due to strain on atom types on the warhead and this was disregarded, therefore total energy was 6) and single torsional strain greater than 2.5 were excluded (155,386 left). Molecules making more than 1 hydrogen bond to the protein, having no hydrogen bond clashes, no unpaired hydrogen bond donors (56,969 left) were prioritized. Remaining molecules were clustered for chemical similarity based on ECFP4-based Tanimoto coefficient (Tc) of 0.5. Viable poses filling the S1’, S1 or S2 sites were selected during visual inspection. A total of 35 aldehydes were selected for make-on-demand synthesis of which 27 were successfully synthesized. For the nitriles, the top 100,000 ranked molecules were evaluated for torsional strain (17,424 left), then filtered for favorable interactions (6,201 left). Lastly, we visually inspected remaining molecules for favorable hydrogen bonds formed with His41, Gly143, Thr26, Glu166, or Cys145. Finally, 41 compounds were ordered for synthesis (31 were successfully obtained). For the ketoamides the top 393,000 ranked molecules with scores less than 0.0 were evaluated for torsional strain (121,234 left), and favorable interactions with the enzyme (37,267 remained). Visual inspection focused on those making hydrogen bonds with His41, Cys145, Gly143, Thr26. In total 21 molecules were prioritized and 16 were successfully synthesized.

### Make-on-demand synthesis

Non-covalent and covalent compounds purchased from docking screens, as well as analogs, were synthesized by Enamine Ltd. (**Table S1**). Purities of molecules were at least 90% and most active compounds were at least 95% (based on LC/MS data) (**Fig. S10**).

### Compound optimization

Optimization of docking hits **ZINC346371112, ZINC301553312, ZINC813360541, ZINC553840273, ‘3620**, **‘0431**, **‘4589**, **‘5103**, **‘5156**, **‘6246**, **‘6792**, **‘0292**, **‘2826/’0892** were attempted (**Table S1**). Analogs were designed for desired chemical perturbations or searched in SmallWorld and Arthor catalogs and synthesized by Enamine Ltd. For **‘3620**, compounds were also designed from the **‘7356** and **‘7021** crystal structures and were modeled with covalent docking or with Maestro (v. 2021-2, Schrödinger, LLC) ligand alignment.

### Protein crystallization

Both covalent and non-covalent compounds including **7021, ‘9121, 8252, ‘9218, 7356, 5548, 6111** and **SG-0001** were co-crystallized with SARS-CoV2 main protease. Before setting up crystals, 10 mg/ml of protein was incubated with either 0.3 mM of covalent compounds or 1.5 mM of non-covalent compounds on ice for 1 hour. Crystals were set using vapor diffusion hanging drop method at 20 C^◦^ in conditions including 0.1 M Tris pH 7.4 and 20% PEG 8000; and 0.1 M MES pH 6.5, 20% PEG 6000. Crystals took 3-4 days to grow for all compounds. Before data collection, crystals were cryo-cooled in a solution containing reservoir solution and 25% glycerol.

### Structure determination and refinement

The MPro-inhibitor compound datasets were either collected at the Advanced Light Source beamline 8.3.1 (Lawrence Berkeley laboratory) or SSRL beamline 12-2 beamline (Stanford, United States) at a temperature of 100K. The diffraction datasets were processed using XDS^76^ and CCP4 software’s suite^77^. AIMLESS^78^ was used for scaling and merging. Molecular replacement was performed either using PHASER^79^ using the protein model from PDB entry 7NG3^80^ as the search model. The bound ligand in the PDB 7NG3 was removed from the search model during molecular replacement, giving unbiased electron density for ligands in the initial electron density maps. The initial model fitting and addition of waters was done in COOT^81^ followed by refinement in REFMAC^82^. Geometry restraints for the ligands were created in eLBOW-PHENIX^83^ and following rounds of refinement were carried out in PHENIX. Geometry for each structure was assessed using Molprobidity and PHENIX polygon. Datasets have been deposited to the PDB with PDB IDs 8DIB, 8DIC, 8DID, 8DIE, 8DIF, 8DIG, 8DIH and 8DII. Statistics for data collection and refinement are in **Table S5**. The ligand symmetry accounted RMSDs between the docked pose and experimental pose were calculated by the Hungarian algorithm in DOCK6^84^.

### Antiviral and cytotoxicity assays

Two thousand (2,000) HeLa-ACE2 cells were seeded into 96-well plates and incubated for 24 h. 2 h before infection, the medium was replaced with a new media containing the compound of interest, including a DMSO control. Plates were then transferred into the biosafety level 3 (BSL-3) facility and 1,000 PFU (MOI = 0.25) of SARS-CoV-2 was added, bringing the final compound concentration to those indicated. SARS-CoV-2/WA1 variant was used as indicated. Plates were then incubated for 48 h. Infectivity was measured by the accumulation of viral NP protein in the nucleus of the HeLa-ACE2 cells (fluorescence accumulation). Percent infection was quantified as ((Infected cells/Total cells) − Background) × 100, and the DMSO control was then set to 100% infection for analysis. Cytotoxicity was also performed at matched concentrations using the MTT assay (Roche), according to the manufacturer’s instructions. Cytotoxicity was performed in uninfected HeLa-ACE2 cells with same compound dilutions and concurrent with viral replication assay. All assays were performed in biologically independent triplicates.

### Dynamic Light Scattering (DLS)

Samples were prepared in filtered 50 mM KPi buffer, pH 7.0 with final DMSO concentration at 1% (v/v). Colloidal particle formation was detected using DynaPro Plate Reader II (Wyatt Technologies). All compounds were screened in triplicate at roughly 2-fold higher than reported IC_50_ (concentrations can be found in **Table S1**). Analysis was performed with GraphPad Prism software version 9.1.1 (San Diego, CA).

### Enzyme Inhibition Assays for Aggregation

Enzyme inhibition assays were performed at room temperature using using CLARIOstar Plate Reader (BMG Labtech). Samples were prepared in 50 mM KPi buffer, pH 7.0 with final DMSO concentration at 1% (v/v). Compounds were incubated with 4 nM AmpC β-lactamase (AmpC) or Malate dehydrogenase (MDH) for 5 minutes. AmpC reactions were initiated by the addition of 50 μM CENTA chromogenic substrate. The change in absorbance was monitored at 405 nm for 1 min 45 sec. MDH reactions were initiated by the addition of 200 μM nicotinamide adenine dinucleotide (NADH) (54839, Sigma Aldrich) and 200 μM oxaloacetic acid (324427, Sigma Aldrich). The change in absorbance was monitored at 340 nm for 1 min 45 sec. Initial rates were divided by the DMSO control rate to determine % enzyme activity. Each compound was screened roughly 2-fold higher than reported IC_50_ value in triplicate (concentrations can be found in **Table S1**). Data was analyzed using GraphPad Prism software version 9.1.1 (San Diego, CA).

For detergent reversibility experiments, inhibition was screened at/near IC75 with or without 0.01% (v/v) Triton X-100 in triplicate. Enzymatic reaction was performed/monitored as previously described^32, 85^.

### Statistical analyses

All statistical analyses were performed on the GraphPad Prism version 8.0 or 9.1.1 software. Changes only at the 95% confidence level (P<0.05) were considered as statistically significant.

### Data availability

All crystallographic structures have been deposited in the PDB as 8DIB (‘7021), 8DIC (‘9121), 8DID (8252), 8DIE (‘9218), 8DIF (‘7356), 8DIG (‘5548), 8DIH (‘6111), 8DII (SG-0001). The identities of compounds docked in non-covalent screens can be found at ZINC15/ZINC20 (http://zinc15.docking.org and http://zinc20.docking.org) and ZINC22 (http://files.docking.org/zinc22/). The covalent compounds have been deposited in http://covalent2022.docking.org along with their DOCKovalent files. Active compounds may be purchased from Enamine Ltd. All other data are available from the corresponding authors on request.

### Code availability

DOCK3.7 and DOCK3.8 are freely available for non-commercial research from the authors; commercial licenses are available via the UC Regents. An open-source web-based version of the program is available without restriction to all (https://blaster.docking.org), as are the Arthor and Small World analoging tools used in this study.

## Acknowledgments

Supported by DARPA grant HR0011-19-2-0020 (B.K.S., J.J.I., A.G.-S.) and by NIH grant R35GM122481 (B.K.S., J.J.I.). This was also funded by NIAID grant U19AI171110 (C.S.C., B.K.S., A.G.-S., A.R.R.) run by Principle Investigator Nevan Krogan. This work was also partly funded by CRIPT (Center for Research on Influenza Pathogenesis and Transmission), an NIAID funded Center of Excellence for Influenza Research and Response (CEIRR, contract #75N93021C00014) and by supplements to DoD grant W81XWH-20-1-0270 and to NIAID grant U19AI135972 to A.G.-S. B.A. received the Covid Catalyst Award from the Center for Emerging and Neglected Diseases (CEND). We gratefully acknowledge OpenEye Software for Omega and related tools, and Schrodinger LLC for the Maestro package. We also thank Dr. Randy Albrecht for support with the BSL3 facility and procedures at the ISMMS as well as Richard Cadagan and Daniel Flores for excellent technical assistance. In addition, we thank Dr. Rolf Hilgenfeld for providing the SARS-CoV-2 MPro plasmid. We acknowledge the contributions of the UCSF Chemical Underpinnings of Biological Systems (CUBS) 2021 cohort which included Siyi Wang, Isabel Lee, Vineet Mathur, Sham Rampersaud, Luis Santiago, Sara Warrington, and Rose Yang.

## Author contributions

S.G. conducted non-covalent docking screens and compound optimization, with input from B.K.S., J.L., S.V., and the 2021 CUBS Cohort. E.A.F., S.Gu, and X.W. performed covalent database building with input from J.T. and B.K.S., covalent docking with input from B.K.S., and compound optimization with input from B.K.S, A.R.R., S.G., and I.S. Enzymatic testing was conducted by C.B., assisted by N.J.Y., and supervised by C.S.C. Antiviral and cytotoxicity assays were performed by K.W., with supervision by A.G.-S. Protein purification was done by I.S., B.A., P.F., with supervision by B.K.S., C.S.C., and A.O. Crystallography was done by I.S. assisted by J.O. and with input from B.K.S. Aggregation testing was performed by I.G. and H.O. with input from B.K.S. ZINC15 and ZINC22 databases were built by J.J.I. Y.S.M. supervised compound synthesis of Enamine compounds, assisted by I.S.K. C.S.C. and B.K.S. supervised the project. E.A.F., S.G., and C.B. wrote the paper with input from all other authors, and primary editing from C.S.C. and B.K.S. C.S.C. and B.K.S. conceived the project.

