## Supplemental Information for "Large library docking for novel SARS-CoV-2 main protease non-covalent and covalent inhibitors"

**This document includes:**

Tables S2 to S5
Figures S1 to S10

**Other Supplementary Material for this manuscript includes the following:**

Table S1 (.xlsx)

**Supplementary Table 2. Analogs of covalent docking hit ‘3620 with improved potencies.**

| Compound | IC50 [µM] | Compound | IC50 [µM] |
|---|---|---|---|
| 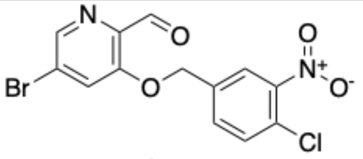 **‘7021** | 1 | 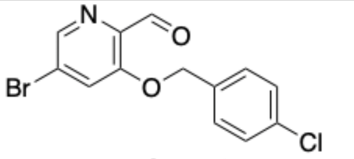 **Analog_12** | 22 |
| 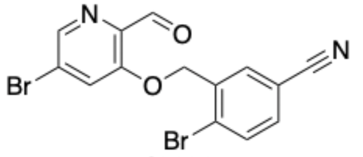 **‘6690** | 2 | 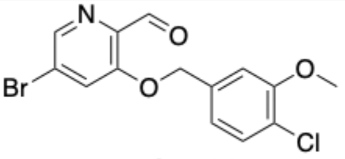 **Analog_20** | 22 |
| 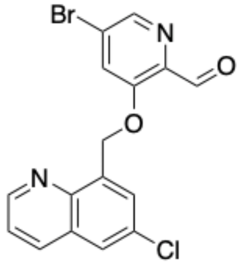 **‘7013** | 5 | 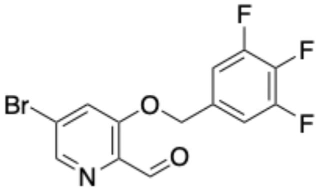 **‘9037** | 23 |
| 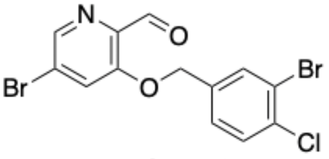 **Analog_18** | 6 | 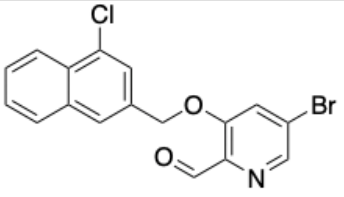 **‘4202** | 24 |
| 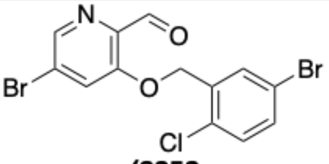 **‘8252** | 6 | 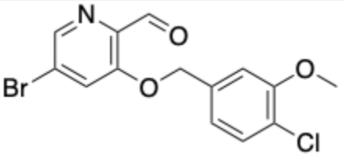 **‘9125** | 25 |
| 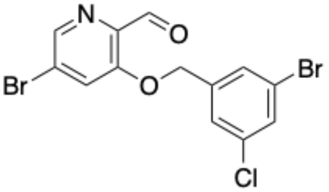 **‘6117** | 7 | 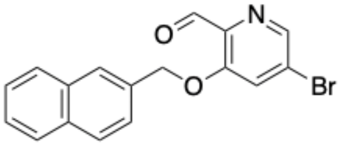 **‘7356** | 26 |
| 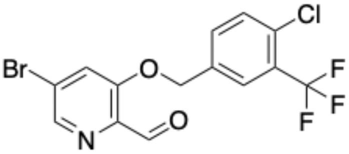 **Analog_19** | 9 | 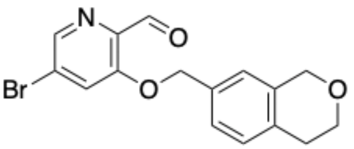 **‘8474** | 27 |

| | | | |
|---|---|---|---|
| 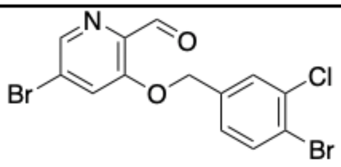 '4217 | 9 | 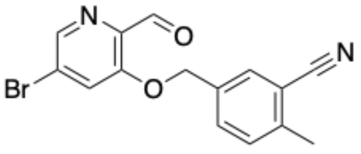 '0481 | 27 |
| 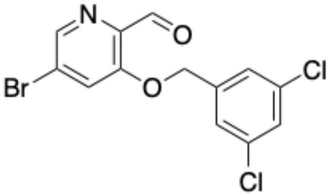 '9033 | 9 | 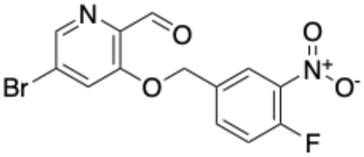 Analog_7 | 28 |
| 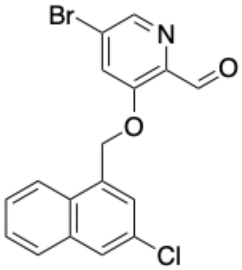 '9122 | 9 | 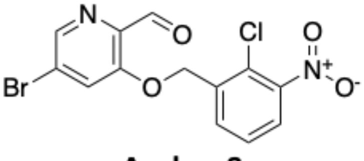 Analog_8 | 33 |
| 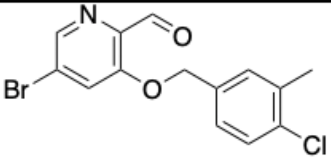 Analog_17 | 11 | 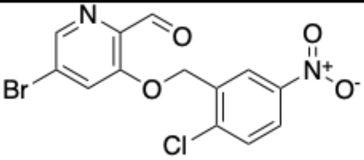 Analog_10 | 34 |
| 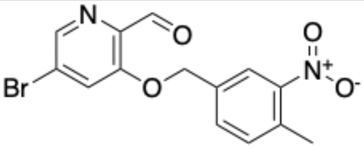 Analog_6 | 12 | 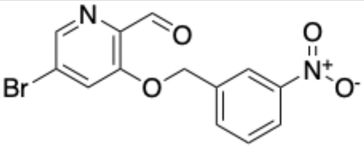 Analog_11 | 35 |
| 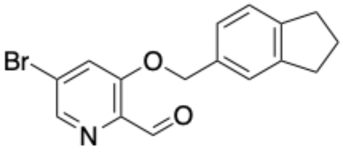 Analog_49 | 13 | 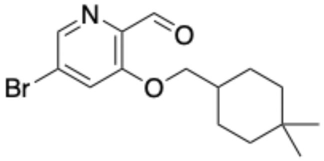 '9009 | 38 |
| 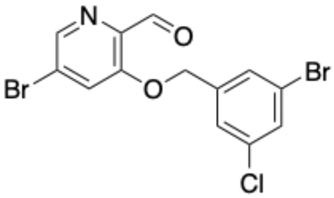 '0156 | 14 | 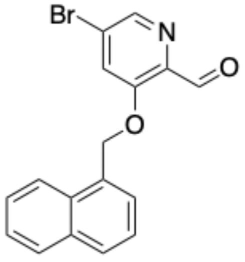 '3251 | 40 |

| | | | |
|---|---|---|---|
| 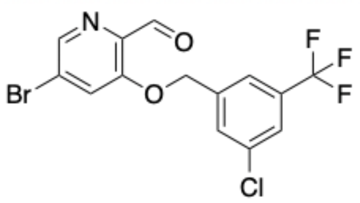 **'9028** | 14 | 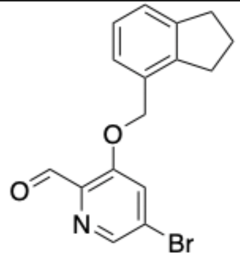 **'3250** | 40 |
| 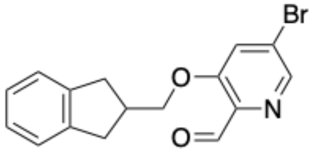 **'4221** | 19 | 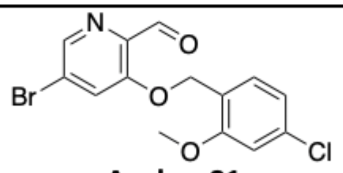 **Analog_21** | 43 |
| 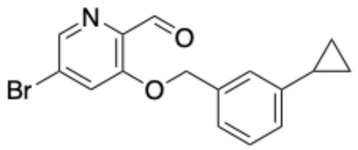 **'3264** | 20 | 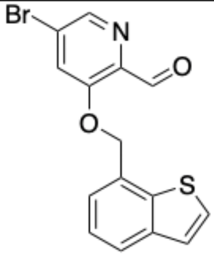 **'3252** | 45 |
| 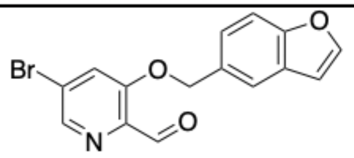 **'4212** | 20 | 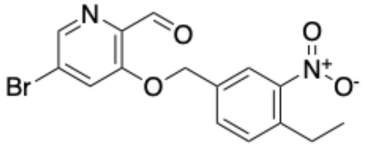 **'3564** | 48 |

**Supplementary Table 3. Antiviral activities.**

| Compound | Antiviral $IC_{50}$[a] [μM] | Antiviral $IC_{90}$[a] [μM] | $CC_{10}$[b] [μM] |
|---|---|---|---|
| **‘7021** | 6.2 | 7.3 | 8.03 |
| **‘7356** | 19.53 | 19.9 | 6.93 |

[a] Efficacy in RT-qPCR viral infectivity assay in X cells.
[b] Reduction in cell viability (cytotoxicity).

**Supplementary Table 4. Pan-viral enzymatic activities of ‘7021.**

| Viral MPro | IC50 [μM] |
|---|---|
| MERS | 50 |
| SARS-CoV-1 | 8 |
| SARS-CoV-2 | 1 |

#### Supplementary Table 5. Crystallographic statistics.

| Ligand | ‘5548 | ‘7356 | ‘6111 | SG-0001 | ‘9121 | ‘7021 | ‘8252 | ‘9218 |
|---|---|---|---|---|---|---|---|---|
| **PDB ID** | **8DIG** | **8DIF** | **8DIH** | **8DII** | **8DIC** | **8DIB** | **8DID** | **8DIE** |
| Resolution range (Å) | 72.62 - 2.45 (2.538 - 2.45) | 48.44 - 1.98 (2.051 - 1.98) | 44.39 - 2.12 (2.196 - 2.12) | 48.6 - 2.59 (2.683 - 2.59) | 38.13 - 2.09 (2.165 - 2.09) | 38.73 - 2.17 (2.248 - 2.17) | 38.22 - 1.95 (2.02 - 1.95) | 38.64 - 1.9 (1.968 - 1.9) |
| Space group | P 21 21 21 | P 1 21 1 | C 1 2 1 | I 1 2 1 | P 1 21 1 | P 1 21 1 | P 1 21 1 | P 1 21 1 |
| *a, b, c* (Å) | 67.779, 102.704, 102.704 | 44.897, 53.624, 115.04 | 114.102, 53.917, 45.258 | 45.041, 53.835, 115.11 | 44.346, 53.814, 115.824 | 44.911, 53.72, 115.527 | 44.238, 53.666, 115.632 | 44.923, 53.751, 115.119 |
| a, b, c (°) | 90, 90, 90 | 90, 101.151, 90 | 90, 101.253, 90 | 90, 101.13, 90 | 90, 100.459, 90 | 90, 100.833, 90 | 90, 100.938, 90 | 90, 101.062, 90 |
| Total reflections | 52786 (4222) | 74588 (6993) | 29895 (2343) | 17080 (1724) | 63019 (6332) | 56799 (5689) | 76834 (7688) | 83998 (8382) |
| Unique reflections | 26433 (2151) | 37368 (3516) | 14998 (1188) | 8552 (864) | 31556 (3170) | 28435 (2848) | 38463 (3848) | 42036 (4194) |
| Multiplicity | 2.0 (2.0) | 2.0 (2.0) | 2.0 (2.0) | 2.0 (2.0) | 2.0 (2.0) | 2.0 (2.0) | 2.0 (2.0) | 2.0 (2.0) |
| Completeness (%) | 97.69 (80.05) | 99.03 (92.87) | 96.99 (77.24) | 99.74 (99.77) | 98.17 (98.84) | 98.18 (98.41) | 98.14 (98.54) | 94.67 (79.45) |
| Mean I/sigma(I) | 10.82 (1.33) | 13.42 (1.00) | 19.51 (1.83) | 8.81 (0.84) | 8.94 (0.98) | 8.68 (0.89) | 7.58 (0.98) | 4.57 (0.59) |
| Wilson B-factor | 44.04 | 36.71 | 48.65 | 56.27 | 39.01 | 40.42 | 30.04 | 33.18 |
| R-merge | 0.05895 (0.6349) | 0.0362 (0.8374) | 0.01881 (0.4432) | 0.06999 (0.9217) | 0.05198 (0.7831) | 0.05847 (0.8779) | 0.05816 (0.7736) | 0.1022 (1.824) |
| R-meas | 0.08337 (0.8979) | 0.0512 (1.184) | 0.0266 (0.6267) | 0.09899 (1.304) | 0.07351 (1.107) | 0.08269 (1.242) | 0.08226 (1.094) | 0.1446 (2.579) |
| R-pim | 0.05895 (0.6349) | 0.0362 (0.8374) | 0.01881 (0.4432) | 0.06999 (0.9217) | 0.05198 (0.7831) | 0.05847 (0.8779) | 0.05816 (0.7736) | 0.1022 (1.824) |
| CC1/2 | 0.997 (0.513) | 1 (0.41) | 1 (0.625) | 0.995 (0.333) | 0.999 (0.474) | 0.998 (0.397) | 0.998 (0.513) | 0.993 (0.201) |
| CC* | 0.999 (0.823) | 1 (0.763) | 1 (0.877) | 0.999 (0.707) | 1 (0.802) | 1 (0.754) | 1 (0.823) | 0.998 (0.579) |
| **Refinement** | | | | | | | | |
| Reflections used in refinement | 26397 (2118) | 37312 (3502) | 14996 (1188) | 8548 (862) | 31513 (3163) | 28394 (2844) | 38390 (3844) | 40479 (3371) |
| Reflections used for R-free | 1317 (116) | 1835 (176) | 723 (54) | 419 (45) | 1615 (173) | 1341 (143) | 1894 (224) | 2128 (174) |
| R-work | 0.2138 (0.3321) | 0.2432 (0.3608) | 0.2114 (0.3256) | 0.2164 (0.3640) | 0.2275 (0.3419) | 0.2267 (0.3265) | 0.2574 (0.3563) | 0.2534 (0.3559) |
| R-free | 0.2717 (0.3771) | 0.2955 (0.3532) | 0.2760 (0.3358) | 0.2672 (0.3837) | 0.2764 (0.3772) | 0.2814 (0.3331) | 0.2915 (0.4119) | 0.3032 (0.3935) |
| CC(work) | 0.954 (0.748) | 0.957 (0.639) | 0.964 (0.698) | 0.957 (0.515) | 0.959 (0.639) | 0.962 (0.651) | 0.954 (0.698) | 0.953 (0.502) |
| CC(free) | 0.922 (0.479) | 0.953 (0.613) | 0.963 (0.738) | 0.947 (0.420) | 0.932 (0.465) | 0.955 (0.574) | 0.948 (0.514) | 0.916 (0.400) |
| Number of non-hydrogen atoms | 4589 | 4365 | 2317 | 2347 | 4502 | 4579 | 4531 | 4604 |
| macromolecules | 4540 | 4269 | 2267 | 2310 | 4433 | 4506 | 4413 | 4509 |
| ligands | 25 | 42 | 25 | 25 | 38 | 42 | 38 | 42 |
| solvent | 24 | 54 | 25 | 12 | 31 | 31 | 80 | 53 |
| Protein residues | 601 | 585 | 303 | 303 | 599 | 599 | 591 | 605 |
| RMS(bonds) | 0.008 | 0.01 | 0.008 | 0.002 | 0.009 | 0.004 | 0.015 | 0.009 |
| RMS(angles) | 0.94 | 0.99 | 1.01 | 0.57 | 0.98 | 0.7 | 1.14 | 0.92 |
| Ramachandran favored (%) | 94.76 | 97.68 | 96.62 | 96.66 | 97.23 | 98.11 | 97.33 | 97.45 |
| Ramachandran allowed (%) | 5.24 | 2.32 | 3.38 | 3.34 | 2.6 | 1.72 | 2.49 | 2.38 |
| Ramachandran outliers (%) | 0 | 0 | 0 | 0 | 0.17 | 0.17 | 0.18 | 0.17 |
| Rotamer outliers (%) | 0 | 0.23 | 0 | 0 | 0.43 | 0 | 0.21 | 0 |
| Clashscore | 7.5 | 3.99 | 4.71 | 5.48 | 6.71 | 3.84 | 8.22 | 5.1 |

| | | | | | | | | |
|---|---|---|---|---|---|---|---|---|
| Average B-factor | 49.4 | 40.26 | 56.53 | 63.66 | 44.1 | 48.58 | 33.91 | 37.93 |
| macromolecules | 49.35 | 40.11 | 56.35 | 63.09 | 43.98 | 48.12 | 33.78 | 37.84 |
| ligands | 60.21 | 54.15 | 73.4 | 122.98 | 61.63 | 99.97 | 55.66 | 50.55 |
| solvent | 48.32 | 41.14 | 55.62 | 49.68 | 39.64 | 46.39 | 30.93 | 35.8 |

(One crystal for each structure)
* Values in parentheses are for the highest-resolution shell.

**A**

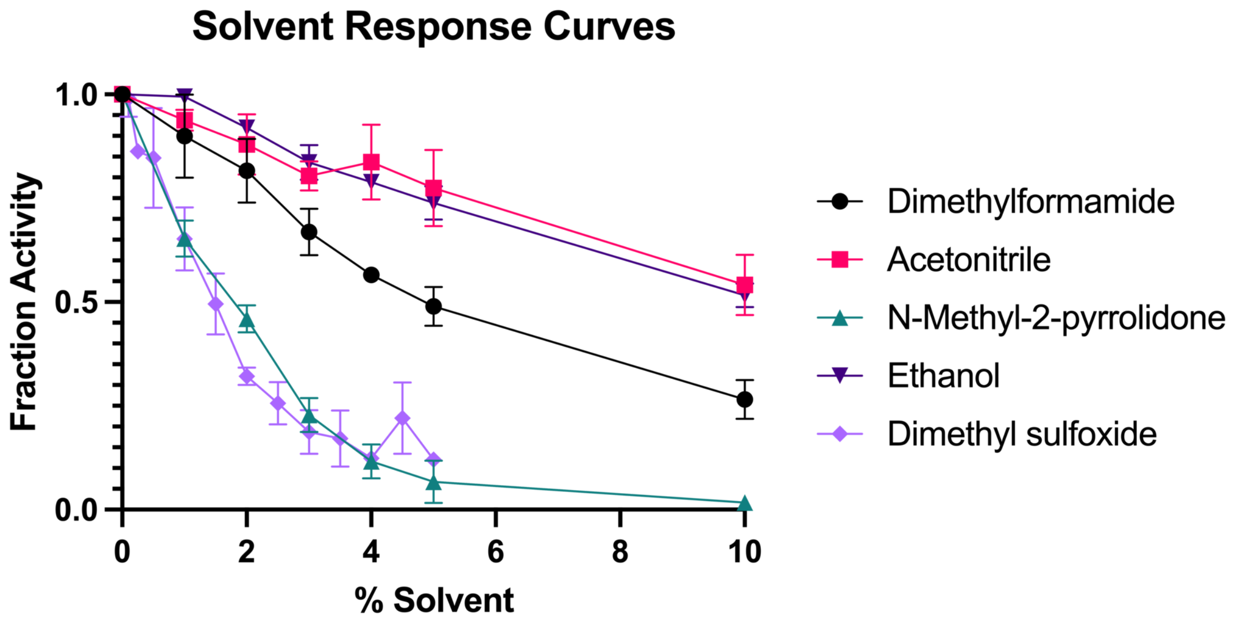


**B**

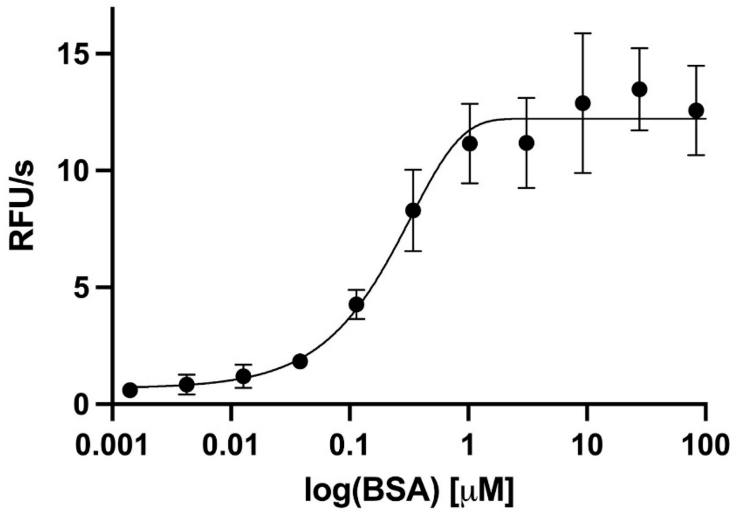


**Supplementary Figure 1. Assay optimization for solvent and detergent.** (**A**) The effect of solvents on enzymatic activity of MPro. (**B**) Activity loss from removal of detergent rescued with addition of BSA.

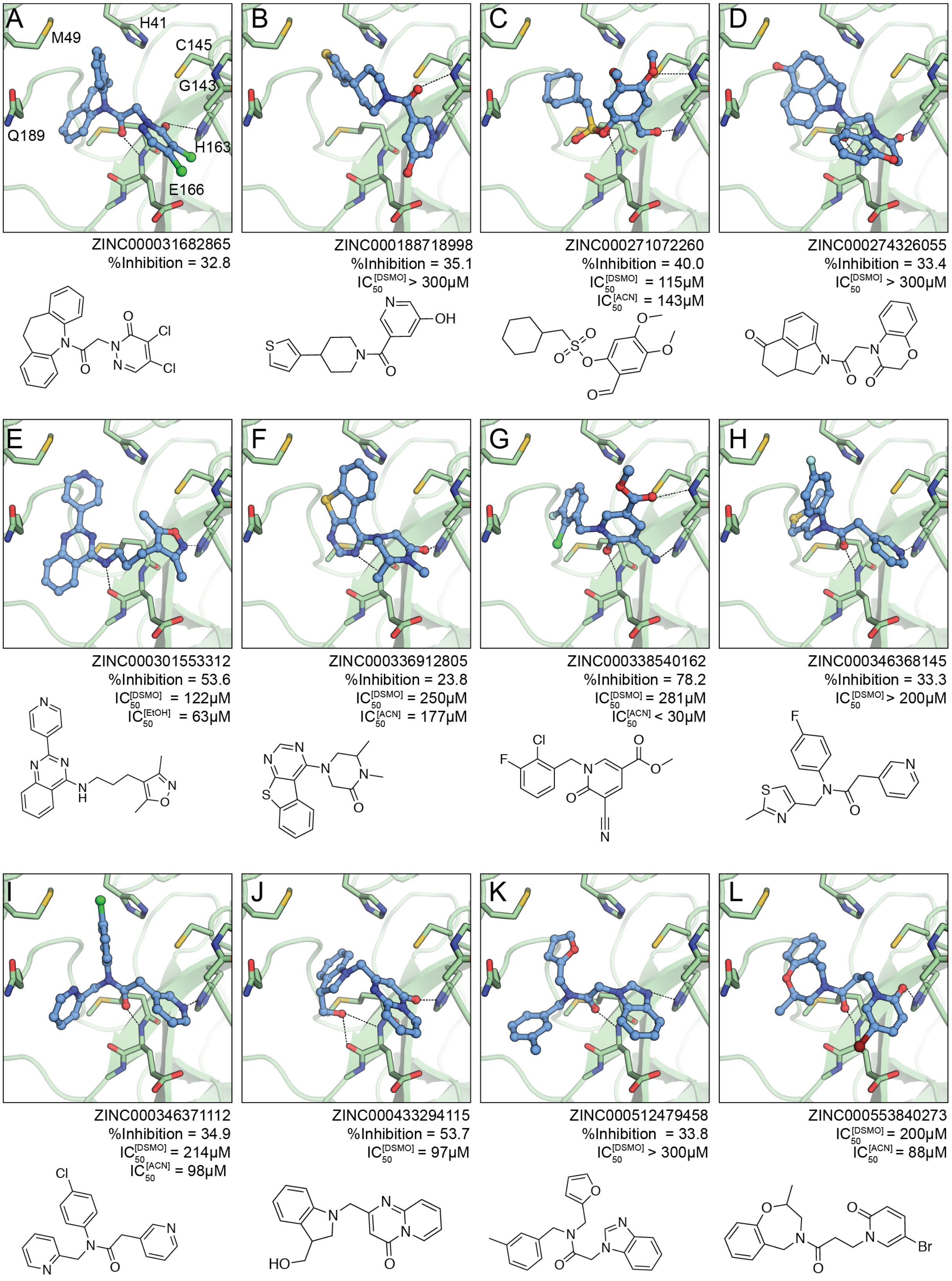


**Supplementary Figure 2.1 Non-covalent docking hits or compounds with >30% inhibition from first virtual screen.**

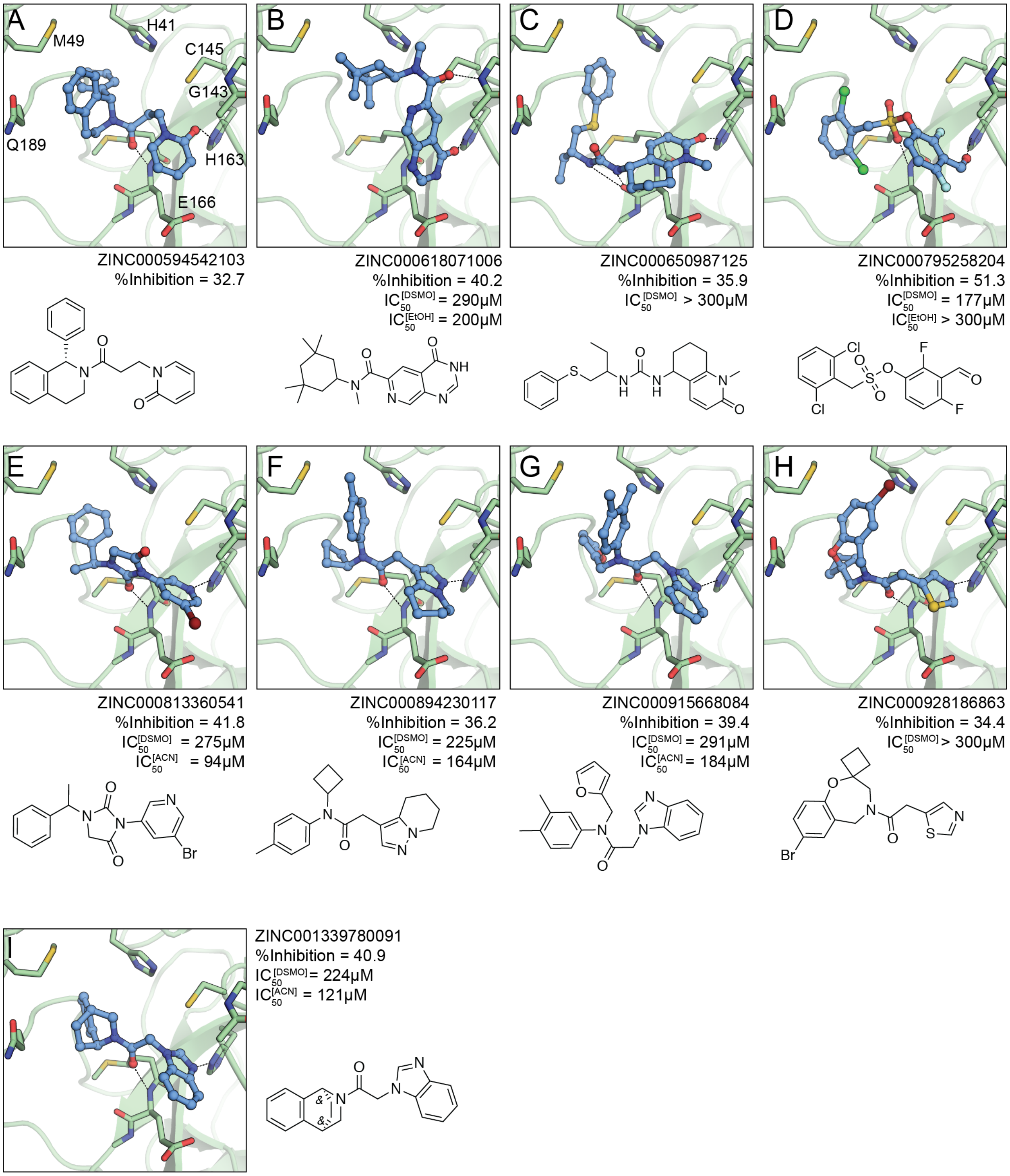


**Supplementary Figure 2.2 Non-covalent docking hits or compounds with >30% inhibition from first virtual screen.**

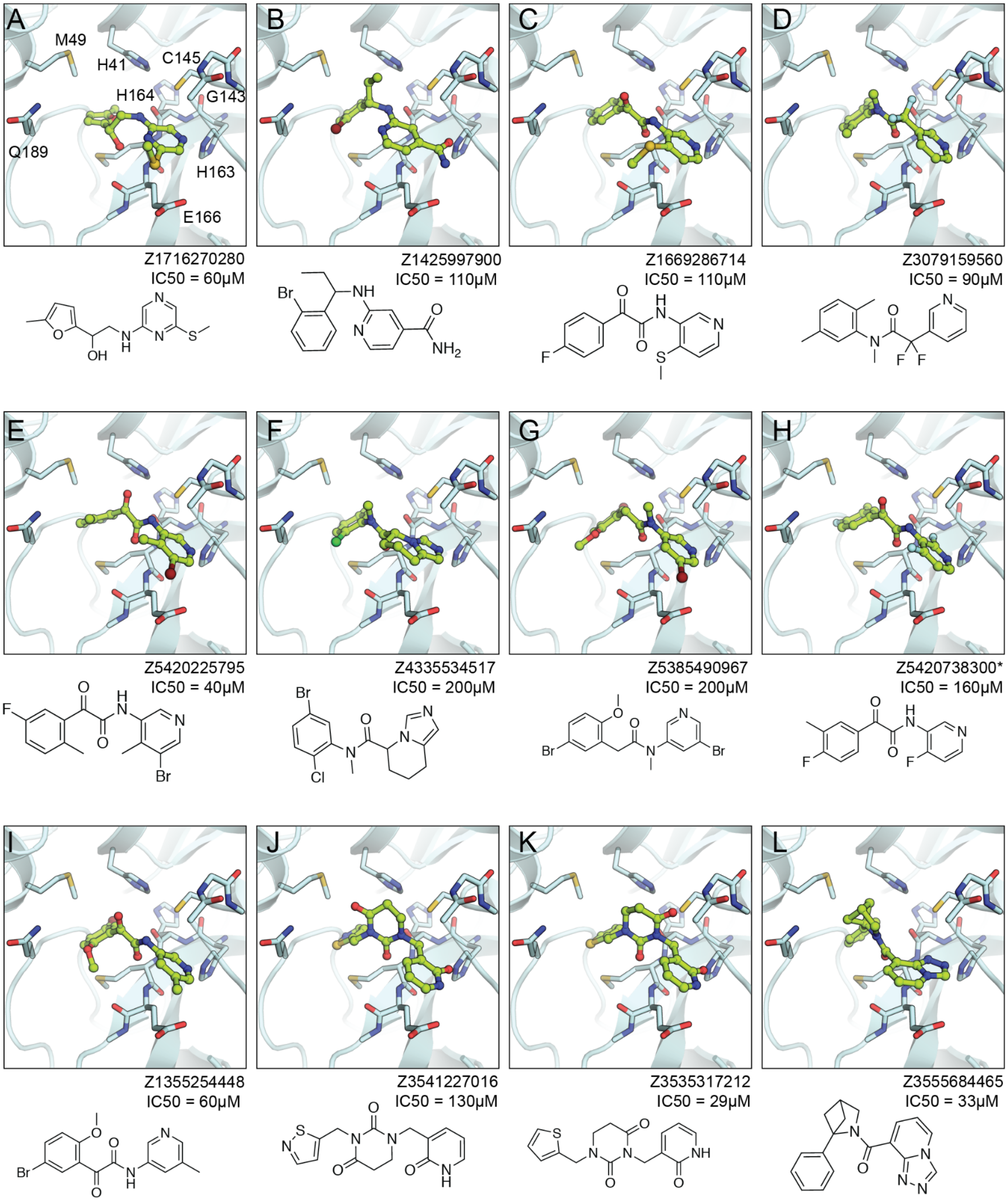


**Supplementary Figure 2.3 Non-covalent docking hits from second virtual screen.**

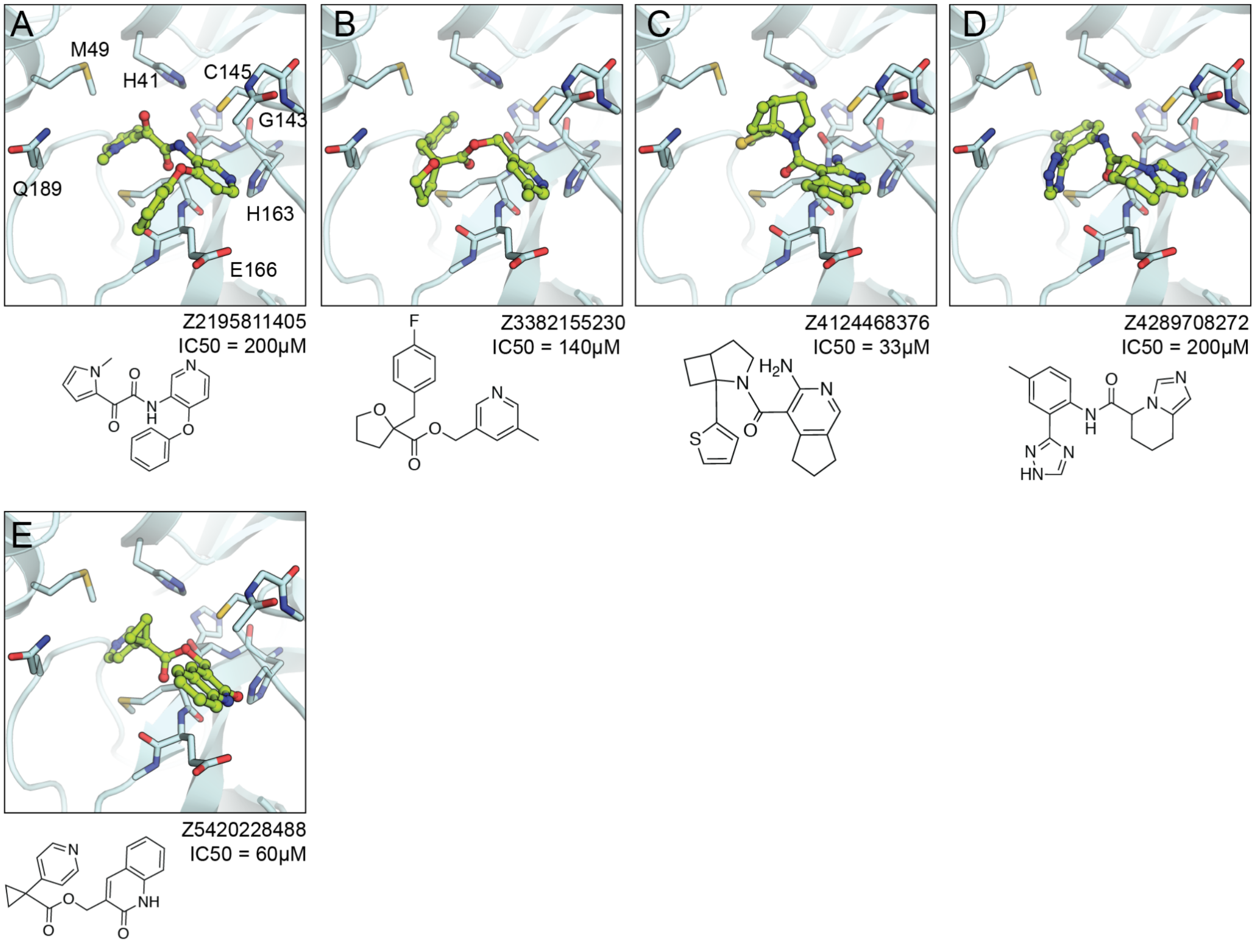


**Supplementary Figure 2.4 Non-covalent docking hits from second virtual screen.**

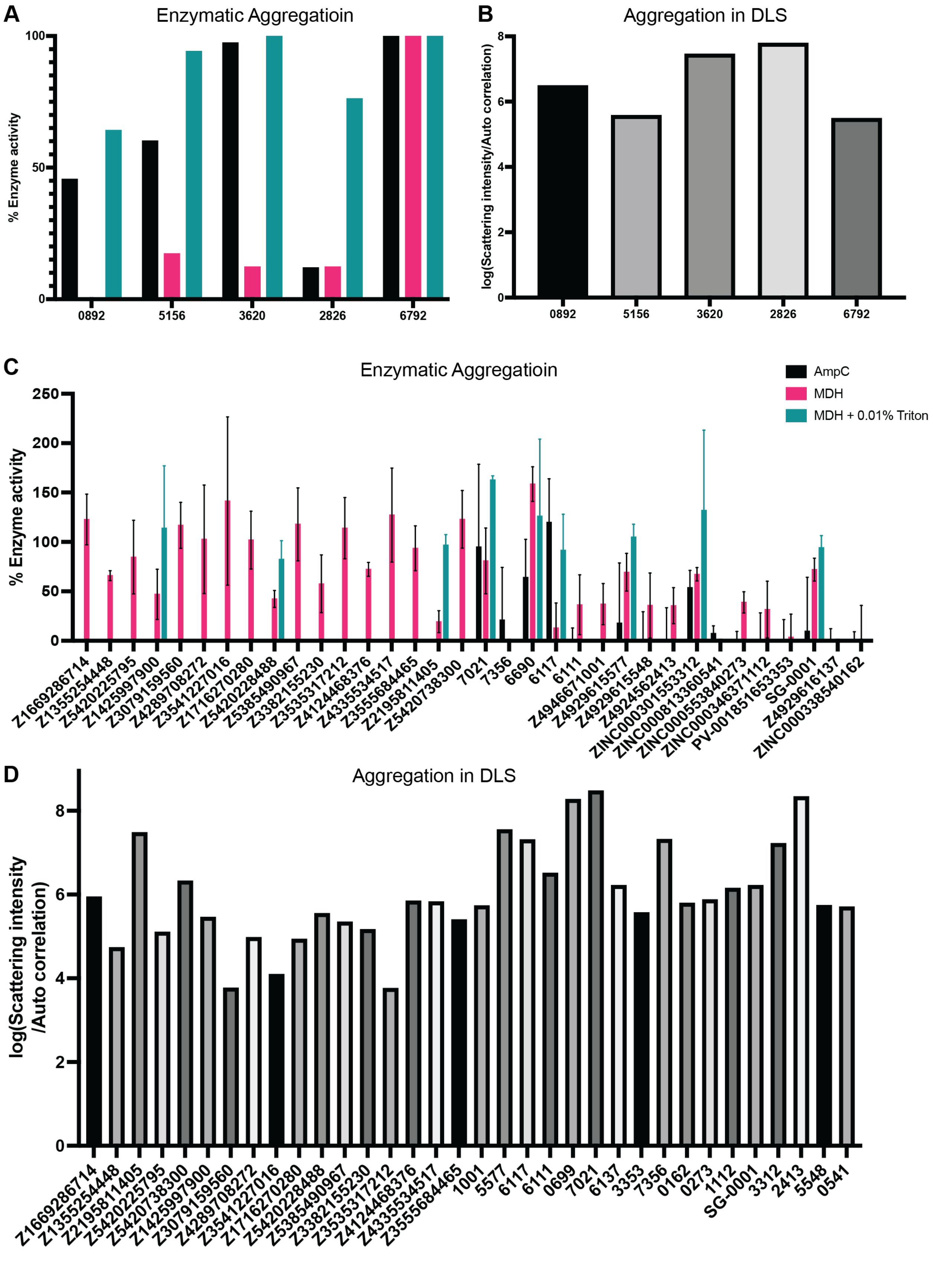
A
Enzymatic Aggregatioin
% Enzyme activity
0
50
100
0892
5156
3620
2826
6792
B
Aggregation in DLS
log(Scattering intensity/Auto correlation)
0
2
4
6
8
0892
5156
3620
2826
6792
C
Enzymatic Aggregatioin
AmpC
MDH
MDH + 0.01% Triton
% Enzyme activity
0
50
100
150
200
250
Z1669286714
Z1355254448
Z5420225795
Z1425997900
Z3079159560
Z4289708272
Z3541227016
Z1716270280
Z5420228488
Z5385490967
Z3382155230
Z3535317212
Z4124468376
Z4335534517
Z3555684465
Z2195811405
Z5420738300
7021
7356
6690
6117
6111
Z4946671001
Z4929615577
Z4929615548
Z4924562413
ZINC000301553312
ZINC000813360541
ZINC000553840273
ZINC000346371112
PV-001851653353
SG-0001
Z4929616137
ZINC000338540162
D
Aggregation in DLS
log(Scattering intensity /Auto correlation)
0
2
4
6
8
Z1669286714
Z1355254448
Z2195811405
Z5420225795
Z5420738300
Z1425997900
Z3079159560
Z4289708272
Z3541227016
Z1716270280
Z5420228488
Z5385490967
Z3382155230
Z3535317212
Z4124468376
Z4335534517
Z3555684465
1001
5577
6117
6111
0699
7021
6137
3353
7356
0162
0273
1112
SG-0001
3312
2413
5548
0541

**Supplementary Figure 3. Evaluating aggregation potential of initial docking hits and potent analogs.** (**A,C**) Enzymatic inhibition by aggregation tested against AmpC and MDH (with or without 0.01% Triton X-100 detergent). (**B,D**) Compounds observed in DLS for forming colloidal-like particles where values above $10^6$ have the potential for aggregating.

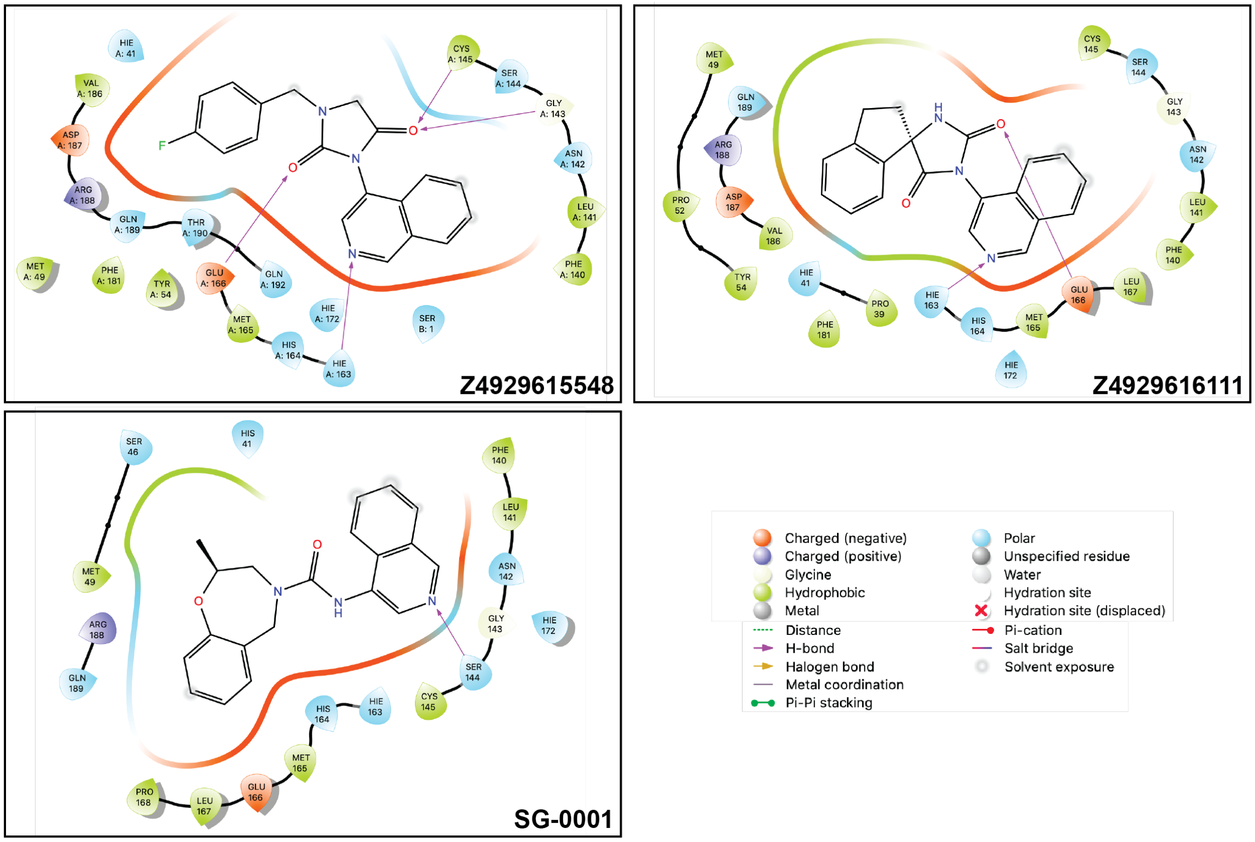


**Supplementary Figure 4. LigPlot visualization of MPro-noncovalent inhibitor interactions in newly solved structures.**

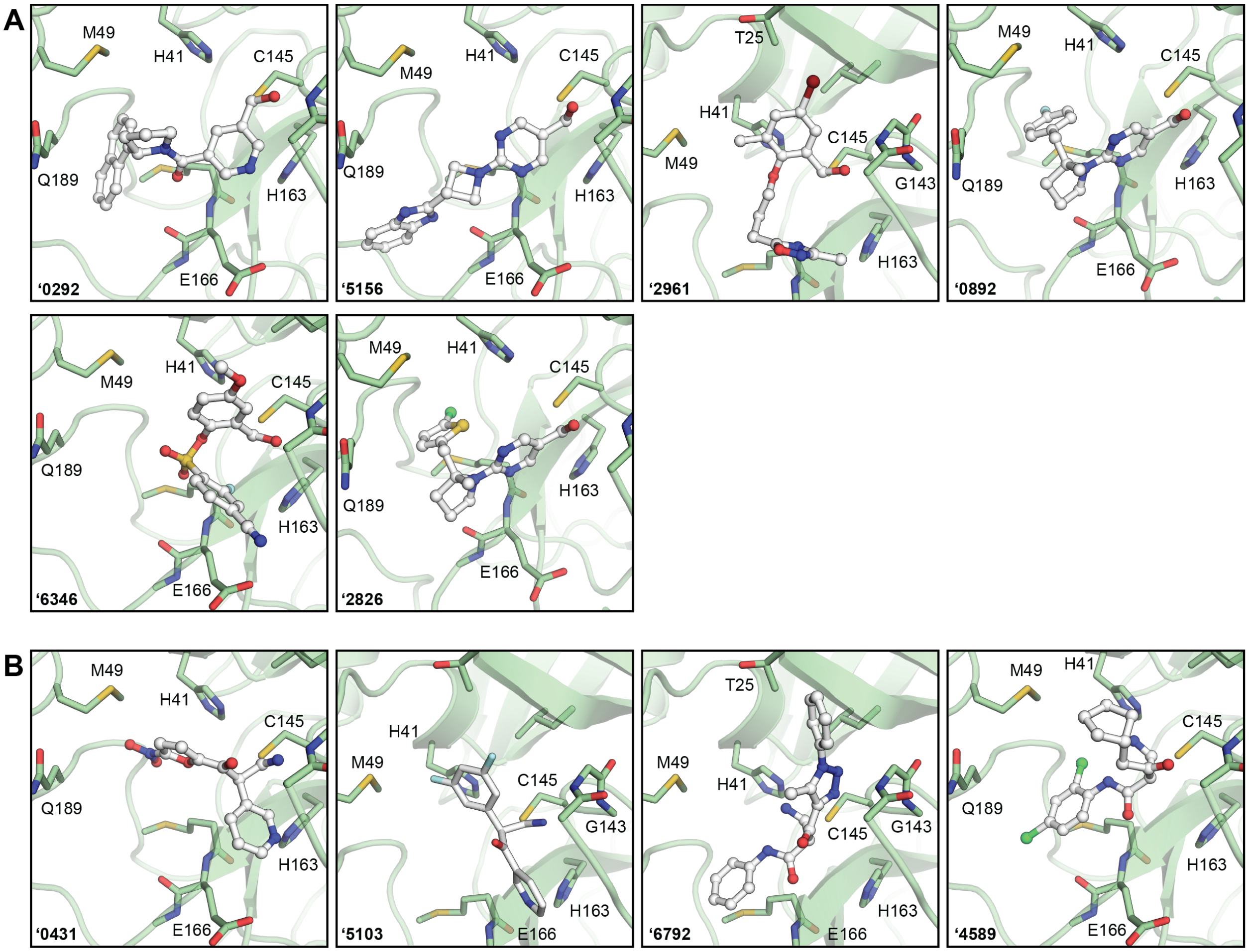


**Supplementary Figure 5. Docked poses of covalent hits.** (**A**) Aldehydes. (**B**) Nitriles. Covalent bonds between the warhead and Cys145 and not explicitly modeled in docking.

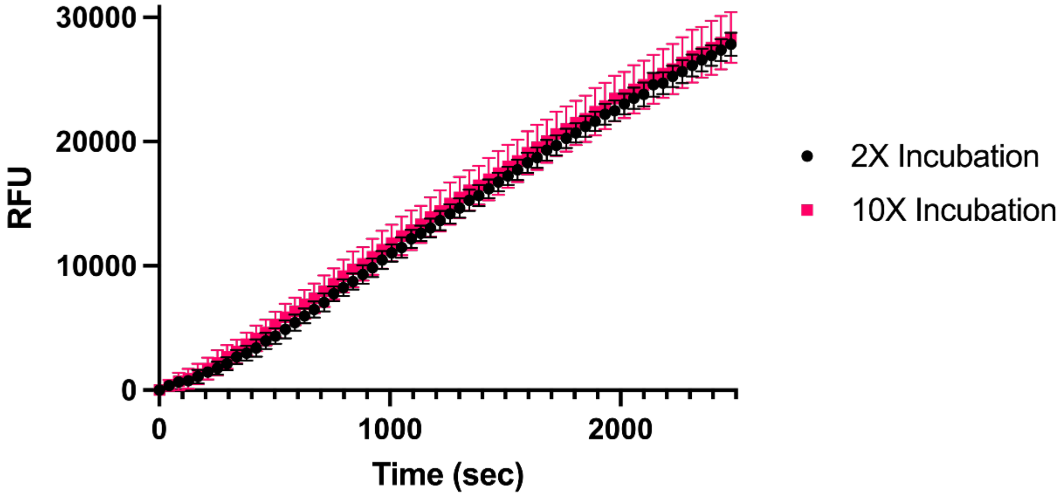


**Supplementary Figure 6. Reversibility of compound '7021.** Both 2X and 10X concentrations of enzyme and '7021 inhibitor were co-incubated for 1-hour. By Le Chatelier's princiciple, the 10X incubation should lead to more inhibited enzyme product. Both 2X and 10X incubations were diluted to 1X concentrations with final substrate concentration of 1X NSP7. Both co-incubations yielded the same velocity, evincing concentration dependent diffusion of the enzyme-inhibitor complex, thus evincing the reversibility of the '7021 compound.

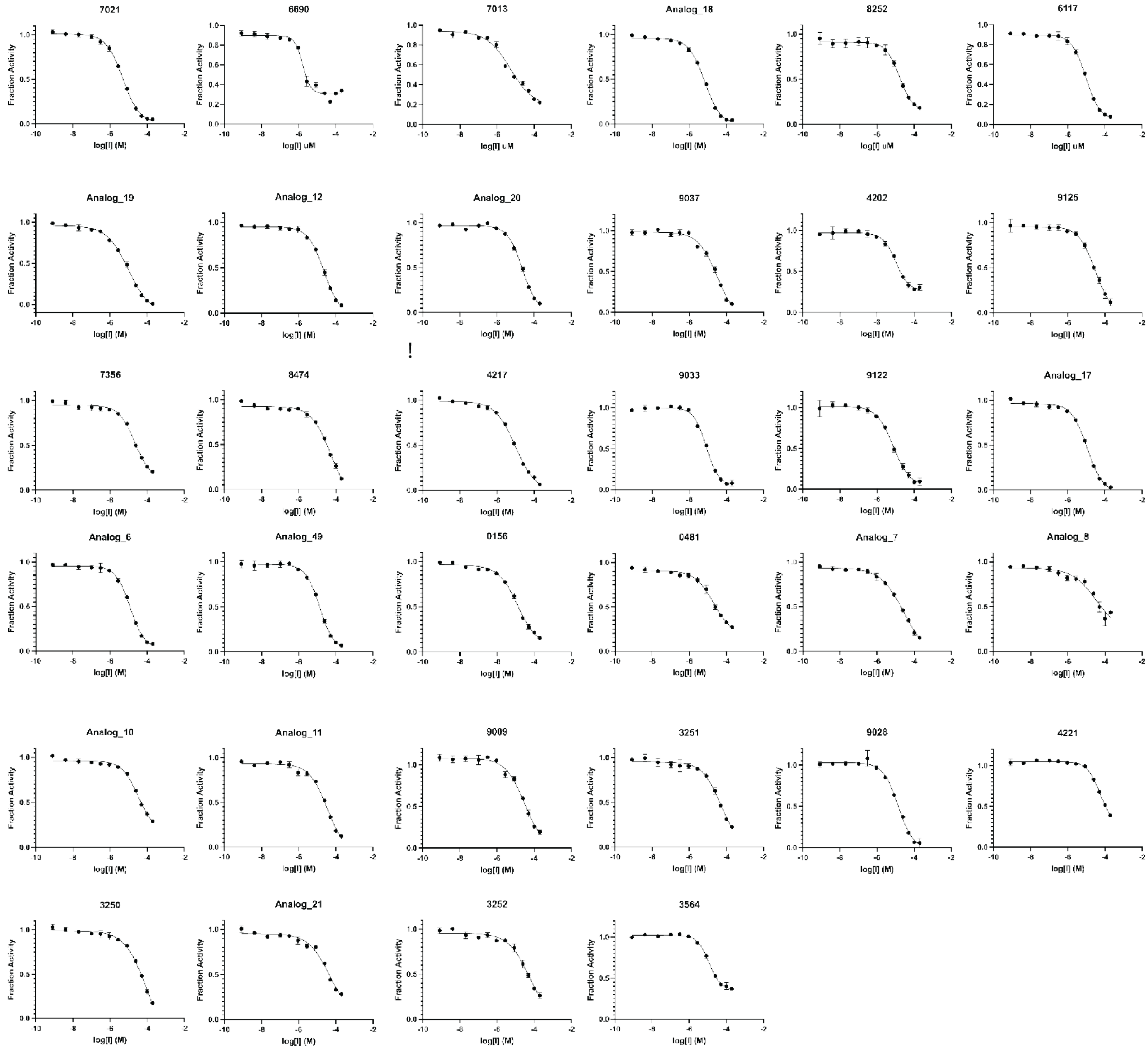


**Supplementary Figure 7. '3620 analogs with improved potencies.** $IC_{50}$ values in **Table S2**.

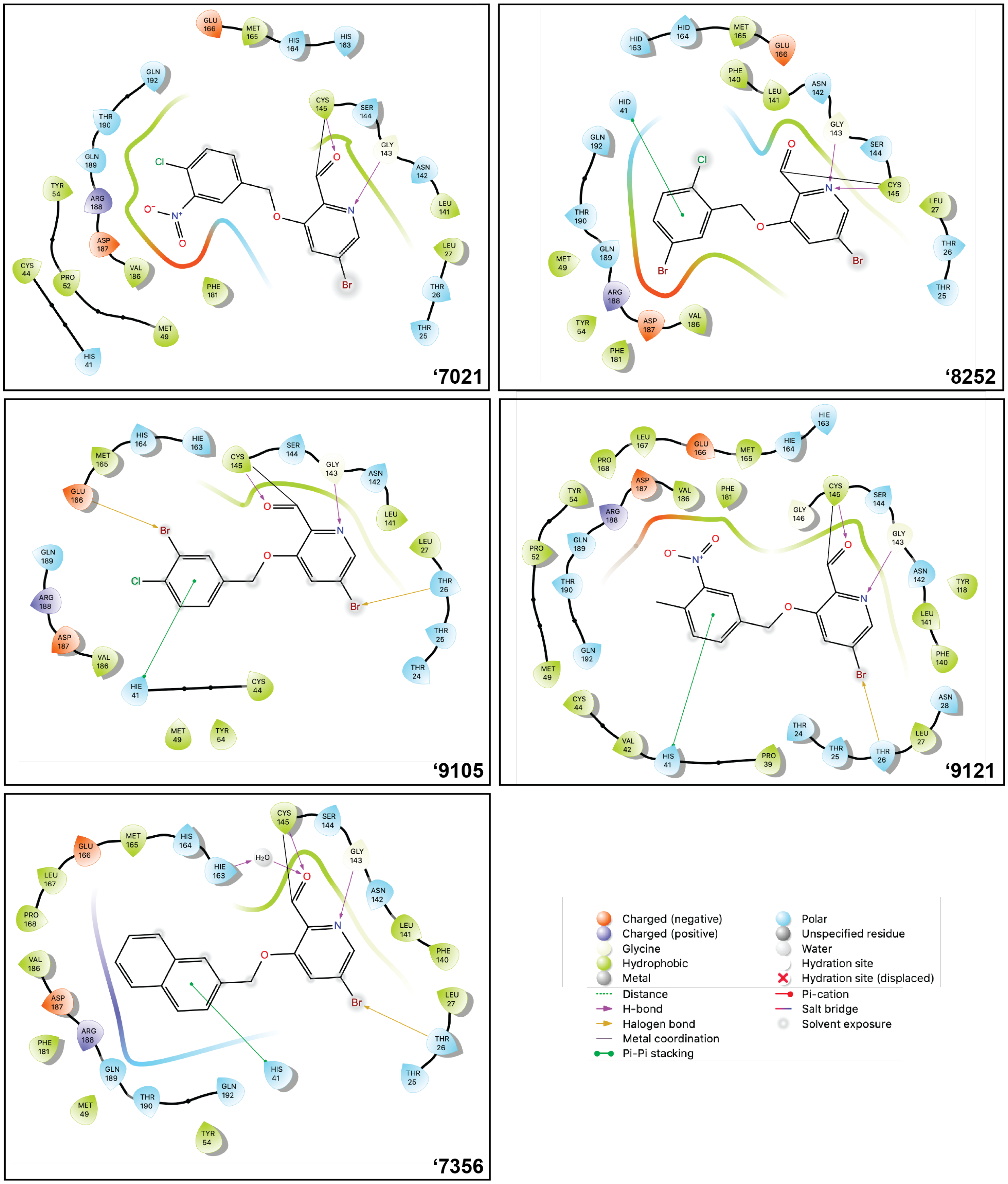


**Supplementary Figure 8. LigPlot visualization of MPro-covalent inhibitor interactions in newly solved structures.**

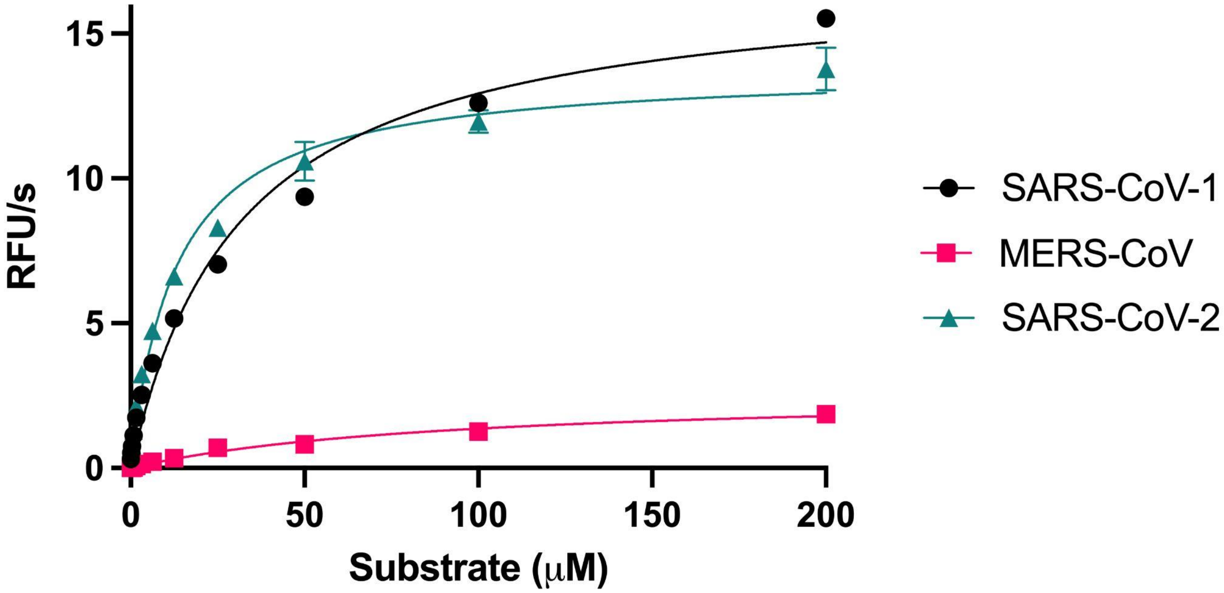


**Supplementary Figure 9. Pan-viral enzymatic activities.** Michaelis-Menten kinetics were ran side-by-side with the optimized NSP7 substrate for SARS-CoV-2 (green), SARS-CoV-1 (black), and MERS (red) major proteases. $K_M$ values were found to be 12μM, 30μM, and 90μM for SARS-CoV-2, SARS-CoV-1, and MERS, respectively. All proceeding inhibitory assays were ran with $K_M$ concentrations of substrate.

MaxPeak: 100.00%
Ret_Time: 0.954 min

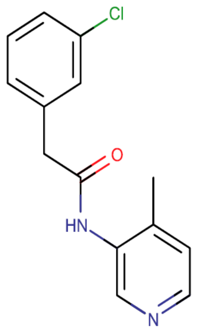


**Mol Wt** 260.72

**Exact Mass** 260.09

| # | Time | Area% |
|---|---|---|
| 1 | 0.954 | 100.00 |

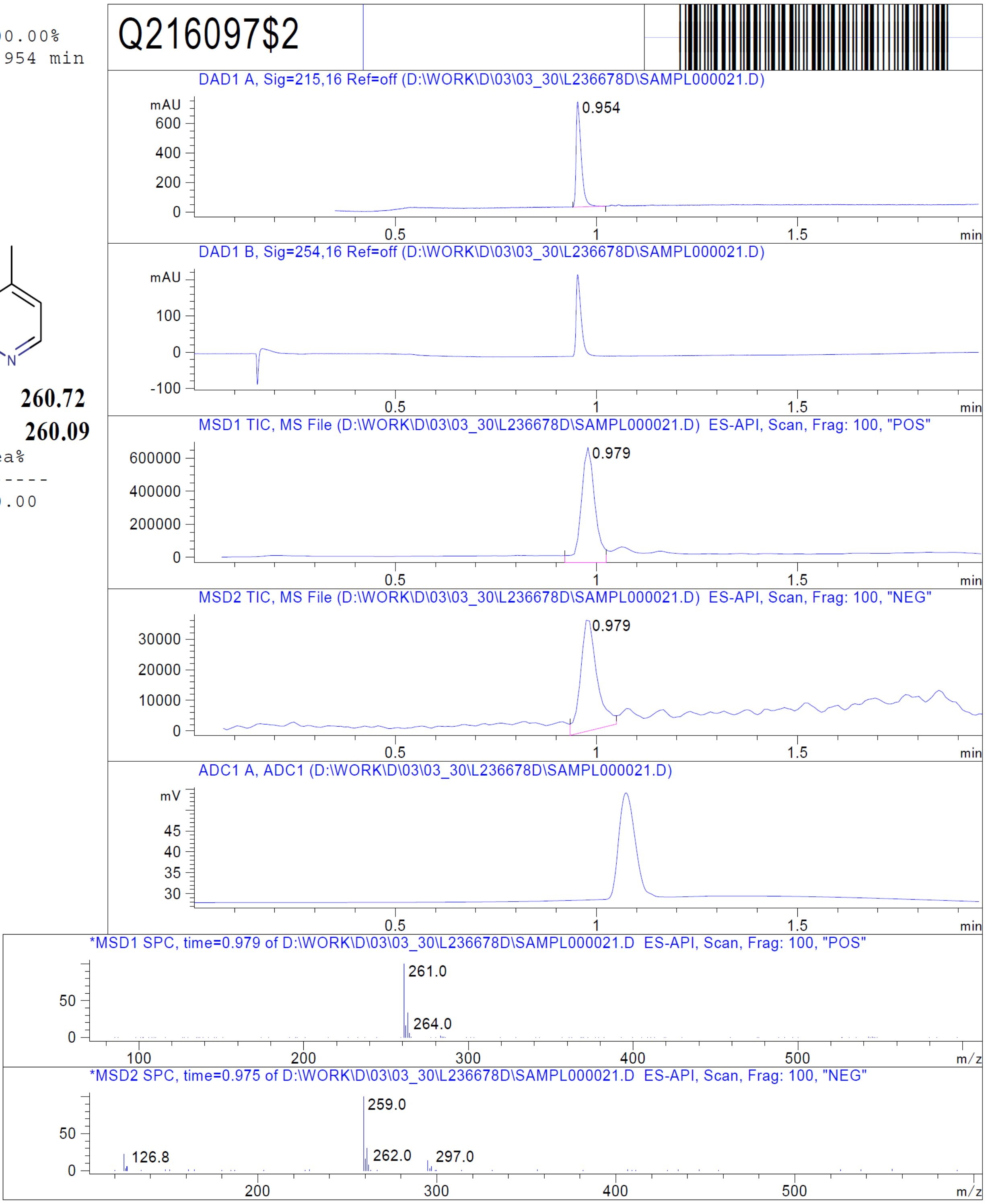


RT 0.979

RT 0.979

Z1129289650

Z1129289650

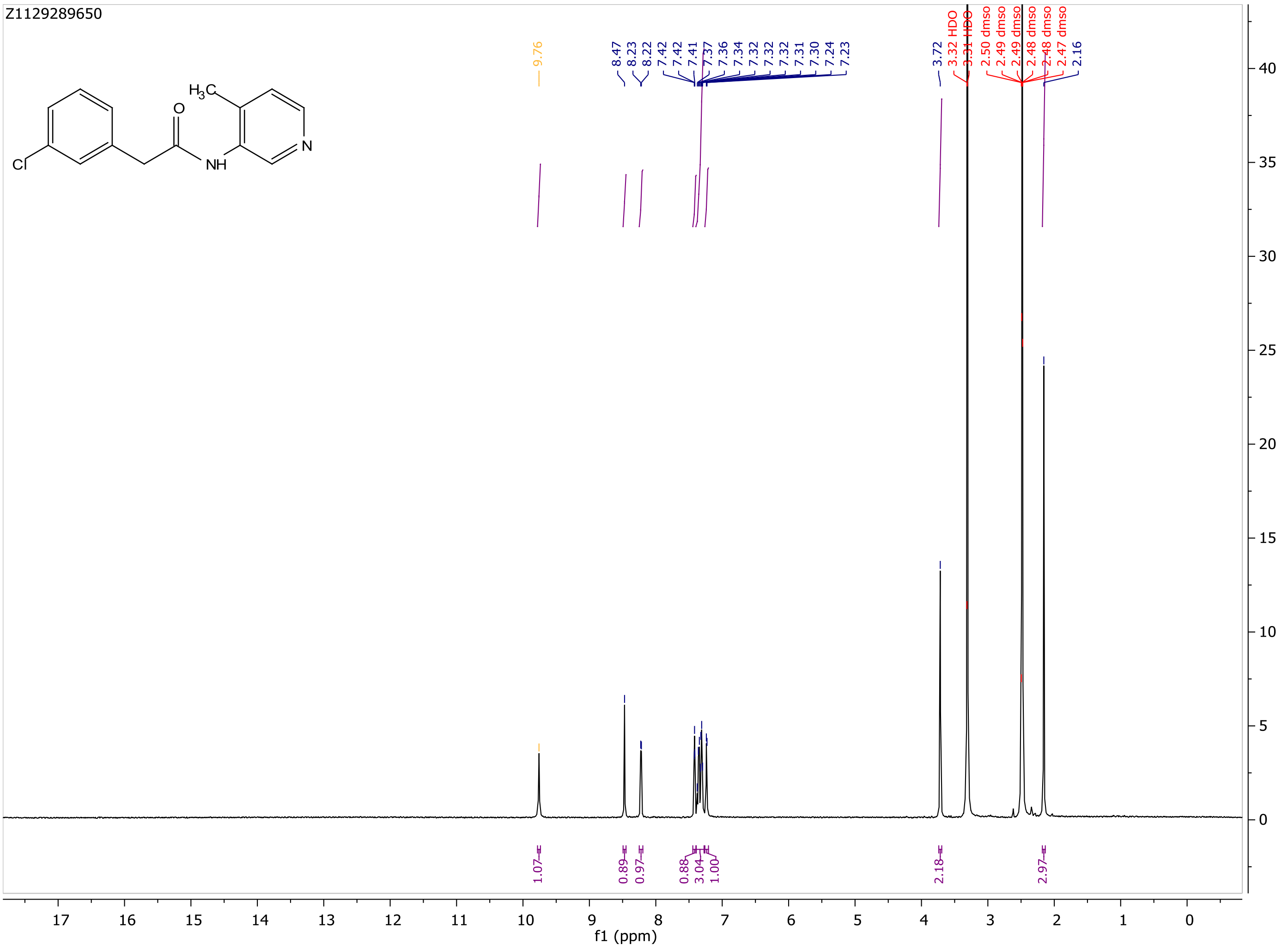

MaxPeak: 100.00%
Ret_Time: 0.887 min

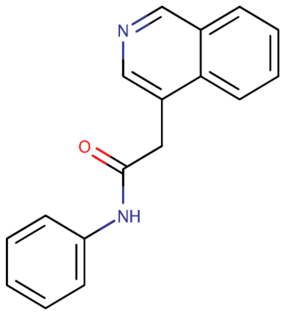


**Mol Wt** 262.31

**Exact Mass** 262.13

| # | Time | Area% |
|---|---|---|
| 1 | 0.887 | 100.00 |

# Q272221$1

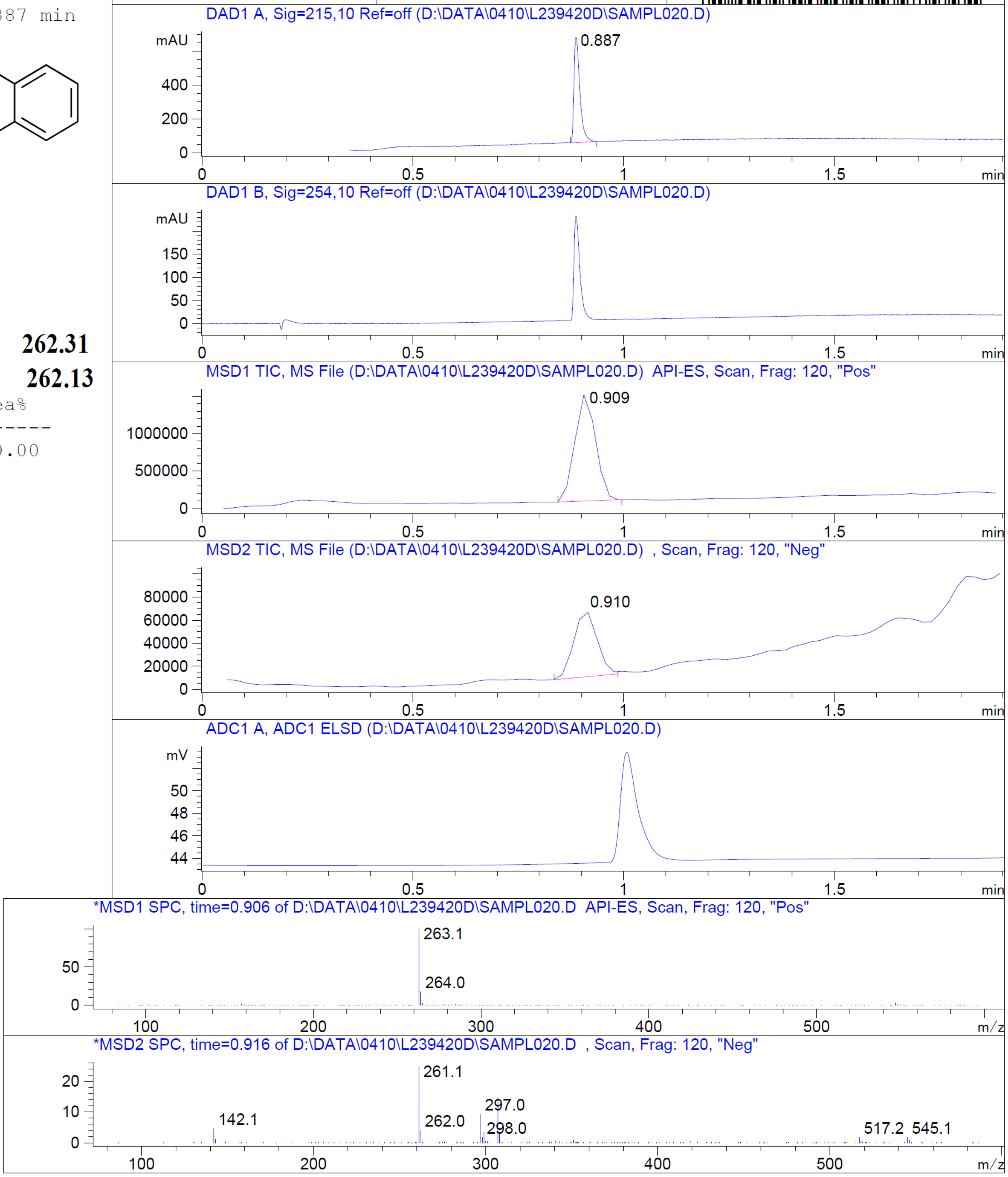


RT 0.909

RT 0.910

Z4444622066

Z4444622066

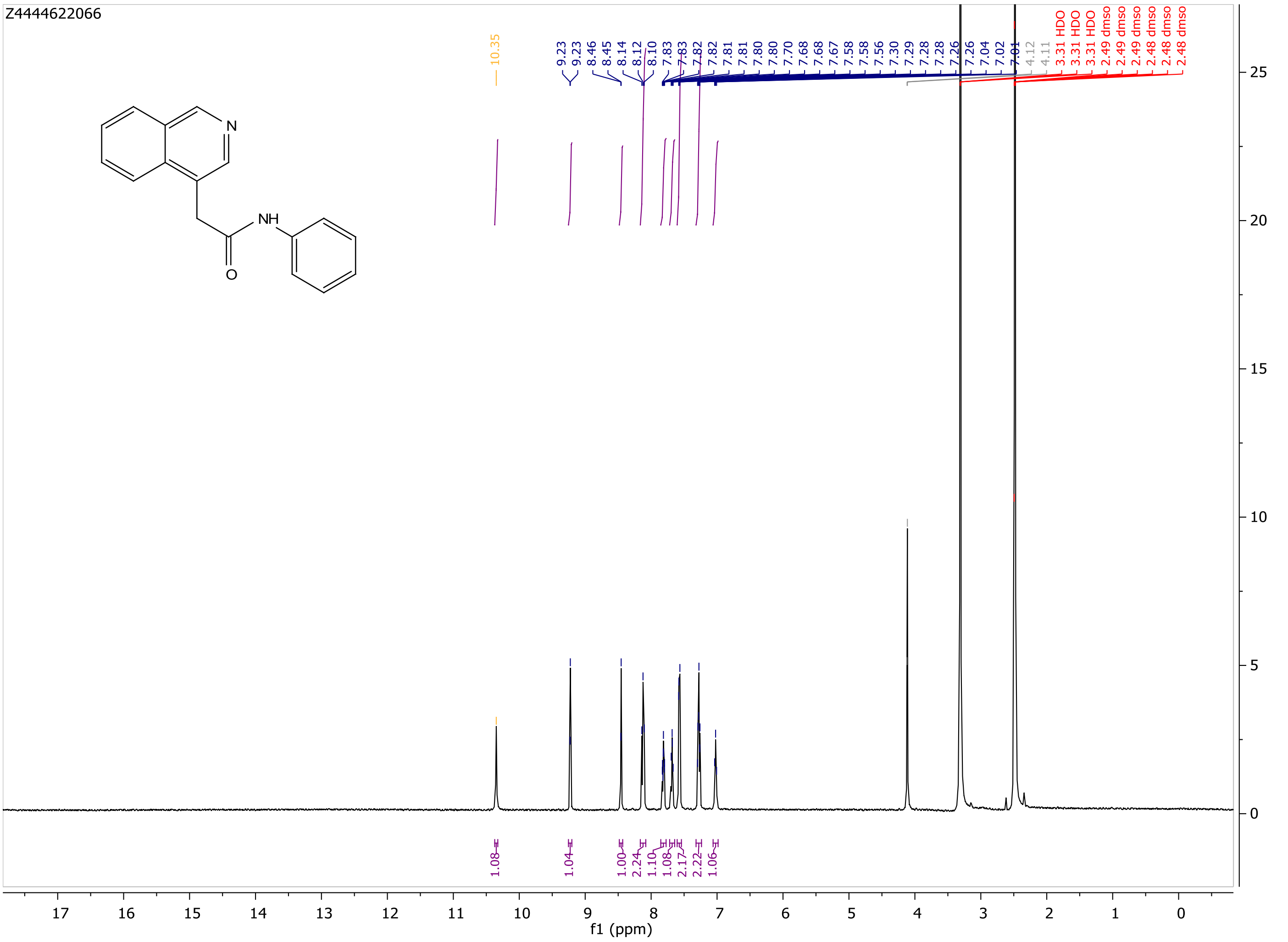

MaxPeak: 100.00%
Ret_Time: 1.173 min

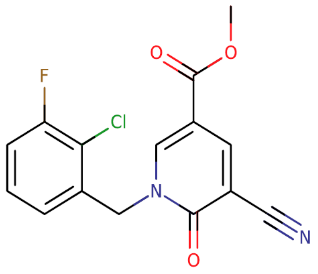


**Mol Wt** 320.7

**Exact Mass** 320.04

| # | Time | Area% |
|---|---|---|
| 1 | 1.173 | 100.00 |

Q227951$2

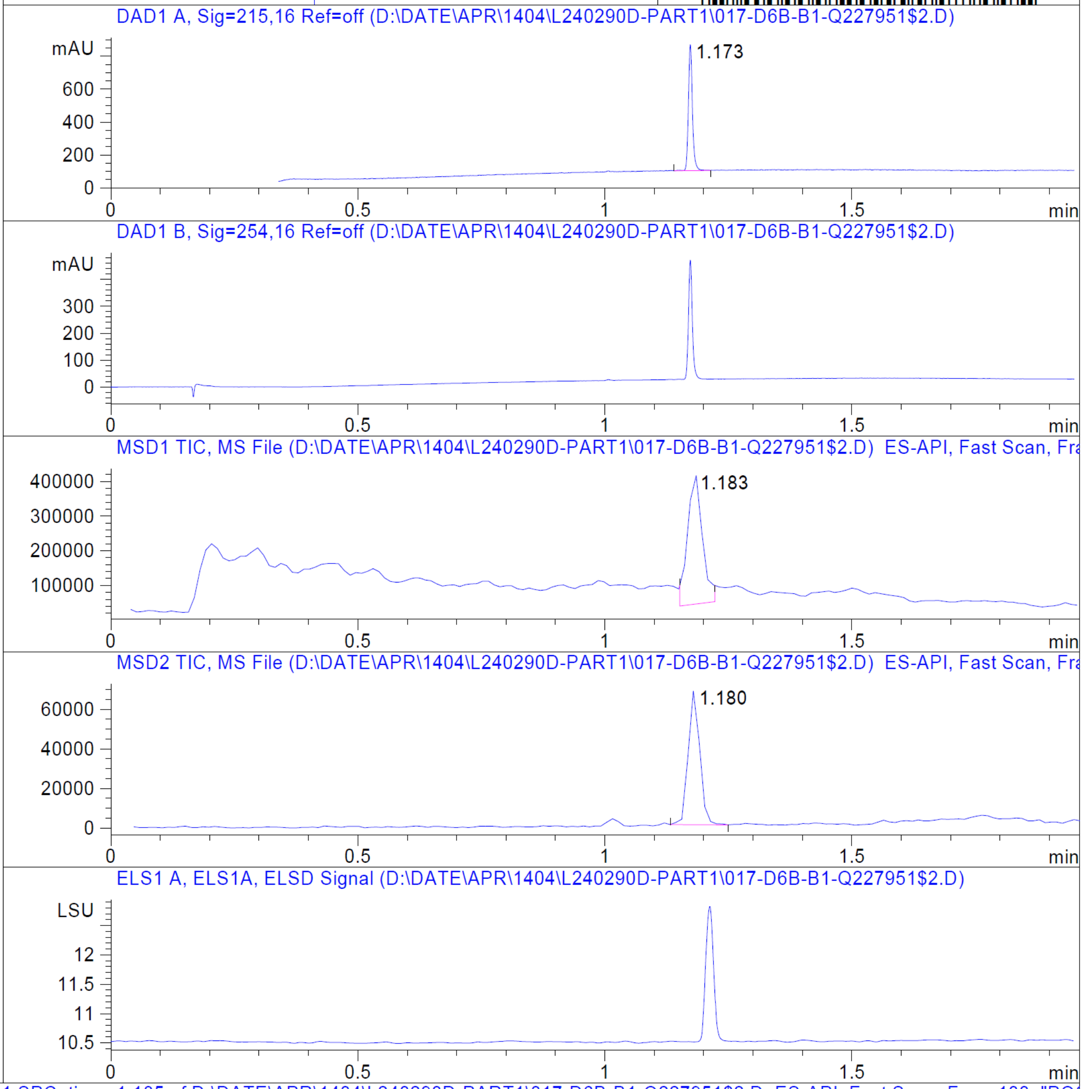


RT 1.183

RT 1.180

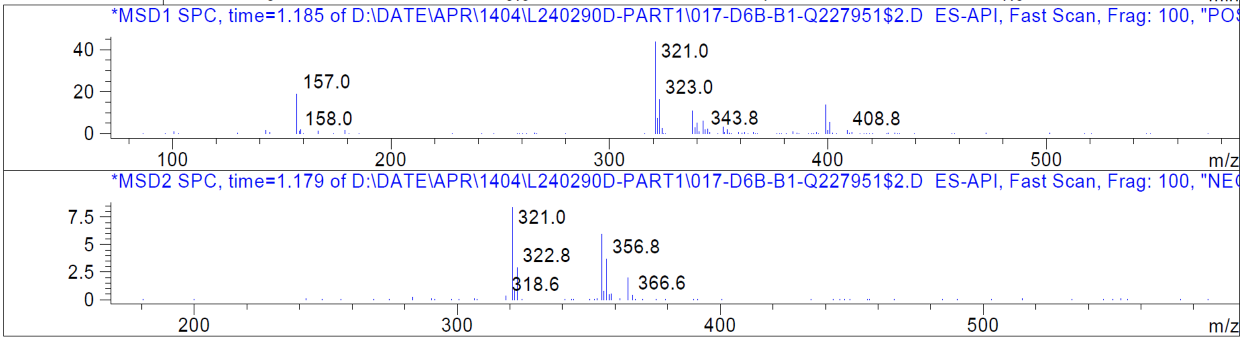


Z1578697783_(ZINC000338540162)

Z1578697783_(ZINC000338540162)

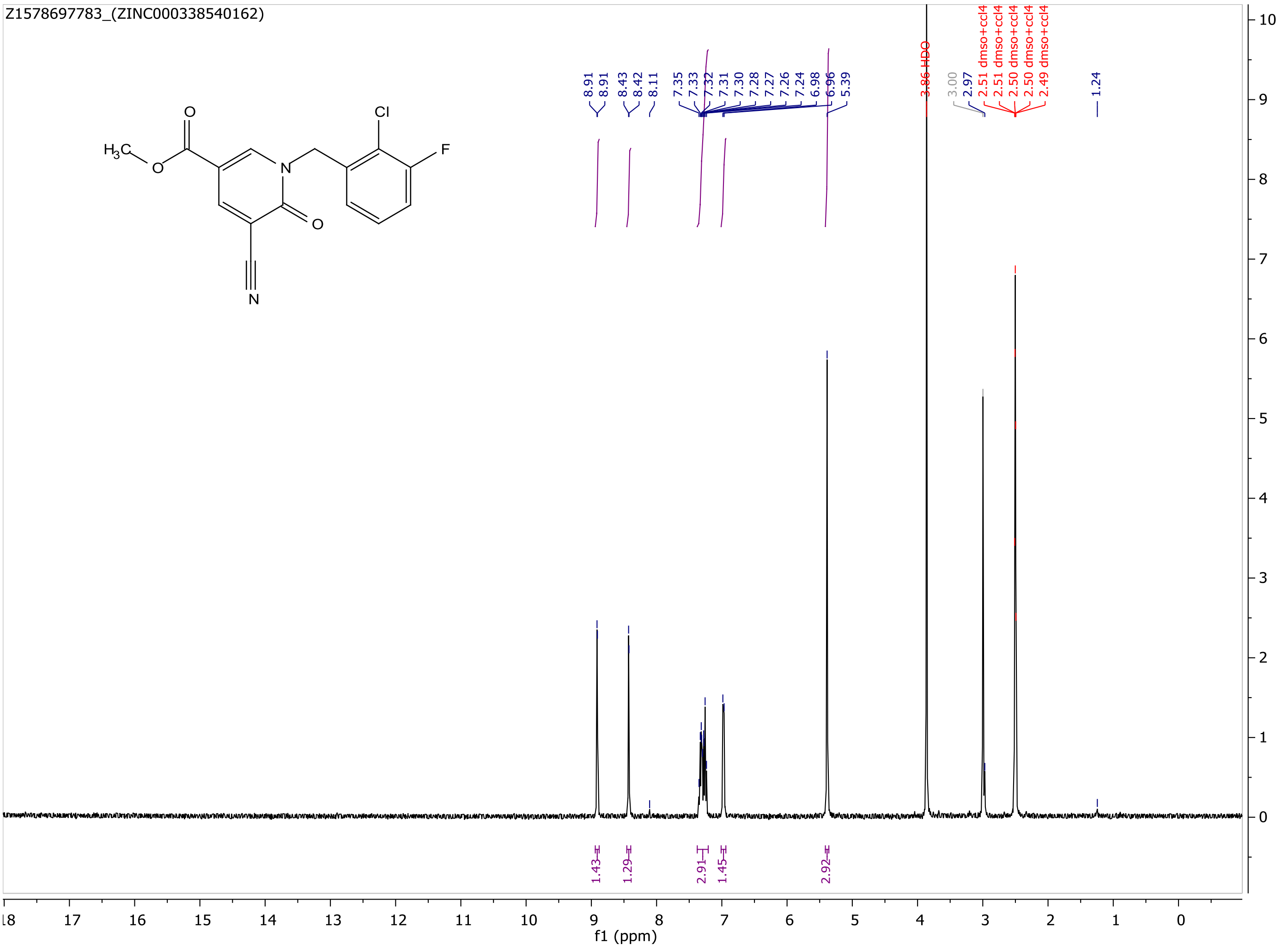

MaxPeak: 100.00%
Ret_Time: 1.333 min

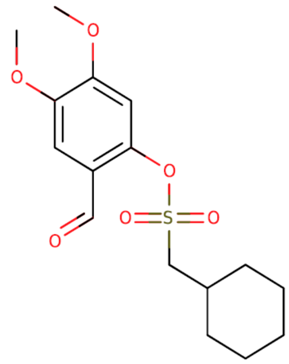

**Mol Wt** 342.41
**Exact Mass** 342.13

| # | Time | Area% |
|---|---|---|
| 1 | 1.333 | 100.00 |

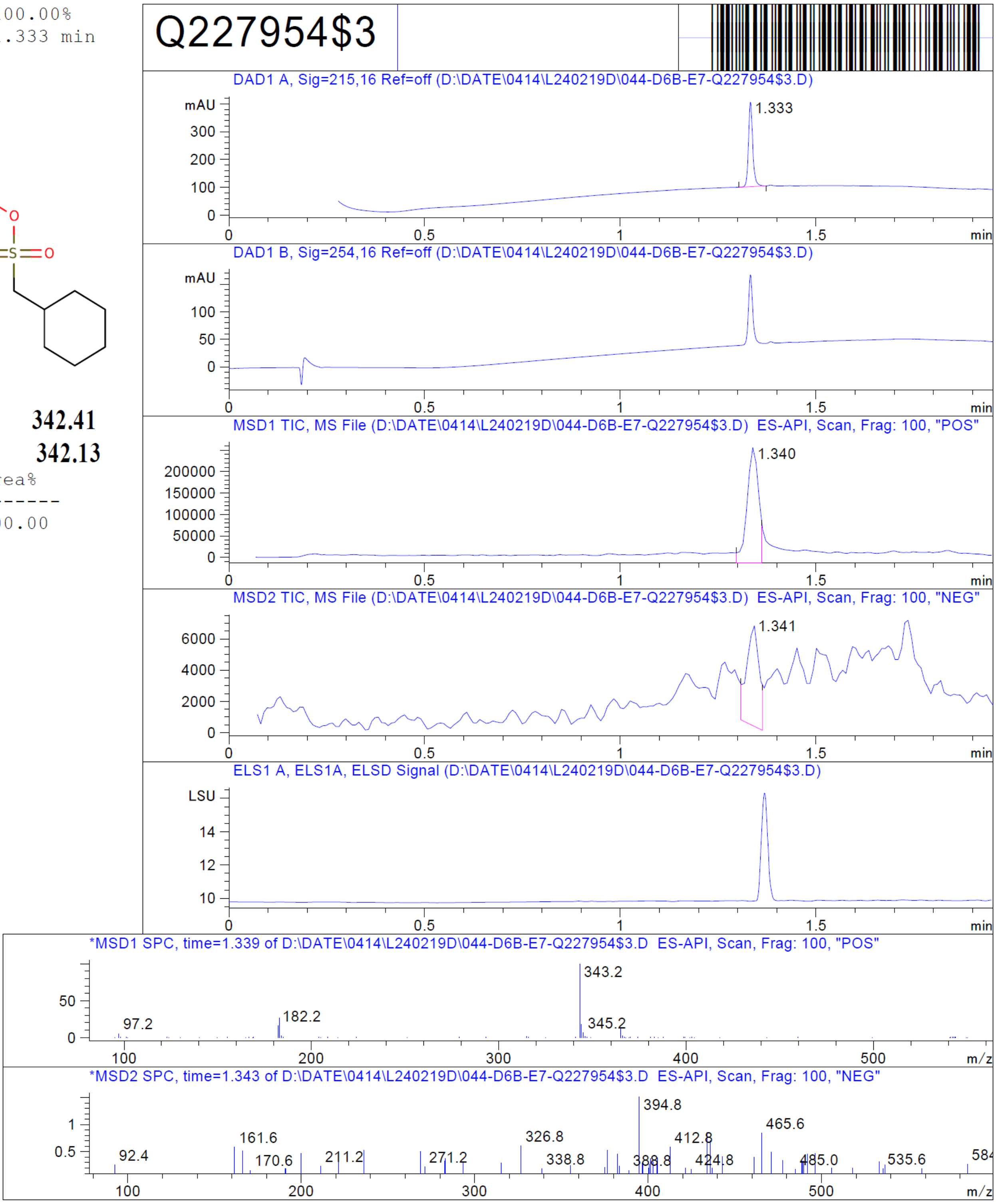


RT 1.340

RT 1.341

Z1564333798_(ZINC000271072260)

Z1564333798_(ZINC000271072260)

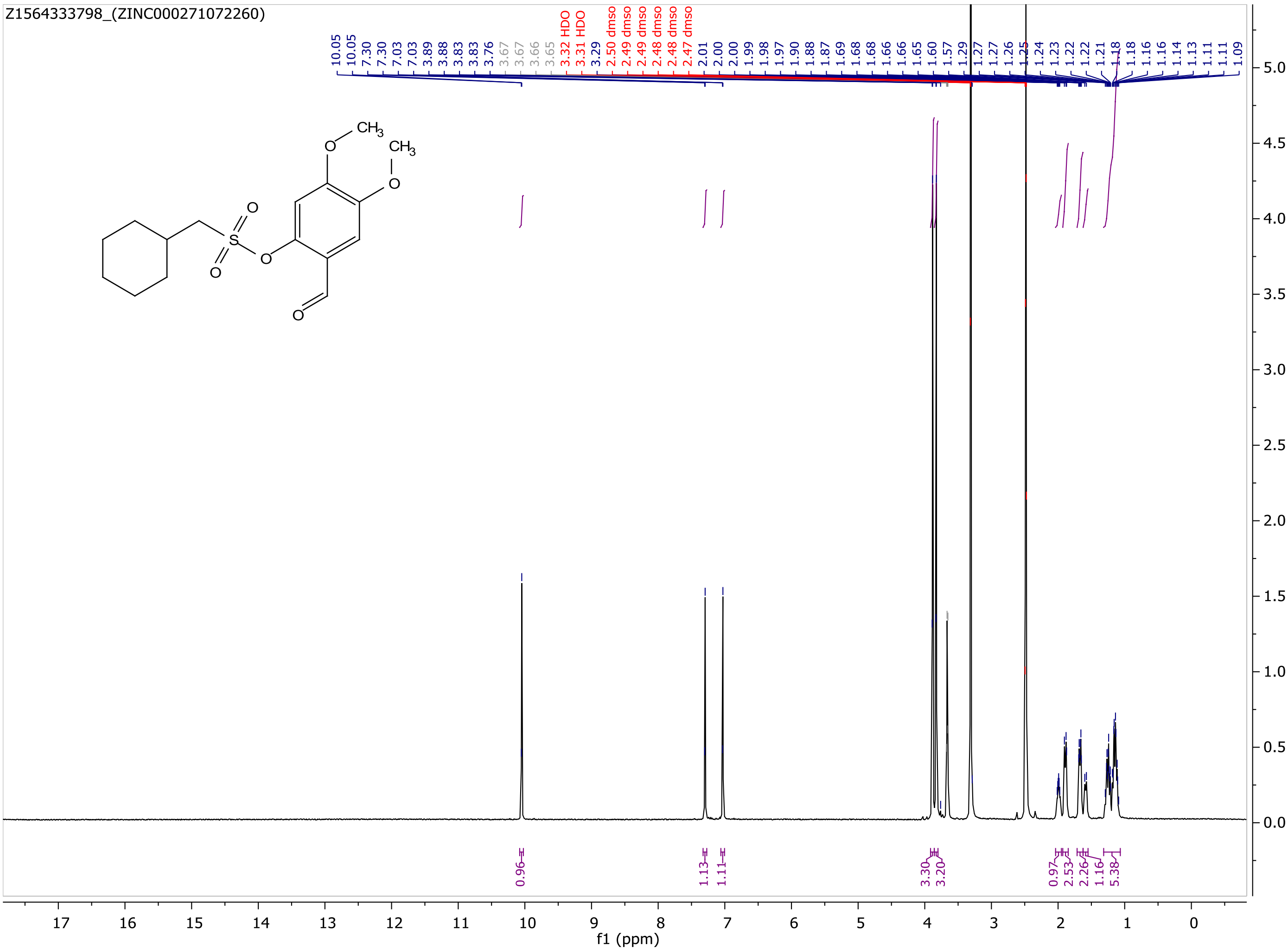

MaxPeak: 91.00%
Ret_Time: 1.297 min

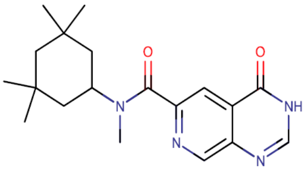

**Mol Wt** 342.44
**Exact Mass** 342.24

| # | Time | Area% |
|---|---|---|
| 1 | 1.297 | 91.00 |
| 2 | 1.448 | 9.00 |

Q228021$4

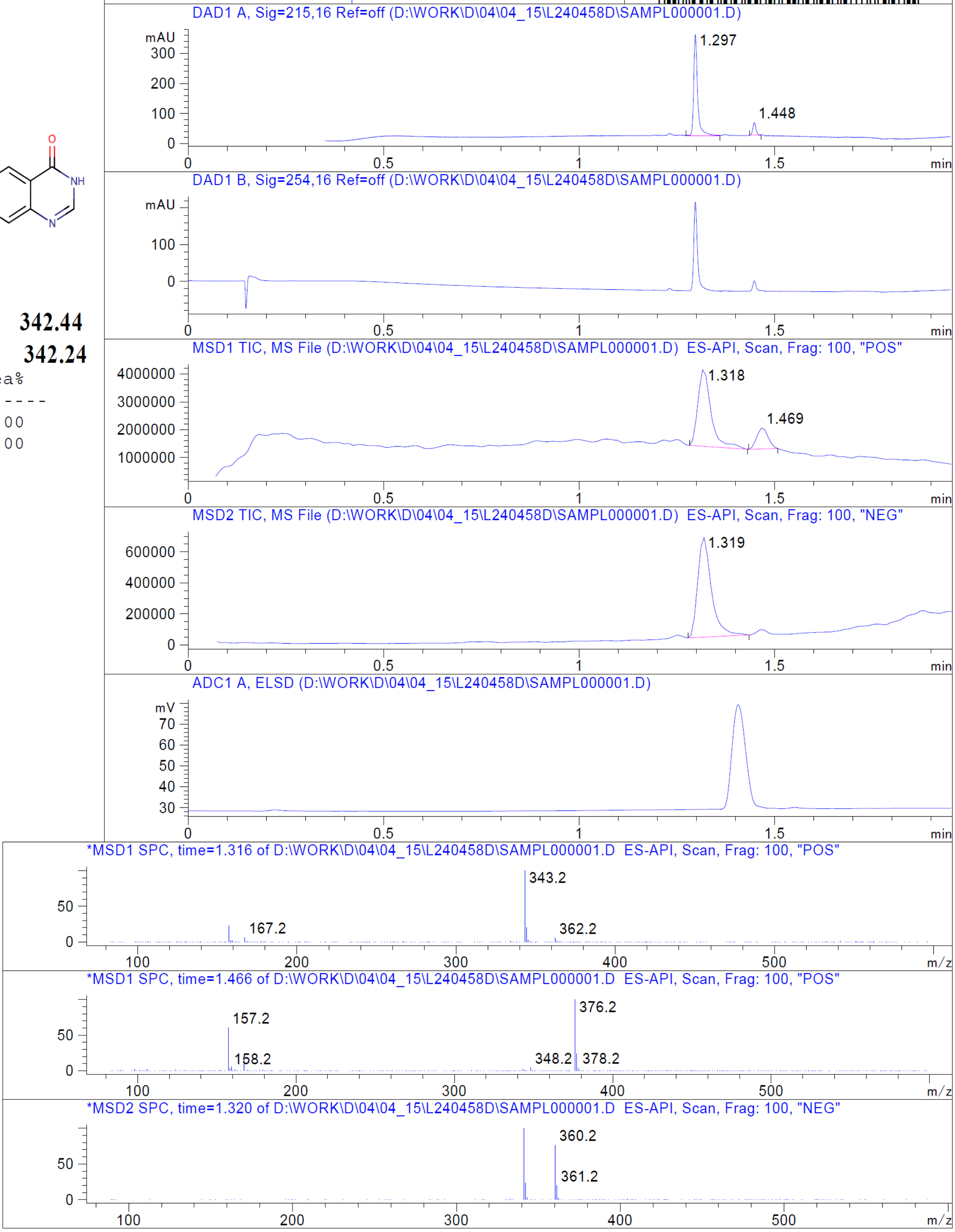


RT 1.318

RT 1.469

RT 1.319

Z2644709353_(ZINC000618071006)

Z2644709353_(ZINC000618071006)

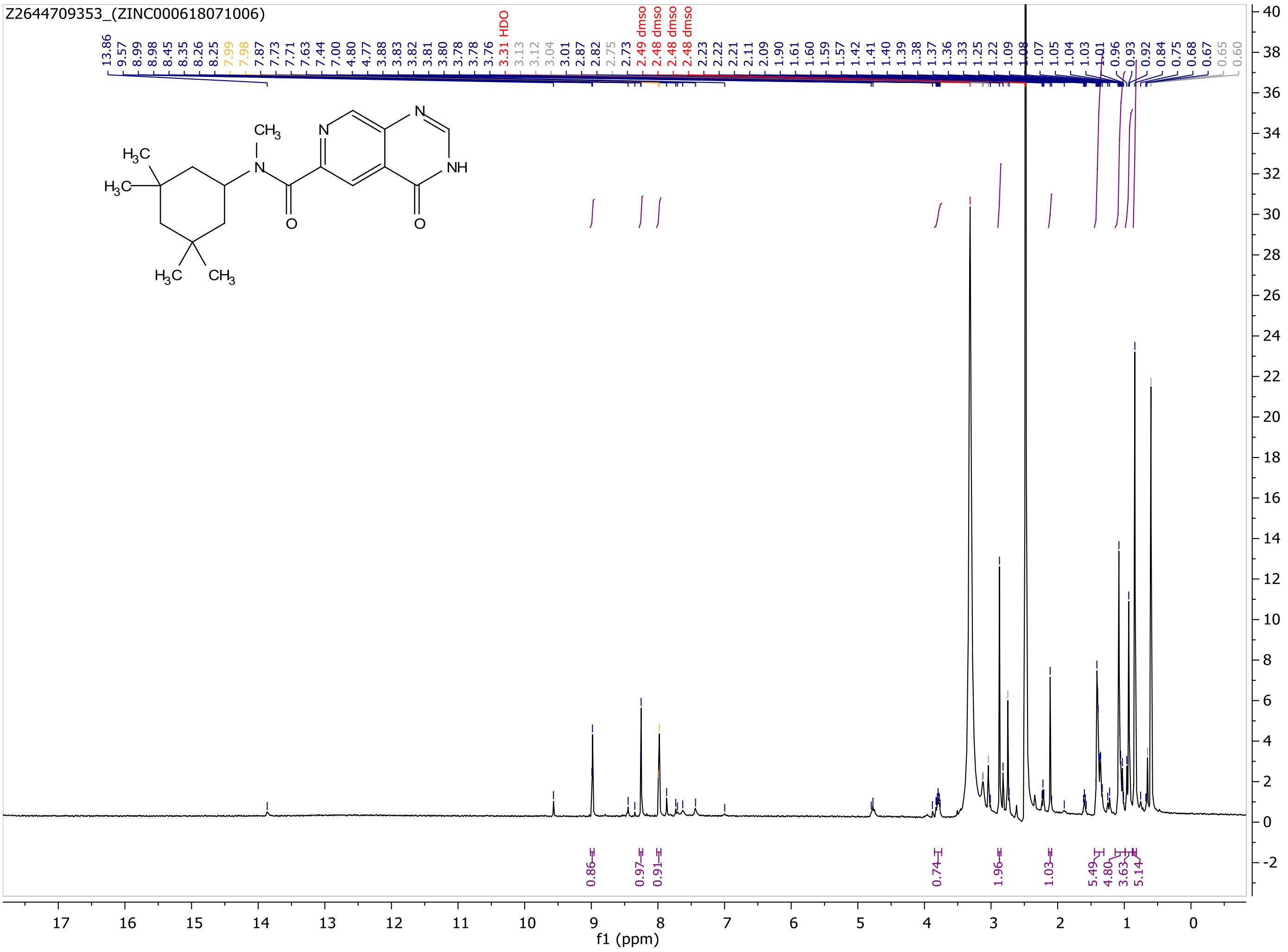

MaxPeak: 100.00%
Ret_Time: 0.736 min

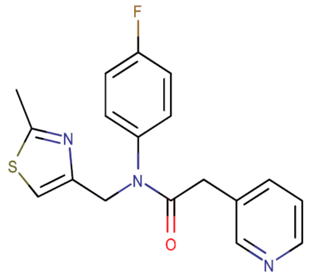

**Mol Wt** 341.4
**Exact Mass** 341.12

| # | Time | Area% |
|---|---|---|
| 1 | 0.736 | 100.00 |

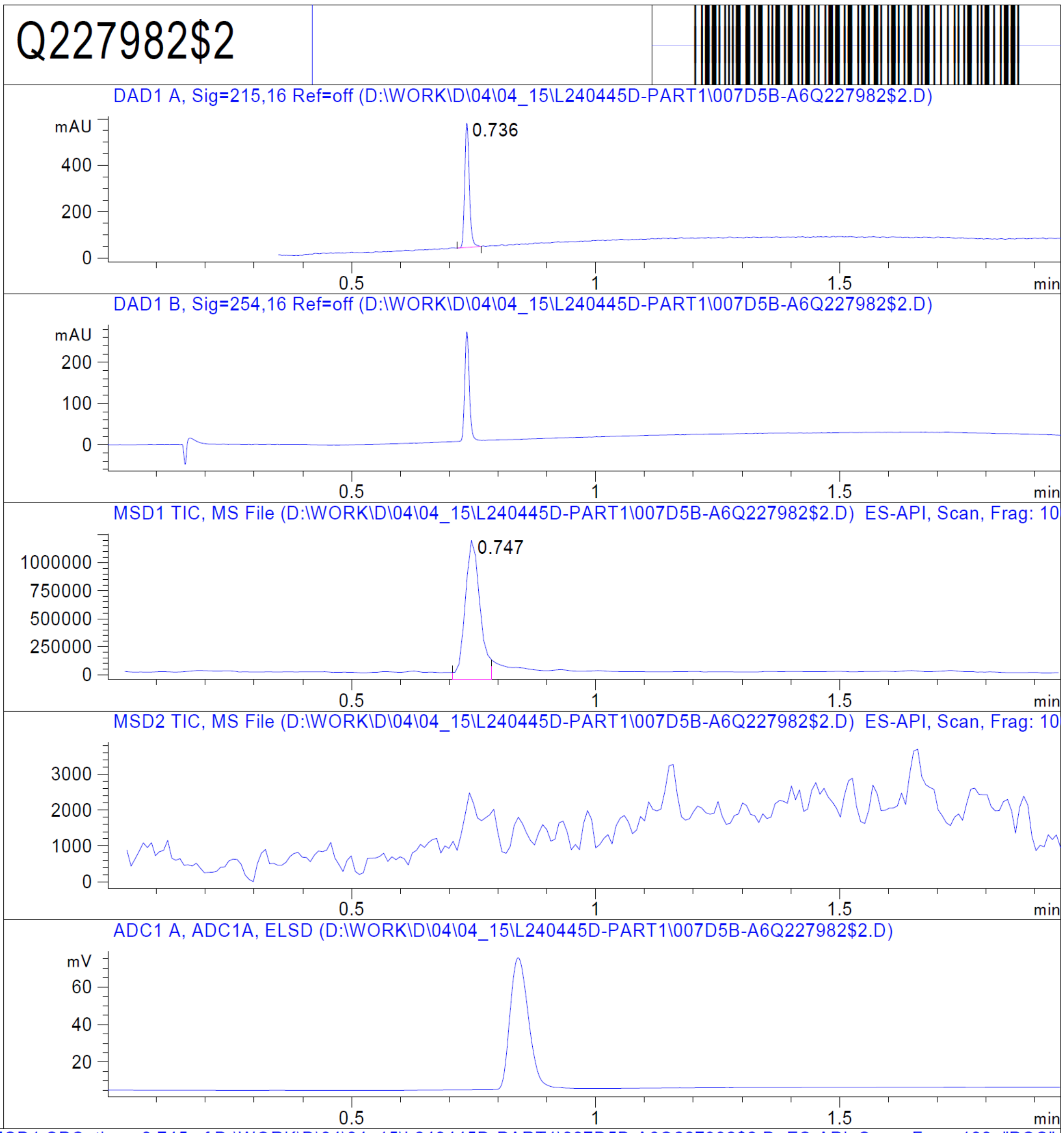


RT 0.747

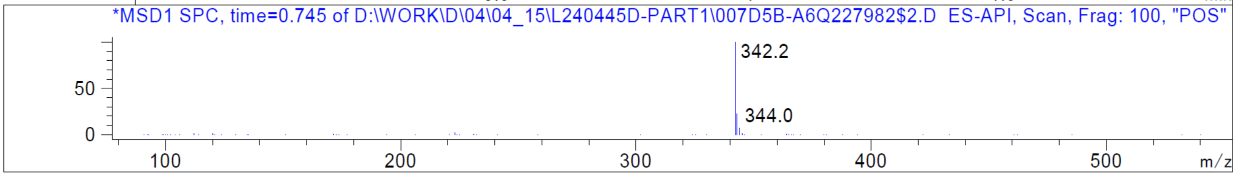


Z11713367722_(ZINC000346368145)

Z1171336722_(ZINC000346368145)

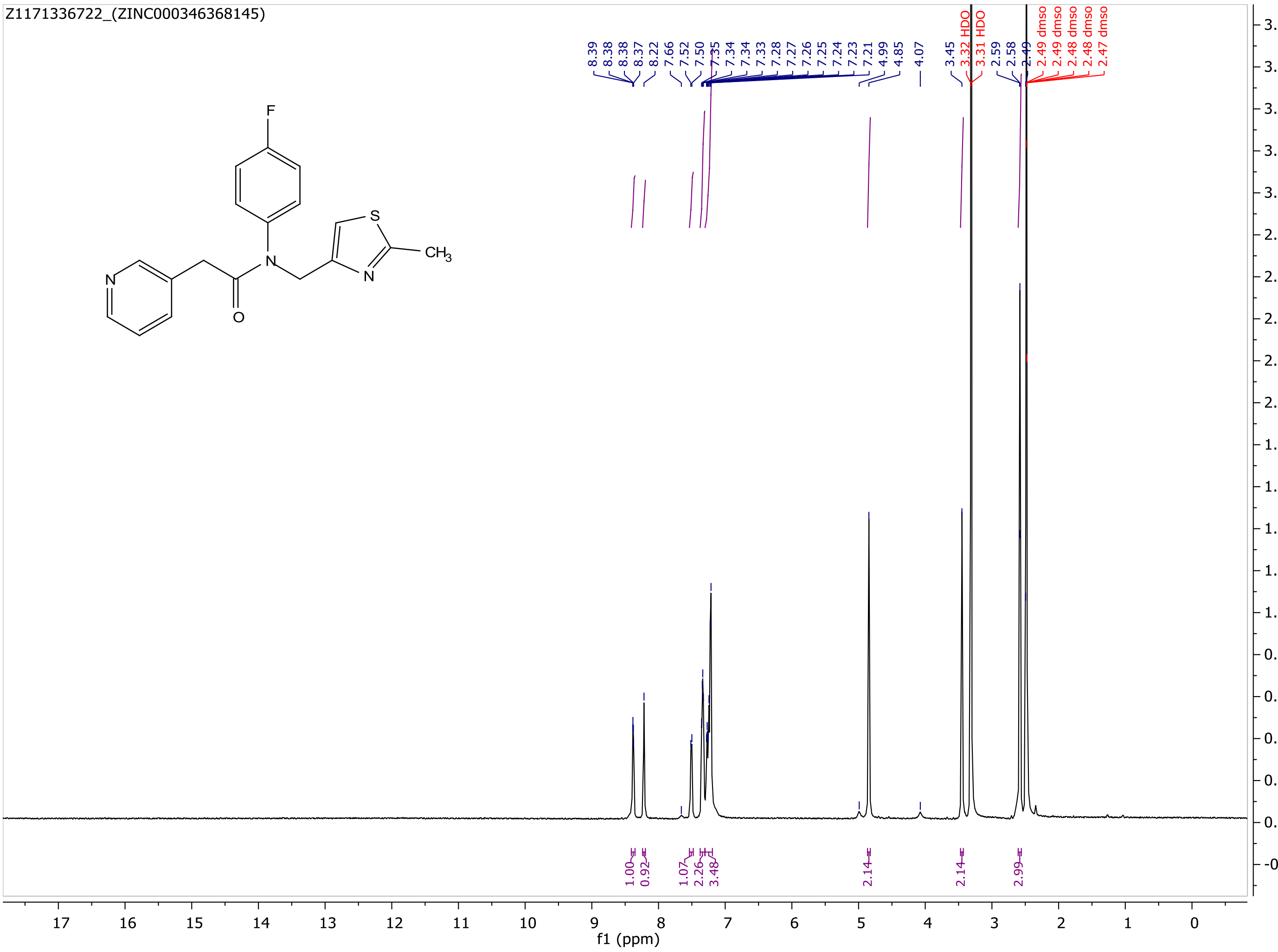

Q227994$1

MaxPeak: 100.00%
Ret_Time: 0.886 min

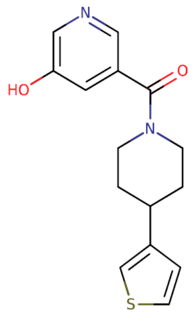


**Mol Wt** 288.36

**Exact Mass** 288.11

| # | Time | Area% |
|---|---|---|
| 1 | 0.886 | 100.00 |

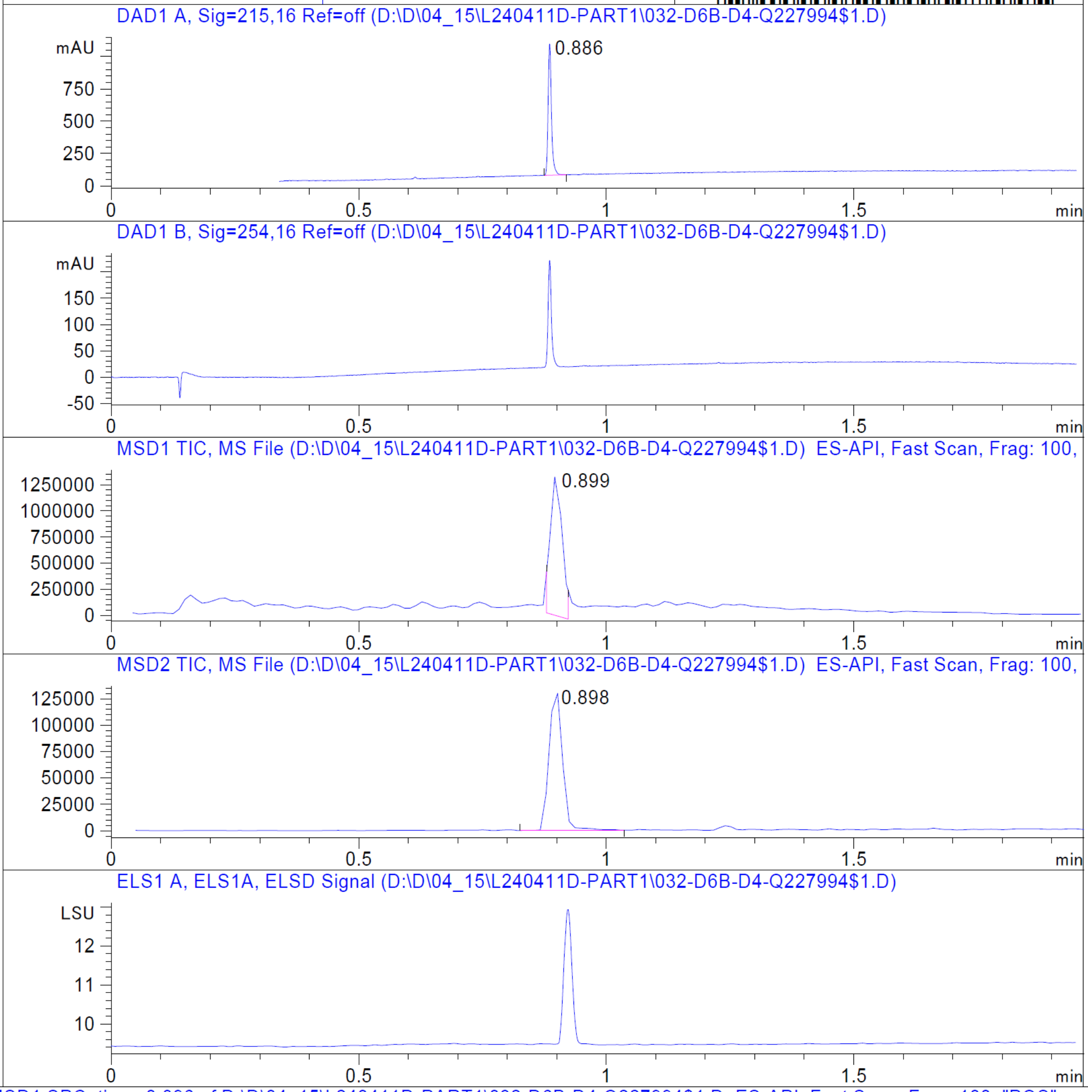


RT 0.899

RT 0.898

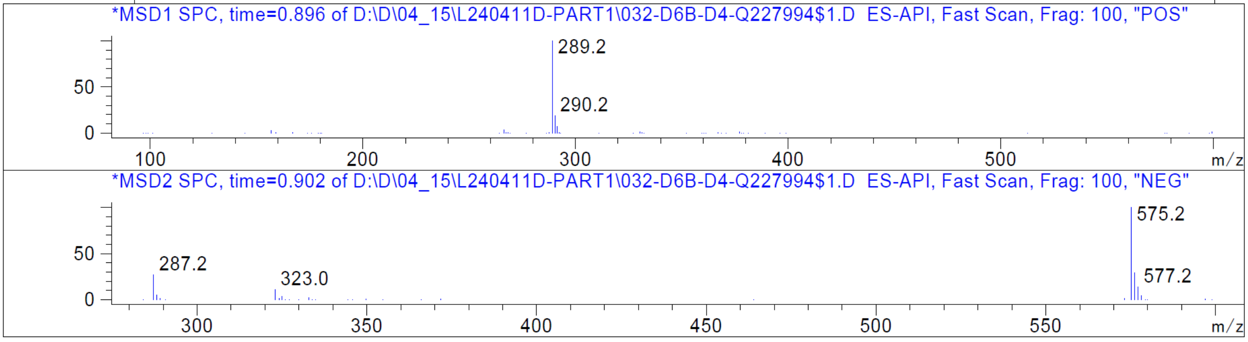


Z1740796453_(ZINC000188718998)

Z1740796453_(ZINC000188718998)

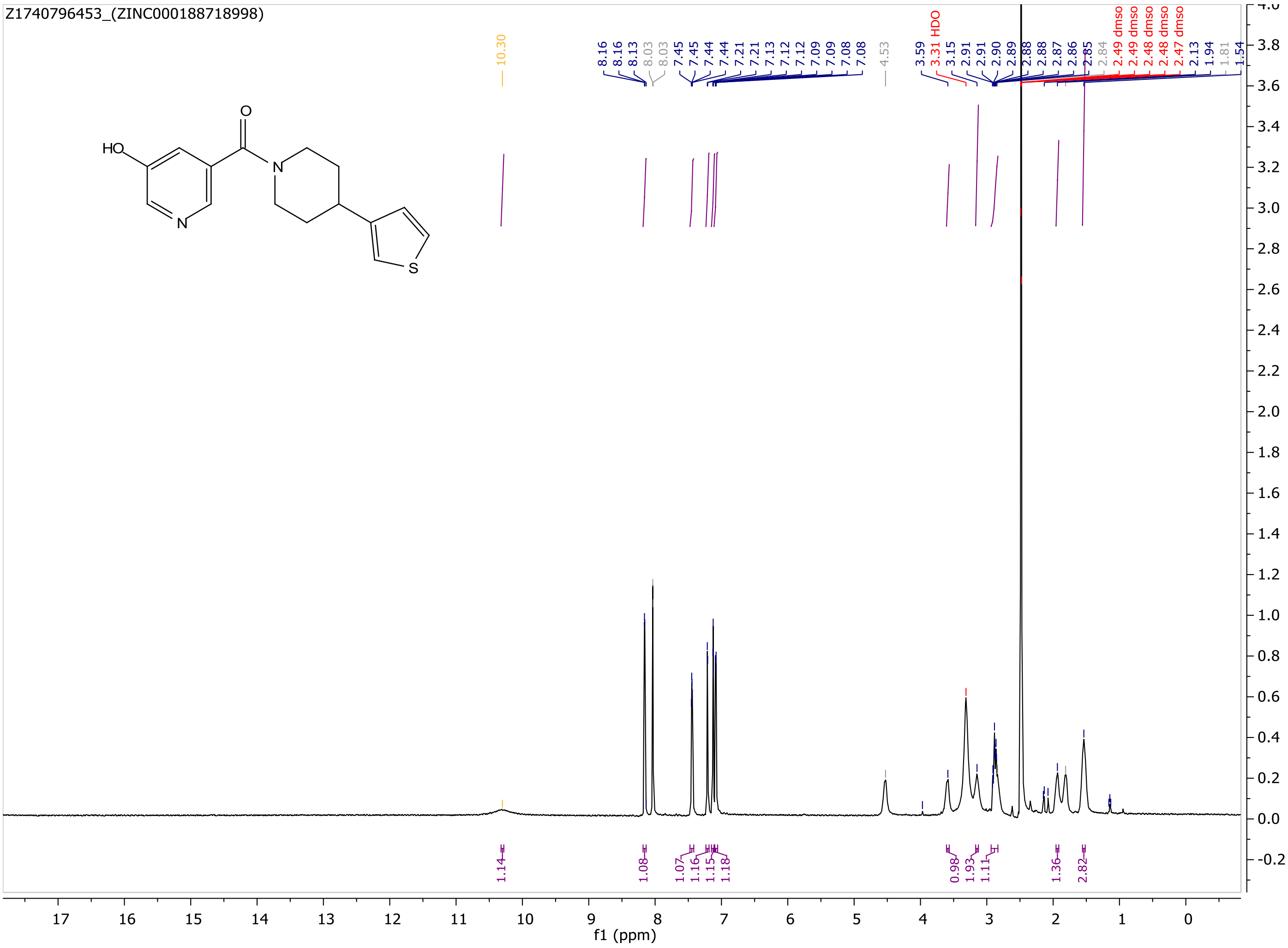

MaxPeak: 100.00%
Ret_Time: 0.772 min

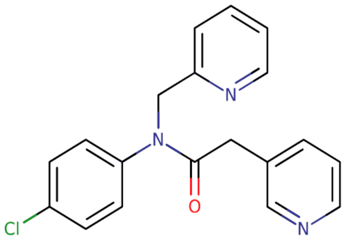

**Mol Wt** 337.8
**Exact Mass** 337.12

| # | Time | Area% |
|---|---|---|
| 1 | 0.772 | 100.00 |

Q227995$4

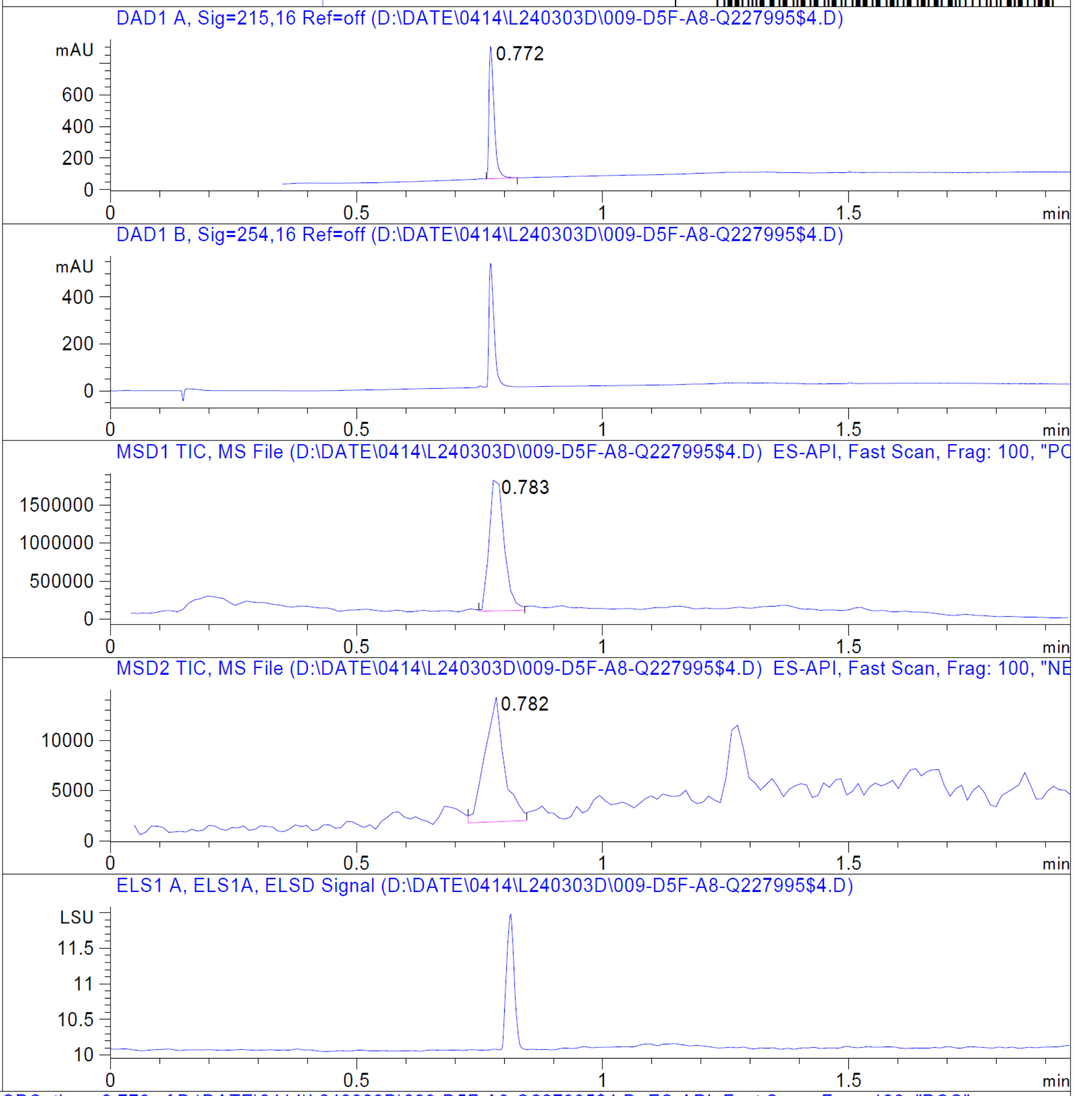


RT 0.783

RT 0.782

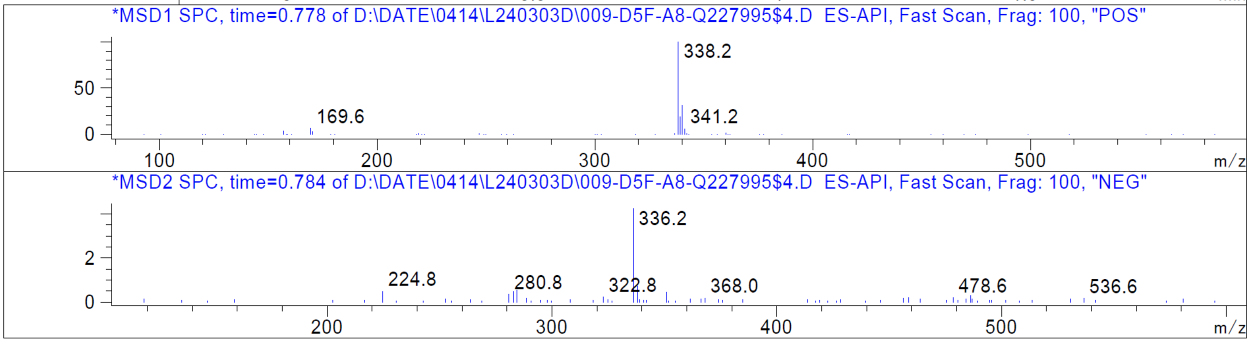


Z1171337460_(ZINC000346371112)

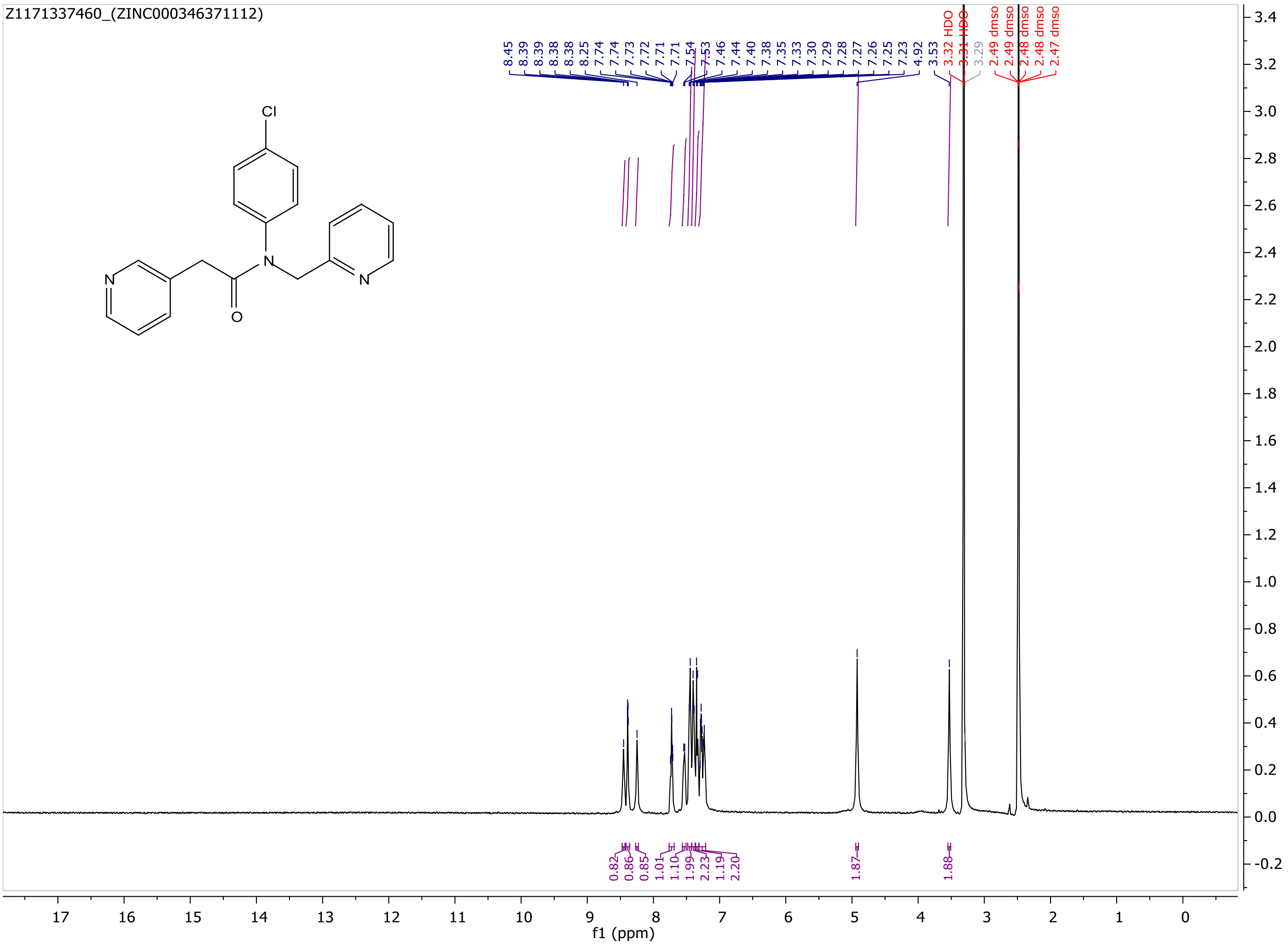
Z1171337460_(ZINC000346371112)
Cl
N
N
O
N
8.45
8.39
8.39
8.38
8.38
8.25
7.74
7.74
7.73
7.72
7.71
7.54
7.53
7.46
7.44
7.40
7.38
7.35
7.33
7.30
7.29
7.28
7.27
7.26
7.25
7.23
4.92
3.53
3.32 HDO
3.31 HDO
3.29
2.49 dmso
2.49 dmso
2.48 dmso
2.48 dmso
2.47 dmso
0.82
0.86
0.85
1.01
1.10
1.99
2.23
1.19
2.20
1.87
1.88
f1 (ppm)

MaxPeak:  100.00%
Ret_Time: 1.043 min

Q227980$2

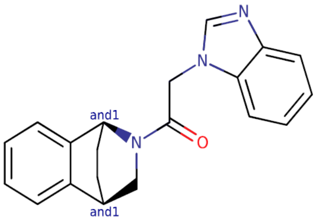

**Mol Wt** 317.38

**Exact Mass** 317.18

| # | Time | Area% |
|---|---|---|
| 1 | 1.043 | 100.00 |

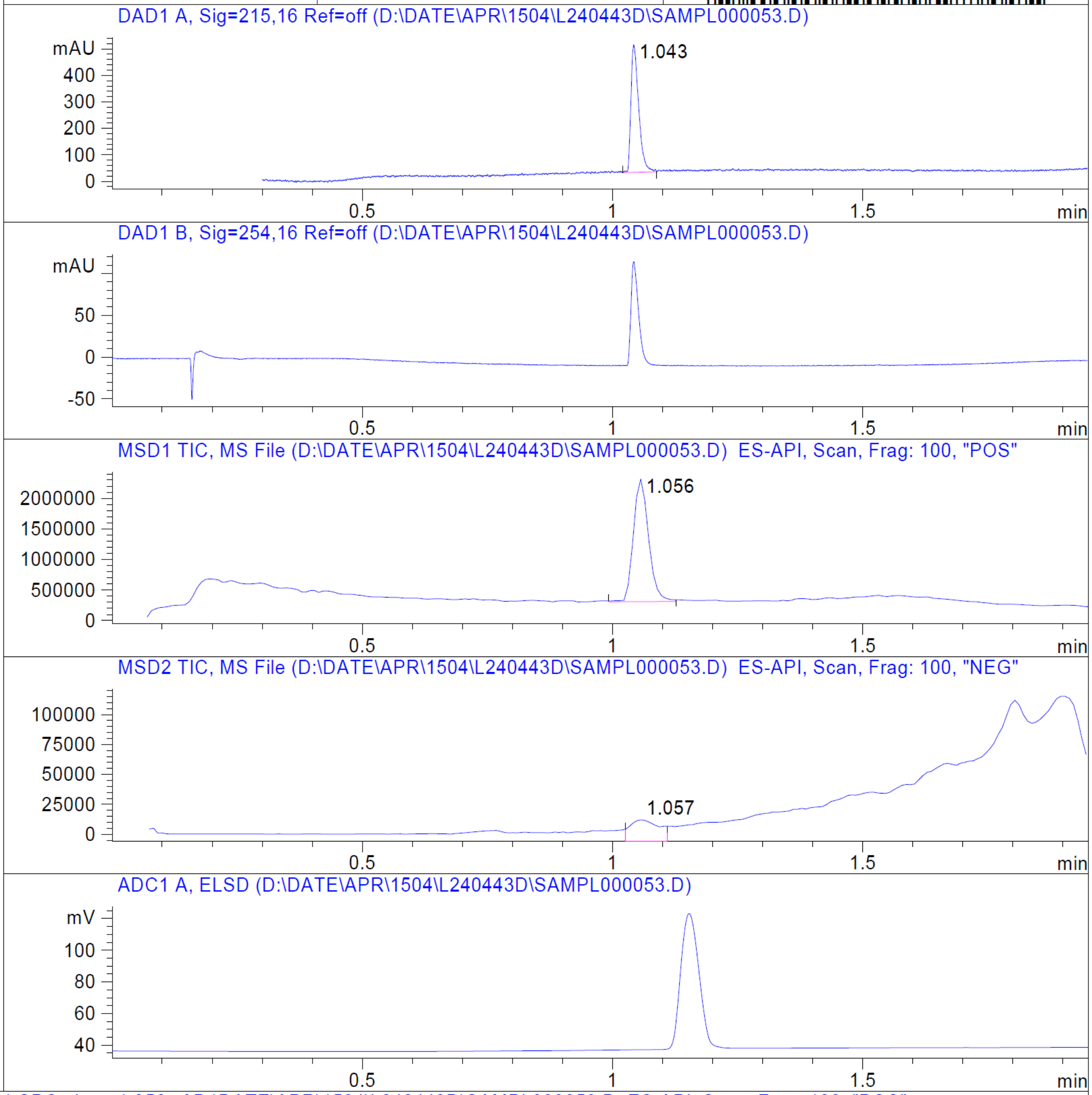


RT 1.056

RT 1.057

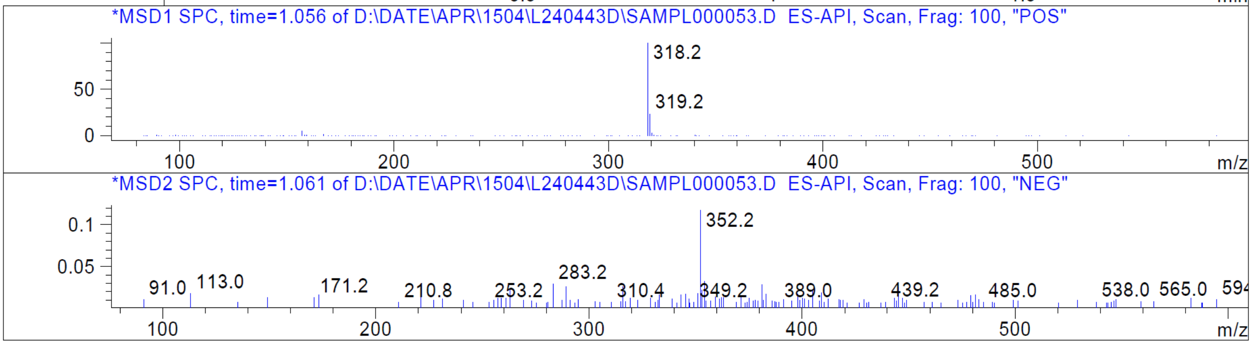


Z2876049675_(ZINC001339780091)

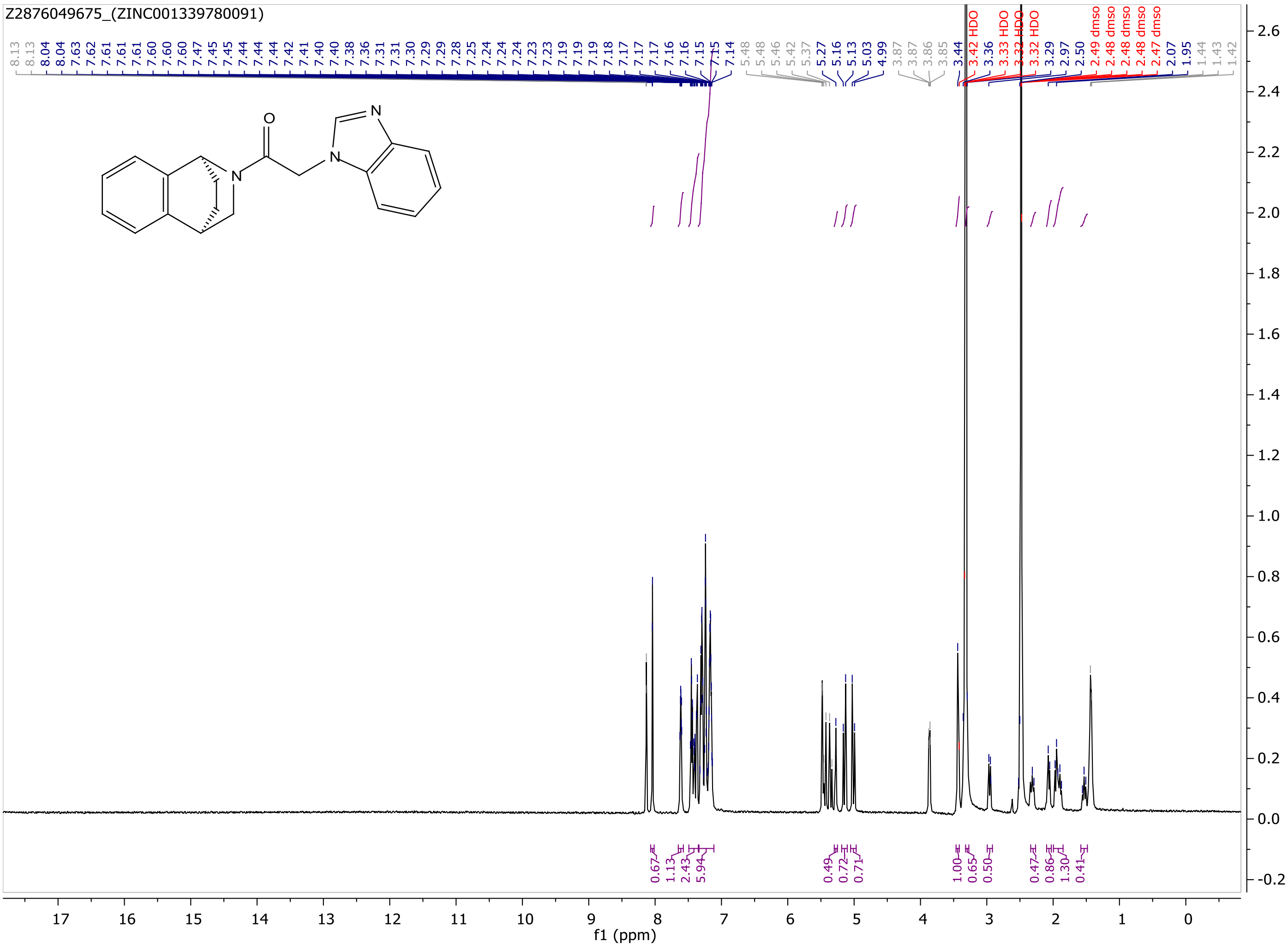

Z2876049675_(ZINC001339780091)
f1 (ppm)

MaxPeak: 100.00%
Ret_Time: 1.255 min

Q228024$1

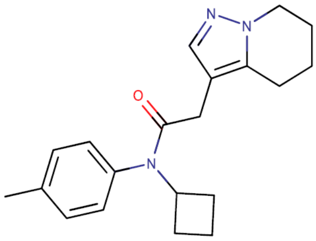


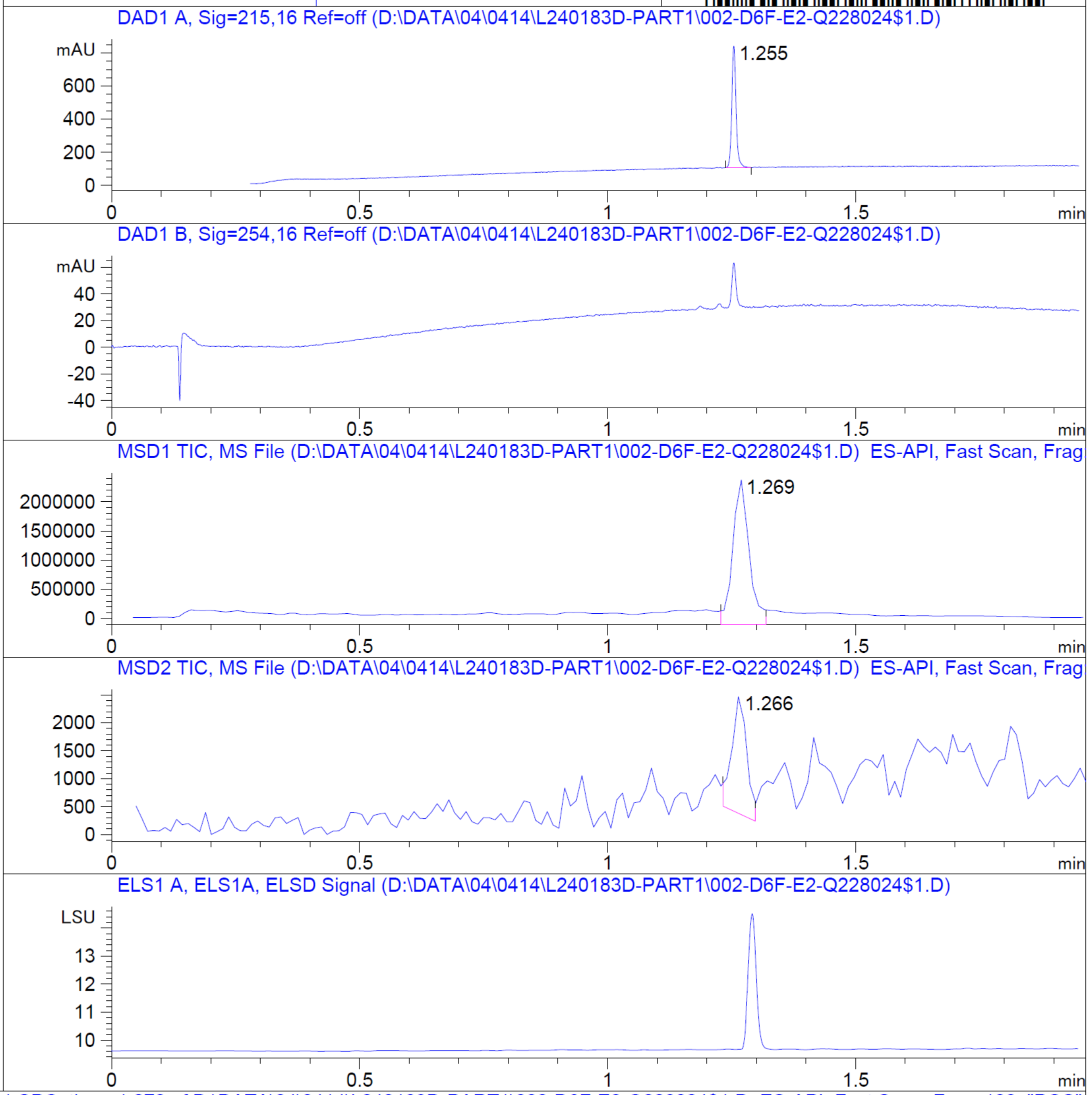


**Mol Wt** 323.43
**Exact Mass** 323.24

| # | Time | Area% |
|---|---|---|
| 1 | 1.255 | 100.00 |

RT 1.269

RT 1.266

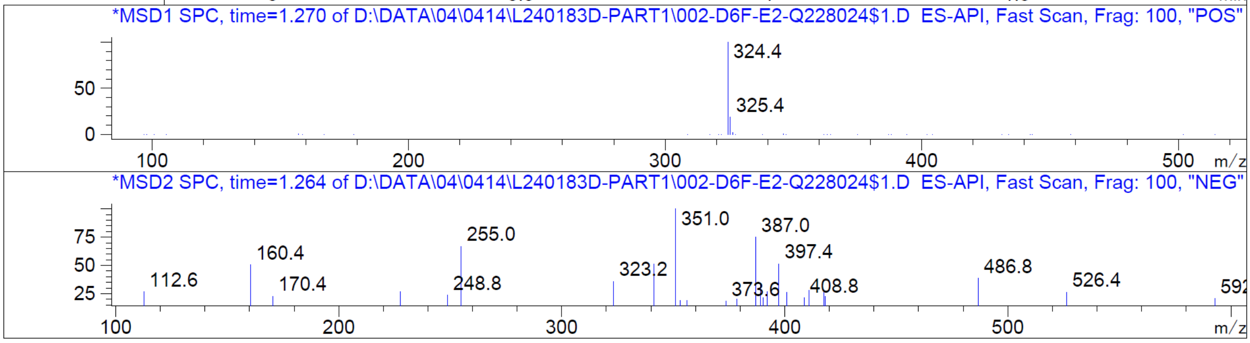


Z2876444855_(ZINC000894230117)

Z2876444855_(ZINC000894230117)

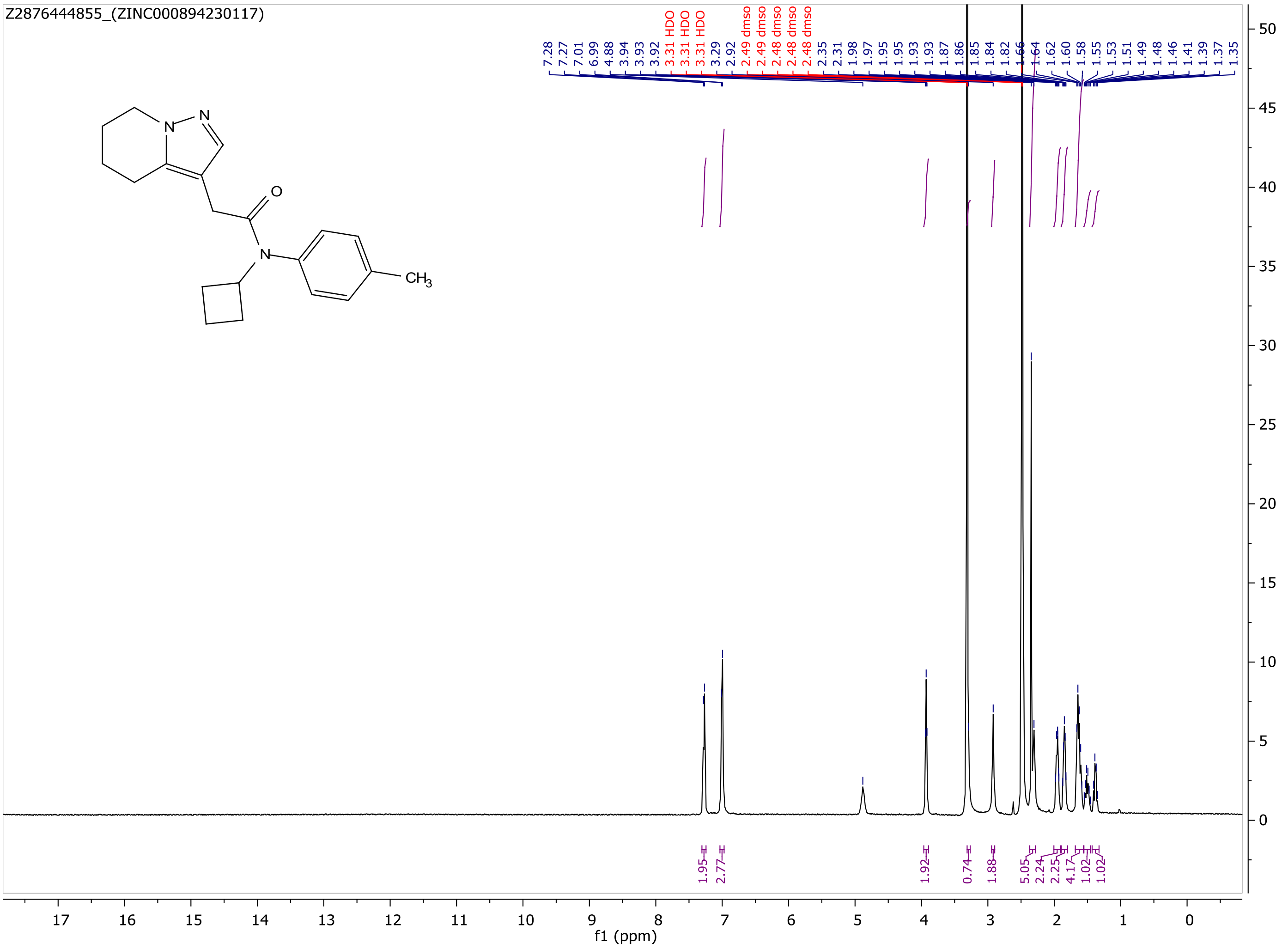

MaxPeak:  100.00%
Ret_Time: 1.053 min

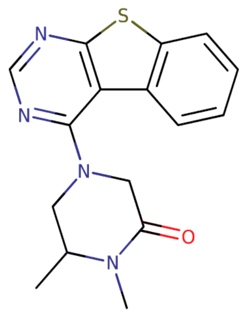


**Mol Wt** 312.39

**Exact Mass** 312.12

| # | Time | Area% |
|---|---|---|
| 1 | 1.053 | 100.00 |

# Q228009$3

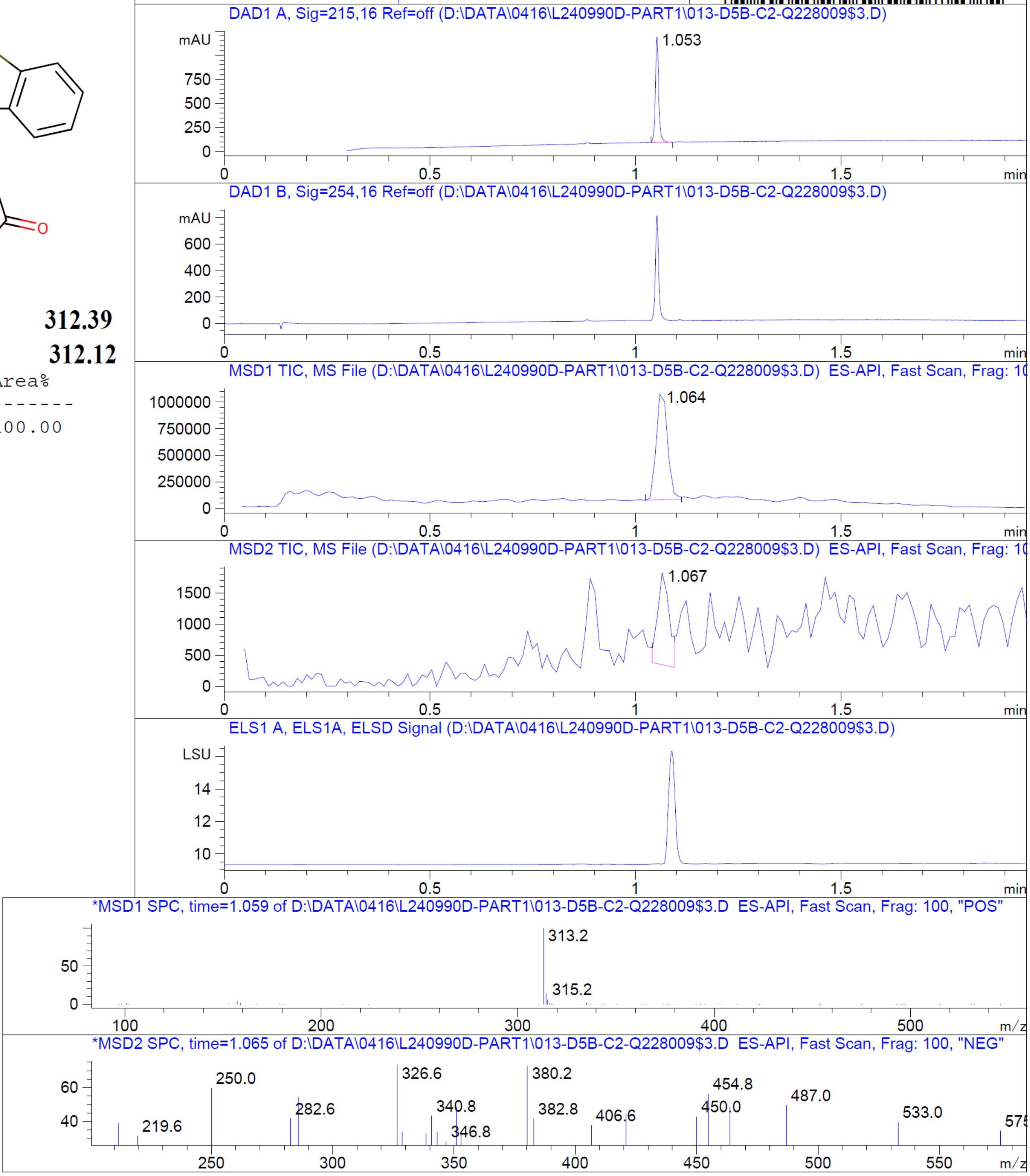


RT 1.064

RT 1.067

Z2000267872_(ZINC000336912805)

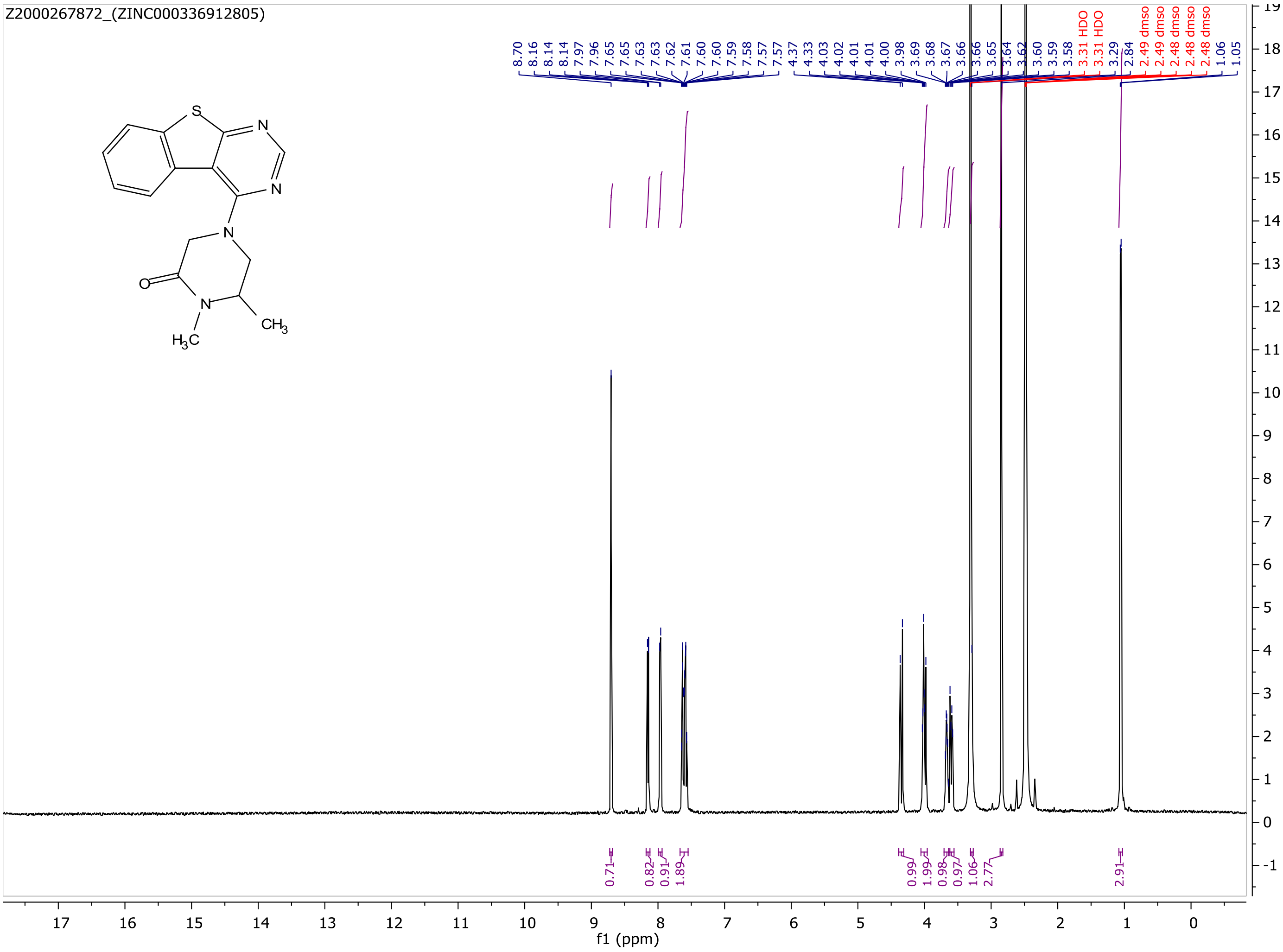
Z2000267872_(ZINC000336912805)
8.70
8.16
8.14
8.14
7.97
7.96
7.65
7.65
7.63
7.63
7.62
7.61
7.60
7.59
7.58
7.57
7.57
4.37
4.33
4.03
4.02
4.01
4.01
4.00
3.98
3.69
3.68
3.67
3.66
3.66
3.65
3.64
3.62
3.60
3.59
3.58
3.31 HDO
3.31 HDO
3.29
2.84
2.49 dmso
2.49 dmso
2.48 dmso
2.48 dmso
2.48 dmso
1.06
1.05
0.71
0.82
0.91
1.89
0.99
1.99
0.98
0.97
1.06
2.77
2.91
f1 (ppm)

MaxPeak: 97.45%
Ret_Time: 1.408 min

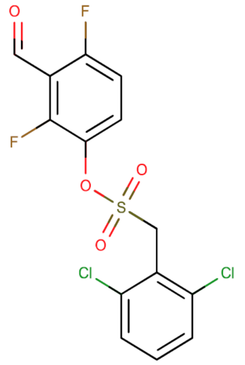

**Mol Wt** 381.18

**Exact Mass** 379.95

| # | Time | Area% |
|---|---|---|
| 1 | 1.408 | 97.45 |
| 2 | 1.461 | 2.55 |

# Q260270$3

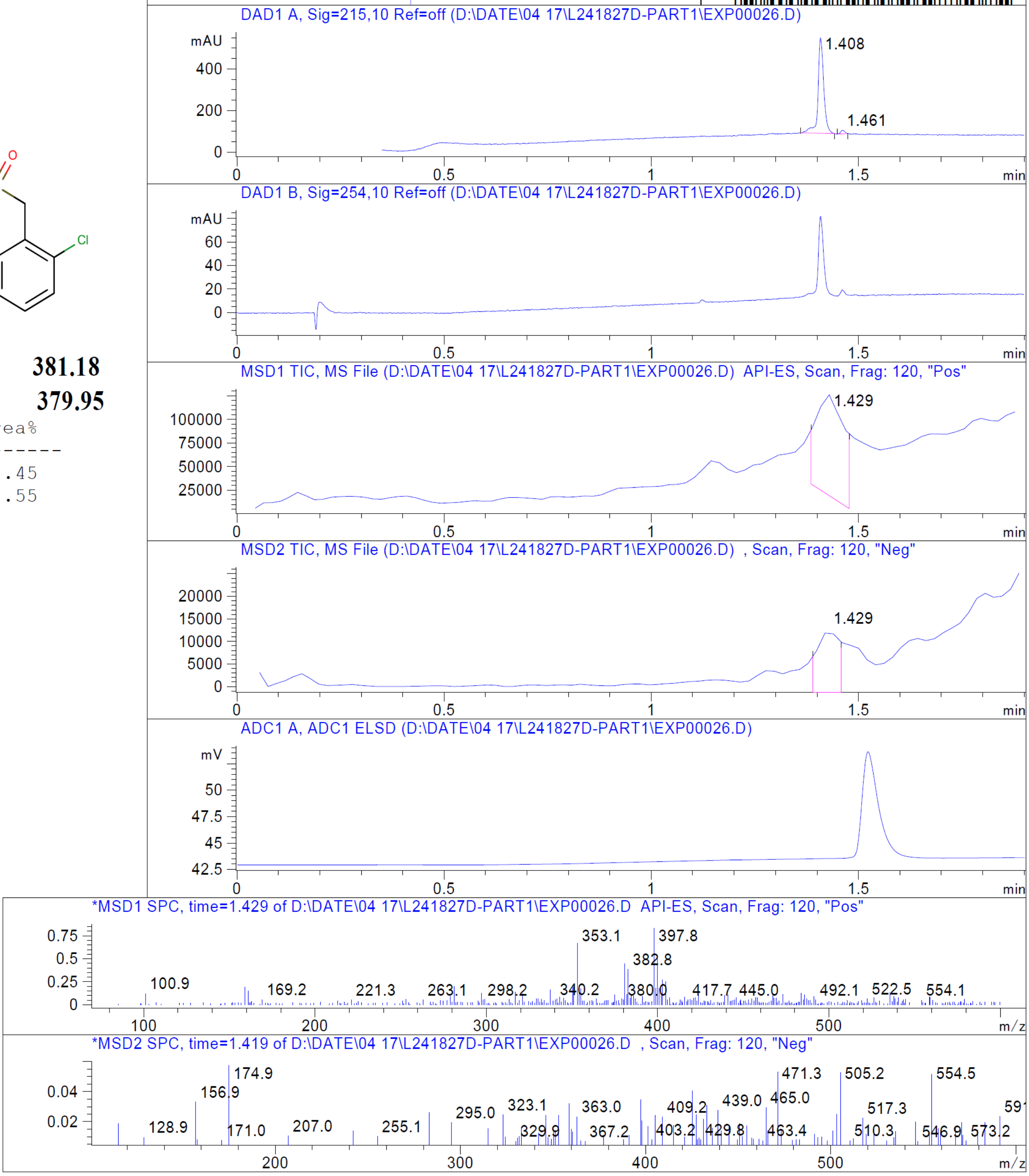


RT 1.429

RT 1.429

Z1958664470_(ZINC000795258204)

Z1958664470_(ZINC000795258204)

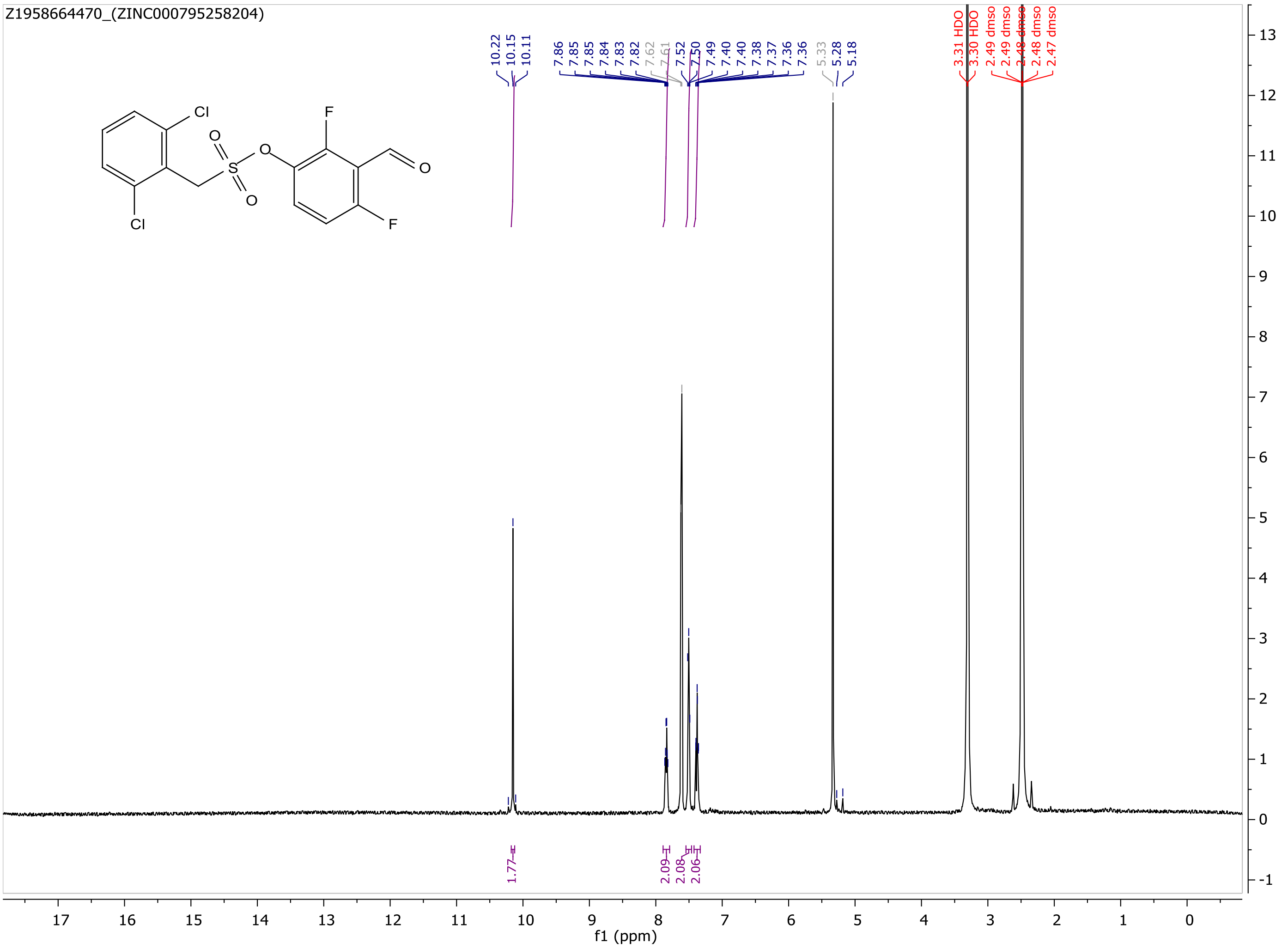

MaxPeak: 100.00%
Ret_Time: 1.212 min

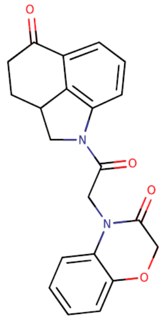

**Mol Wt** 362.38

**Exact Mass** 362.14

| # | Time | Area% |
|---|---|---|
| 1 | 1.212 | 100.00 |

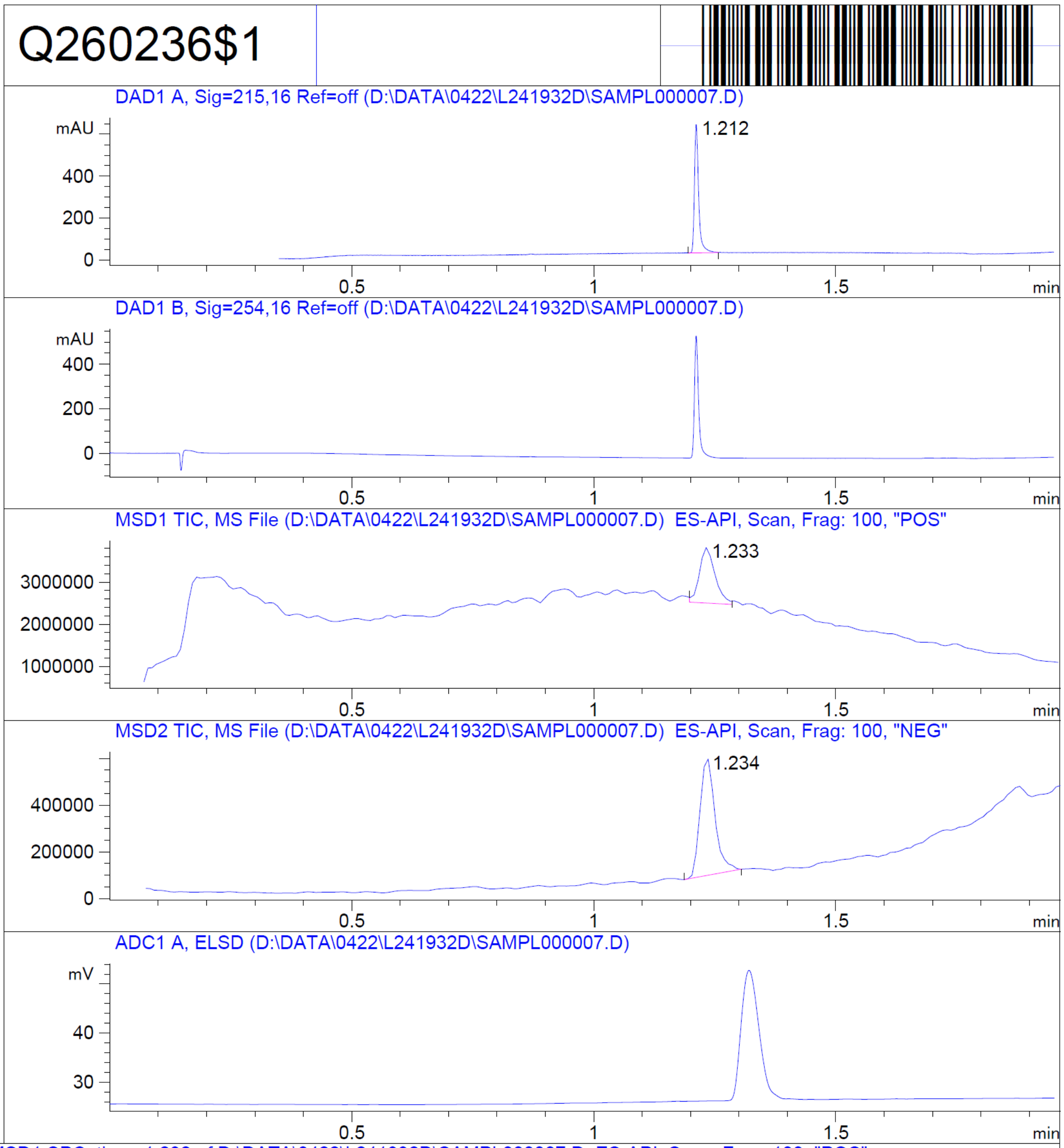


RT 1.233

RT 1.234

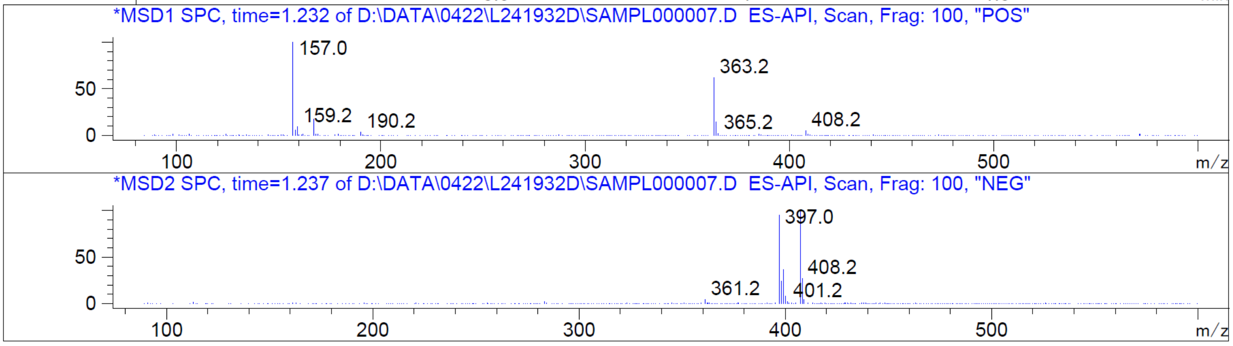


Z1697490576_(ZINC000274326055)

Z1697490576_(ZINC000274326055)

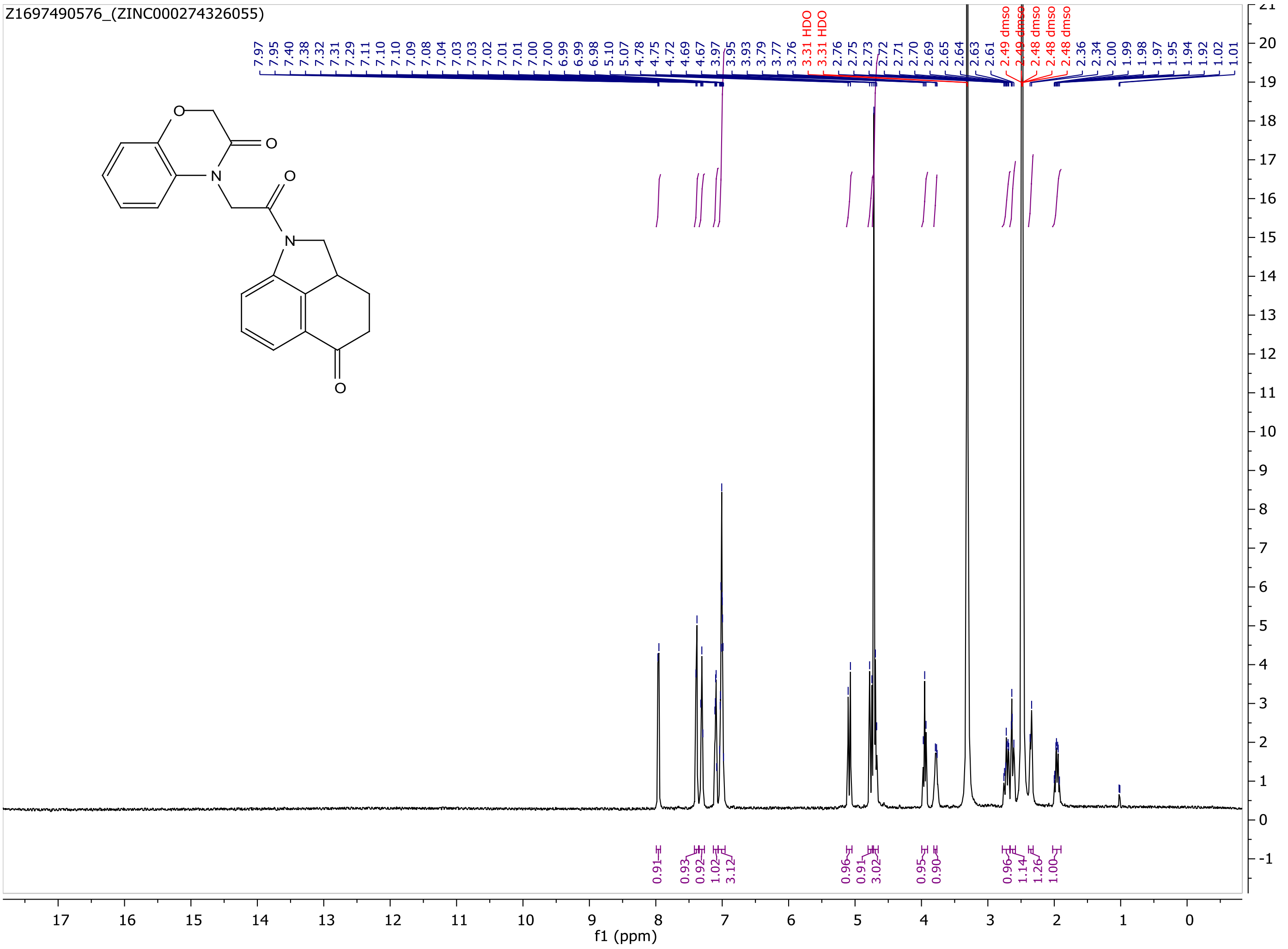

MaxPeak: 100.00%
Ret_Time: 1.204 min

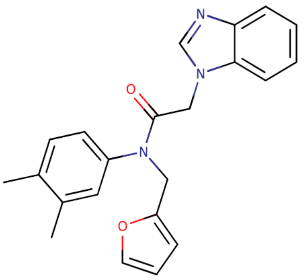


**Mol Wt** 359.42
**Exact Mass** 359.19

| # | Time | Area% |
|---|---|---|
| 1 | 1.204 | 100.00 |

Q260188$1

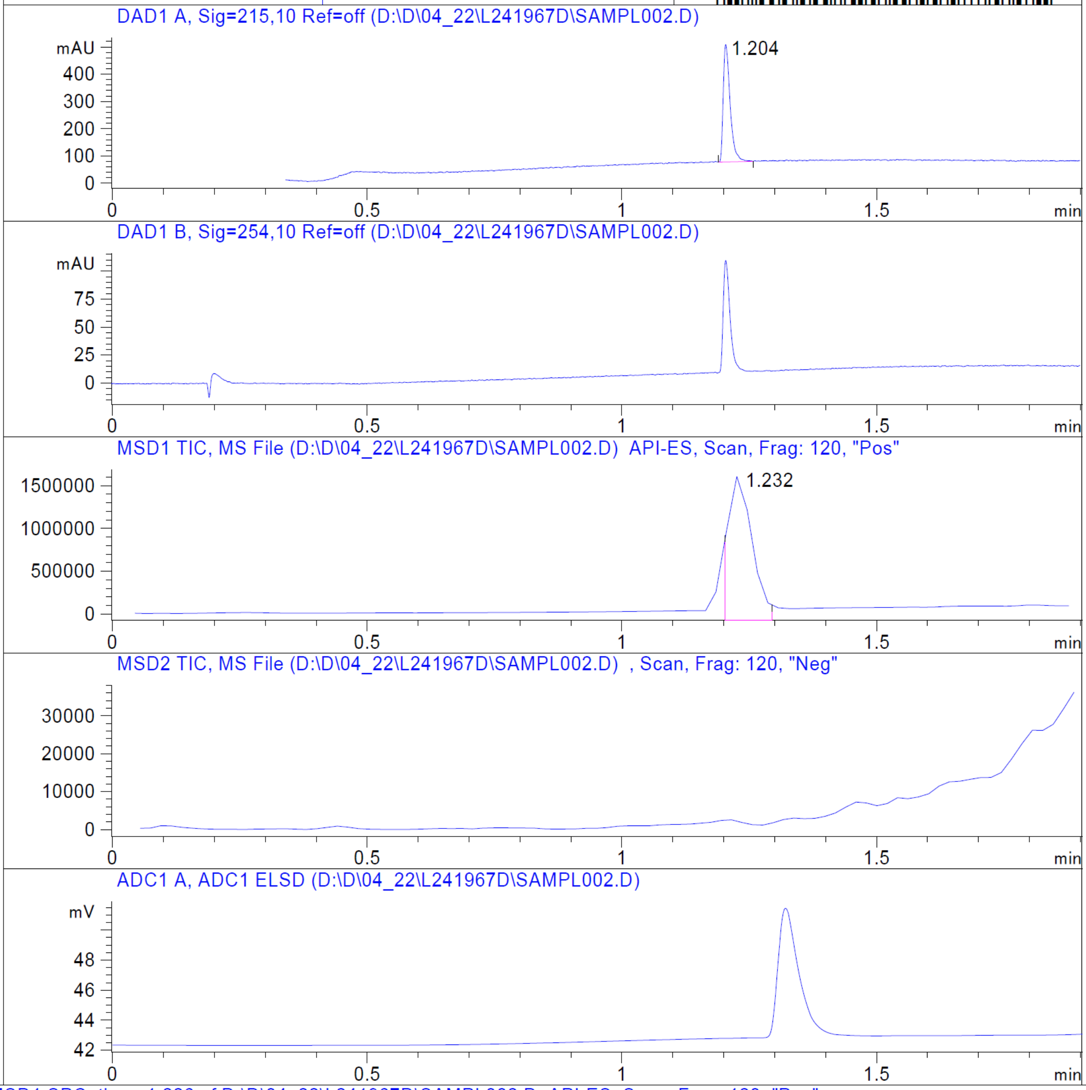


RT 1.232

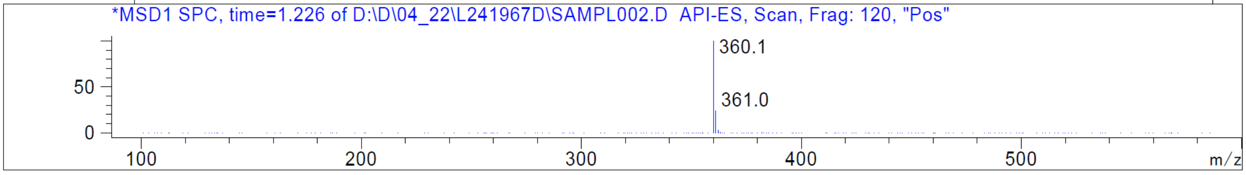


Z430141838_(ZINC000915668084)

Z430141838_(ZINC000915668084)

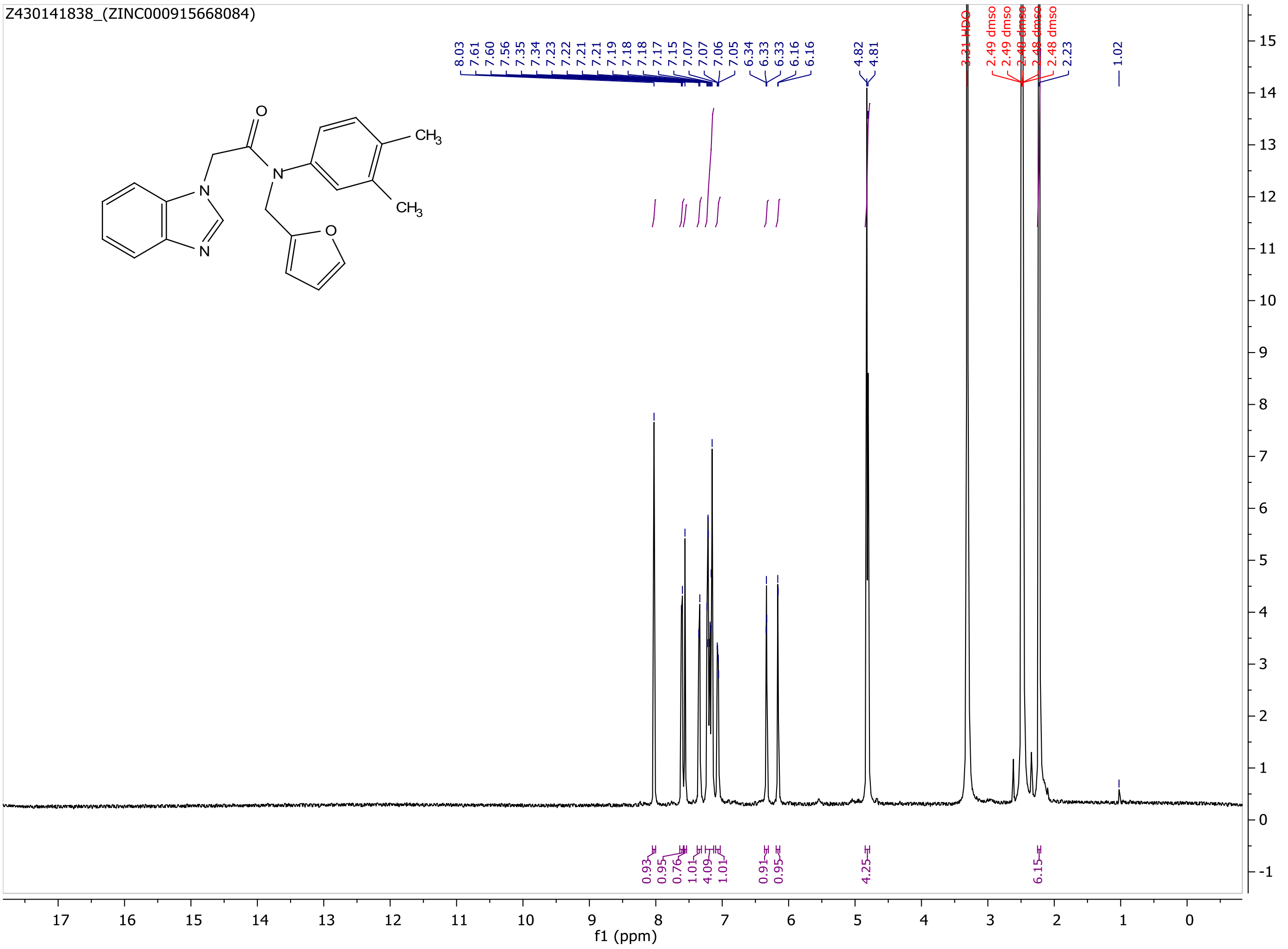

MaxPeak: 96.66%
Ret_Time: 1.411 min

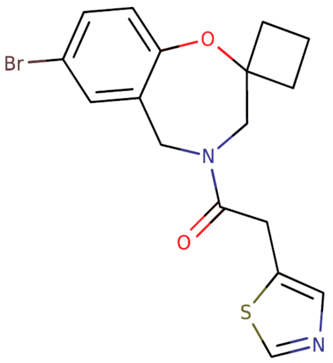


**Mol Wt** 393.3

**Exact Mass** 394.04

| # | Time | Area% |
|---|---|---|
| 1 | 1.411 | 96.66 |
| 2 | 1.585 | 3.34 |

Q260276$1

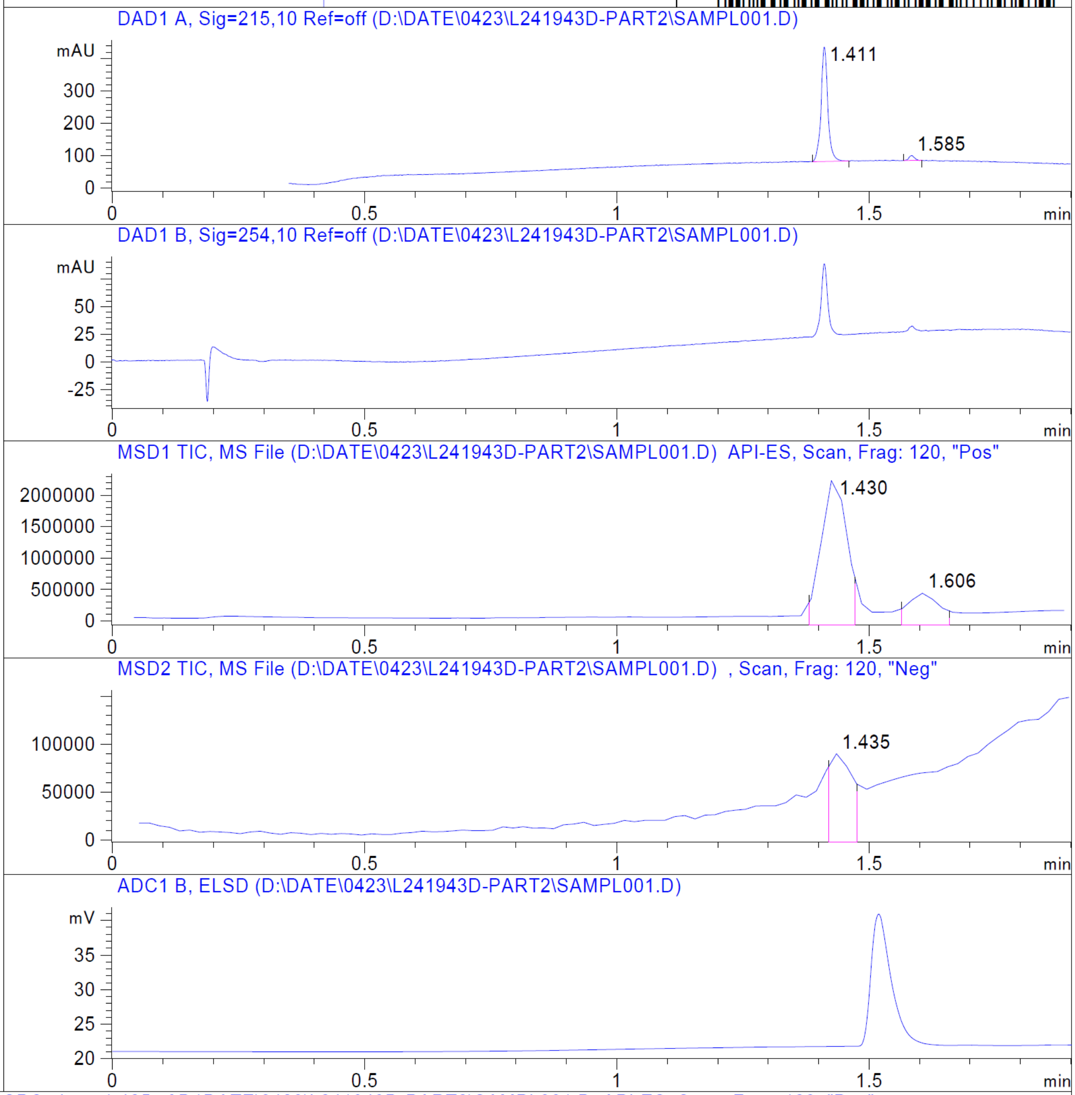


RT 1.430

RT 1.606

RT 1.435

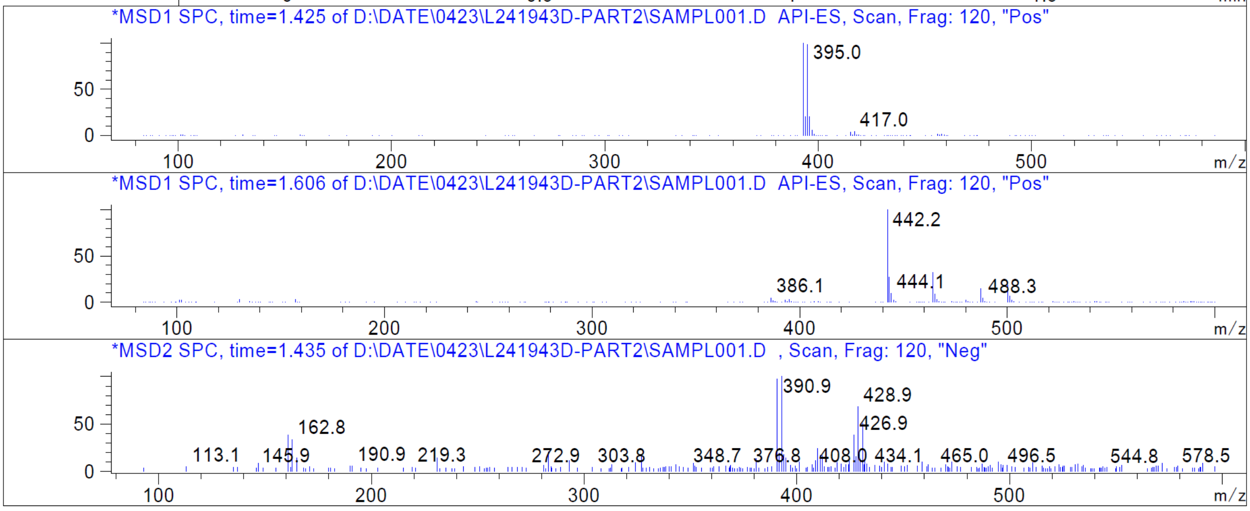


Z2931822383_(ZINC000928186863)

Z2931822383_(ZINC000928186863)

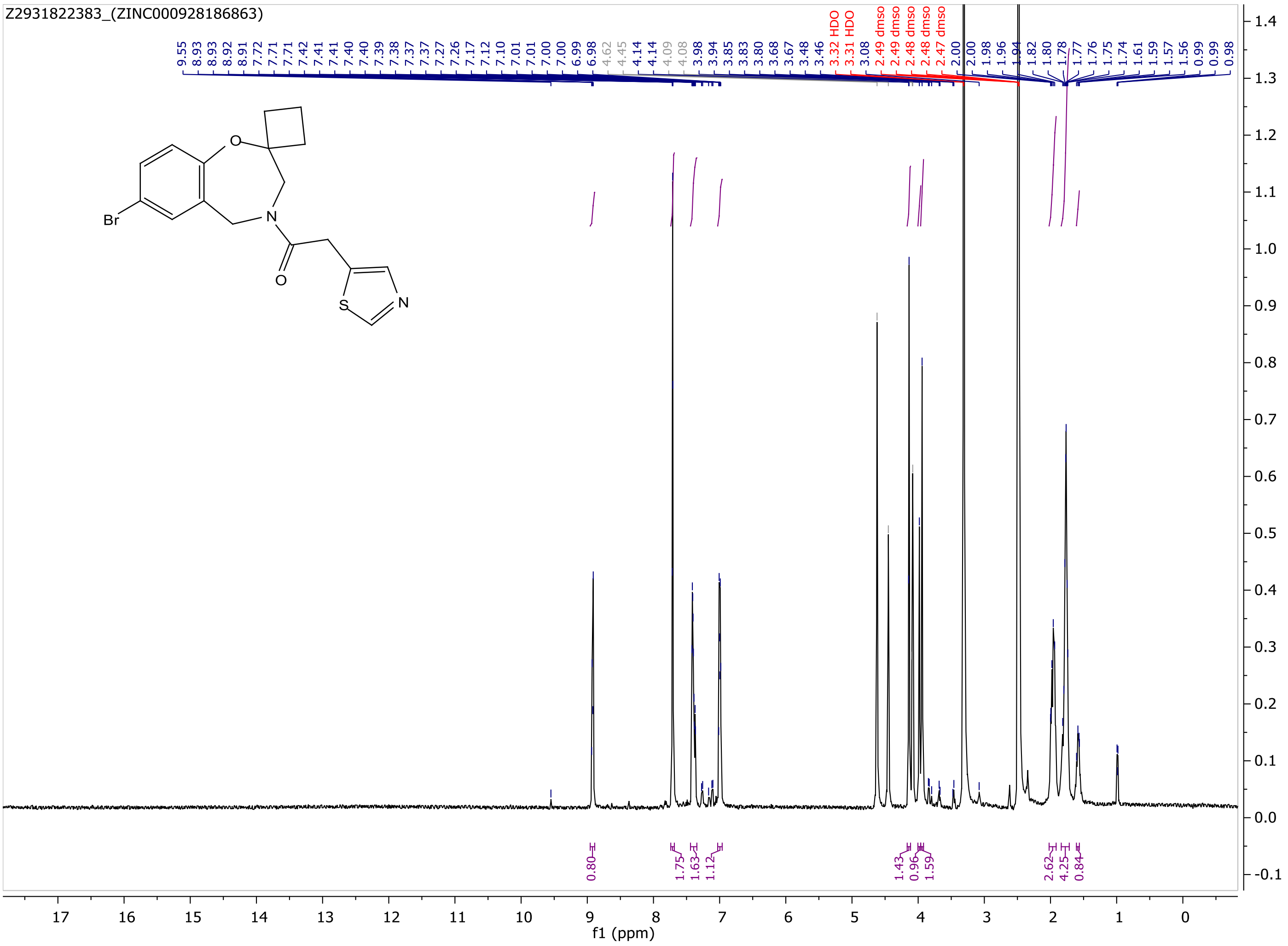

MaxPeak: 100.00%
Ret_Time: 1.041 min

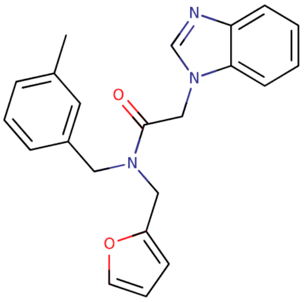


**Mol Wt** 359.42

**Exact Mass** 359.19

| # | Time | Area% |
|---|---|---|
| 1 | 1.041 | 100.00 |

# H2334205

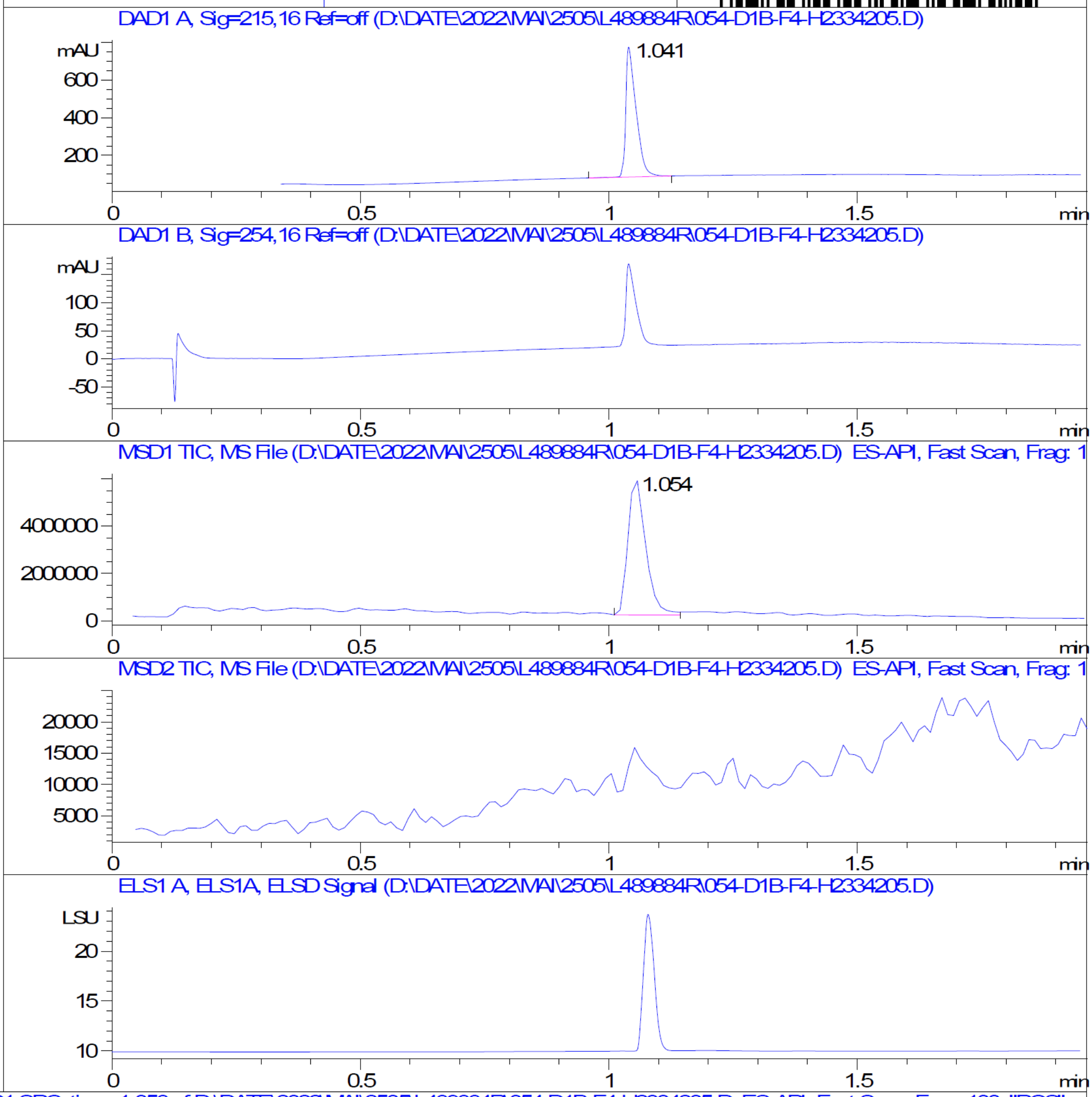


RT 1.054

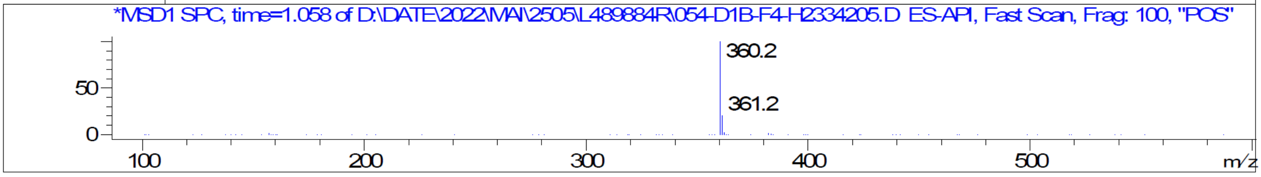


Z430143532_(ZINC000512479458)

Z430143532_(ZINC000512479458)

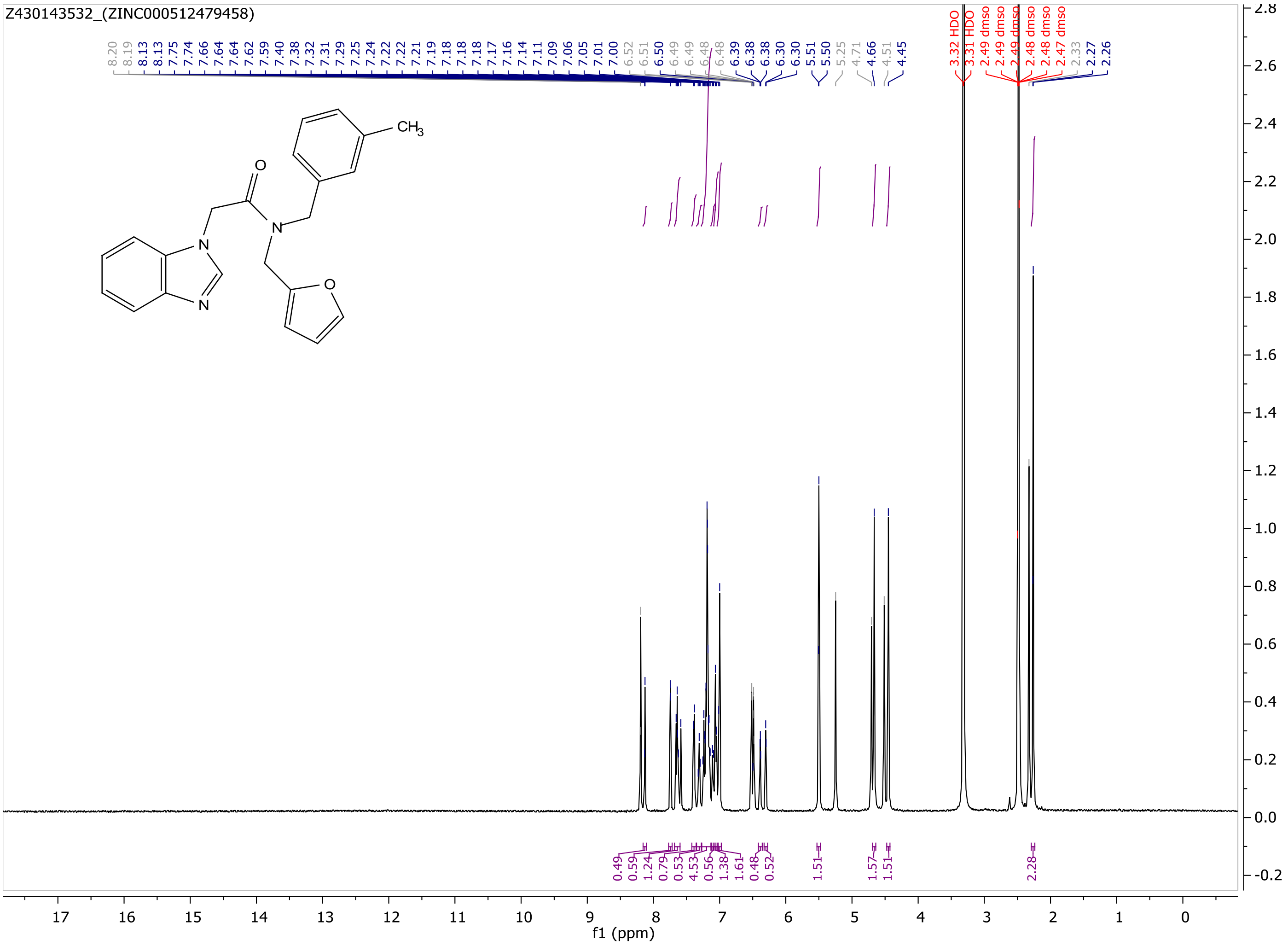

MaxPeak: 100.00%
Ret_Time: 1.099 min

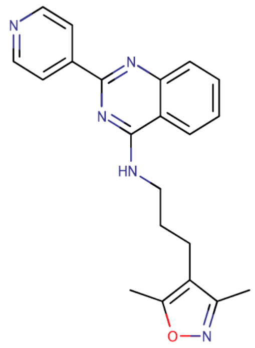

**Mol Wt** 359.42

**Exact Mass** 359.2

| # | Time | Area% |
|---|---|---|
| 1 | 1.099 | 100.00 |

Q260300$2

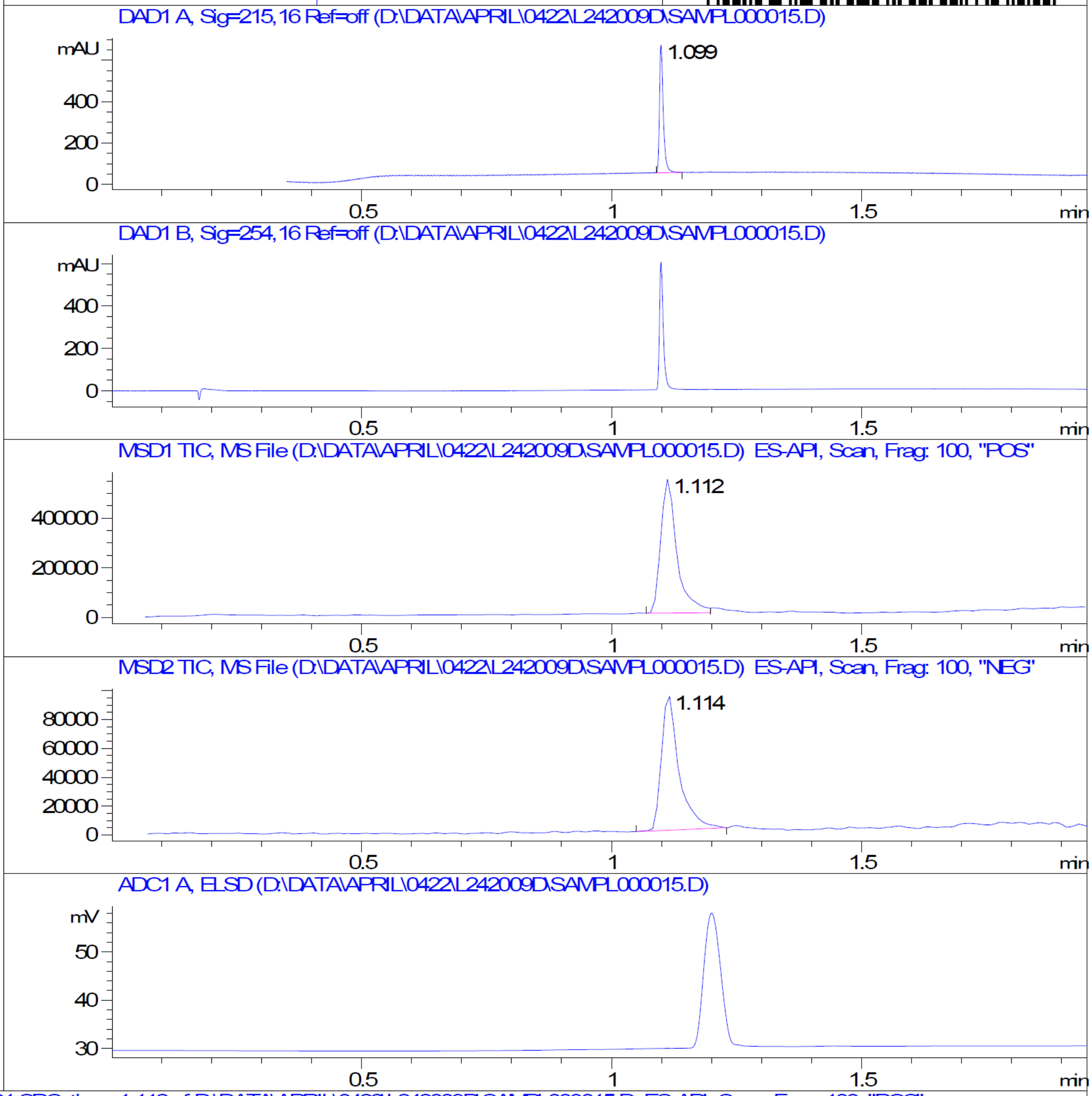


RT 1.112

RT 1.114

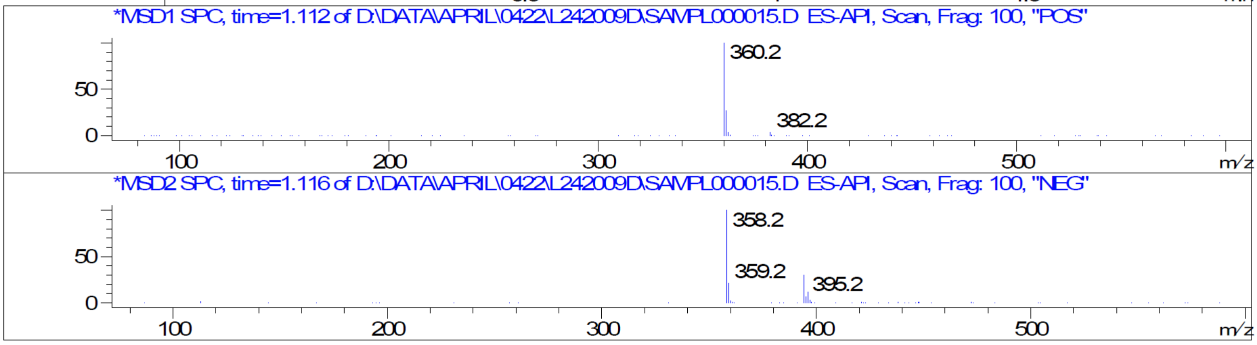


Z1378403257_(ZINC000301553312)

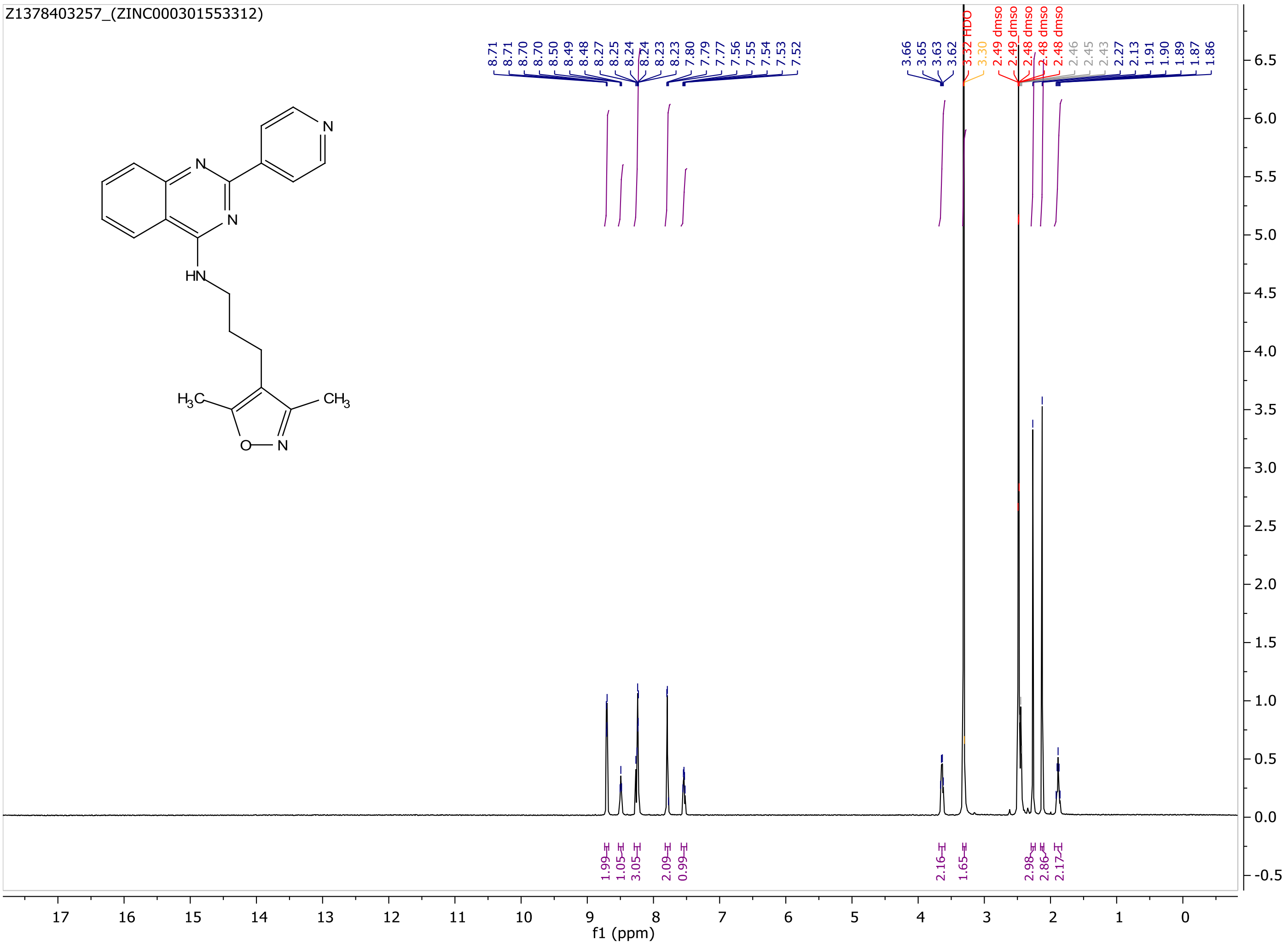
Z1378403257_(ZINC000301553312)
8.71 8.71 8.70 8.70 8.50 8.49 8.48 8.27 8.25 8.24 8.24 8.23 7.80 7.79 7.77 7.56 7.55 7.54 7.53 7.52
3.66 3.65 3.63 3.62 3.32 HDO 3.30 2.49 dmso 2.49 dmso 2.48 dmso 2.48 dmso 2.48 dmso 2.46 2.45 2.43 2.27 2.13 1.91 1.90 1.89 1.87 1.86
N
N
N
HN
$H_3C$
$CH_3$
O
N
1.99
1.05
3.05
2.09
0.99
2.16
1.65
2.98
2.86
2.17
f1 (ppm)

MaxPeak: 100.00%
Ret_Time: 1.386 min

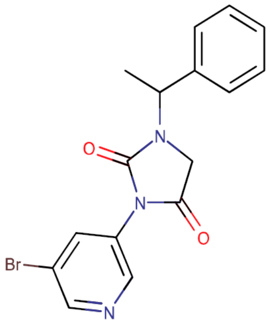

**Mol Wt** 360.2
**Exact Mass** 359.04

| # | Time | Area% |
|---|---|---|
| 1 | 1.386 | 100.00 |

Q260256$3

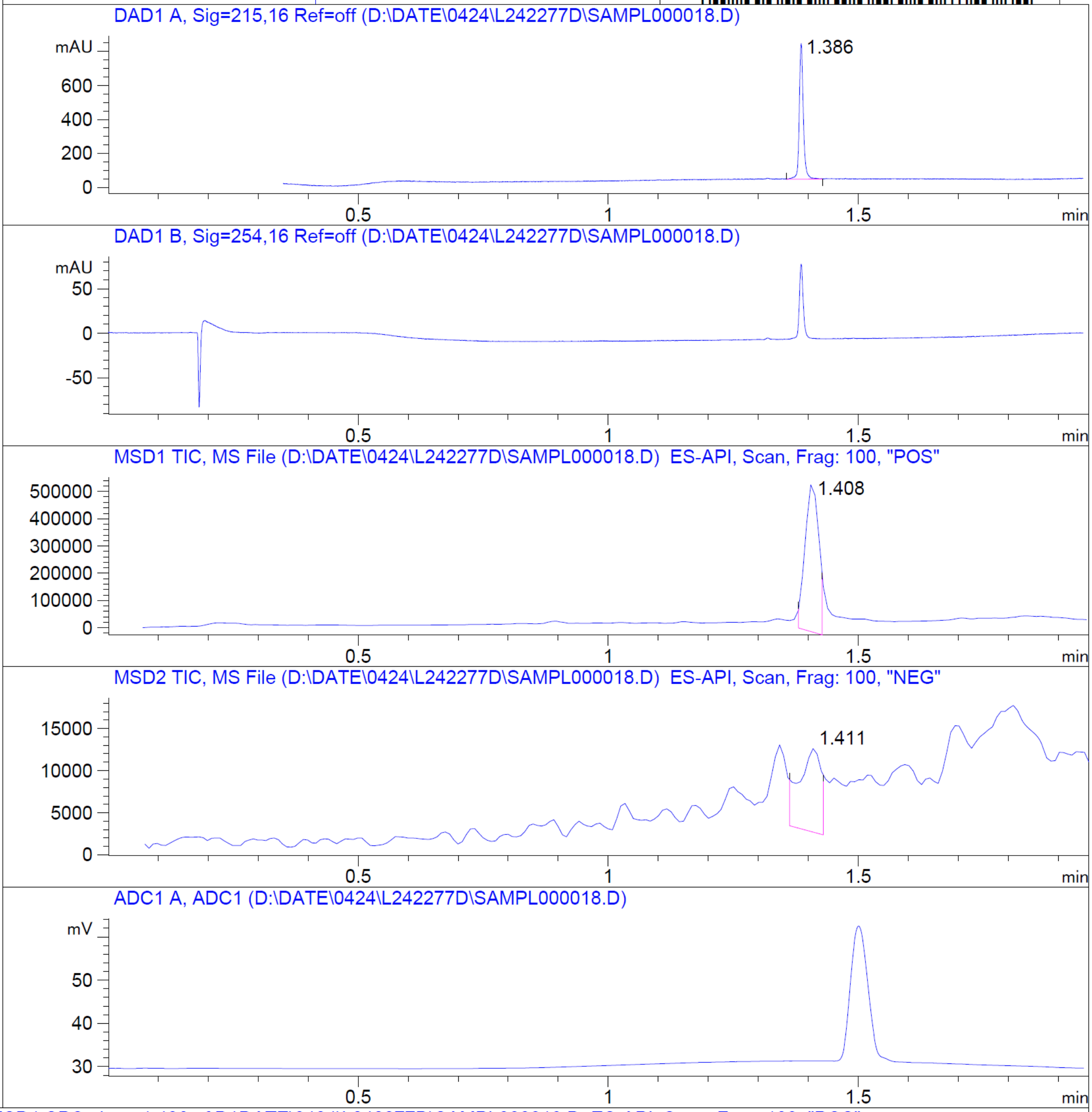


RT 1.408

RT 1.411

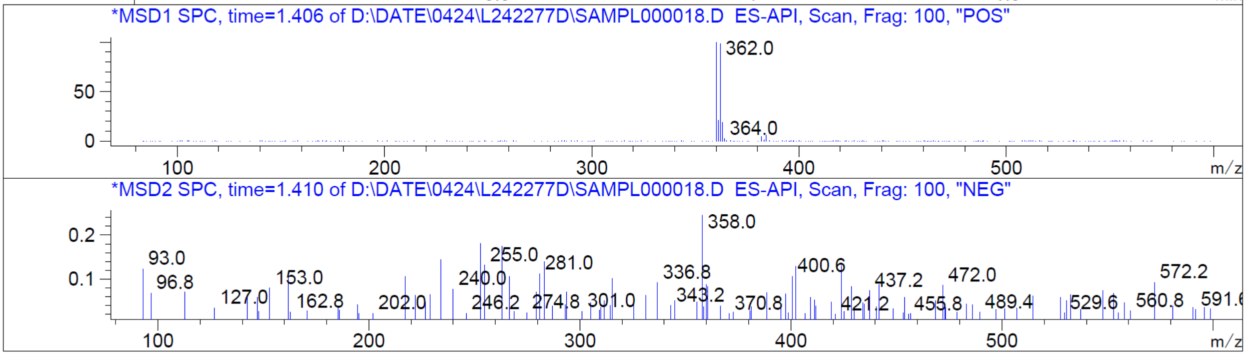


Z2193994640_(ZINC000813360541)

Z2193994640_(ZINC000813360541)

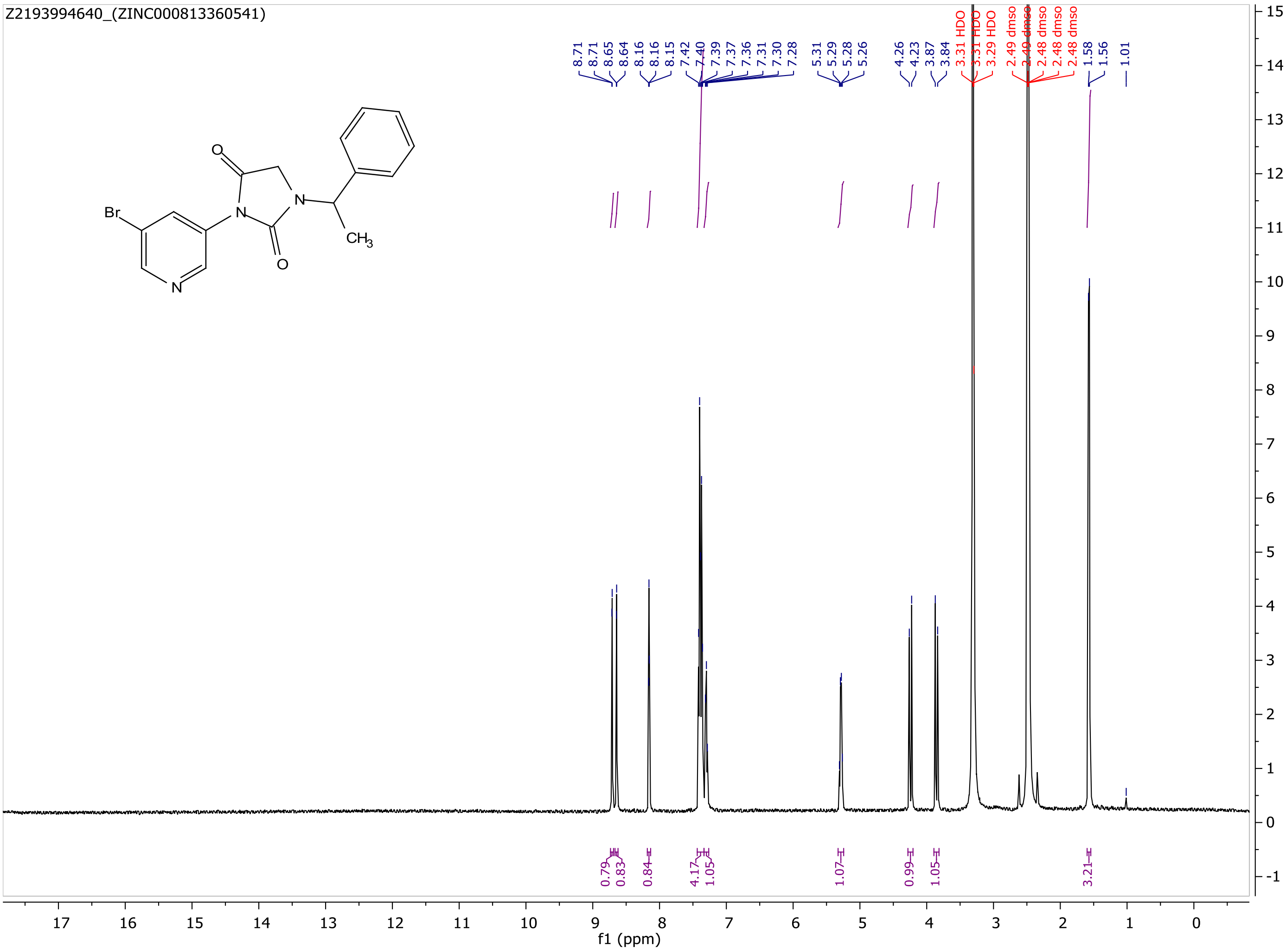

MaxPeak: 100.00%
Ret_Time: 1.291 min

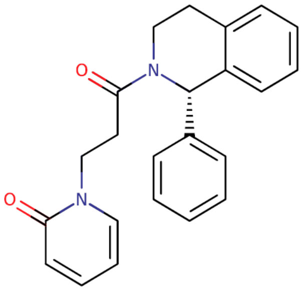


**Mol Wt 358.43**
**Exact Mass 358.2**

| # | Time | Area% |
|---|---|---|
| 1 | 1.291 | 100.00 |

Q260233$2

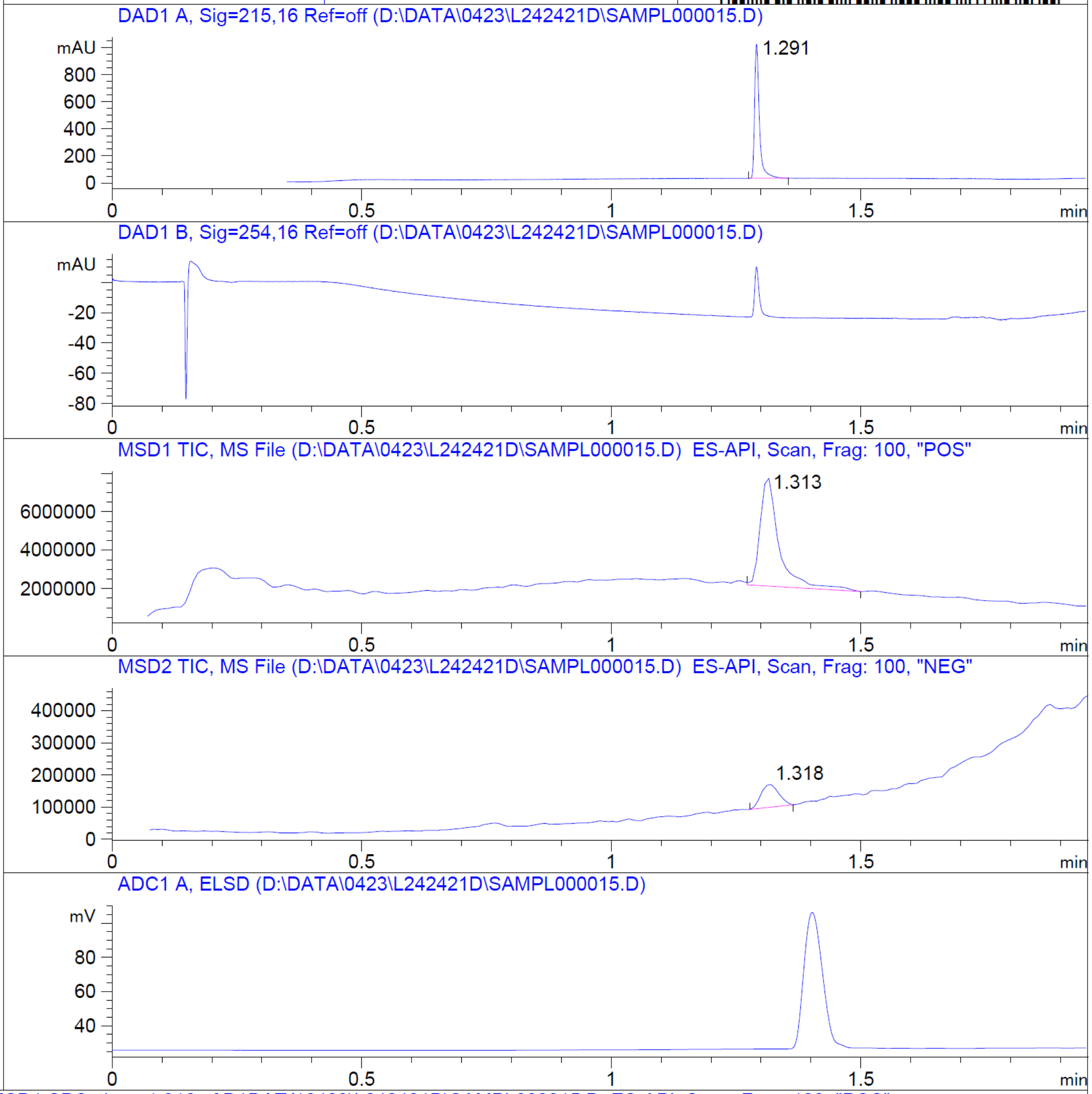


RT 1.313

RT 1.318

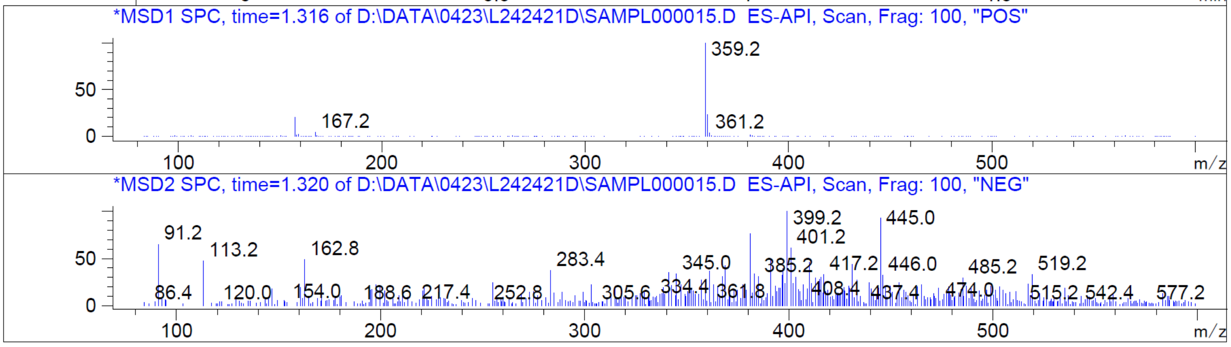


Z2232119003_(ZINC000594542103)

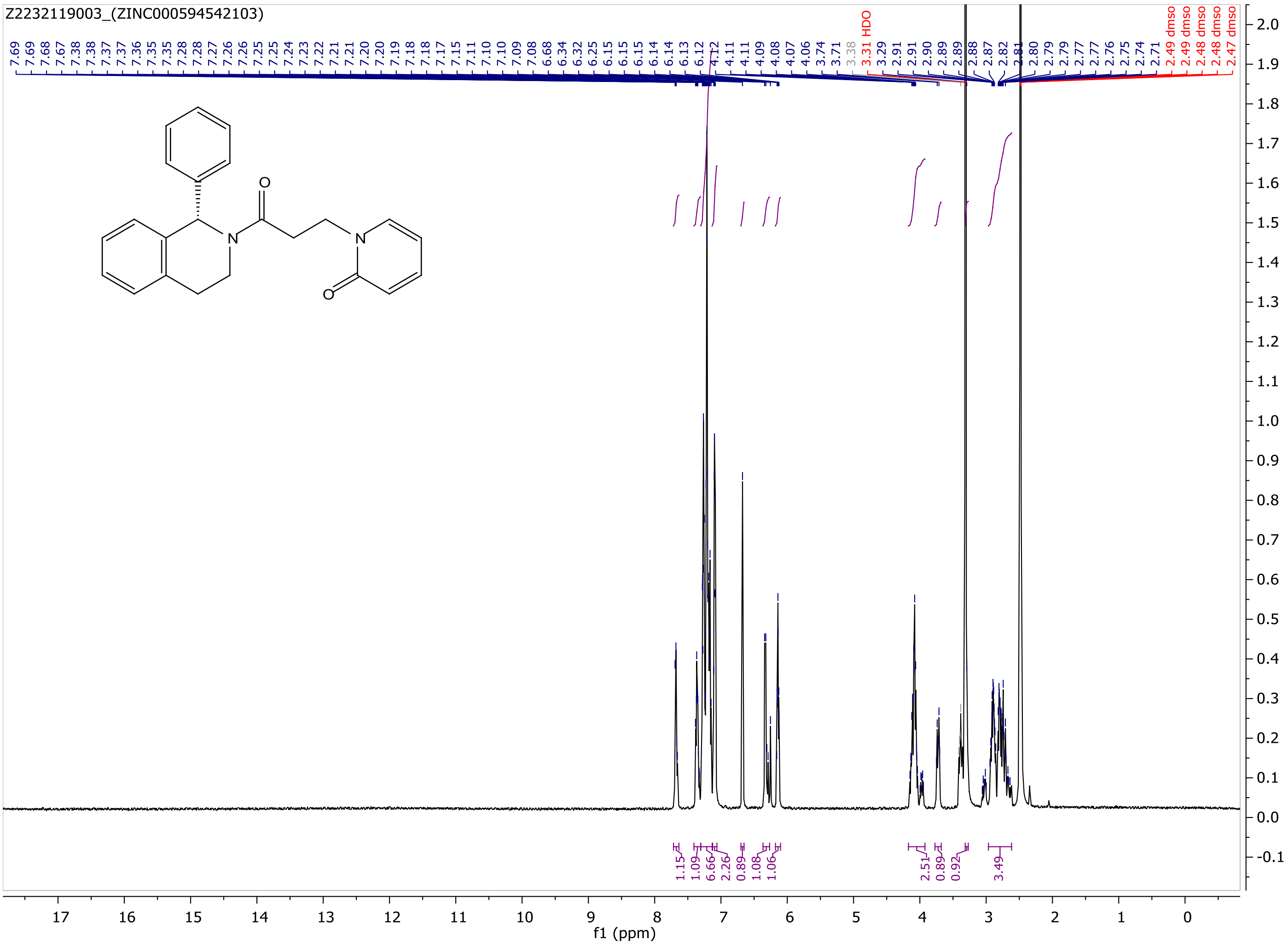
Z2232119003_(ZINC000594542103)
7.69 7.69 7.68 7.67 7.38 7.38 7.37 7.37 7.36 7.35 7.35 7.28 7.28 7.27 7.26 7.26 7.25 7.25 7.24 7.23 7.22 7.21 7.21 7.20 7.20 7.19 7.18 7.18 7.17 7.15 7.11 7.10 7.10 7.09 7.08 6.68 6.34 6.32 6.25 6.15 6.15 6.15 6.14 6.14 6.13 6.12 4.12 4.11 4.11 4.09 4.08 4.07 4.06 3.74 3.71 3.38 3.31 HDO 3.29 2.91 2.91 2.90 2.89 2.89 2.88 2.87 2.82 2.81 2.80 2.79 2.79 2.77 2.77 2.76 2.75 2.74 2.71 2.49 dmso 2.49 dmso 2.48 dmso 2.48 dmso 2.47 dmso
O
N
N
O
1.15 1.09 6.66 2.26 0.89 1.08 1.06 2.51 0.89 0.92 3.49
f1 (ppm)

MaxPeak: 98.15%
Ret_Time: 1.258 min

Q260286$3

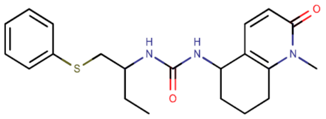

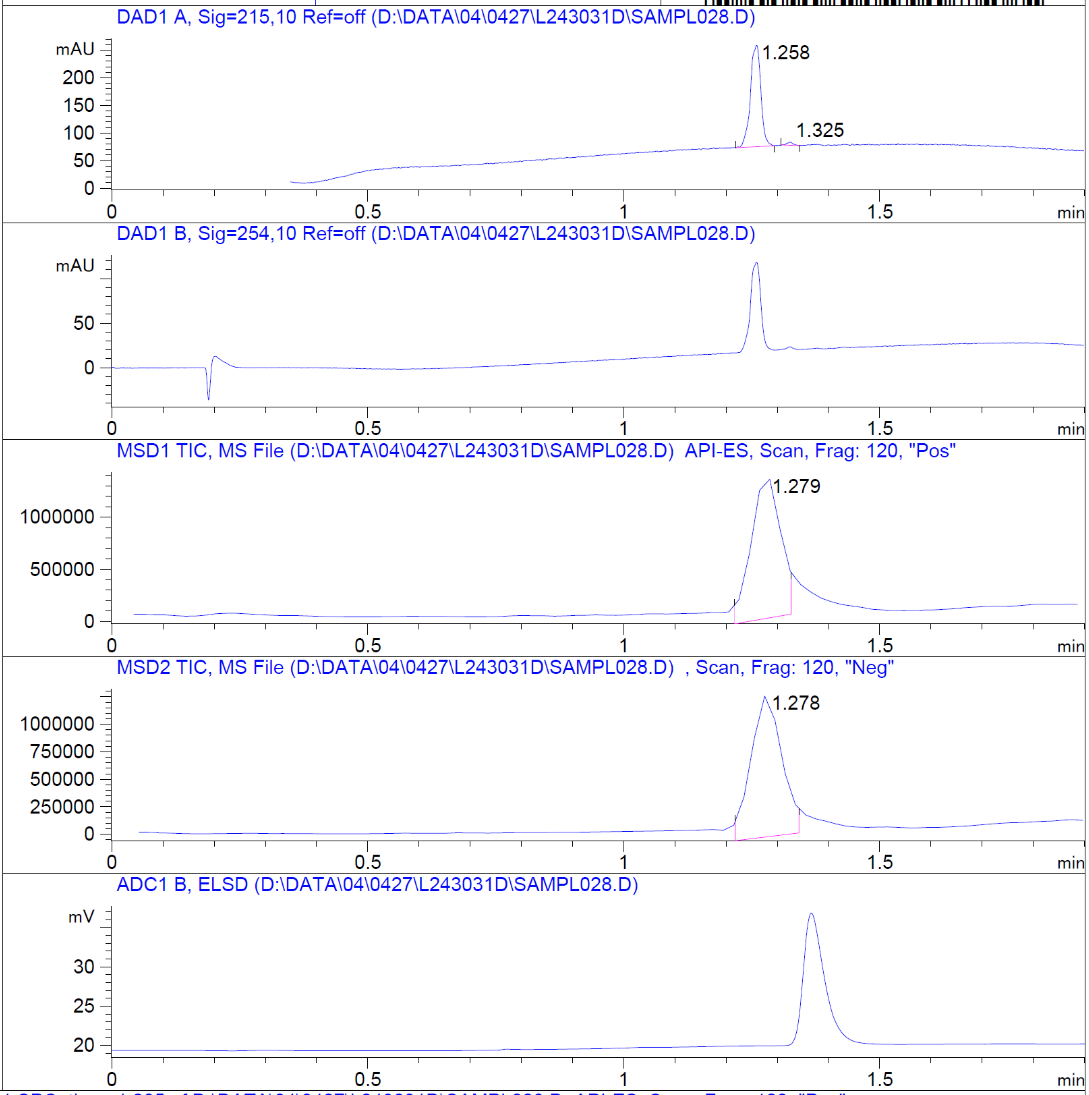


**Mol Wt** **385.52**

**Exact Mass** **385.22**

| # | Time | Area% |
|---|---|---|
| 1 | 1.258 | 98.15 |
| 2 | 1.325 | 1.85 |

RT 1.279

RT 1.278

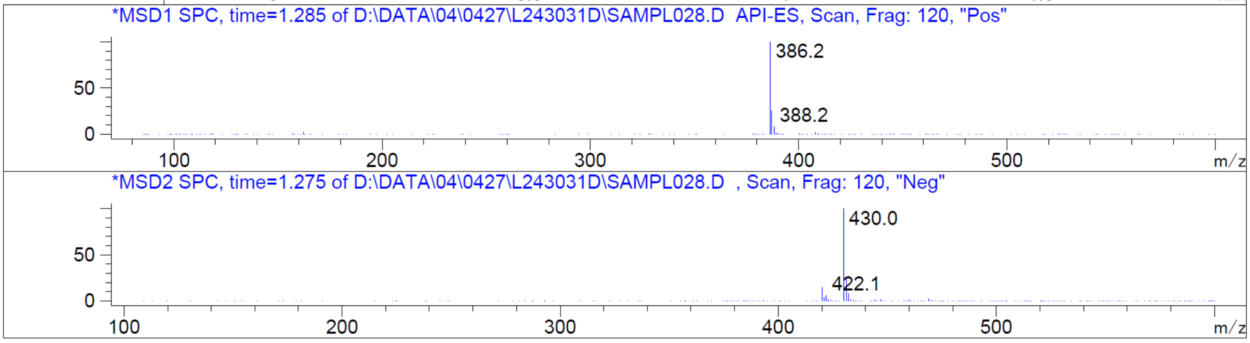


Z2477360481_(ZINC000650987125)

Z2477360481_(ZINC000650987125)

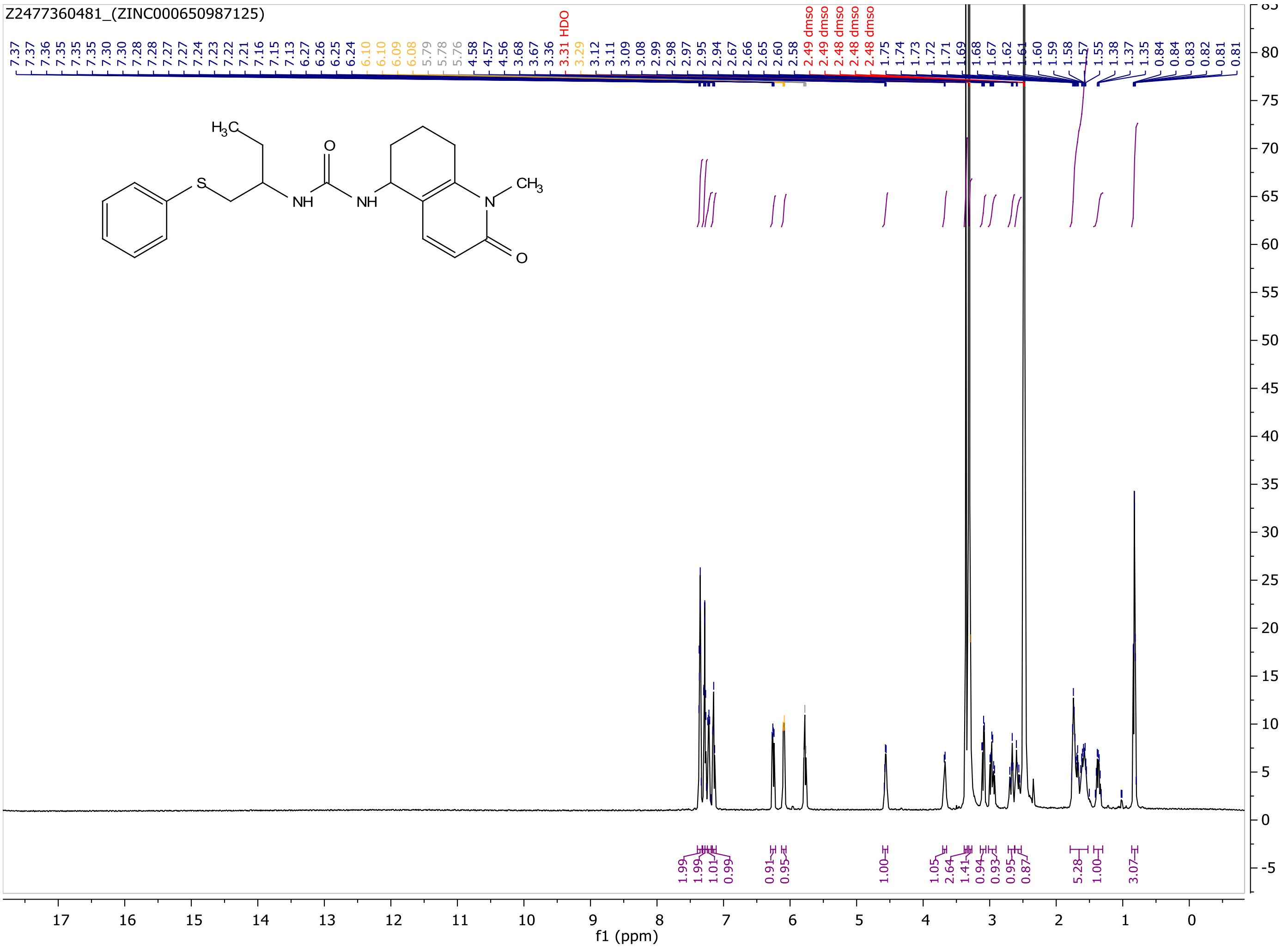

MaxPeak: 100.00%
Ret_Time: 1.300 min

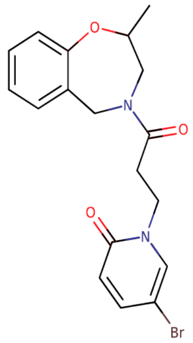


**Mol Wt** 391.26

**Exact Mass** 390.08

| # | Time | Area% |
|---|---|---|
| 1 | 1.300 | 100.00 |

Q260214$114

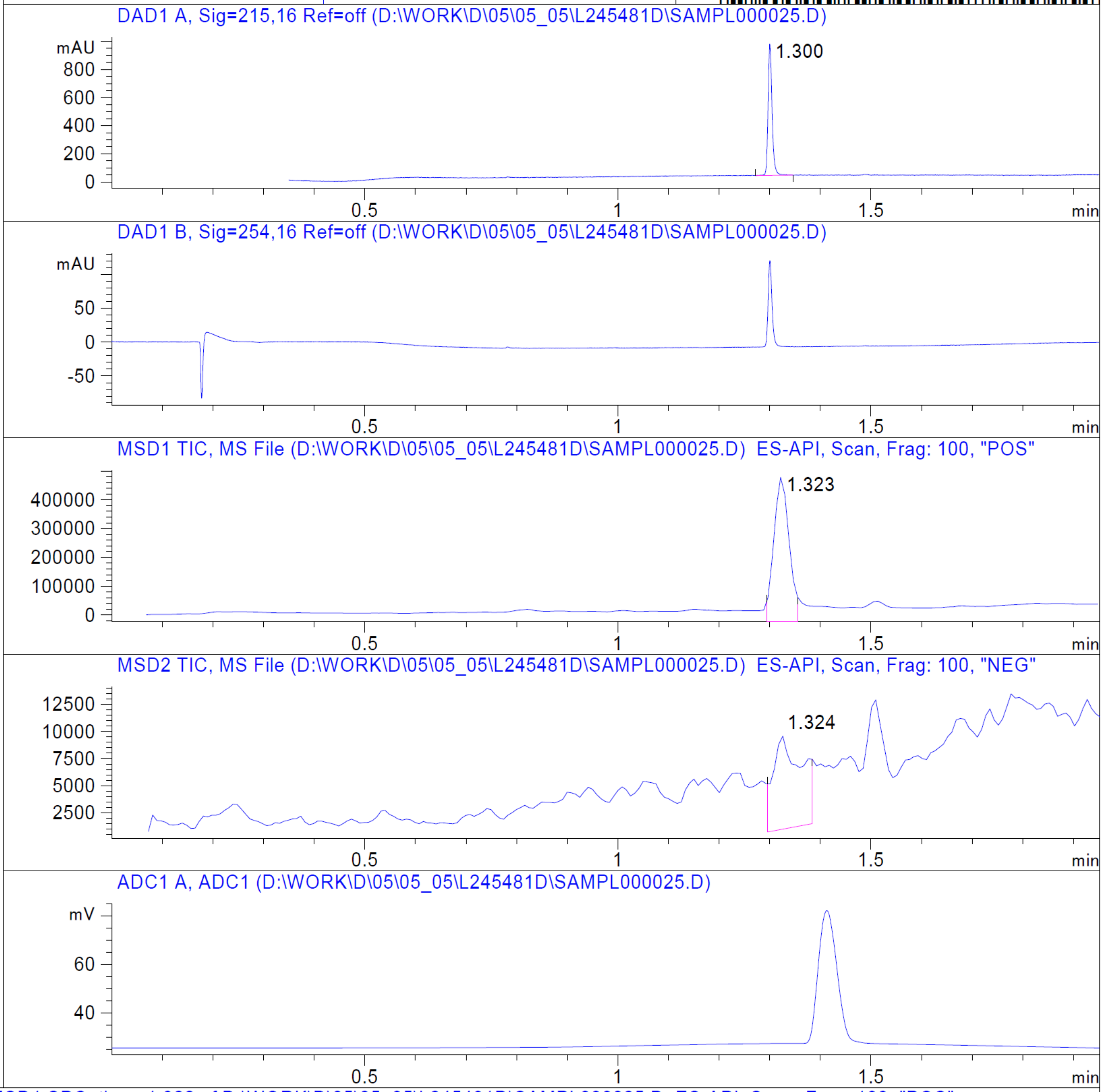


RT 1.323

RT 1.324

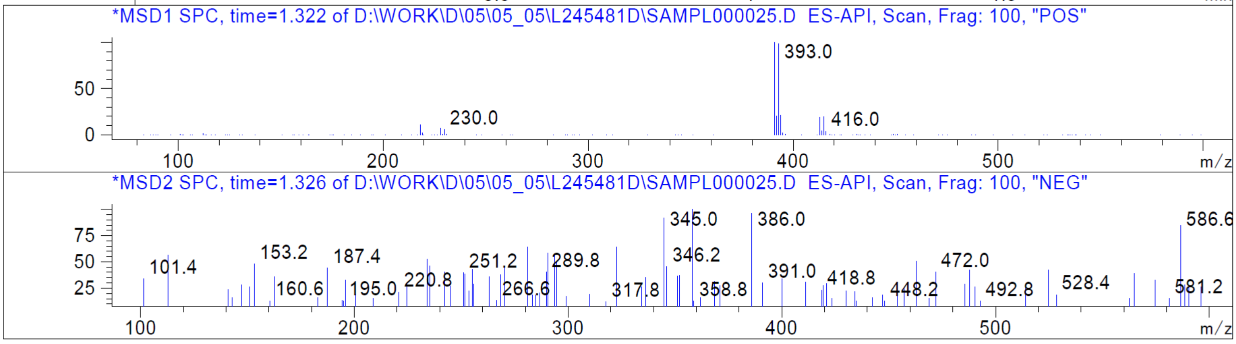


Z2230170963_(ZINC000553840273)

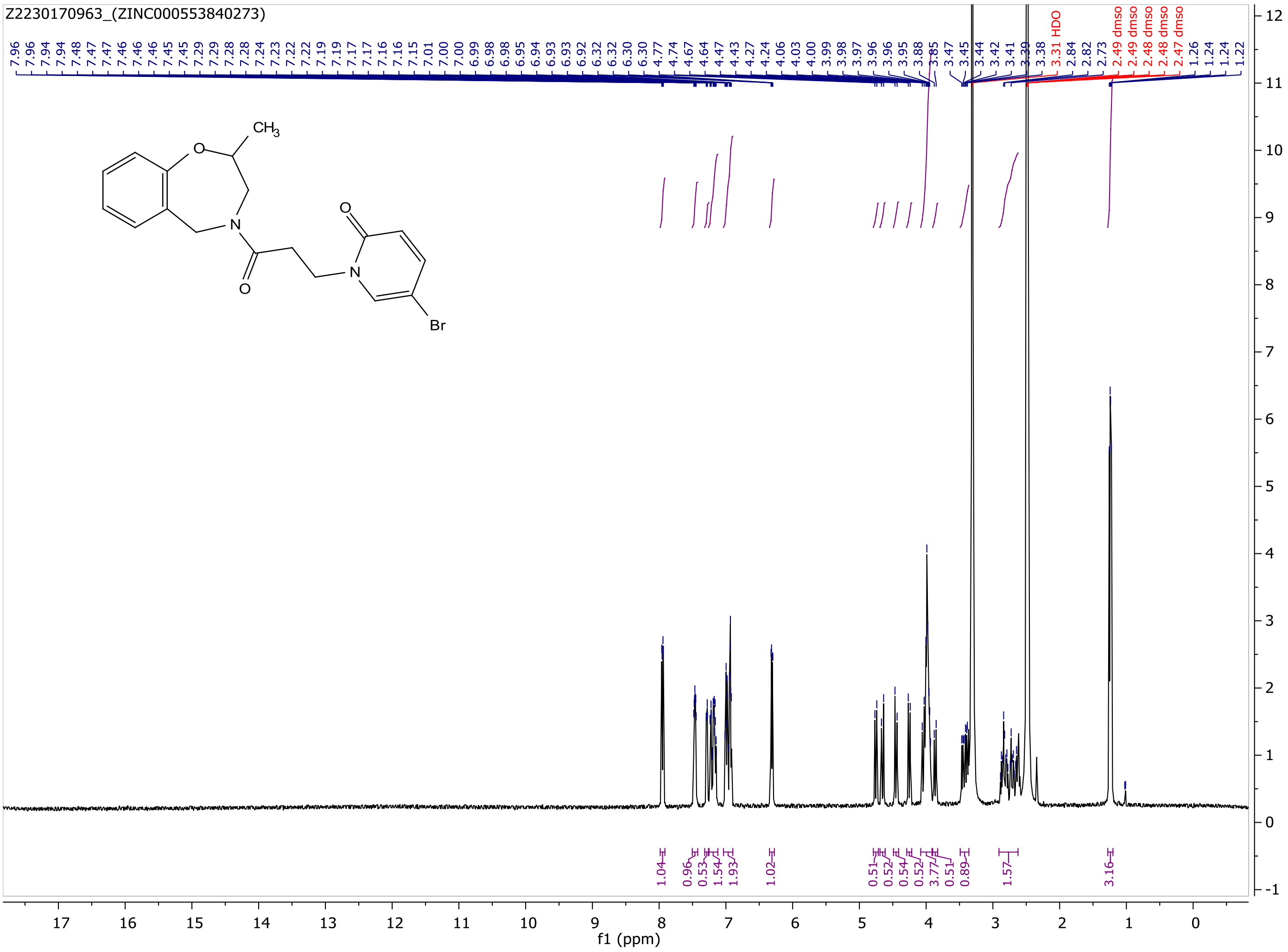
Z2230170963_(ZINC000553840273)
CH3
O
N
Br
HDO
dmso
f1 (ppm)

MaxPeak: 100.00%
Ret_Time: 1.498 min

# Q784757$G

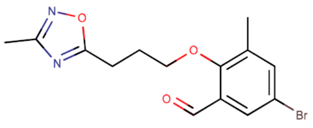

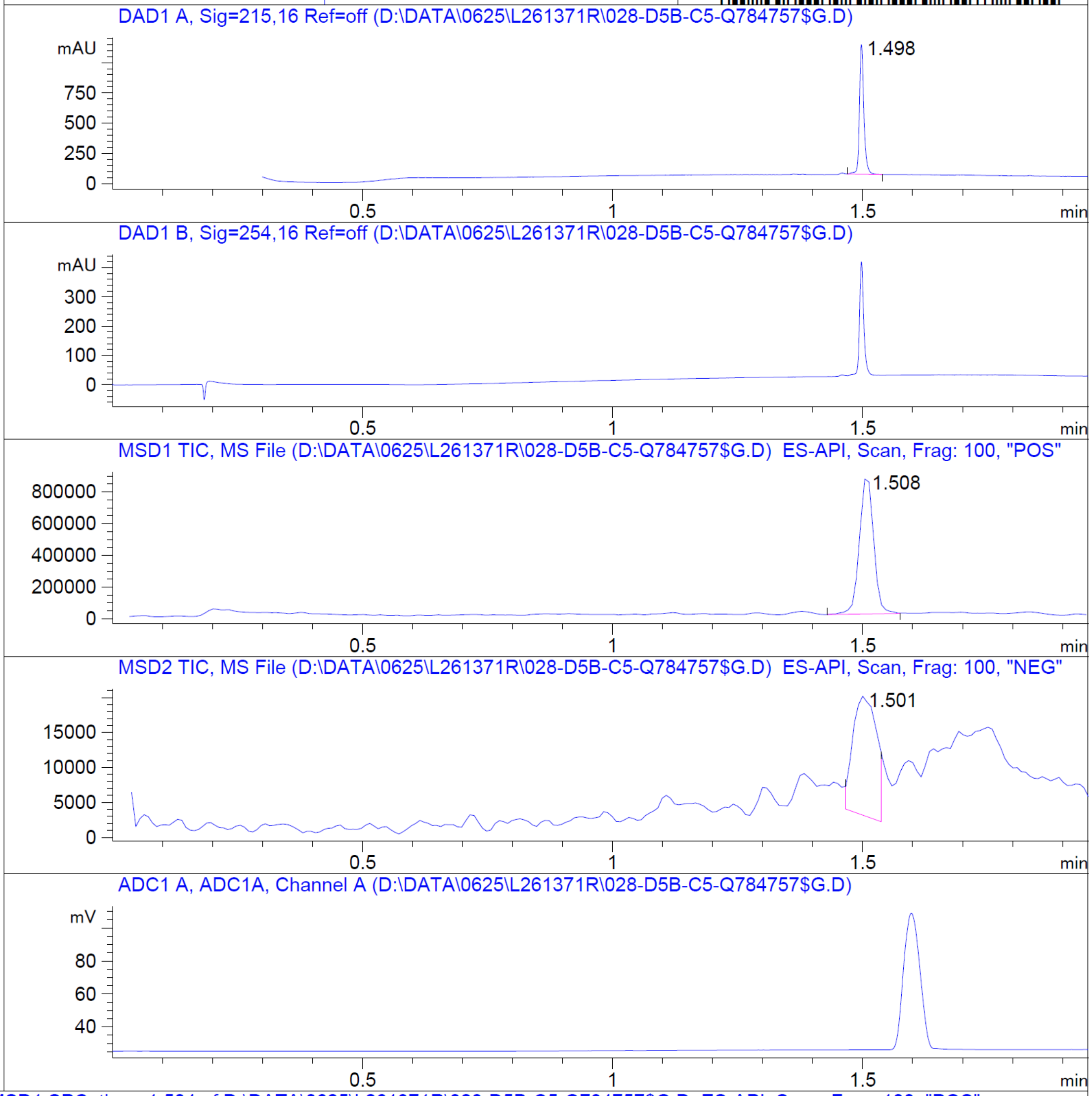


**Mol Wt** 339.18

**Exact Mass** 338.04

| # | Time | Area% |
|---|---|---|
| 1 | 1.498 | 100.00 |

RT 1.508

RT 1.501

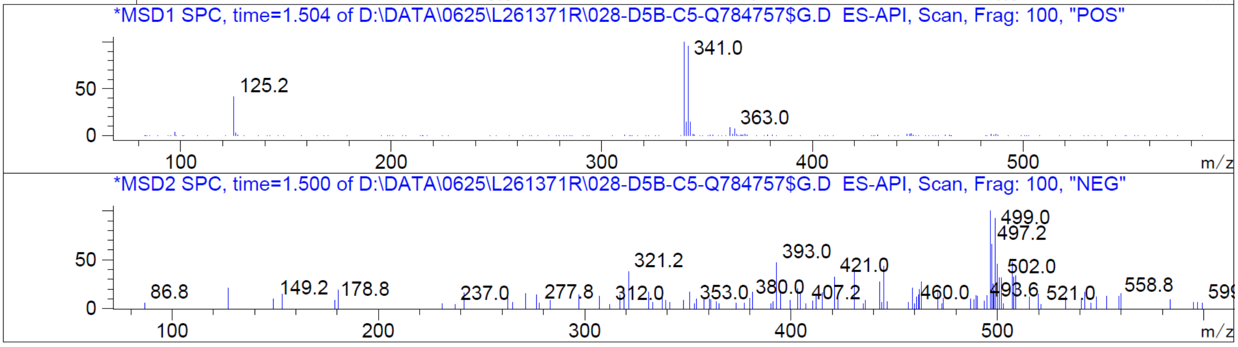


Z2686795662

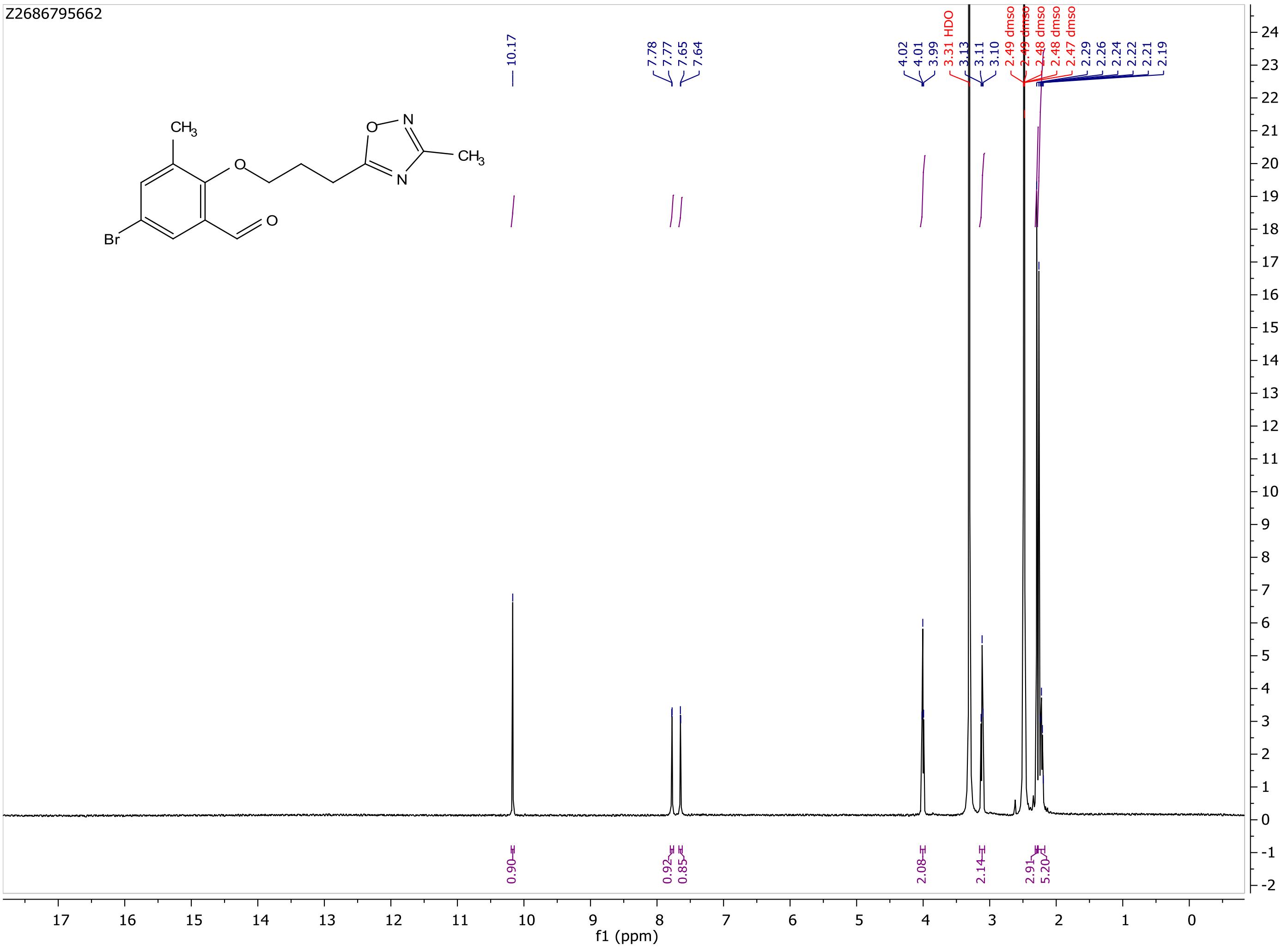
Z2686795662
CH3
O
O–N
N
CH3
Br
O
10.17
7.78
7.77
7.65
7.64
4.02
4.01
3.99
3.31 HDO
3.13
3.11
3.10
2.49 dmso
2.49 dmso
2.48 dmso
2.48 dmso
2.47 dmso
2.29
2.26
2.24
2.22
2.21
2.19
0.90
0.92
0.85
2.08
2.14
2.91
5.20
f1 (ppm)

MaxPeak: 100.00%
Ret_Time: 1.142 min

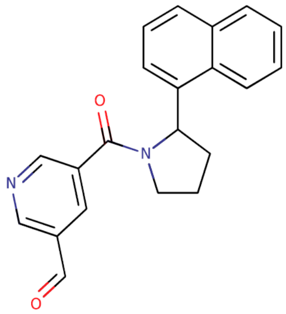

**Mol Wt** 330.38

**Exact Mass** 330.16

| # | Time | Area% |
|---|---|---|
| 1 | 1.142 | 100.00 |

Q763572$3

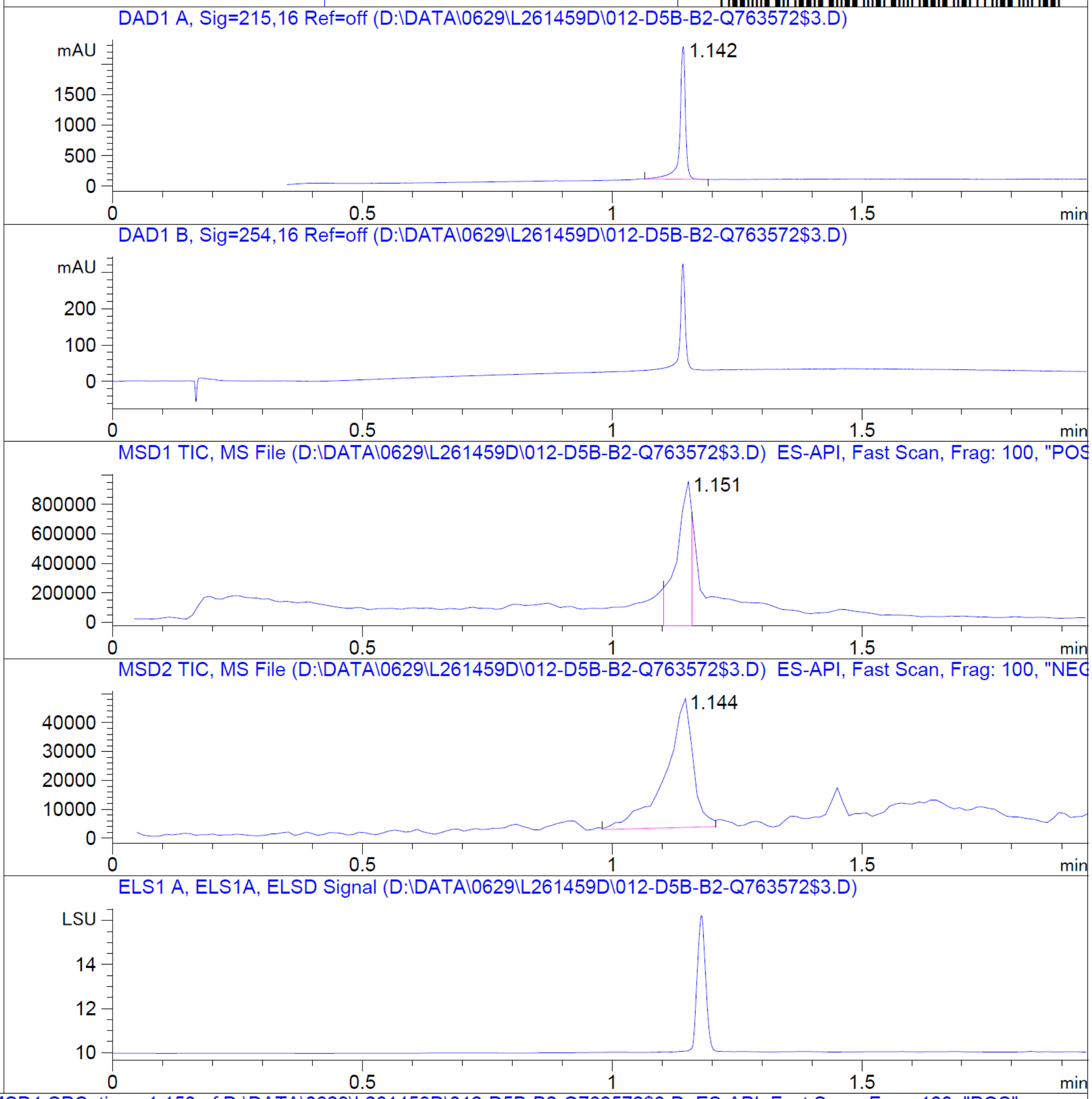


RT 1.151

RT 1.144

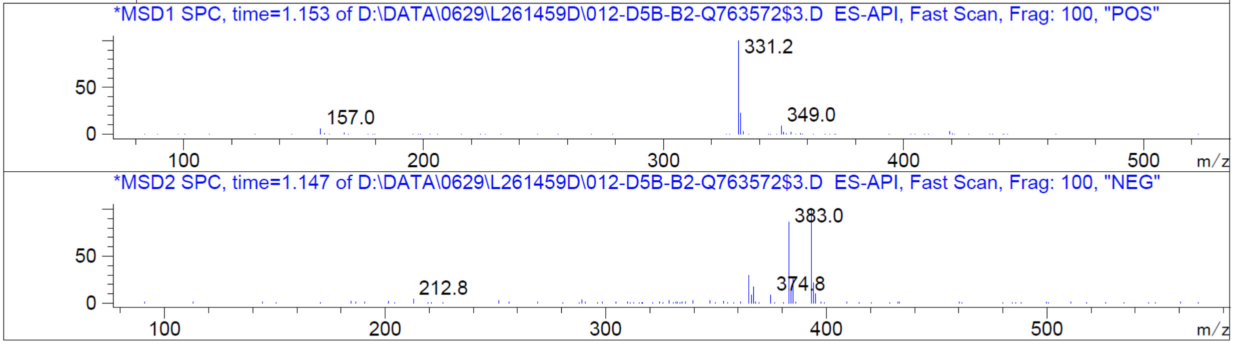


Z4497434908

Z4497434908

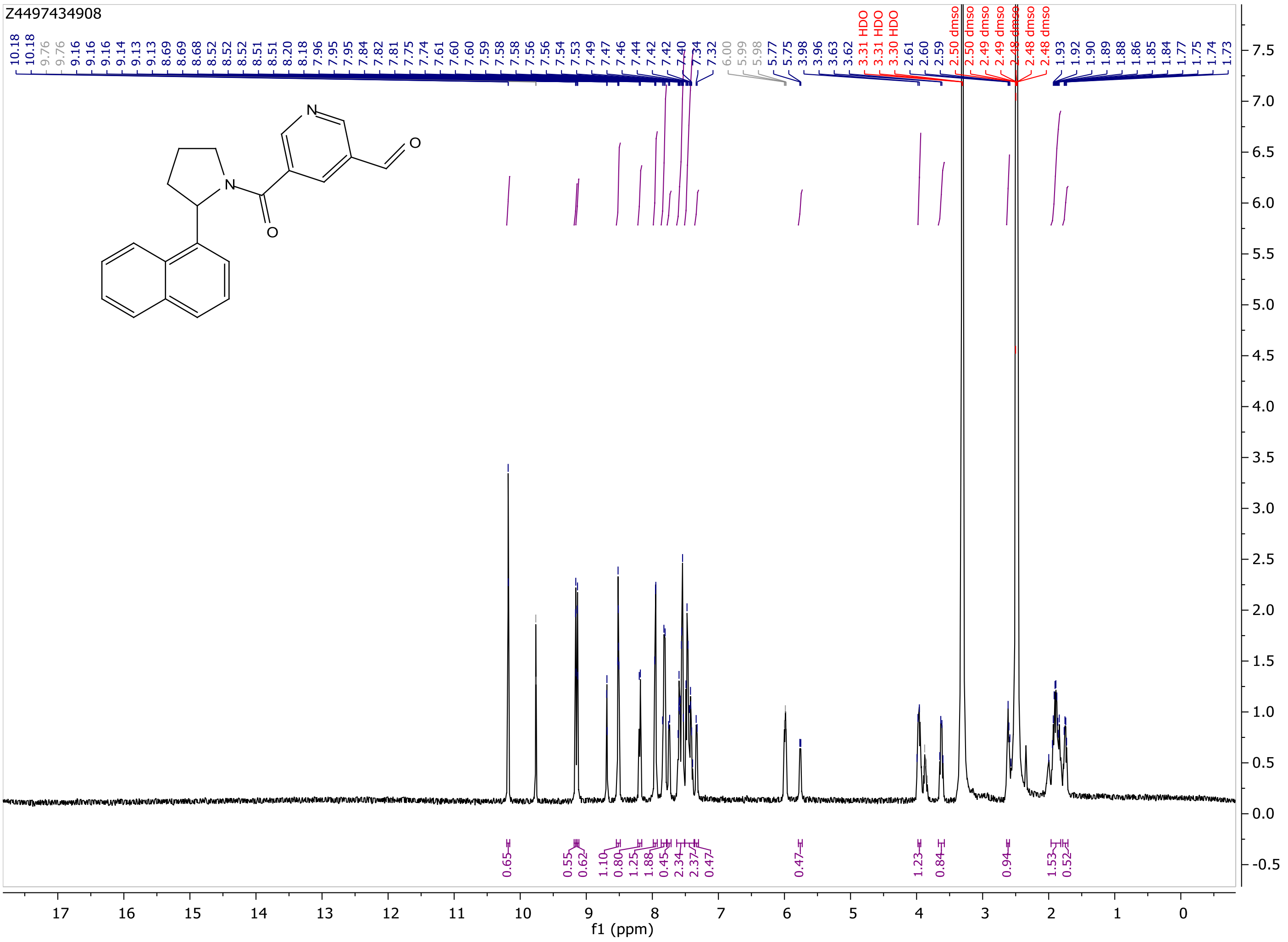

MaxPeak: 100.00%
Ret_Time: 1.477 min

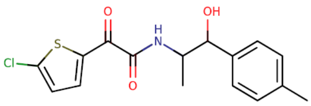

**Mol Wt** 337.82
**Exact Mass** 337.07

| # | Time | Area% |
|---|---|---|
| 1 | 1.477 | 100.00 |

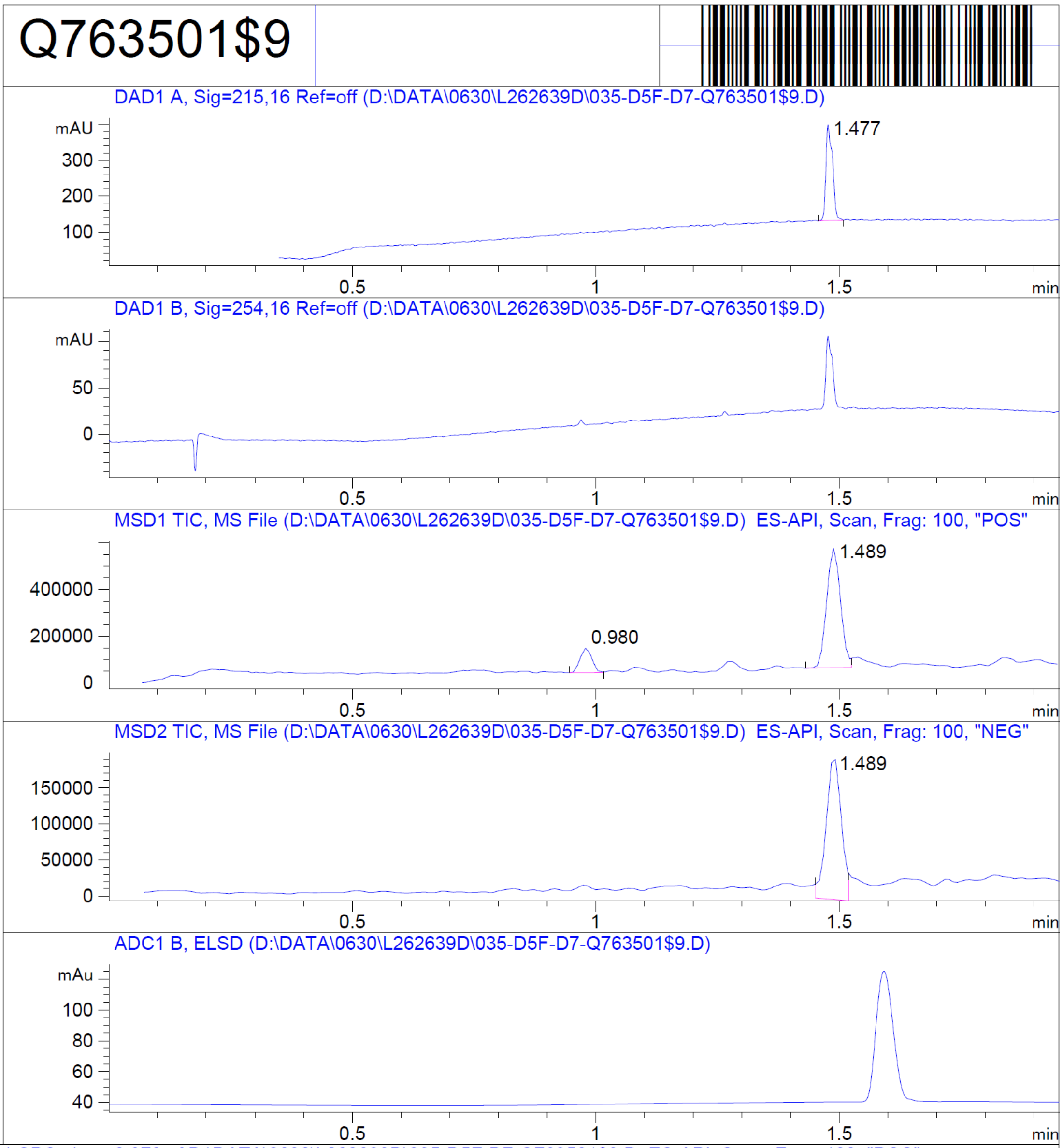


RT 0.980

RT 1.489

RT 1.489

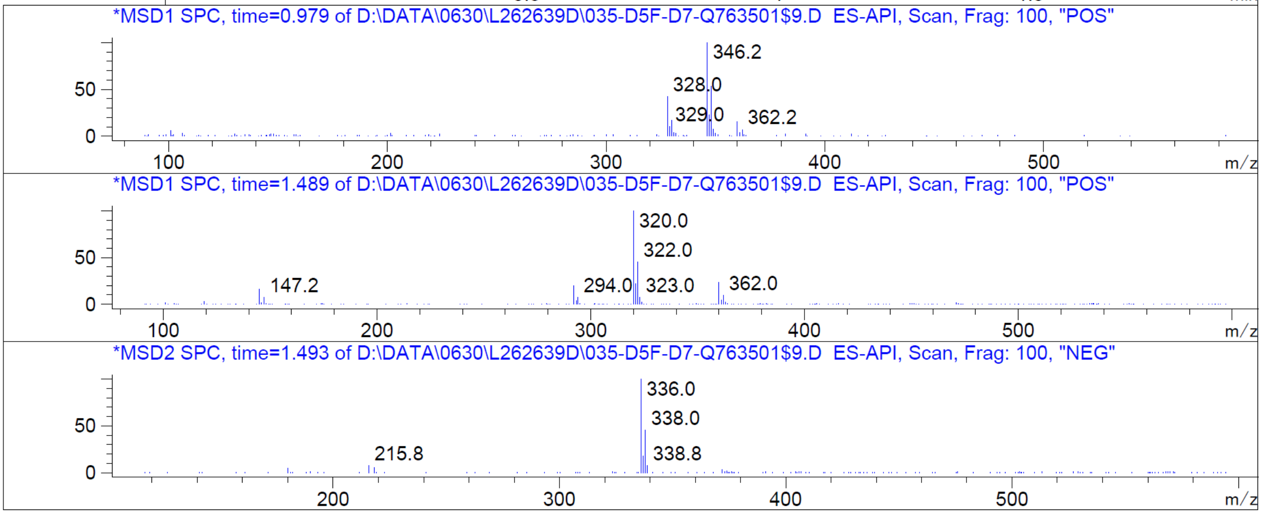


Z2091248890

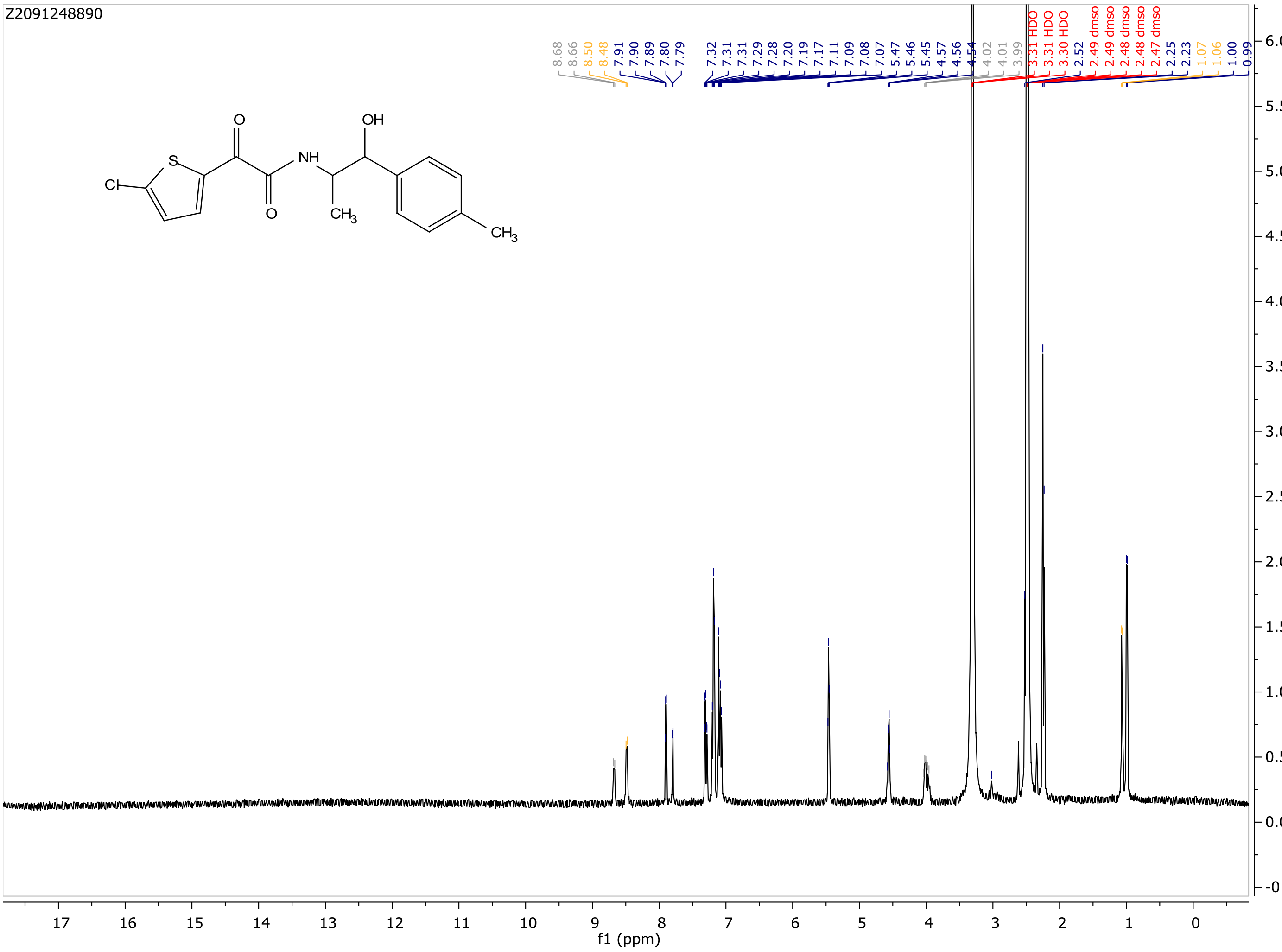
Z2091248890
8.68
8.66
8.50
8.48
7.91
7.90
7.89
7.80
7.79
7.32
7.31
7.31
7.29
7.28
7.20
7.19
7.17
7.11
7.09
7.08
7.07
5.47
5.46
5.45
4.57
4.56
4.54
4.02
4.01
3.99
3.31 HDO
3.31 HDO
3.30 HDO
2.52
2.49 dmso
2.49 dmso
2.48 dmso
2.48 dmso
2.47 dmso
2.25
2.23
1.07
1.06
1.00
0.99
O
S
Cl
NH
OH
O
CH3
CH3
f1 (ppm)

MaxPeak: 100.00%
Ret_Time: 1.053 min

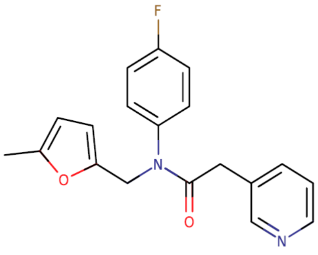

**Mol Wt** 324.35
**Exact Mass** 324.15

| # | Time | Area% |
|---|---|---|
| 1 | 1.053 | 100.00 |

Q797696$1

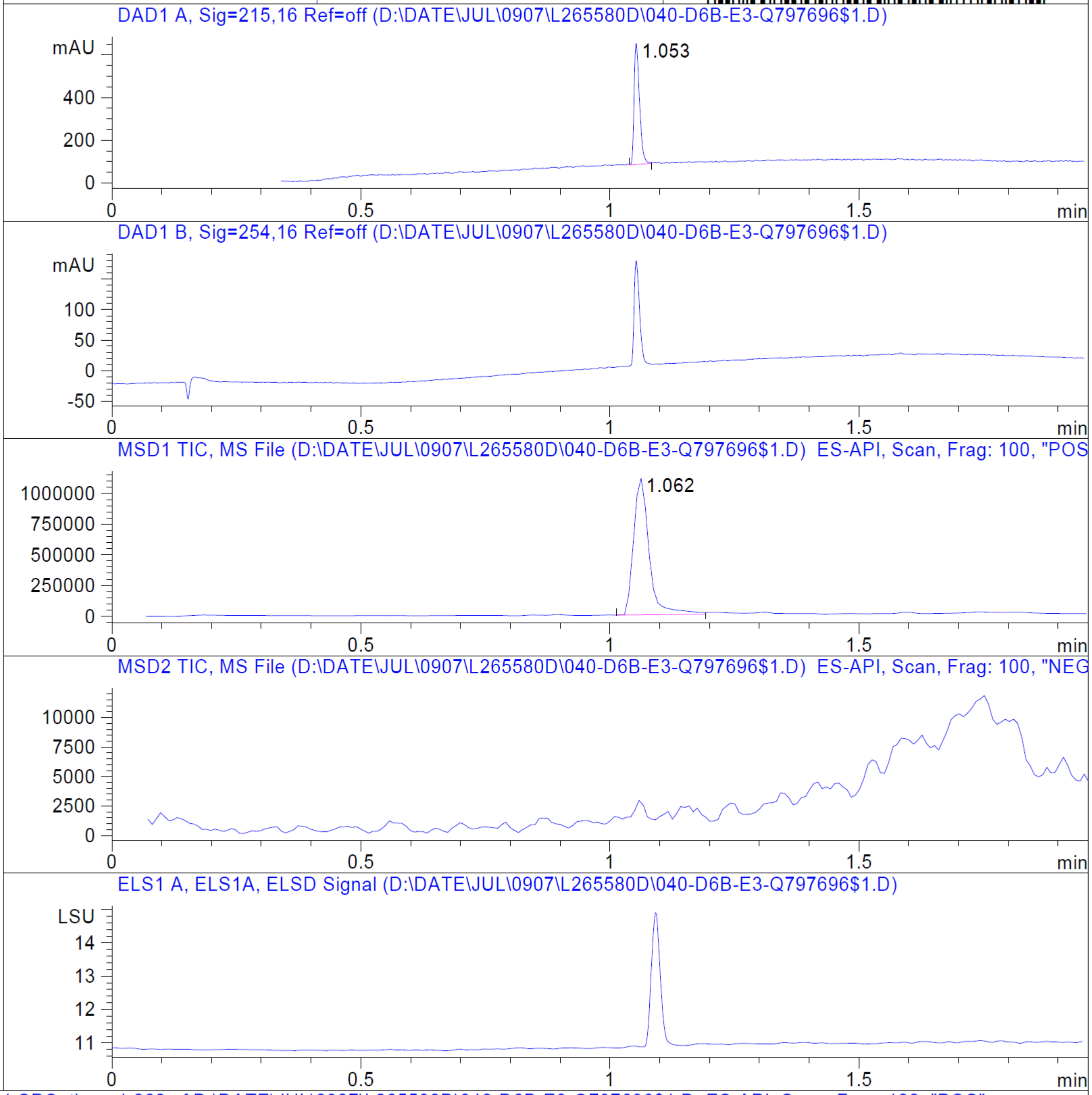


RT 1.062

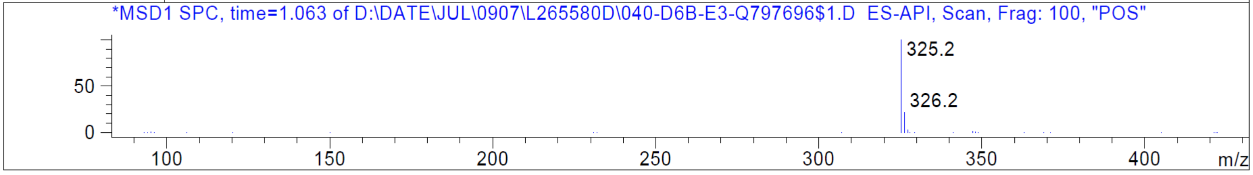


Z1173082912

Z1173082912

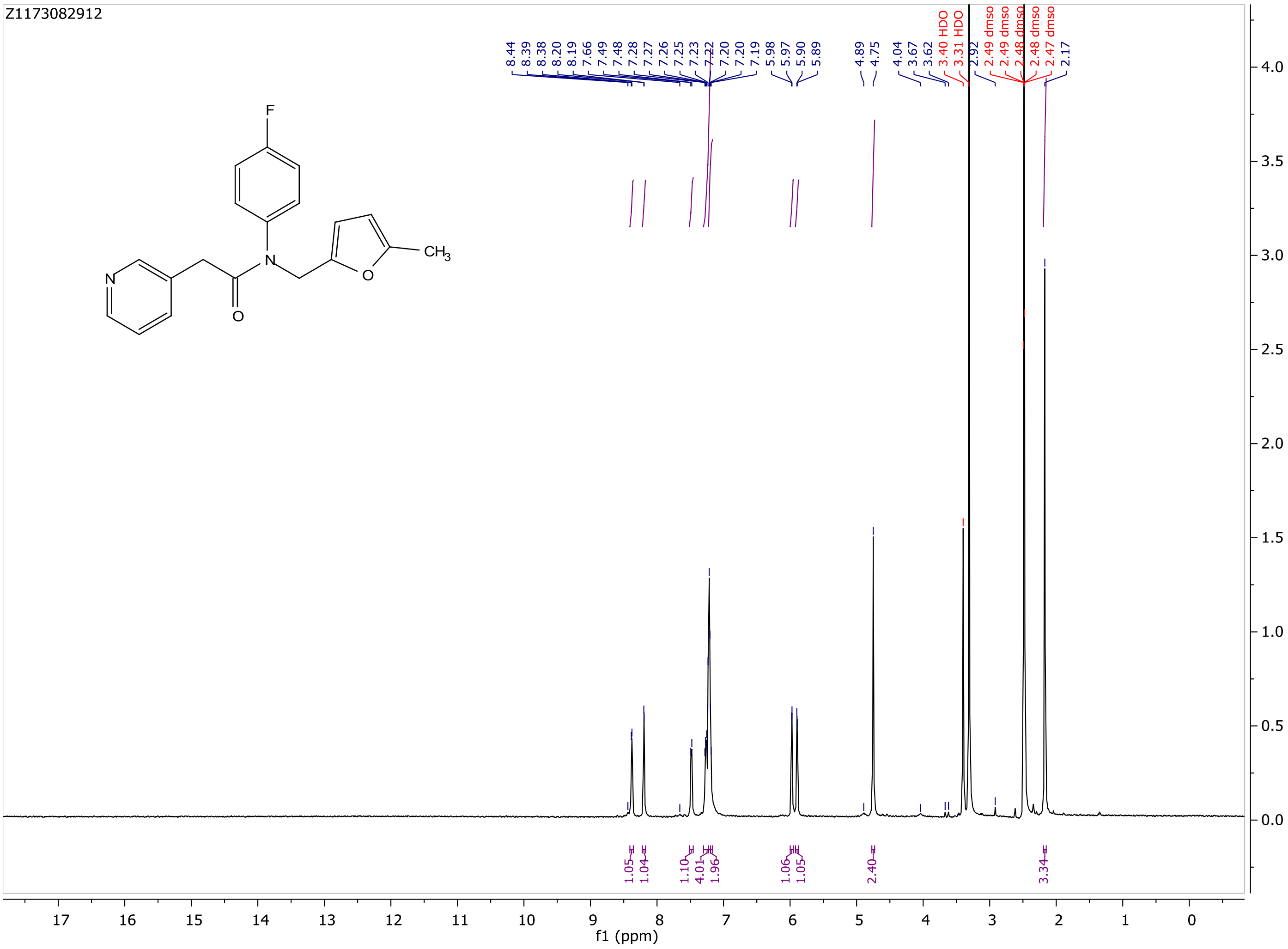

MaxPeak: 100.00%
Ret_Time: 1.436 min

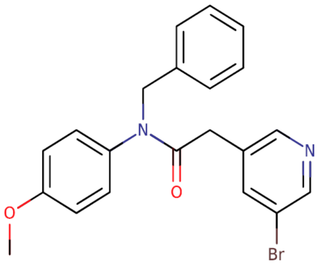


**Mol Wt** 411.29

**Exact Mass** 412.09

| # | Time | Area% |
|---|---|---|
| 1 | 1.436 | 100.00 |

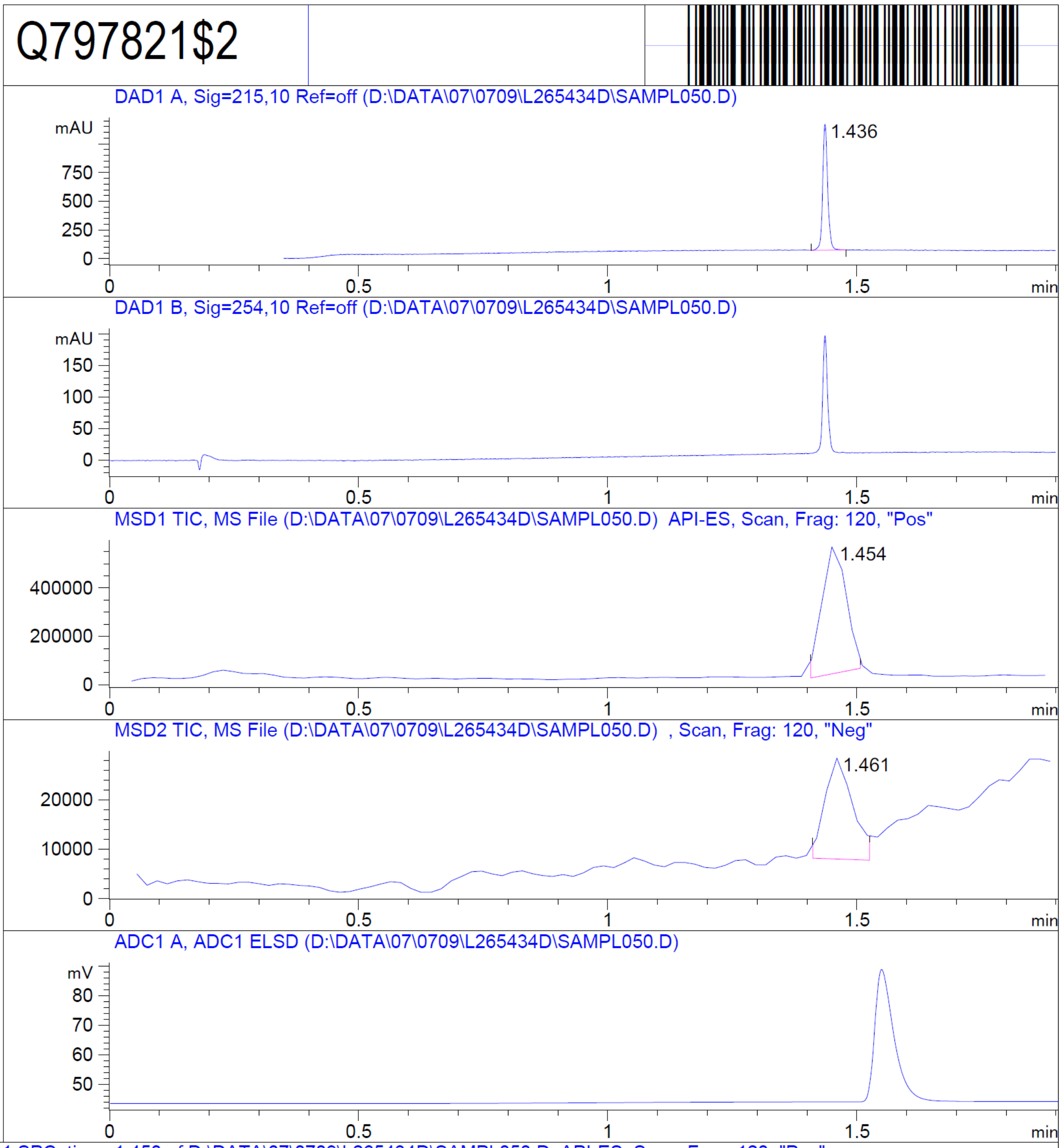


RT 1.454

RT 1.461

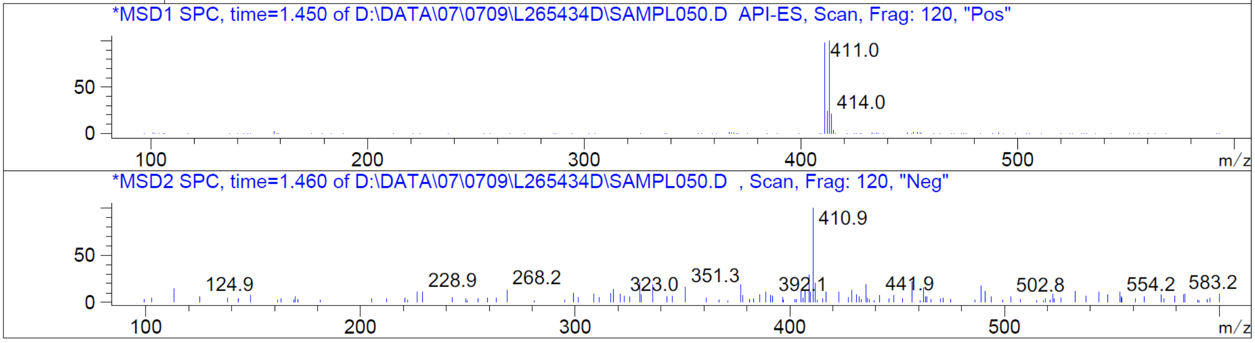


Z4521849273

Z4521849273

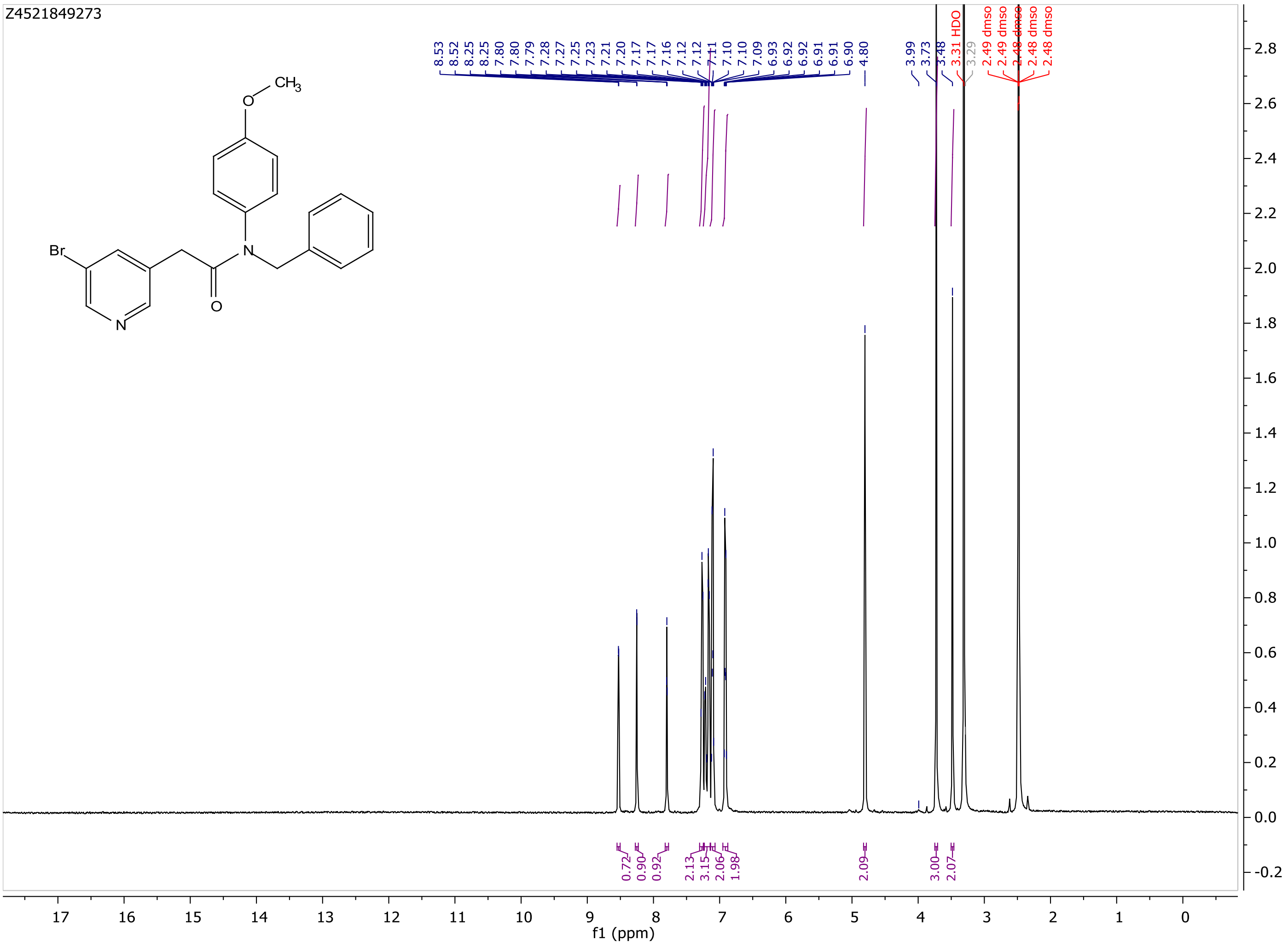

MaxPeak: 100.00%
Ret_Time: 1.177 min

# Q797685$1

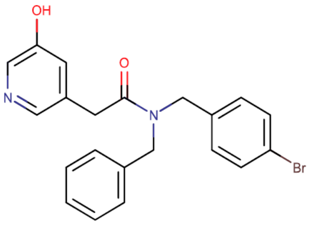

**Mol Wt** 411.29

**Exact Mass** 412.09

| # | Time | Area% |
|---|---|---|
| 1 | 1.177 | 100.00 |

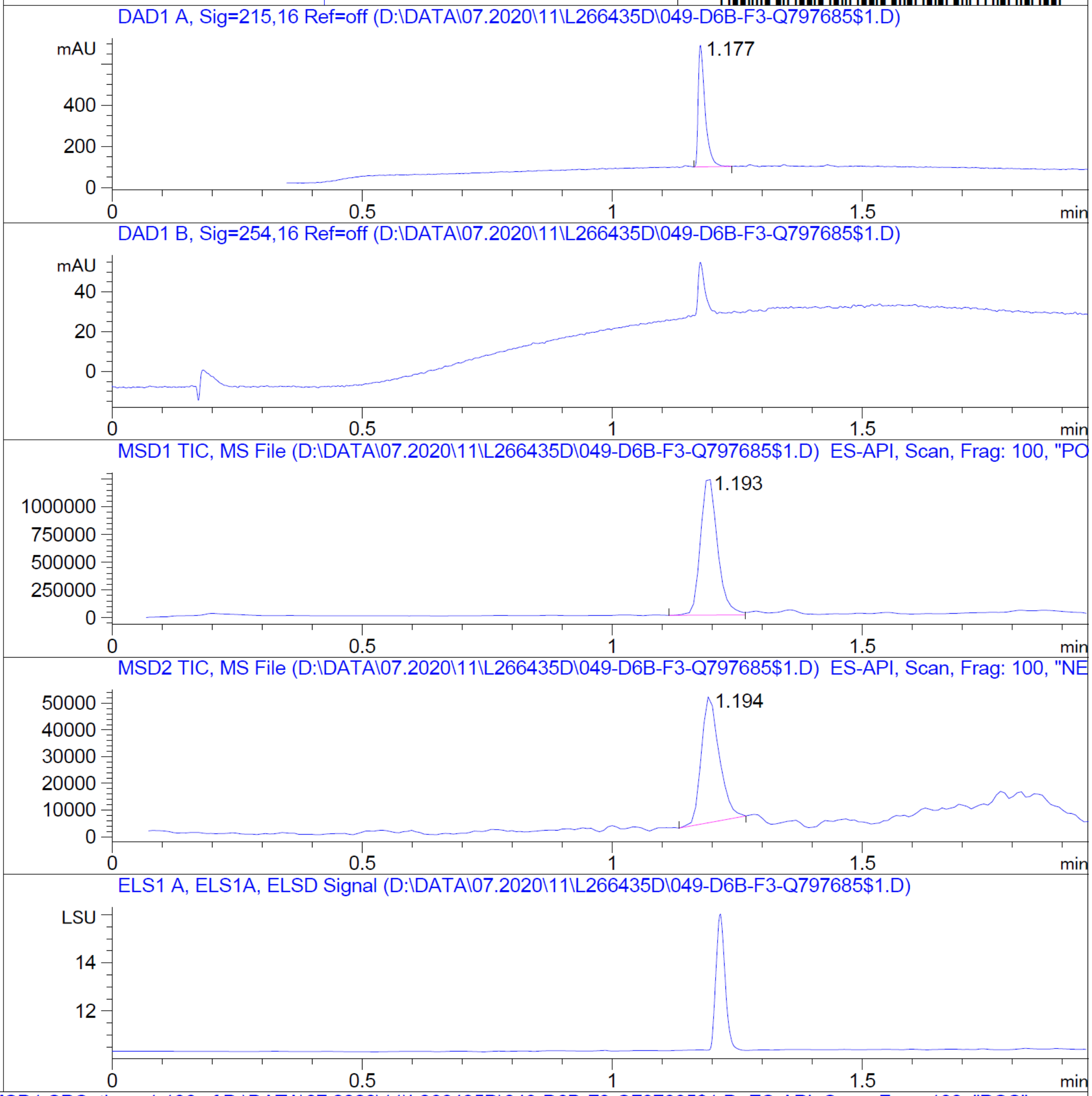


RT 1.193

RT 1.194

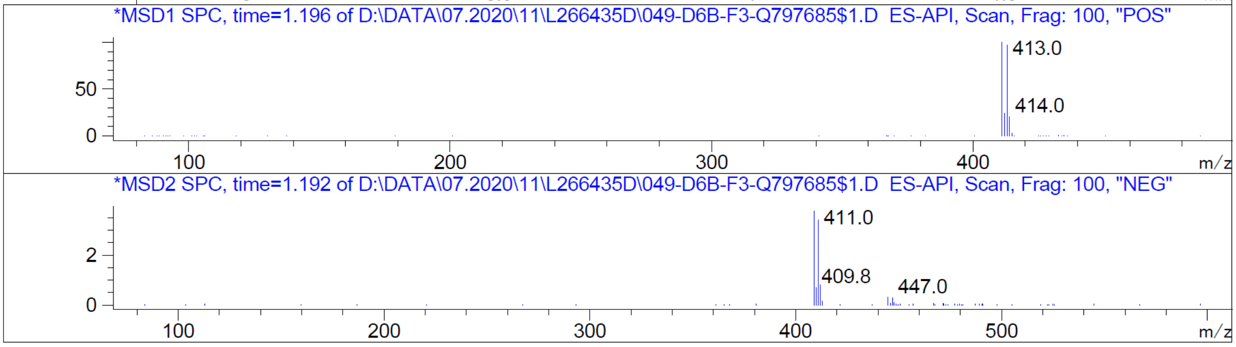


Z4521849300

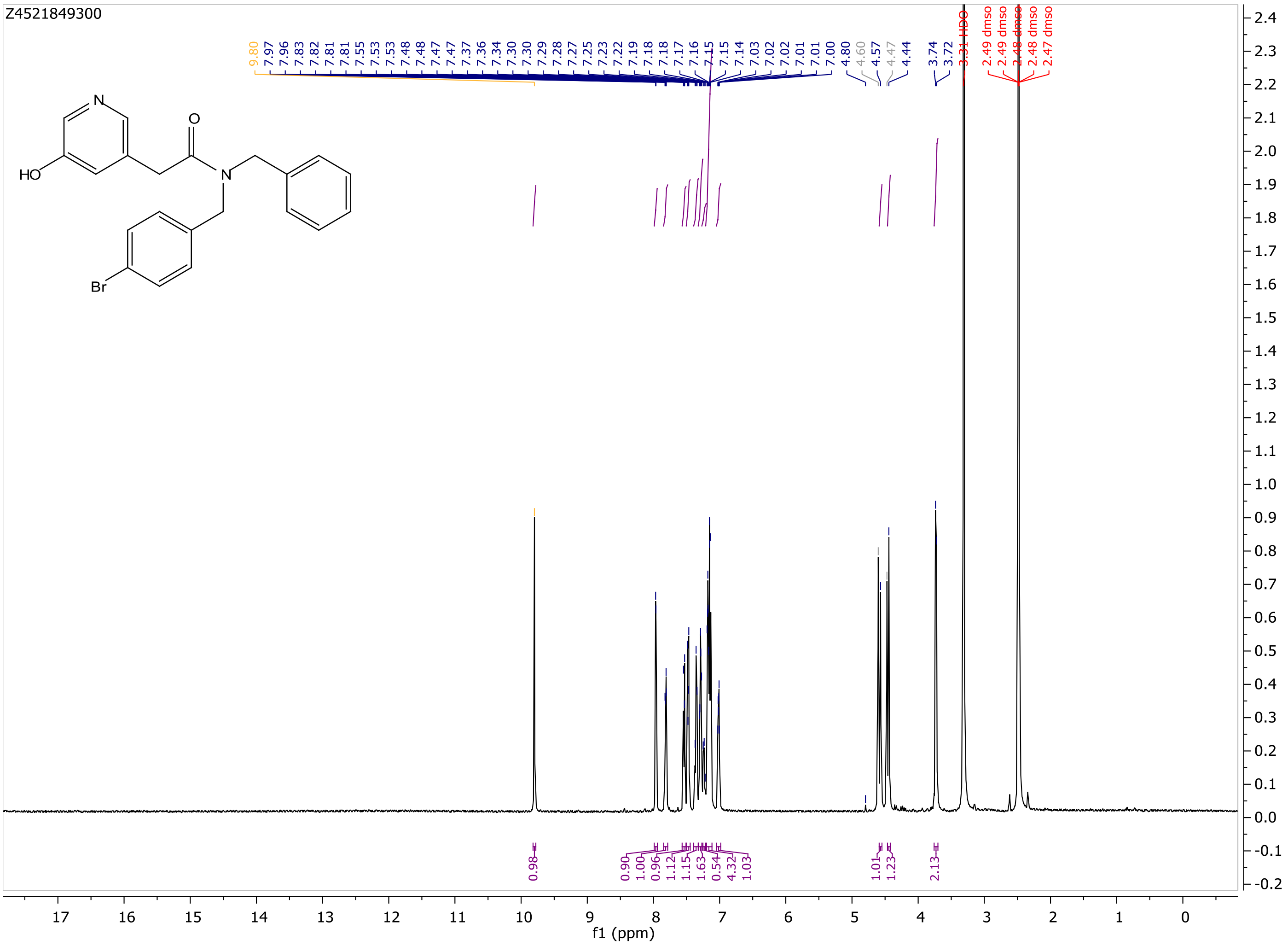
Z4521849300
HO
N
O
Br
9.80
7.97
7.96
7.83
7.82
7.81
7.81
7.55
7.53
7.53
7.48
7.48
7.47
7.47
7.37
7.36
7.34
7.30
7.30
7.29
7.28
7.27
7.25
7.23
7.22
7.19
7.18
7.18
7.17
7.16
7.15
7.15
7.14
7.03
7.02
7.02
7.01
7.01
7.00
4.80
4.60
4.57
4.47
4.44
3.74
3.72
3.31 HDO
2.49 dmso
2.49 dmso
2.48 dmso
2.48 dmso
2.47 dmso
0.98
0.90
1.00
0.96
1.12
1.15
1.63
0.54
4.32
1.03
1.01
1.23
2.13
f1 (ppm)

MaxPeak: 100.00%
Ret_Time: 2.429 min

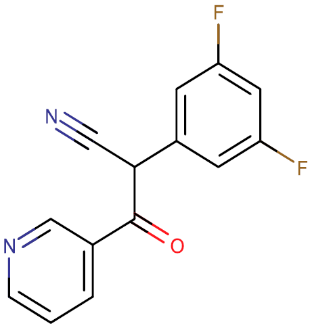

**Mol Wt** 258.22

**Exact Mass** 258.07

| # | Time | Area% |
|---|---|---|
| 1 | 2.429 | 100.00 |

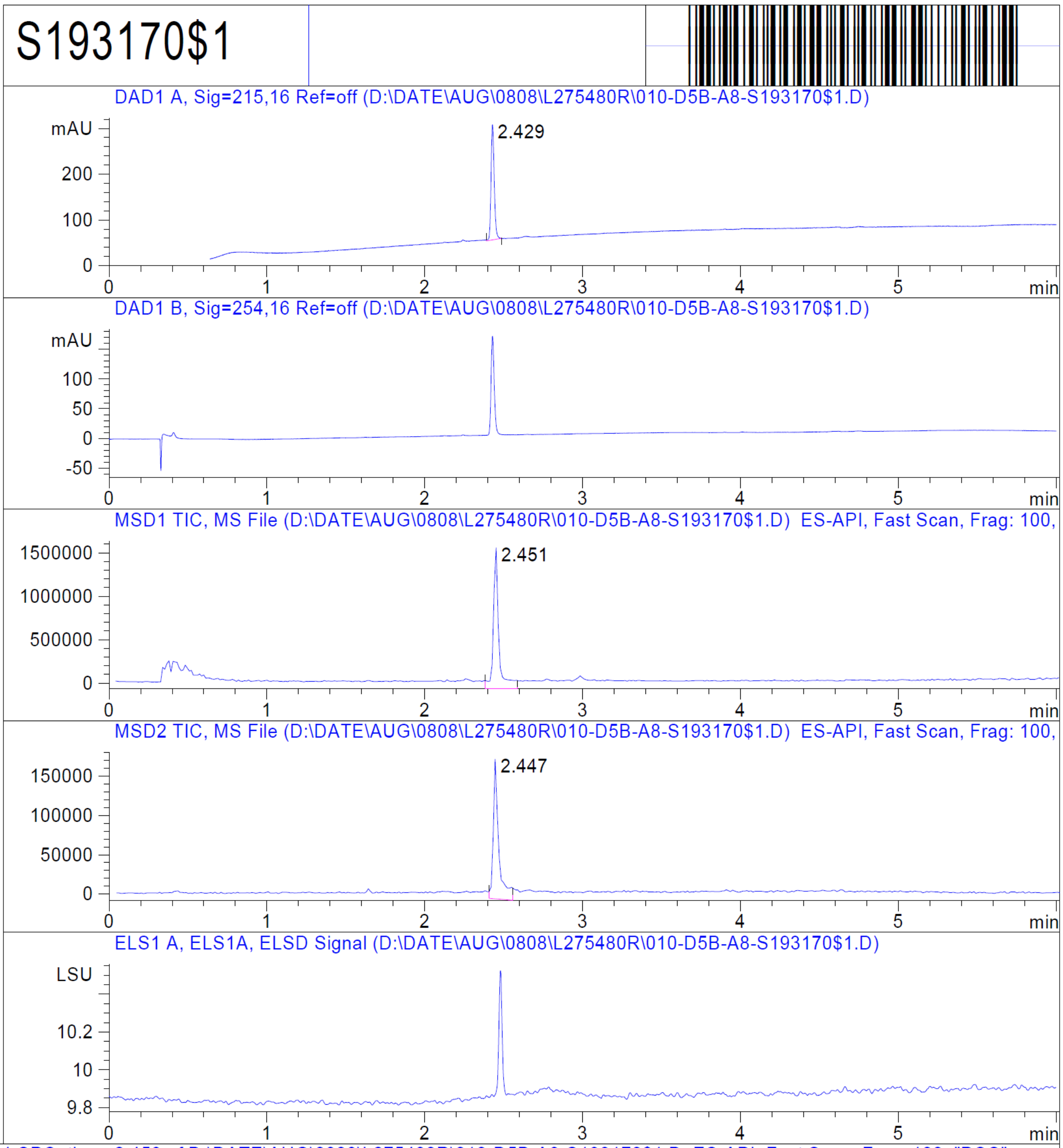


RT 2.451

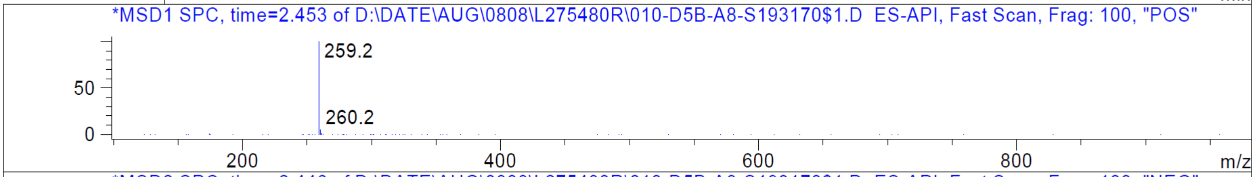


RT 2.447

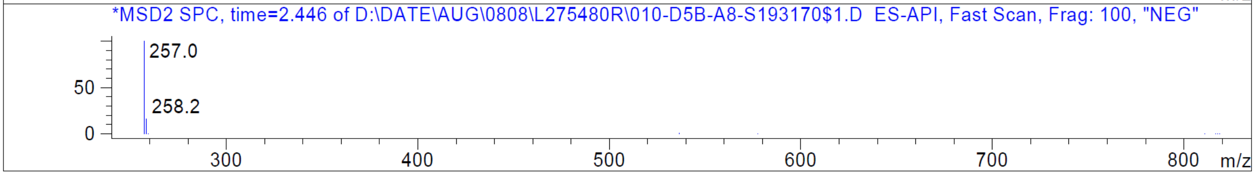


Z821056998

Z821056998

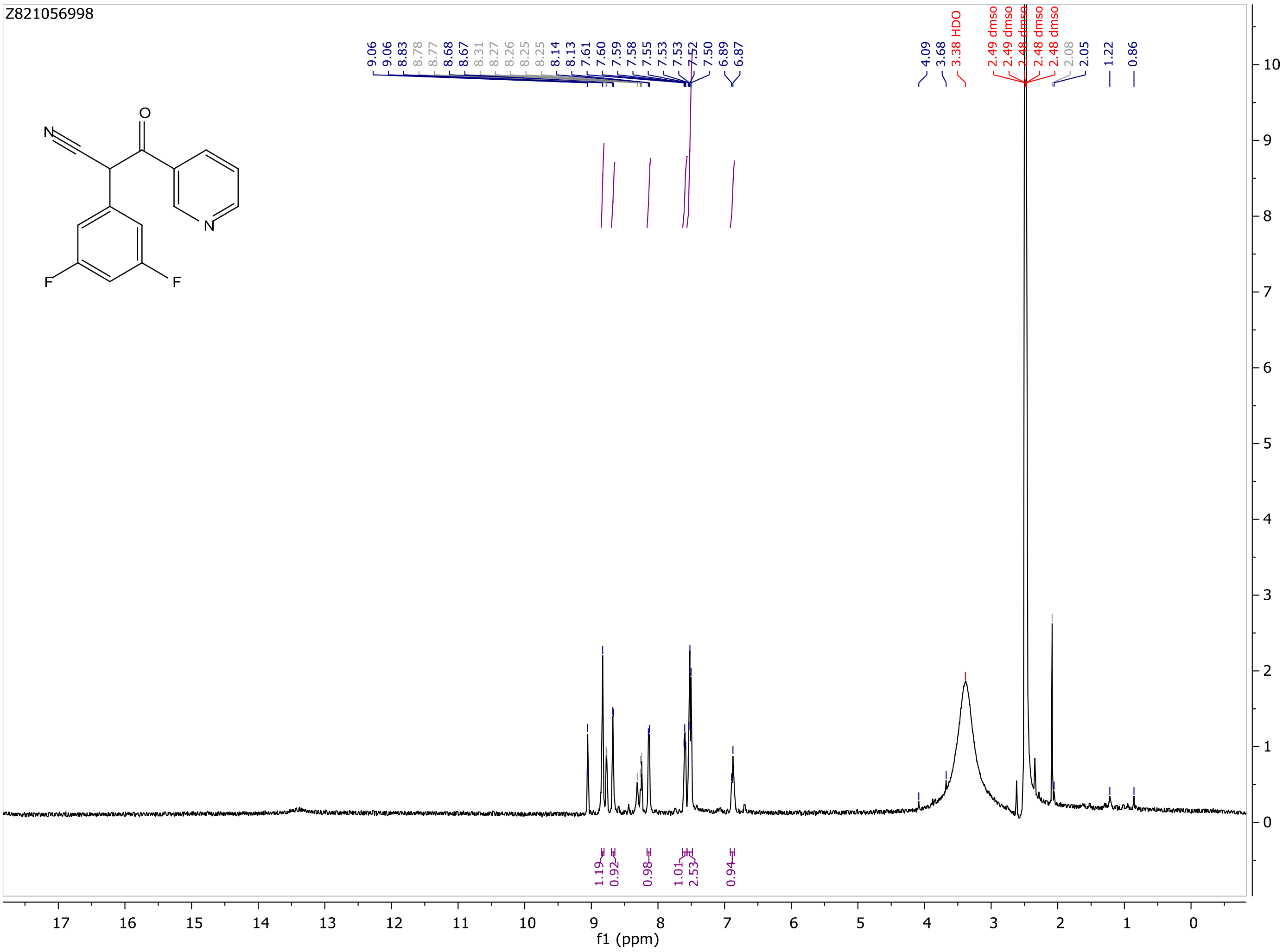

9.06
9.06
8.83
8.78
8.77
8.68
8.67
8.31
8.27
8.26
8.25
8.25
8.14
8.13
7.61
7.60
7.59
7.58
7.55
7.53
7.53
7.52
7.50
6.89
6.87
4.09
3.68
3.38 HDO
2.49 dmso
2.49 dmso
2.48 dmso
2.48 dmso
2.48 dmso
2.08
2.05
1.22
0.86
1.19
0.92
0.98
1.01
2.53
0.94
f1 (ppm)

MaxPeak:  100.00%
Ret_Time: 1.081 min

S081650$1

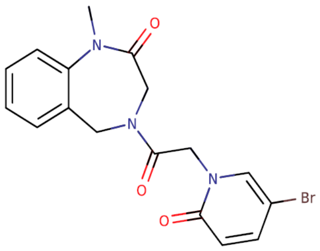


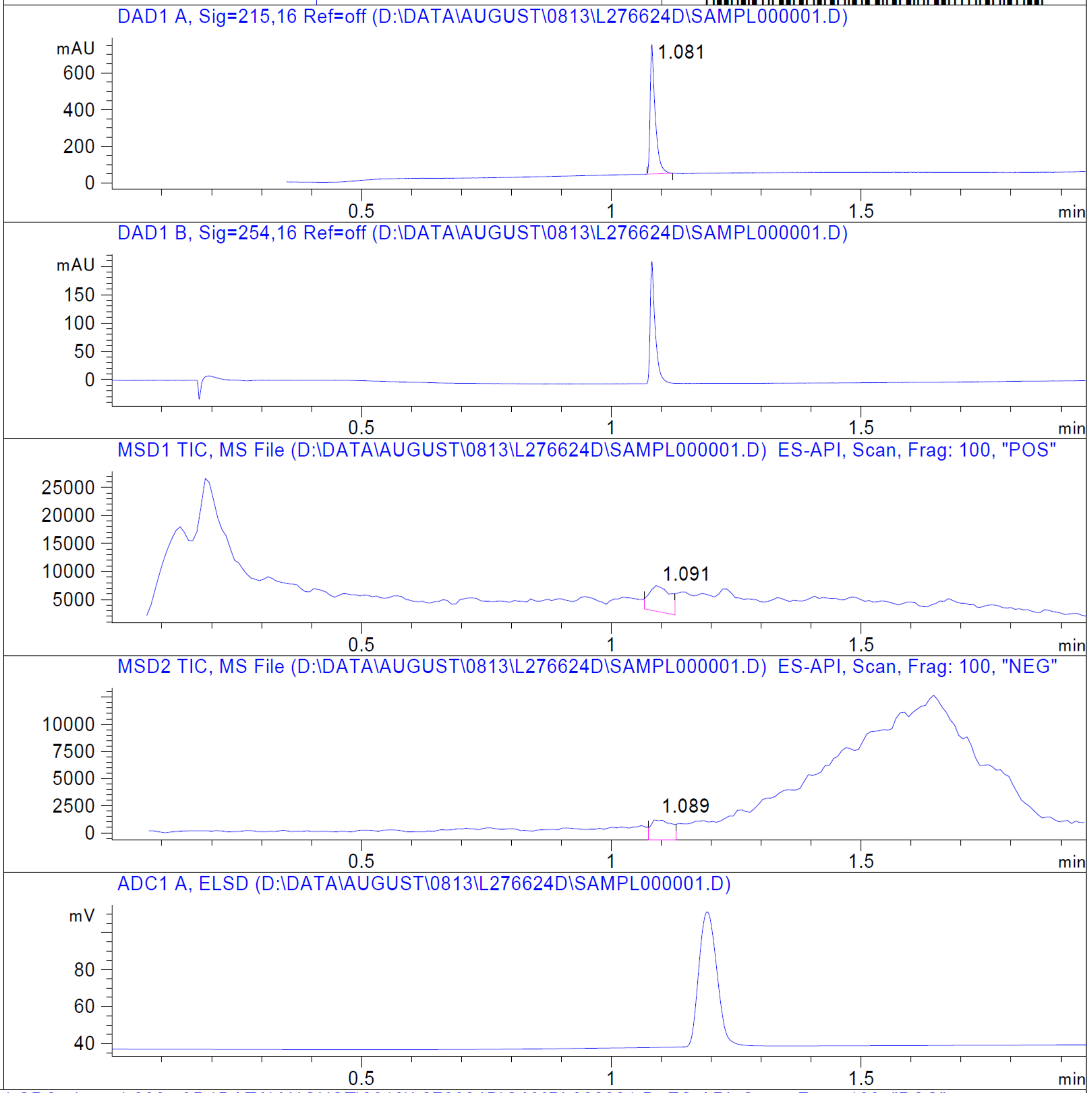


**Mol Wt** 390.23

**Exact Mass** 389.05

| # | Time | Area% |
| --- | --- | --- |
| 1 | 1.081 | 100.00 |

RT 1.091

RT 1.089

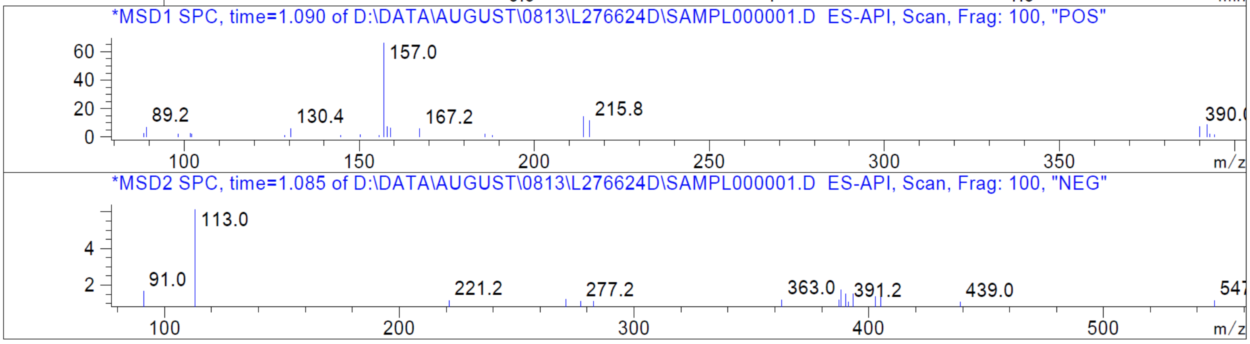


Z4594384266_(0273num9044)

Z4594384266_(0273num9044)

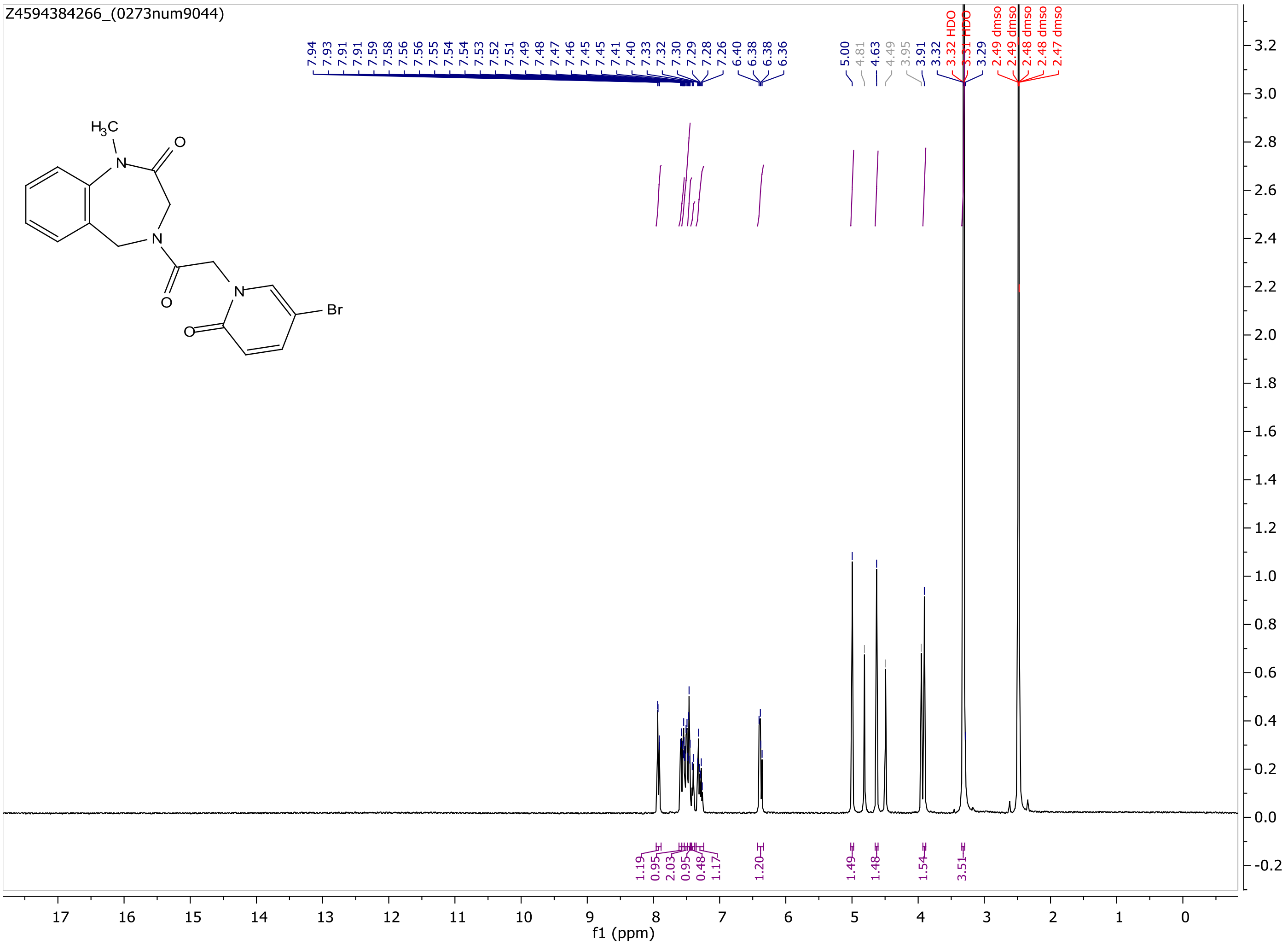

MaxPeak: 100.00%
Ret_Time: 0.941 min

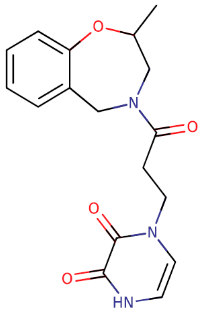

**Mol Wt** 329.35

**Exact Mass** 329.15

| # | Time | Area% |
|---|---|---|
| 1 | 0.941 | 100.00 |

S081659$1

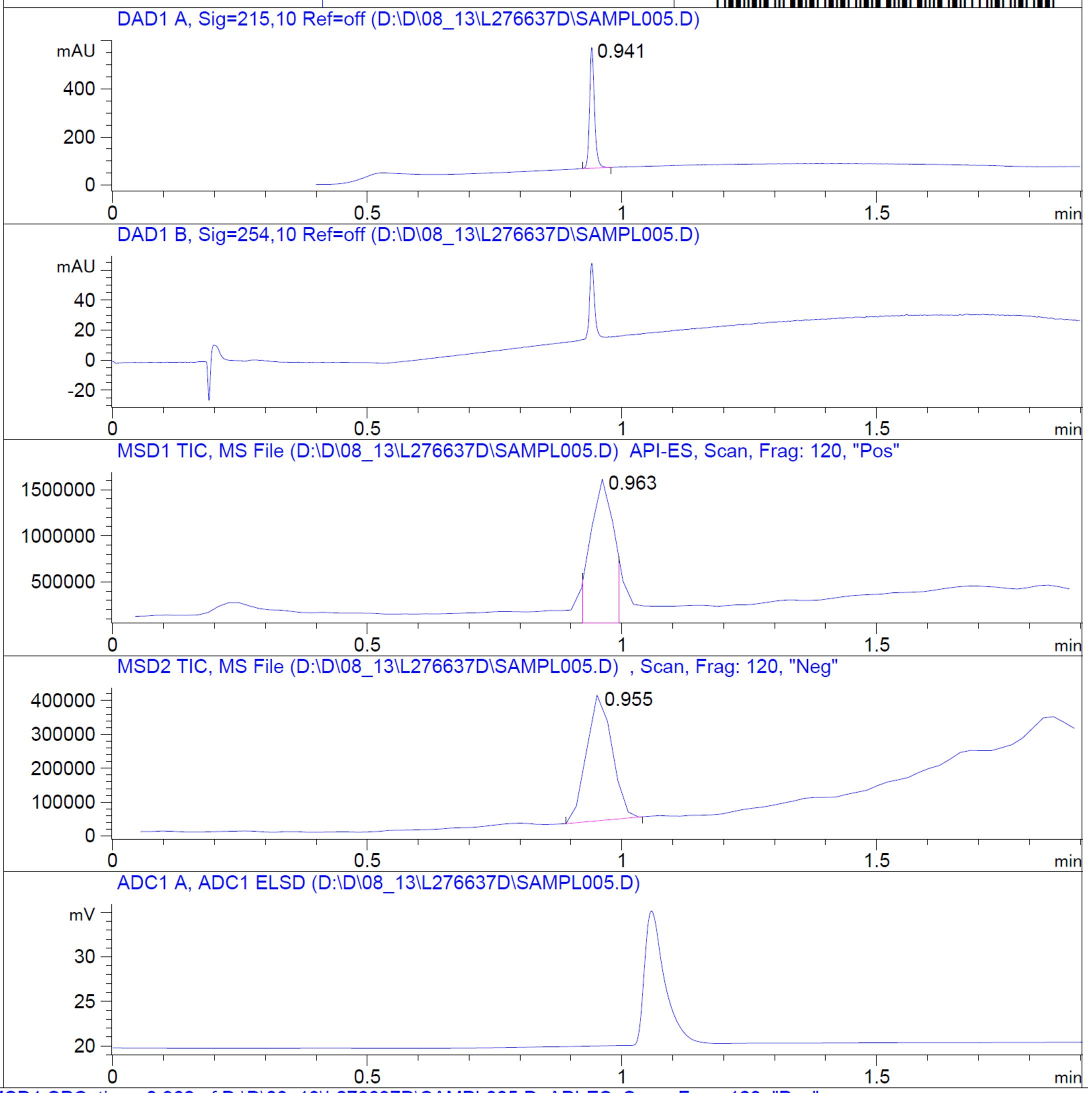


RT 0.963

RT 0.955

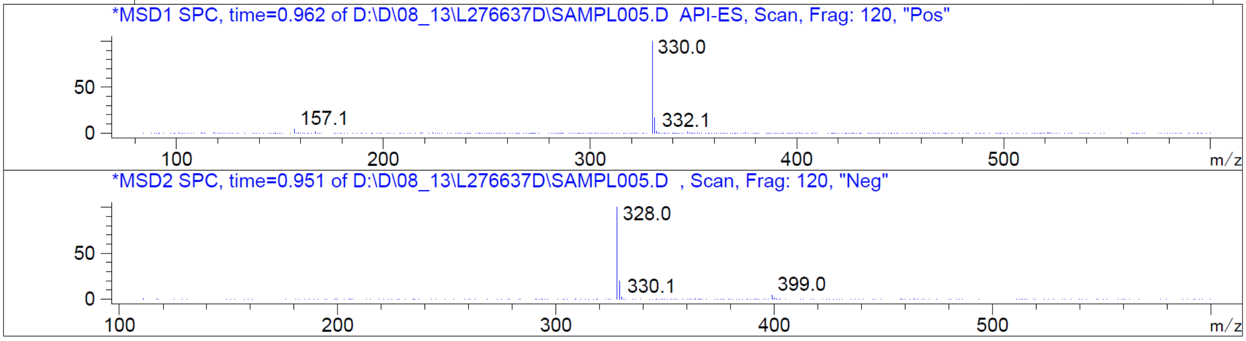


Z2230166392_(0273num3613.2)

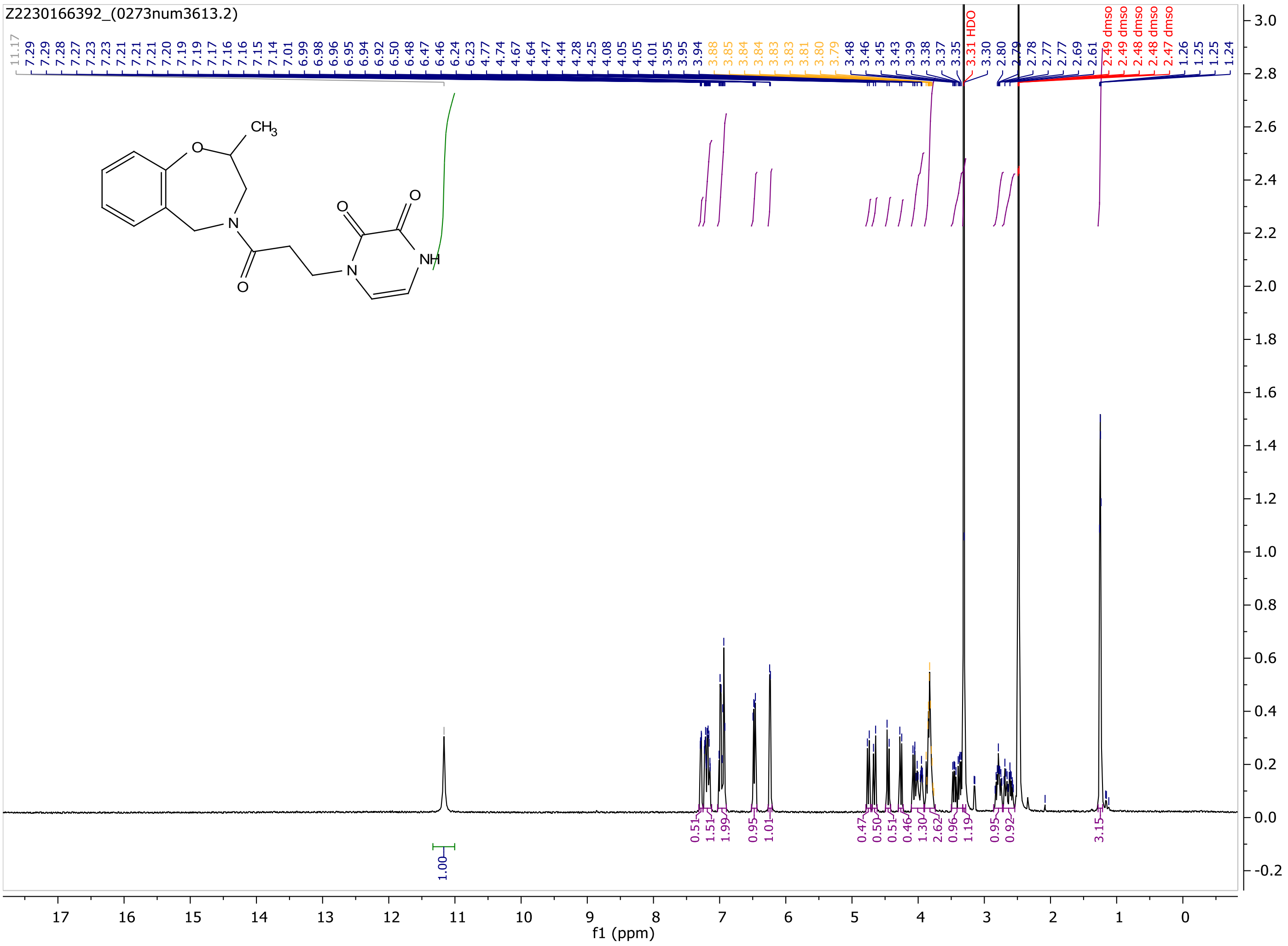
Z2230166392_(0273num3613.2)
f1 (ppm)
3.31 HDO
2.49 dmso
CH3
NH
O
N

MaxPeak: 100.00%
Ret_Time: 1.280 min

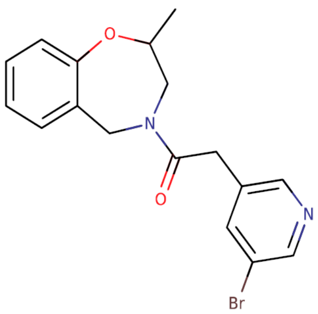


**Mol Wt** 361.23
**Exact Mass** 360.07

| # | Time | Area% |
| --- | --- | --- |
| 1 | 1.280 | 100.00 |

# S081639$2

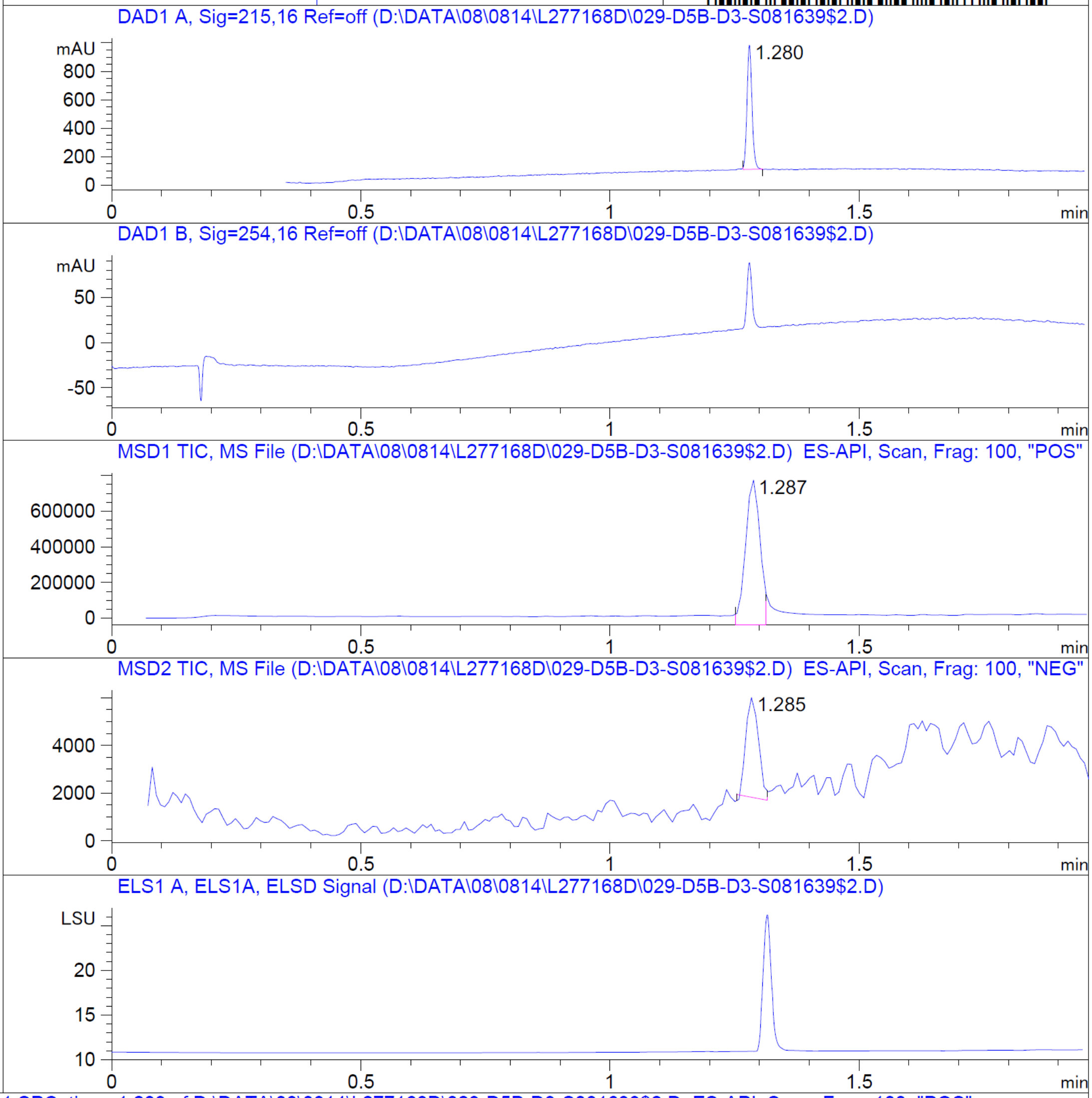


RT 1.287

RT 1.285

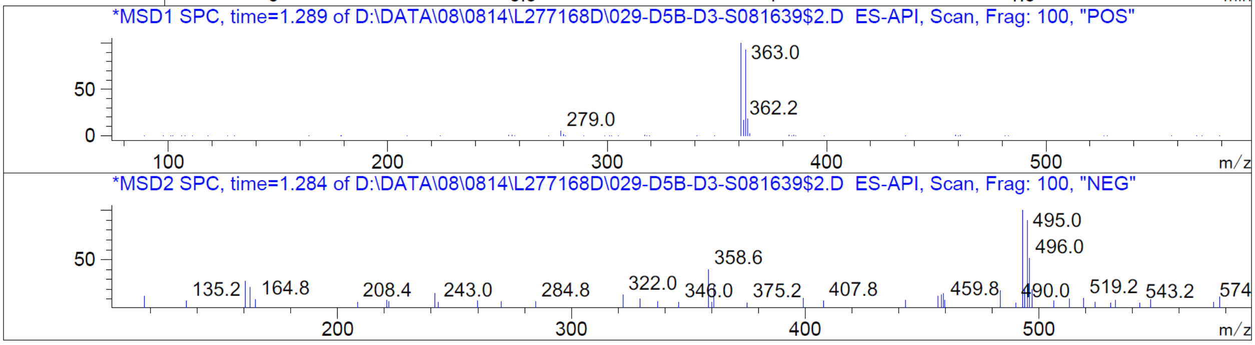


Z3201310989_(0273num3247.2)

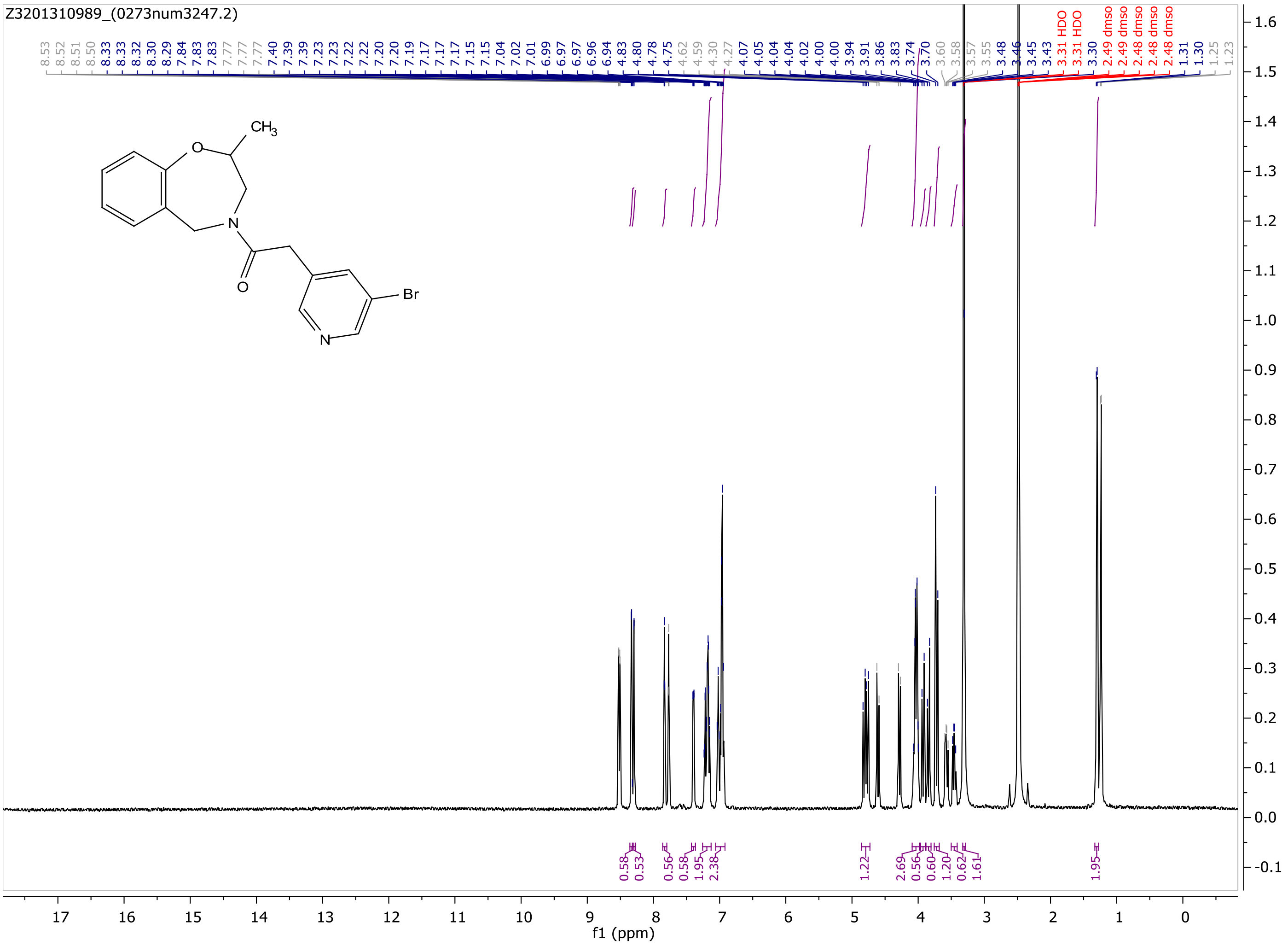
Z3201310989_(0273num3247.2)
CH3
O
N
Br
f1 (ppm)

MaxPeak: 100.00%
Ret_Time: 1.122 min

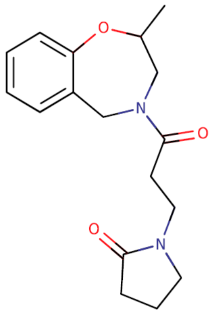


**Mol Wt** 302.37
**Exact Mass** 302.19

| # | Time | Area% |
|---|---|---|
| 1 | 1.122 | 100.00 |

S081643$1

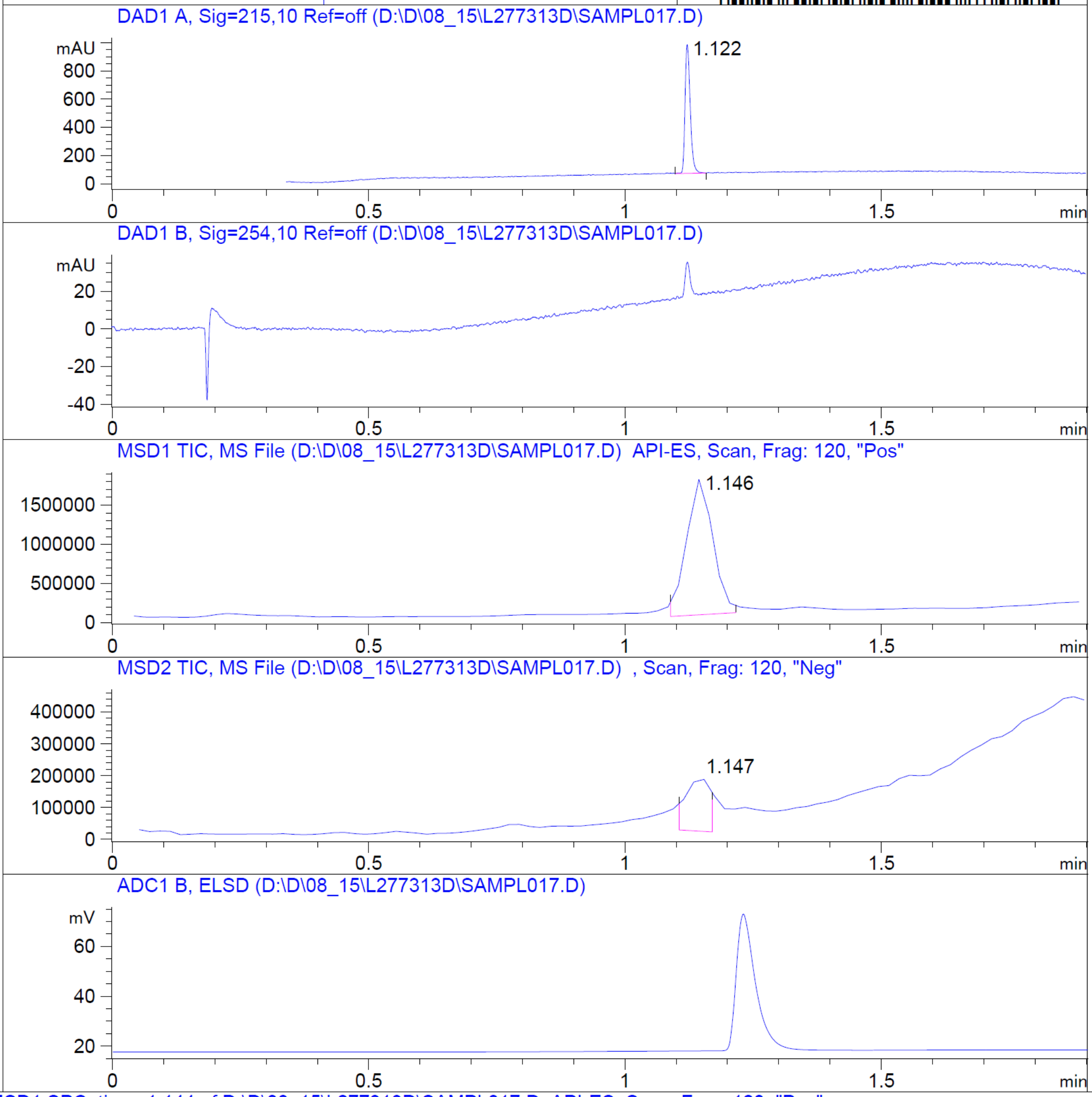


RT 1.146

RT 1.147

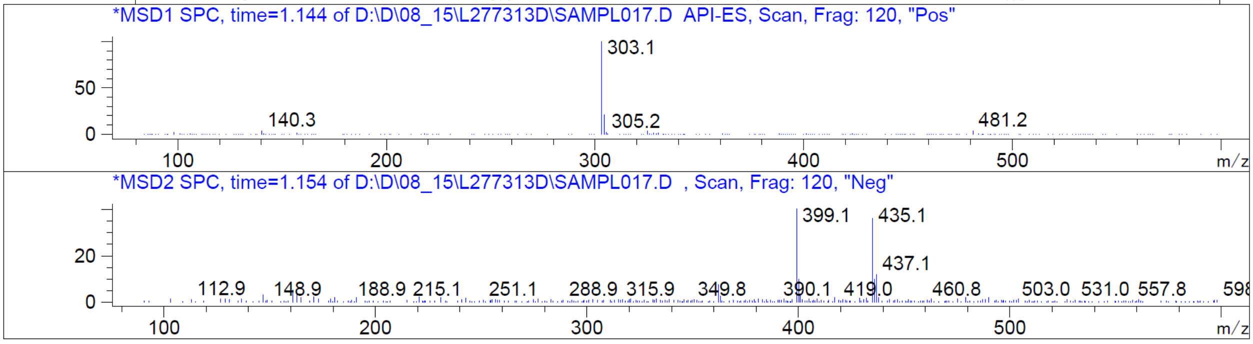


Z2227611595_(0273num3604.2)

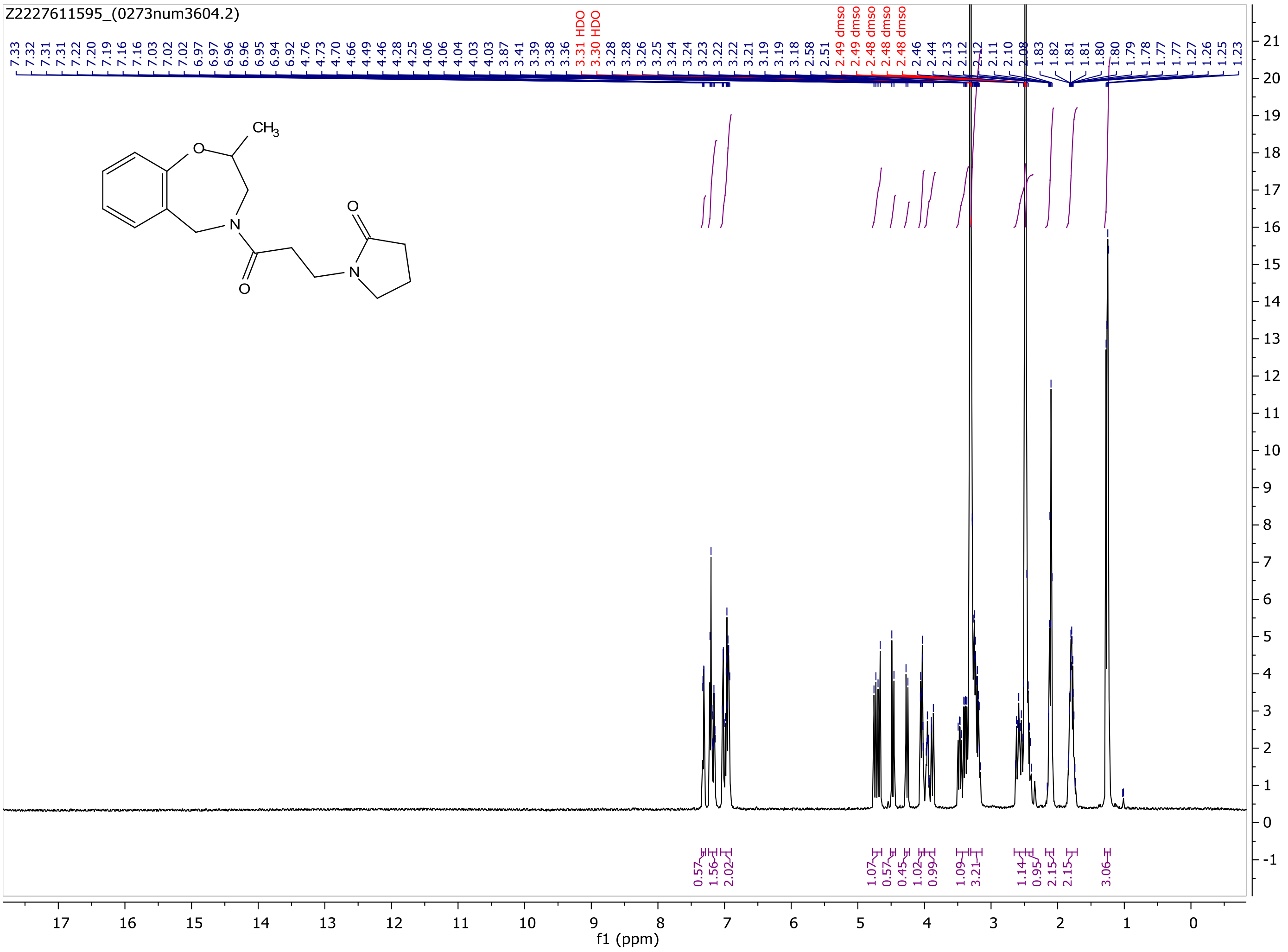
Z2227611595_(0273num3604.2)
7.33 7.32 7.31 7.31 7.22 7.20 7.19 7.16 7.16 7.03 7.02 7.02 6.97 6.97 6.96 6.96 6.95 6.94 6.92 4.76 4.73 4.70 4.66 4.49 4.46 4.28 4.25 4.06 4.06 4.04 4.03 4.03 3.87 3.41 3.39 3.38 3.36 3.31 HDO 3.30 HDO 3.28 3.28 3.26 3.25 3.24 3.24 3.23 3.22 3.22 3.21 3.19 3.19 3.18 2.58 2.51 2.49 dmso 2.49 dmso 2.48 dmso 2.48 dmso 2.48 dmso 2.46 2.44 2.13 2.12 2.12 2.11 2.10 2.08 1.83 1.82 1.81 1.81 1.80 1.80 1.79 1.78 1.77 1.77 1.27 1.26 1.25 1.23
CH3
O
N
0.57 1.56 2.02 1.07 0.57 0.45 1.02 0.99 1.09 3.21 1.14 0.95 2.15 2.15 3.06
f1 (ppm)

MaxPeak:  100.00%
Ret_Time: 1.205 min

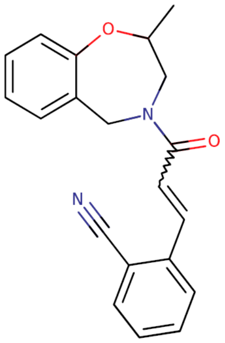


**Mol Wt** 318.37

**Exact Mass** 318.16

| # | Time | Area% |
|---|---|---|
| 1 | 1.205 | 100.00 |

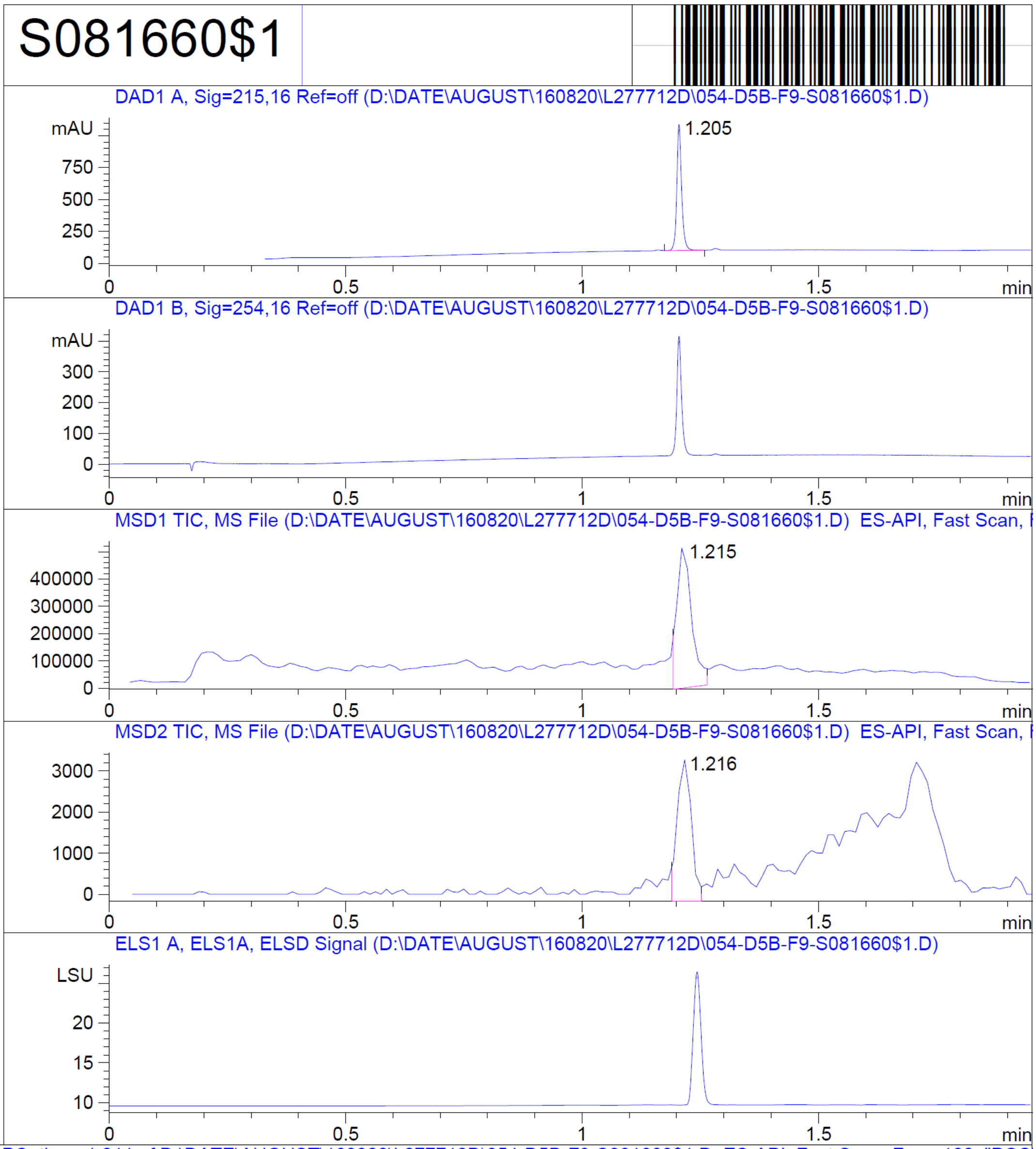


RT 1.215

RT 1.216

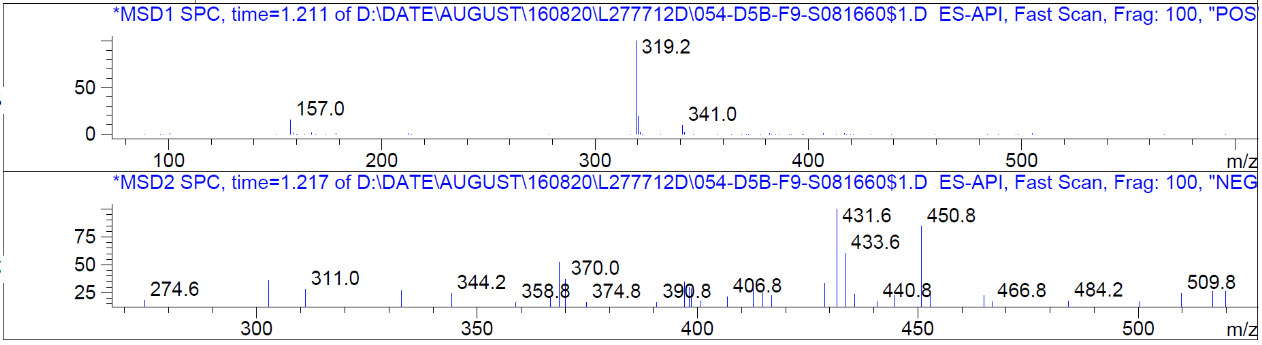


Z2230172883_(0273num3343.2)

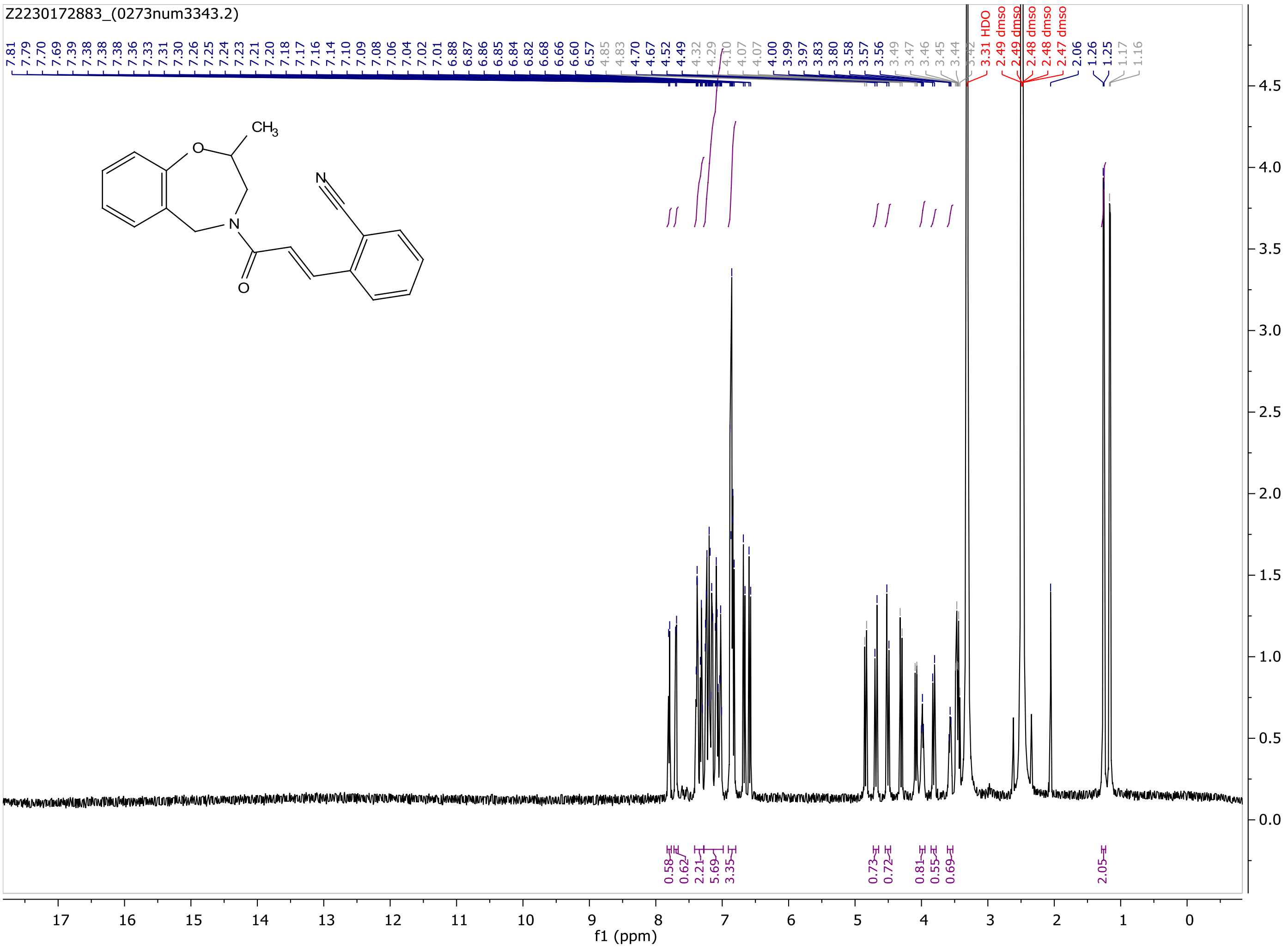
Z2230172883_(0273num3343.2)
f1 (ppm)

MaxPeak: 98.21%
Ret_Time: 1.386 min

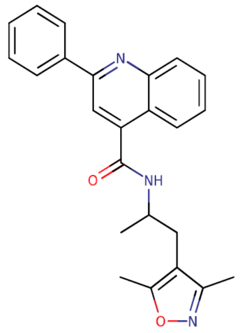

**Mol Wt 385.46**

**Exact Mass 385.21**

| # | Time | Area% |
|---|---|---|
| 1 | 1.386 | 98.21 |
| 2 | 1.465 | 1.79 |

S117856$2

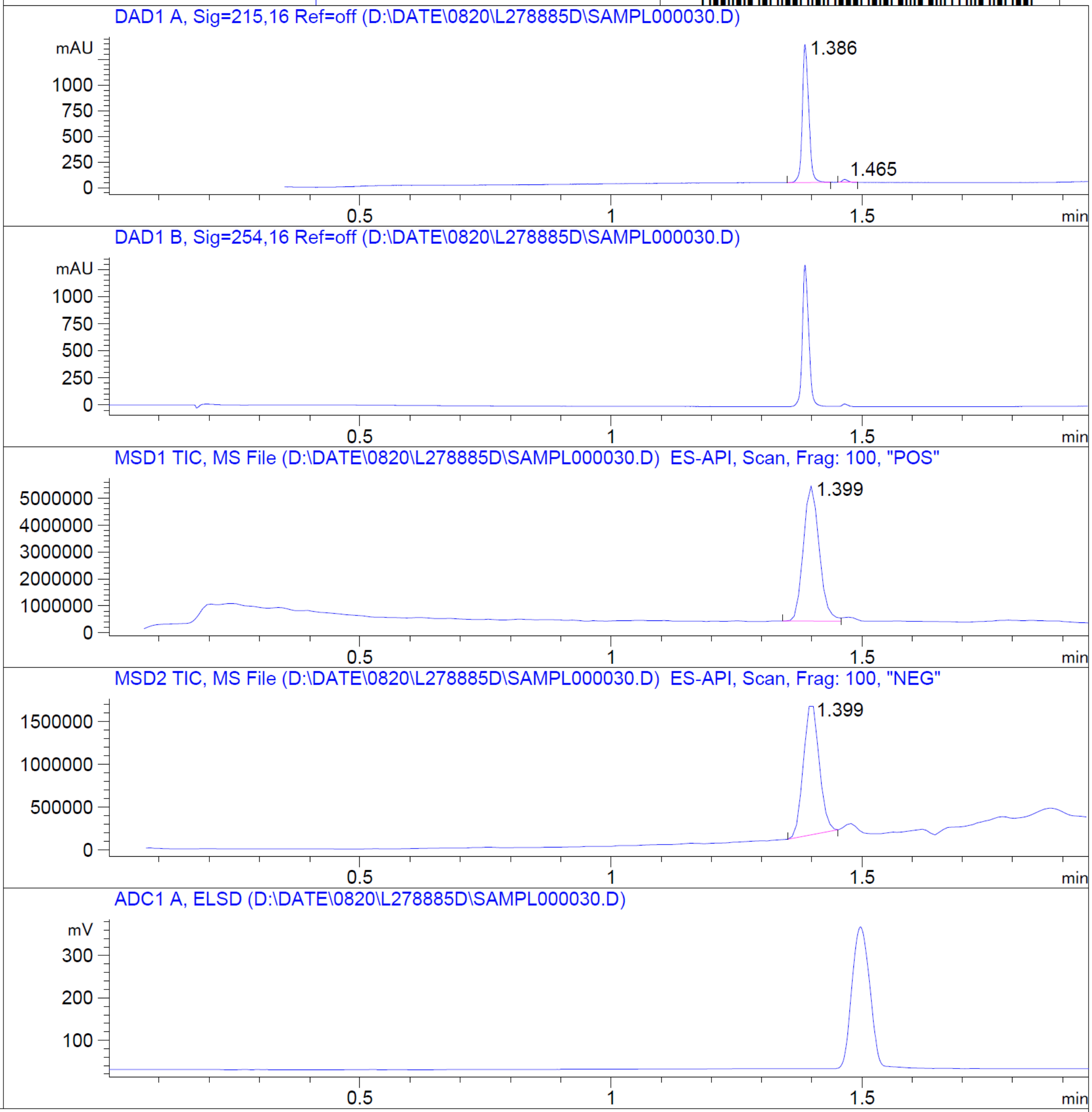


RT 1.399

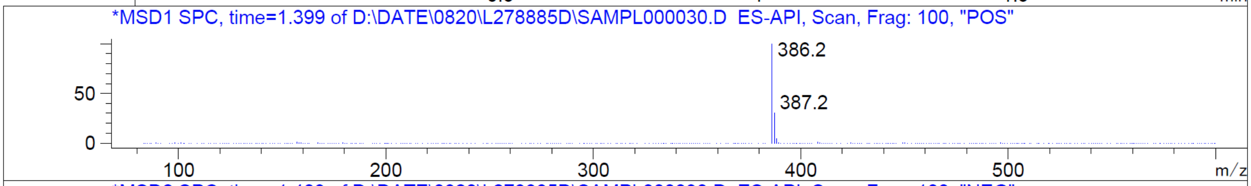


RT 1.399

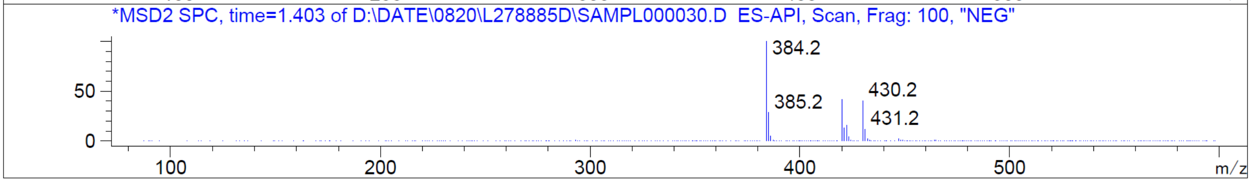


Z1239545406_(3312num225.1)

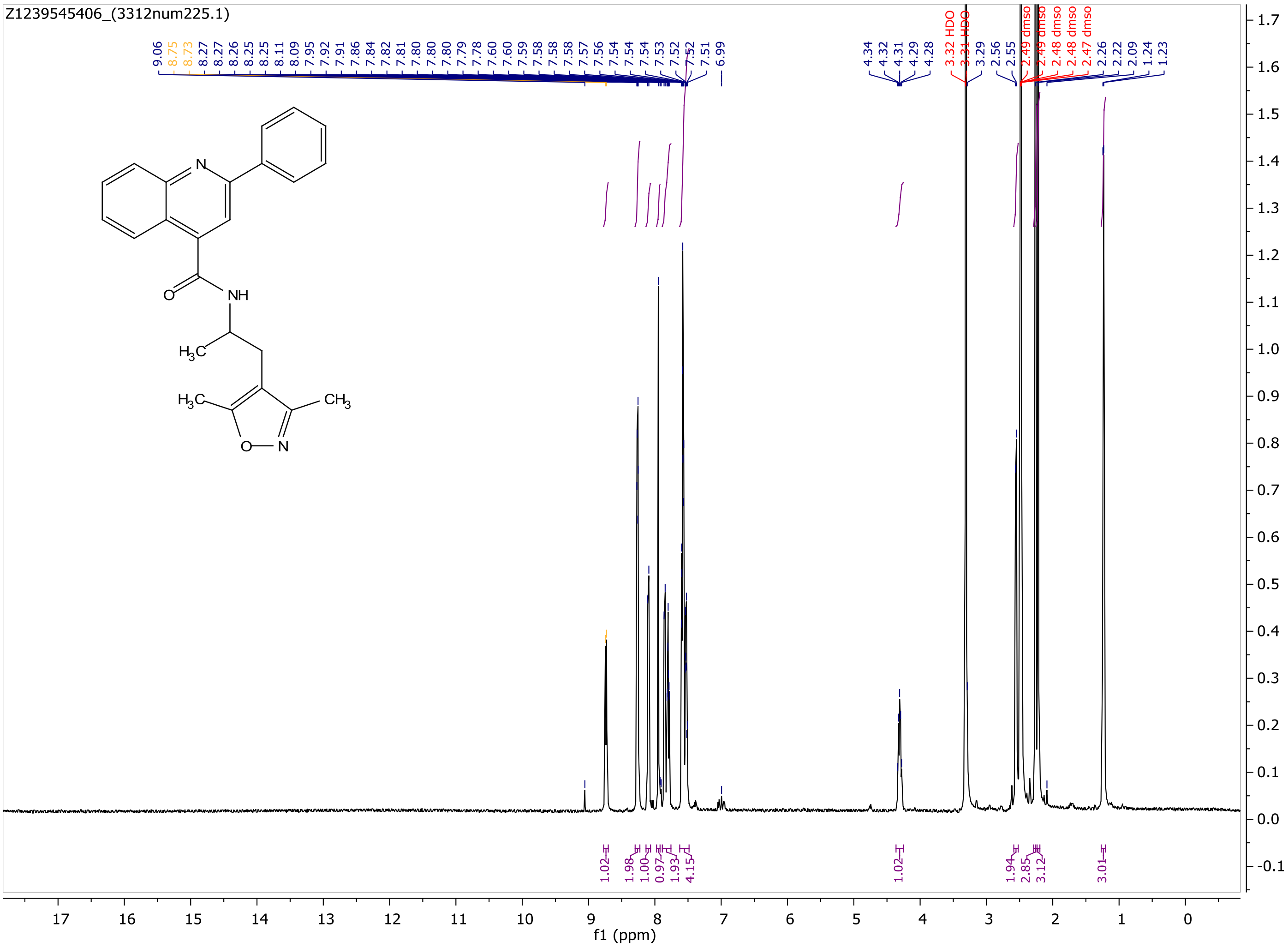
Z1239545406_(3312num225.1)
9.06 8.75 8.73 8.27 8.27 8.26 8.25 8.25 8.11 8.09 7.95 7.92 7.91 7.86 7.84 7.82 7.81 7.80 7.80 7.80 7.79 7.78 7.60 7.60 7.59 7.58 7.58 7.58 7.57 7.56 7.54 7.54 7.54 7.53 7.52 7.52 7.51 6.99
4.34 4.32 4.31 4.29 4.28
3.32 HDO
3.31 HDO
3.29 2.56 2.55
2.49 dmso
2.49 dmso
2.48 dmso
2.48 dmso
2.47 dmso
2.26 2.22 2.09 1.24 1.23
N
NH
O
$H_3C$
$H_3C$
$CH_3$
O
N
1.02 1.98 1.00 0.97 1.93 4.15 1.02 1.94 2.85 3.12 3.01
f1 (ppm)

MaxPeak: 92.07%
Ret_Time: 0.916 min

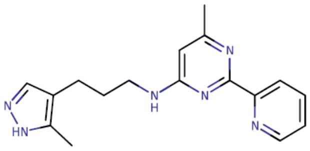


**Mol Wt** 308.38
**Exact Mass** 308.2

| # | Time | Area% |
|---|---|---|
| 1 | 0.875 | 7.93 |
| 2 | 0.916 | 92.07 |

S117850$4

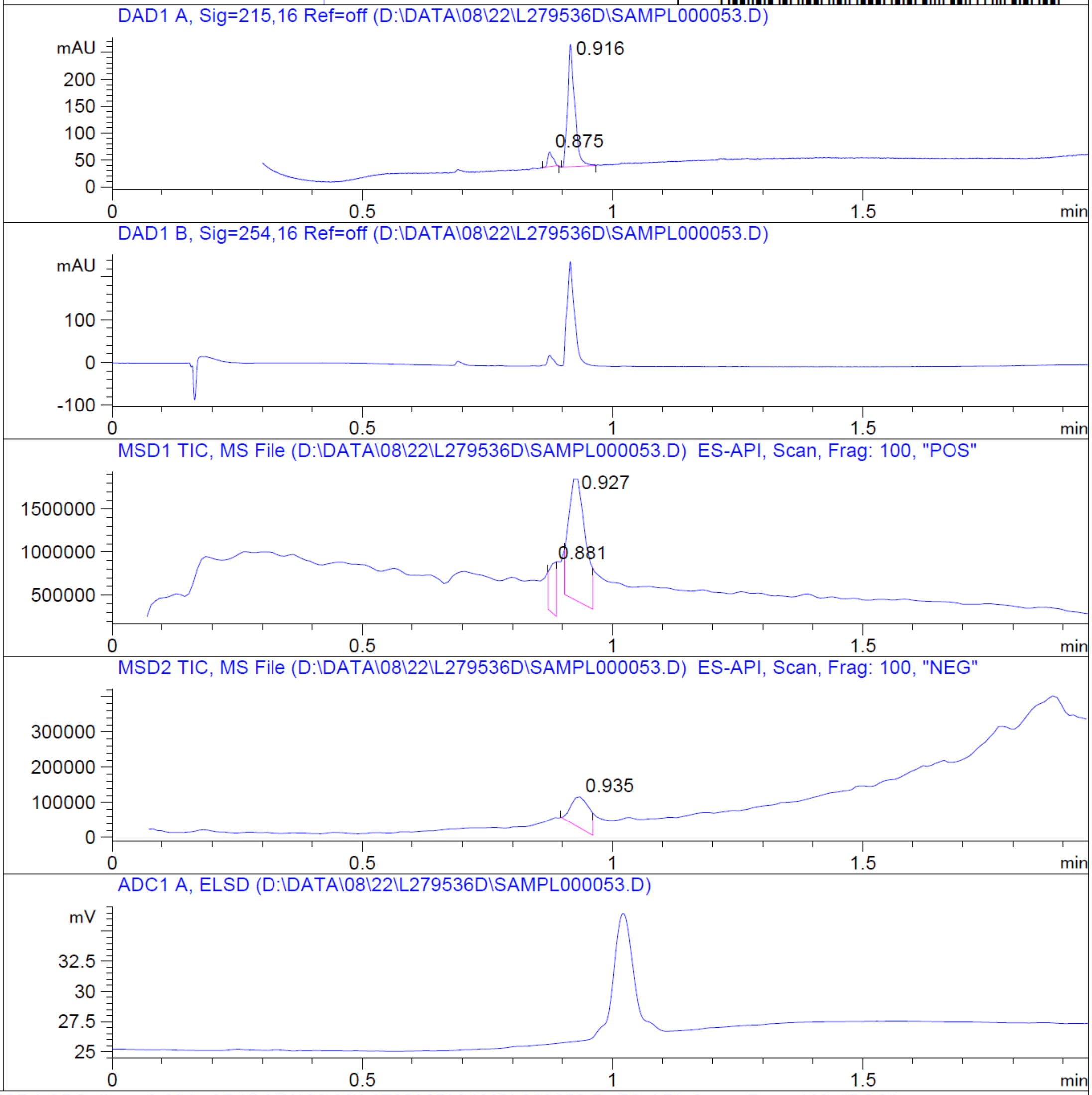


RT 0.881

RT 0.927

RT 0.935

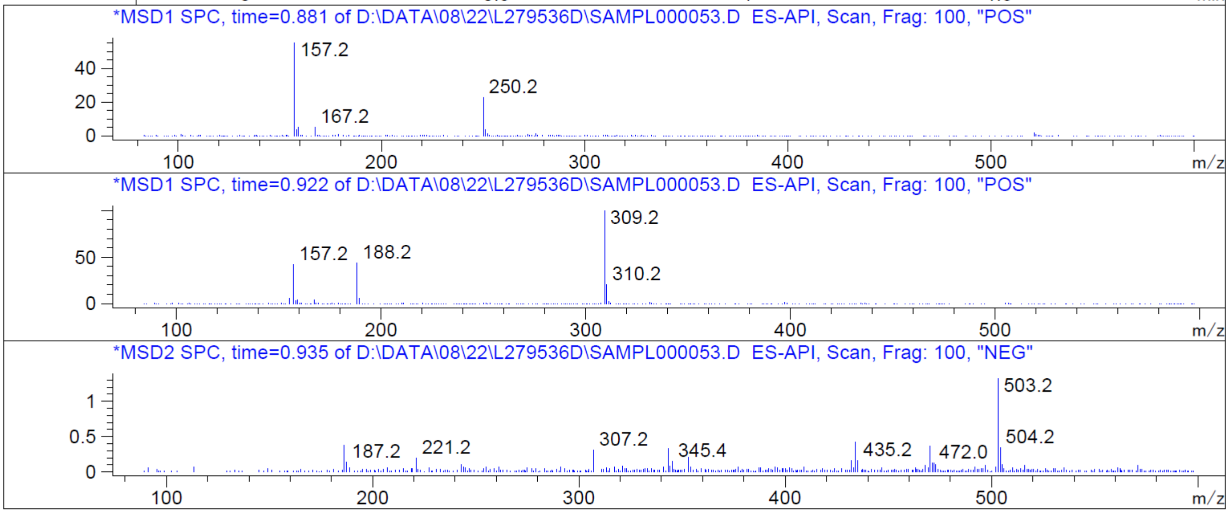


Z1168173619_(3312num339)

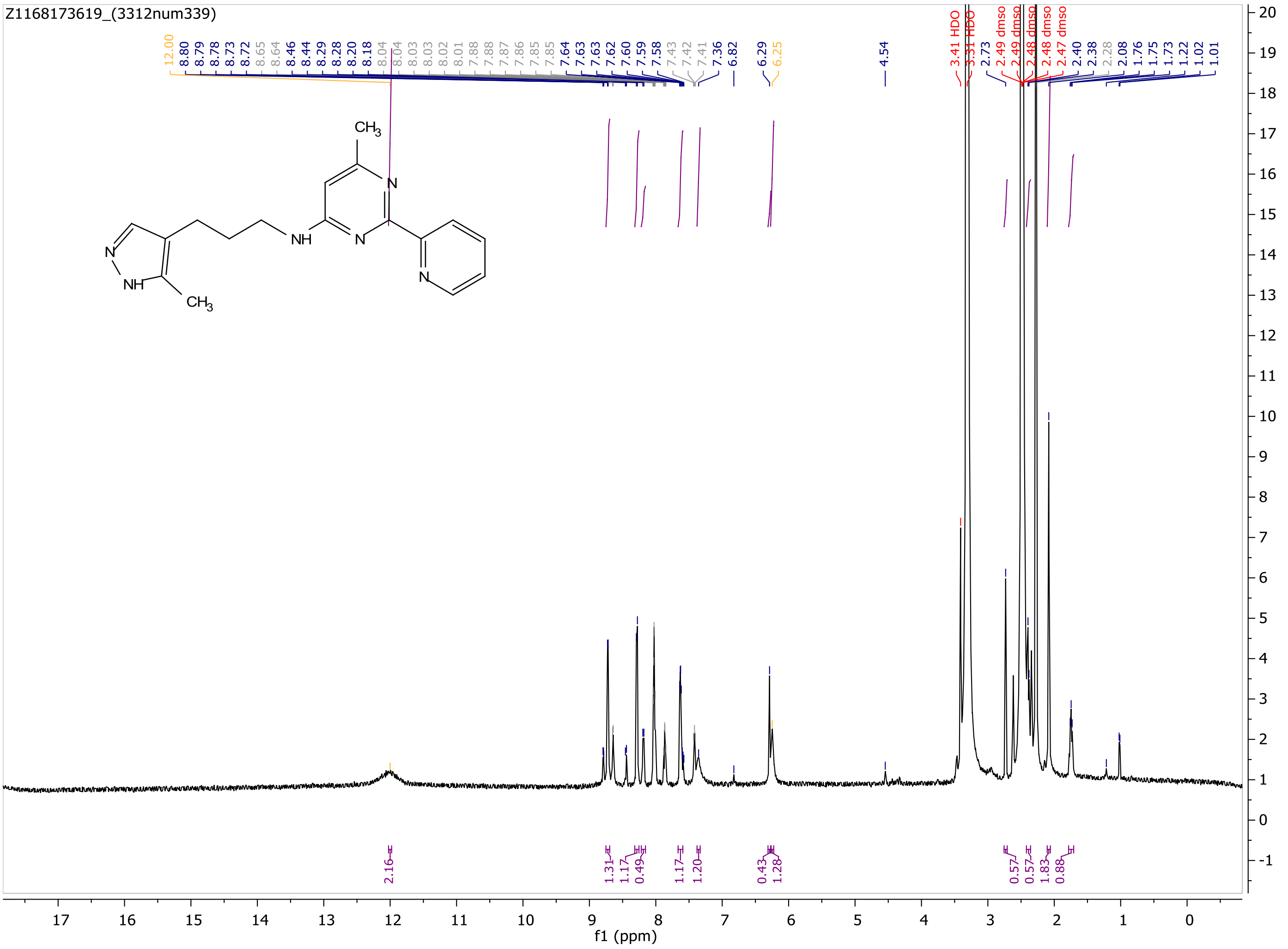

Z1168173619_(3312num339)
12.00 8.80 8.79 8.78 8.73 8.72 8.65 8.64 8.46 8.44 8.29 8.28 8.20 8.18 8.04 8.04 8.03 8.03 8.02 8.01 7.88 7.88 7.87 7.86 7.85 7.85 7.64 7.63 7.62 7.60 7.59 7.58 7.43 7.42 7.41 7.36 6.82 6.29 6.25 4.54
3.41 HDO
3.31 HDO
2.73
2.49 dmso
2.49 dmso
2.48 dmso
2.48 dmso
2.47 dmso
2.40 2.38 2.28 2.08 1.76 1.75 1.73 1.22 1.02 1.01
CH3
N
NH
N
N
N
NH
CH3
2.16 1.31 1.17 0.49 1.17 1.20 0.43 1.28 0.57 0.57 1.83 0.88
f1 (ppm)

MaxPeak: 100.00%
Ret_Time: 1.247 min

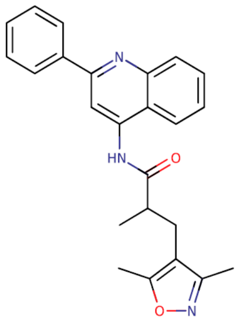

**Mol Wt** 385.46
**Exact Mass** 385.21

| # | Time | Area% |
|---|---|---|
| 1 | 1.247 | 100.00 |

# S117851$3

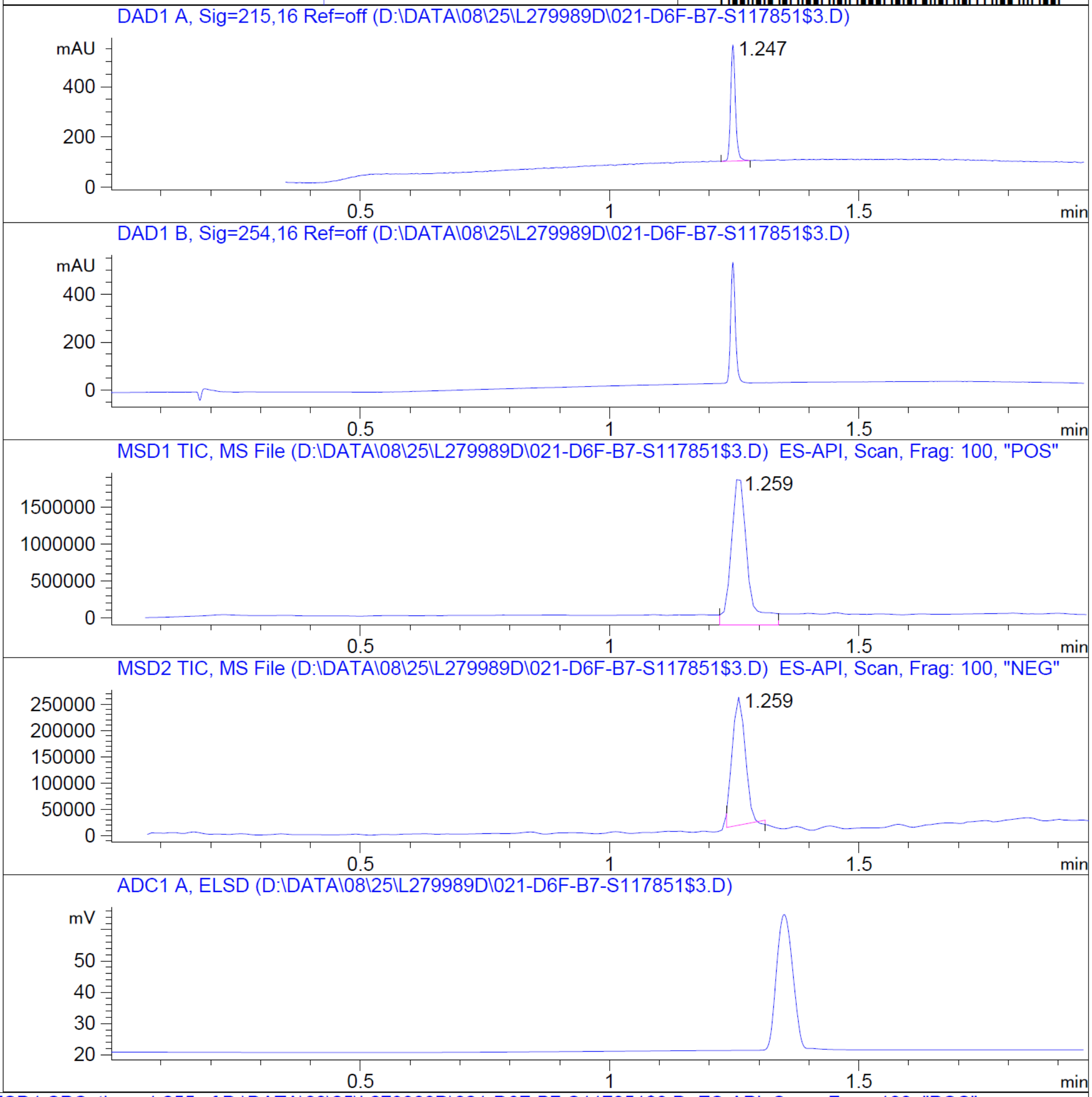


RT 1.259

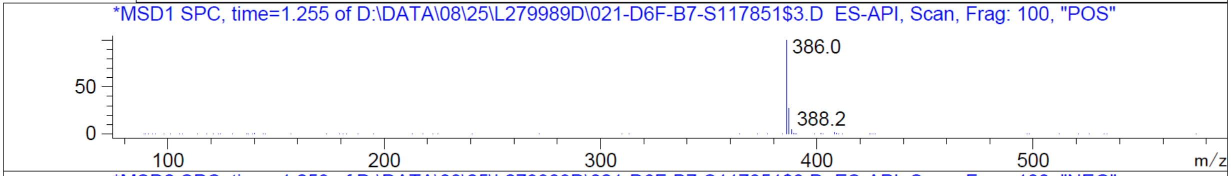


RT 1.259

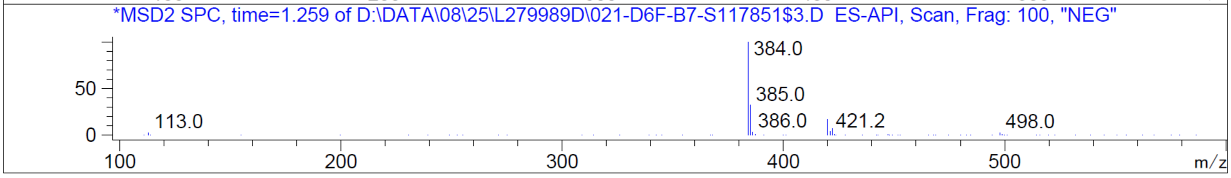


Z4570447323_(3312num215.1)

Z4570447323_(3312num215.1)

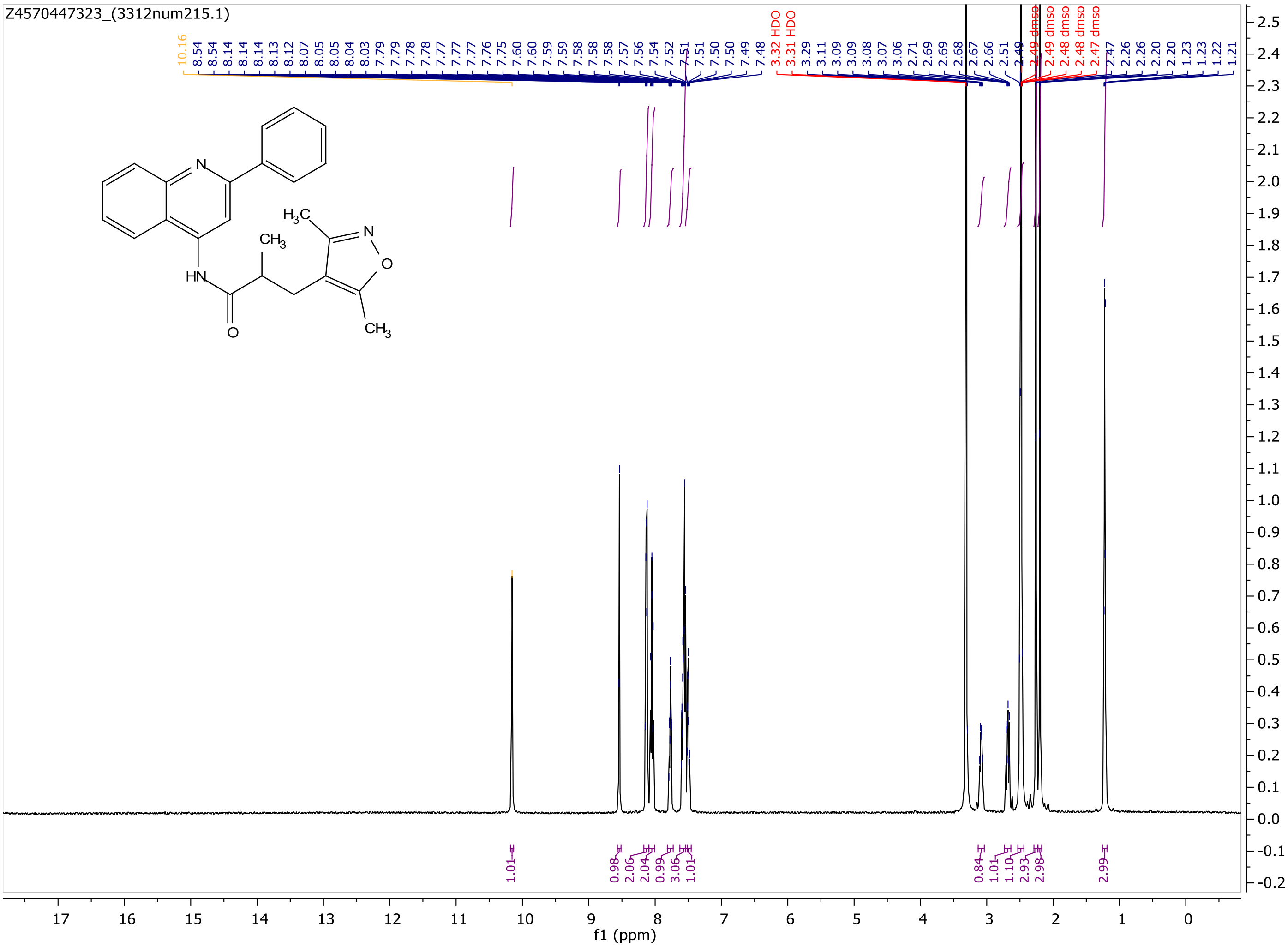

MaxPeak: 100.00%
Ret_Time: 1.035 min

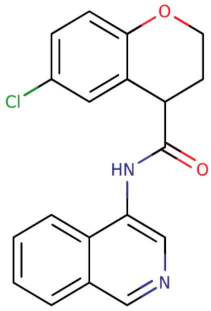


**Mol Wt** 338.79
**Exact Mass** 338.1

| # | Time | Area% |
|---|---|---|
| 1 | 1.035 | 100.00 |

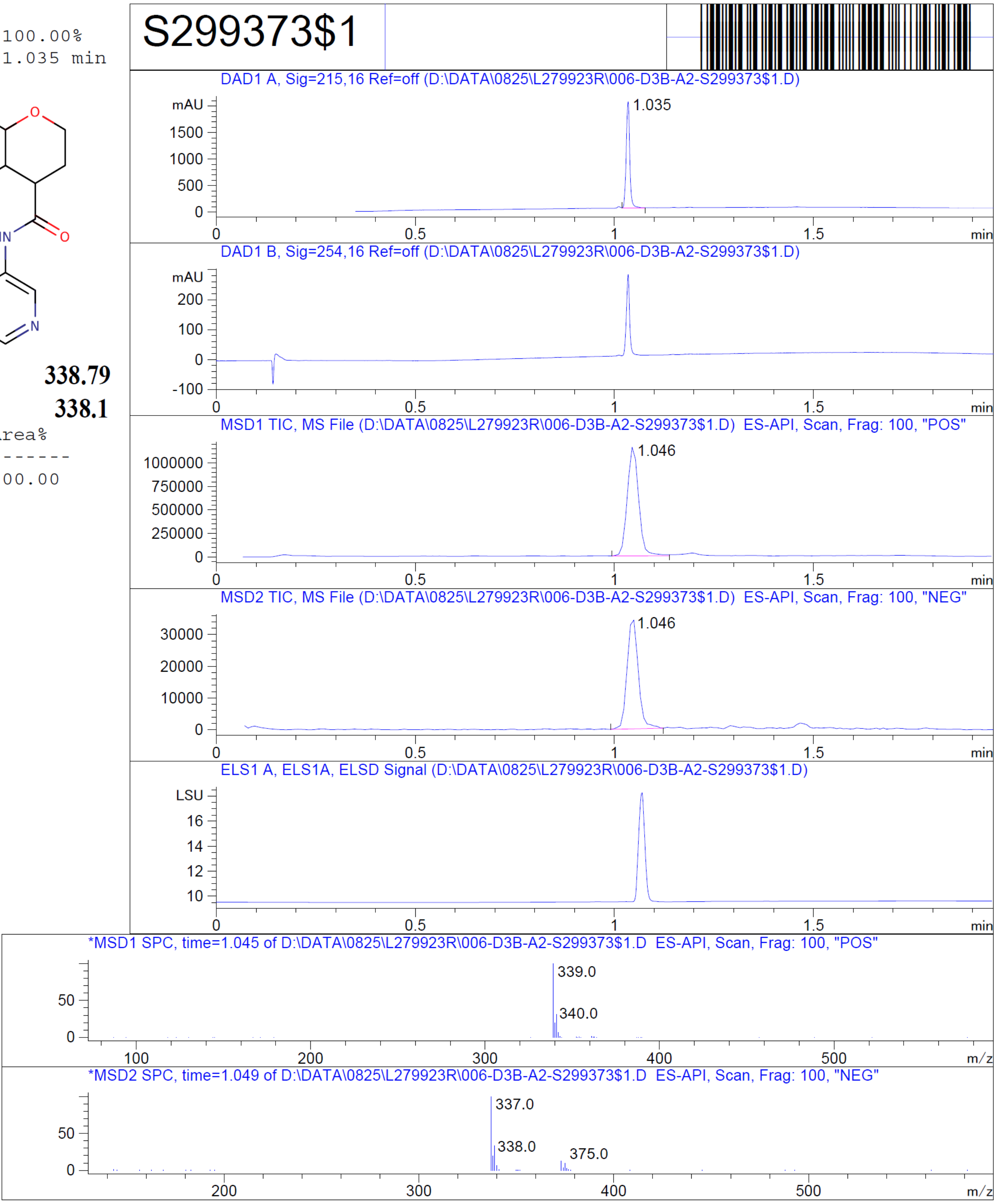


RT 1.046

RT 1.046

Z1675865540_(EN300-27120704)

Z1675865540_(EN300-27120704)

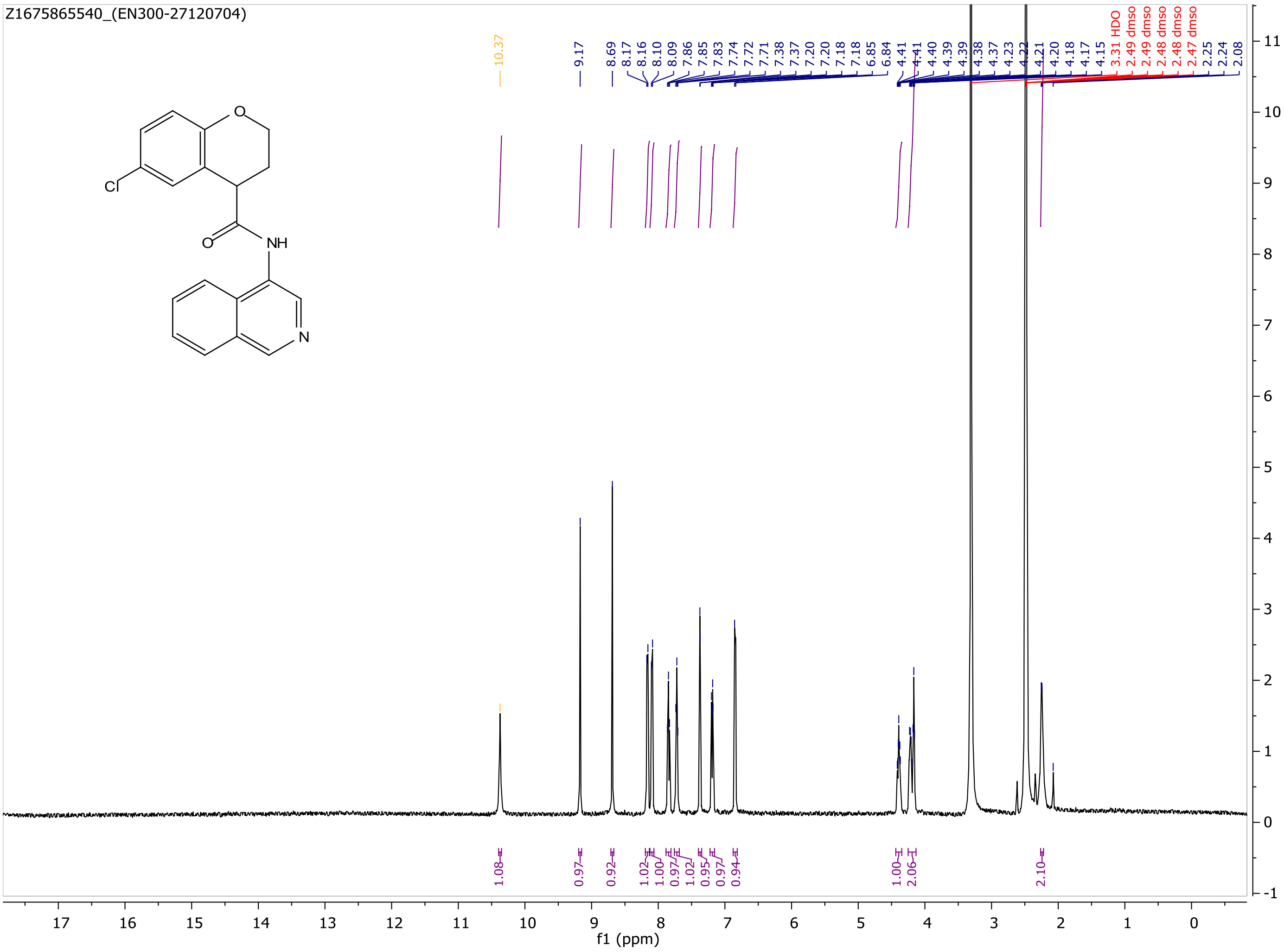

MaxPeak:  100.00%
Ret_Time: 1.012 min

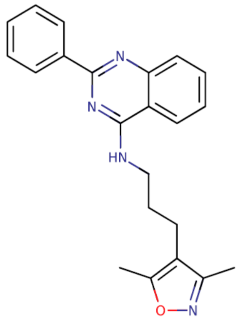

**Mol Wt** 358.44

**Exact Mass** 358.21

| # | Time | Area% |
| --- | --- | --- |
| 1 | 1.012 | 100.00 |

# S231075$A

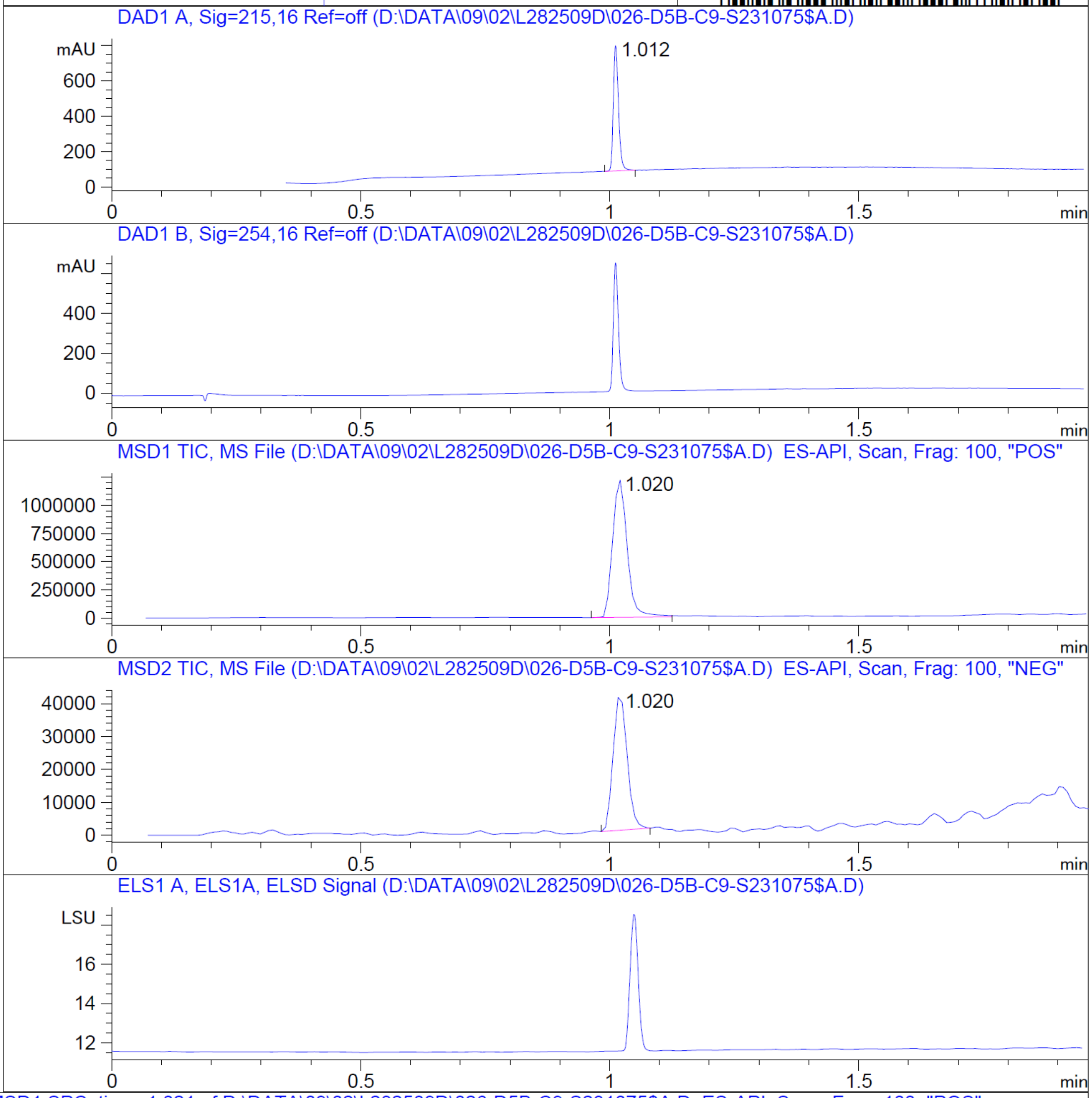


RT 1.020

RT 1.020

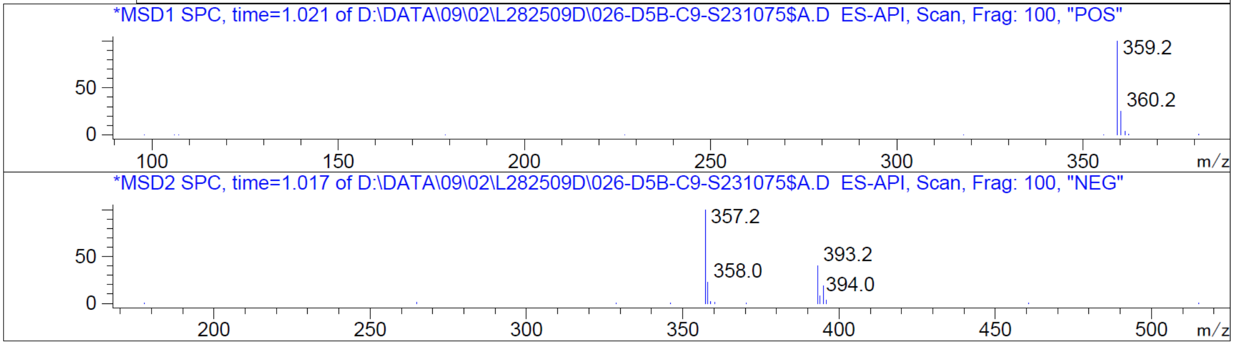


Z4600752345

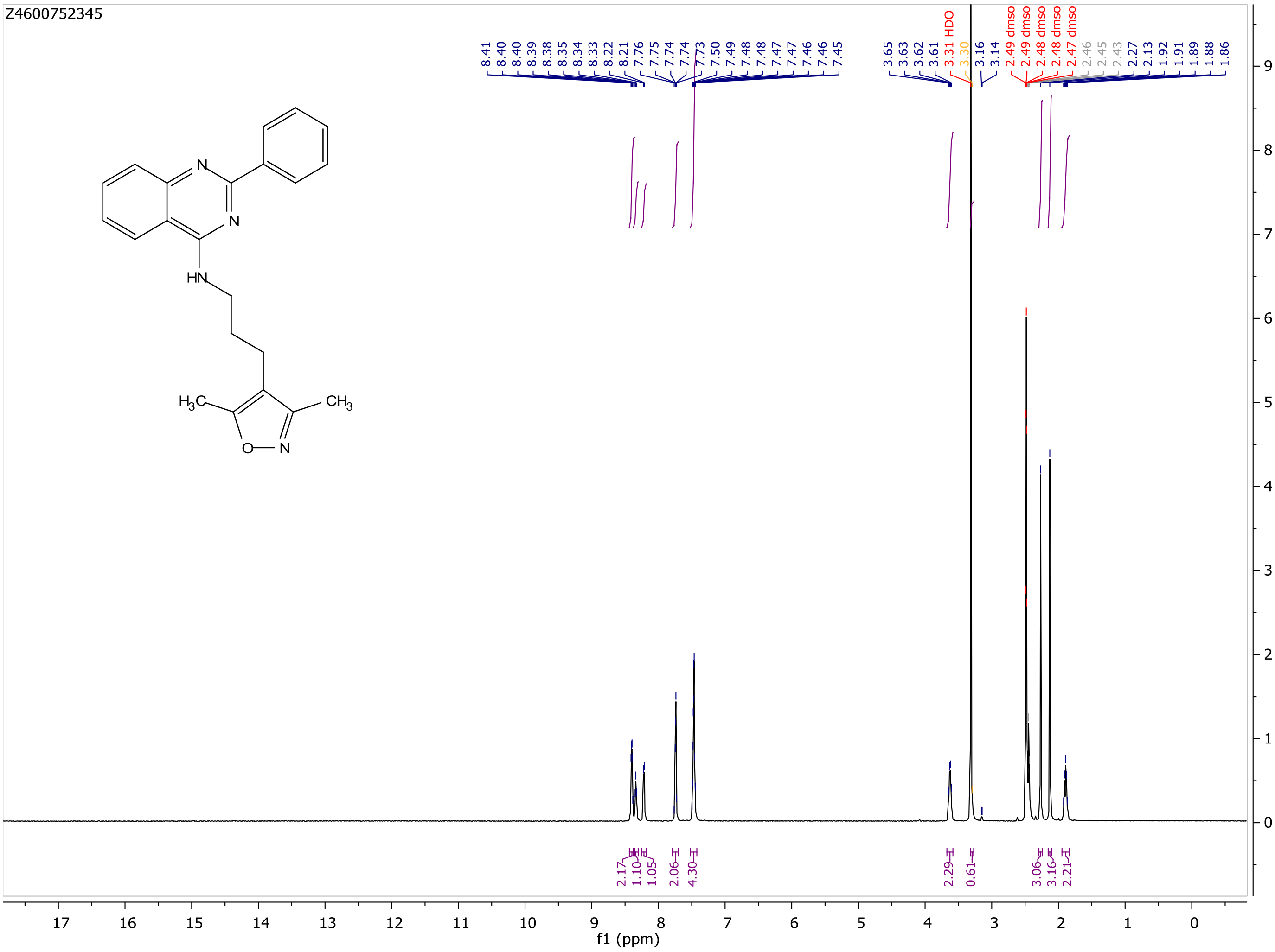
Z4600752345
8.41 8.40 8.40 8.39 8.38 8.35 8.34 8.33 8.22 8.21 7.76 7.75 7.74 7.74 7.73 7.50 7.49 7.48 7.48 7.47 7.47 7.46 7.46 7.45
3.65 3.63 3.62 3.61 3.31 HDO 3.30 3.16 3.14 2.49 dmso 2.49 dmso 2.48 dmso 2.48 dmso 2.47 dmso 2.46 2.45 2.43 2.27 2.13 1.92 1.91 1.89 1.88 1.86
2.17 1.10 1.05 2.06 4.30 2.29 0.61 3.06 3.16 2.21
f1 (ppm)

MaxPeak: 100.00%
Ret_Time: 0.788 min

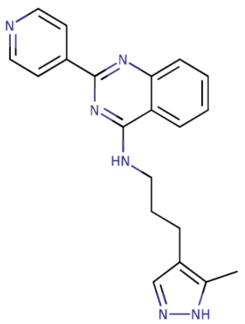

**Mol Wt** 344.41
**Exact Mass** 344.2

| # | Time | Area% |
|---|---|---|
| 1 | 0.788 | 100.00 |

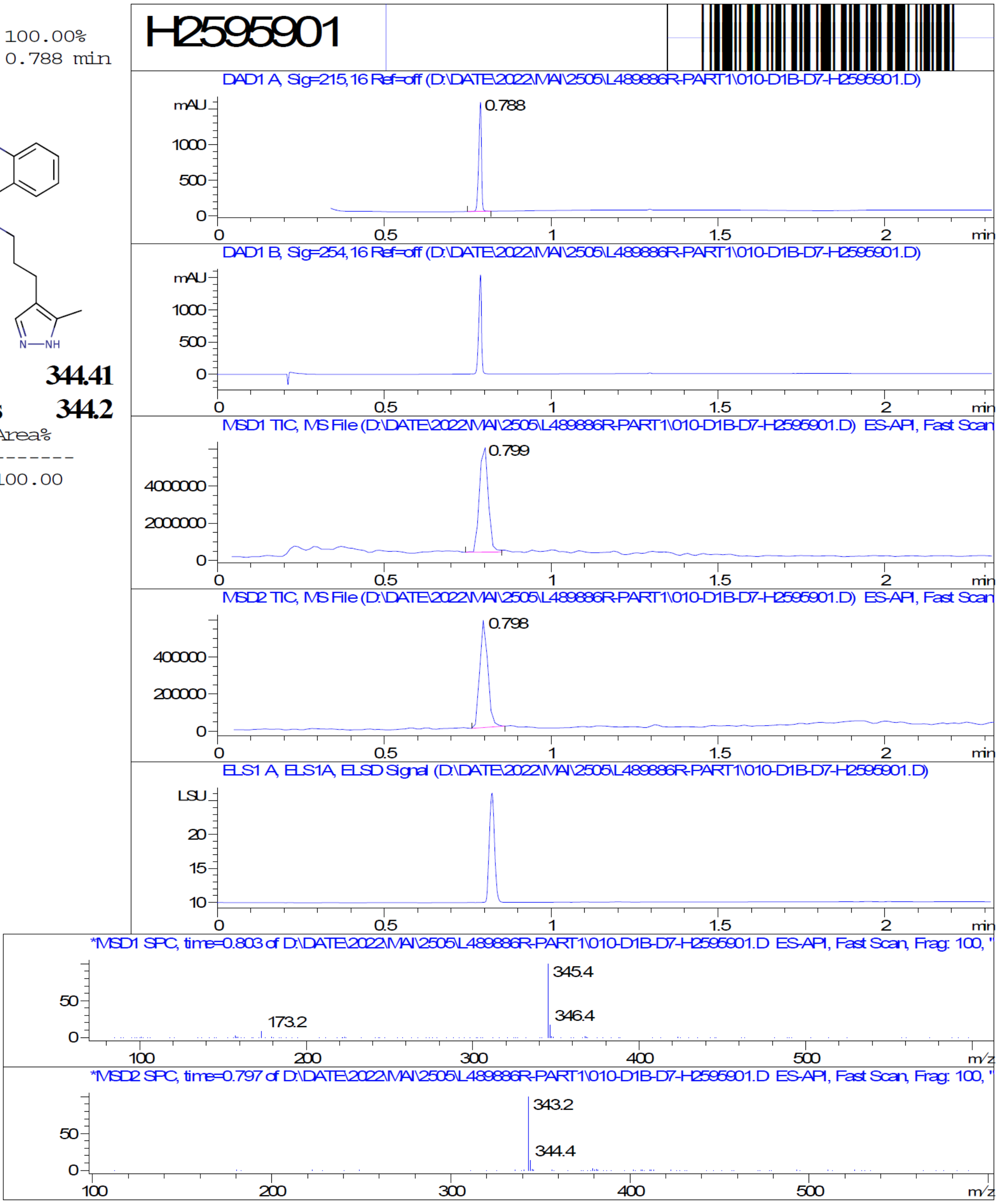


Z285889704

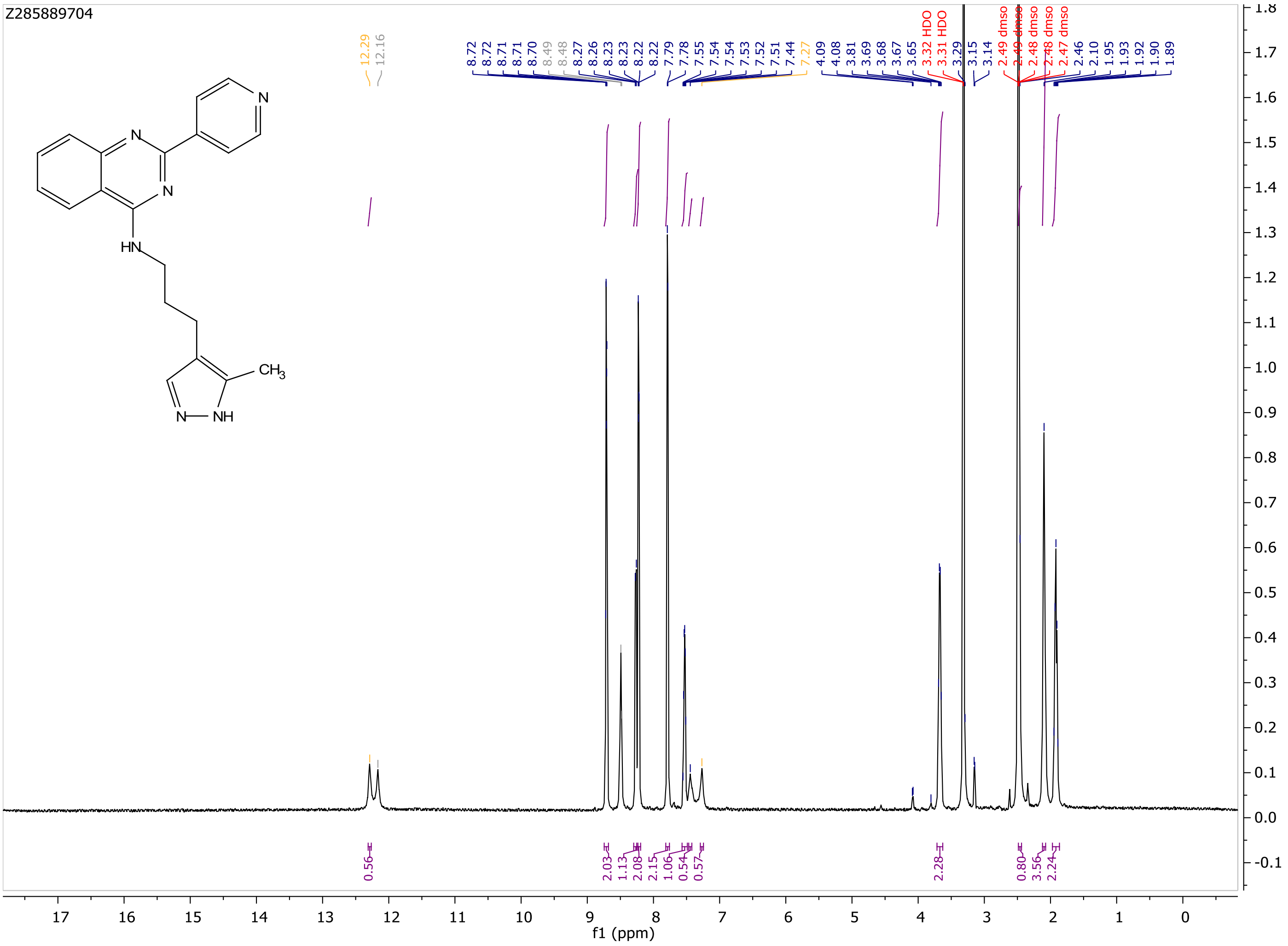
Z285889704
12.29
12.16
8.72
8.72
8.71
8.71
8.70
8.49
8.48
8.27
8.26
8.23
8.23
8.22
8.22
7.79
7.78
7.55
7.54
7.54
7.53
7.52
7.51
7.44
7.27
4.09
4.08
3.81
3.69
3.68
3.67
3.65
3.32 HDO
3.31 HDO
3.29
3.15
3.14
2.49 dmso
2.49 dmso
2.48 dmso
2.48 dmso
2.47 dmso
2.46
2.10
1.95
1.93
1.92
1.90
1.89
0.56
2.03
1.13
2.08
2.15
1.06
0.54
0.57
2.28
0.80
3.56
2.24
f1 (ppm)

MaxPeak: 100.00%
Ret_Time: 1.274 min

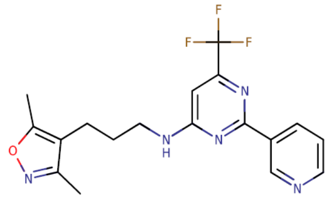


**Mol Wt** 377.36

**Exact Mass** 377.17

| # | Time | Area% |
|---|---|---|
| 1 | 1.274 | 100.00 |

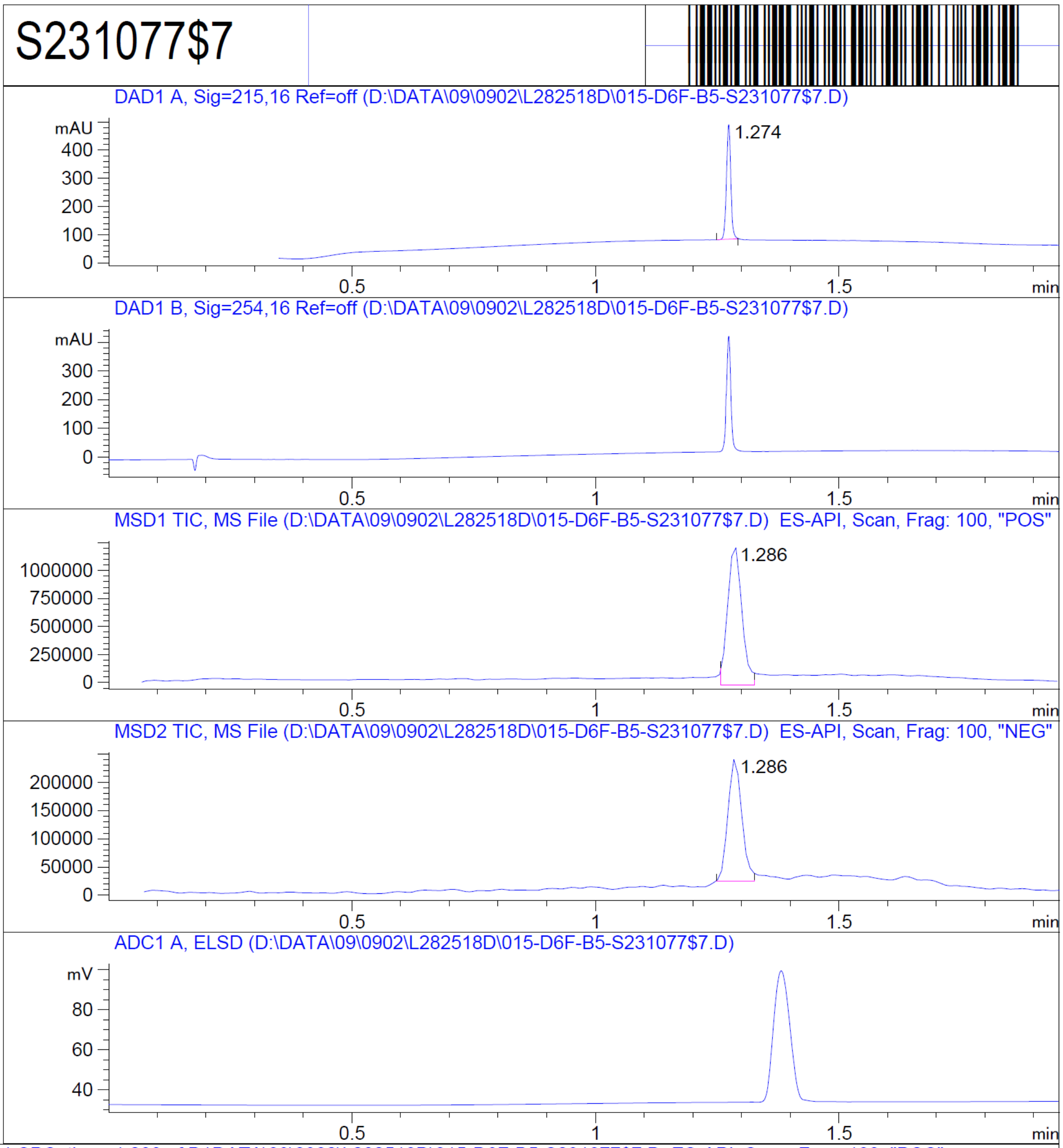


RT 1.286

RT 1.286

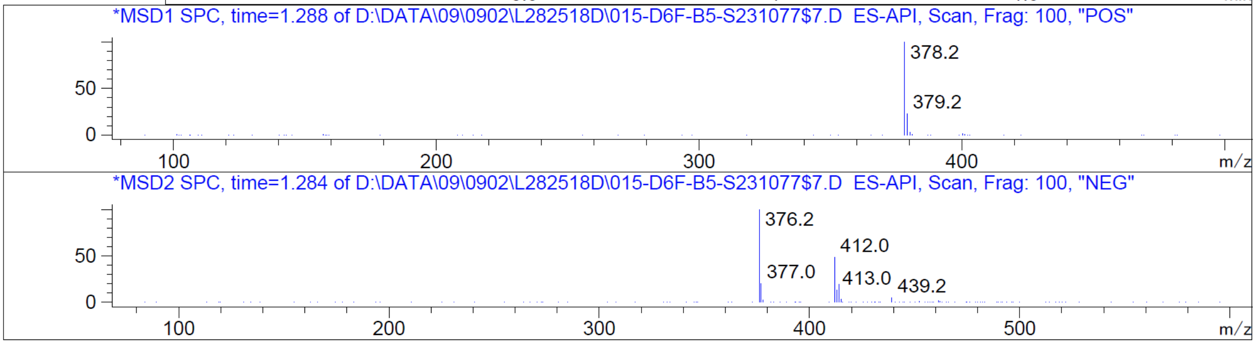


Z1378402602

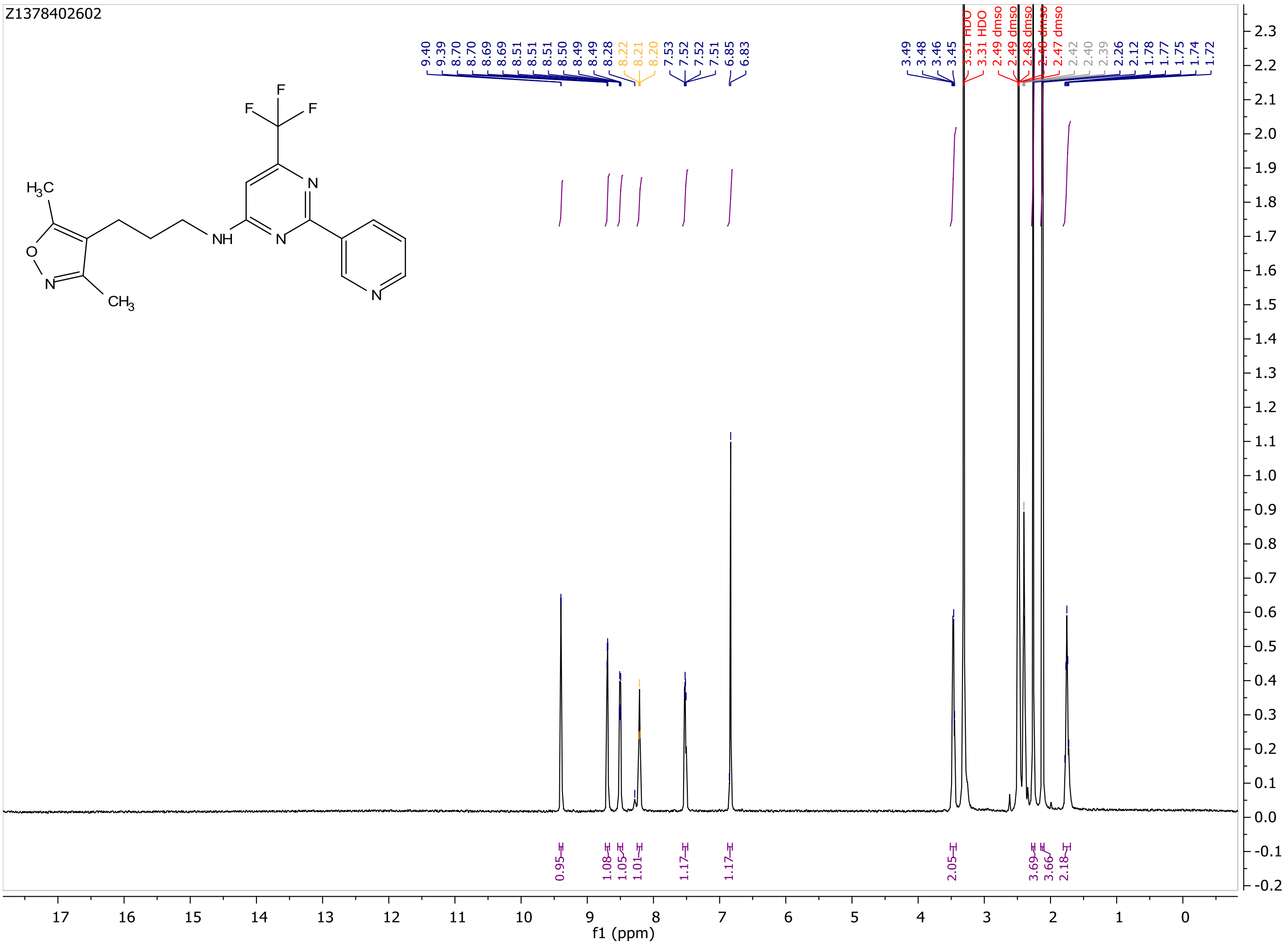
Z1378402602
9.40
9.39
8.70
8.69
8.69
8.51
8.51
8.51
8.50
8.49
8.49
8.28
8.22
8.21
8.20
7.53
7.52
7.52
7.51
6.85
6.83
3.49
3.48
3.46
3.45
3.31 HDO
3.31 HDO
2.49 dmso
2.49 dmso
2.48 dmso
2.48 dmso
2.47 dmso
2.42
2.40
2.39
2.26
2.12
1.78
1.77
1.75
1.74
1.72
0.95
1.08
1.05
1.01
1.17
1.17
2.05
3.69
3.66
2.18
f1 (ppm)

MaxPeak: 100.00%
Ret_Time: 0.863 min

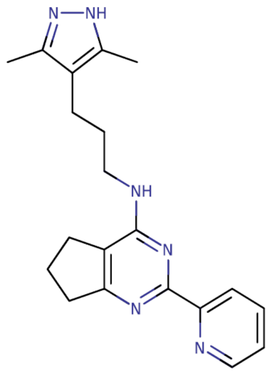

**Mol Wt 348.44**

**Exact Mass 348.24**

| # | Time | Area% |
|---|---|---|
| 1 | 0.863 | 100.00 |

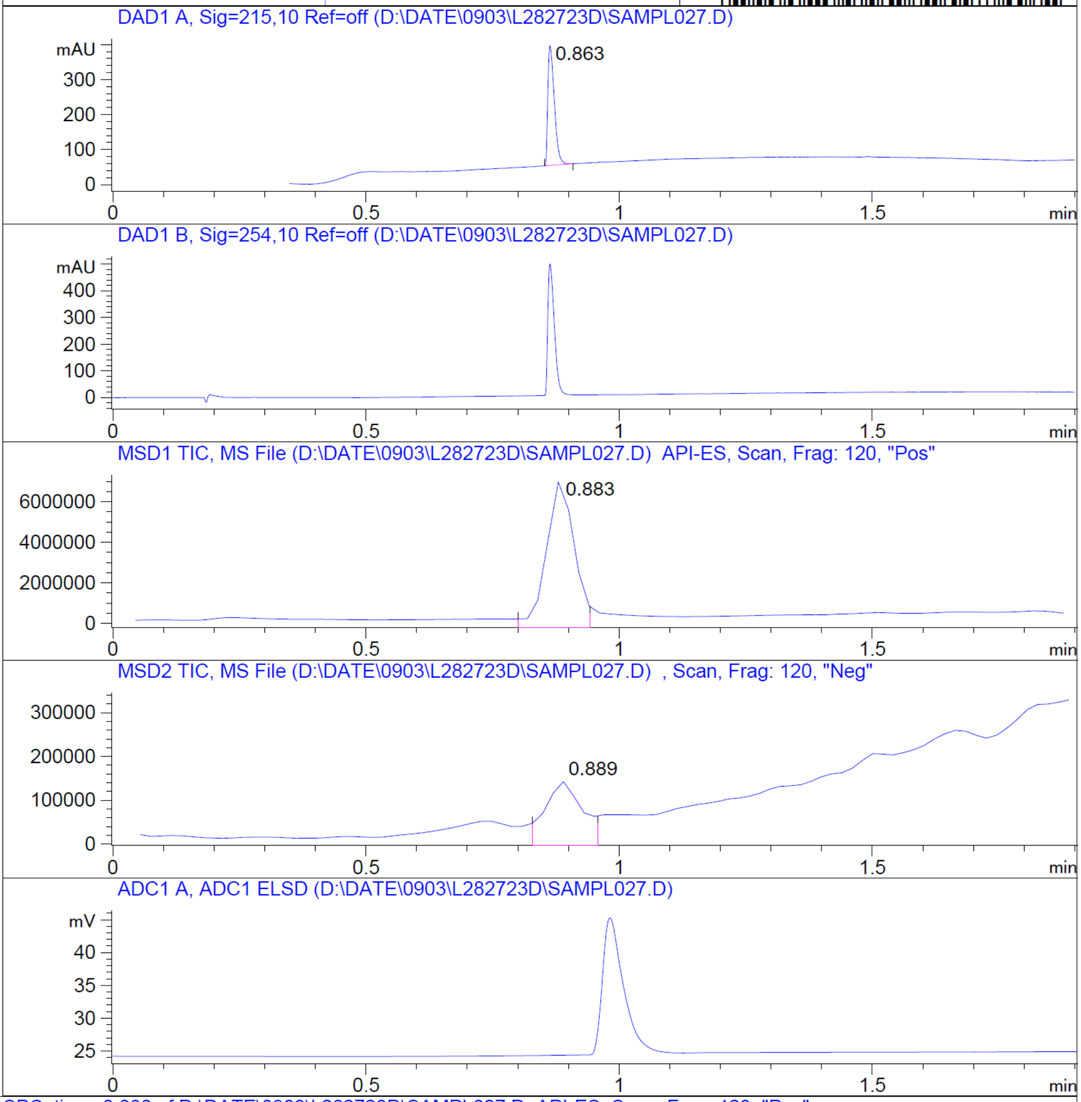


RT 0.883

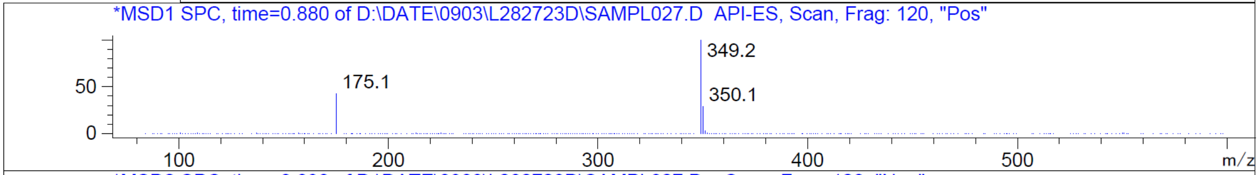


RT 0.889

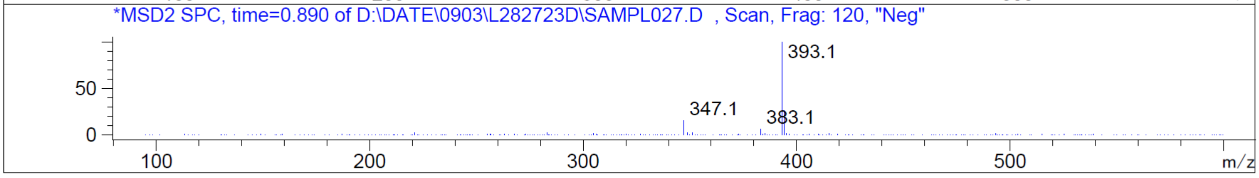


Z4576452292

Z4576452292

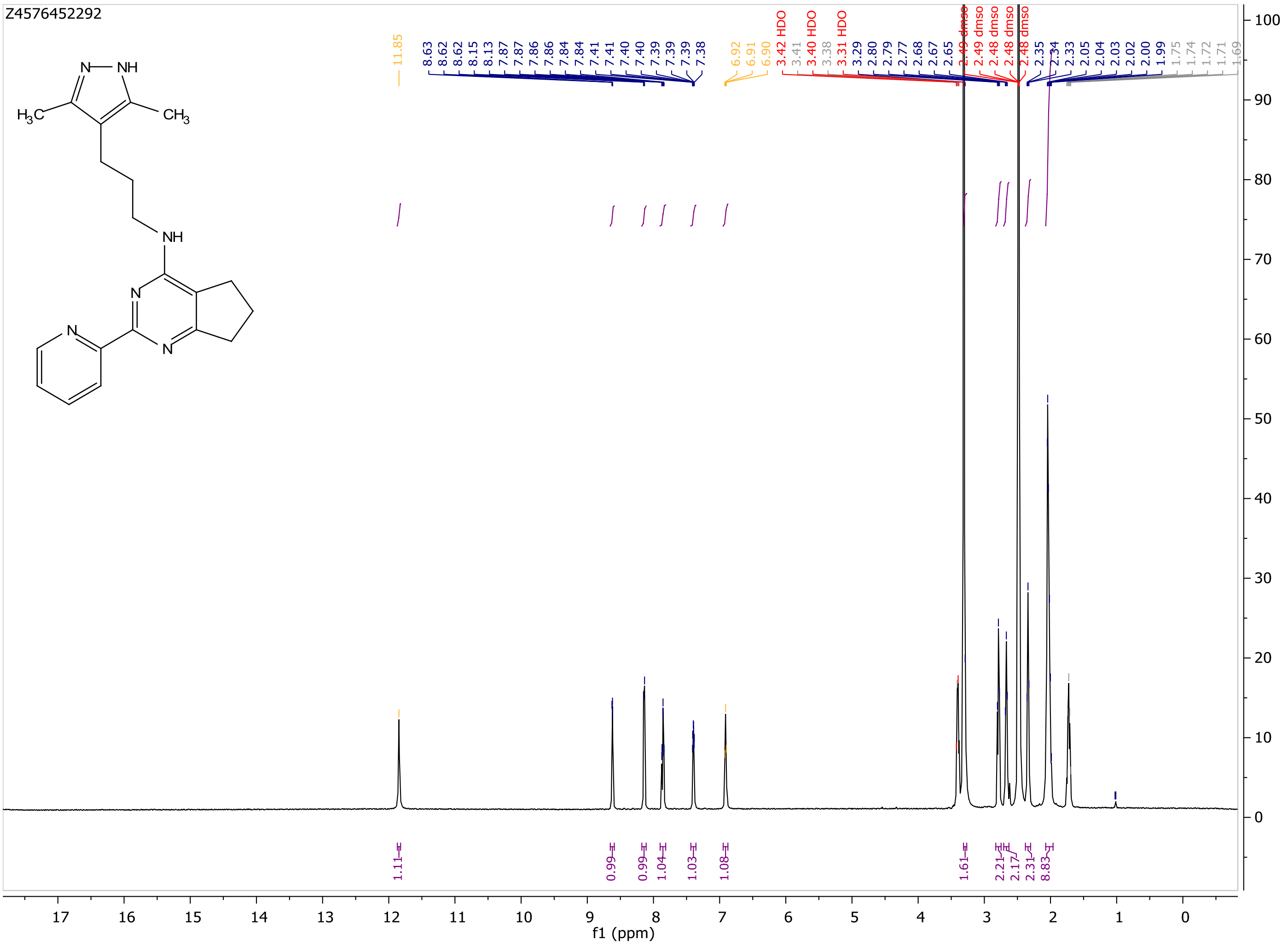

MaxPeak: 100.00%
Ret_Time: 1.128 min

# S238059$1

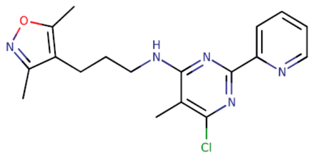


**Mol Wt** 357.84
**Exact Mass** 357.16

| # | Time | Area% |
|---|---|---|
| 1 | 1.128 | 100.00 |

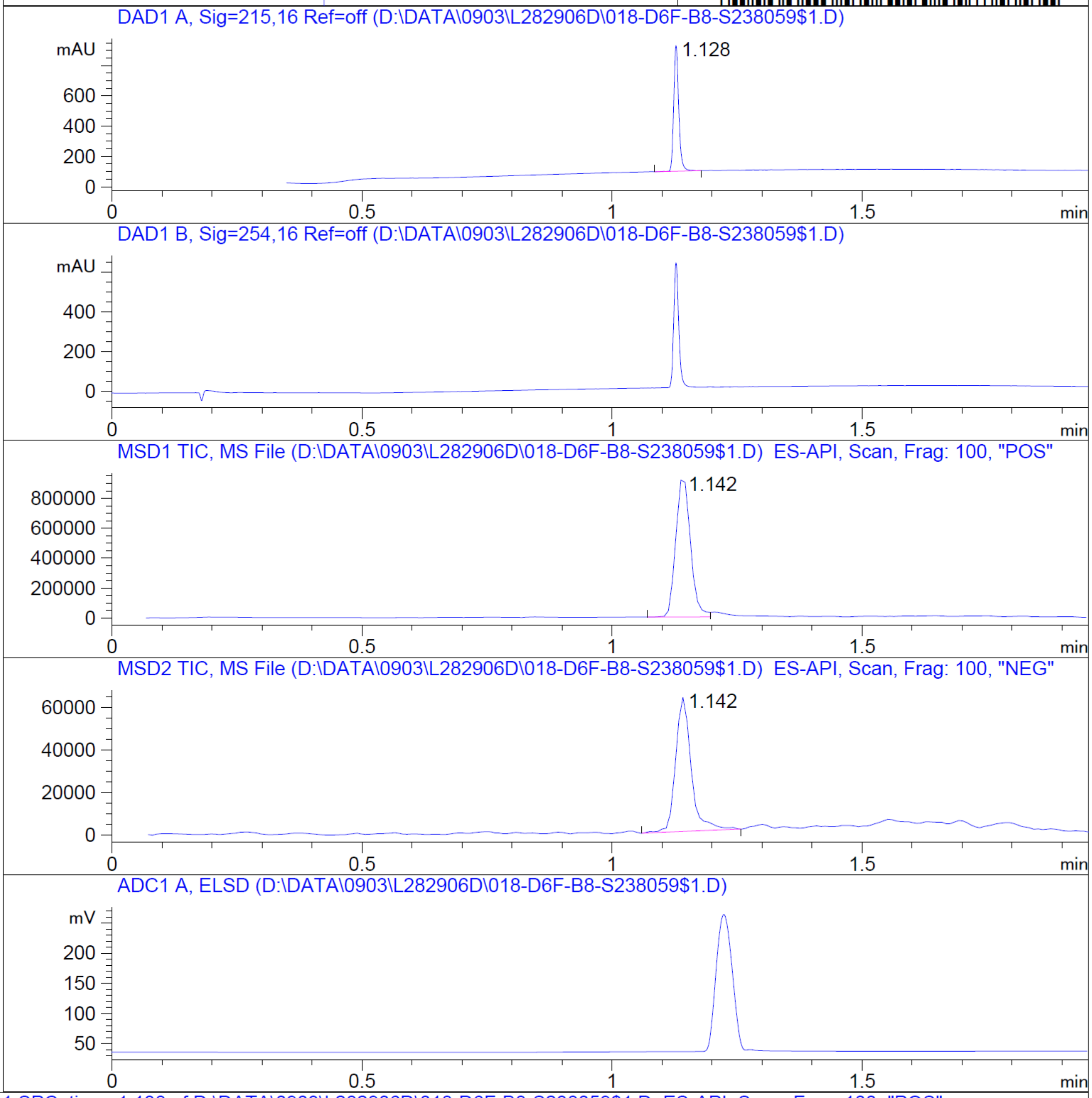


RT 1.142

RT 1.142

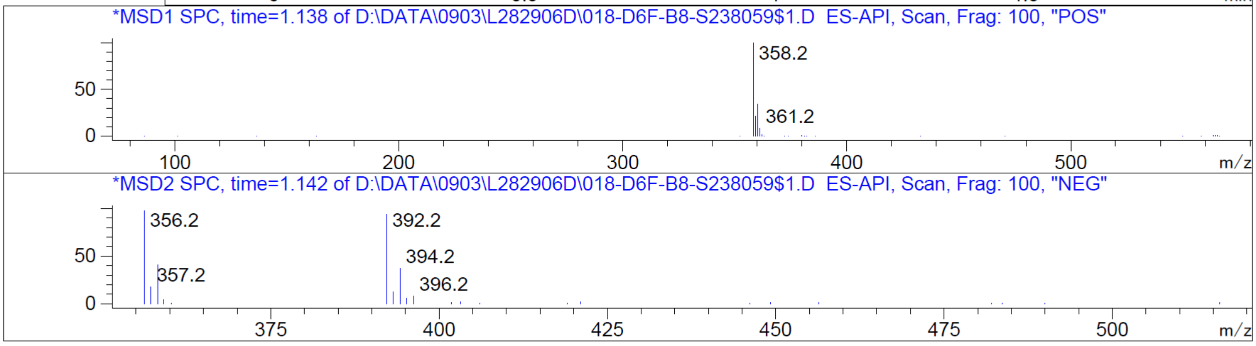


Z4607408926

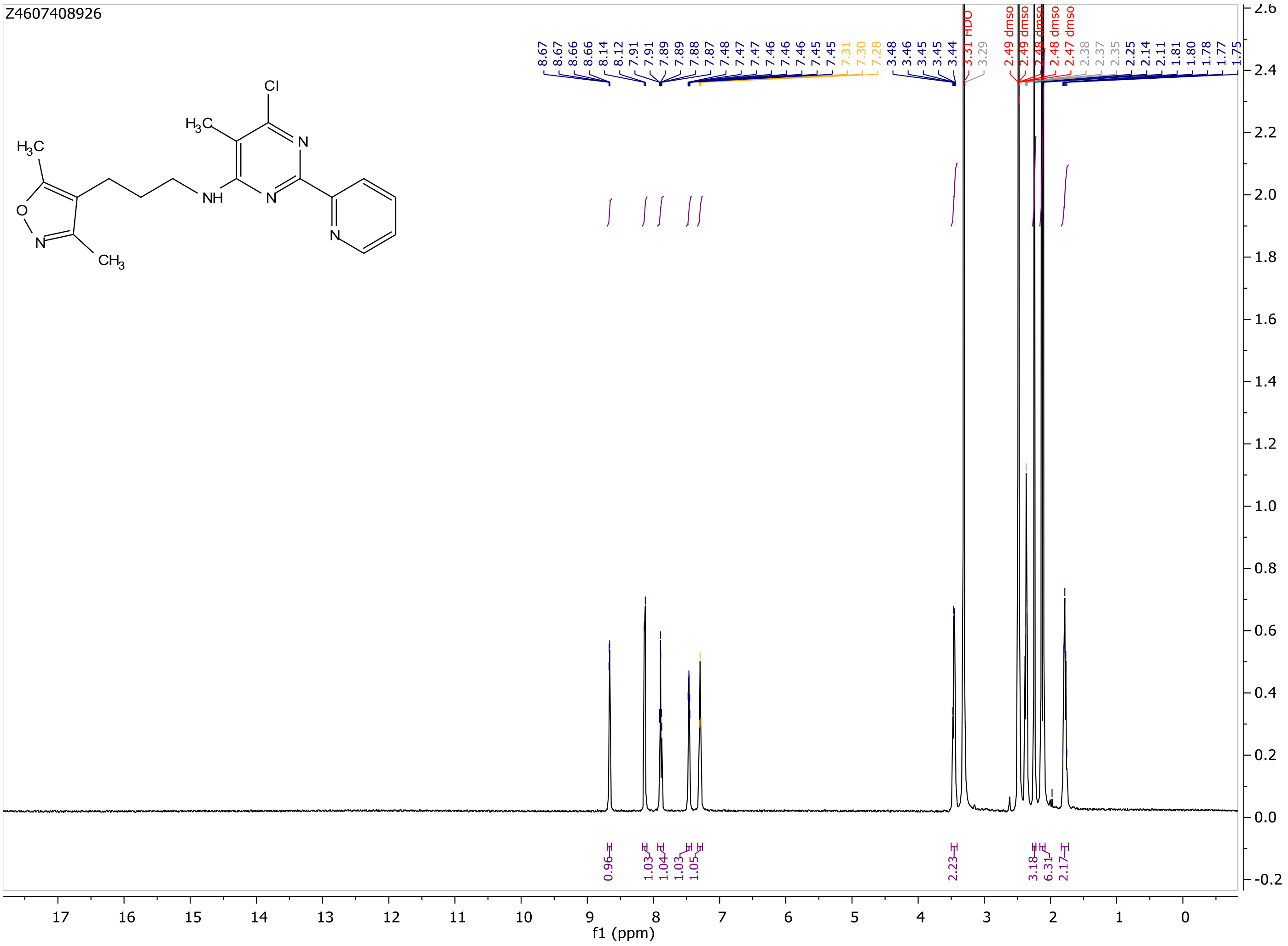
Z4607408926
Cl
$H_3C$
N
NH
N
N
$H_3C$
O
N
$CH_3$
8.67
8.67
8.66
8.66
8.14
8.12
7.91
7.89
7.89
7.88
7.87
7.48
7.47
7.47
7.46
7.46
7.46
7.45
7.45
7.31
7.30
7.28
3.48
3.46
3.45
3.45
3.44
3.31 HDO
3.29
2.49 dmso
2.49 dmso
2.48 dmso
2.48 dmso
2.47 dmso
2.38
2.37
2.35
2.25
2.14
2.11
1.81
1.80
1.78
1.77
1.75
0.96
1.03
1.04
1.03
1.05
2.23
3.18
6.31
2.17
f1 (ppm)

MaxPeak: 91.57%
Ret_Time: 1.147 min

S568076$2

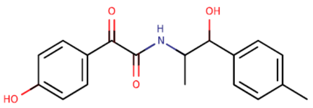

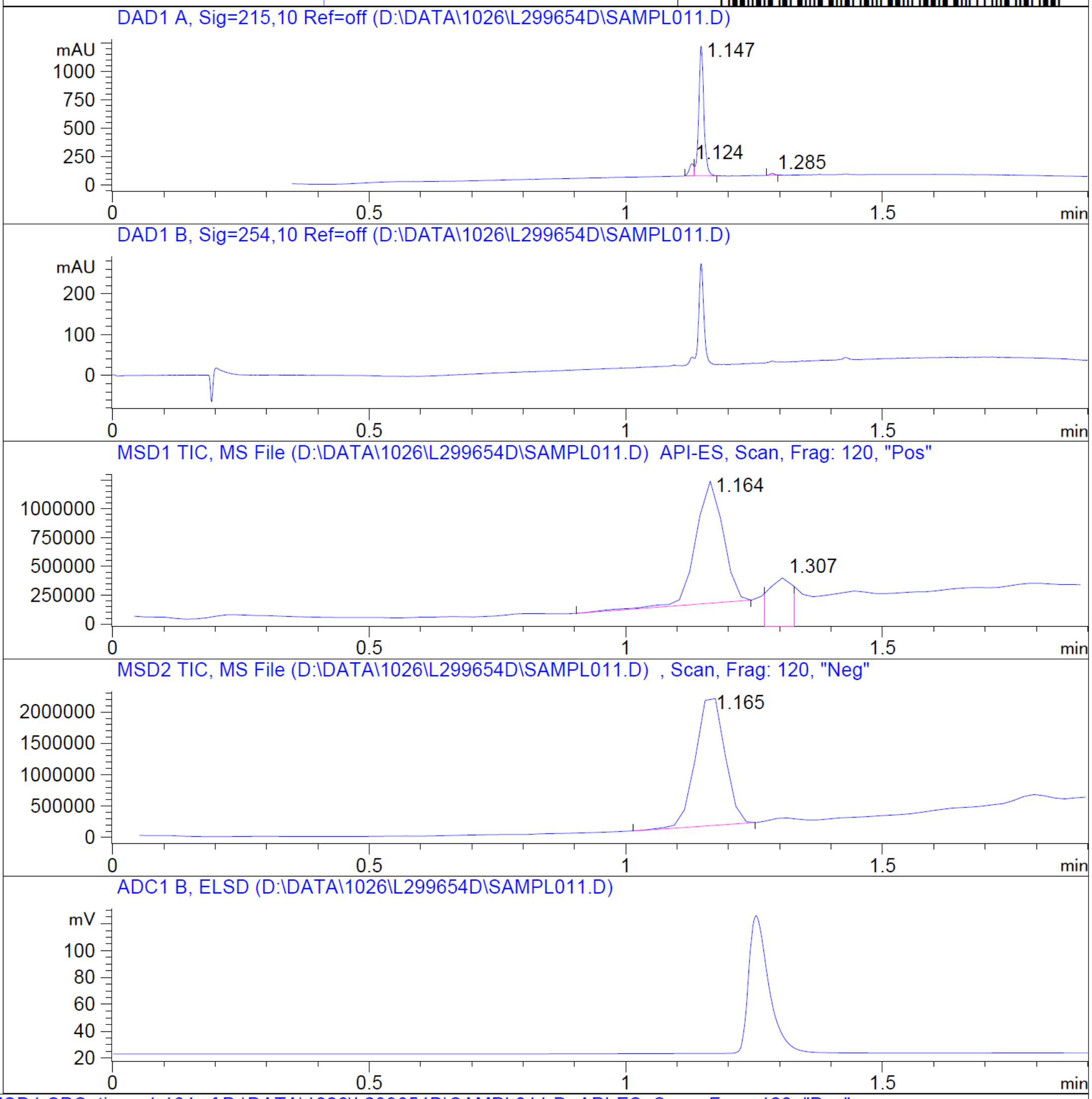


**Mol Wt** 313.35

**Exact Mass** 313.15

| # | Time | Area% |
|---|---|---|
| 1 | 1.124 | 7.28 |
| 2 | 1.147 | 91.57 |
| 3 | 1.285 | 1.15 |

RT 1.164

RT 1.307

RT 1.165

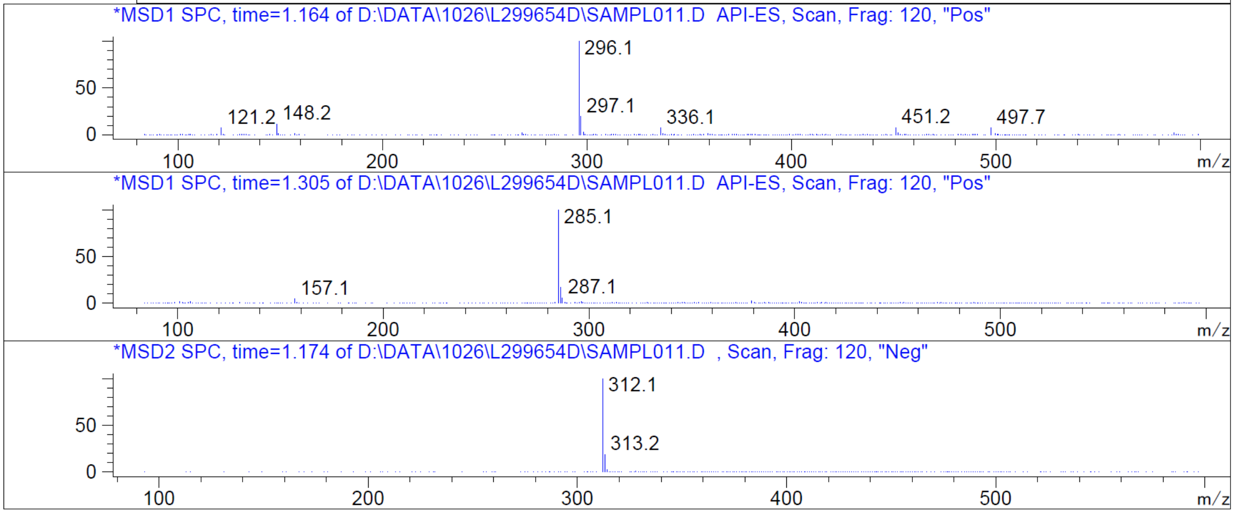


Z2091253662_(6634_3)

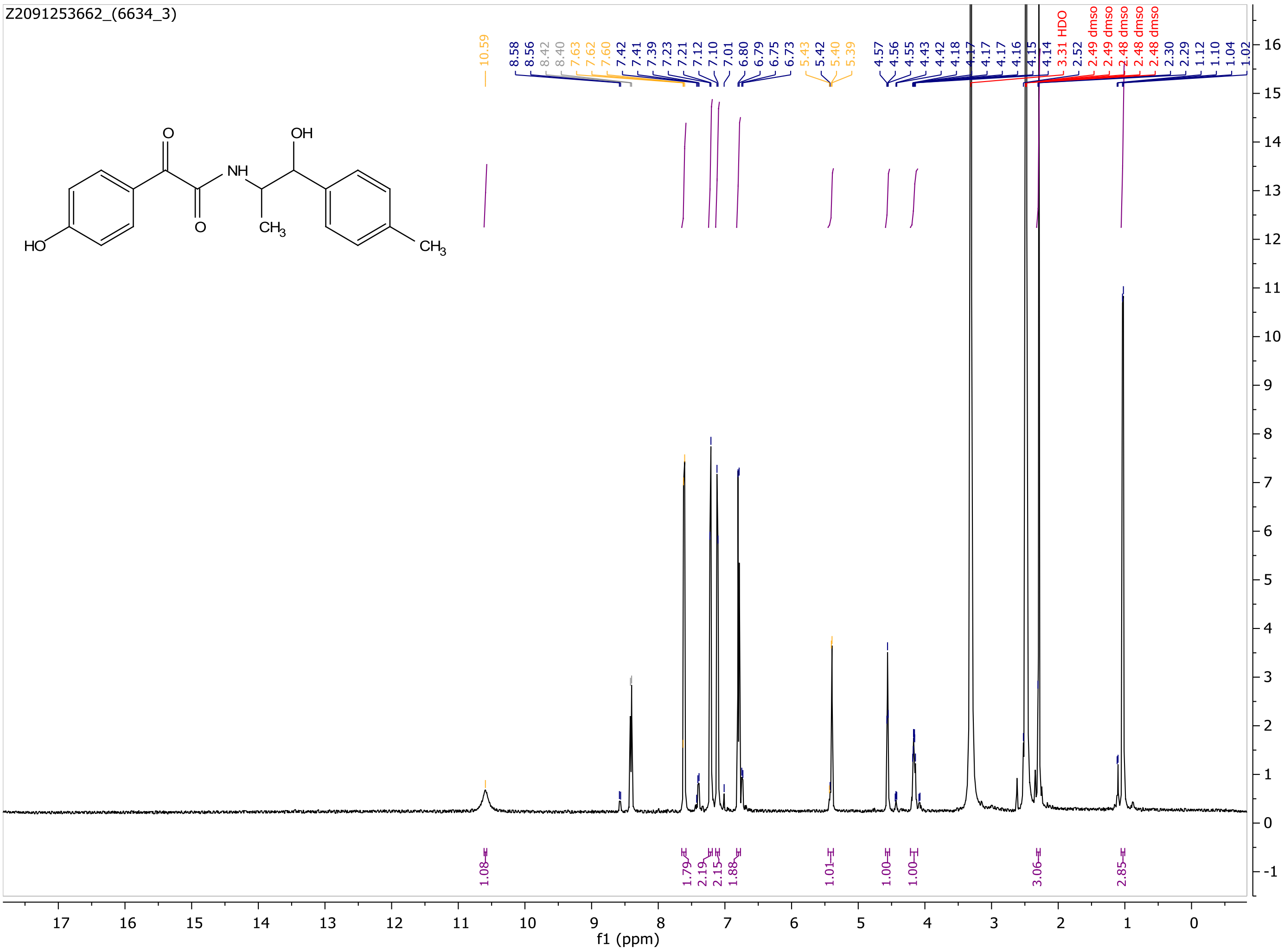
Z2091253662_(6634_3)
10.59
8.58 8.56 8.42 8.40 7.63 7.62 7.60 7.42 7.41 7.39 7.23 7.21 7.12 7.10 7.01 6.80 6.79 6.75 6.73 5.43 5.42 5.40 5.39 4.57 4.56 4.55 4.43 4.42 4.18 4.17 4.17 4.17 4.16 4.15 4.14 3.31 HDO 2.52 2.49 dmso 2.49 dmso 2.48 dmso 2.48 dmso 2.48 dmso 2.30 2.29 1.12 1.10 1.04 1.02
1.08 1.79 2.19 2.15 1.88 1.01 1.00 1.00 3.06 2.85
f1 (ppm)

MaxPeak: 98.42%
Ret_Time: 1.286 min

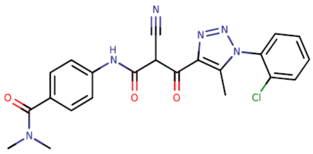

**Mol Wt** 450.88

**Exact Mass** 450.13

| # | Time | Area% |
|---|---|---|
| 1 | 0.808 | 1.58 |
| 2 | 1.286 | 98.42 |

# S919151$5

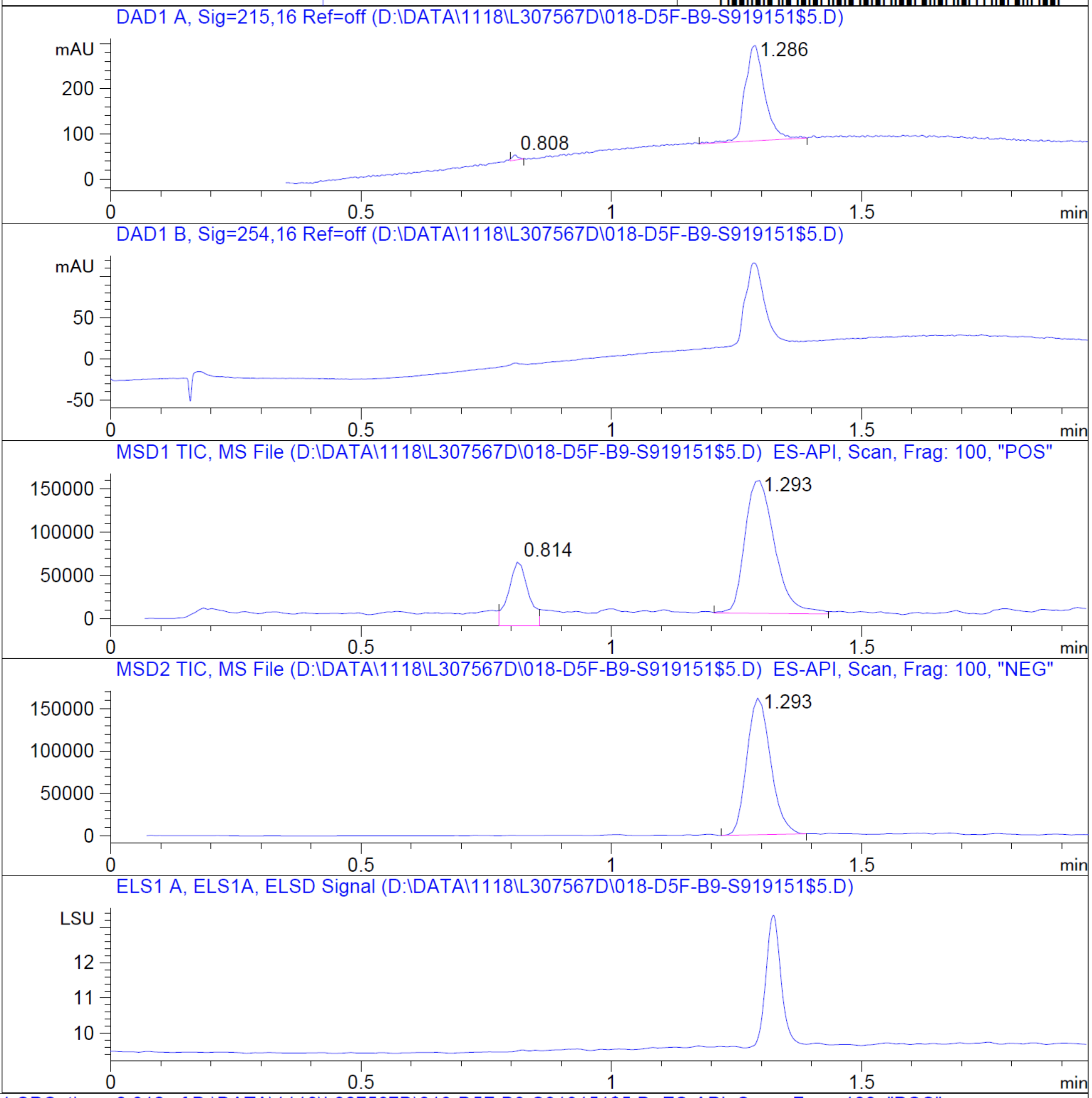


RT 0.814

RT 1.293

RT 1.293

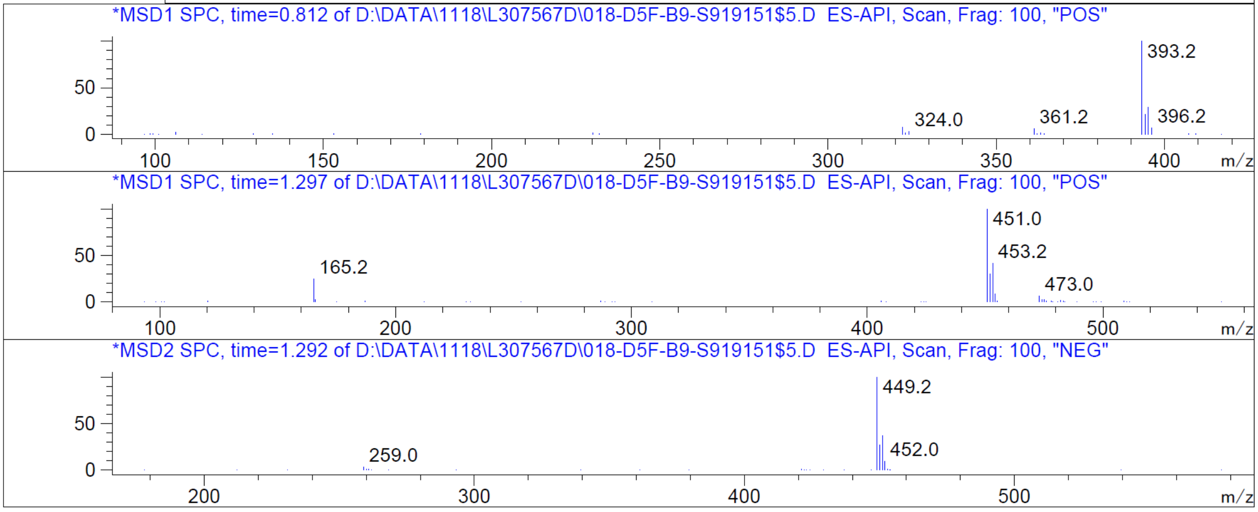


Z1123377193_(6792_6)

Z1123377193_(6792_6)

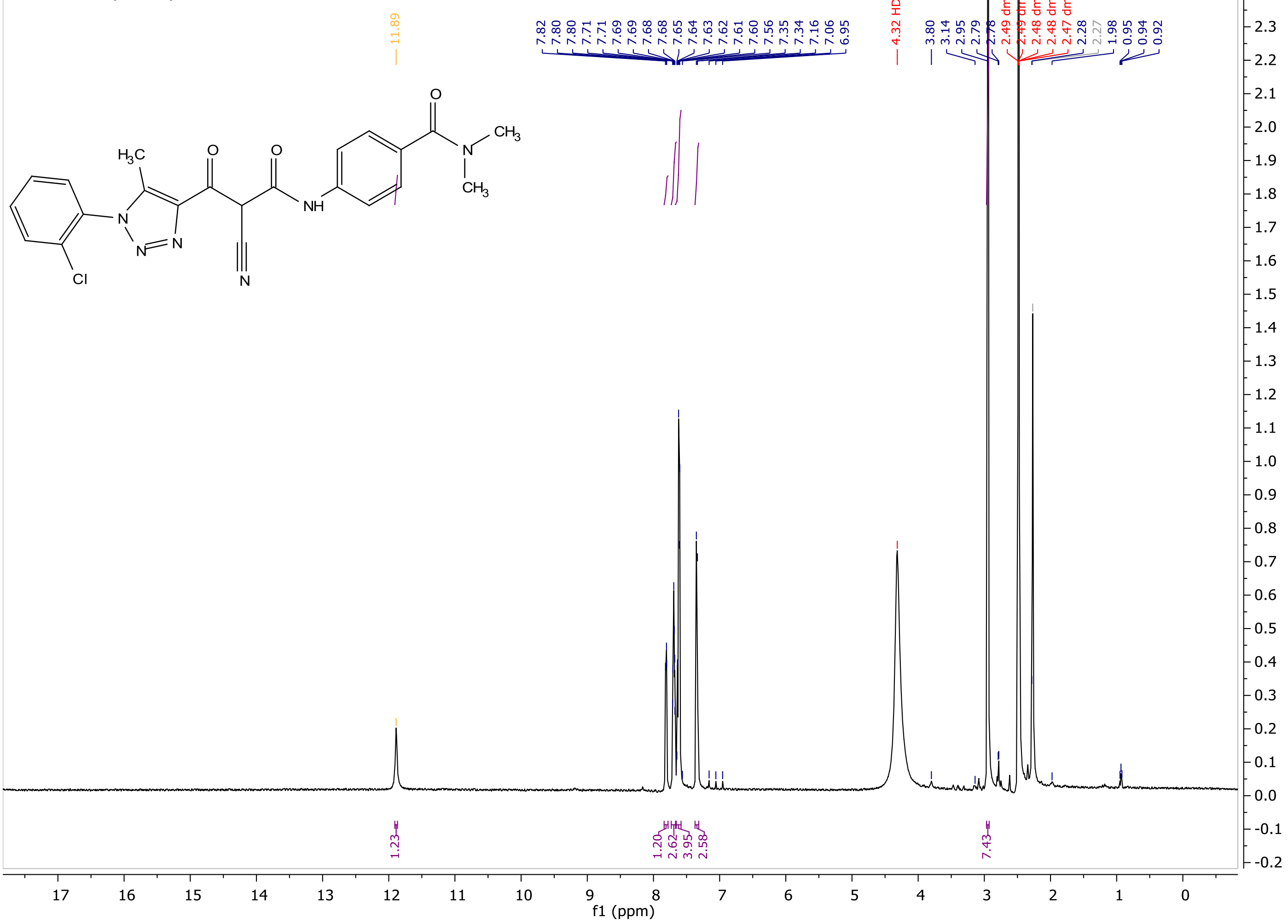

MaxPeak: 100.00%
Ret_Time: 1.281 min

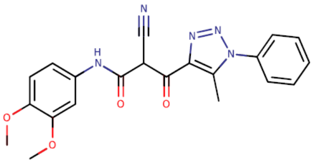

**Mol Wt** 405.41
**Exact Mass** 405.15

| # | Time | Area% |
|---|---|---|
| 1 | 1.281 | 100.00 |

# S910024$A

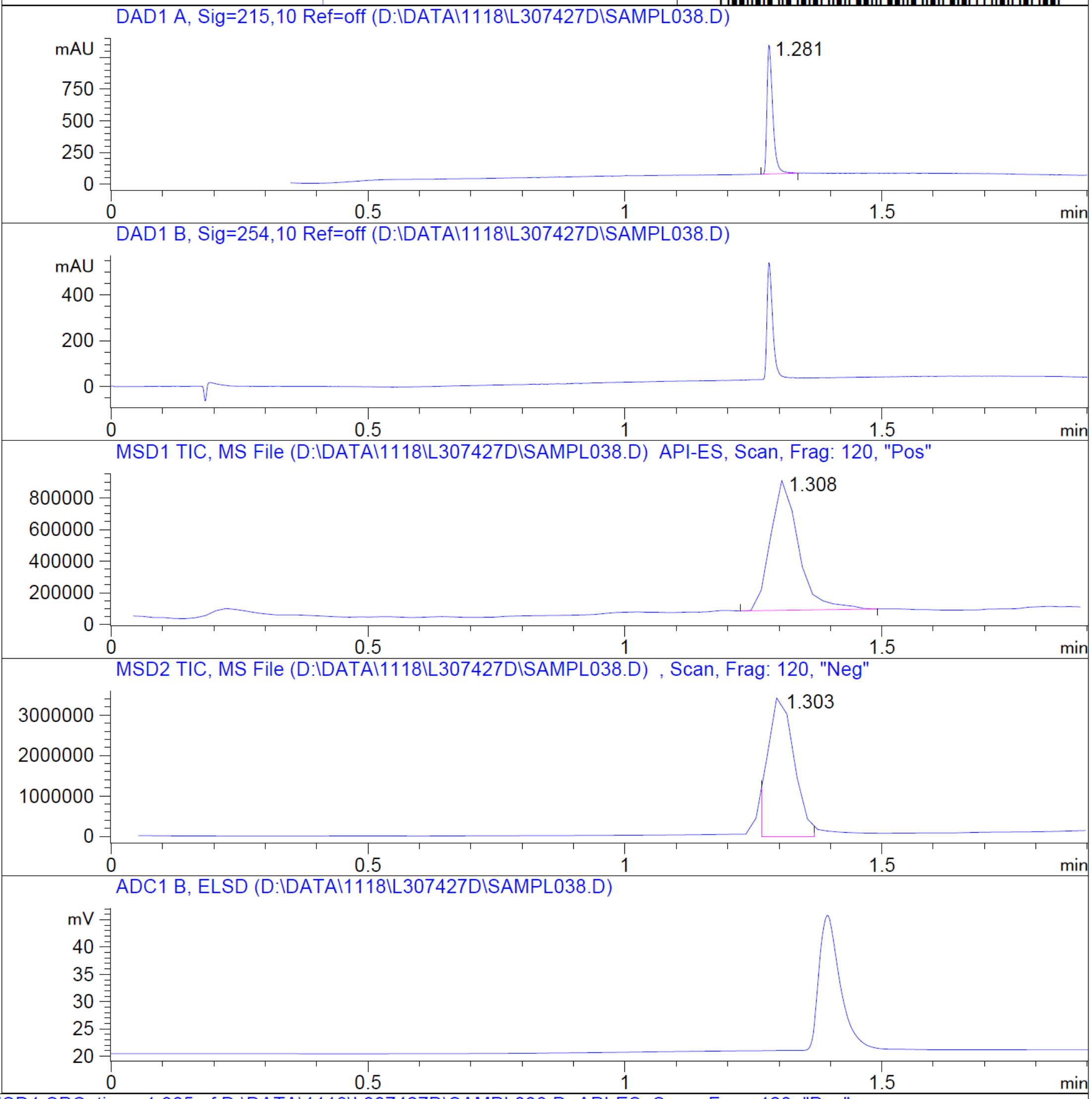


RT 1.308

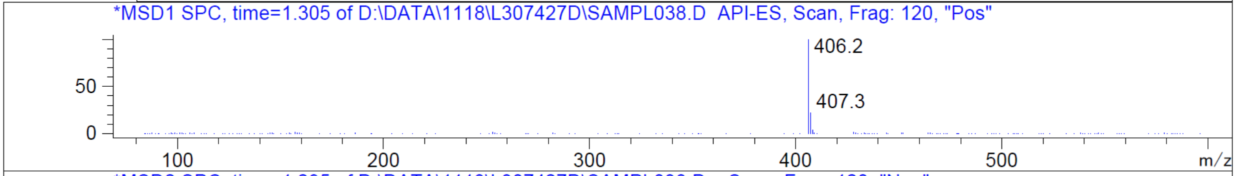


RT 1.303

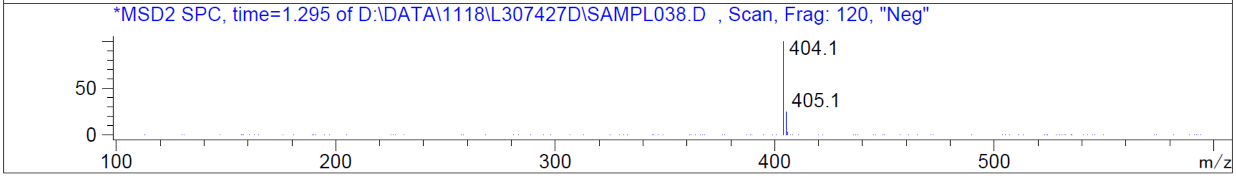


Z762211478_(6792_5)

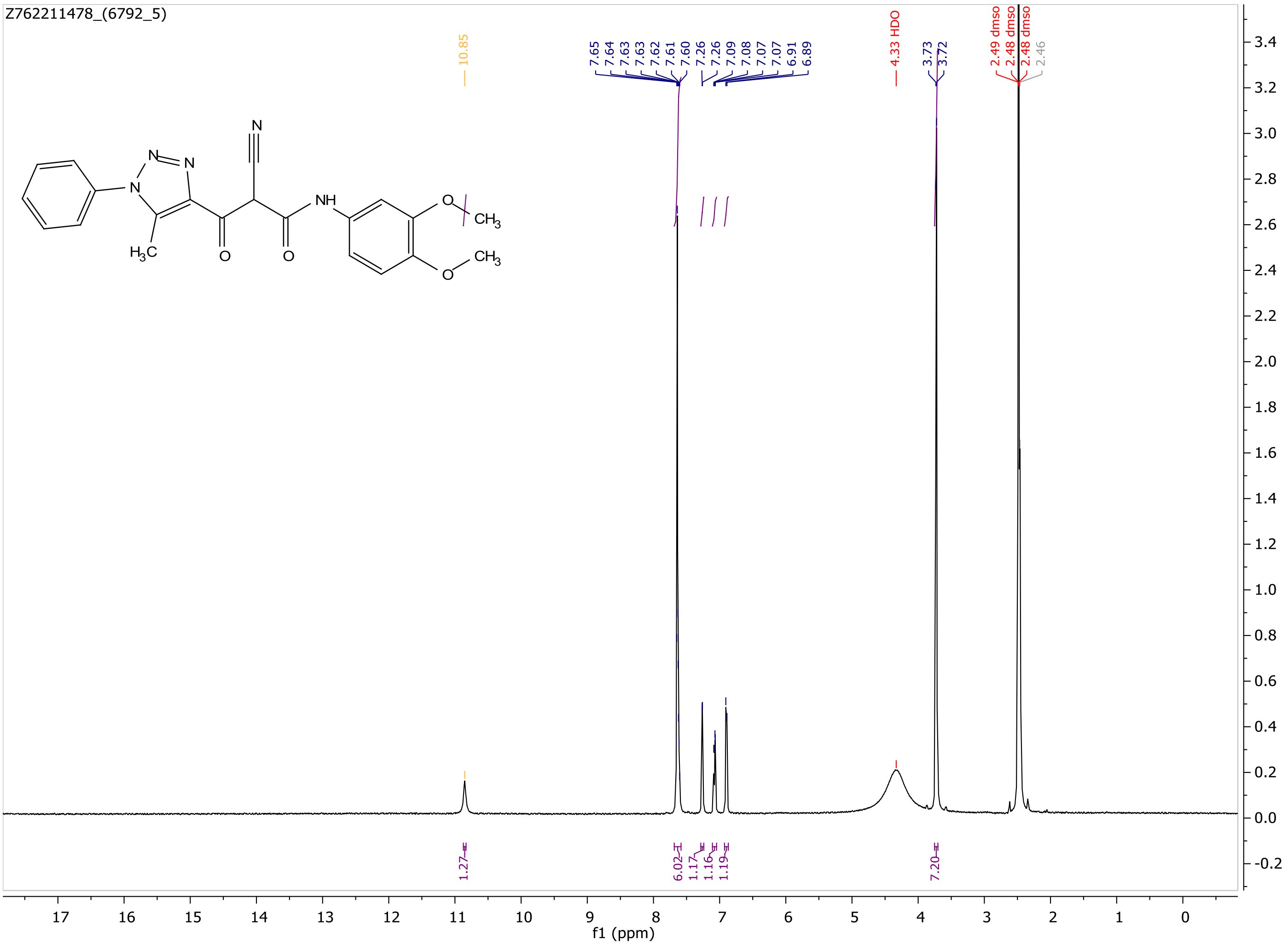
Z762211478_(6792_5)
10.85
7.65
7.64
7.63
7.63
7.62
7.61
7.60
7.26
7.26
7.09
7.08
7.07
7.07
6.91
6.89
4.33 HDO
3.73
3.72
2.49 dmso
2.48 dmso
2.48 dmso
2.46
1.27
6.02
1.17
1.16
1.19
7.20
f1 (ppm)

MaxPeak: 98.46%
Ret_Time: 1.428 min

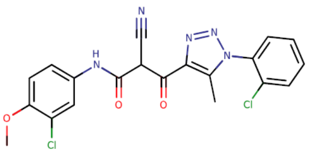

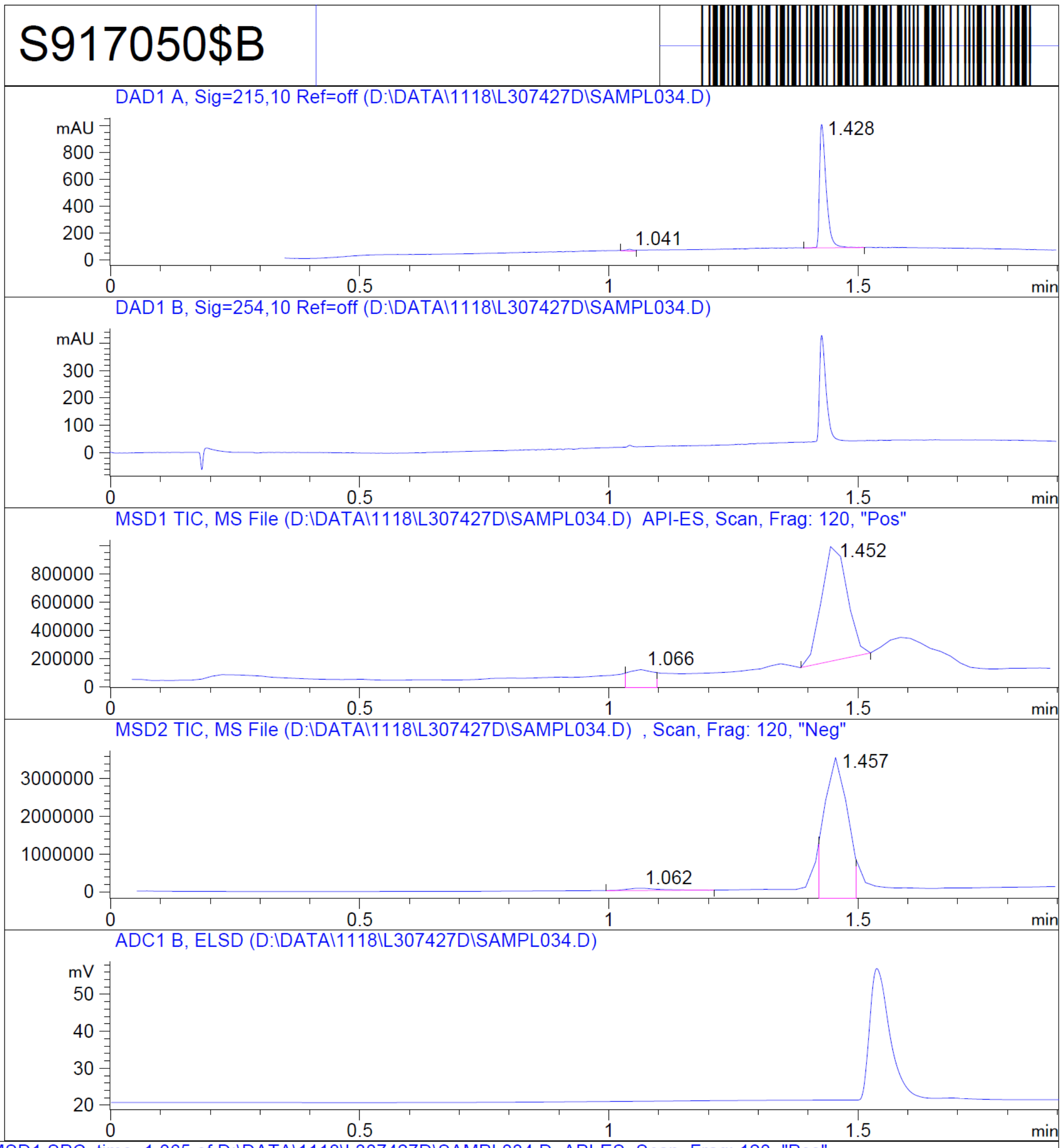


**Mol Wt** 444.27
**Exact Mass** 443.06

| # | Time | Area% |
|---|---|---|
| 1 | 1.041 | 1.54 |
| 2 | 1.428 | 98.46 |

RT 1.066

RT 1.452

RT 1.062

RT 1.457

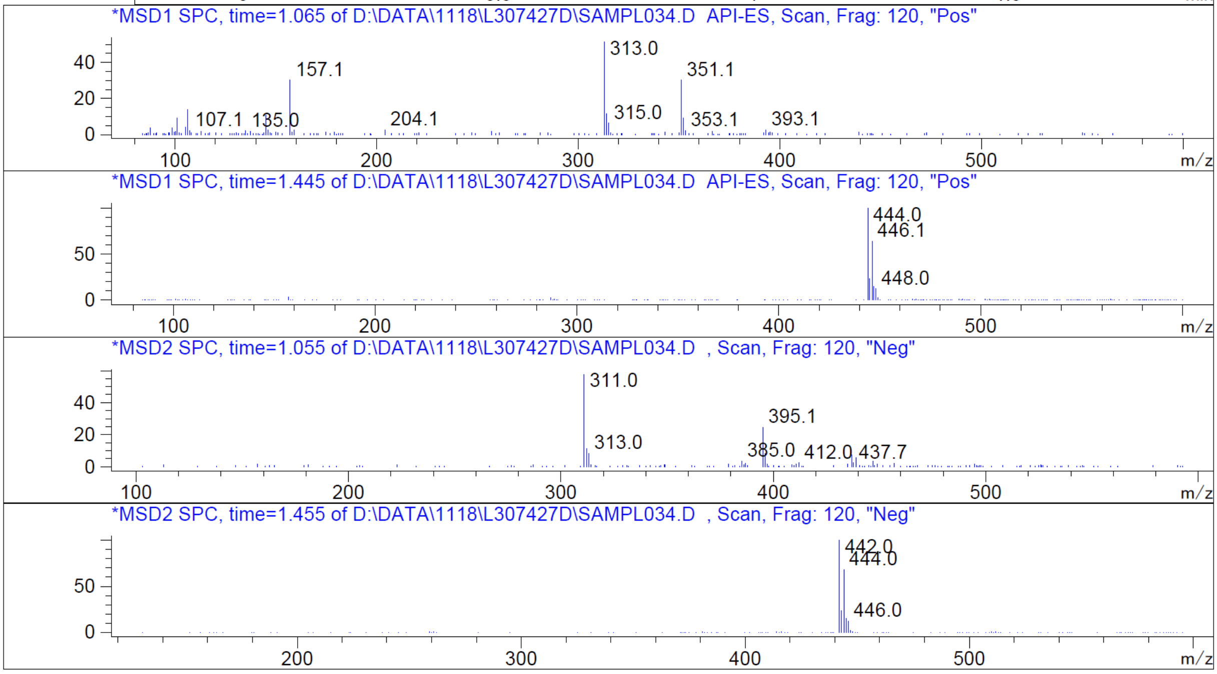


Z1123371957_(6792_3)

Z1123371957_(6792_3)

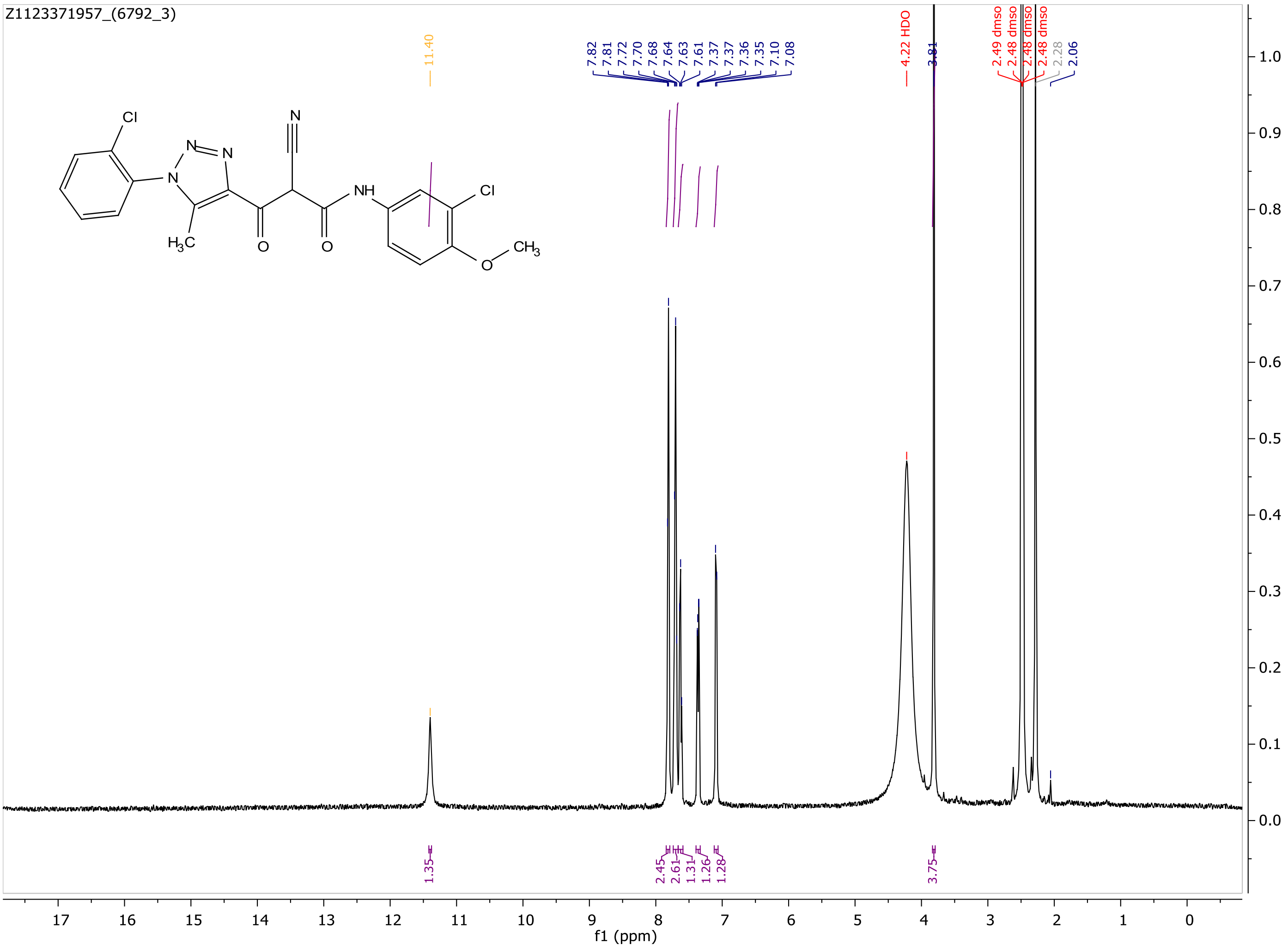

MaxPeak: 100.00%
Ret_Time: 1.448 min

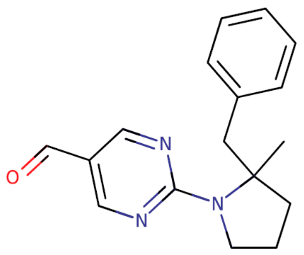

**Mol Wt** 281.35
**Exact Mass** 281.18

| # | Time | Area% |
|---|---|---|
| 1 | 1.448 | 100.00 |

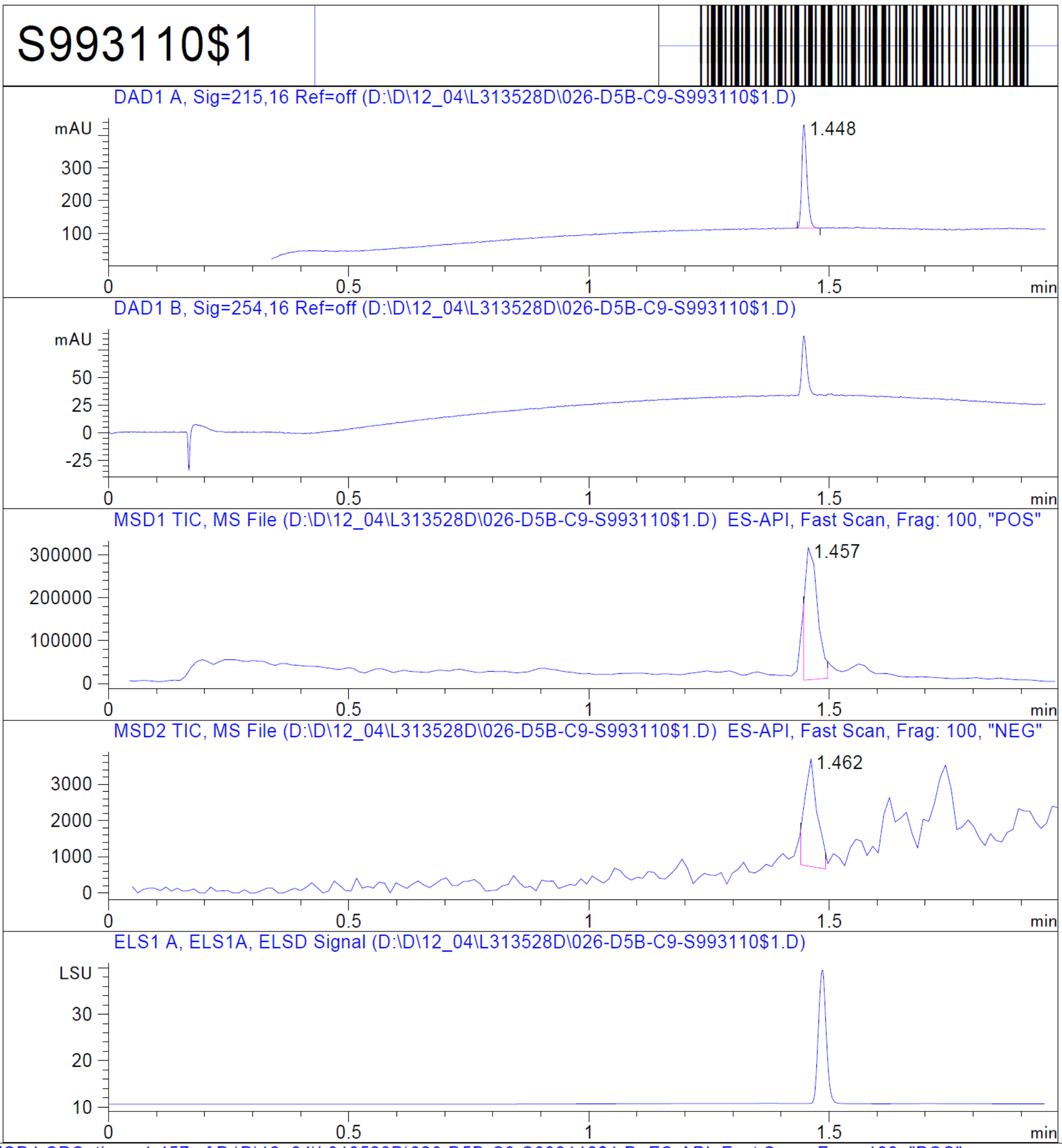


RT 1.457

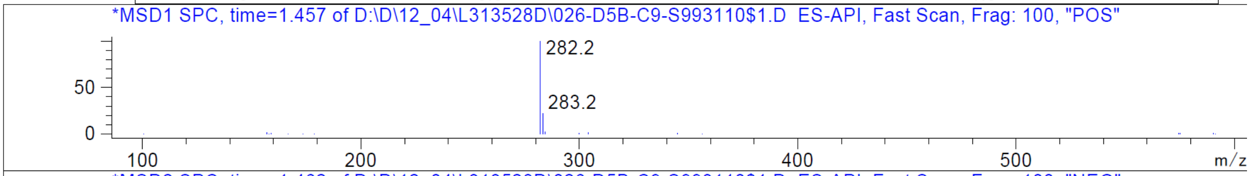


RT 1.462

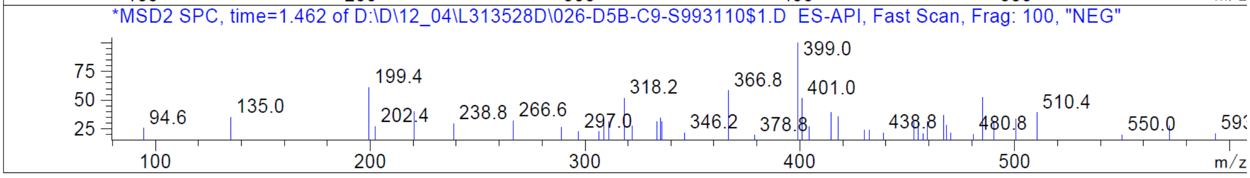


Z2851084476_(ZINC882196186)

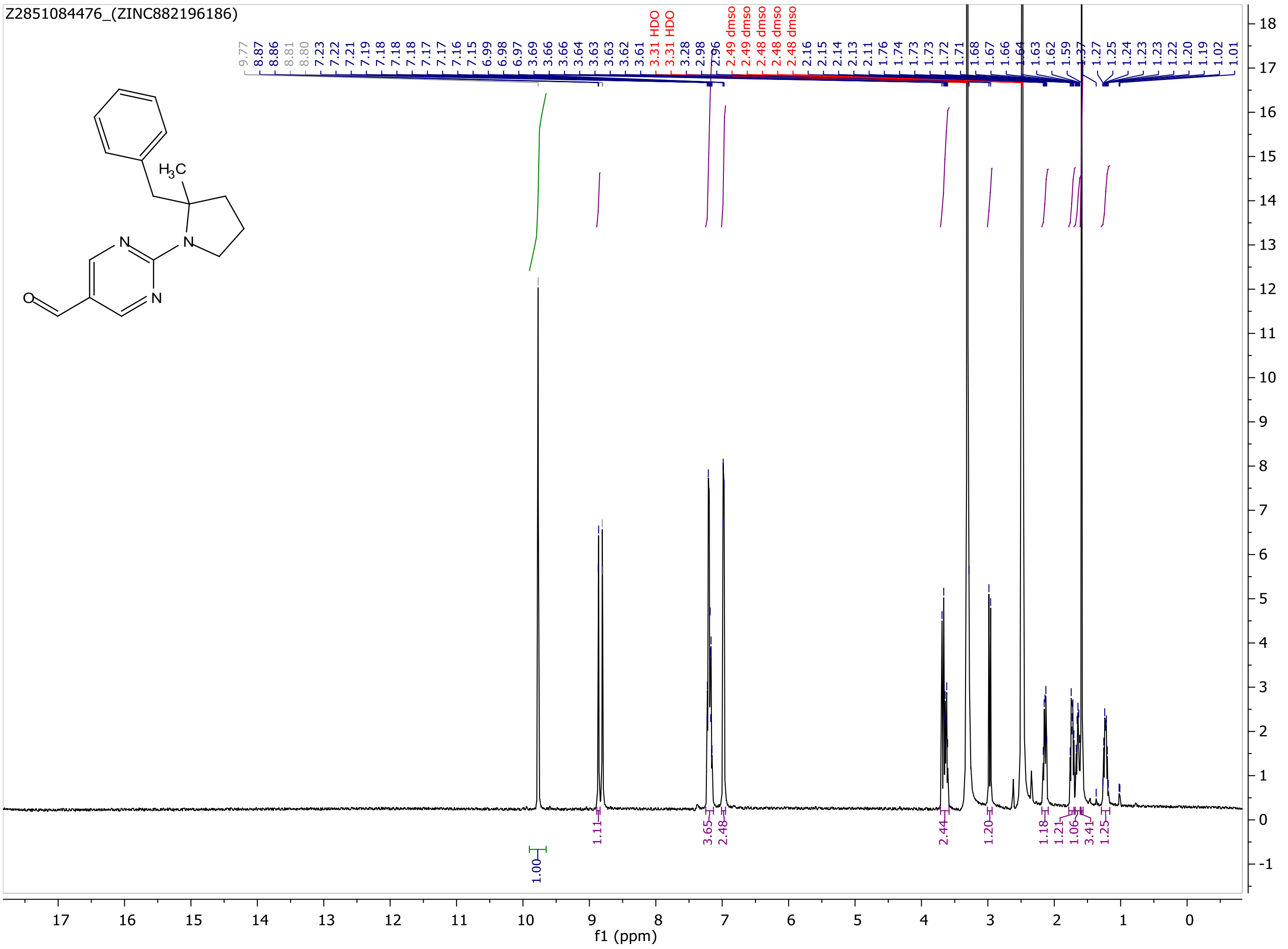
Z2851084476_(ZINC882196186)
H3C
N
N
N
O
f1 (ppm)

MaxPeak: 100.00%
Ret_Time: 1.308 min

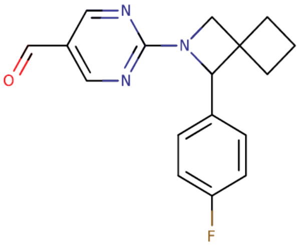


**Mol Wt** 297.33

**Exact Mass** 297.15

| # | Time | Area% |
|---|---|---|
| 1 | 1.308 | 100.00 |

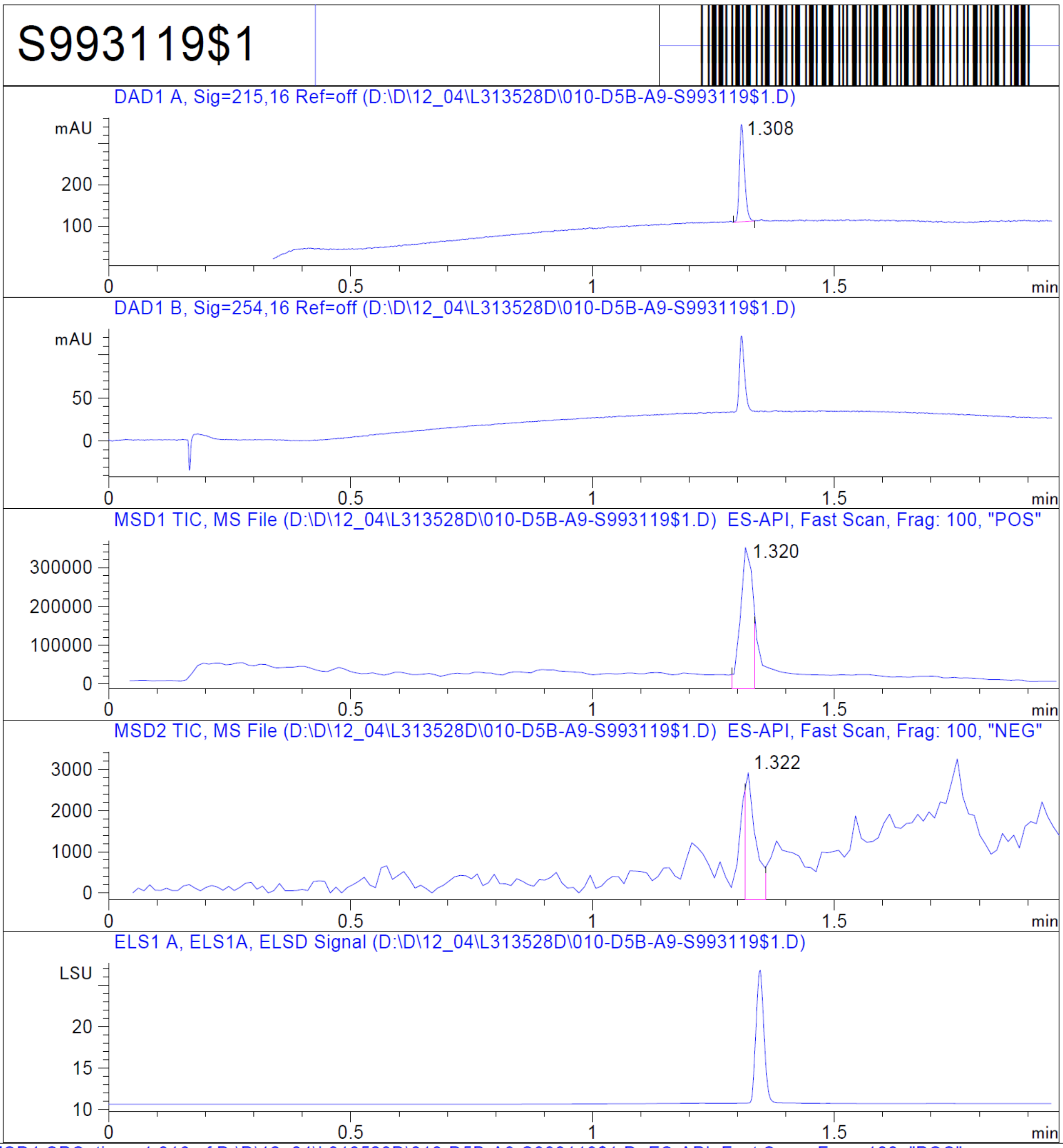


RT 1.320

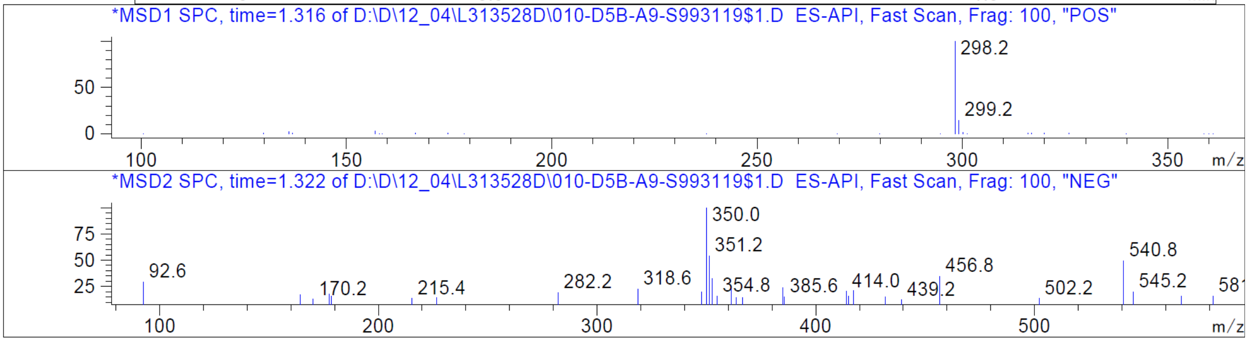


RT 1.322

Z2851916546

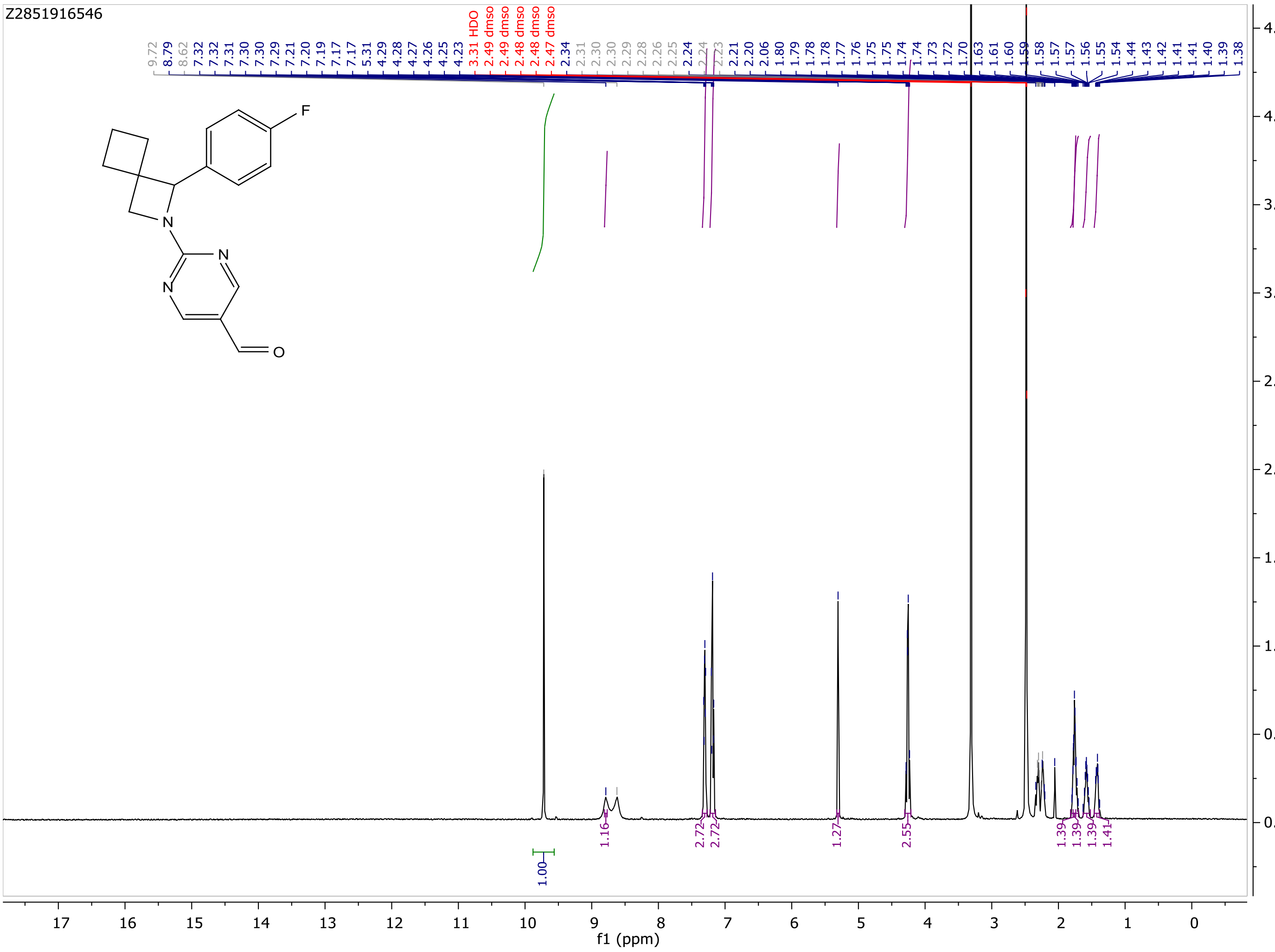
Z2851916546
9.72 8.79 8.62 7.32 7.32 7.31 7.30 7.30 7.29 7.21 7.20 7.19 7.17 7.17 5.31 4.29 4.28 4.27 4.26 4.25 4.23
3.31 HDO 2.49 dmso 2.49 dmso 2.48 dmso 2.48 dmso 2.47 dmso
2.34 2.31 2.30 2.30 2.29 2.28 2.26 2.25 2.24 2.24 2.23 2.21 2.20 2.06 1.80 1.79 1.78 1.78 1.77 1.76 1.75 1.75 1.74 1.74 1.73 1.72 1.70 1.63 1.61 1.60 1.59 1.58 1.57 1.57 1.56 1.55 1.54 1.44 1.43 1.42 1.41 1.41 1.40 1.39 1.38
F
N
N
N
O
1.00
1.16
2.72
2.72
1.27
2.55
1.39
1.39
1.39
1.41
f1 (ppm)

MaxPeak: 90.85%
Ret_Time: 1.344 min

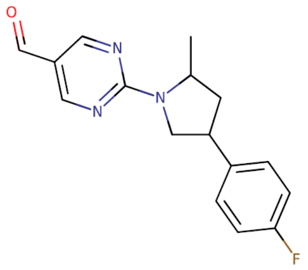


**Mol Wt** 285.32
**Exact Mass** 285.15

| # | Time | Area% |
|---|---|---|
| 1 | 1.324 | 9.15 |
| 2 | 1.344 | 90.85 |

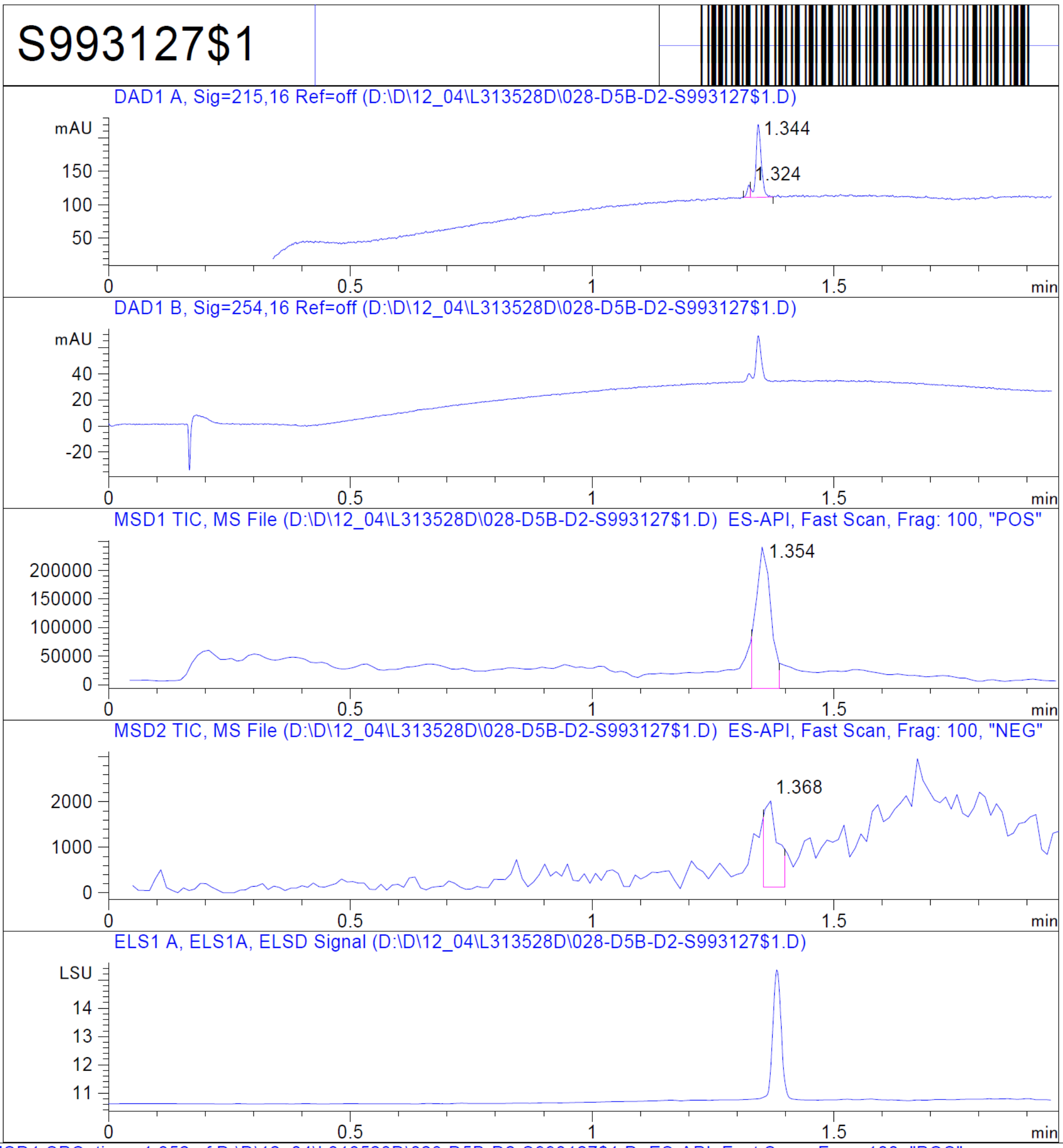


RT 1.354

RT 1.368

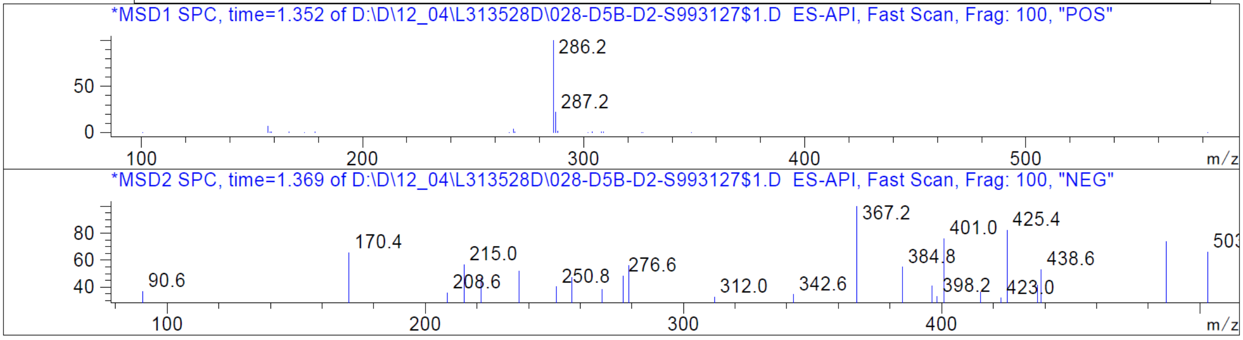


Z2850584231_(ZINC881997690)

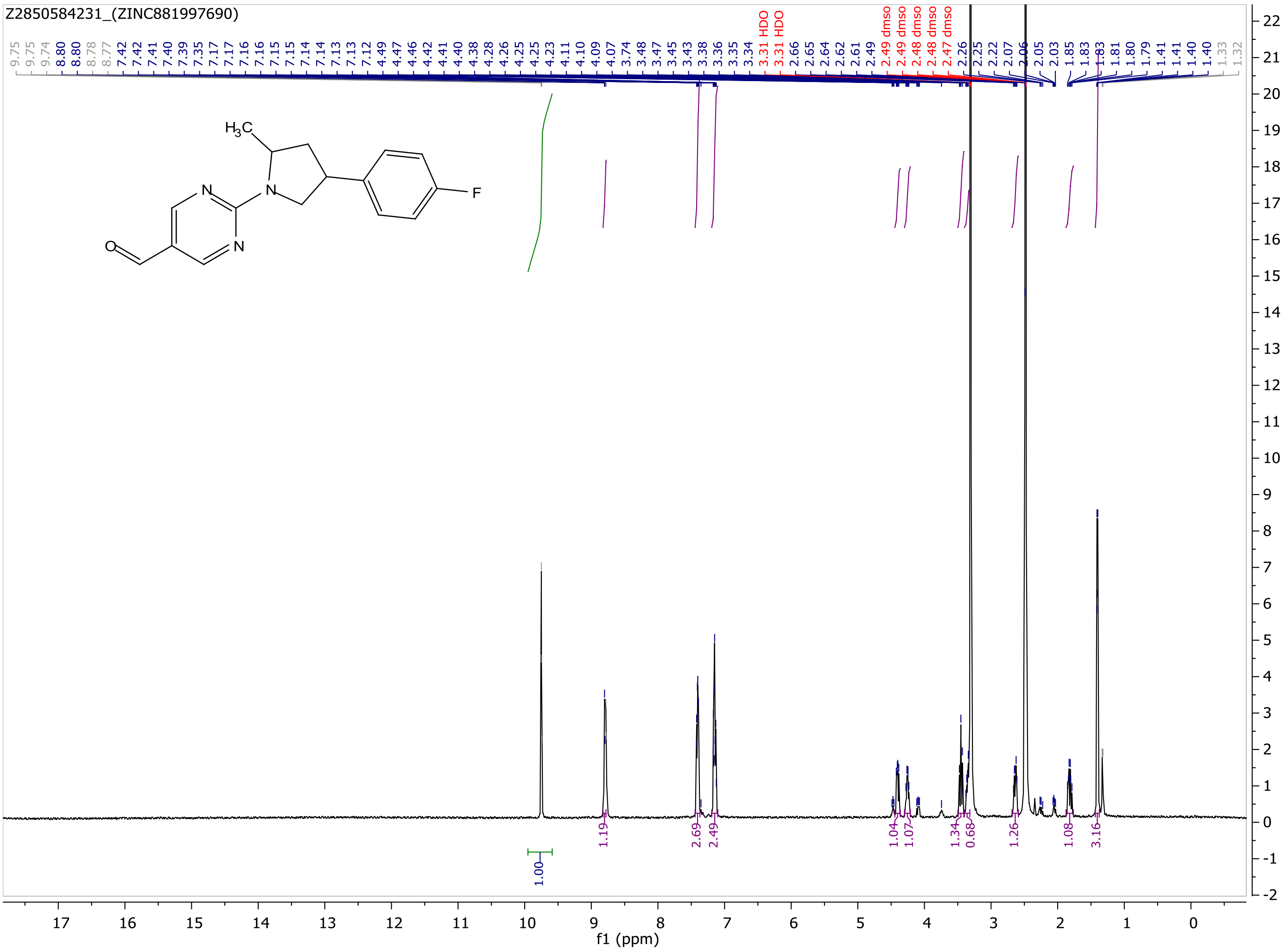
Z2850584231_(ZINC881997690)
9.75 9.75 9.74 8.80 8.80 8.78 8.77 7.42 7.42 7.41 7.40 7.39 7.35 7.17 7.17 7.16 7.15 7.15 7.14 7.14 7.13 7.13 7.12 4.49 4.47 4.46 4.42 4.41 4.40 4.38 4.28 4.26 4.25 4.25 4.23 4.11 4.10 4.09 4.07 3.74 3.48 3.47 3.45 3.43 3.38 3.36 3.35 3.34 3.31 HDO 3.31 HDO 2.66 2.65 2.64 2.62 2.61 2.49 2.49 dmso 2.49 dmso 2.48 dmso 2.48 dmso 2.47 dmso 2.26 2.25 2.22 2.07 2.06 2.05 2.03 1.85 1.83 1.83 1.81 1.80 1.79 1.41 1.41 1.40 1.40 1.33 1.32
H3C
N
N
N
O
F
1.00
1.19
2.69
2.49
1.04
1.07
1.34
0.68
1.26
1.08
3.16
f1 (ppm)

MaxPeak: 100.00%
Ret_Time: 1.625 min

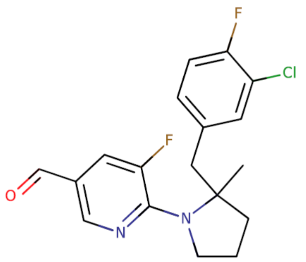


**Mol Wt** 350.79

**Exact Mass** 350.13

| # | Time | Area% |
|---|---|---|
| 1 | 1.625 | 100.00 |

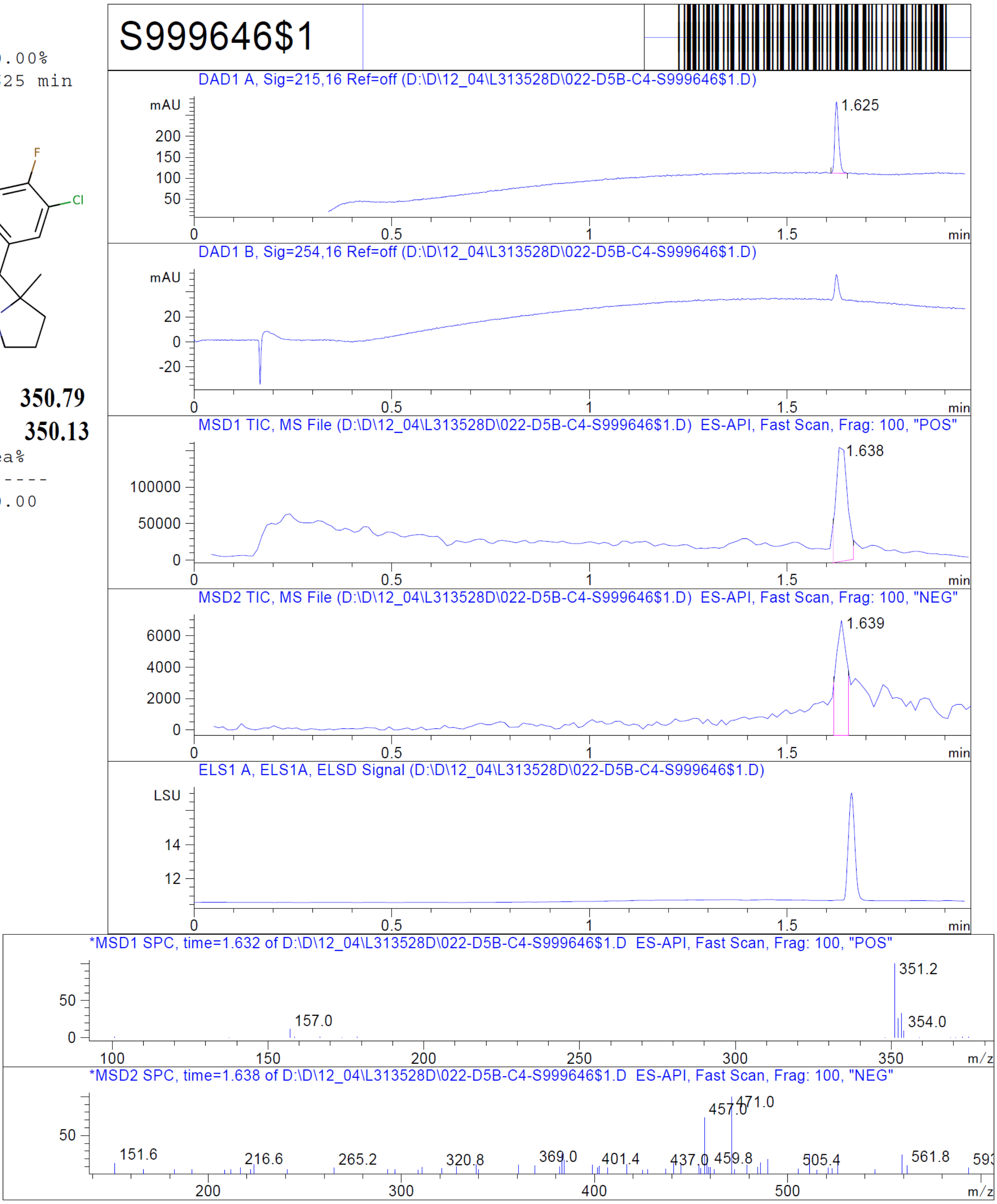


RT 1.638

RT 1.639

Z4767502333

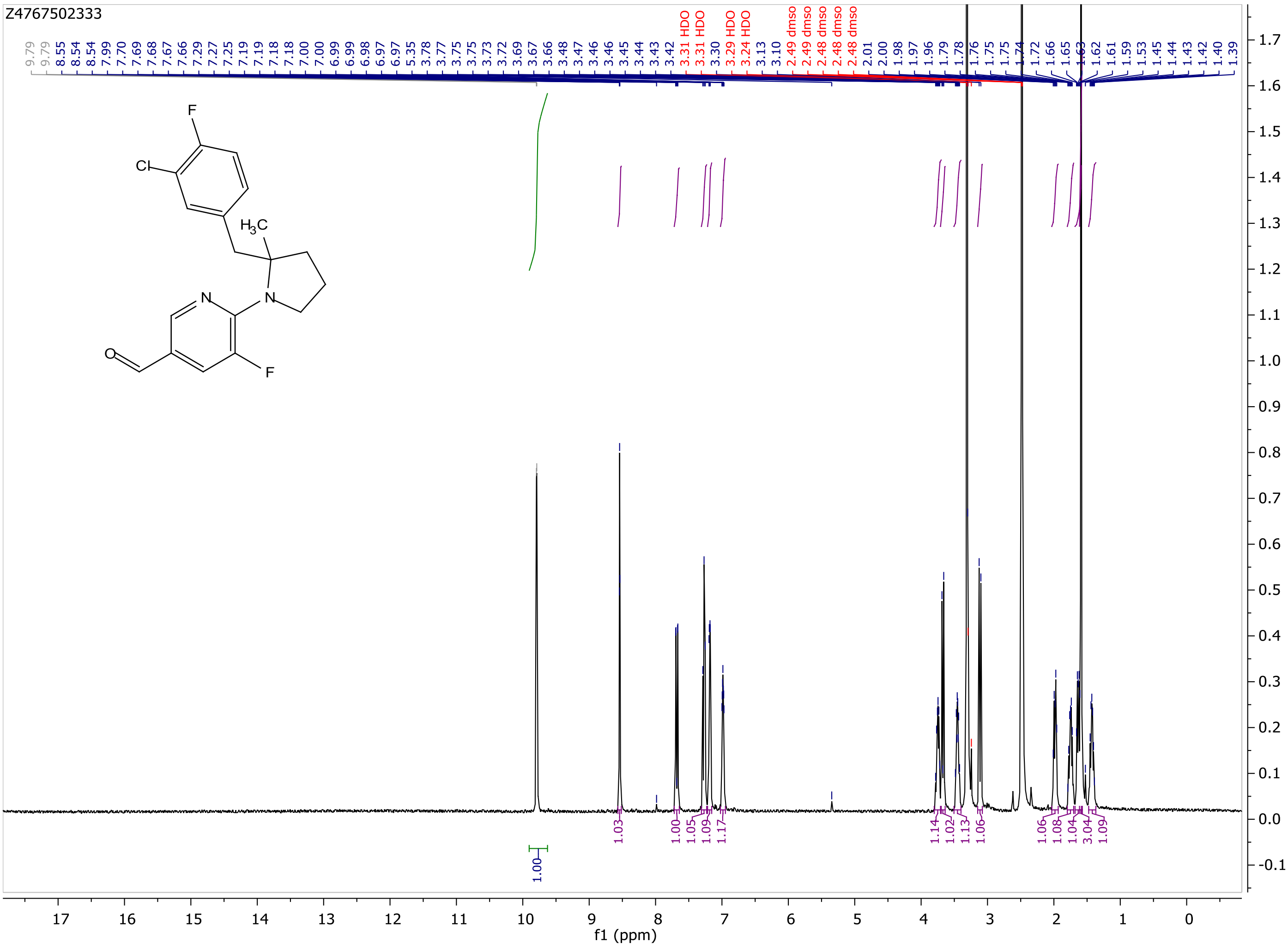
Z4767502333
f1 (ppm)

MaxPeak: 100.00%
Ret_Time: 1.569 min

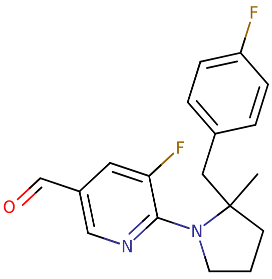


**Mol Wt** 316.35
**Exact Mass** 316.17

| # | Time | Area% |
|---|---|---|
| 1 | 1.569 | 100.00 |

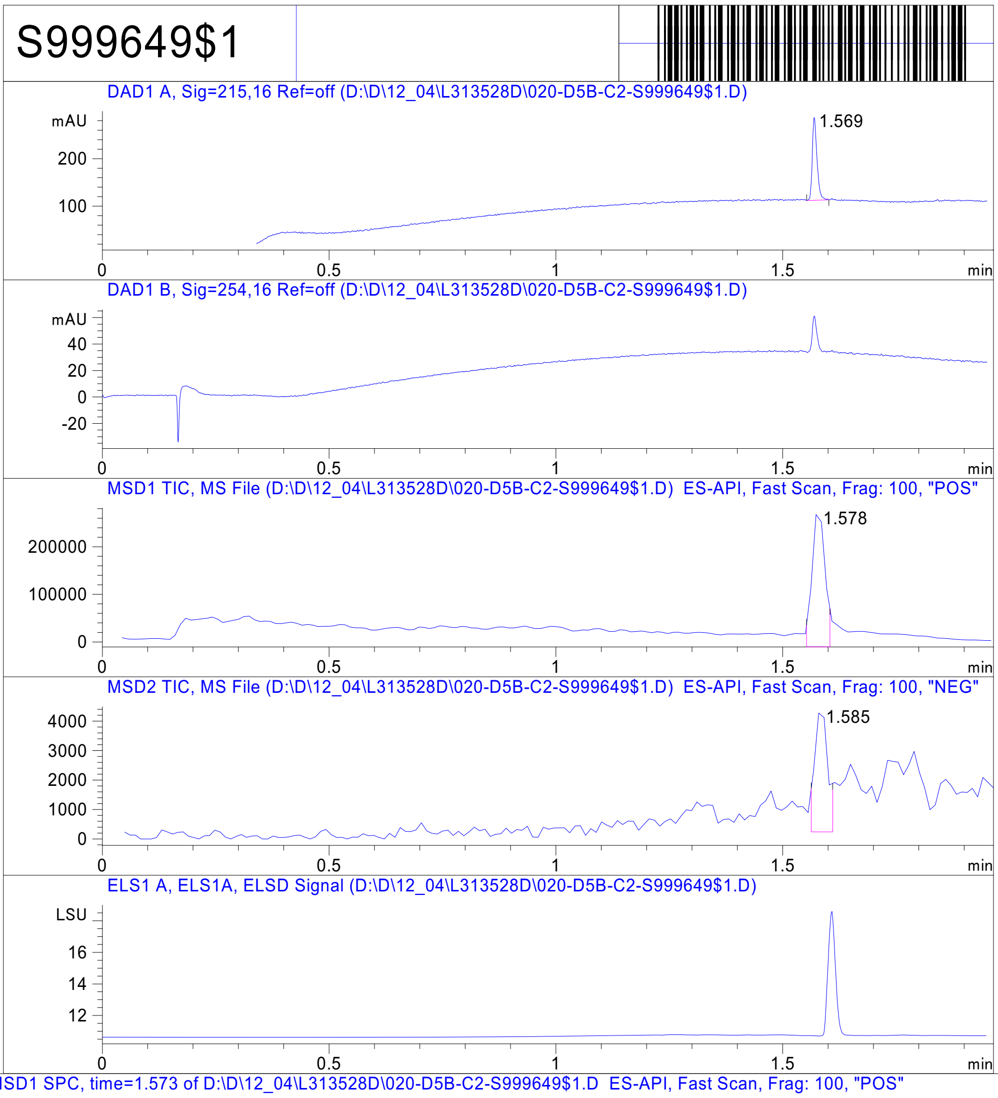


RT 1.578

RT 1.585

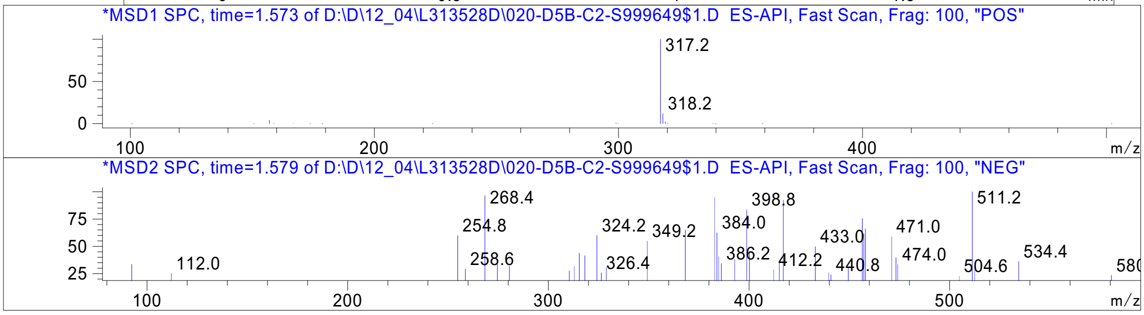


Inj.Date 12/3/2020 N Acq. Method C:\Users\ -> ->

Z4767502336_

MaxPeak: 100.00%
Ret_Time: 1.062 min

S993149$3

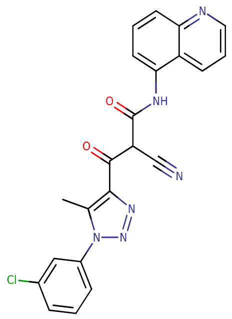

**Mol Wt** 430.85

**Exact Mass** 430.1

| # | Time | Area% |
| --- | --- | --- |
| 1 | 1.062 | 100.00 |

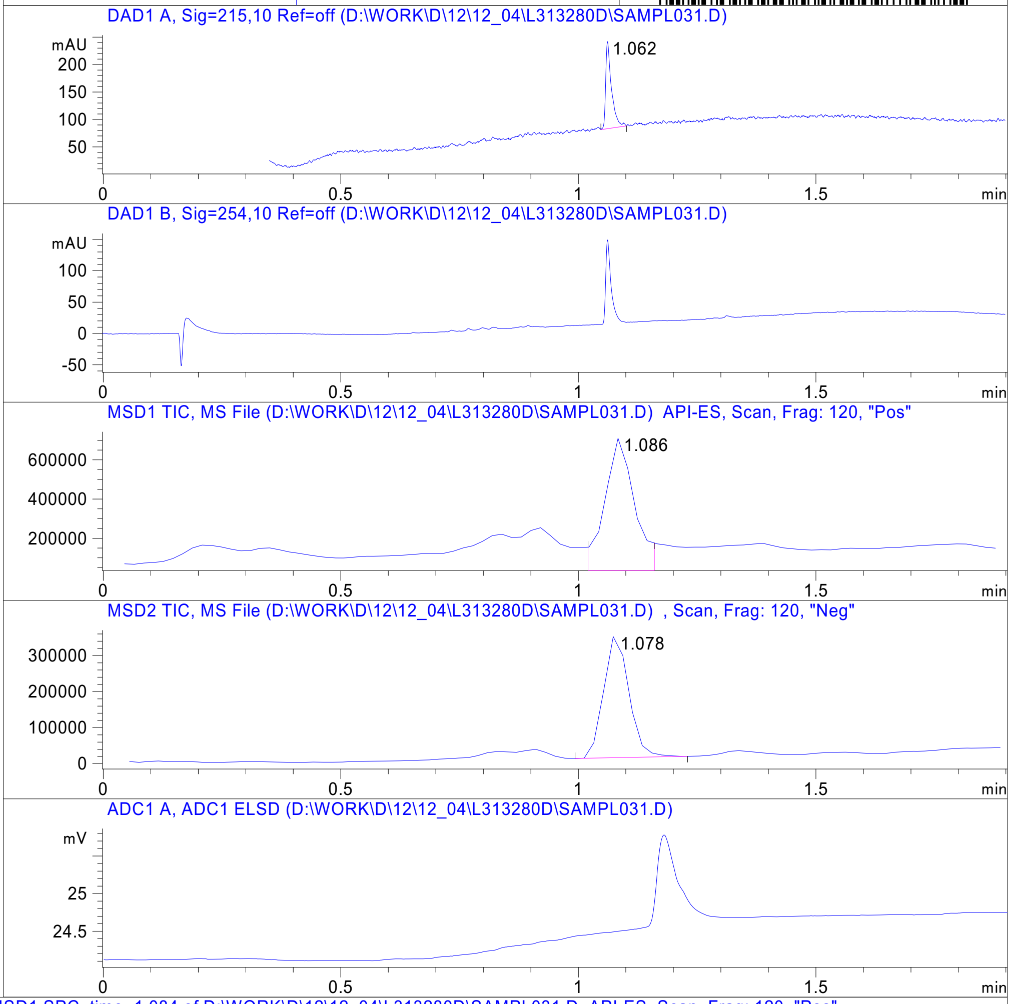


RT 1.086

RT 1.078

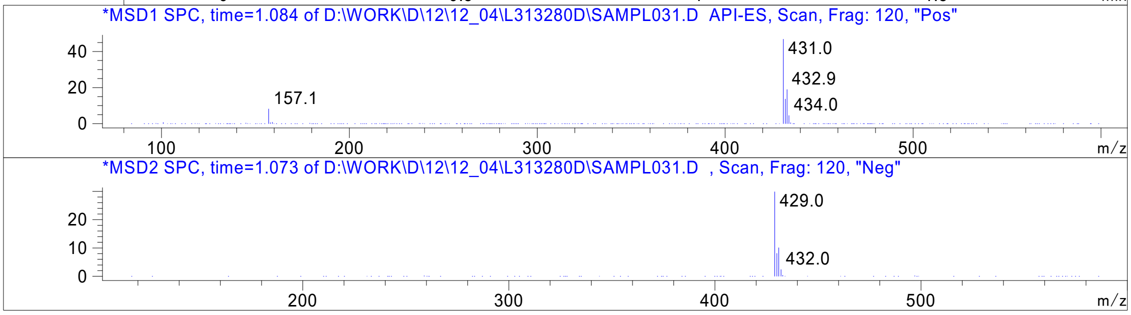


Inj.Date 12/4/2020 K -VL- Acq. Method C:\HPCHEM\-> ->

Z762205396_

MaxPeak: 100.00%
Ret_Time: 1.171 min

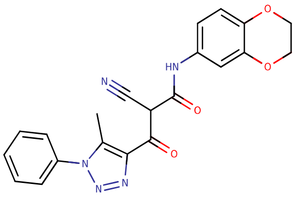


**Mol Wt** 403.39

**Exact Mass** 403.13

| # | Time | Area% |
|---|---|---|
| 1 | 1.171 | 100.00 |

S993131$5A

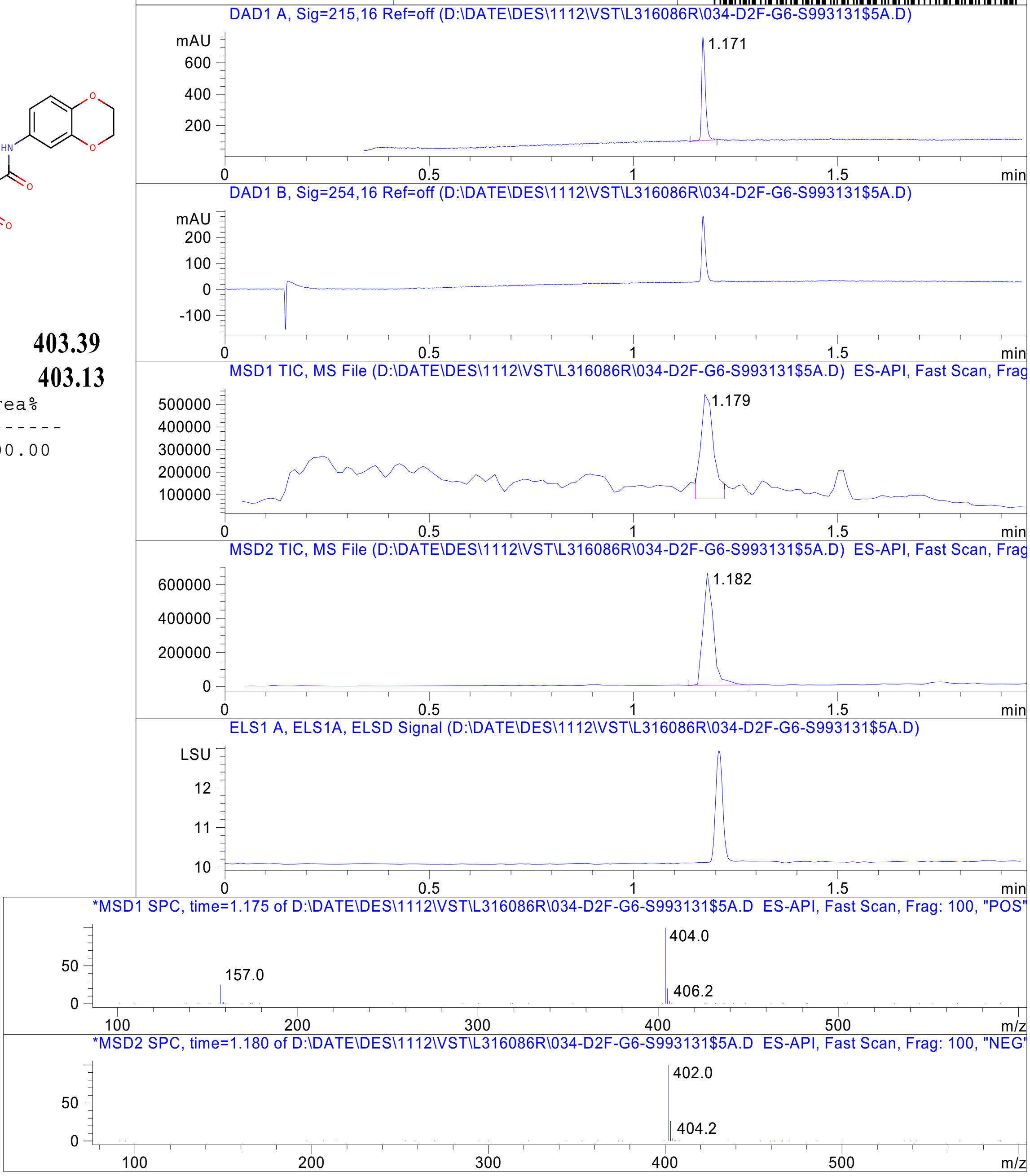


RT 1.179

RT 1.182

Inj.Date 12/11/2020 Y Acq. Method C:\Users\ -> ->

Z762217606_

MaxPeak: 94.15%
Ret_Time: 1.361 min

S993150$56

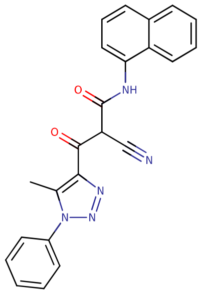


**Mol Wt** 395.41

**Exact Mass** 395.15

| # | Time | Area% |
|---|---|---|
| 1 | 0.791 | 1.38 |
| 2 | 0.924 | 1.34 |
| 3 | 1.361 | 94.15 |
| 4 | 1.401 | 3.13 |

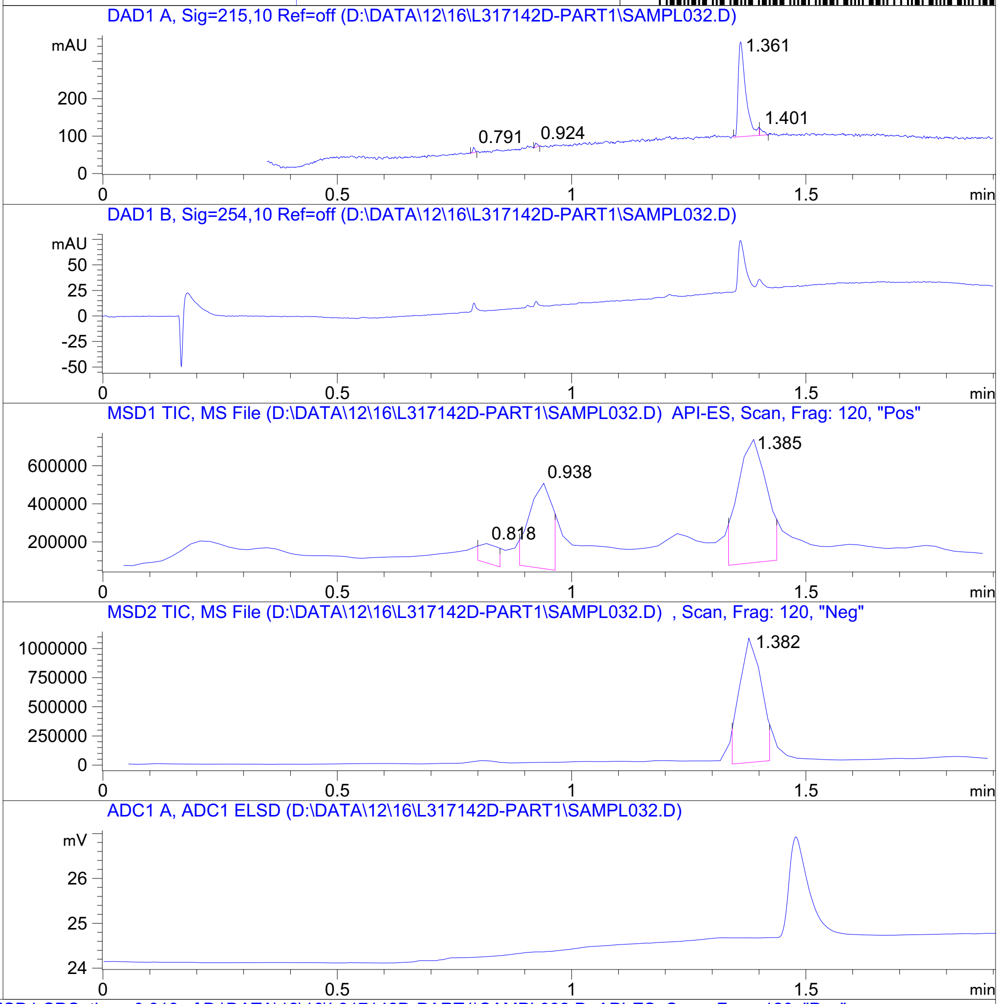


RT 0.818

RT 0.938

RT 1.385

RT 1.382

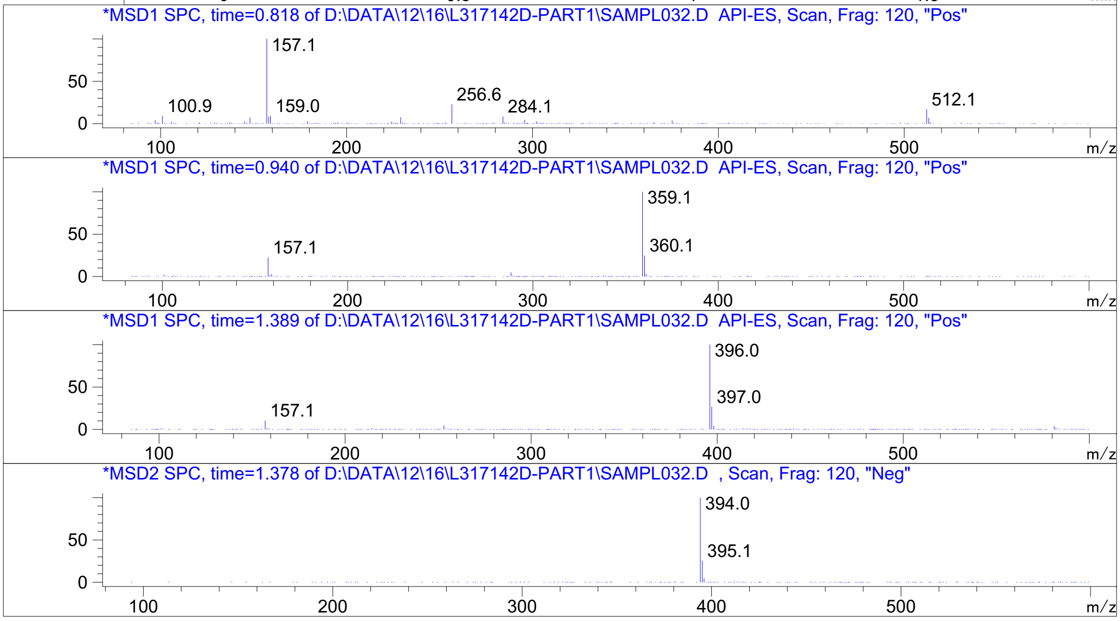


Inj.Date 12/16/2020 LB -VL- Acq. Method C:\HPCHEM\-> ->

Z762209308_

MaxPeak: 100.00%
Ret_Time: 1.246 min

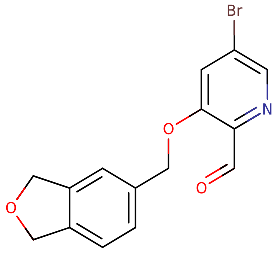

**Mol Wt** 334.16
**Exact Mass** 333.01

| # | Time | Area% |
|---|---|---|
| 1 | 1.246 | 100.00 |

# U211101$2

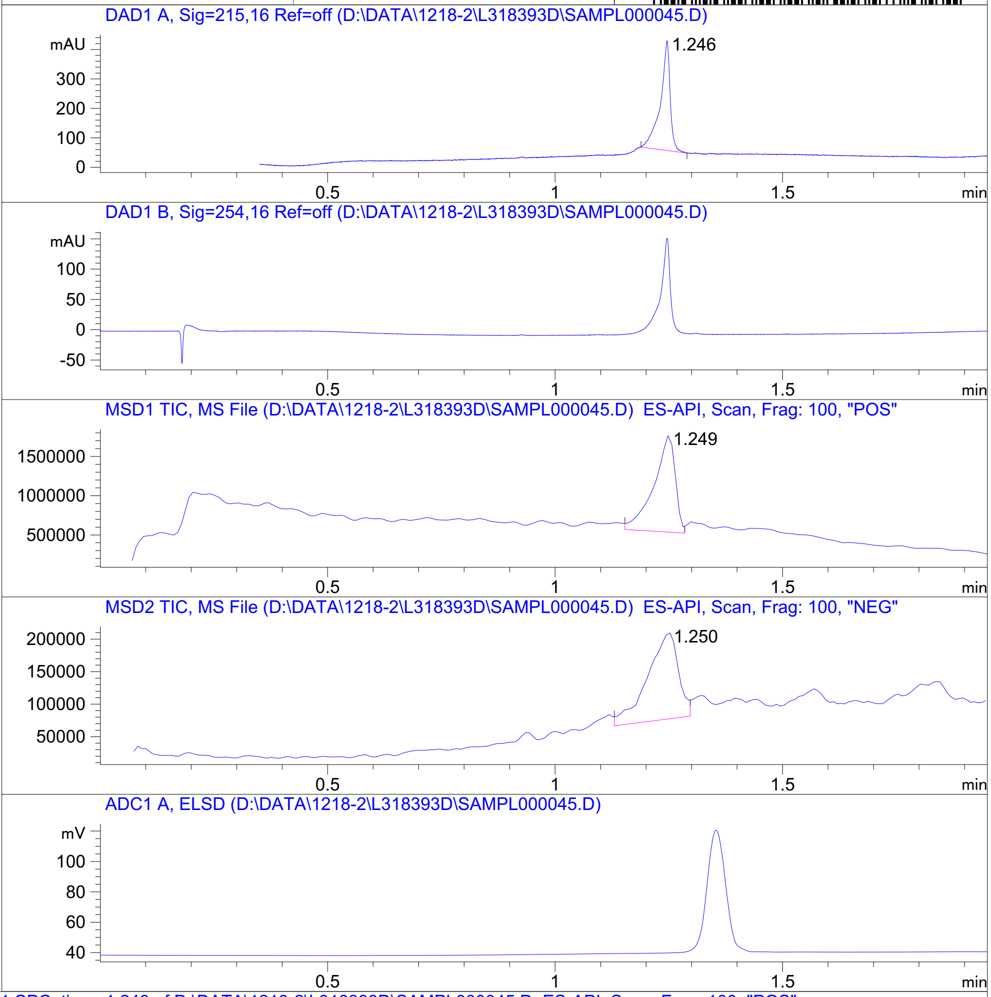


RT 1.249

RT 1.250

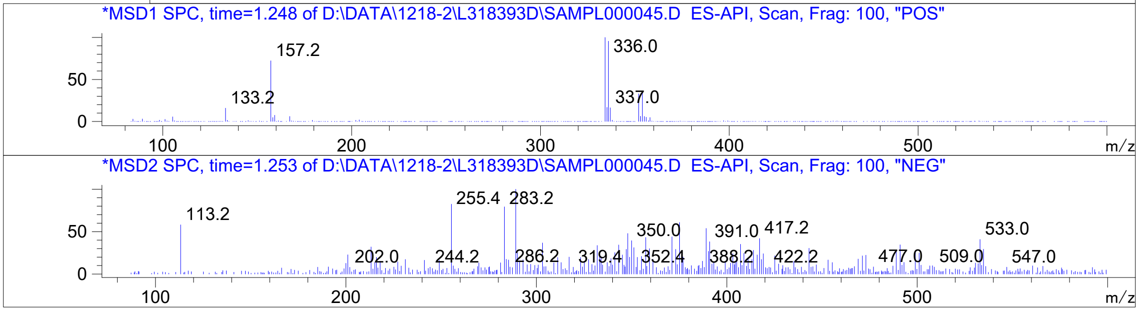




Z4497433897_

MaxPeak: 100.00%
Ret_Time: 3.558 min

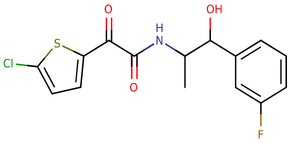


**Mol Wt** 341.78

**Exact Mass** 341.04

| # | Time | Area% |
|---|---|---|
| 1 | 3.558 | 100.00 |

U211102$9

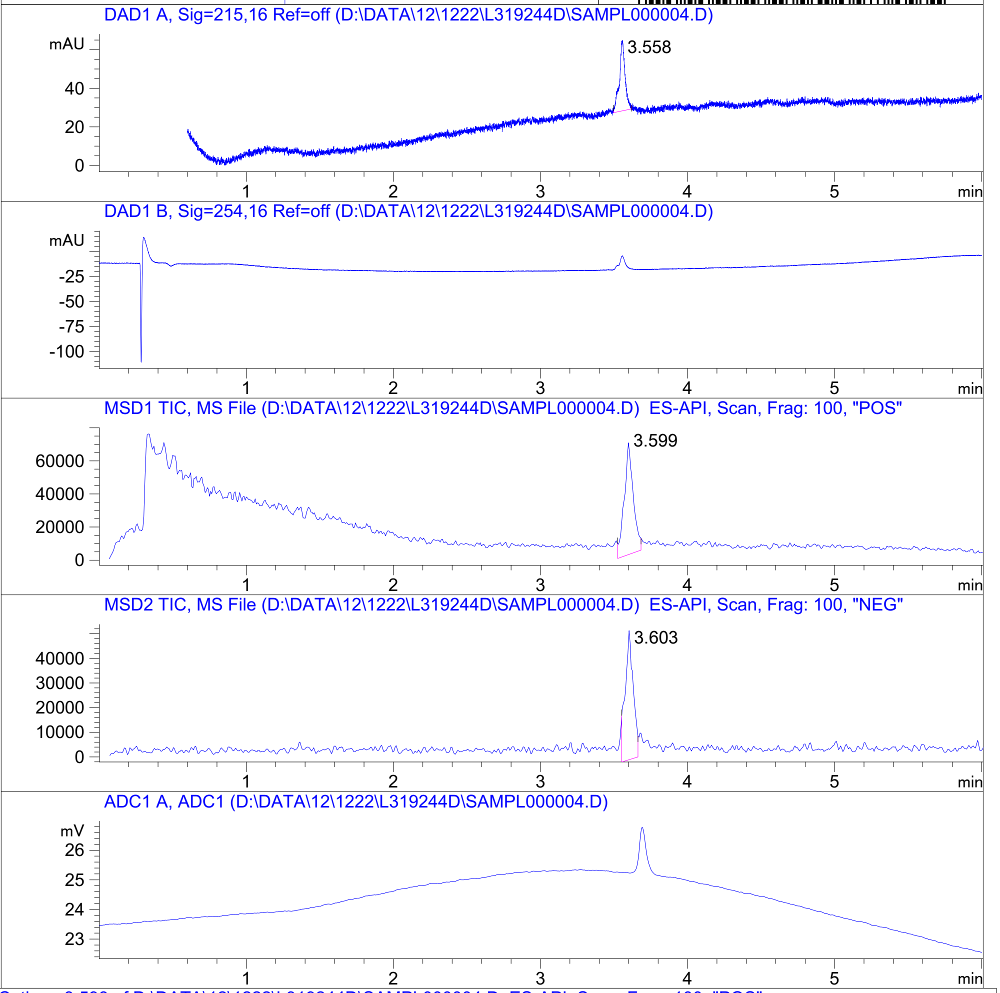


RT 3.599

RT 3.603

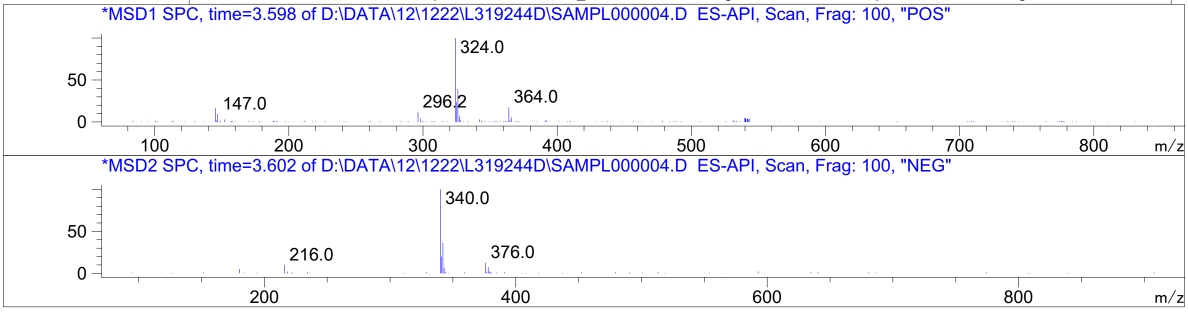


Inj.Date 12/21/2020 LT - 4 - Acq. Method C:\CHEM32\-> ->

Z1866628450_

U173081$3

MaxPeak: 96.04%
Ret_Time: 1.098 min

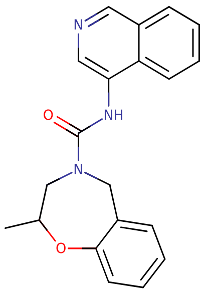

**Mol Wt** 333.38
**Exact Mass** 333.17

| # | Time | Area% |
|---|---|---|
| 1 | 0.815 | 1.16 |
| 2 | 1.098 | 96.04 |
| 3 | 1.542 | 2.79 |

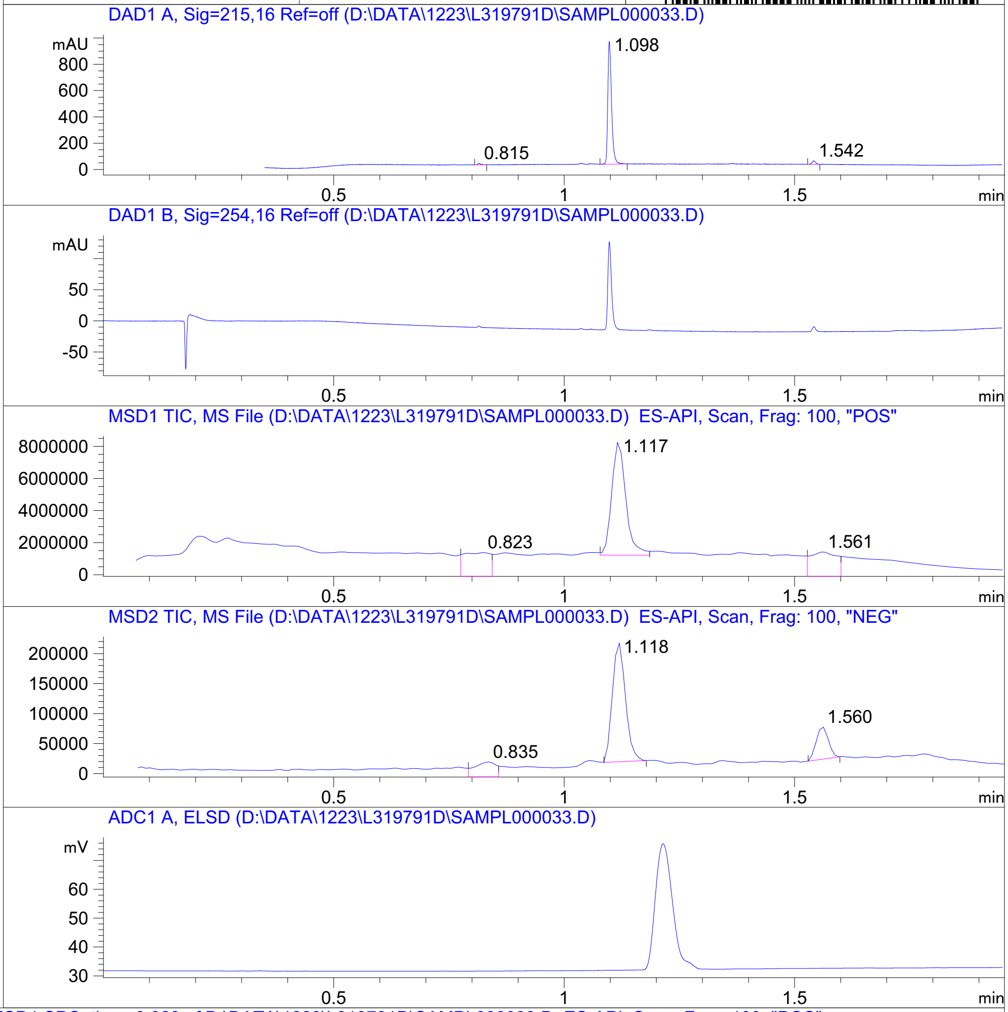


RT 0.823

RT 1.117

RT 1.561

RT 0.835

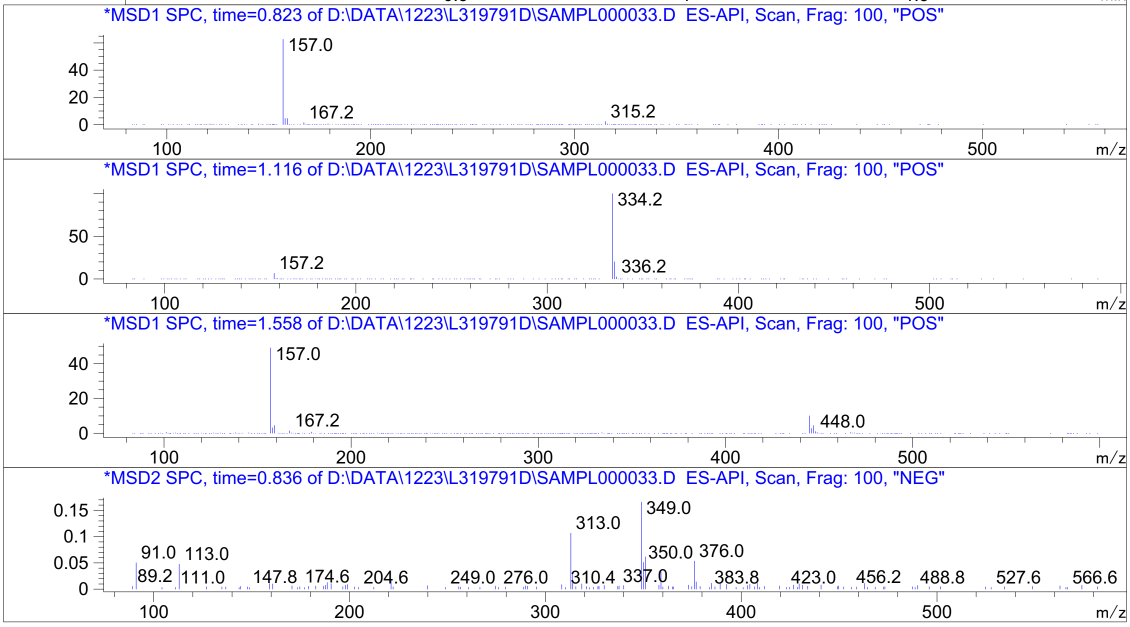




Z4872561628_

MaxPeak: 98.86%
Ret_Time: 0.874 min

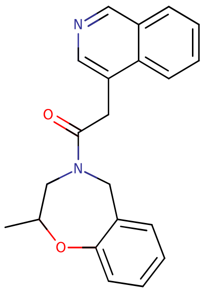

**Mol Wt** 332.4
**Exact Mass** 332.18

| # | Time | Area% |
|---|---|---|
| 1 | 0.874 | 98.86 |
| 2 | 1.182 | 1.14 |

# U172997$2

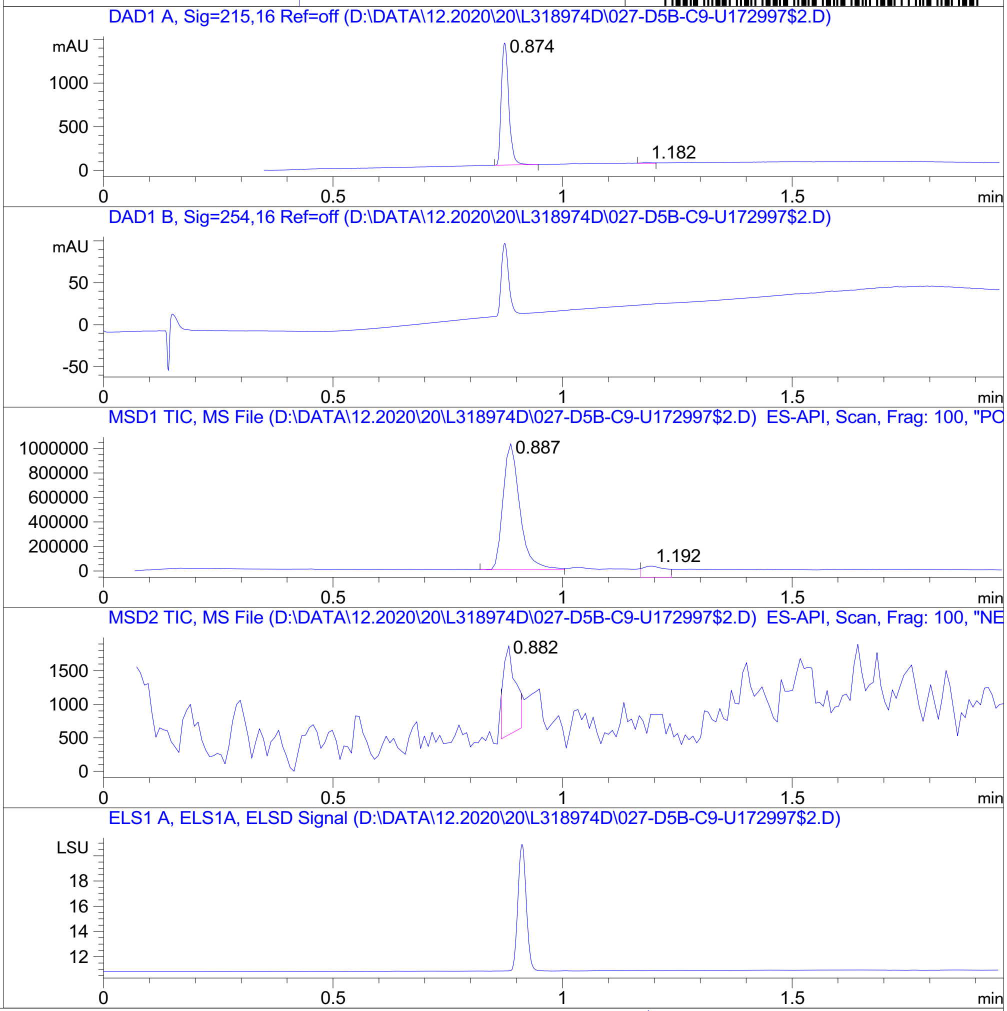


RT 0.887

RT 1.192

RT 0.882

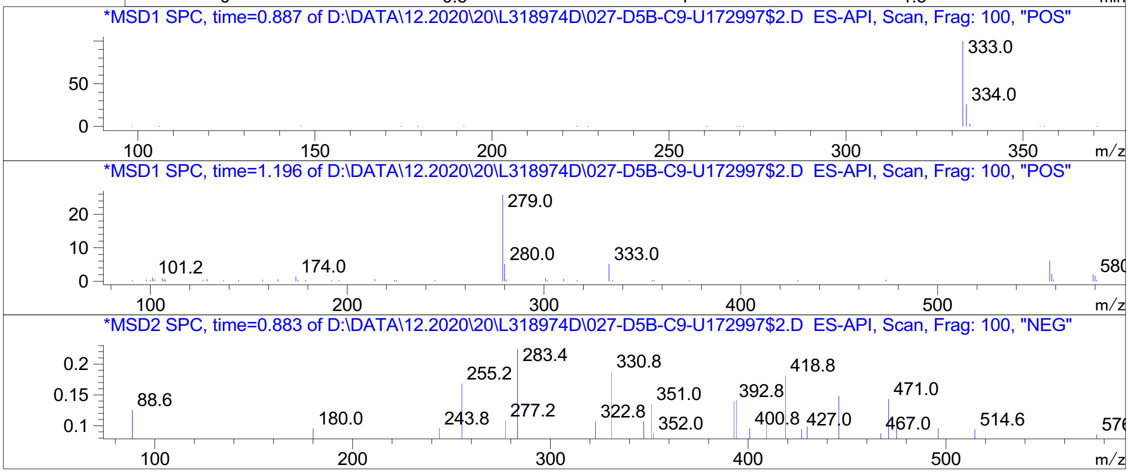


Inj.Date 12/19/2020 CH <invalid> 13 Acq. Method C:\Chem32\-> ->

Z4872561612_

MaxPeak: 100.00%
Ret_Time: 1.524 min

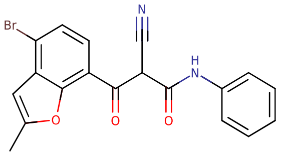

**Mol Wt** 397.22
**Exact Mass** 396.02

| # | Time | Area% |
|---|---|---|
| 1 | 1.524 | 100.00 |

U217010$A

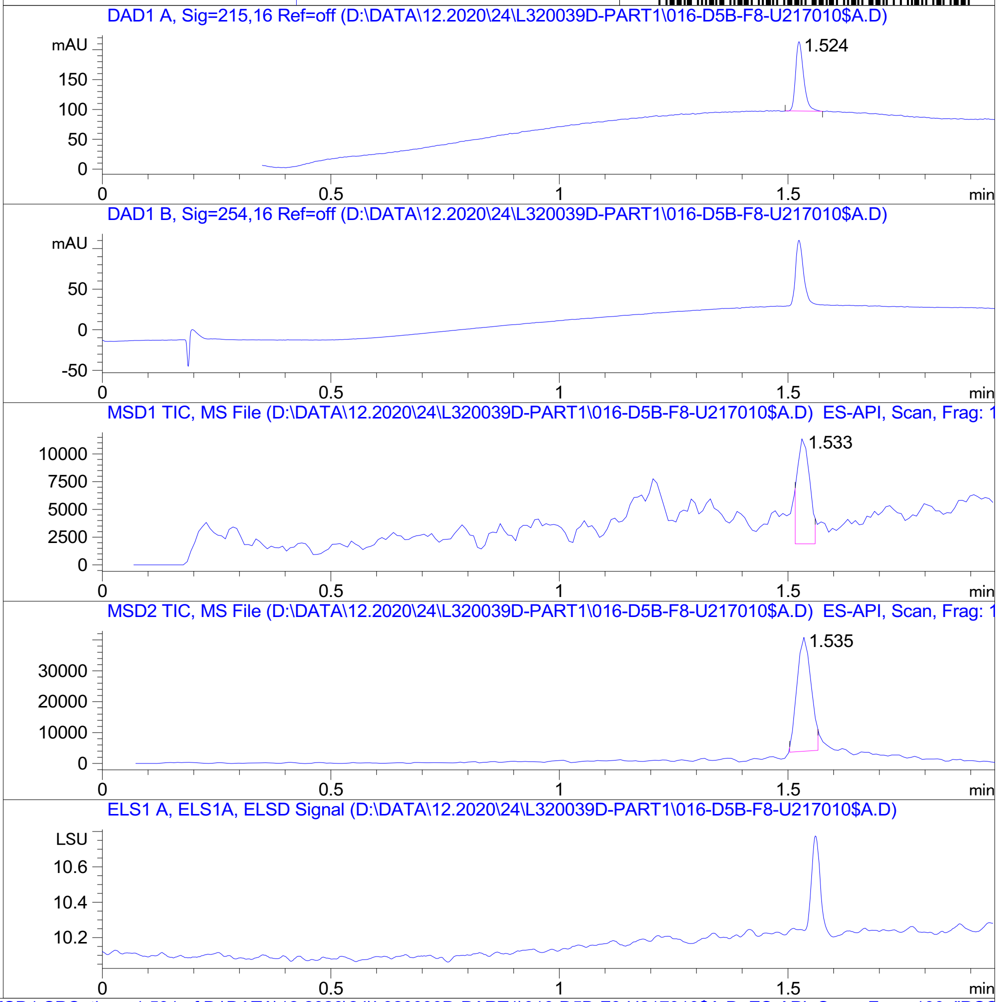


RT 1.533

RT 1.535

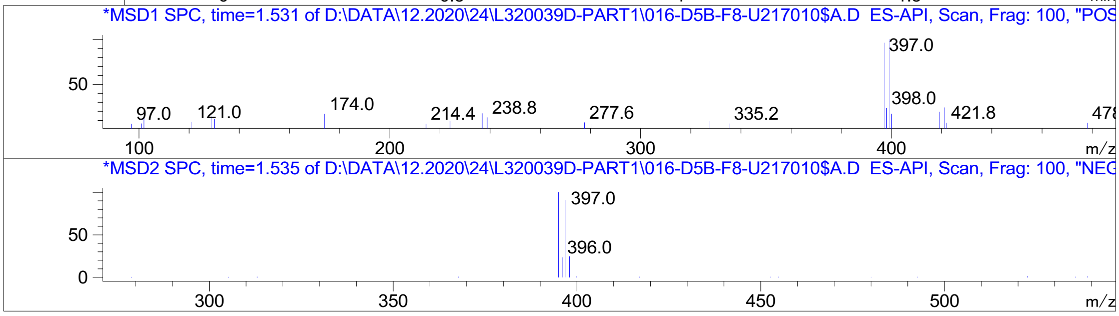


Inj.Date 12/24/2020 CH <invalid> -15- Acq. Method C:\Chem32\-> ->

Z1798873704_

MaxPeak: 100.00%
Ret_Time: 1.488 min

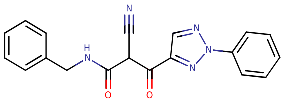


**Mol Wt** 345.36
**Exact Mass** 345.13

| # | Time | Area% |
|---|---|---|
| 1 | 1.488 | 100.00 |

# U253697$G

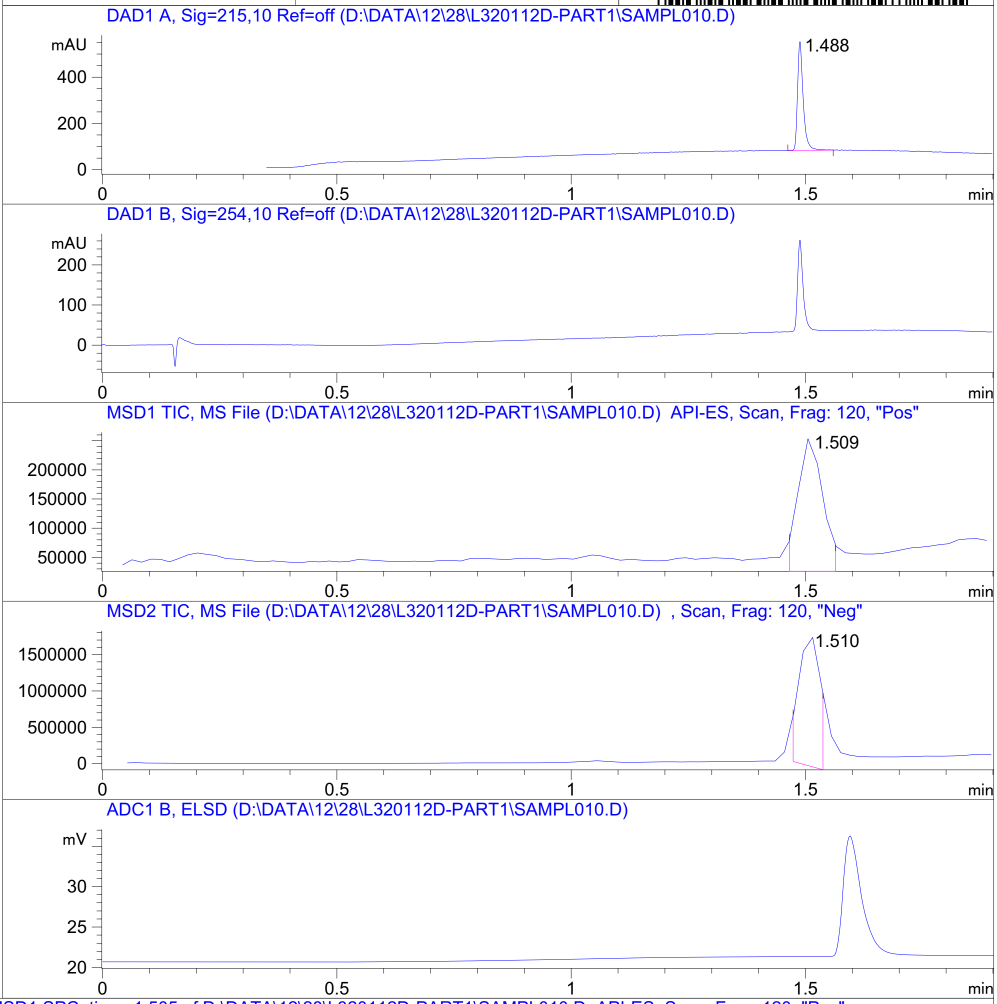


RT 1.509

RT 1.510

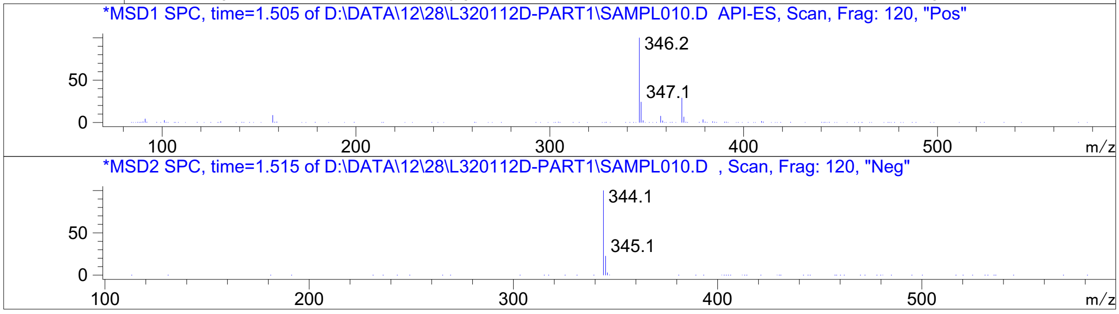


Inj.Date 12/25/2020 LB -SL- Acq. Method C:\HPCHEM\-> ->

Z638691128_

MaxPeak: 100.00%
Ret_Time: 1.307 min

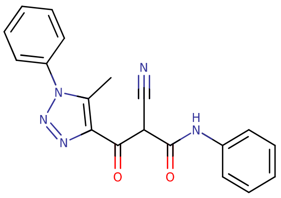

**Mol Wt** 345.36
**Exact Mass** 345.13

| # | Time | Area% |
|---|---|---|
| 1 | 1.307 | 100.00 |

7165188710

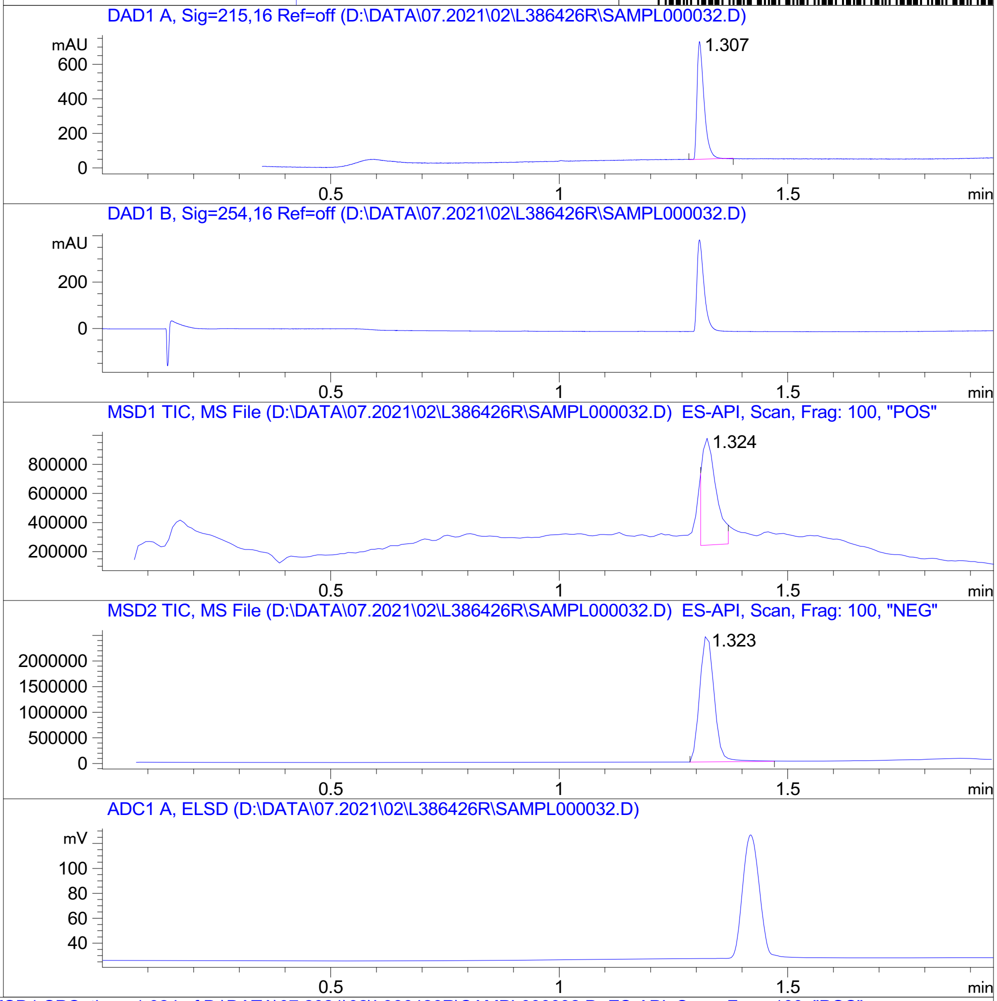


RT 1.324

RT 1.323

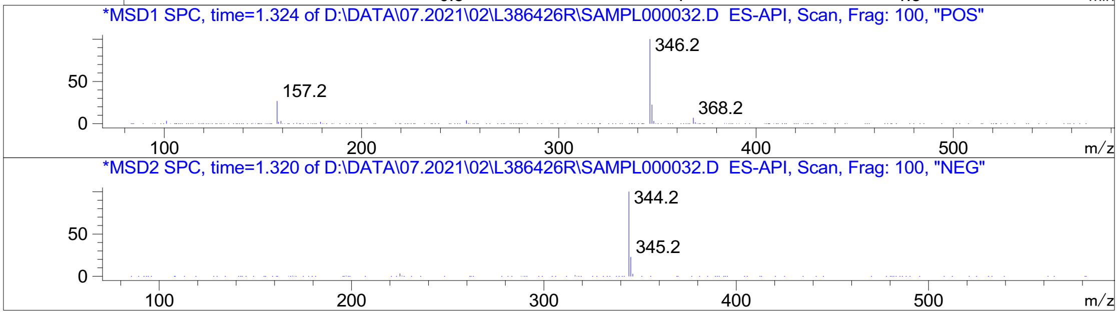


Inj.Date 7/2/2021 CH P2-C-01 -5- Acq. Method C:\CHEM32\-> ->

Z762207984_

MaxPeak: 97.01%
Ret_Time: 1.631 min

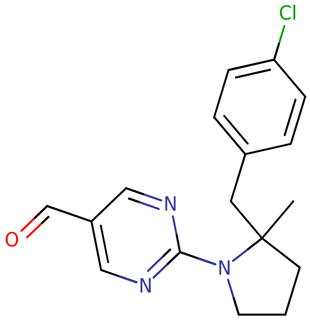

**Mol Wt** 315.8
**Exact Mass** 315.14

| # | Time | Area% |
|---|---|---|
| 1 | 0.942 | 2.99 |
| 2 | 1.631 | 97.01 |

U341347$1

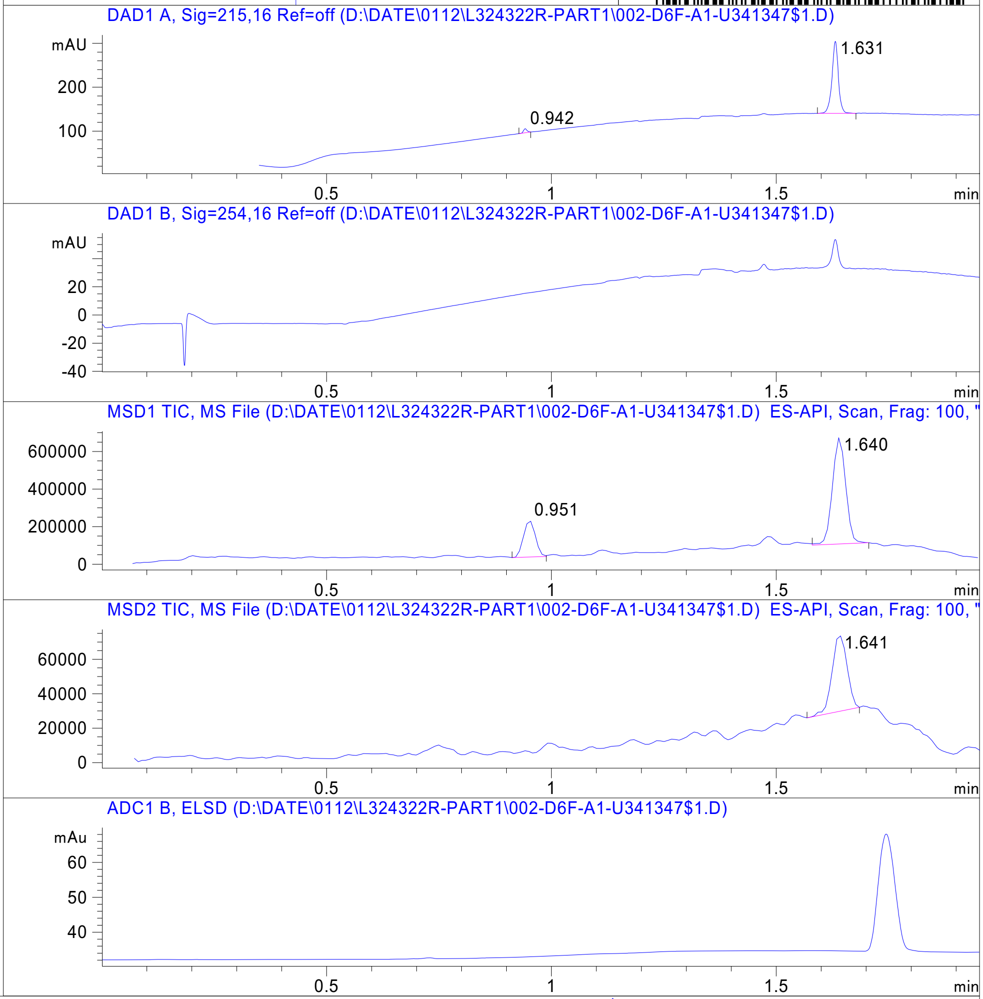


RT 0.951

RT 1.640

RT 1.641

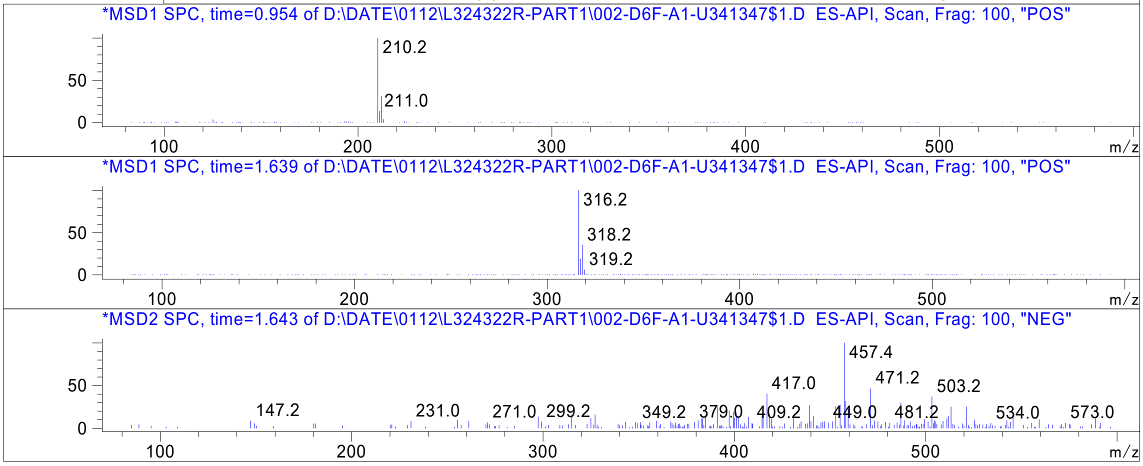


Inj.Date 1/12/2021 E -10- Acq. Method C:\Chem32\-> ->

Z4658847372_

MaxPeak: 100.00%
Ret_Time: 1.565 min

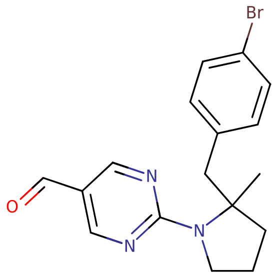

**Mol Wt 360.25**
**Exact Mass 359.09**

| # | Time | Area% |
|---|---|---|
| 1 | 1.565 | 100.00 |

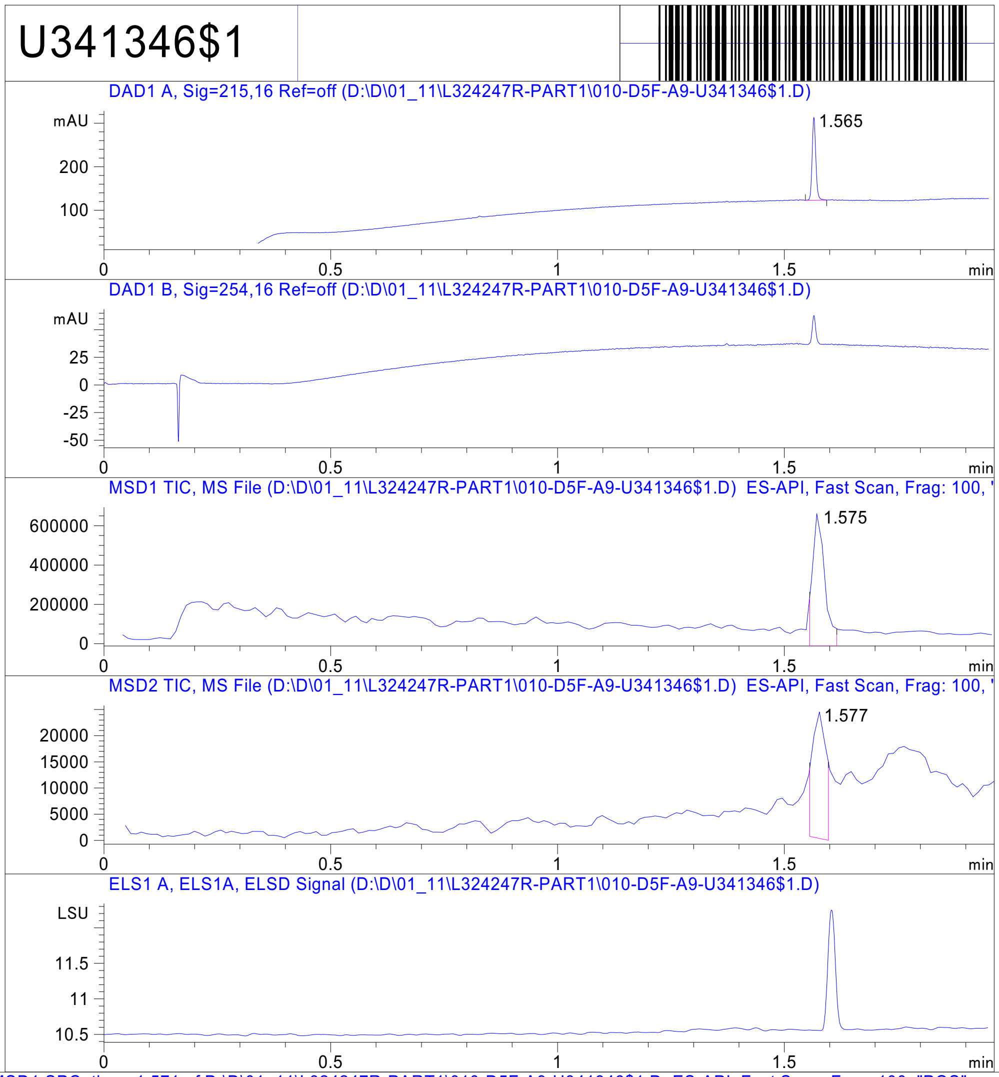


RT 1.575

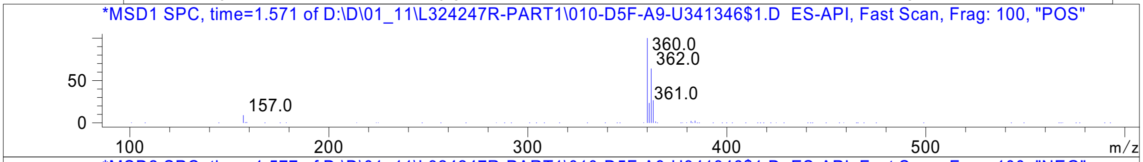


RT 1.577

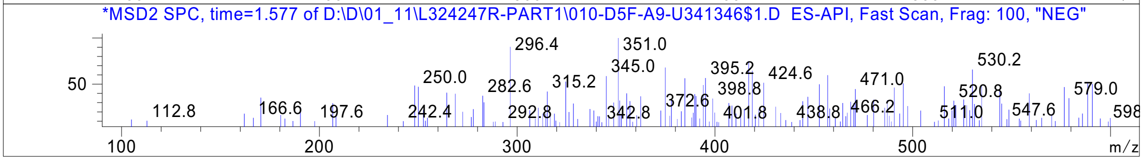


Inj.Date 1/11/2021 N Acq. Method C:\Users\ -> ->

Z4658847373_

MaxPeak: 100.00%
Ret_Time: 1.332 min

# U340878$35

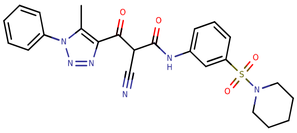

**Mol Wt** 492.55

**Exact Mass** 492.17

| # | Time | Area% |
|---|---|---|
| 1 | 1.332 | 100.00 |

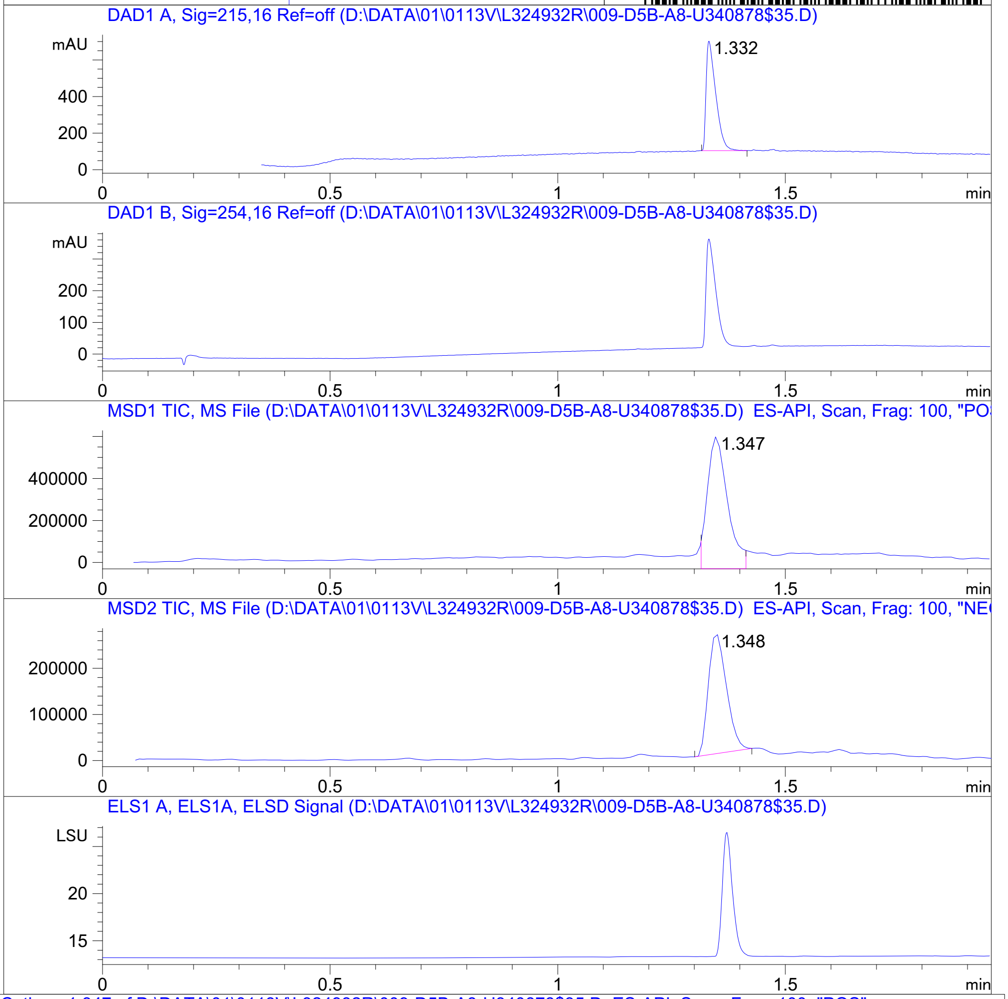


RT 1.347

RT 1.348

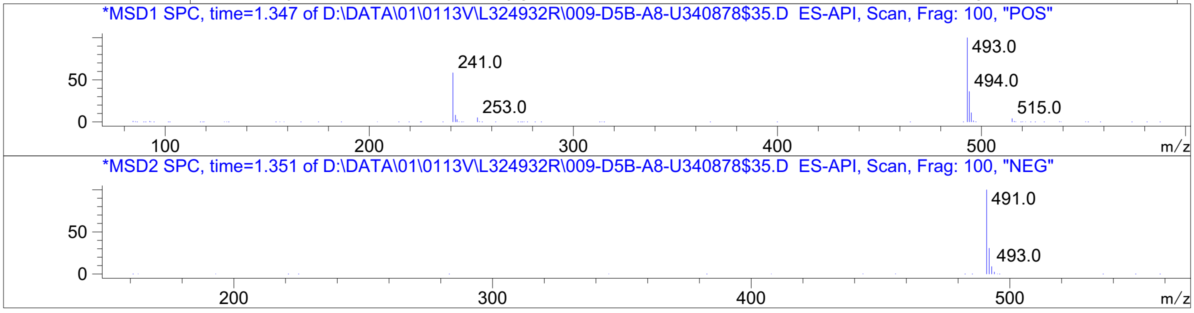




Z762213918_

# U804751$32

MaxPeak: 100.00%
Ret_Time: 1.166 min

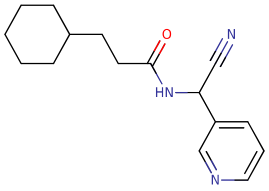


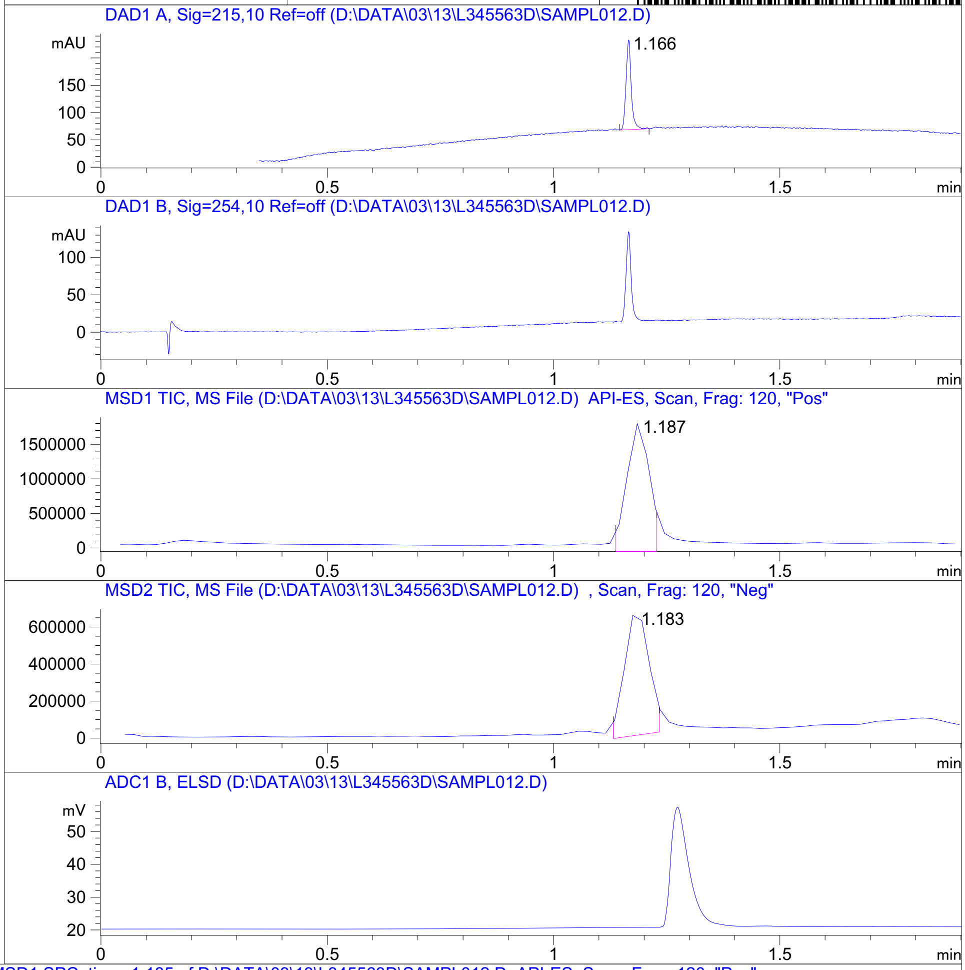


**Mol Wt** **271.36**
**Exact Mass** **271.2**

| # | Time | Area% |
|---|---|---|
| 1 | 1.166 | 100.00 |

RT 1.187

RT 1.183

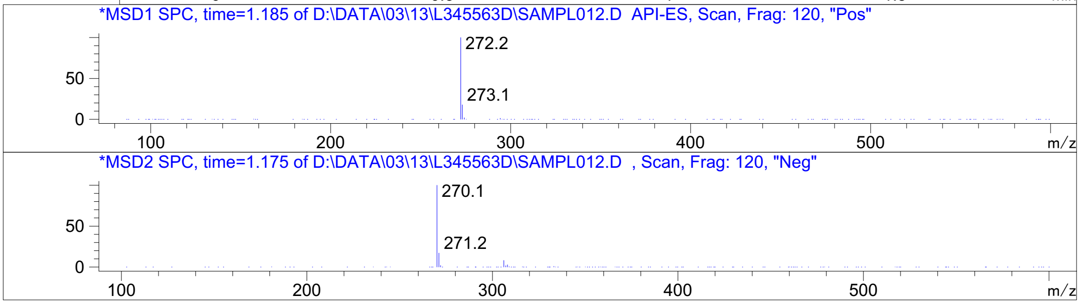


Inj.Date 3/13/2021 LB -SL- Acq. Method C:\HPCHEM\-> ->

Z4466780750_

MaxPeak:  90.21%
Ret_Time: 1.359 min

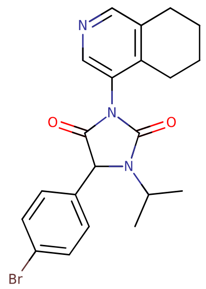

**Mol Wt** 428.32

**Exact Mass** 429.12

| # | Time | Area% |
|---|---|---|
| 1 | 0.936 | 9.79 |
| 2 | 1.359 | 90.21 |

U853197$1

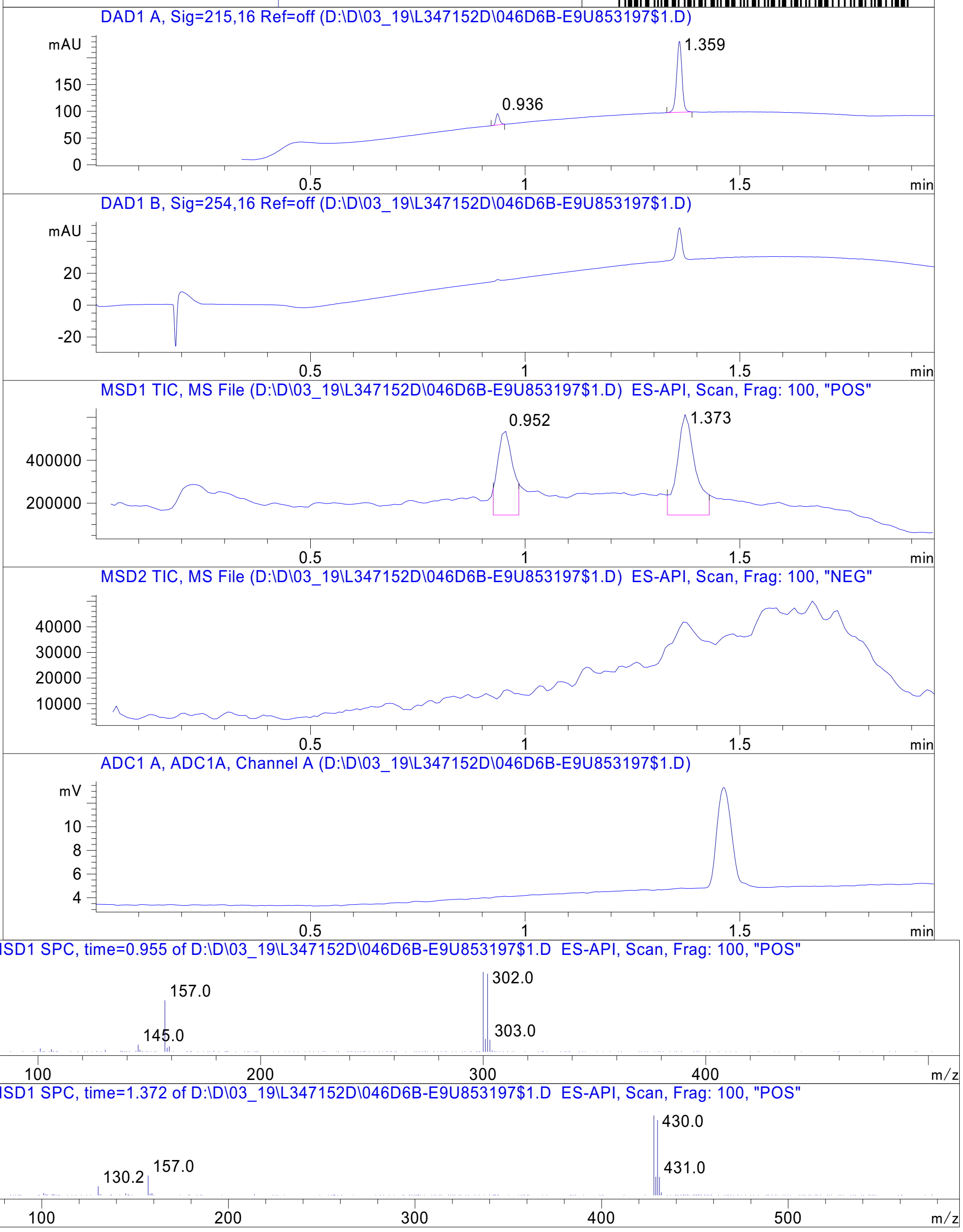


Inj.Date 3/19/2021    N    -11-    Acq. Method C:\Chem32\->  ->

Z4929616252_

MaxPeak: 100.00%
Ret_Time: 1.211 min

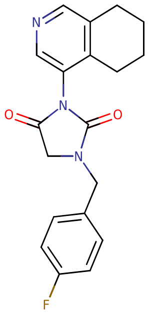

**Mol Wt** 339.36
**Exact Mass** 339.16

| # | Time | Area% |
|---|---|---|
| 1 | 1.211 | 100.00 |

U853195$2

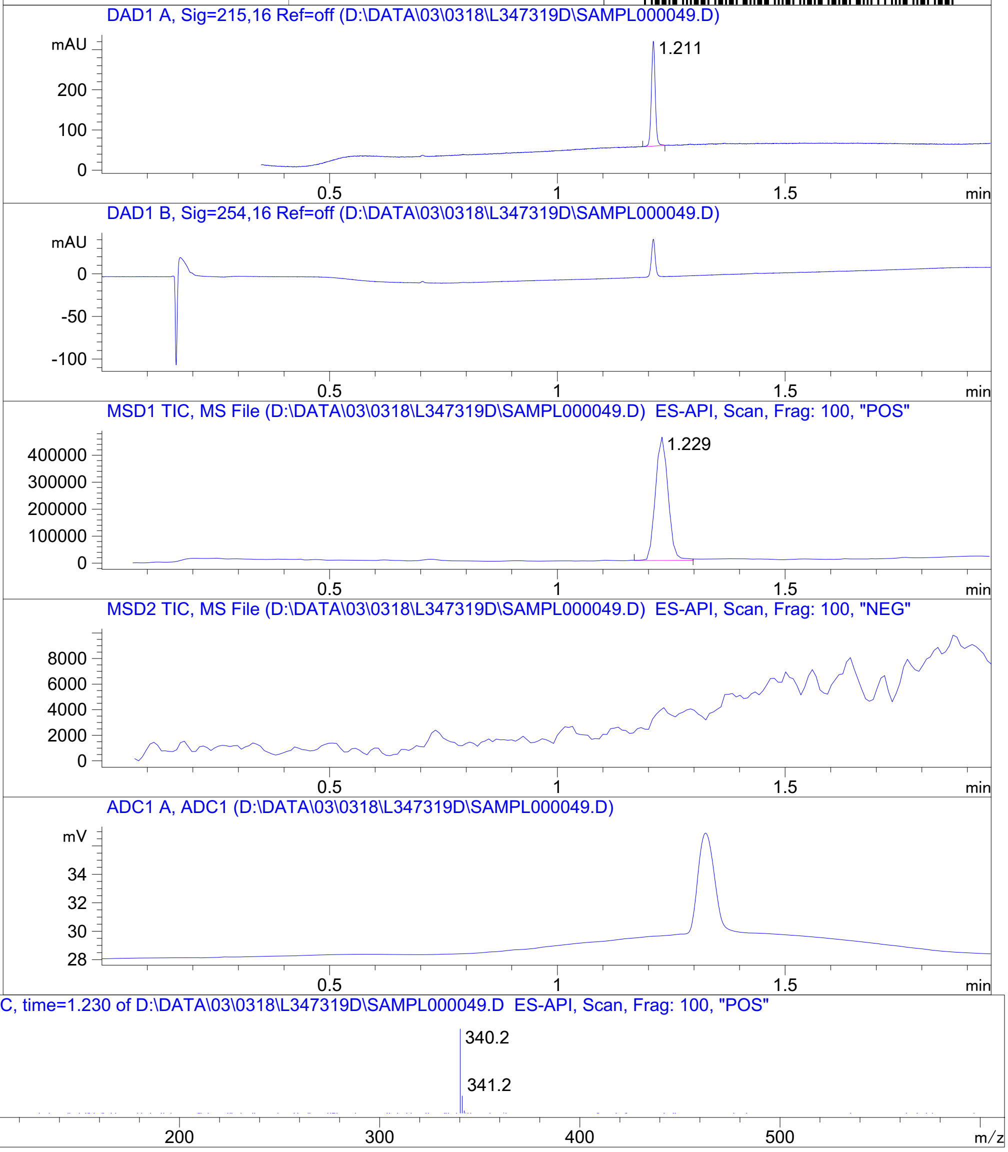


RT 1.229

Inj.Date 3/18/2021 LT - 4 - Acq. Method C:\CHEM32\-> ->

Z4929616328_

MaxPeak: 98.91%
Ret_Time: 1.087 min

# U853206$2

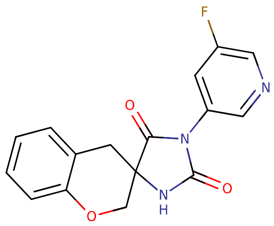


**Mol Wt** 313.28
**Exact Mass** 313.09

| # | Time | Area% |
|---|---|---|
| 1 | 0.961 | 1.09 |
| 2 | 1.087 | 98.91 |

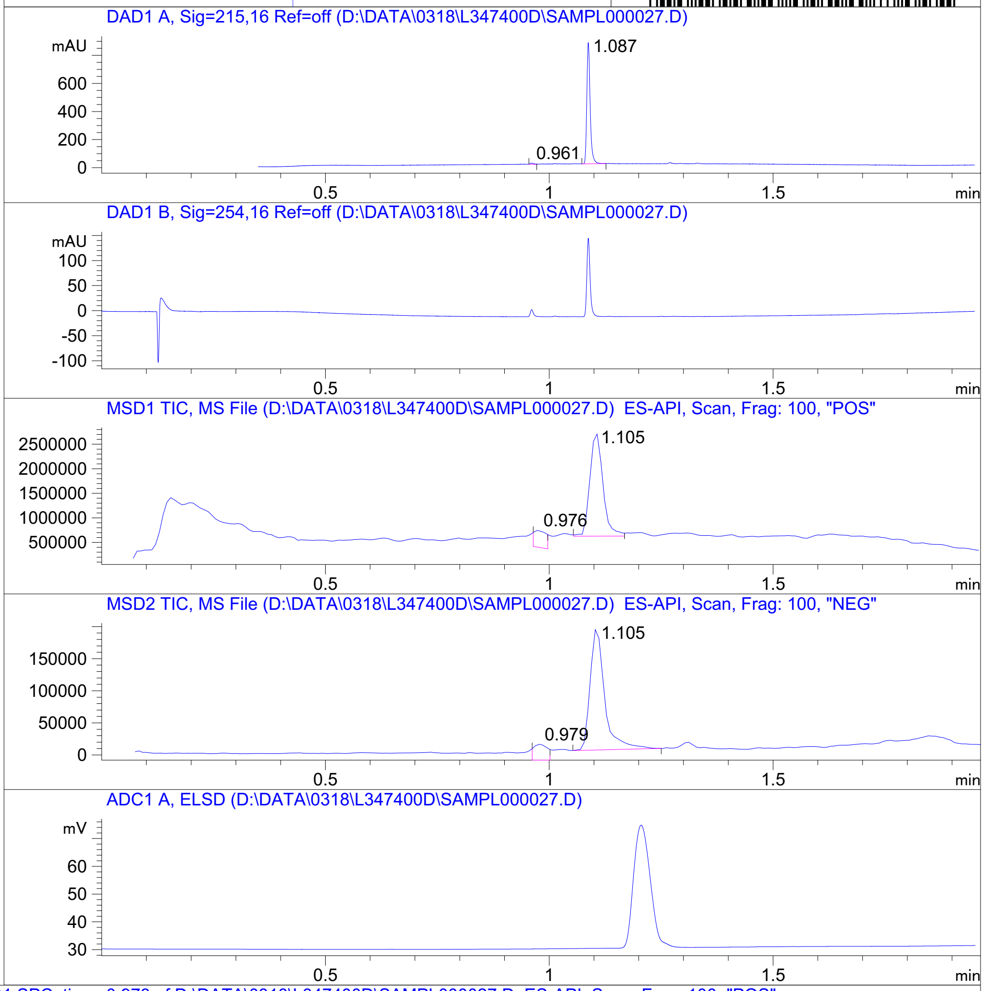


RT 0.976

RT 1.105

RT 0.979

RT 1.105

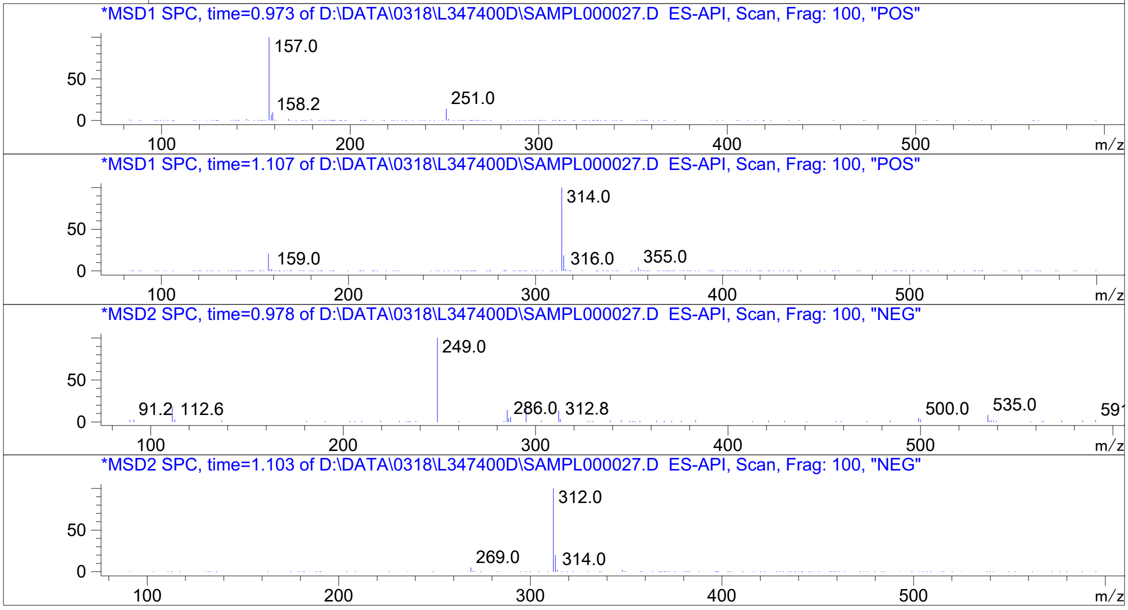




Z4739648085_

MaxPeak: 100.00%
Ret_Time: 1.520 min

# 7158466344

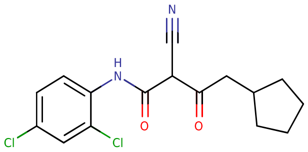


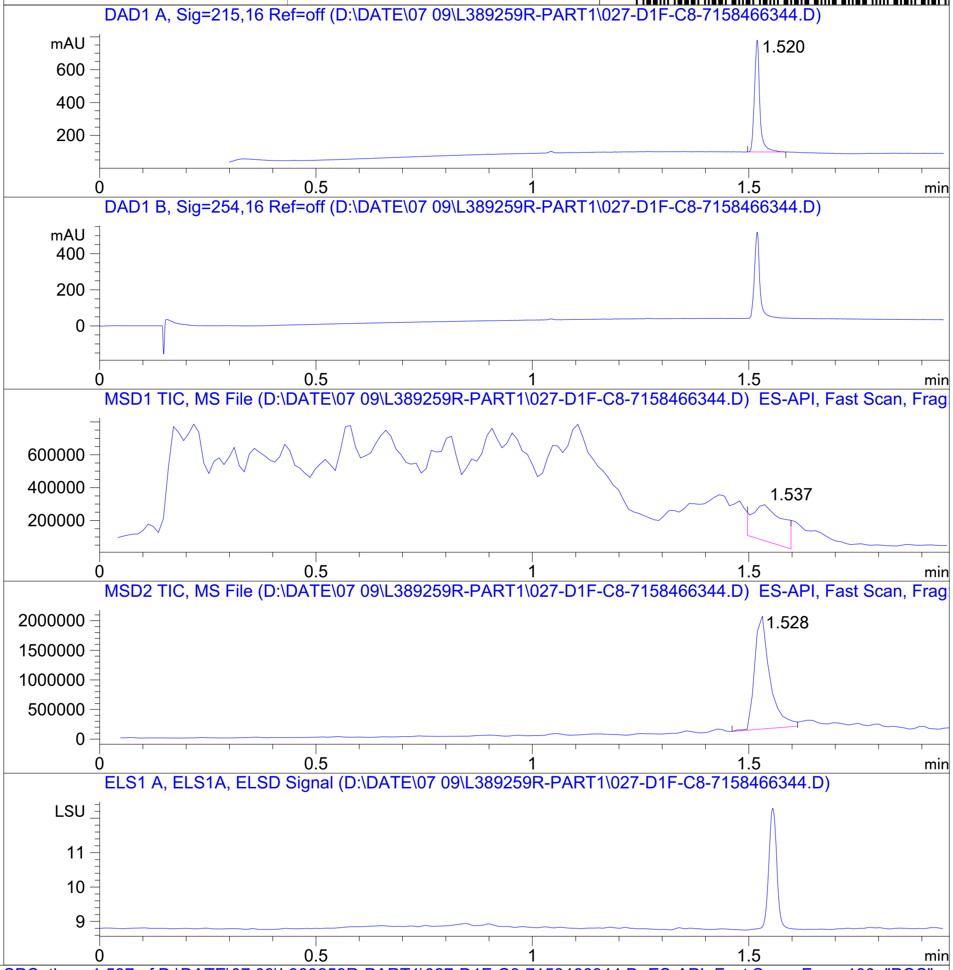


**Mol Wt** 339.22
**Exact Mass** 338.08

| # | Time | Area% |
|---|---|---|
| 1 | 1.520 | 100.00 |

RT 1.537

RT 1.528

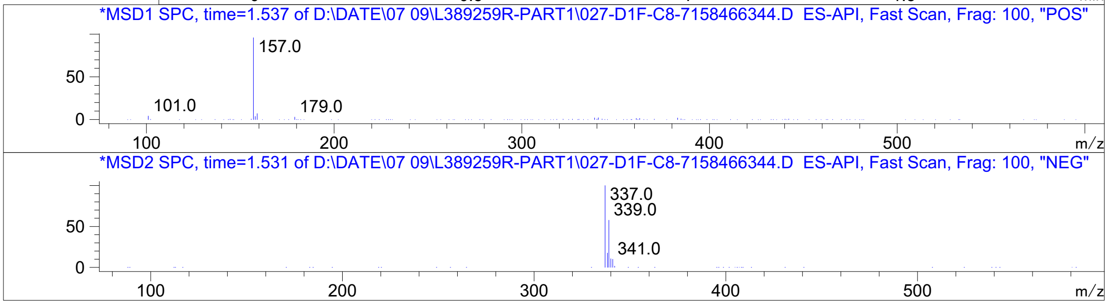


Inj.Date 7/9/2021 E Acq. Method C:\Users\ -> ->

Z639519658_

```
MaxPeak:  98.16%
Ret_Time: 1.052 min
```

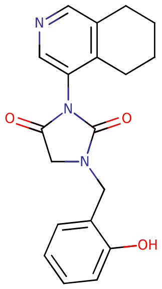

**Mol Wt** 337.37

**Exact Mass** 337.16

| # | Time | Area% |
|---|---|---|
| 1 | 0.725 | 1.84 |
| 2 | 1.052 | 98.16 |

U853196$2

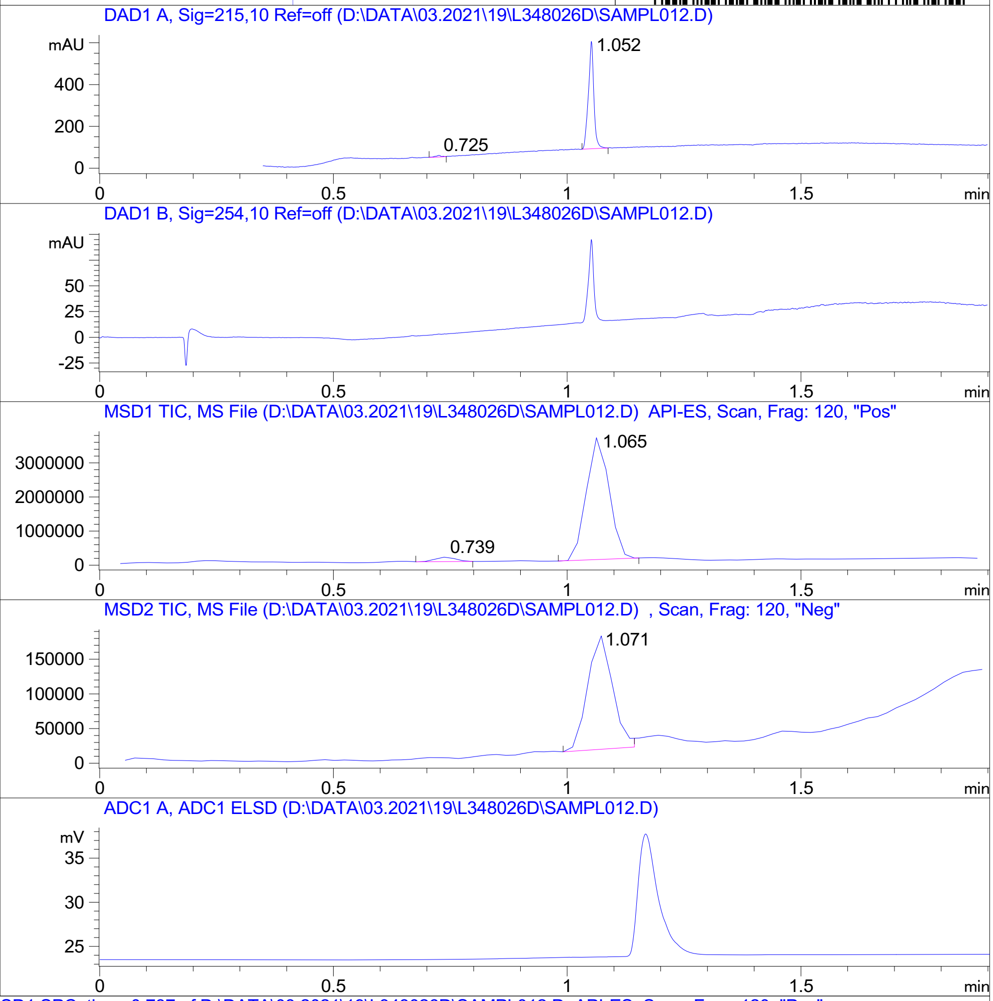


RT 0.739

RT 1.065

RT 1.071

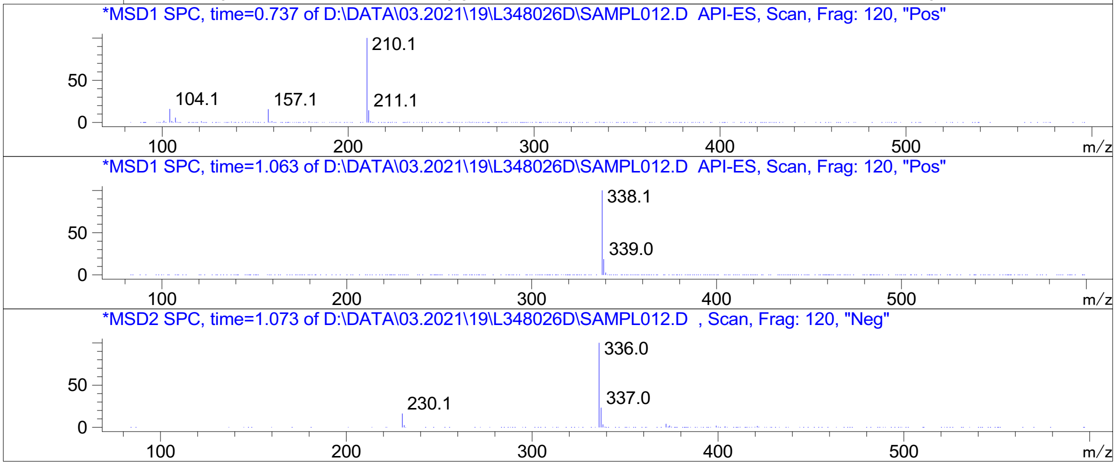


Inj.Date 3/19/2021 CH P2-B-03 -VL- Acq. Method C:\HPCHEM\-> ->

Z4929616577_

MaxPeak: 92.69%
Ret_Time: 1.096 min

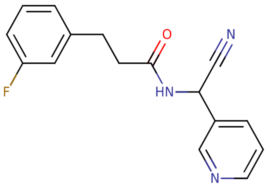


**Mol Wt** 283.3
**Exact Mass** 283.13

| # | Time | Area% |
|---|---|---|
| 1 | 1.096 | 92.69 |
| 2 | 1.183 | 7.31 |

U812291$23

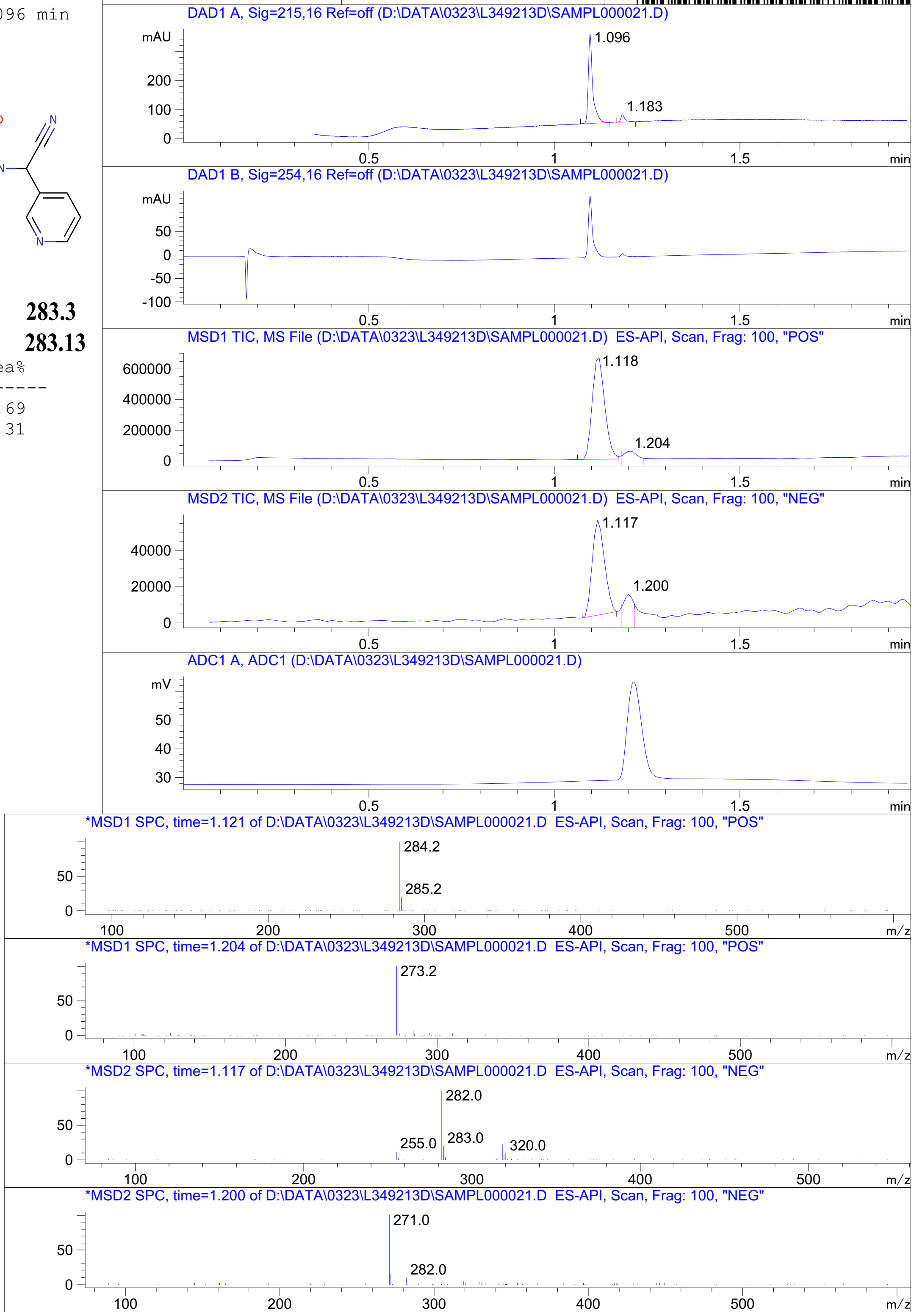




Z4934489626_

MaxPeak: 95.50%
Ret_Time: 1.473 min

# U853208$2

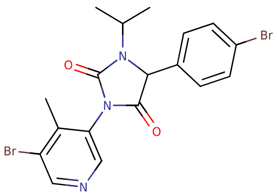


**Mol Wt** 467.15

**Exact Mass** 466.99

| # | Time | Area% |
|---|---|---|
| 1 | 1.452 | 4.50 |
| 2 | 1.473 | 95.50 |

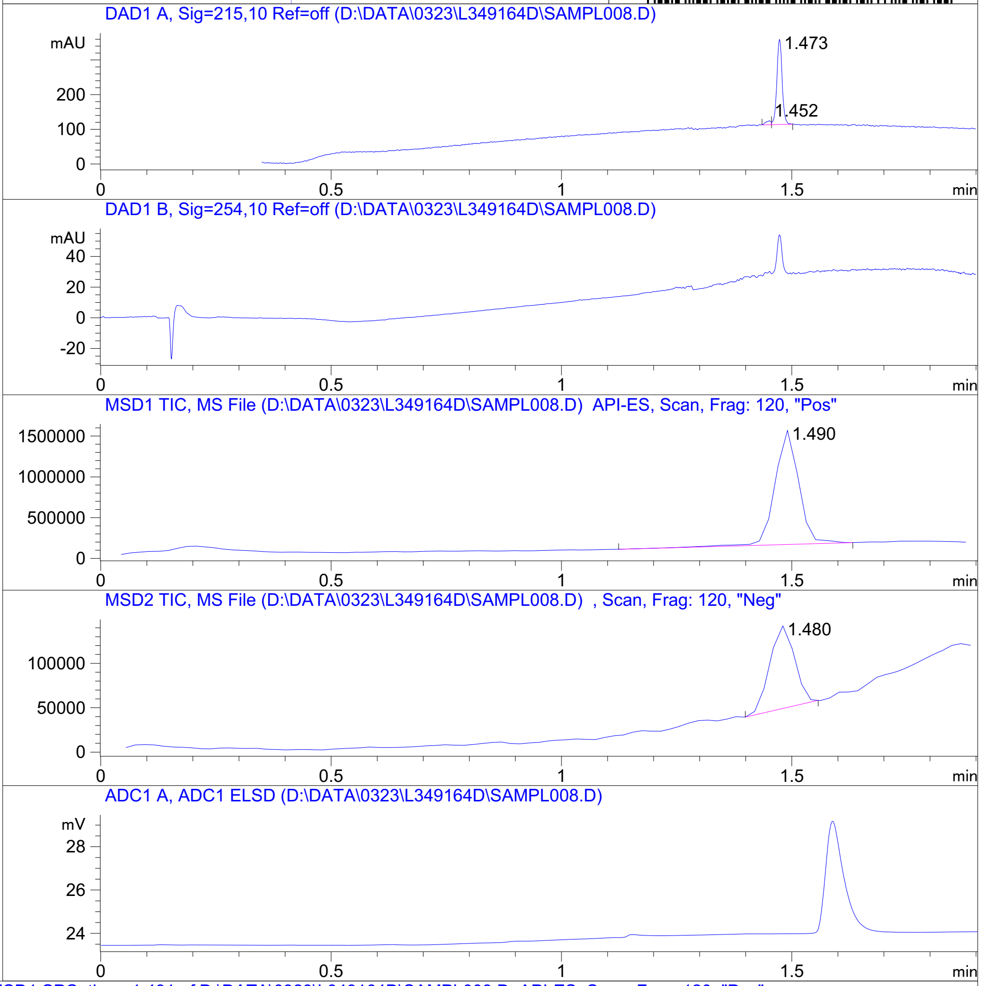


RT 1.490

RT 1.480

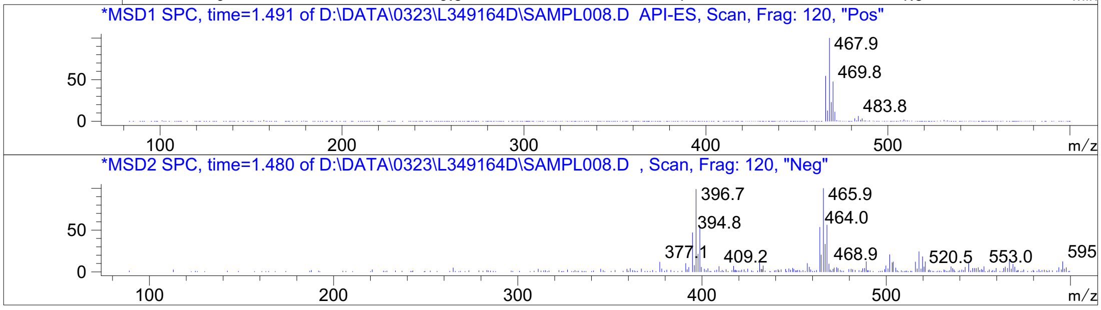


Inj.Date 3/23/2021 OA -VL- Acq. Method C:\HPCHEM\-> ->

Z4927215427_

MaxPeak: 83.37%
Ret_Time: 1.485 min

# U950663$1

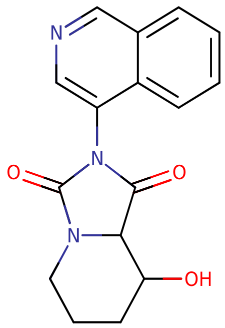


**Mol Wt** 297.31

**Exact Mass** 297.12

| # | Time | Area% |
|---|---|---|
| 1 | 1.485 | 83.37 |
| 2 | 1.541 | 11.66 |
| 3 | 2.085 | 4.97 |

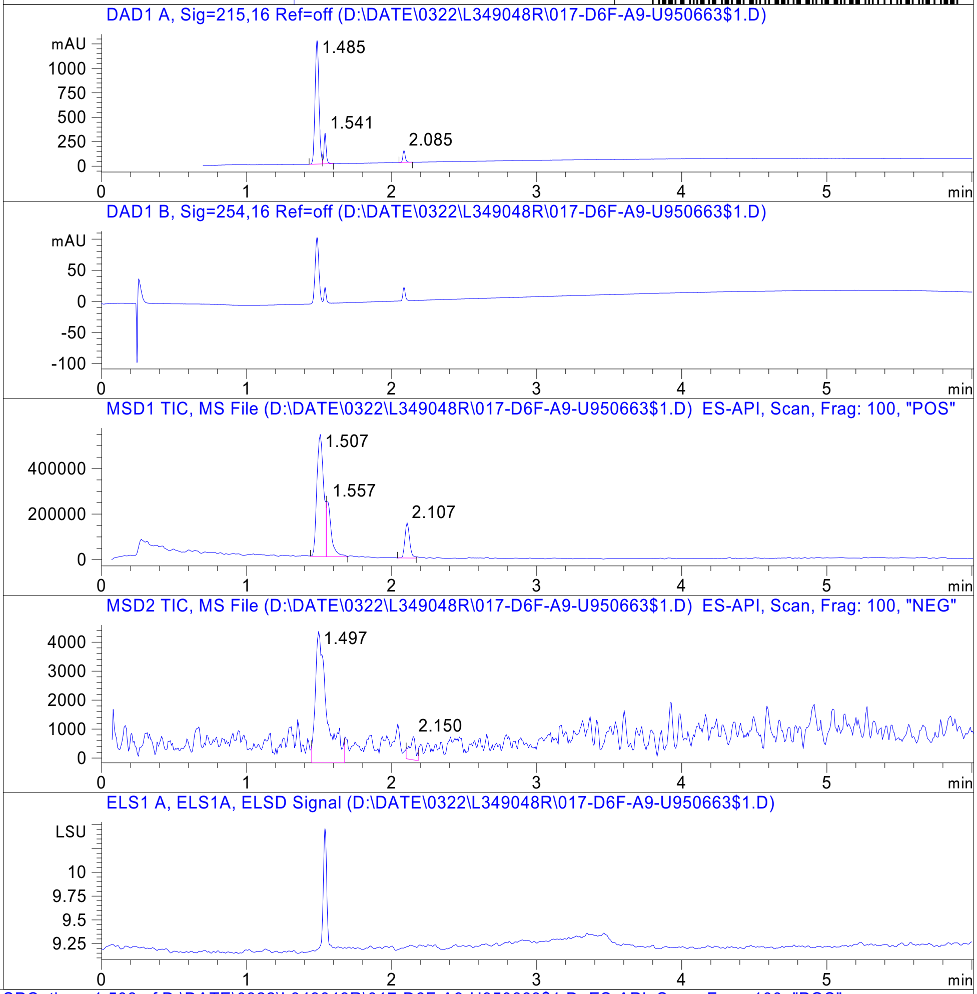


RT 1.507

RT 1.557

RT 2.107

RT 1.497

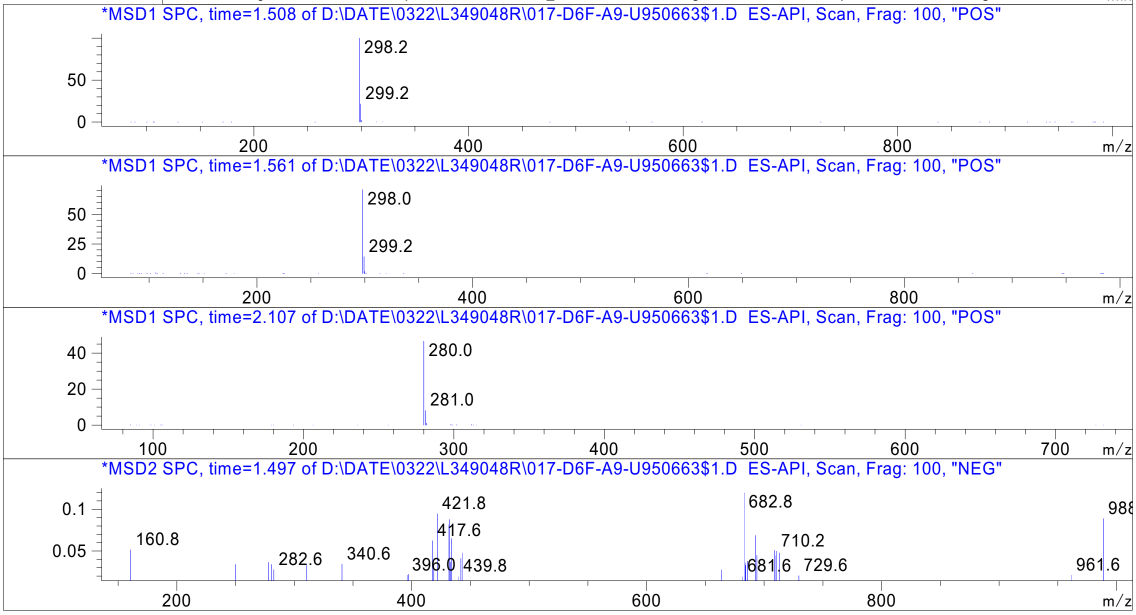


Inj.Date 3/22/2021 E 13 Acq. Method C:\Chem32\-> ->

Z4929616088_

MaxPeak: 100.00%
Ret_Time: 3.550 min

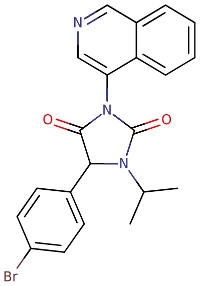


**Mol Wt** 424.29
**Exact Mass** 425.08

| # | Time | Area% |
|---|---|---|
| 1 | 3.550 | 100.00 |

# U950662$2

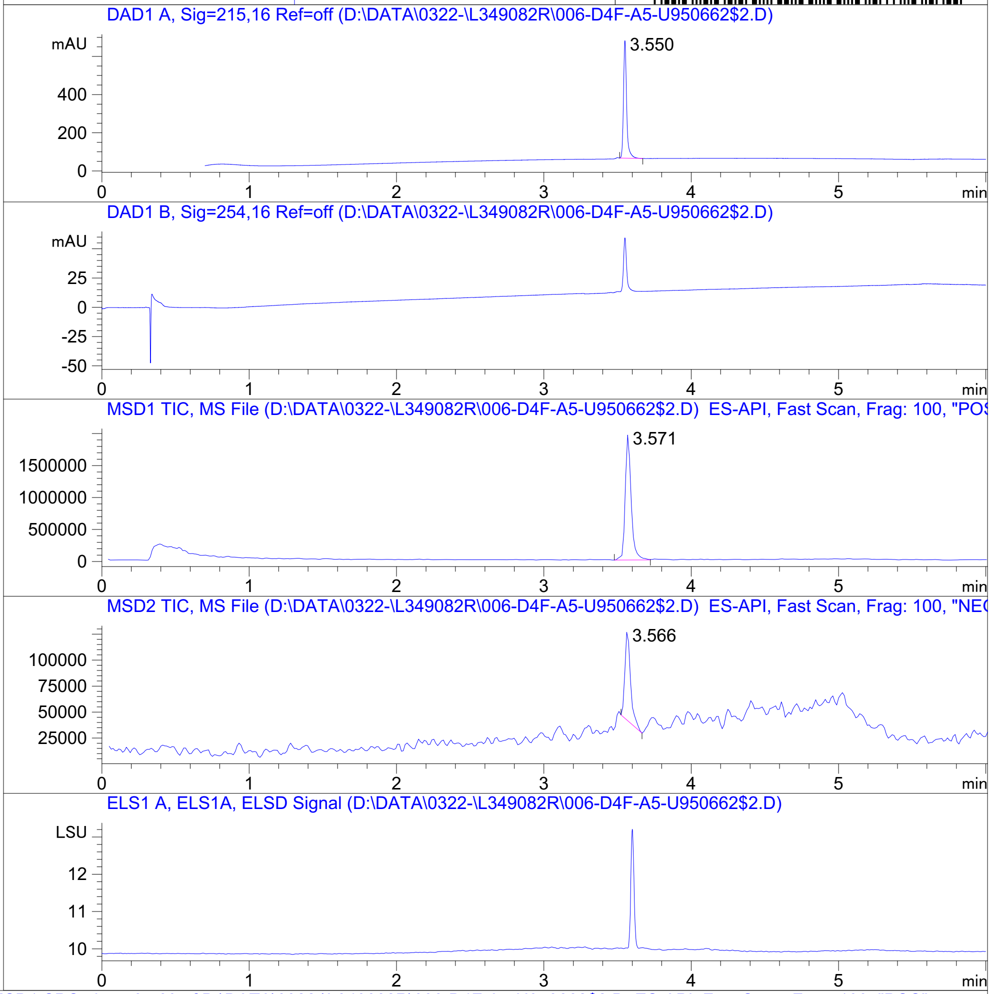


RT 3.571

RT 3.566

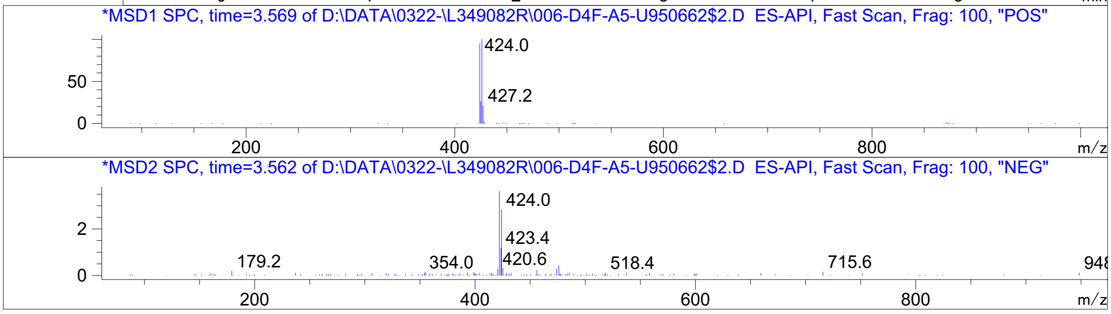


Inj.Date 3/22/2021 OA Acq. Method C:\Users\ -> ->

Z4925123630_

MaxPeak: 100.00%
Ret_Time: 0.937 min

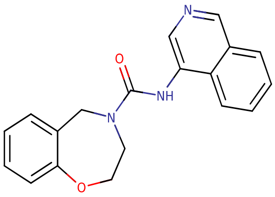


**Mol Wt** 319.36

**Exact Mass** 319.15

| # | Time | Area% |
|---|---|---|
| 1 | 0.937 | 100.00 |

U689907$2

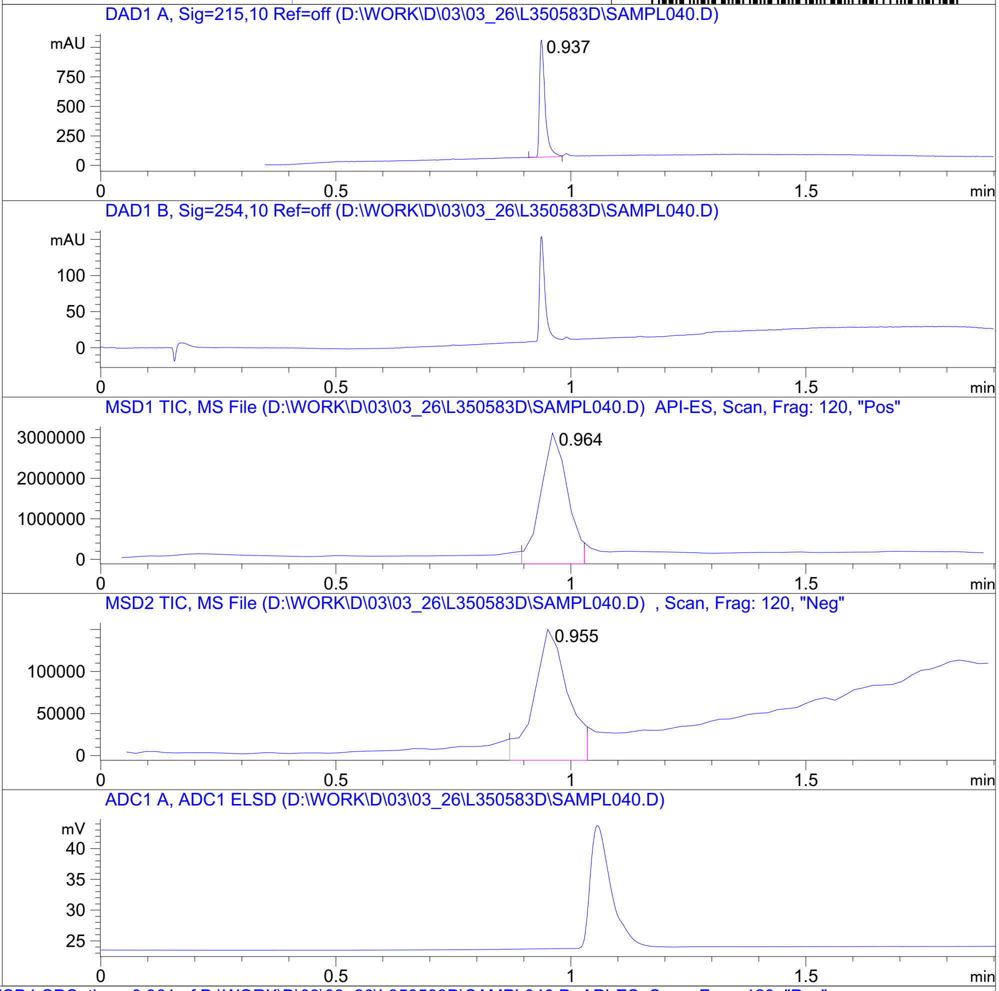


RT 0.964

RT 0.955

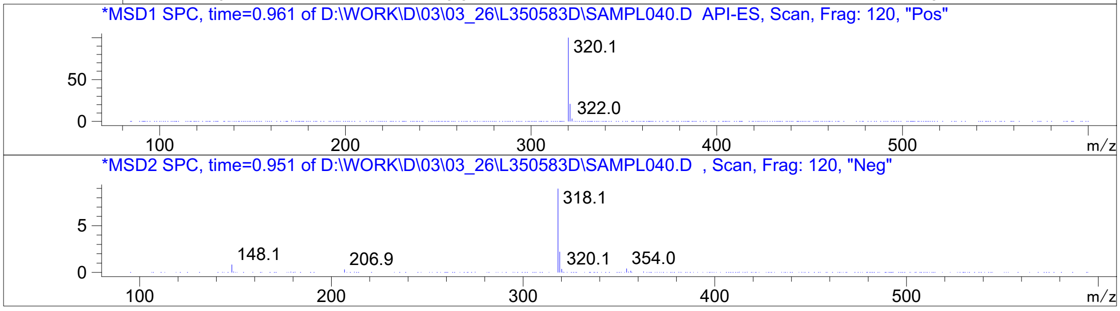


Inj.Date 3/26/2021 K -VL- Acq. Method C:\HPCHEM\-> ->

Z1531604940_

MaxPeak: 100.00%
Ret_Time: 0.968 min

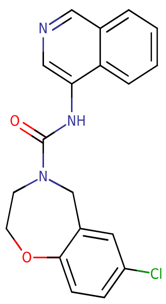

**Mol Wt** 353.8
**Exact Mass** 353.11

| # | Time | Area% |
| --- | --- | --- |
| 1 | 0.968 | 100.00 |

U693154$1

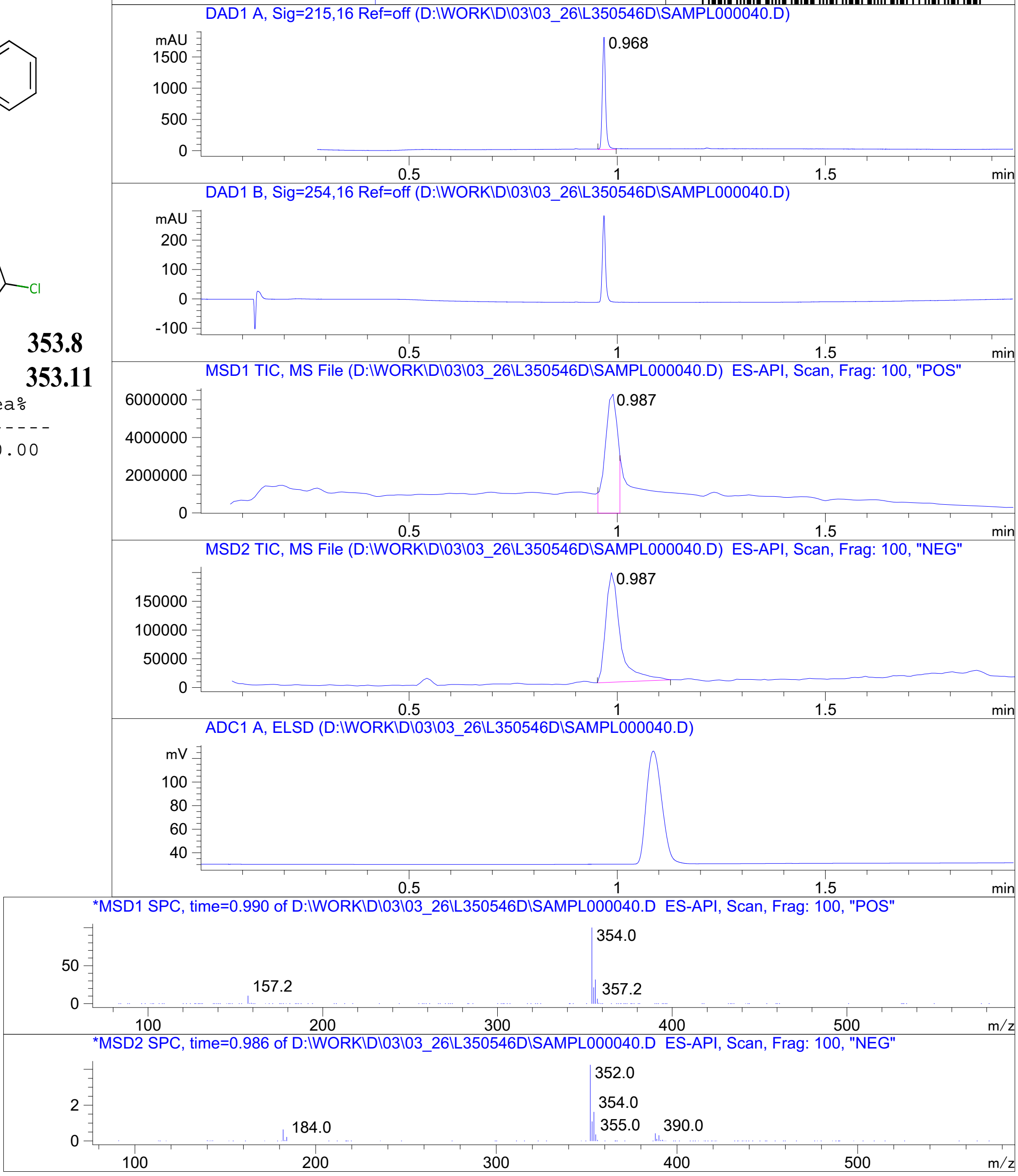


RT 0.987

RT 0.987

Inj.Date 3/26/2021 K  Acq. Method C:\CHEM32\-> ->

Z4924562414_

MaxPeak: 98.19%
Ret_Time: 1.054 min

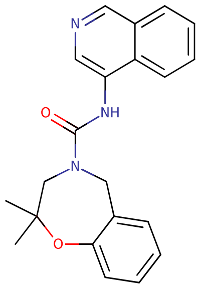


**Mol Wt** 347.41
**Exact Mass** 347.19

| # | Time | Area% |
|---|---|---|
| 1 | 0.681 | 1.81 |
| 2 | 1.054 | 98.19 |

# U689906$3

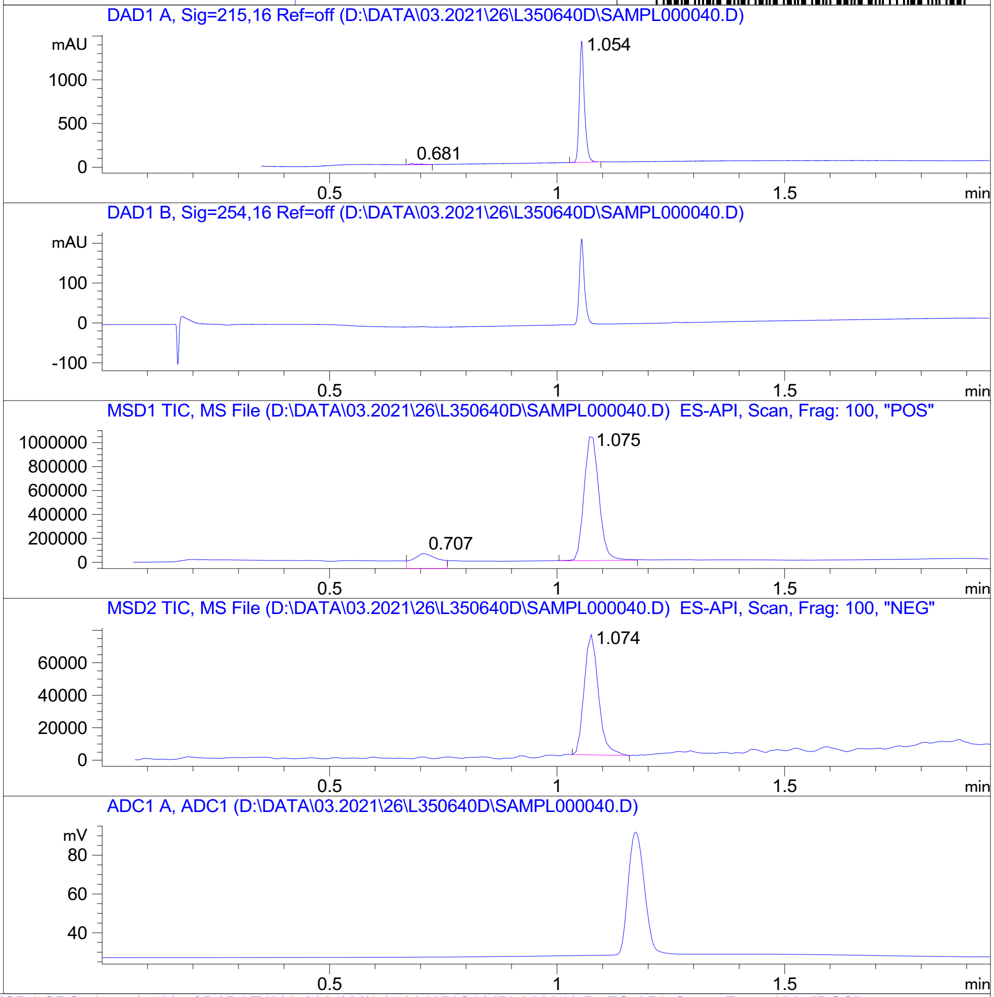


RT 0.707

RT 1.075

RT 1.074

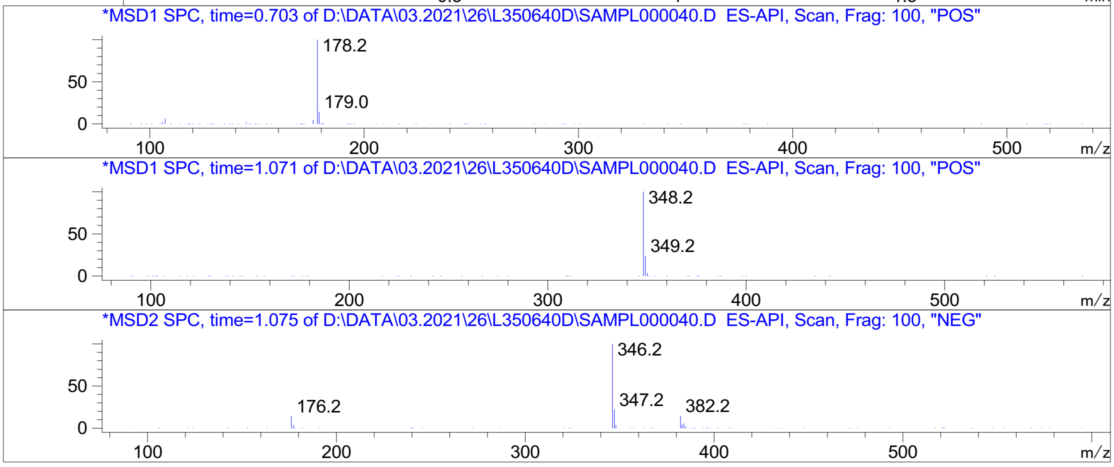


Inj.Date 3/26/2021 CH P2-E-04 - 4 - Acq. Method C:\CHEM32\-> ->

Z2150969870_

MaxPeak: 100.00%
Ret_Time: 0.990 min

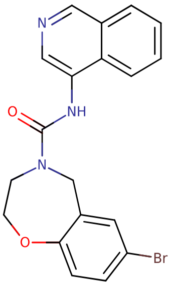


**Mol Wt** 398.25

**Exact Mass** 397.06

| # | Time | Area% |
|---|---|---|
| 1 | 0.990 | 100.00 |

U689908$1

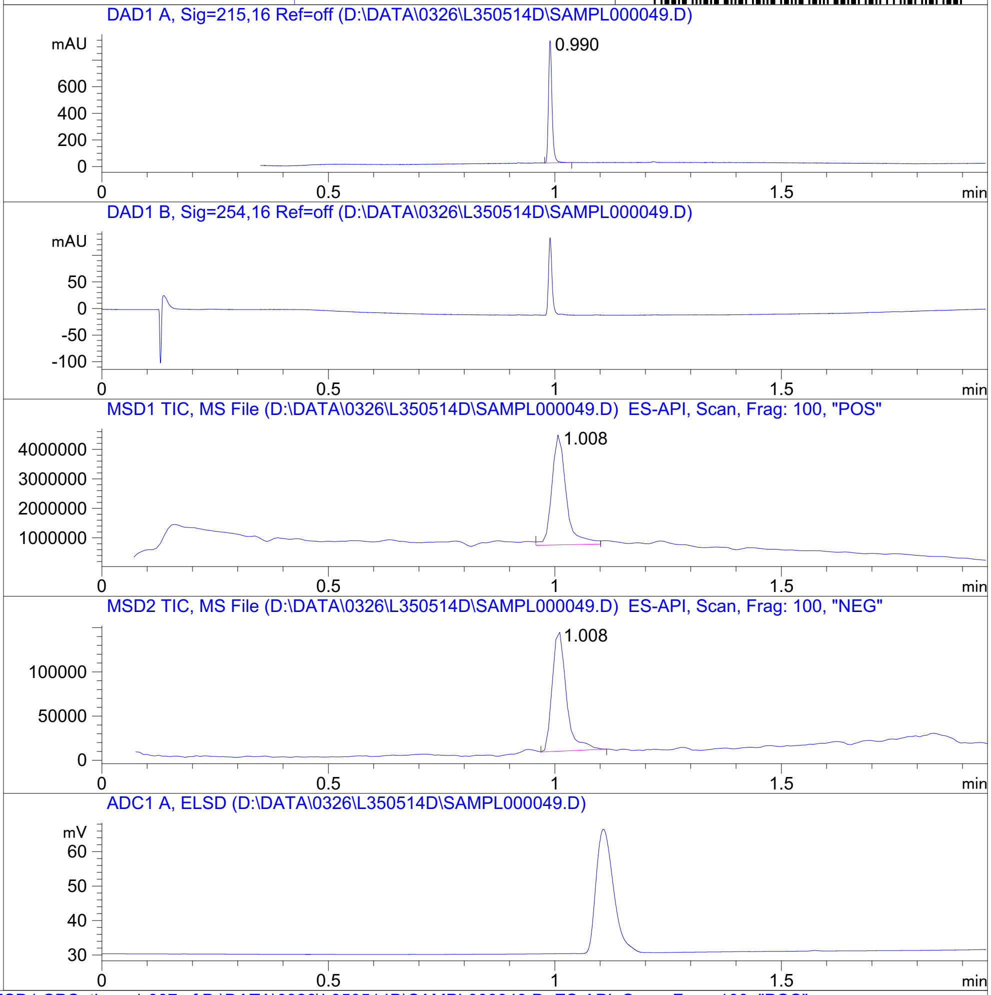


RT 1.008

RT 1.008

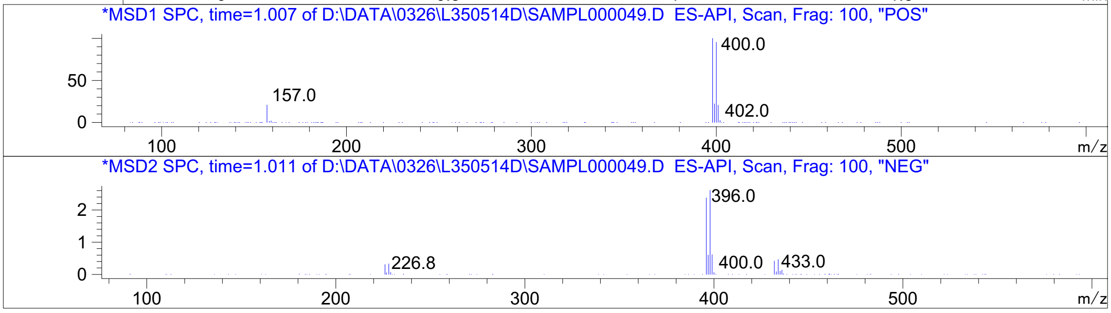




Z2017308172_

U963858$1

MaxPeak:  100.00%
Ret_Time: 1.100 min

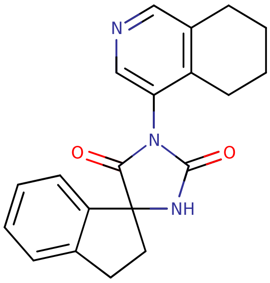


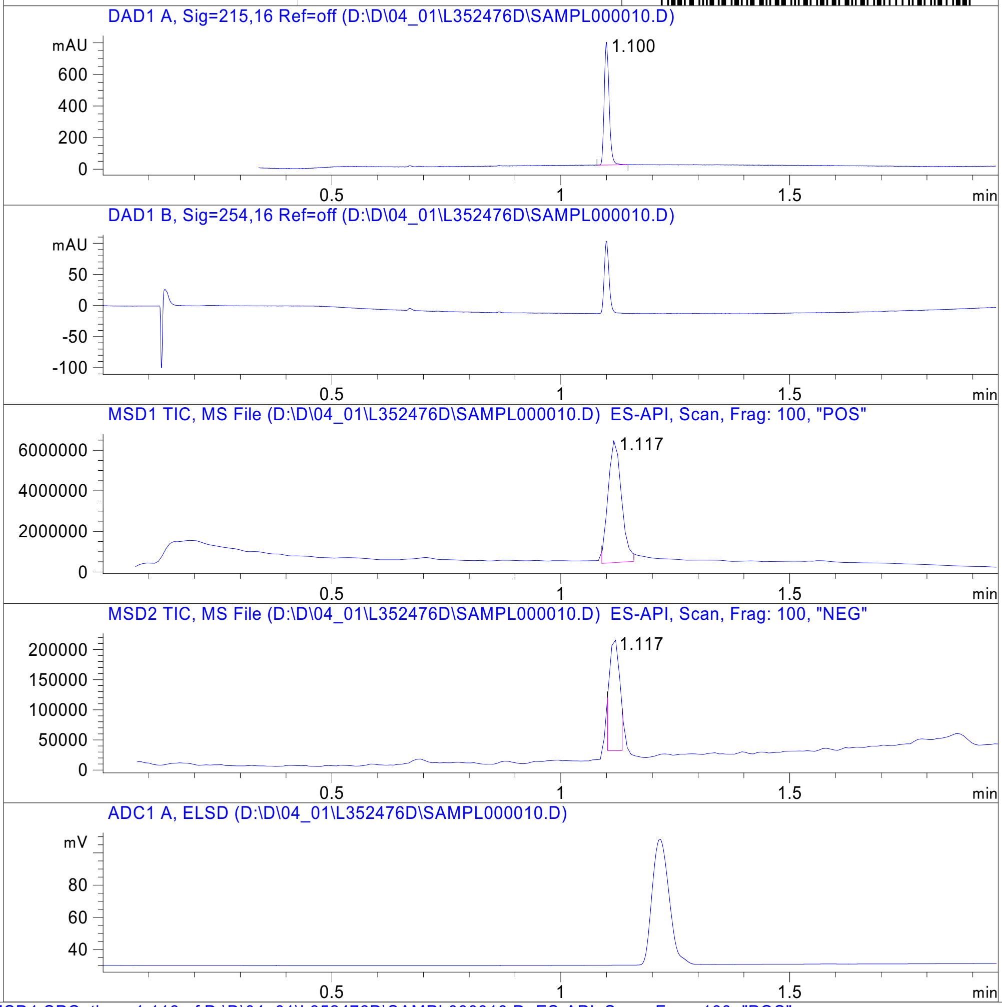


**Mol Wt** 333.38

**Exact Mass** 333.17

```
#  Time   Area%
-----------------
1  1.100  100.00
```

RT 1.117

RT 1.117

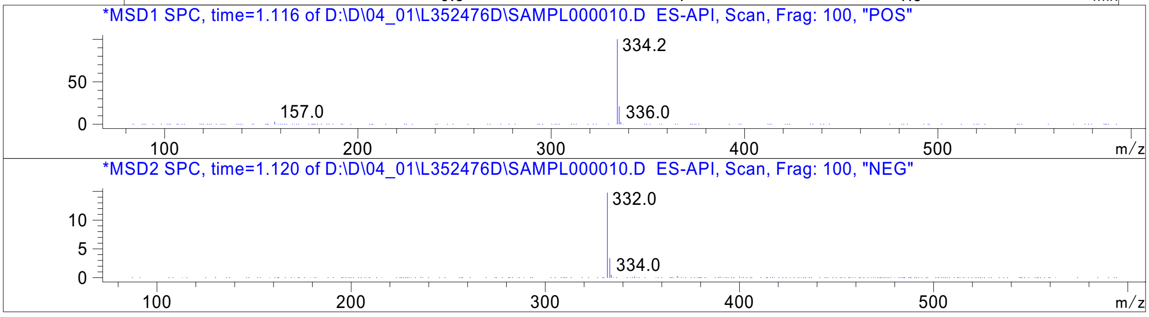




Z4929616344_

MaxPeak: 100.00%
Ret_Time: 1.332 min

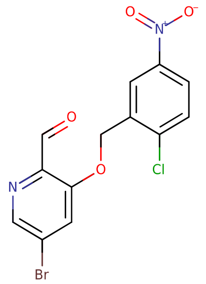


**Mol Wt** 371.57
**Exact Mass** 371.93

| # | Time | Area% |
|---|---|---|
| 1 | 1.332 | 100.00 |

U920096$2

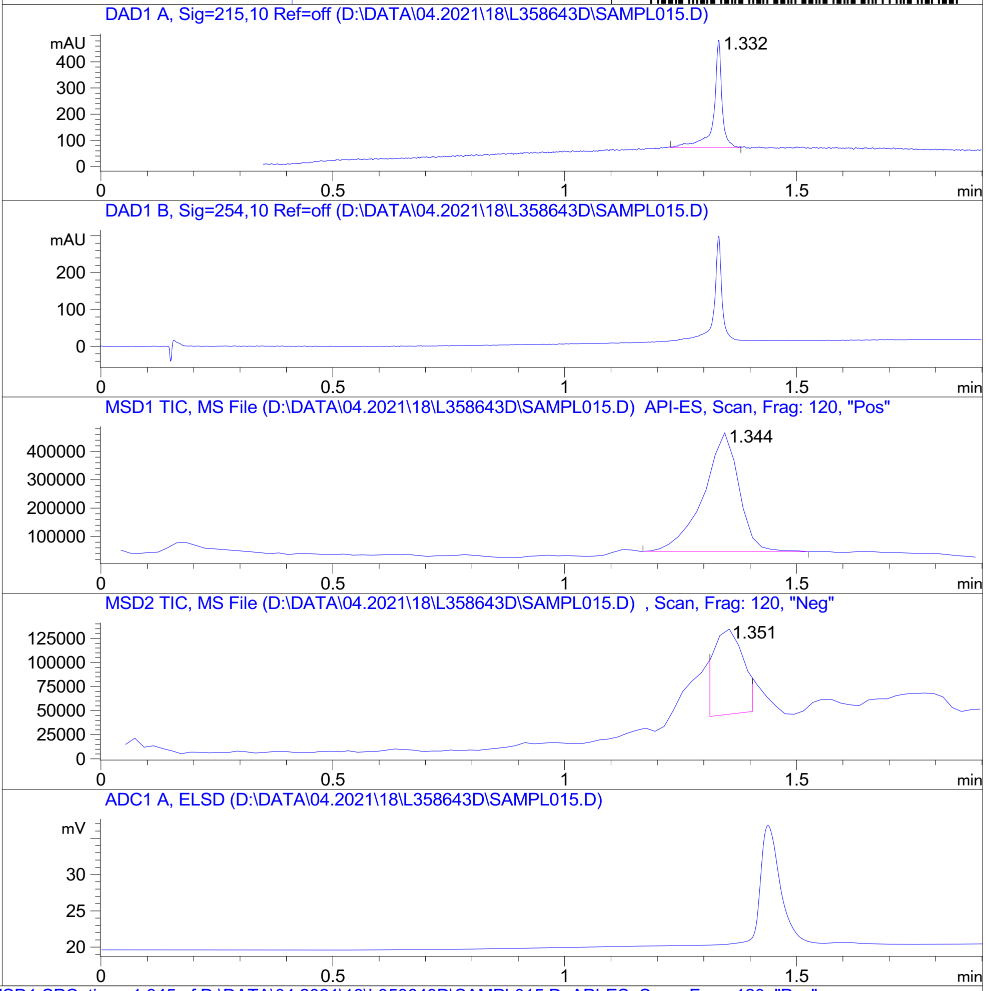


RT 1.344

RT 1.351

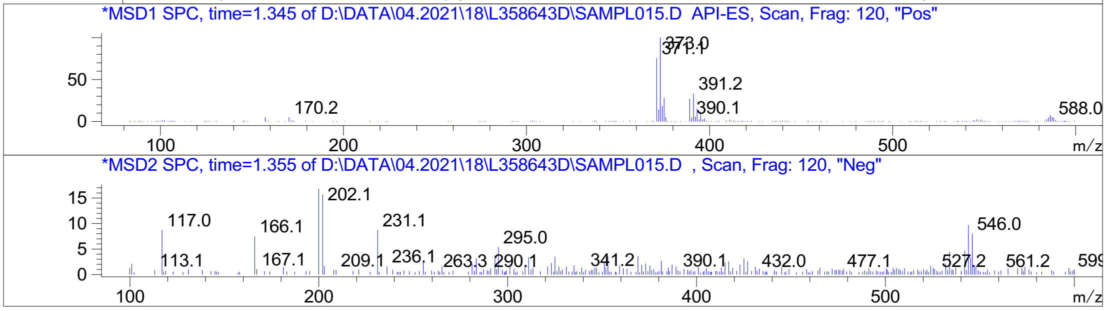


Inj.Date 4/18/2021 CH P2-B-06 -SL- Acq. Method C:\HPCHEM\-> ->

Z4954569105_

# U920109$3

MaxPeak: 100.00%
Ret_Time: 1.465 min

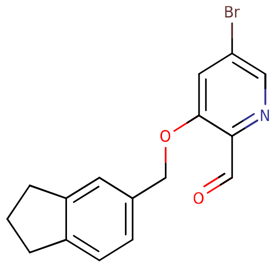


**Mol Wt** 332.19

**Exact Mass** 331.04

| # | Time | Area% |
|---|---|---|
| 1 | 1.465 | 100.00 |

RT 1.480

RT 1.493

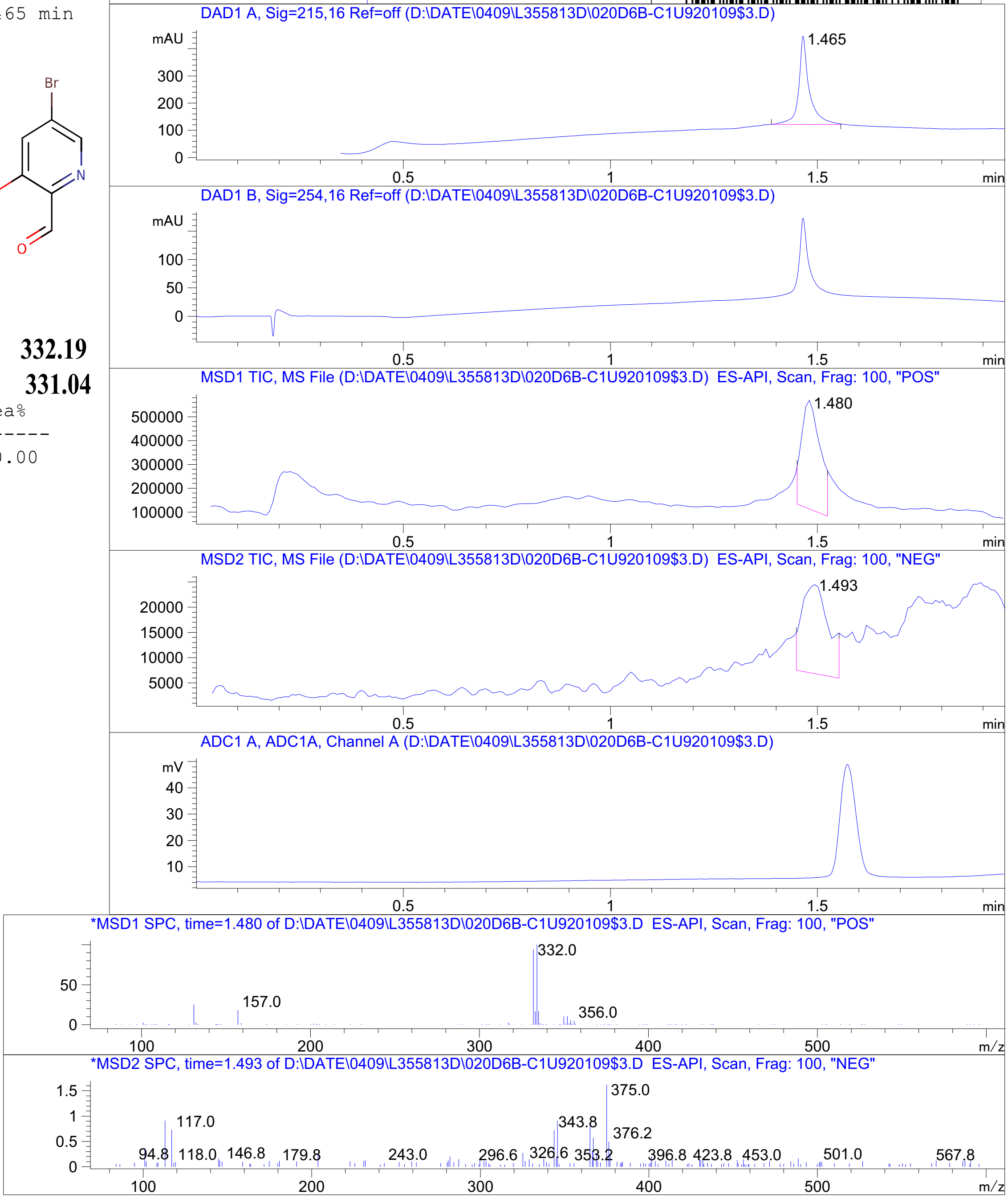




Z4954569202_

MaxPeak: 98.52%
Ret_Time: 0.661 min

# V005787$4

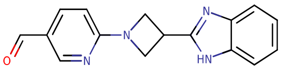

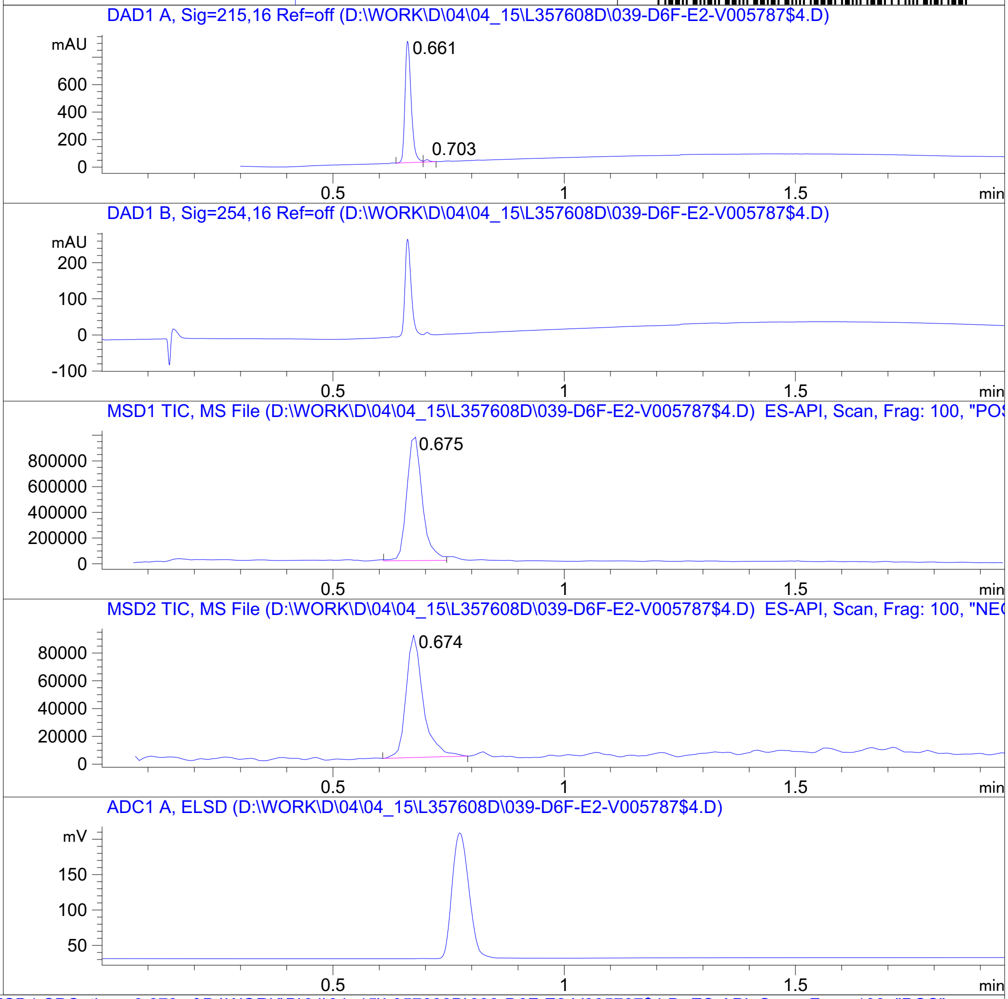


**Mol Wt** 278.31

**Exact Mass** 278.13

| # | Time | Area% |
|---|---|---|
| 1 | 0.661 | 98.52 |
| 2 | 0.703 | 1.48 |

RT 0.675

RT 0.674

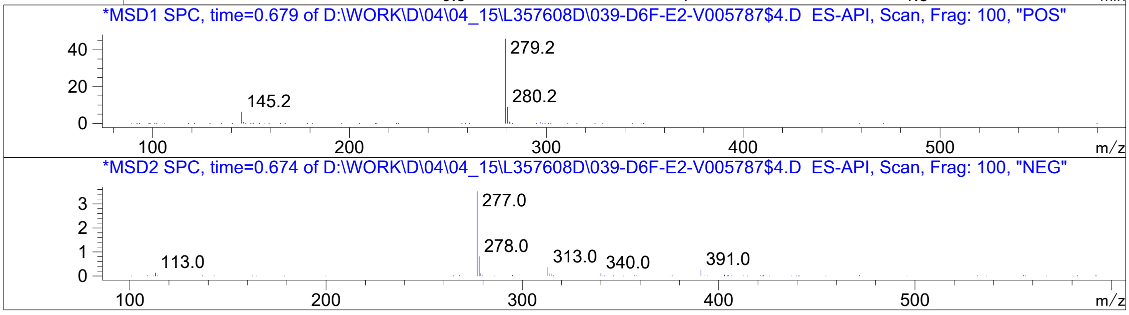




Z4968112353_

MaxPeak: 100.00%
Ret_Time: 2.310 min

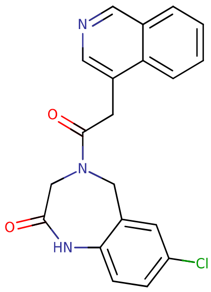


**Mol Wt 365.81**

**Exact Mass 365.11**

| # | Time | Area% |
|---|---|---|
| 1 | 2.310 | 100.00 |

# V123973$1

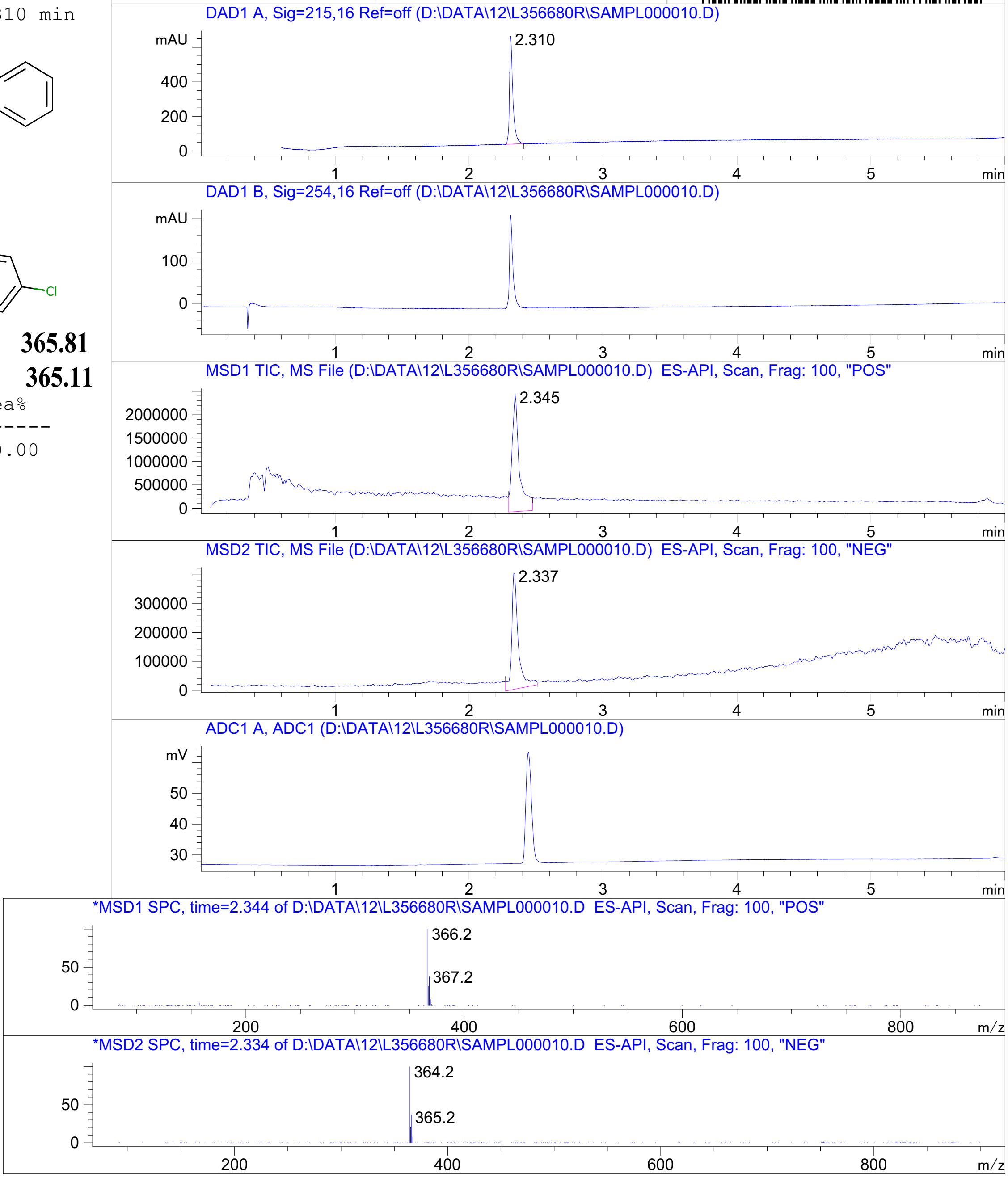


RT 2.345

RT 2.337



Z4924562394_

MaxPeak: 100.00%
Ret_Time: 1.483 min

# U920101$22

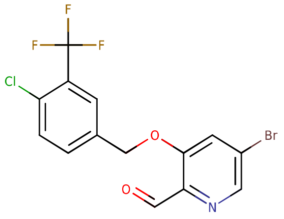


**Mol Wt** 394.57

**Exact Mass** 394.95

| # | Time | Area% |
|---|---|---|
| 1 | 1.483 | 100.00 |

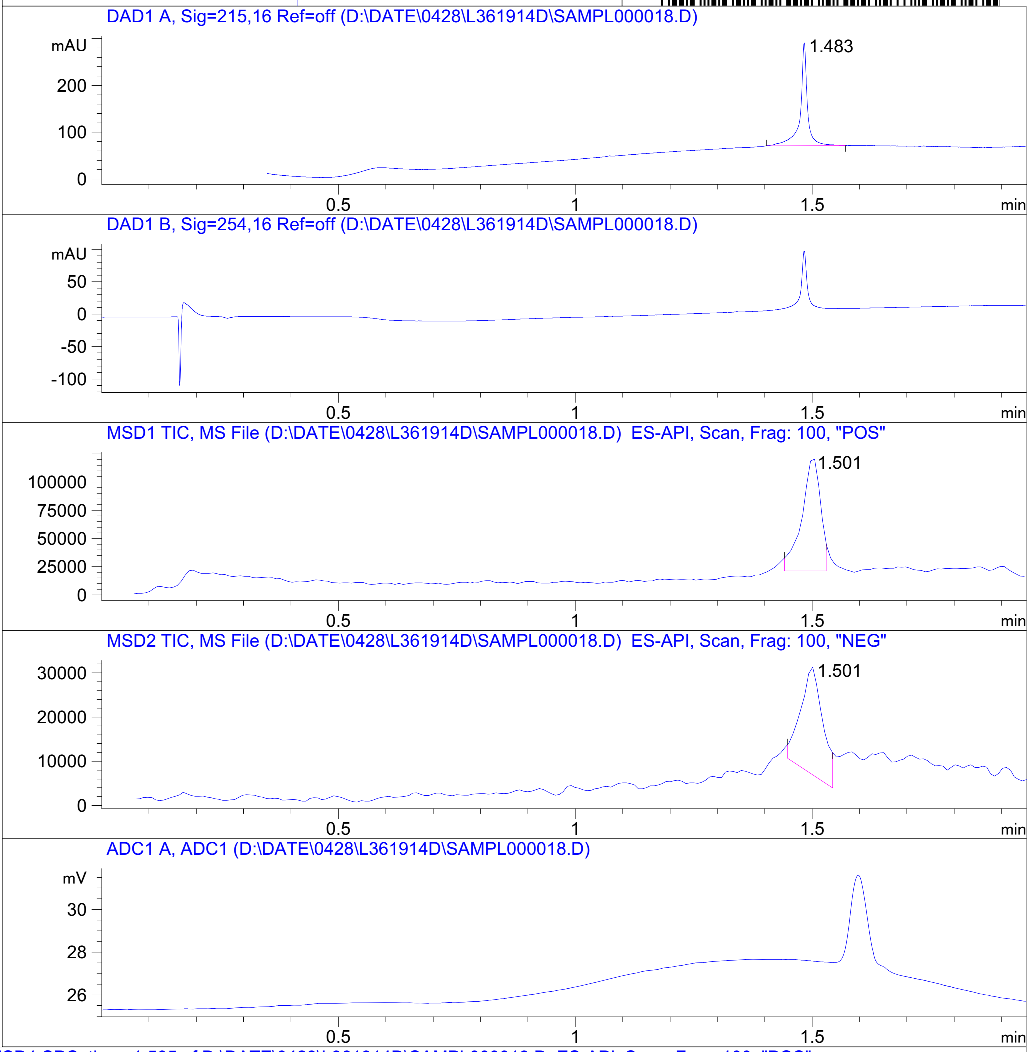


RT 1.501

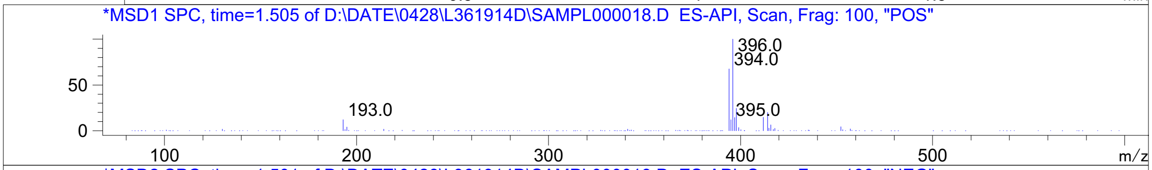


RT 1.501

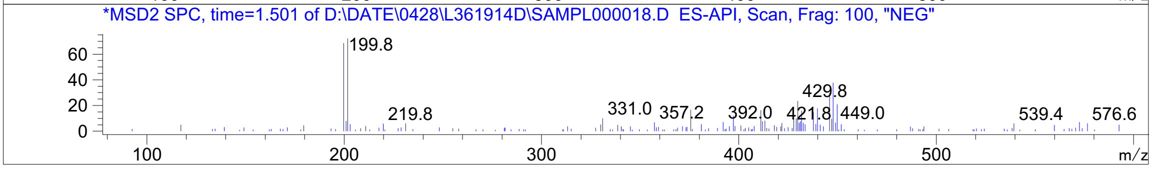




Z4954569122_

MaxPeak:  96.26%
Ret_Time: 2.010 min

# V245837$1

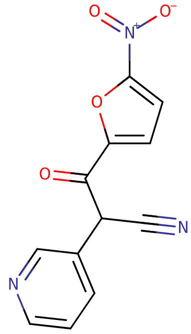


**Mol Wt** 257.2

**Exact Mass** 257.03

| # | Time | Area% |
|---|---|---|
| 1 | 0.971 | 3.74 |
| 2 | 2.010 | 96.26 |

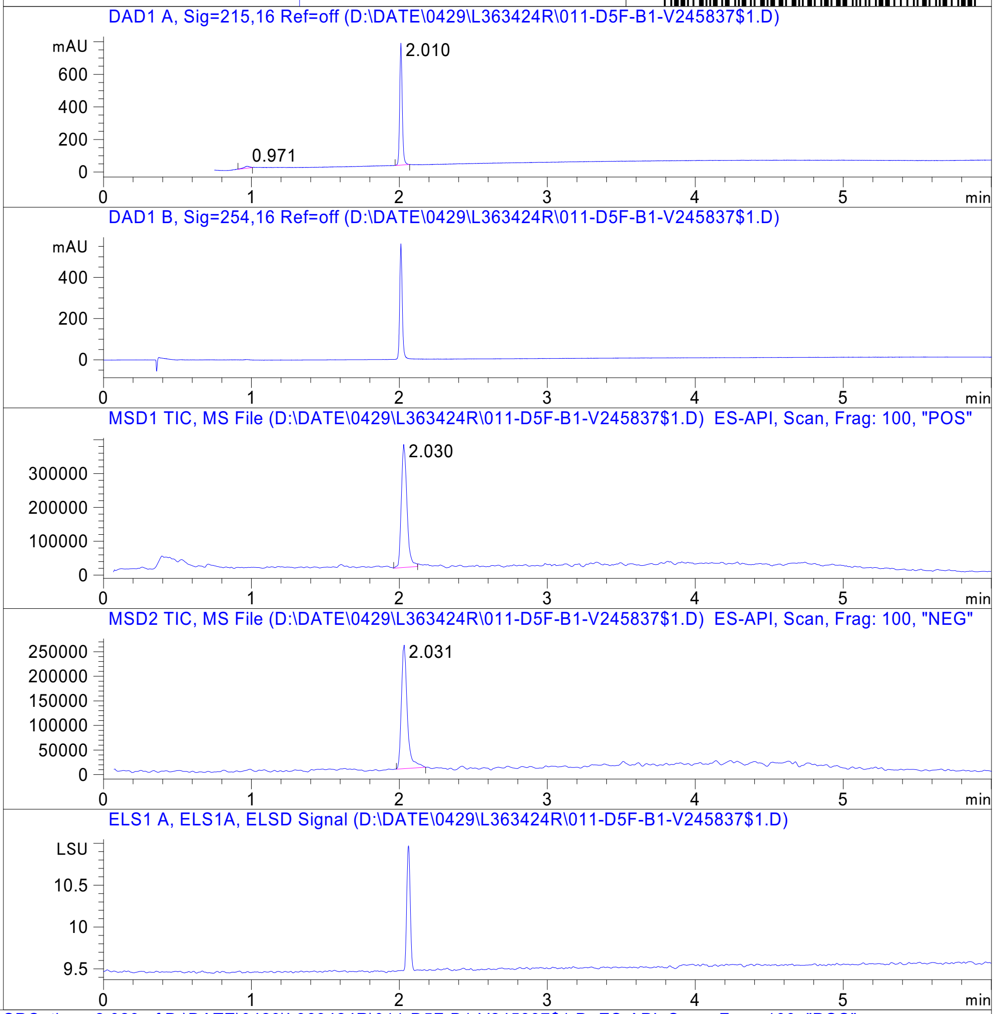


RT 2.030

RT 2.031

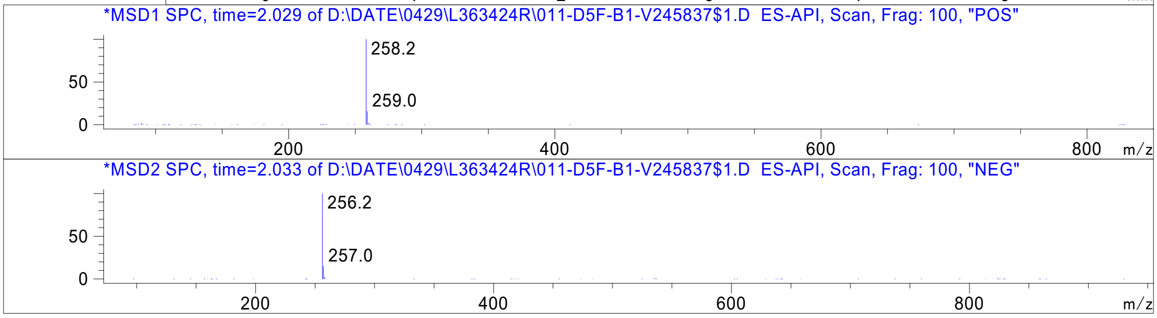




Z1410957731_

MaxPeak: 100.00%
Ret_Time: 3.908 min

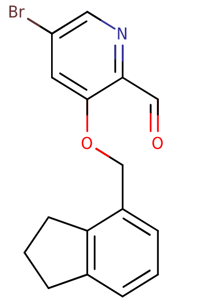

**Mol Wt** 332.19
**Exact Mass** 331.04

| # | Time | Area% |
|---|---|---|
| 1 | 3.908 | 100.00 |

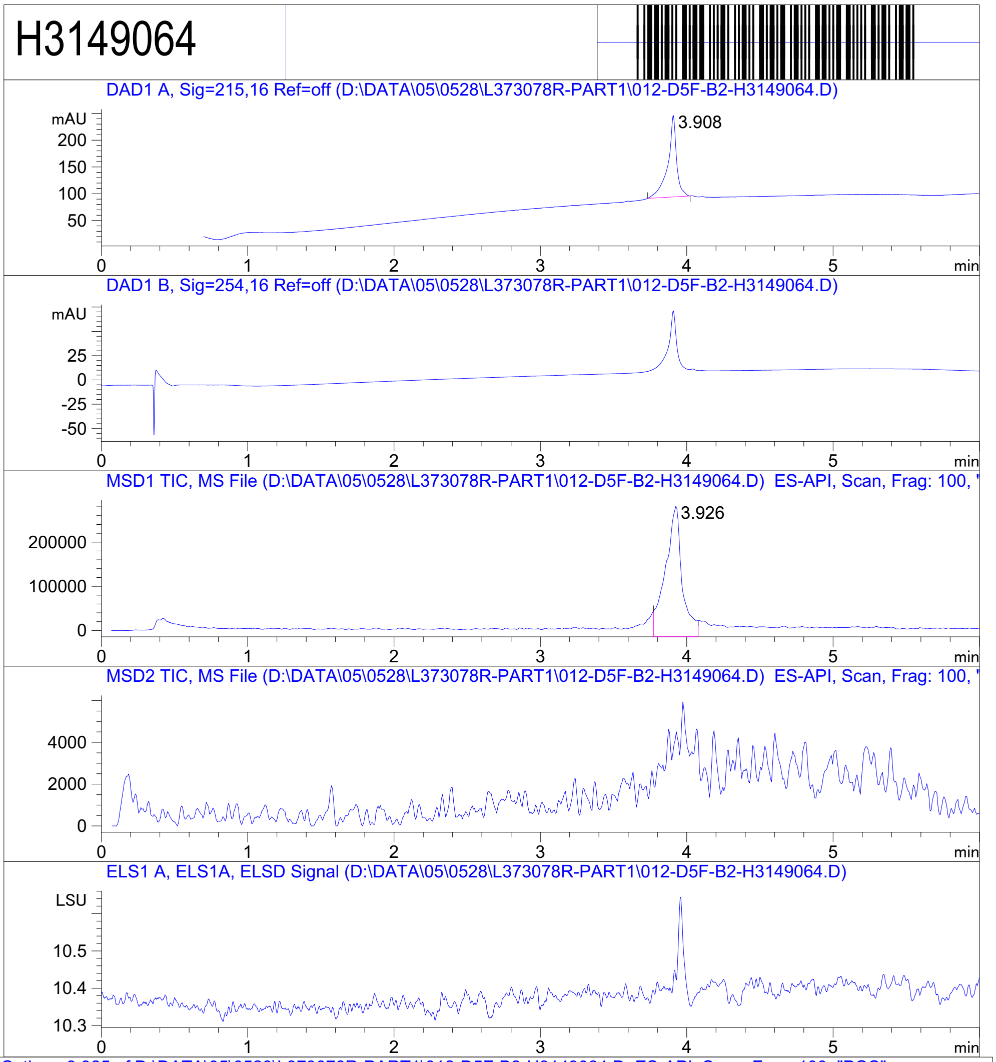


RT 3.926

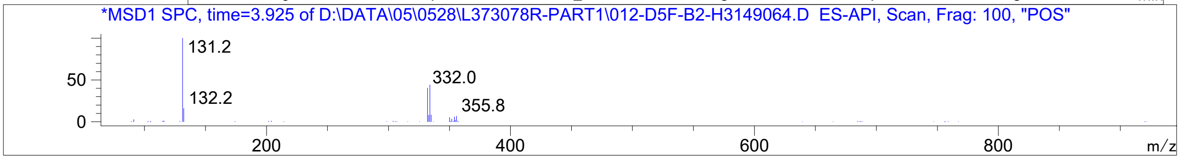


Inj.Date 5/28/2021 LT -15- Acq. Method C:\Chem32\-> ->

Z4989203250_

MaxPeak: 99.08%
Ret_Time: 1.100 min

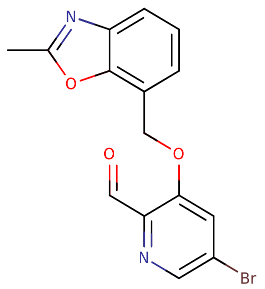

**Mol Wt** 347.16
**Exact Mass** 346

| # | Time | Area% |
|---|---|---|
| 1 | 1.100 | 99.08 |
| 2 | 1.159 | 0.92 |

V296344$7

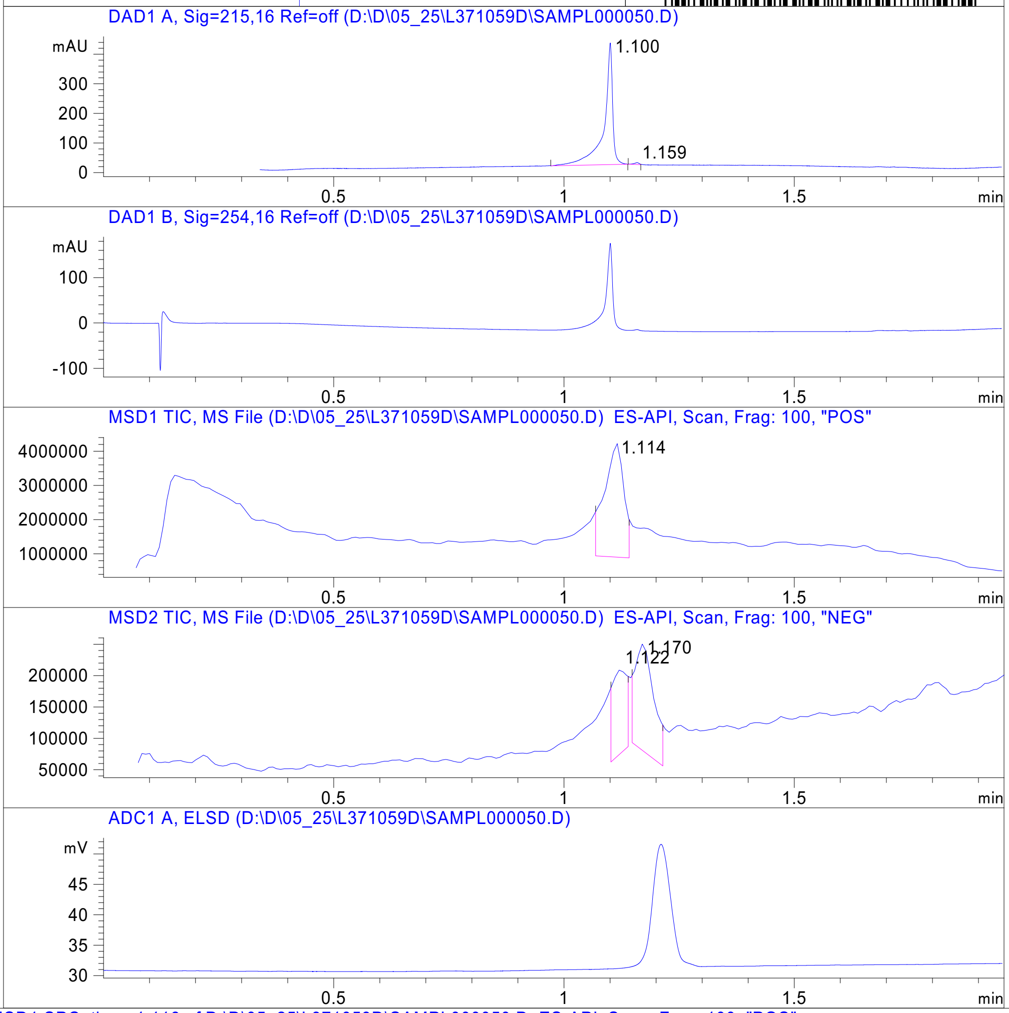


RT 1.114

RT 1.122

RT 1.170

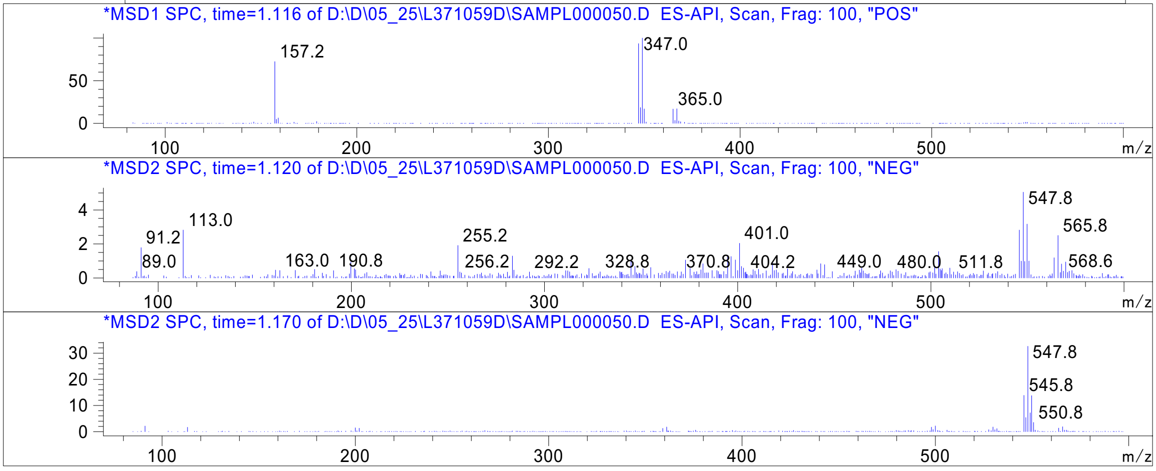


Inj.Date 5/23/2021 N -3- Acq. Method C:\CHEM32\-> ->

Z4989203262_

MaxPeak: 100.00%
Ret_Time: 1.568 min

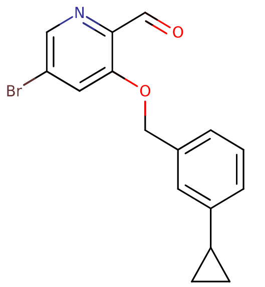


**Mol Wt** 332.19
**Exact Mass** 331.04

| # | Time | Area% |
|---|---|---|
| 1 | 1.568 | 100.00 |

V211811$A

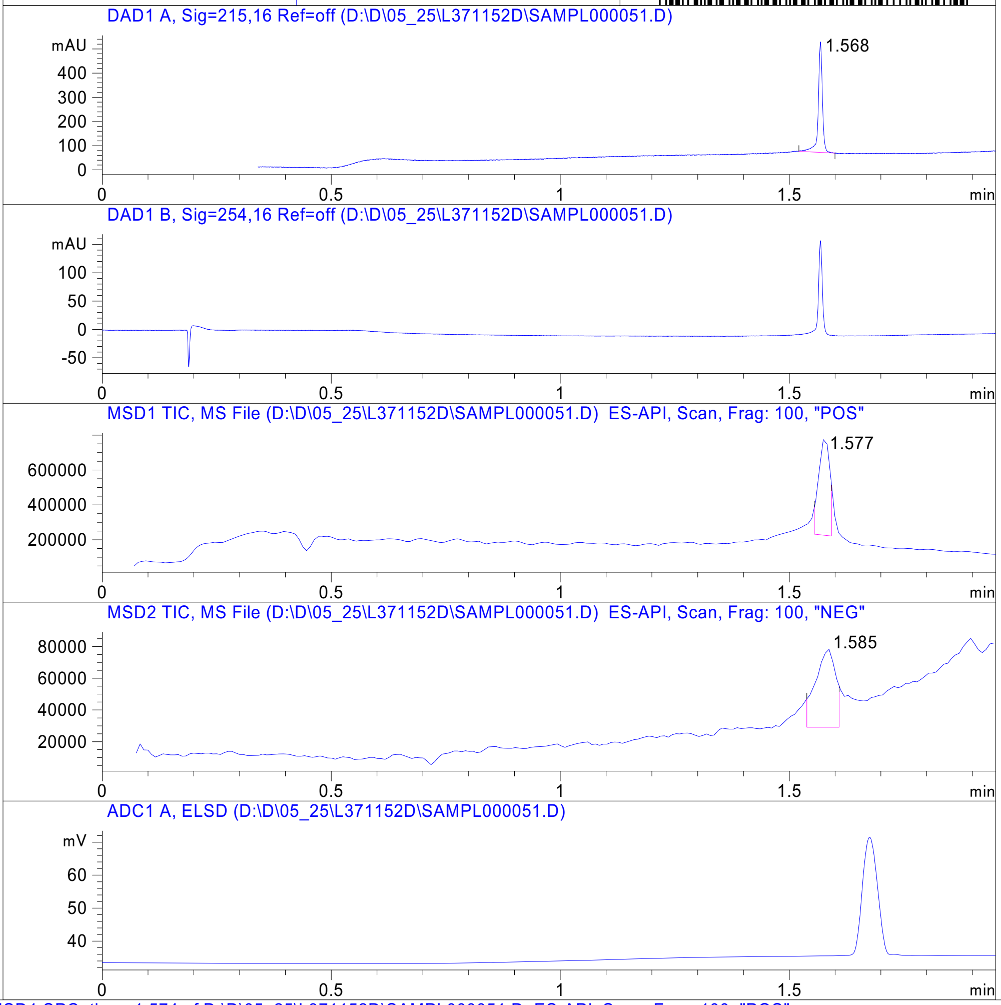


RT 1.577

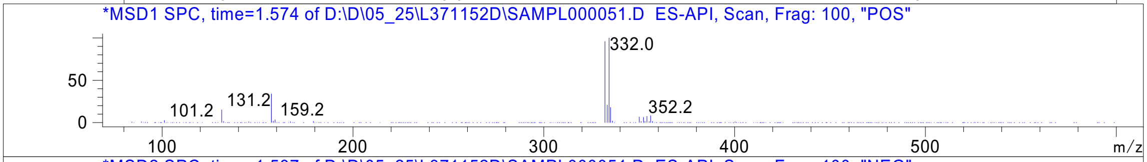


RT 1.585

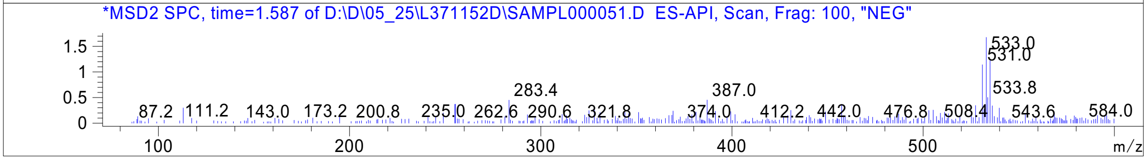




Z4989203264_

MaxPeak: 100.00%
Ret_Time: 1.128 min

V233168$9

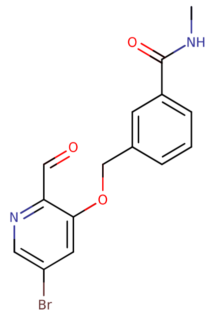

**Mol Wt** 349.18

**Exact Mass** 348.02

| # | Time | Area% |
|---|---|---|
| 1 | 1.128 | 100.00 |

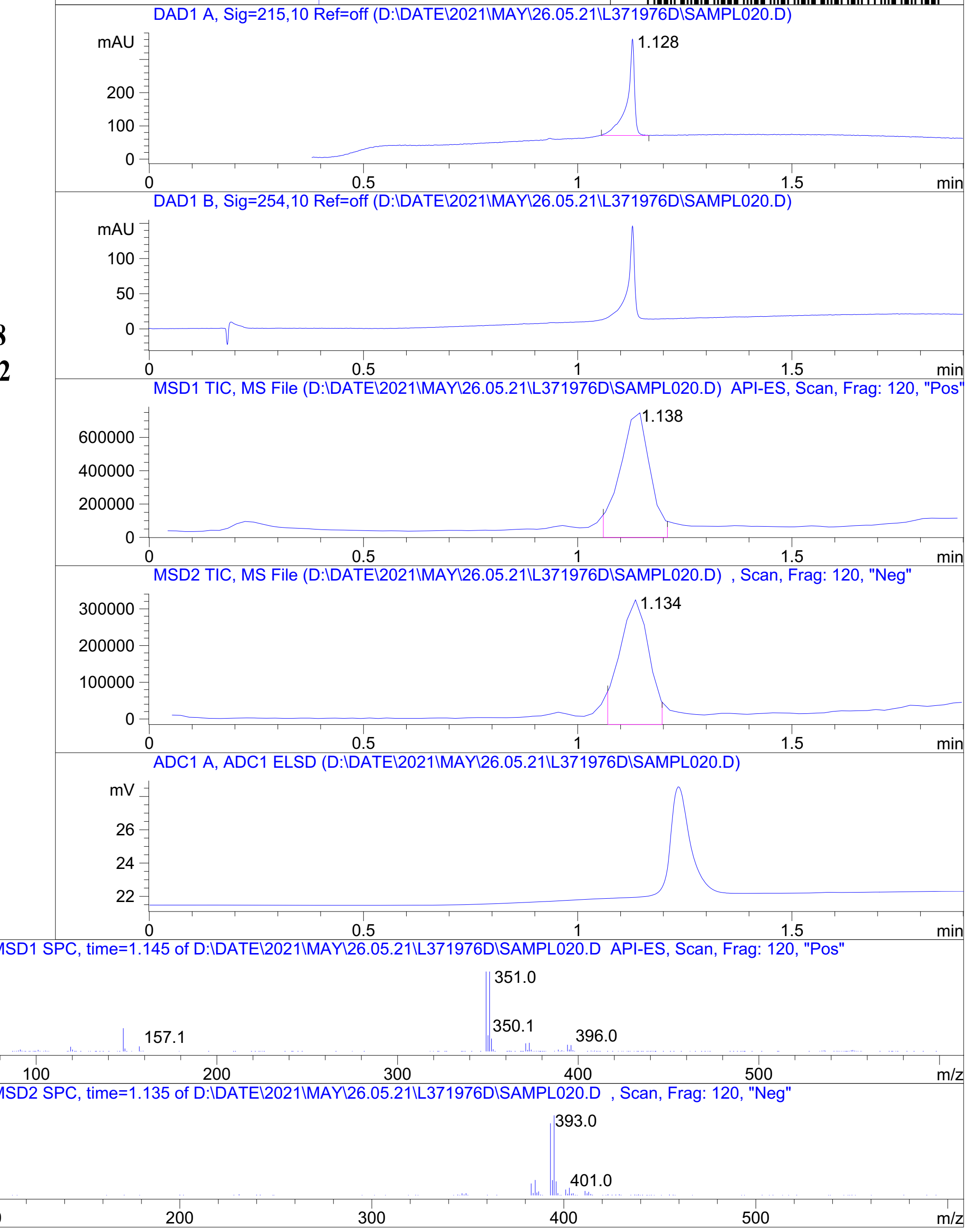


RT 1.138

RT 1.134

Inj.Date 5/26/2021 T P2-C-03 -SL- Acq. Method C:\HPCHEM\-> ->

Z5009000157_

MaxPeak: 100.00%
Ret_Time: 1.476 min

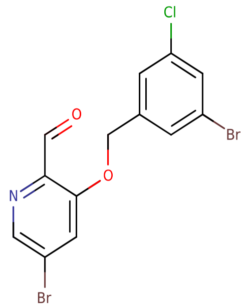


**Mol Wt 405.47**
**Exact Mass 404.87**

| # | Time | Area% |
|---|---|---|
| 1 | 1.476 | 100.00 |

V233167$2

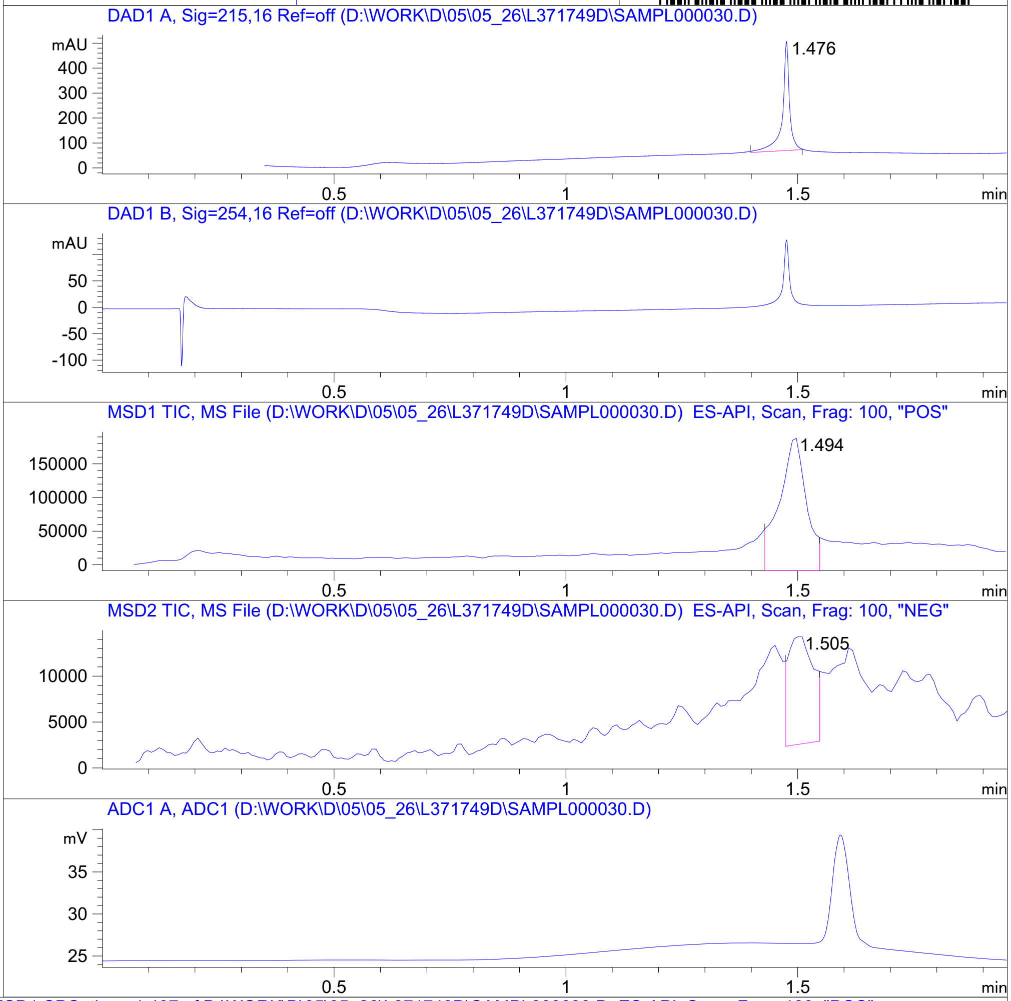


RT 1.494

RT 1.505

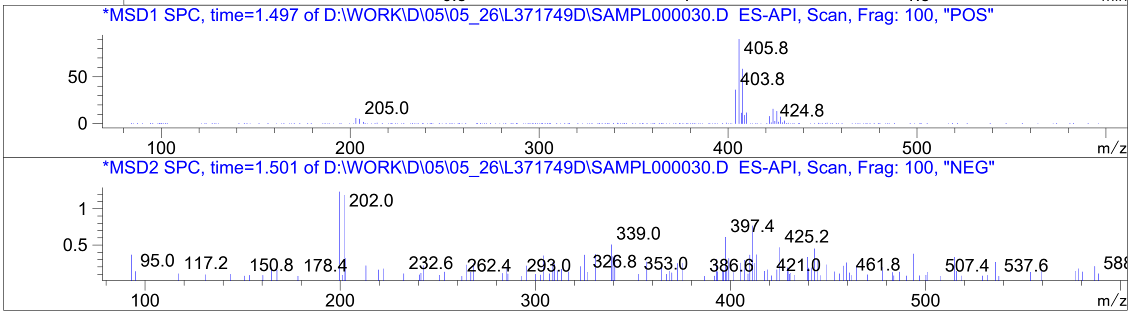


Inj.Date 5/26/2021 K - 4 - Acq. Method C:\CHEM32\-> ->

Z5009000156_

MaxPeak: 100.00%
Ret_Time: 3.583 min

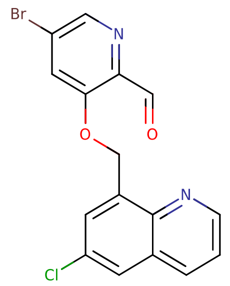


**Mol Wt** 377.62
**Exact Mass** 377.97

| # | Time | Area% |
|---|---|---|
| 1 | 3.583 | 100.00 |

V233165$6

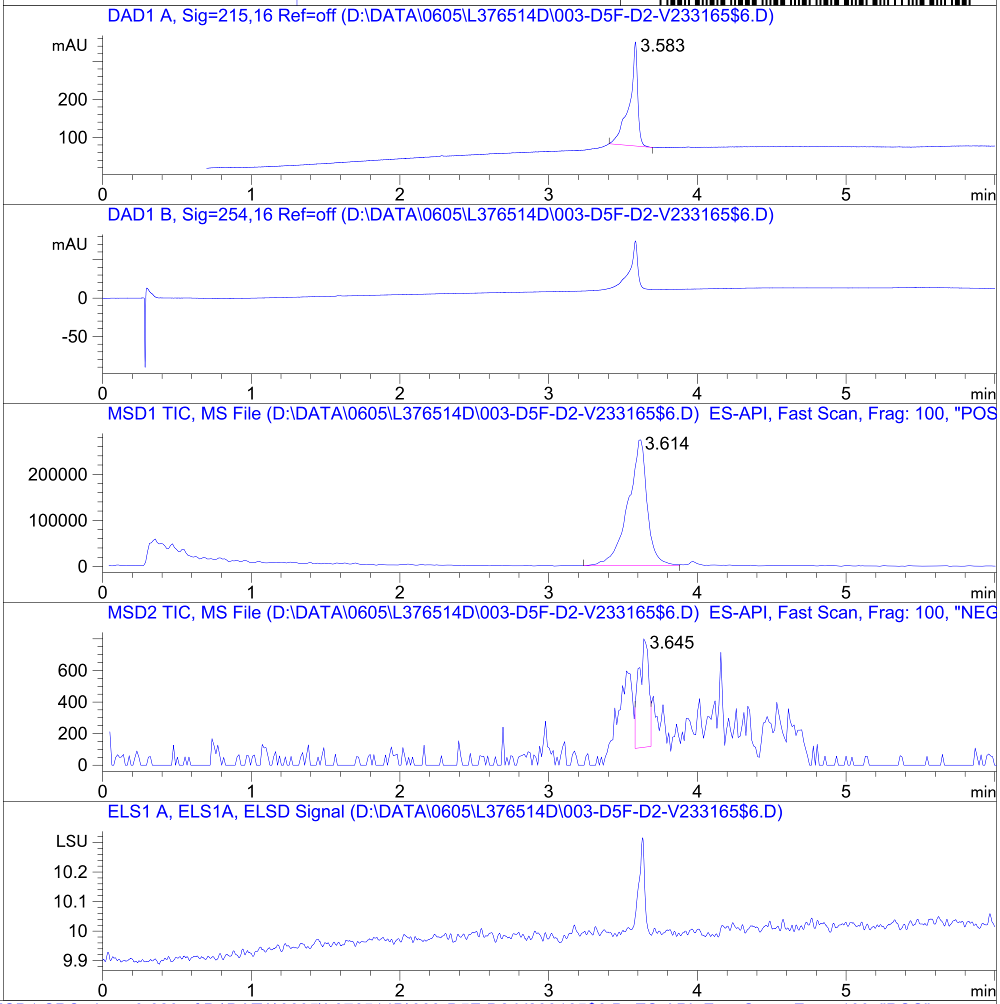


RT 3.614

RT 3.645

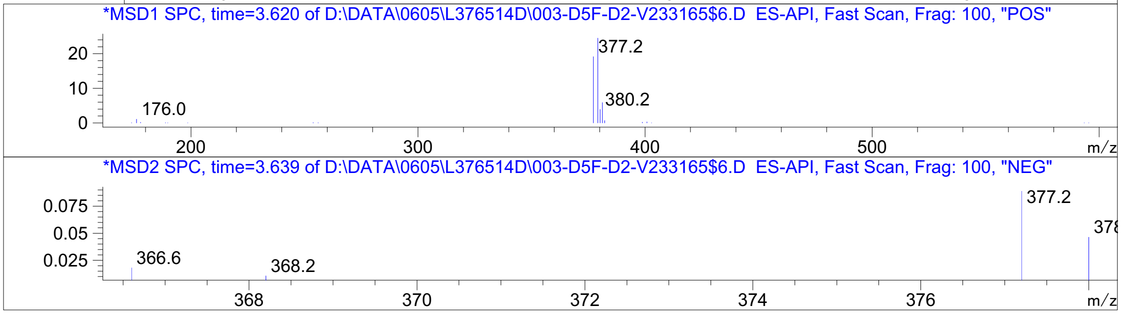


Inj.Date 6/5/2021 OA 24 Acq. Method C:\Users\ -> ->

Z5009000154_

MaxPeak: 90.89%
Ret_Time: 1.542 min

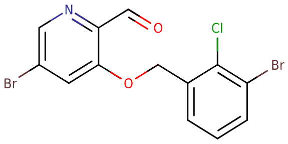


**Mol Wt** 405.47
**Exact Mass** 404.87

| # | Time | Area% |
|---|---|---|
| 1 | 0.954 | 4.30 |
| 2 | 0.990 | 4.81 |
| 3 | 1.542 | 90.89 |

V211815$4

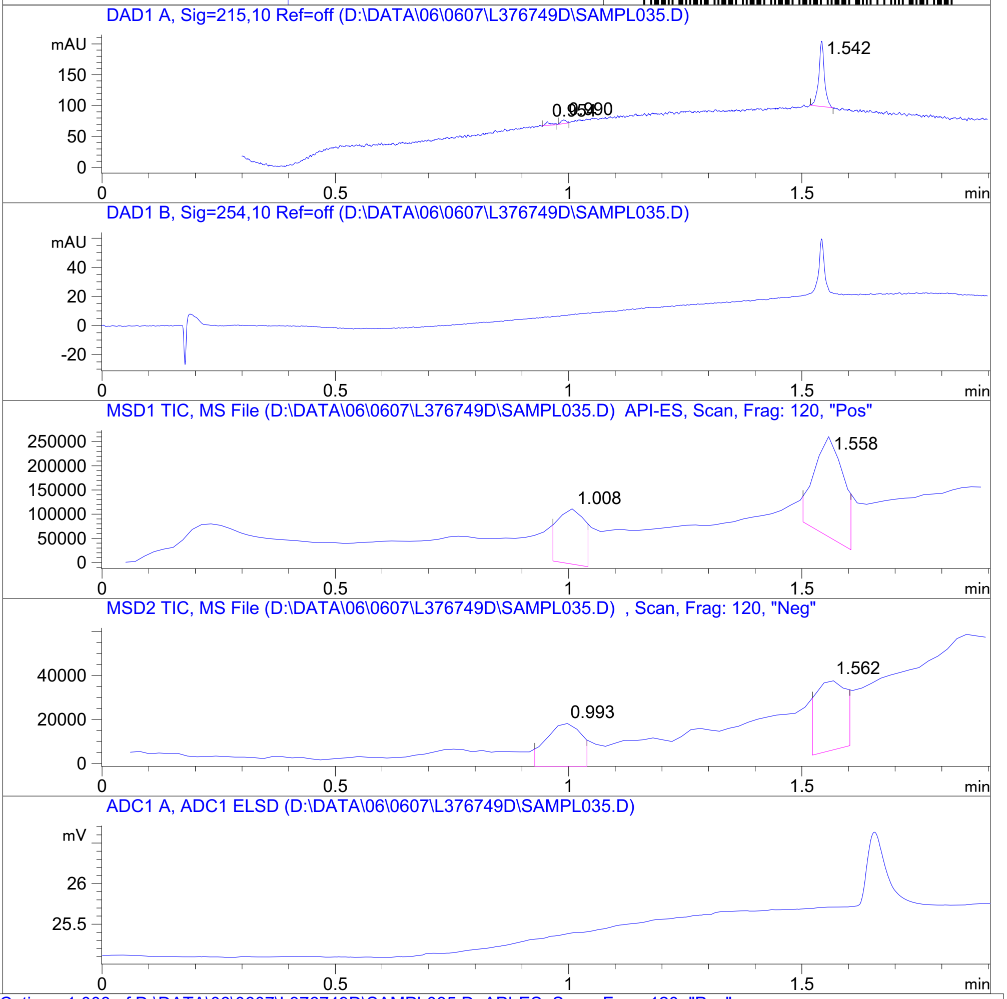


RT 1.008

RT 1.558

RT 0.993

RT 1.562

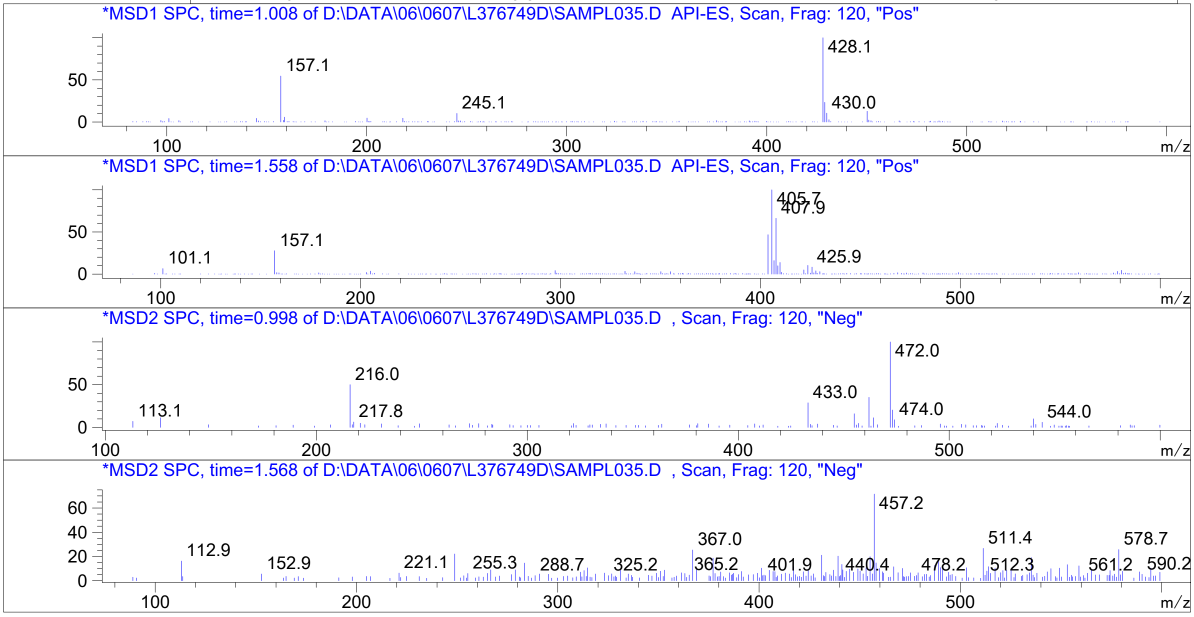


Inj.Date 6/6/2021 LT -VL- Acq. Method C:\HPCHEM\-> ->

Z4989203260_

MaxPeak: 100.00%
Ret_Time: 1.245 min

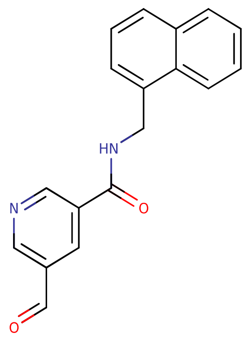


**Mol Wt** 290.32
**Exact Mass** 290.12

| # | Time | Area% |
|---|---|---|
| 1 | 1.245 | 100.00 |

V220413$3

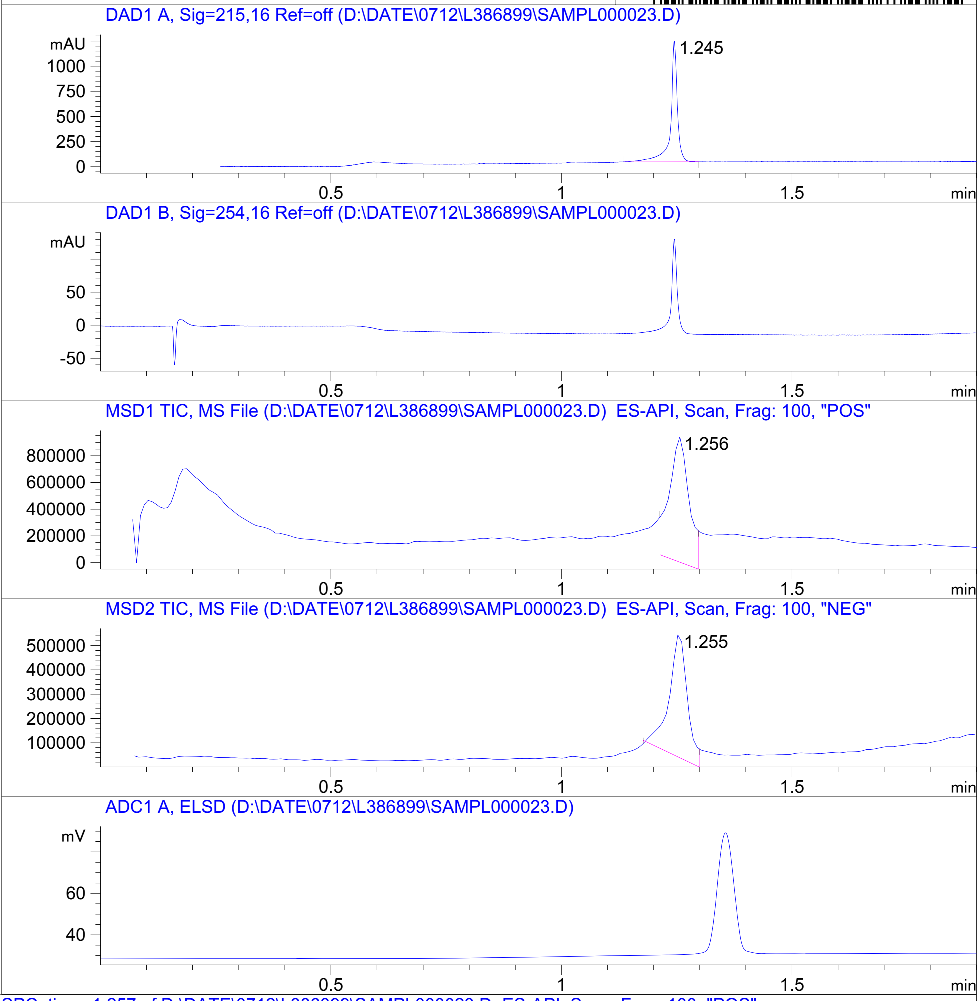


RT 1.256

RT 1.255

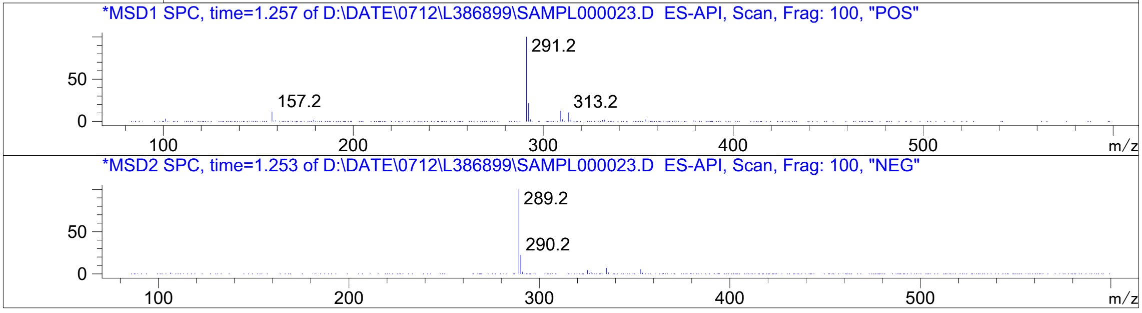




Z4968112276_

MaxPeak: 100.00%
Ret_Time: 1.237 min

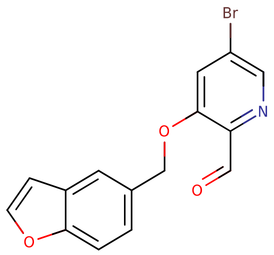


**Mol Wt** 332.15
**Exact Mass** 330.99

| # | Time | Area% |
|---|---|---|
| 1 | 1.237 | 100.00 |

# V816207$5

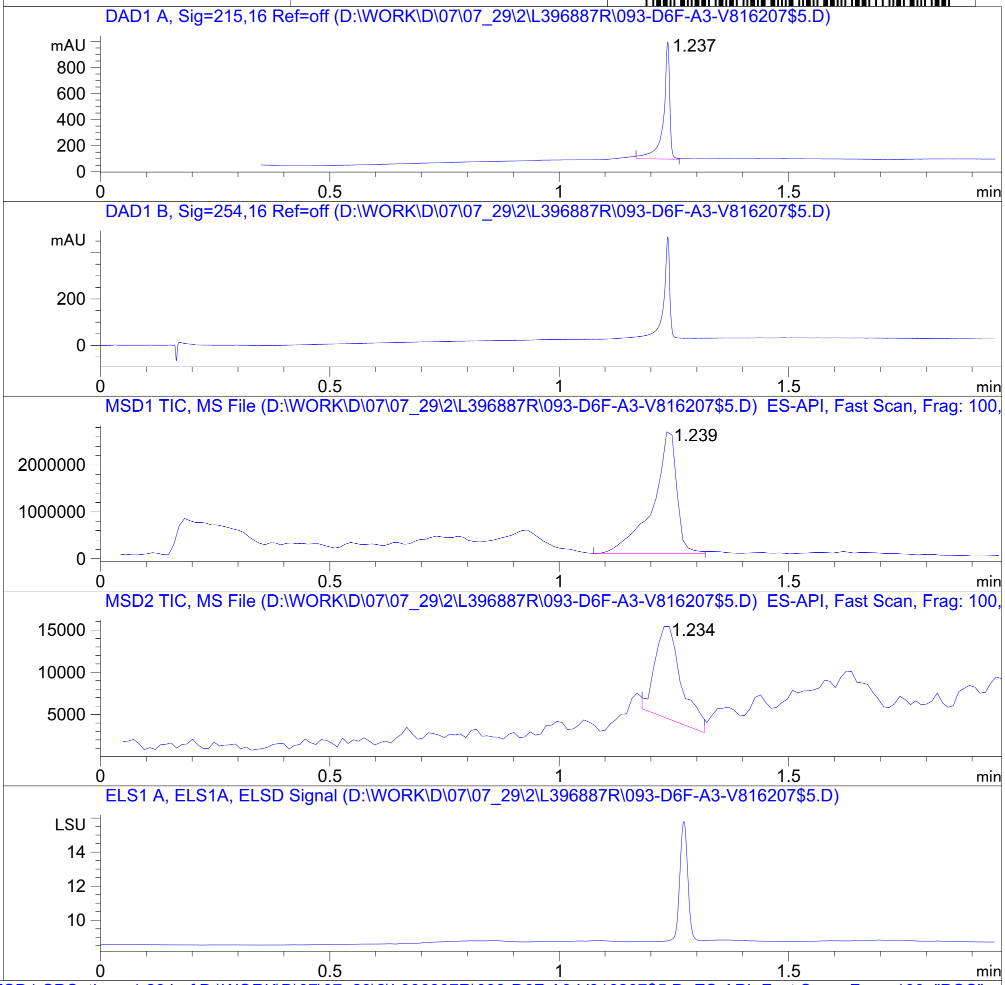


RT 1.239

RT 1.234

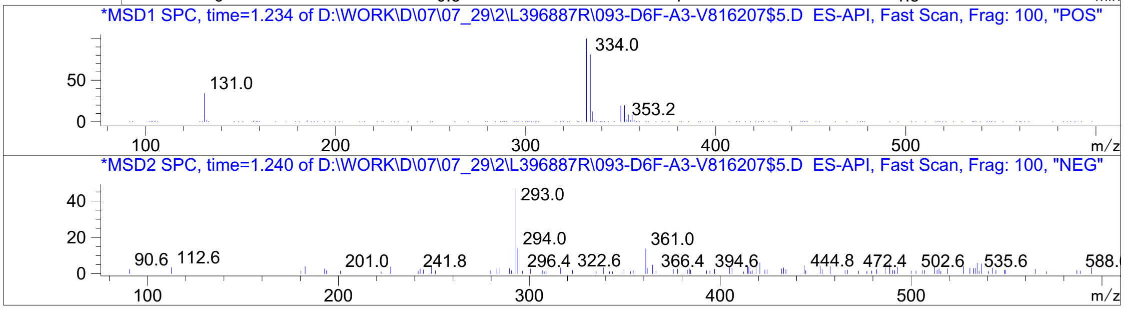


Inj.Date 7/29/2021 K <invalid> Acq. Method C:\Users\ -> ->

Z5084954212_

# V609074$4

MaxPeak: 100.00%
Ret_Time: 1.504 min

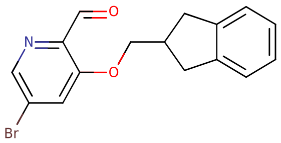


**Mol Wt** 332.19
**Exact Mass** 331.04

| # | Time | Area% |
|---|---|---|
| 1 | 1.504 | 100.00 |

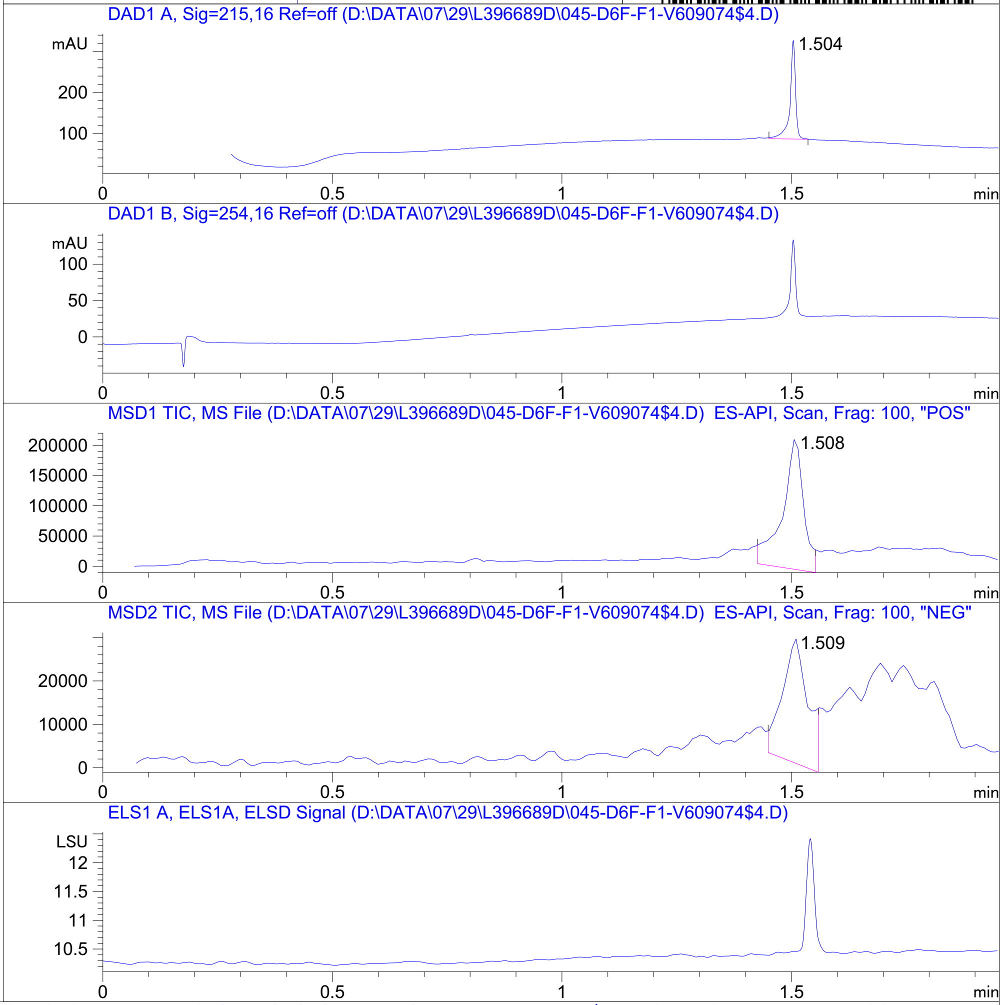


RT 1.508

RT 1.509

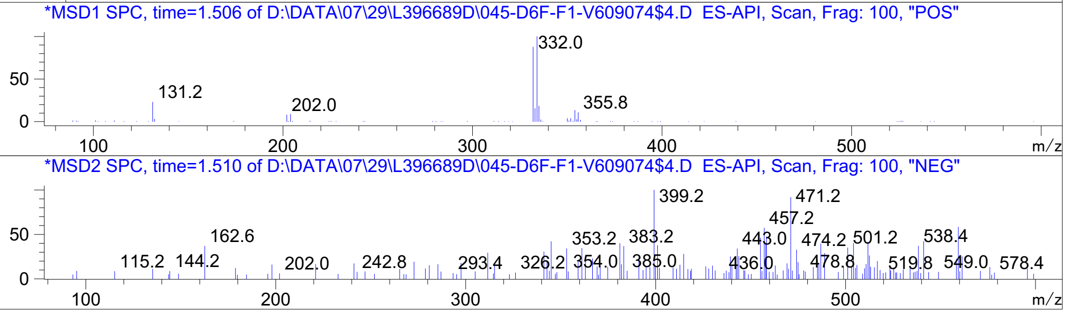




Z5084954221_

MaxPeak: 100.00%
Ret_Time: 1.551 min

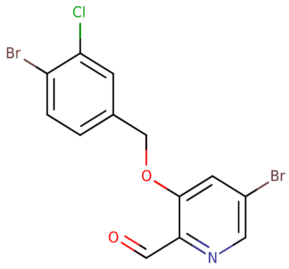


**Mol Wt** 405.47

**Exact Mass** 404.87

| # | Time | Area% |
|---|---|---|
| 1 | 1.551 | 100.00 |

V609072$7

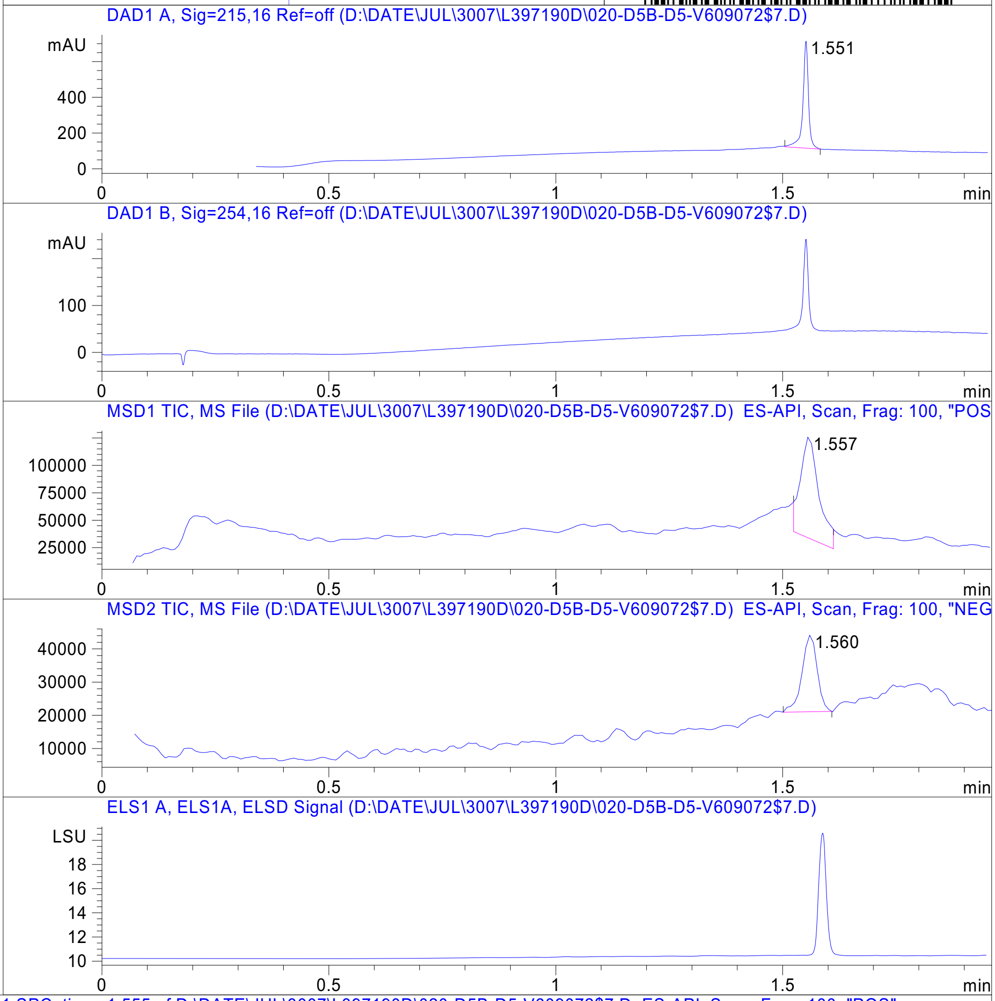


RT 1.557

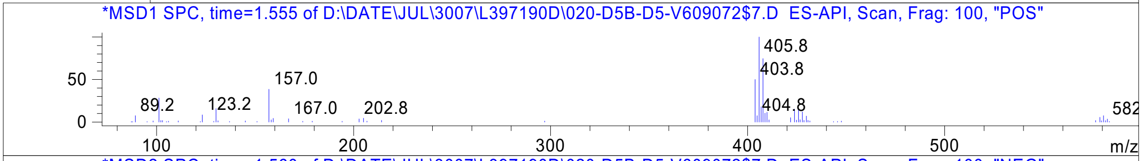


RT 1.560

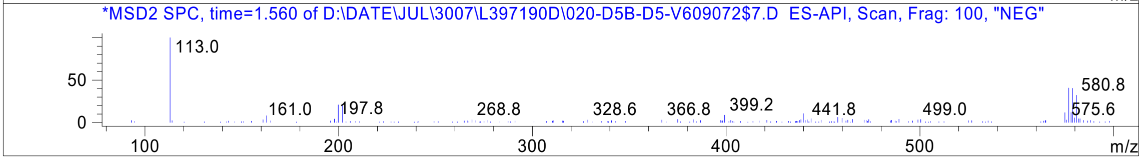




Z5084954217_

MaxPeak: 100.00%
Ret_Time: 1.516 min

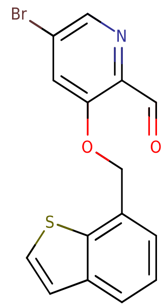


**Mol Wt** 348.21
**Exact Mass** 348.97

| # | Time | Area% |
|---|---|---|
| 1 | 1.516 | 100.00 |

V296340$9

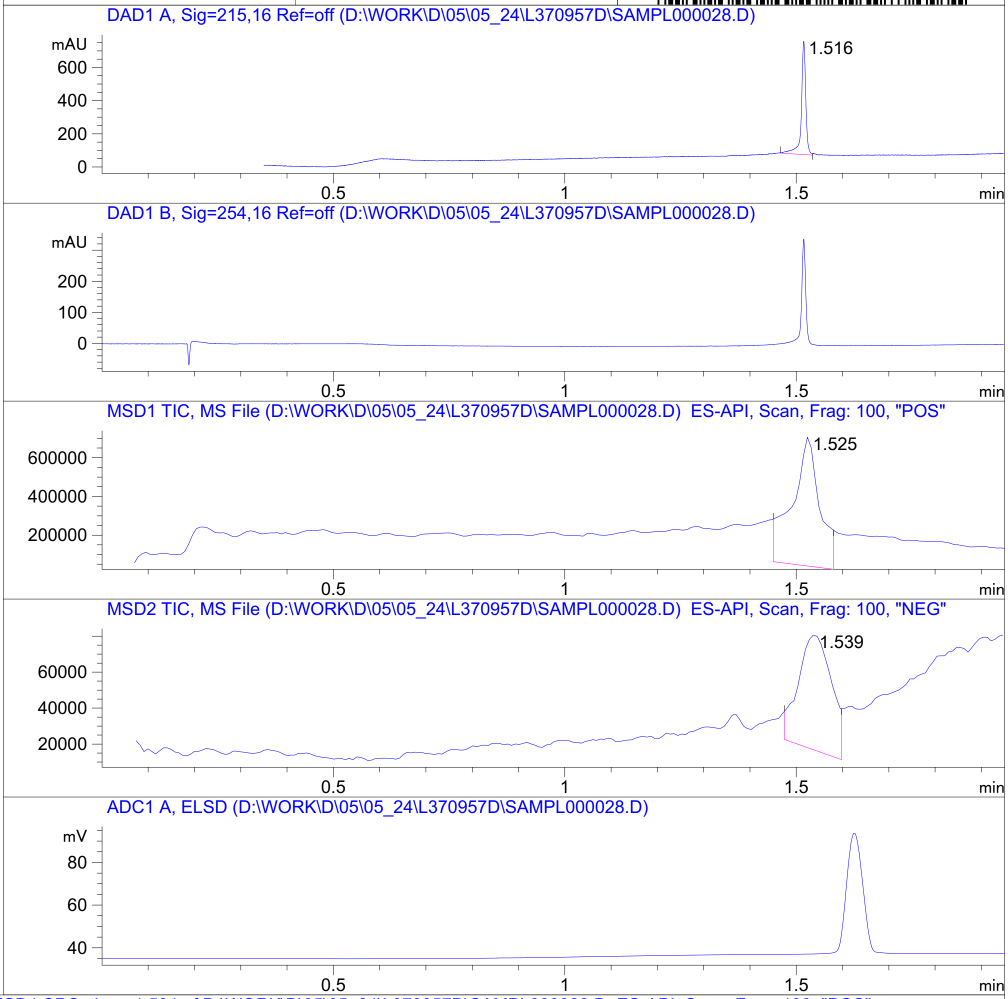


RT 1.525

RT 1.539

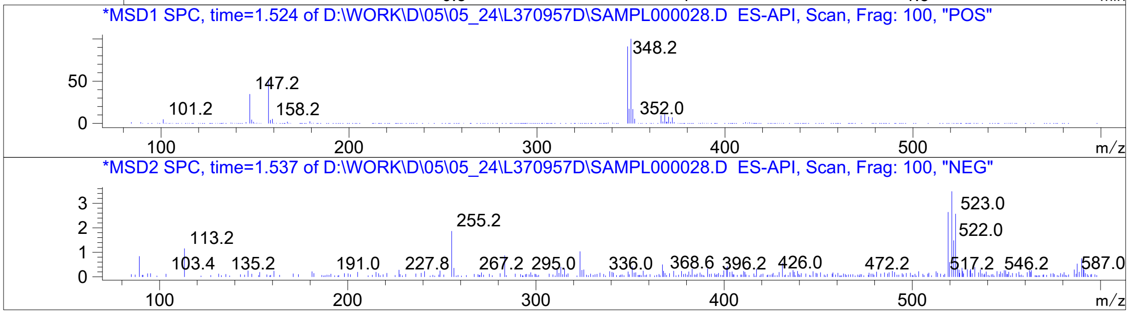




Z4989203252_

MaxPeak: 100.00%
Ret_Time: 1.650 min

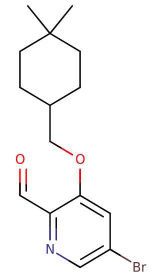

**Mol Wt** 326.23

**Exact Mass** 325.1

| # | Time | Area% |
| --- | --- | --- |
| 1 | 1.650 | 100.00 |

W179753$3

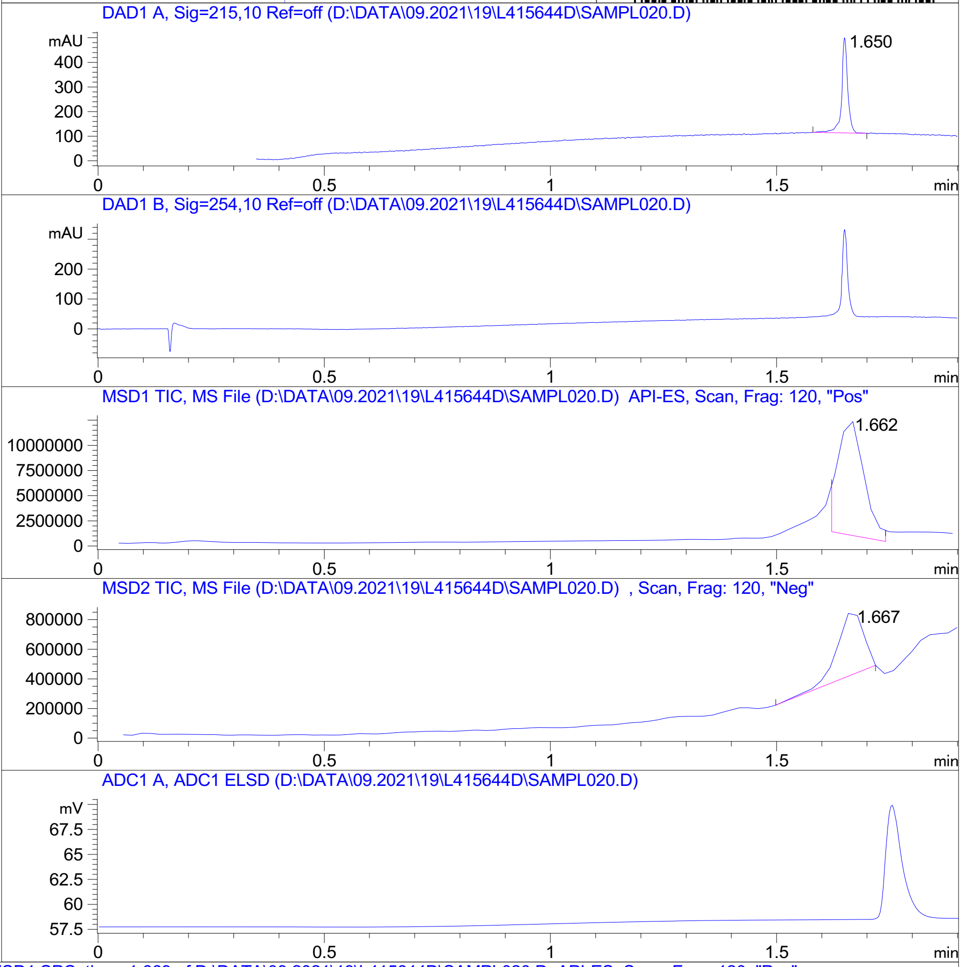


RT 1.662

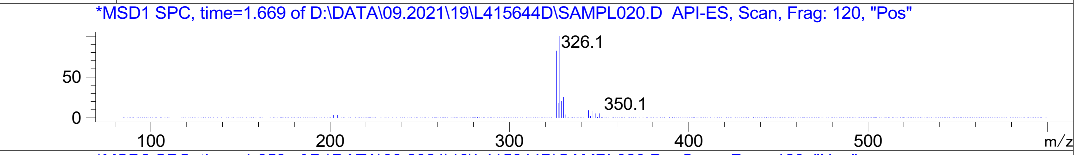


RT 1.667

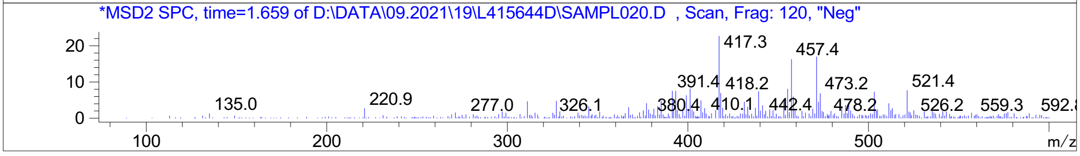


Inj.Date 9/19/2021 CH P2-C-03 -SL- Acq. Method C:\HPCHEM\-> ->

Z5192779009_

MaxPeak: 100.00%
Ret_Time: 1.193 min

# W182833$4

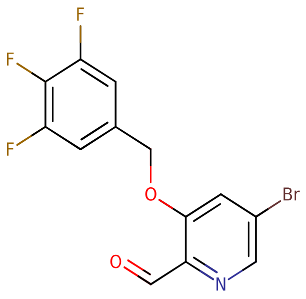


**Mol Wt** 346.1

**Exact Mass** 344.97

| # | Time | Area% |
|---|---|---|
| 1 | 1.193 | 100.00 |

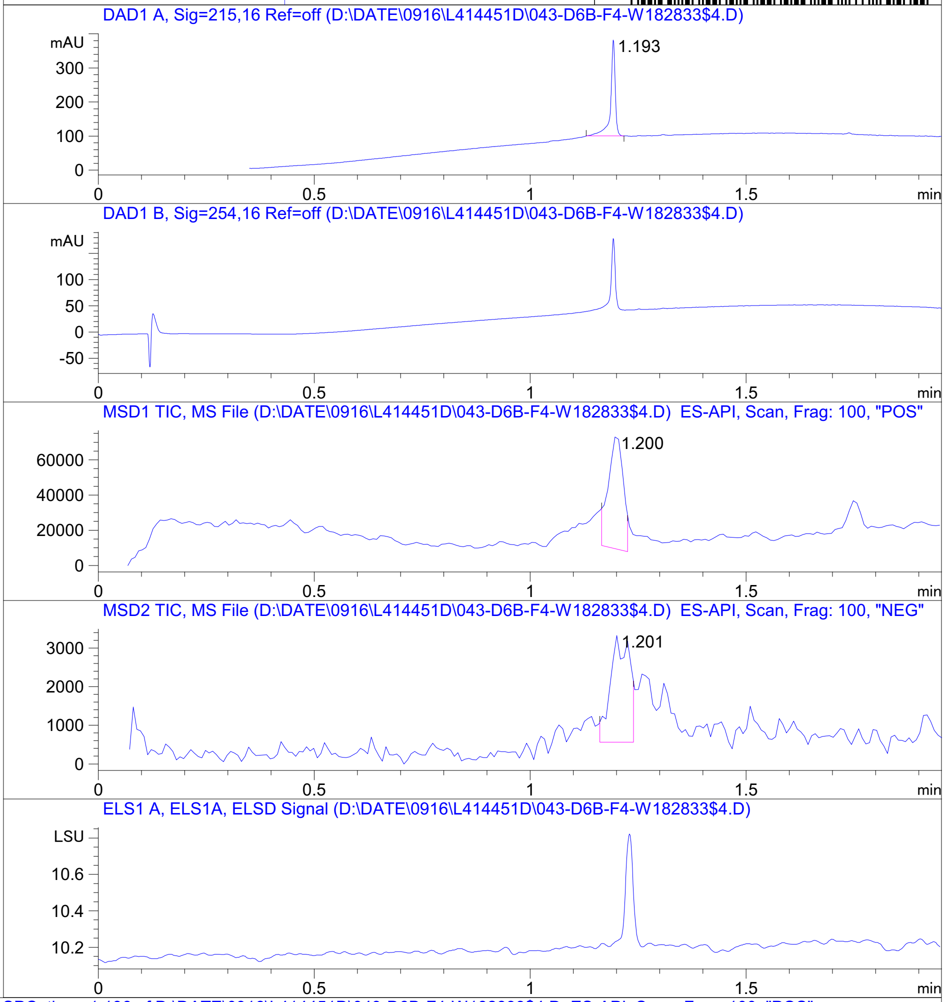


RT 1.200

RT 1.201

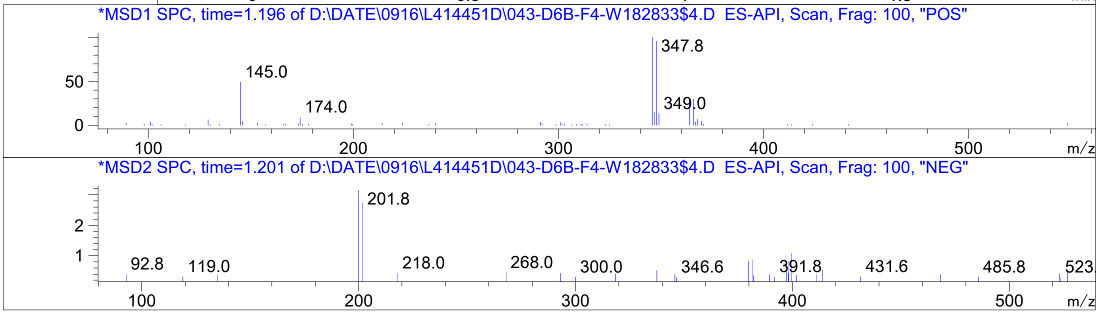


Inj.Date 9/16/2021 E 13 Acq. Method C:\Chem32\-> ->

Z5192779037_

MaxPeak: 98.96%
Ret_Time: 1.146 min

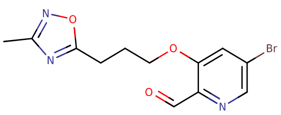


**Mol Wt** 326.15
**Exact Mass** 325.01

| # | Time | Area% |
|---|---|---|
| 1 | 0.909 | 1.04 |
| 2 | 1.146 | 98.96 |

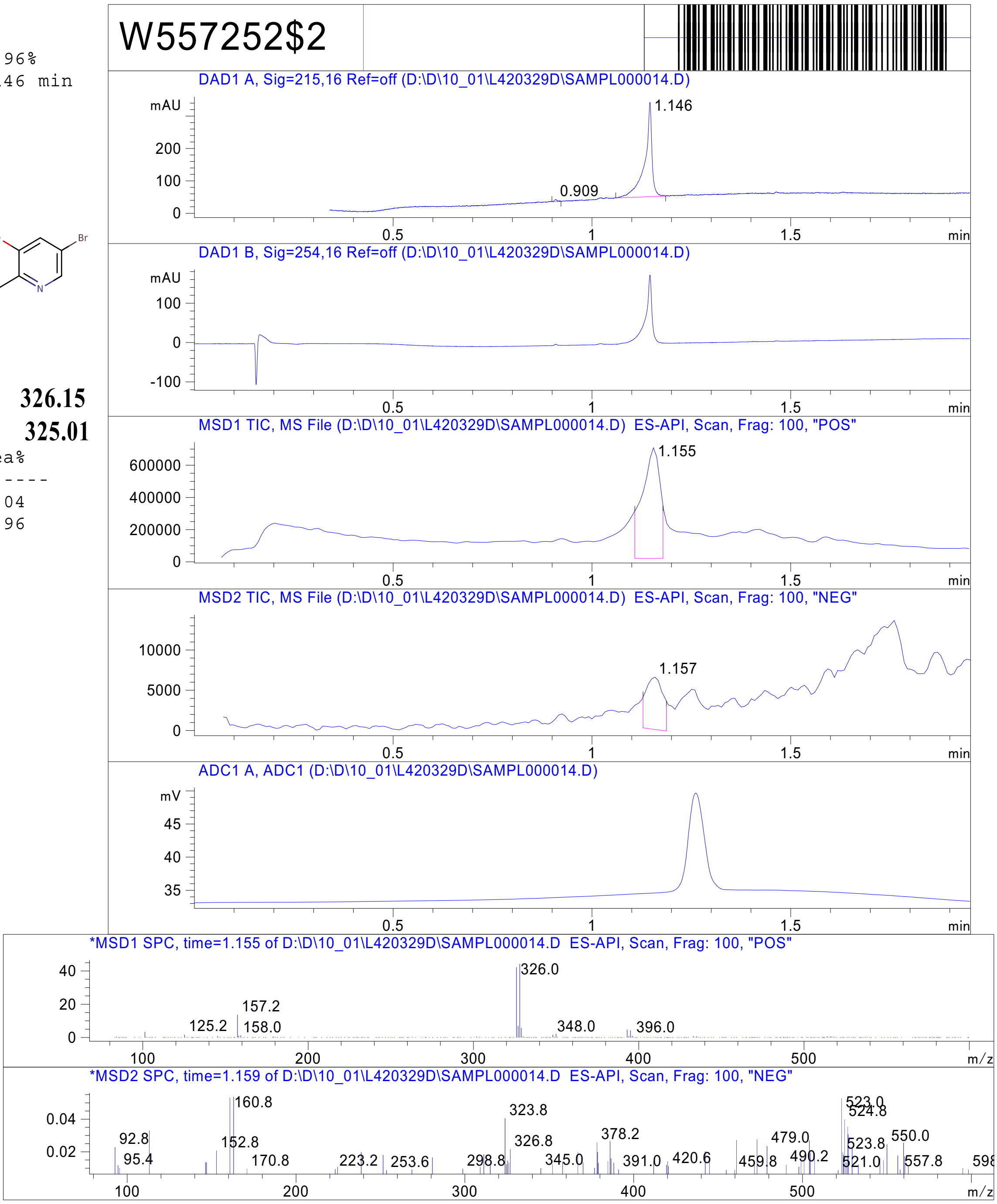


RT 1.155

RT 1.157

Inj.Date 10/1/2021 N - 4 - Acq. Method C:\CHEM32\-> ->

Z2991877909_

W557642$B

MaxPeak: 100.00%
Ret_Time: 1.476 min

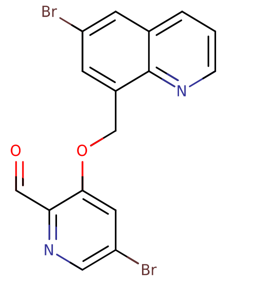


**Mol Wt** 422.07
**Exact Mass** 421.92

| # | Time | Area% |
|---|---|---|
| 1 | 1.476 | 100.00 |

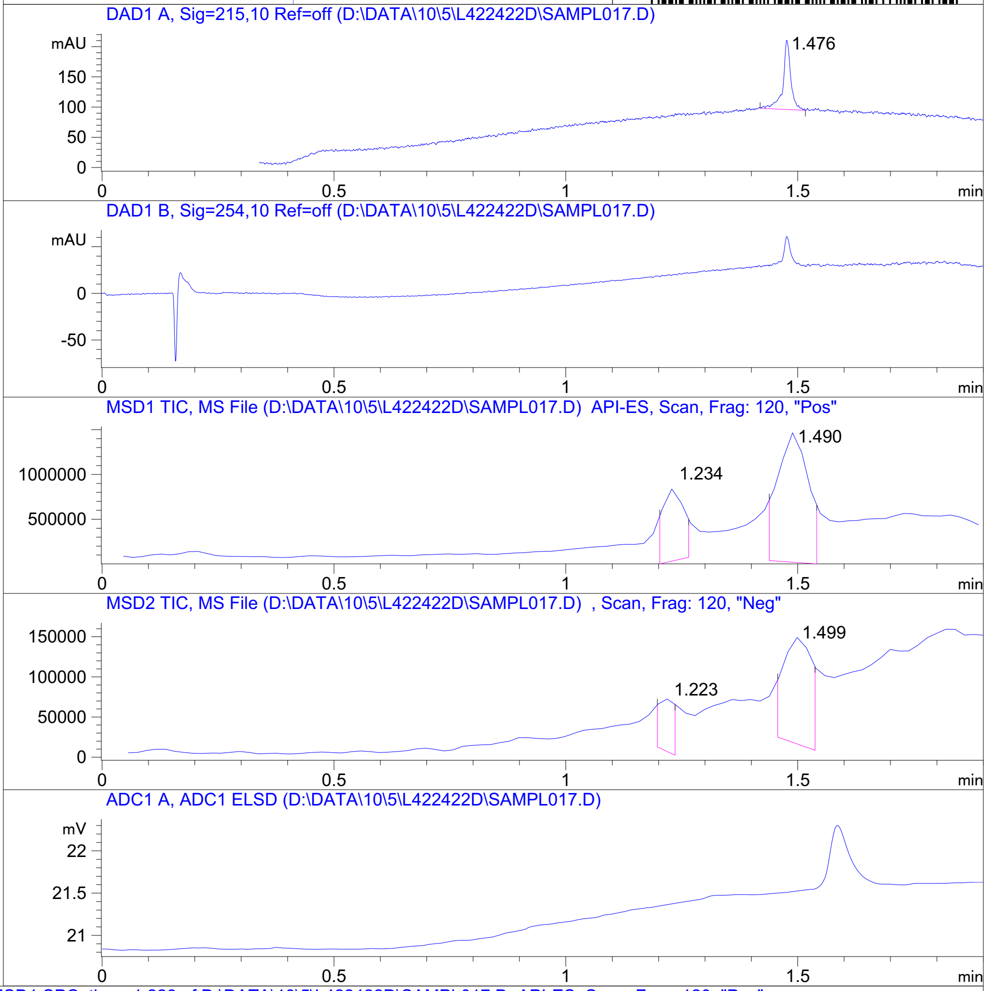


RT 1.234

RT 1.490

RT 1.223

RT 1.499

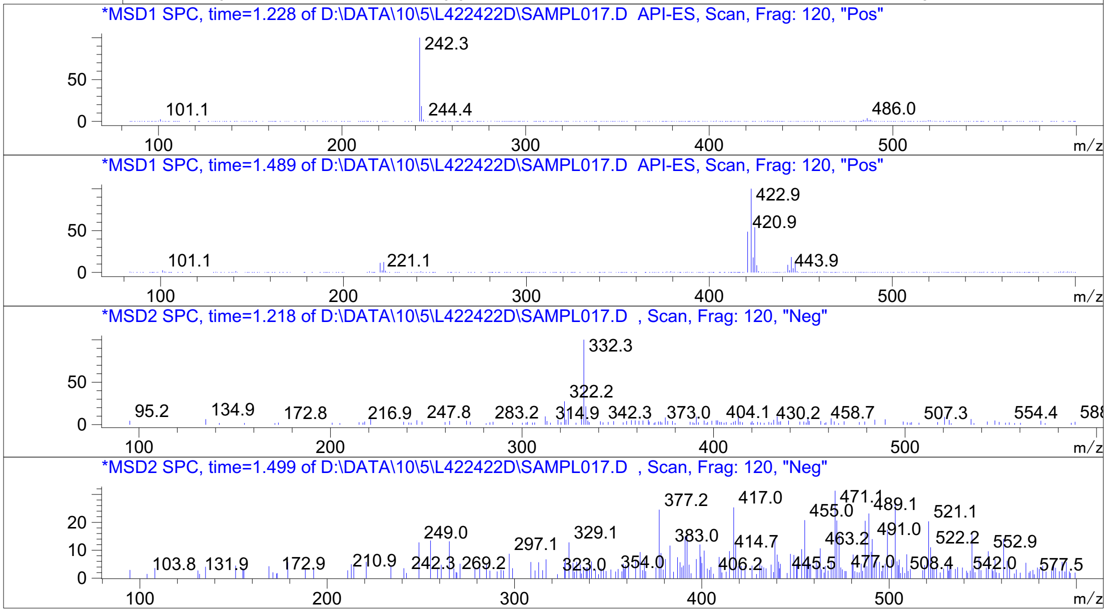


Inj.Date 10/6/2021 LB -SL- Acq. Method C:\HPCHEM\-> ->

Z5192779150_

MaxPeak: 100.00%
Ret_Time: 2.908 min

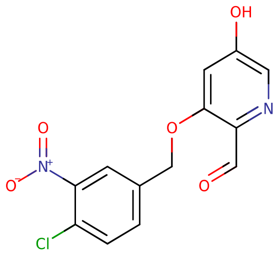


**Mol Wt** 308.67
**Exact Mass** 308.01

| # | Time | Area% |
|---|---|---|
| 1 | 2.908 | 100.00 |

W656530$2

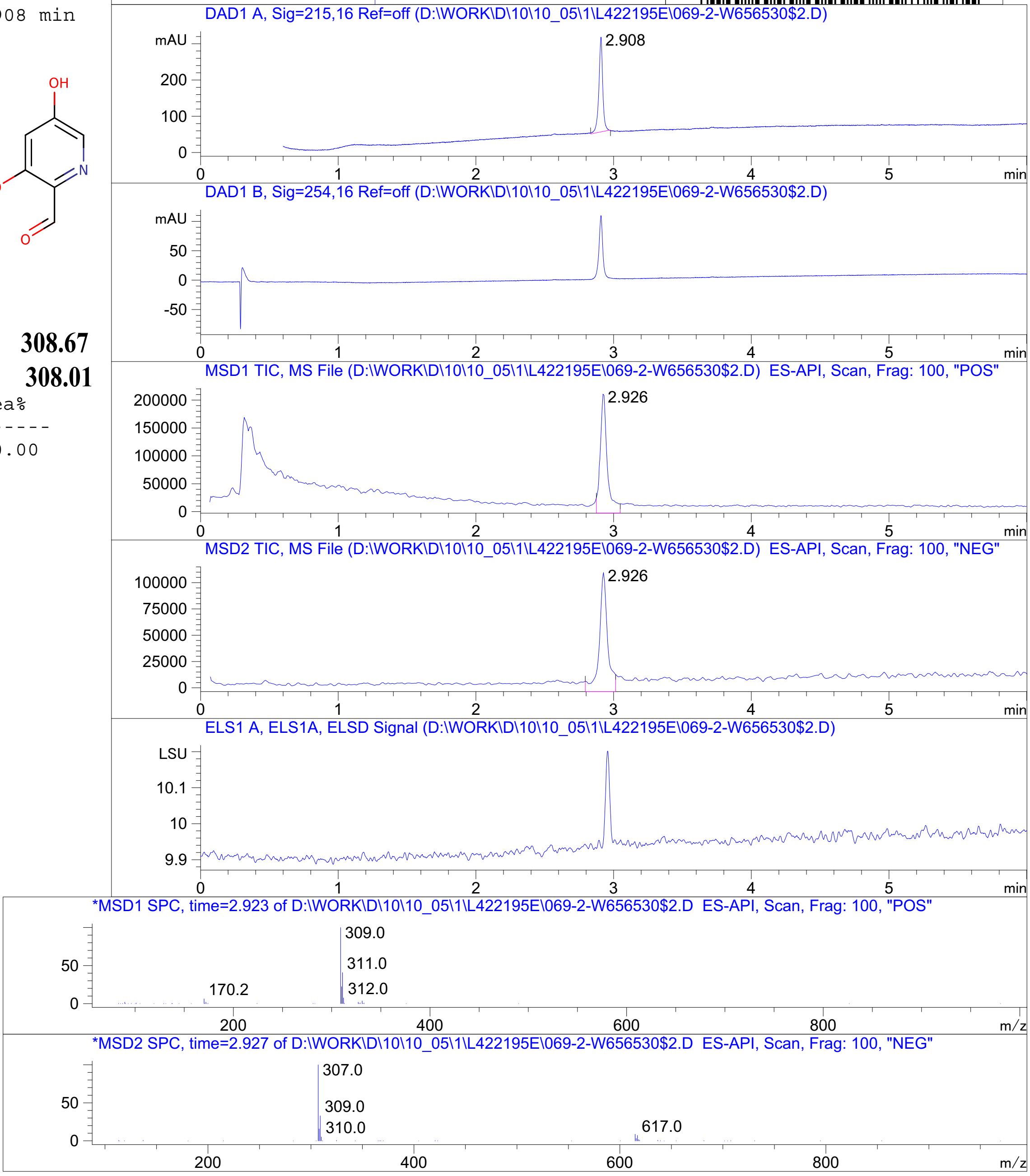


RT 2.926

RT 2.926

Inj.Date 10/5/2021 K Vial 2 -18- Acq. Method C:\Chem32\-> ->

Z4954569176_

W847046$2

MaxPeak: 100.00%
Ret_Time: 1.326 min

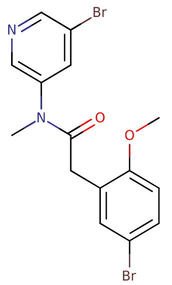


**Mol Wt** 414.09
**Exact Mass** 413.96

| # | Time | Area% |
|---|---|---|
| 1 | 1.326 | 100.00 |

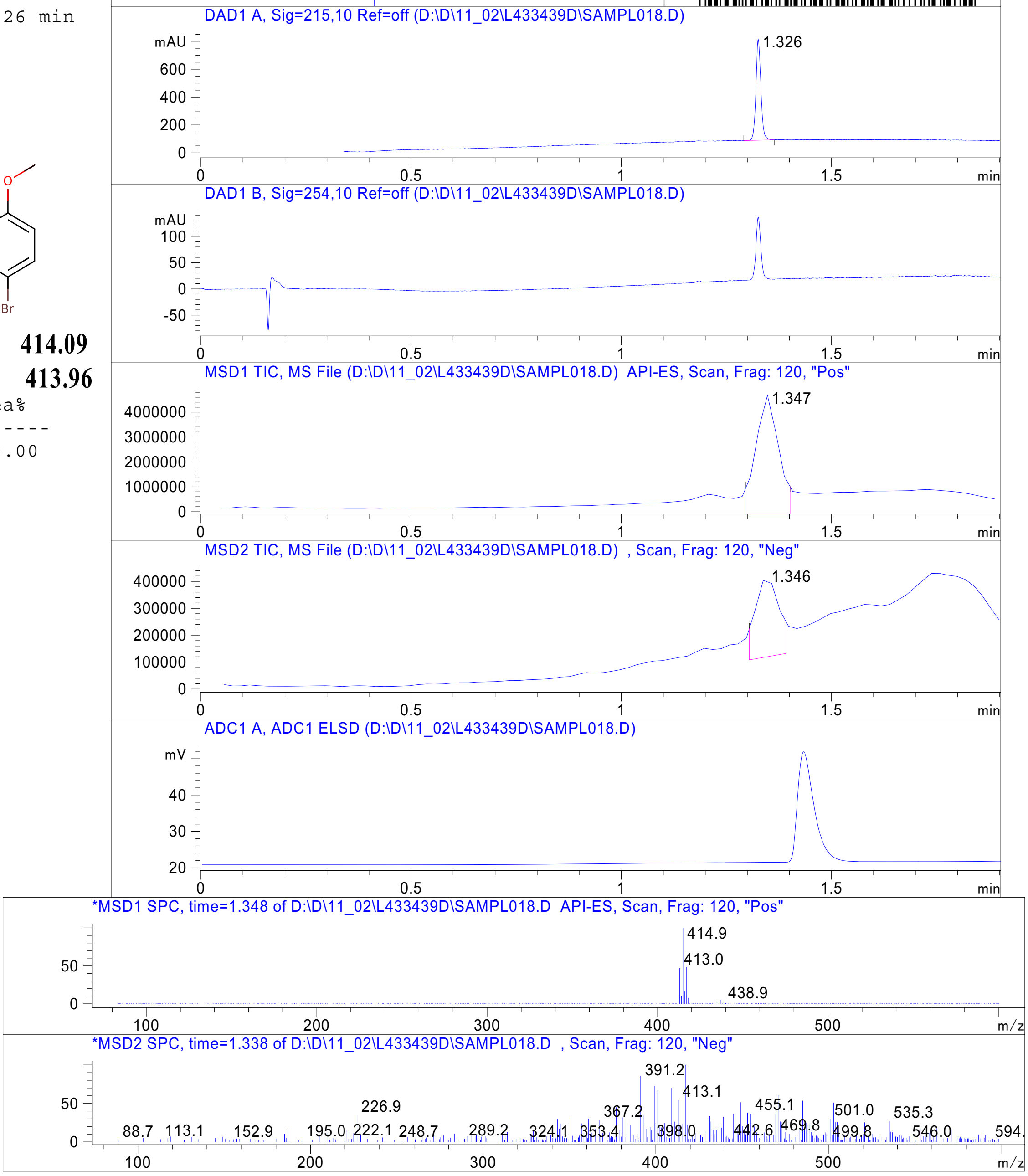


RT 1.347

RT 1.346

Inj.Date 11/2/2021 N -SL- Acq. Method C:\HPCHEM\-> ->

Z5385490967_

MaxPeak: 100.00%
Ret_Time: 1.134 min

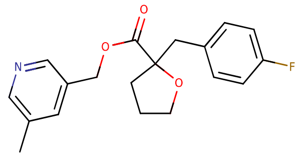

**Mol Wt** 329.36
**Exact Mass** 329.17

| # | Time | Area% |
|---|---|---|
| 1 | 1.134 | 100.00 |

W847028$3

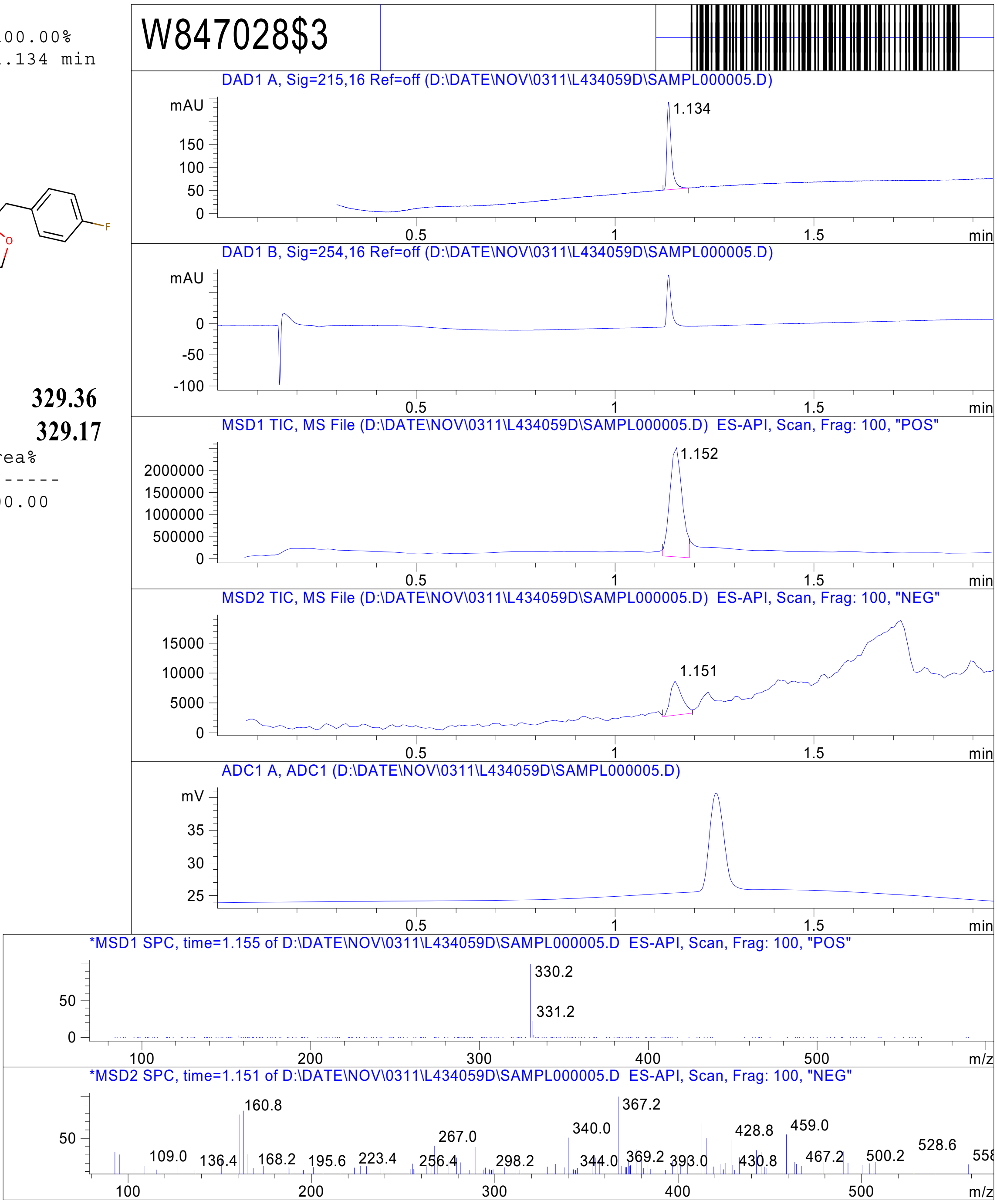


RT 1.152

RT 1.151

Inj.Date 11/3/2021 Y - 4 - Acq. Method C:\CHEM32\-> ->

Z3382155230_

MaxPeak: 100.00%
Ret_Time: 1.373 min

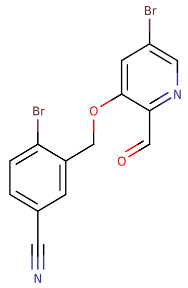


**Mol Wt** 396.03
**Exact Mass** 395.9

| # | Time | Area% |
|---|---|---|
| 1 | 1.373 | 100.00 |

# W753440$1

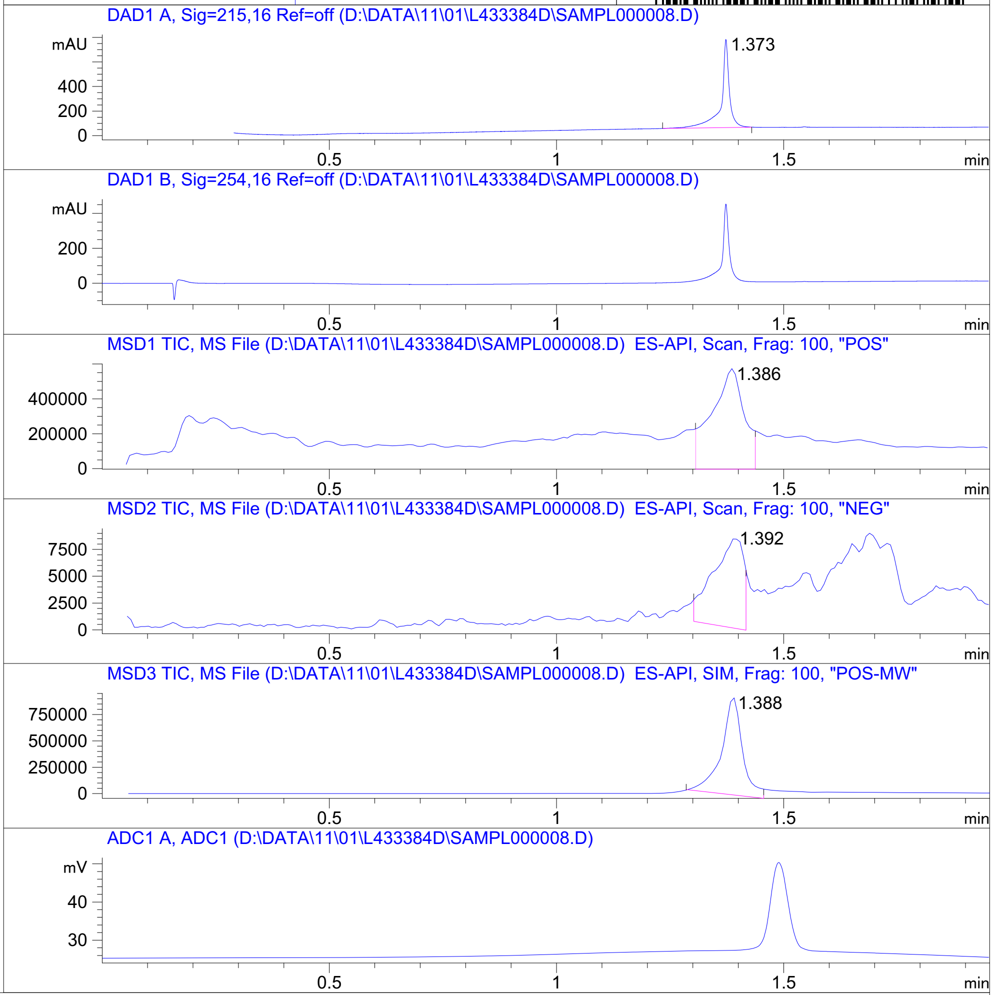


RT 1.386

RT 1.392

RT 1.388

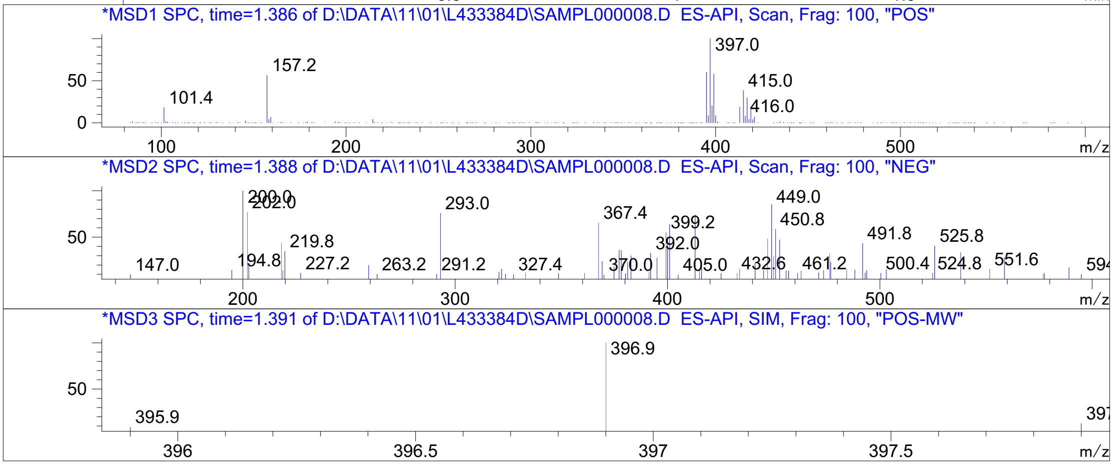




Z5009000158_

MaxPeak: 100.00%
Ret_Time: 0.976 min

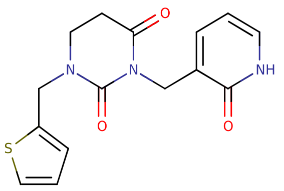


**Mol Wt** 317.36
**Exact Mass** 317.09

| # | Time | Area% |
|---|---|---|
| 1 | 0.976 | 100.00 |

W847029$1

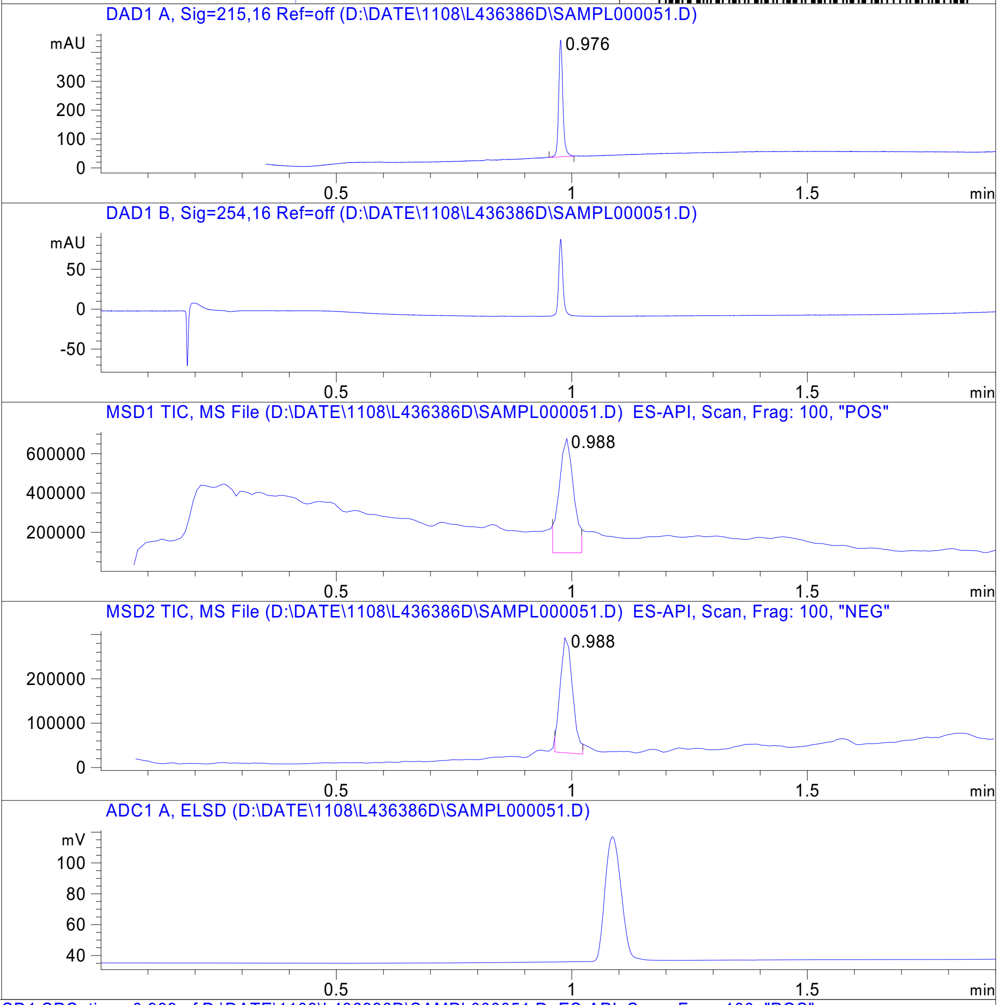


RT 0.988

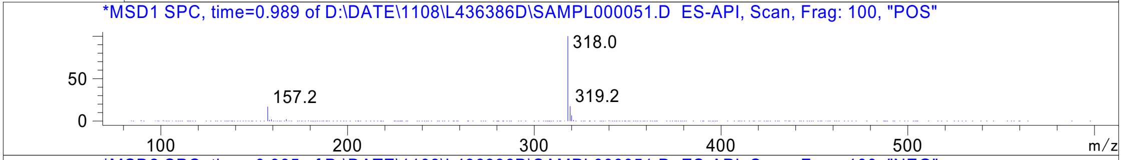


RT 0.988

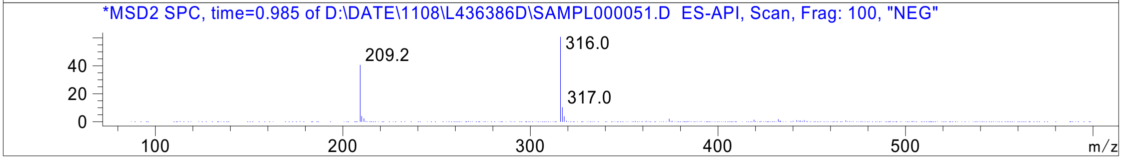




Z3535317212_

MaxPeak: 92.20%
Ret_Time: 0.990 min

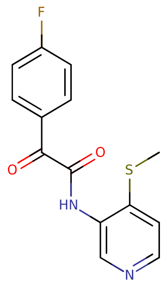


**Mol Wt** 290.31
**Exact Mass** 290.06

| # | Time | Area% |
|---|---|---|
| 1 | 0.852 | 7.80 |
| 2 | 0.990 | 92.20 |

W980739$2

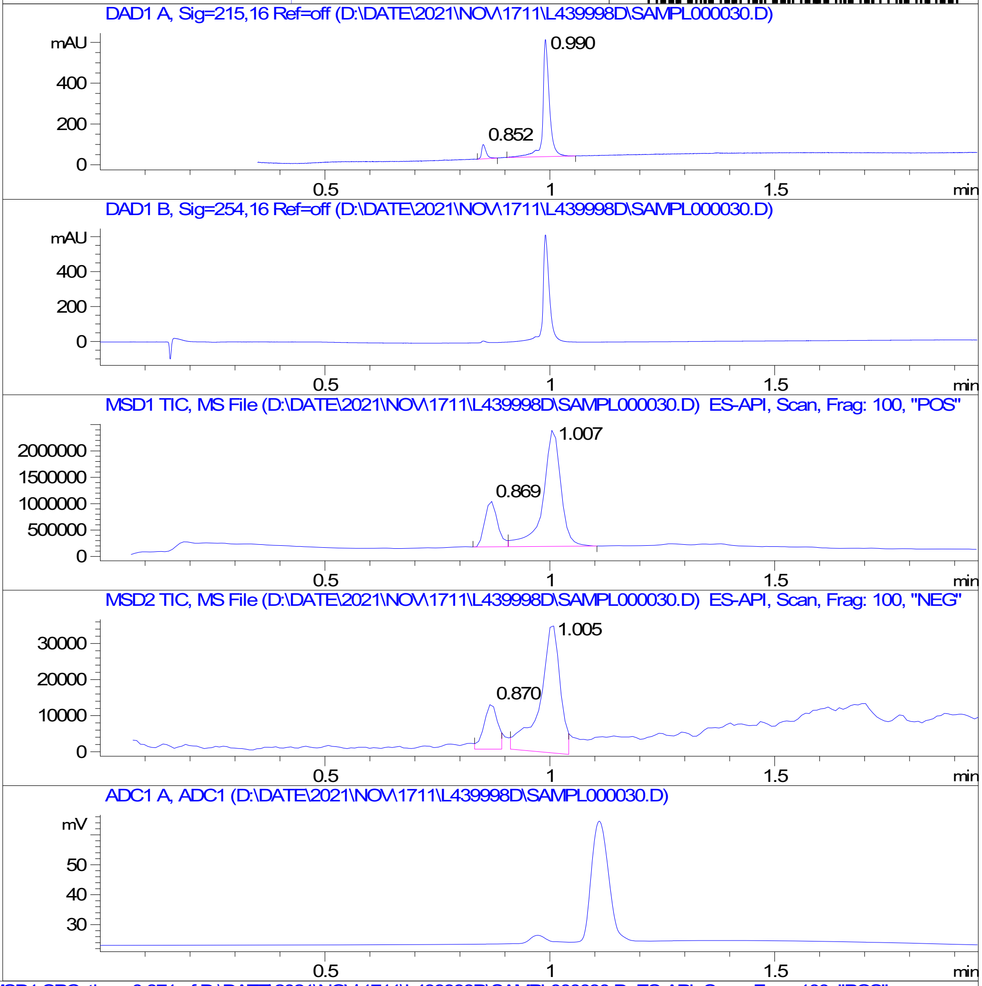


RT 0.869

RT 1.007

RT 0.870

RT 1.005

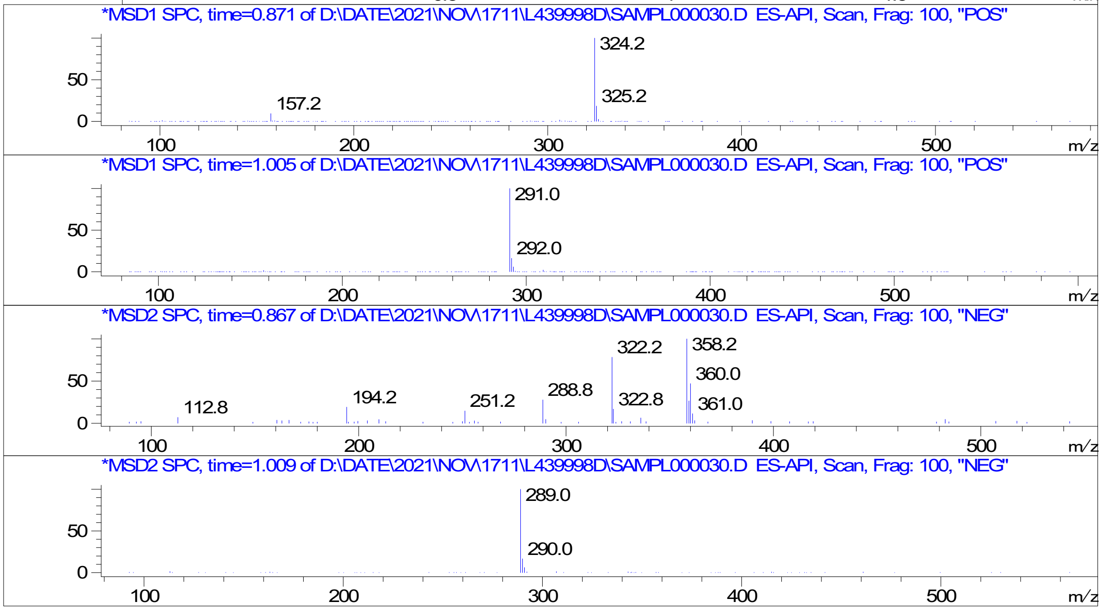


Inj.Date 17-Nov-21 A P2-D-03 - 4 - Acq. Method C:\CHEM32\-> ->

Z1669286714_

MaxPeak: 97.88%
Ret_Time: 1.052 min

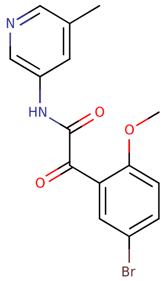


**Mol Wt** 349.18

**Exact Mass** 348.02

| # | Time | Area% |
|---|---|---|
| 1 | 1.052 | 97.88 |
| 2 | 1.180 | 2.12 |

W980702$6

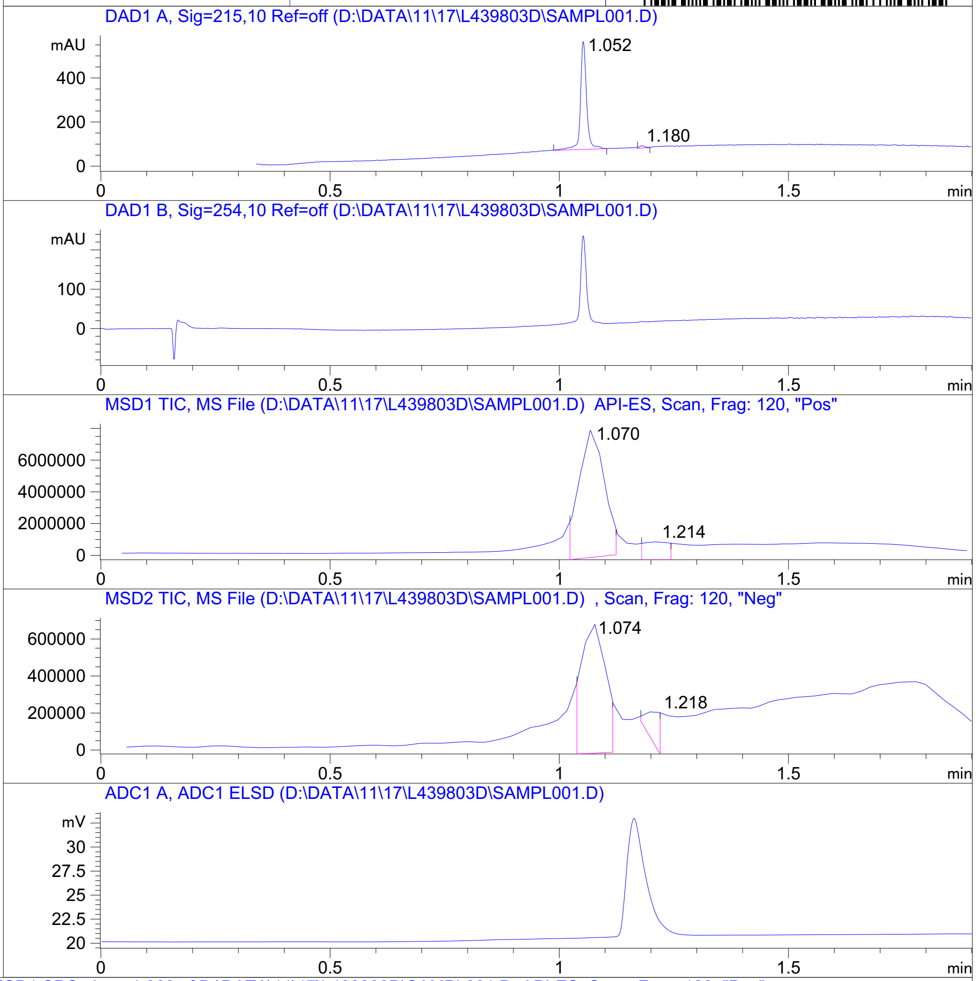


RT 1.070

RT 1.214

RT 1.074

RT 1.218

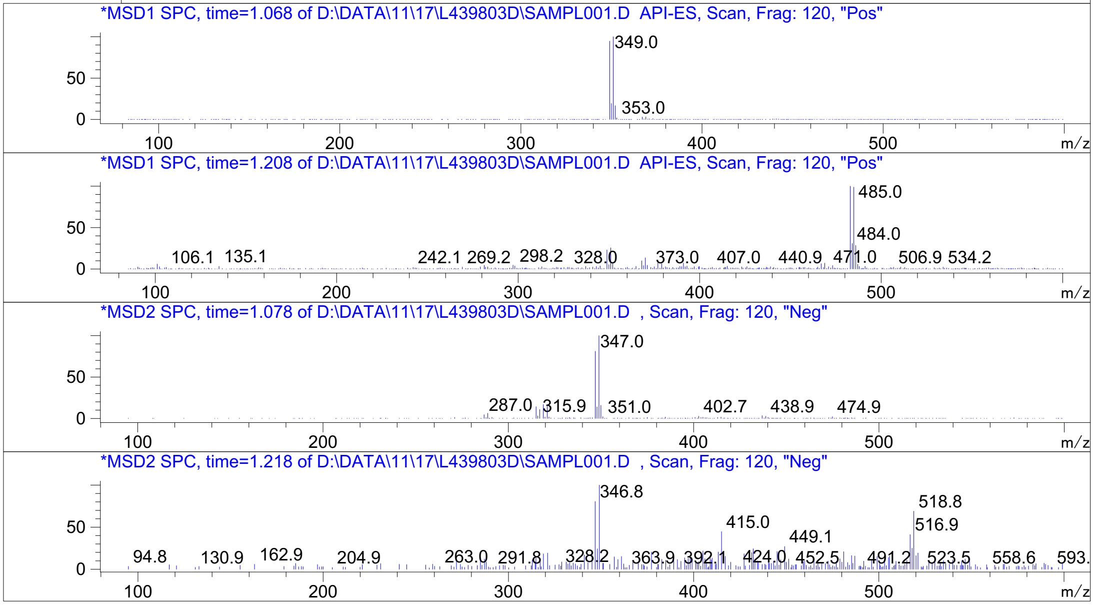


Inj.Date 11/17/2021 LB -SL- Acq. Method C:\HPCHEM\-> ->

Z1355254448_

MaxPeak: 100.00%
Ret_Time: 1.368 min

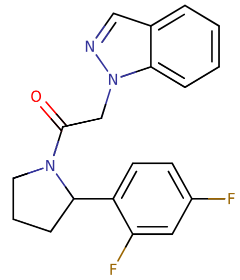


**Mol Wt** 341.36
**Exact Mass** 341.16

| # | Time | Area% |
|---|---|---|
| 1 | 1.368 | 100.00 |

W980607$1

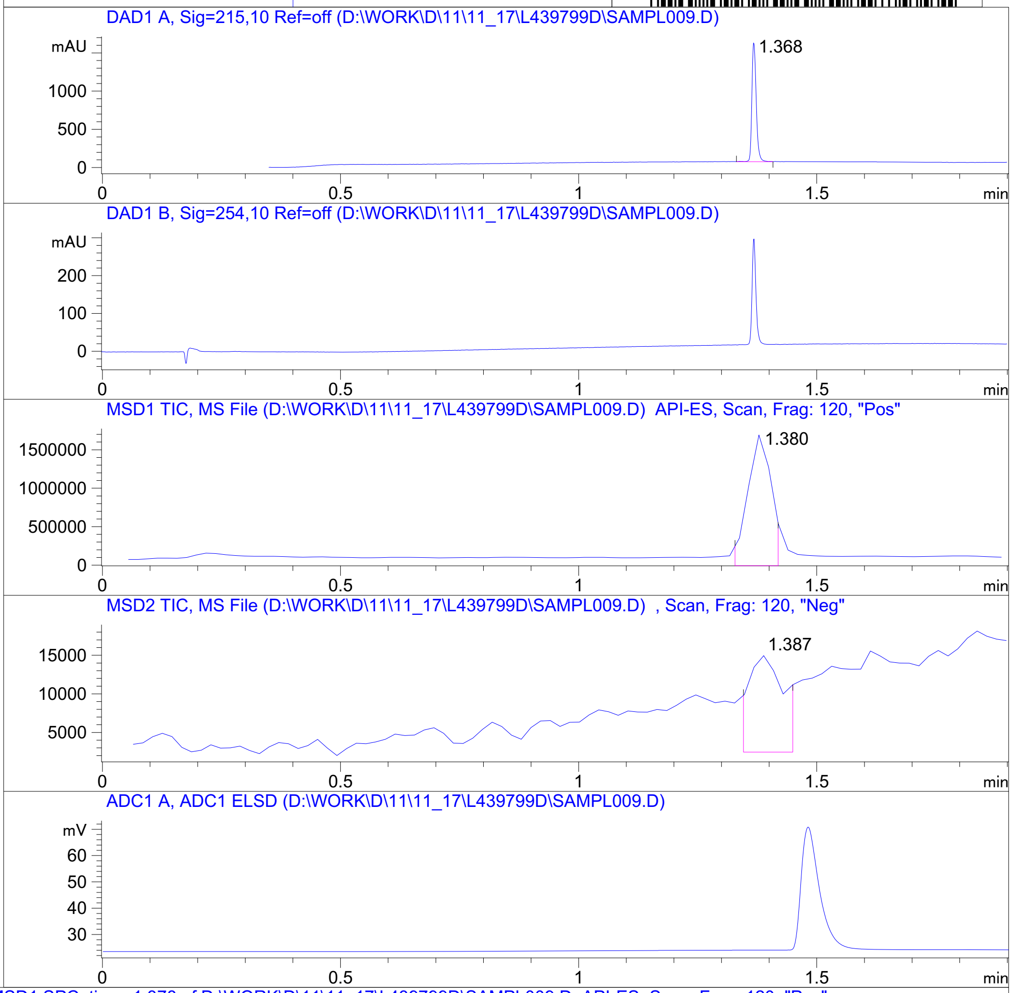


RT 1.380

RT 1.387

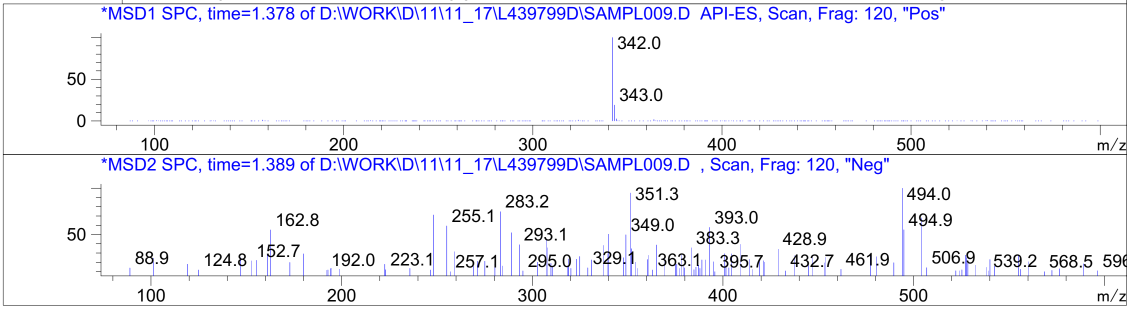


Inj.Date 11/17/2021 K P2-A-09 -VL- Acq. Method C:\HPCHEM\-> ->

Z22988060 57_

MaxPeak: 100.00%
Ret_Time: 1.258 min

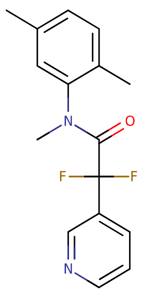

**Mol Wt** 290.31
**Exact Mass** 290.15

| # | Time | Area% |
|---|---|---|
| 1 | 1.258 | 100.00 |

W980611$2

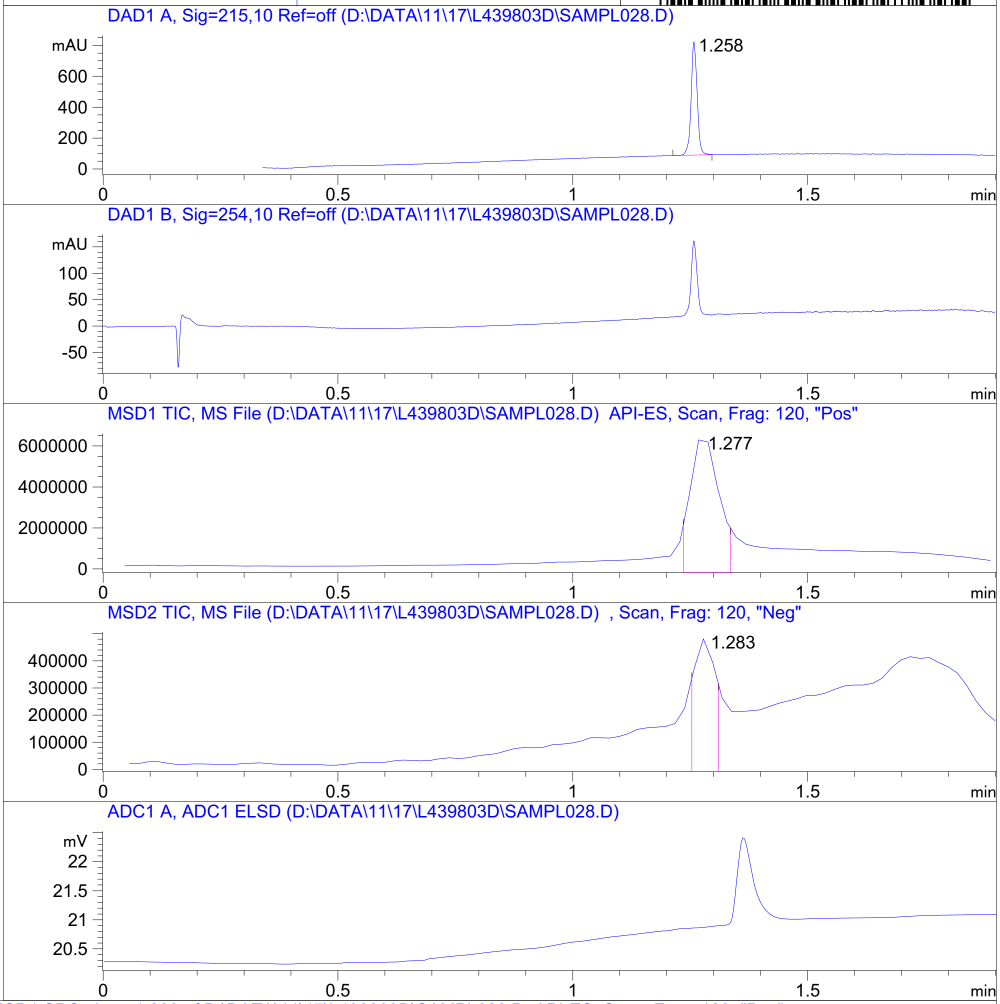


RT 1.277

RT 1.283

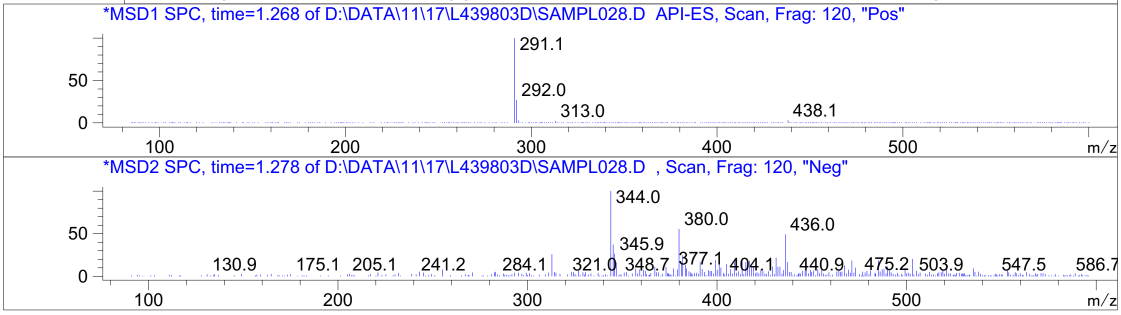


Inj.Date 11/17/2021 LB -SL- Acq. Method C:\HPCHEM\-> ->

Z3079159560_

MaxPeak: 100.00%
Ret_Time: 1.369 min

# W980703$1

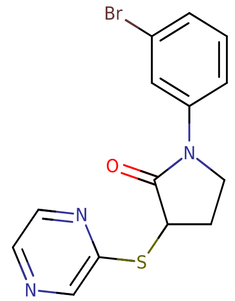


**Mol Wt** 350.23

**Exact Mass** 351

| # | Time | Area% |
|---|---|---|
| 1 | 1.369 | 100.00 |

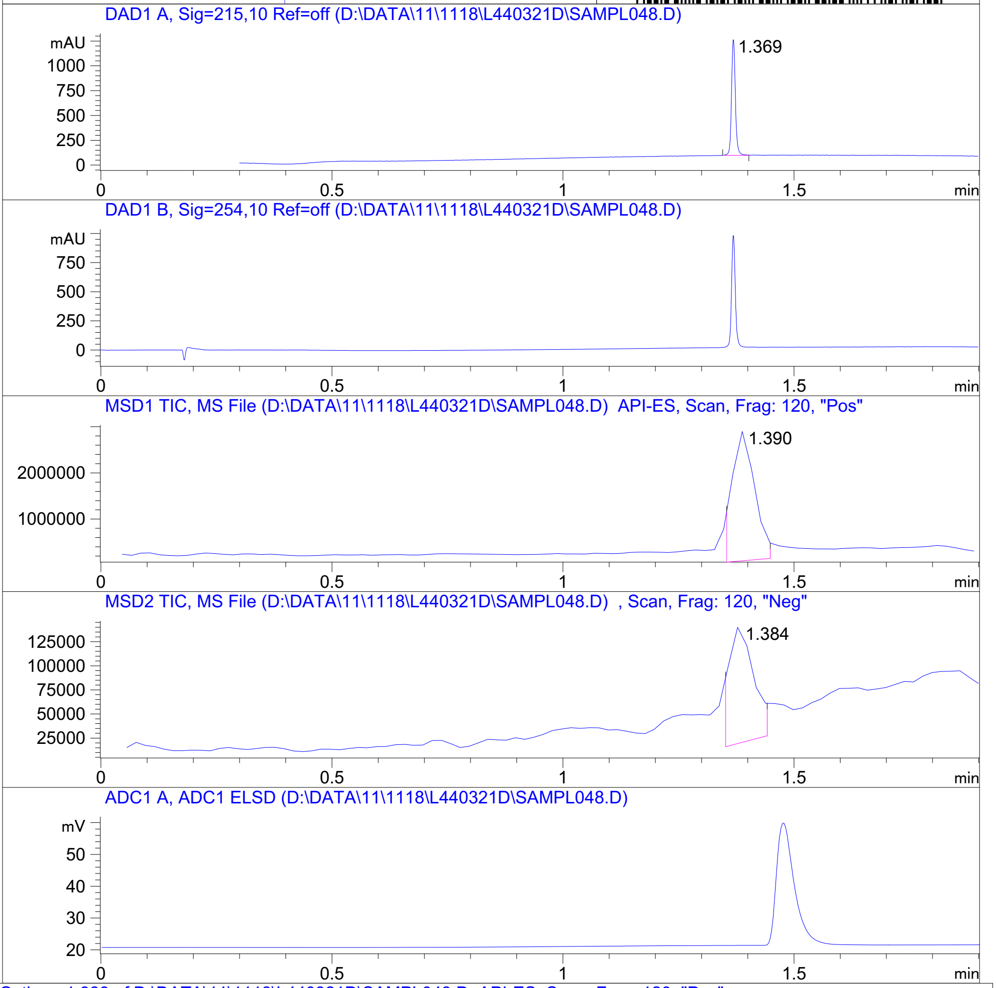


RT 1.390

RT 1.384

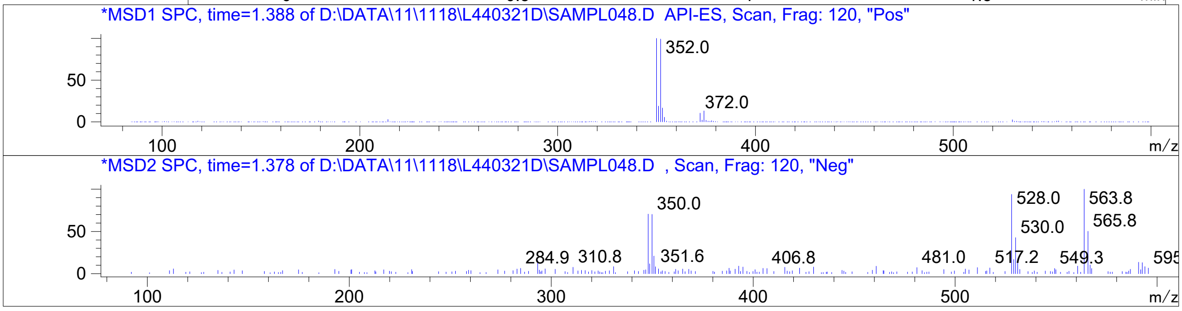


Inj.Date 11/18/2021 LT -SL- Acq. Method C:\HPCHEM\-> ->

Z1872042935_

MaxPeak: 100.00%
Ret_Time: 1.079 min

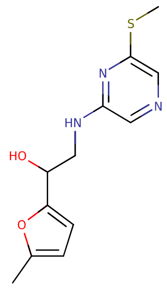


**Mol Wt** 265.33
**Exact Mass** 265.1

| # | Time | Area% |
|---|---|---|
| 1 | 1.079 | 100.00 |

W980679$3

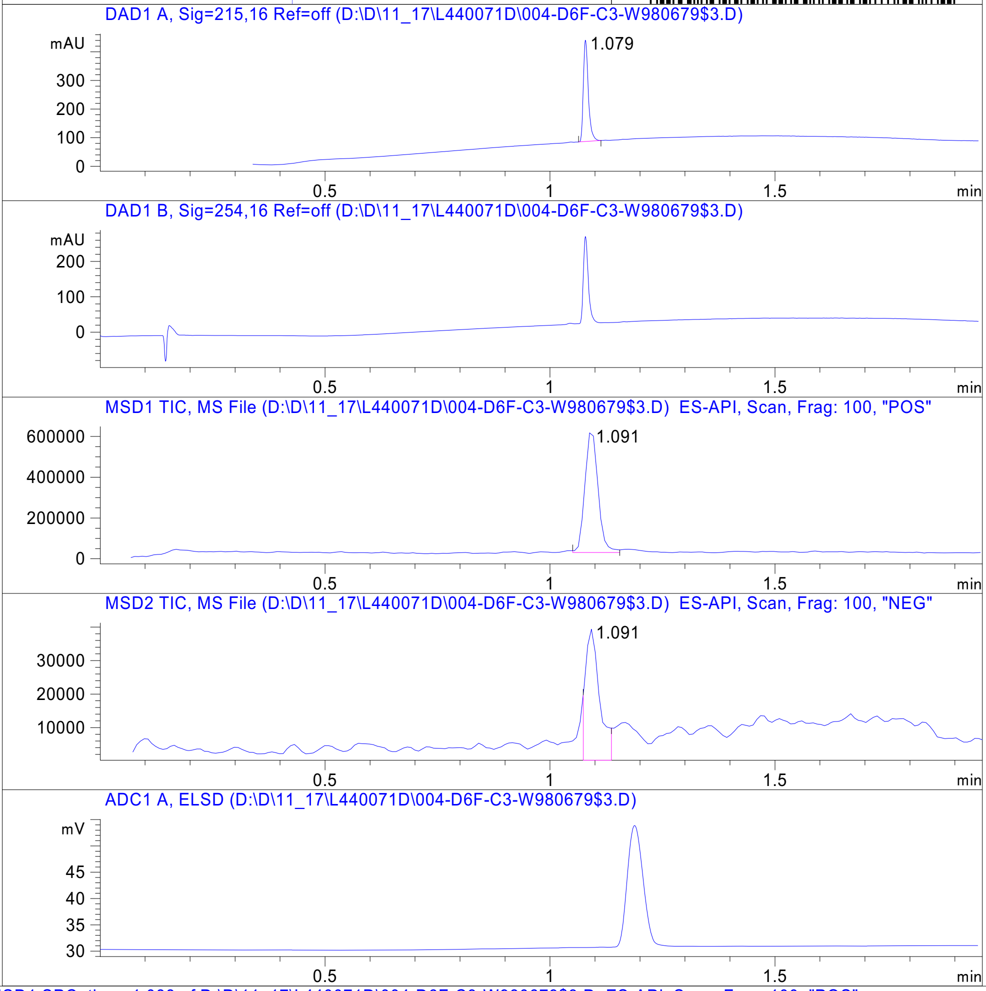


RT 1.091

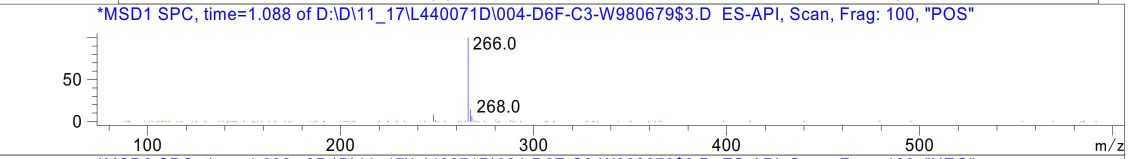


RT 1.091

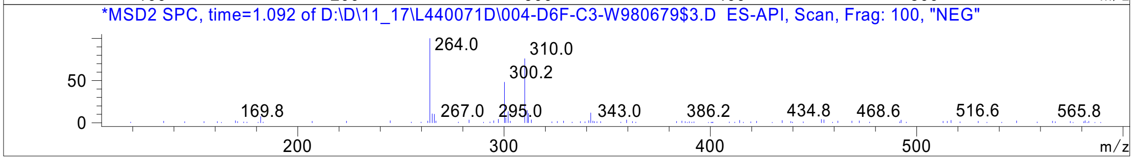




Z1716270280_

MaxPeak: 98.67%
Ret_Time: 0.847 min

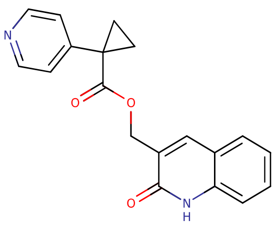

# W980675$3

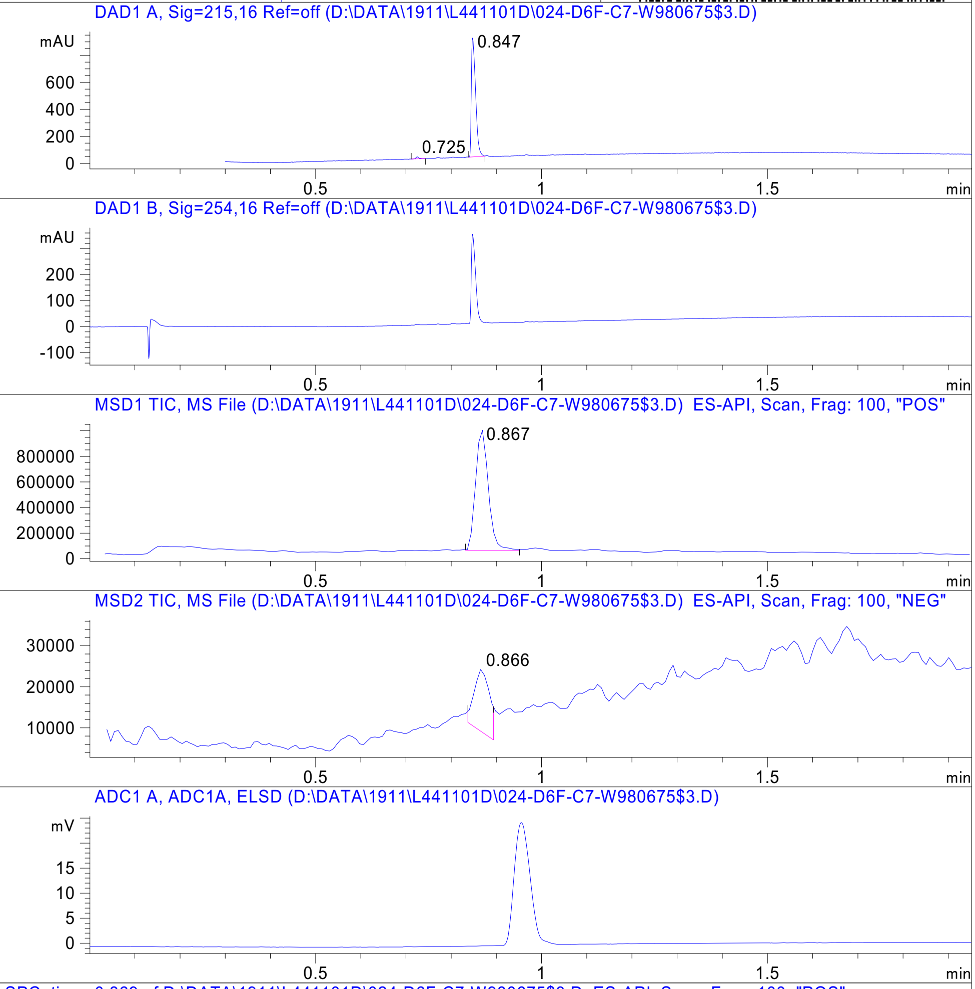


**Mol Wt** 320.34
**Exact Mass** 320.13

| # | Time | Area% |
|---|---|---|
| 1 | 0.725 | 1.33 |
| 2 | 0.847 | 98.67 |

RT 0.867

RT 0.866

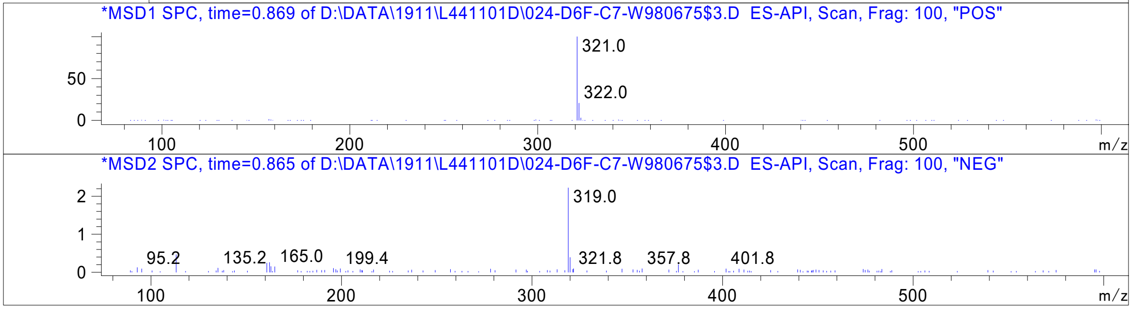




Z5420228488_

MaxPeak: 100.00%
Ret_Time: 1.150 min

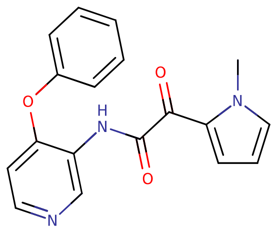


**Mol Wt** 321.33

**Exact Mass** 321.12

| # | Time | Area% |
|---|---|---|
| 1 | 1.150 | 100.00 |

W980606$8

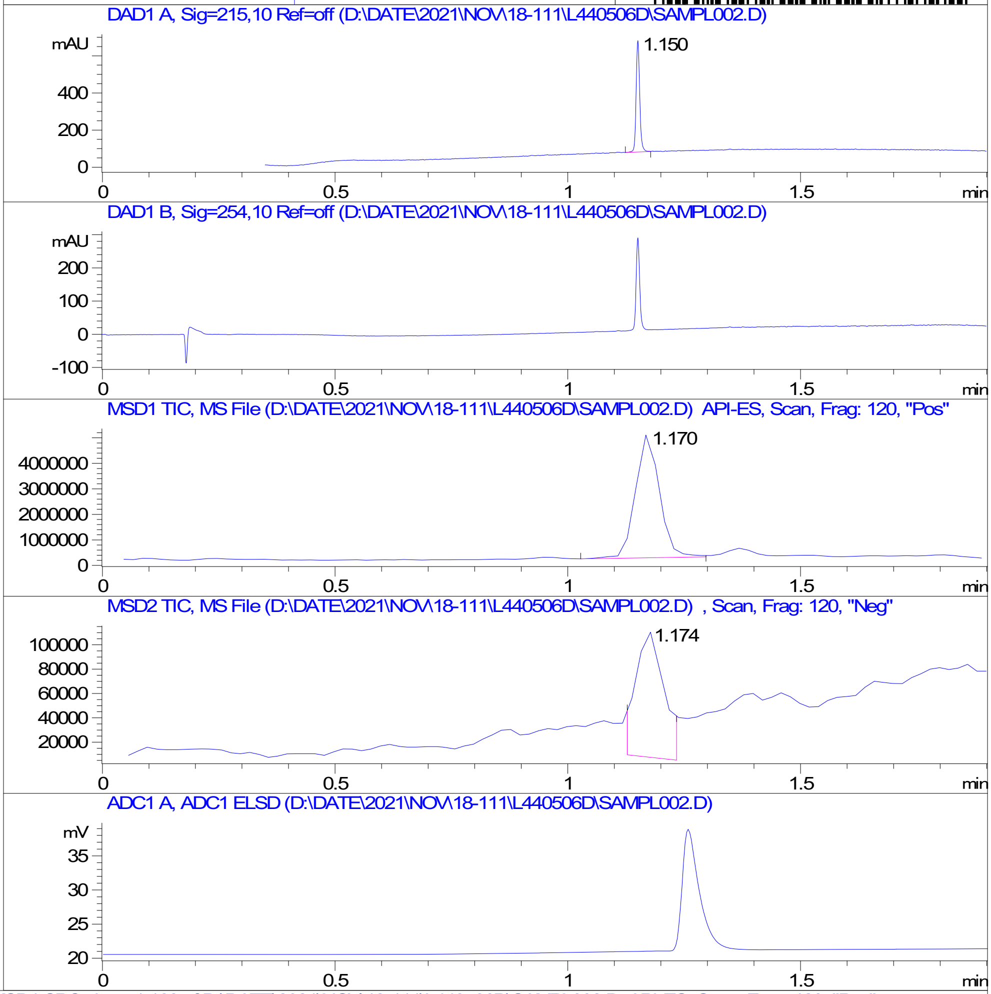


RT 1.170

RT 1.174

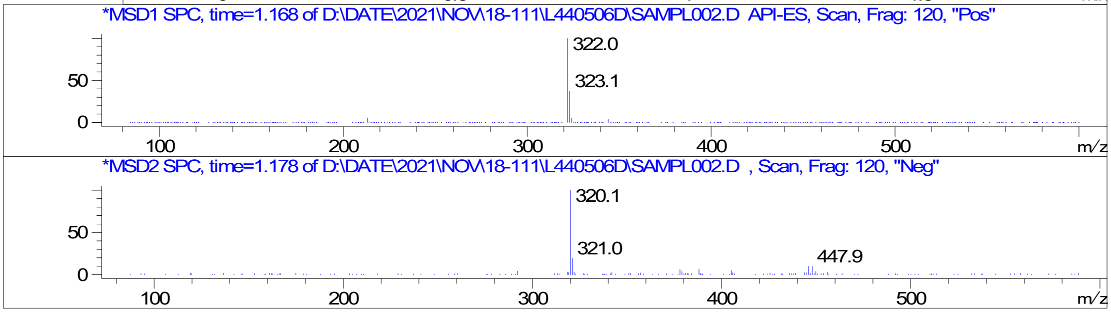


Inj.Date 18-Nov-21 A P2-A-02 -SL- Acq. Method C:\HPCHEM\-> ->

Z2195811405_

MaxPeak: 100.00%
Ret_Time: 1.363 min

# W980620$3

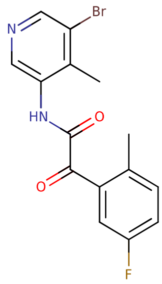


**Mol Wt** 351.17
**Exact Mass** 350.02

| # | Time | Area% |
|---|---|---|
| 1 | 1.363 | 100.00 |

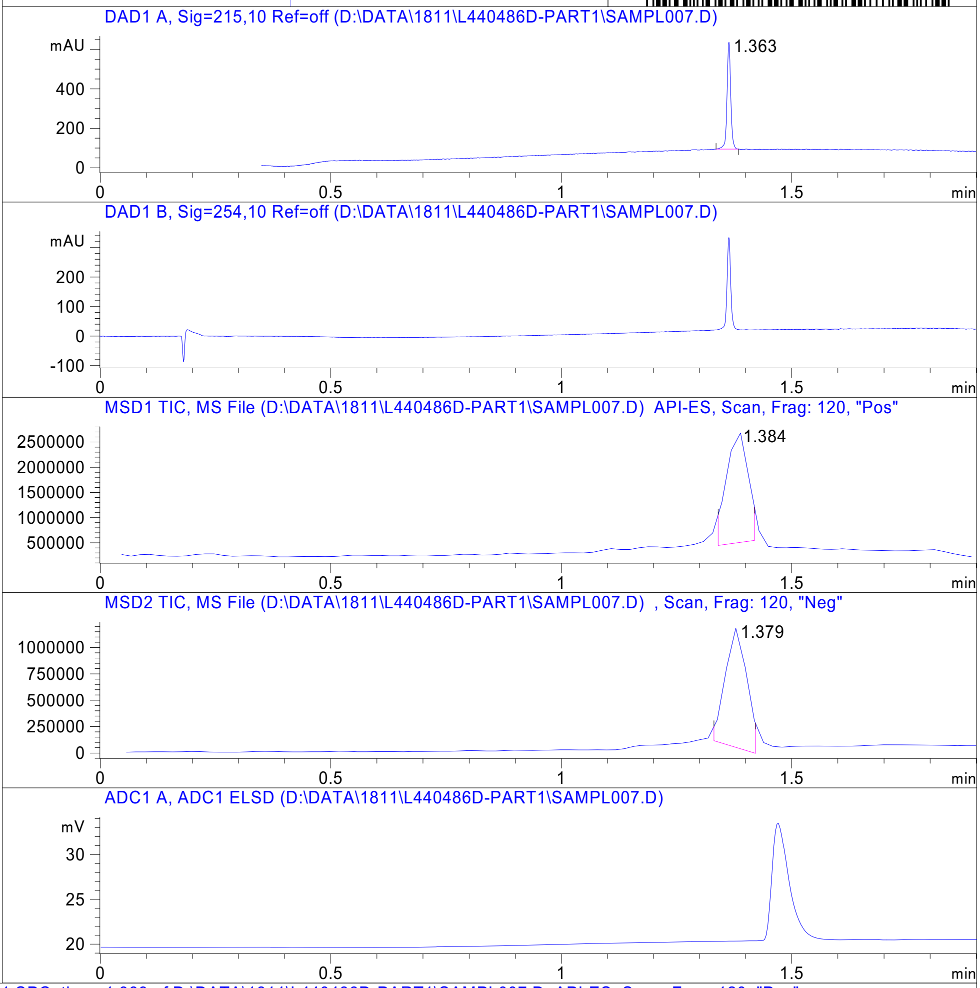


RT 1.384

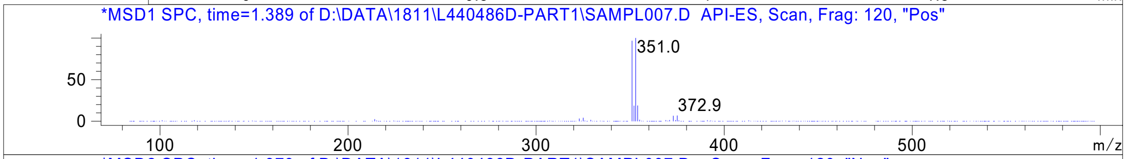


RT 1.379

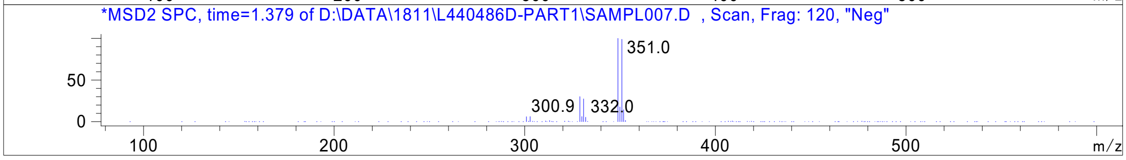


Inj.Date 11/18/2021 H -SL- Acq. Method C:\HPCHEM\-> ->

Z5420225795_

MaxPeak: 100.00%
Ret_Time: 0.876 min

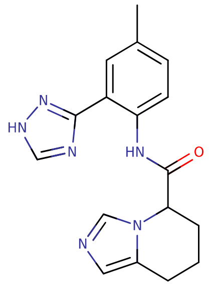


**Mol Wt** 322.36
**Exact Mass** 322.17

| # | Time | Area% |
|---|---|---|
| 1 | 0.876 | 100.00 |

W980749$4

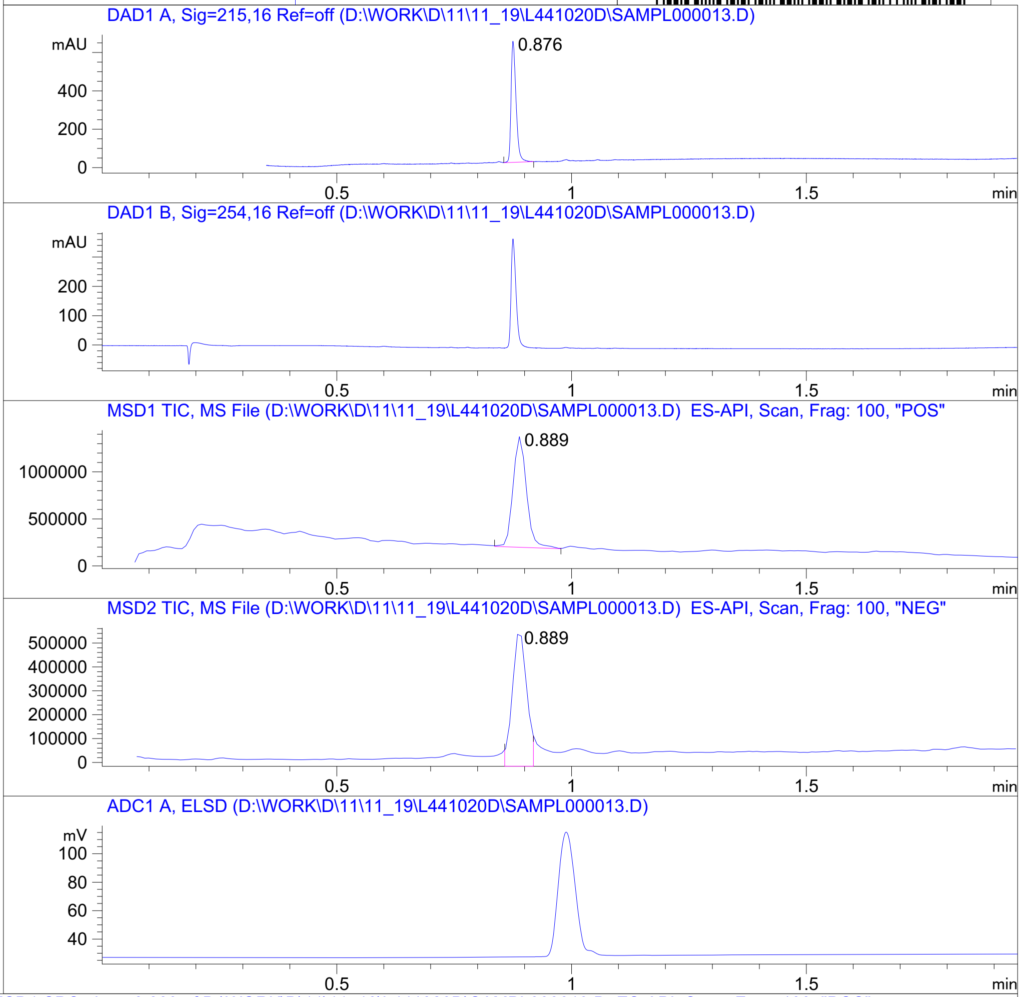


RT 0.889

RT 0.889

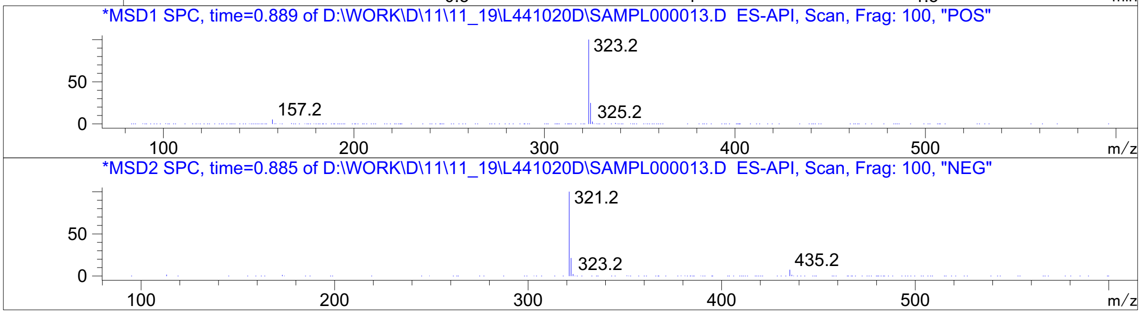


Inj.Date 11/19/2021 K P2-B-04 -5- Acq. Method C:\CHEM32\-> ->

Z4289708272_

MaxPeak: 98.55%
Ret_Time: 1.202 min

# W980729

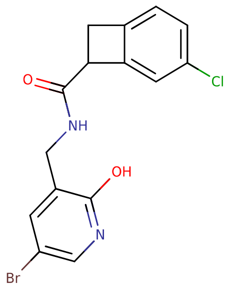


**Mol Wt** 367.62
**Exact Mass** 367.99

| # | Time | Area% |
|---|---|---|
| 1 | 1.080 | 1.45 |
| 2 | 1.202 | 98.55 |

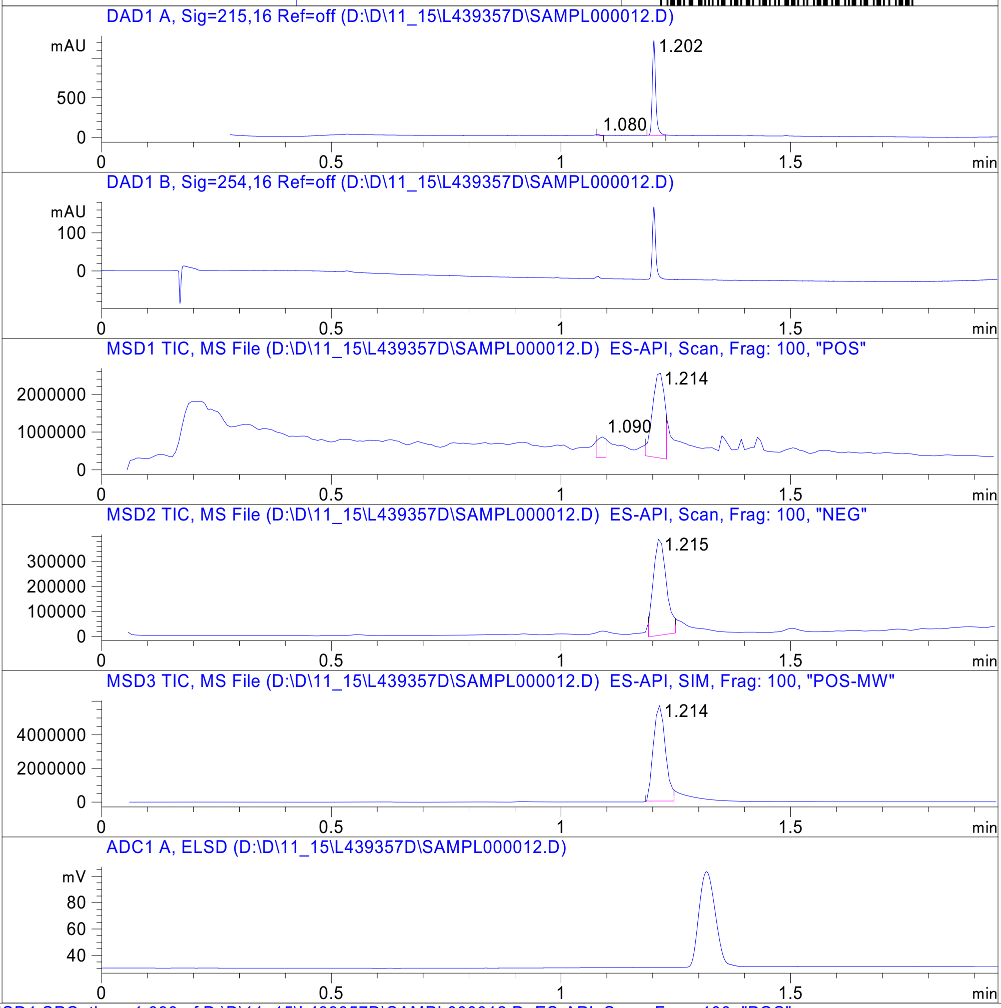


RT 1.090

RT 1.214

RT 1.215

RT 1.214

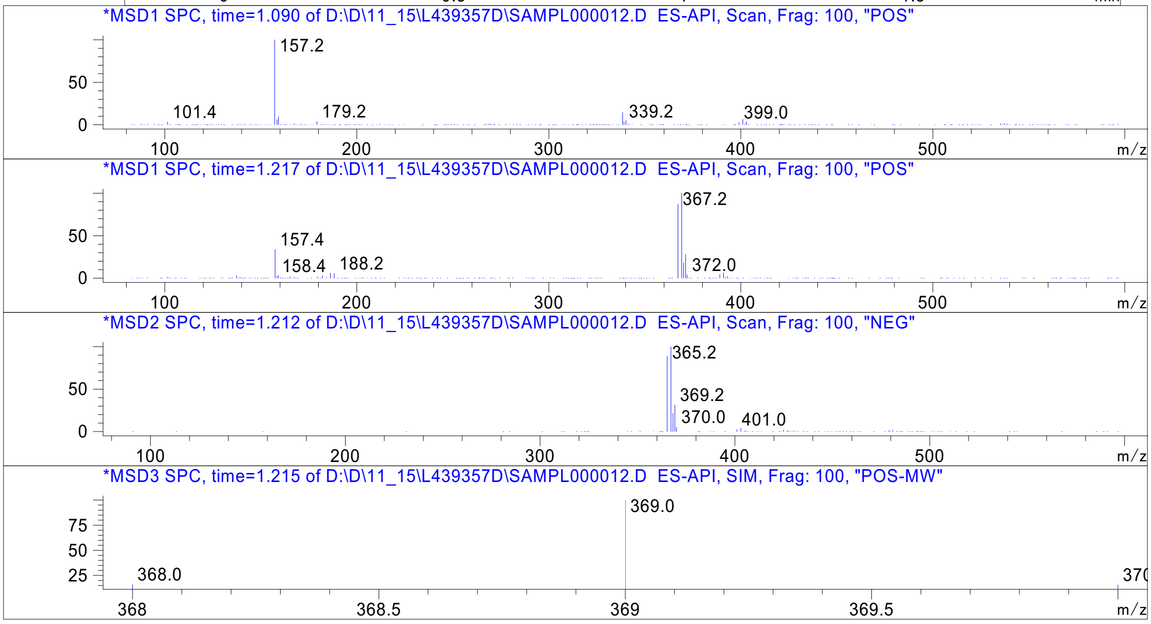




Z5420225639_

MaxPeak: 96.28%
Ret_Time: 1.319 min

X175457$4

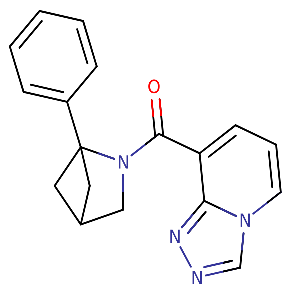


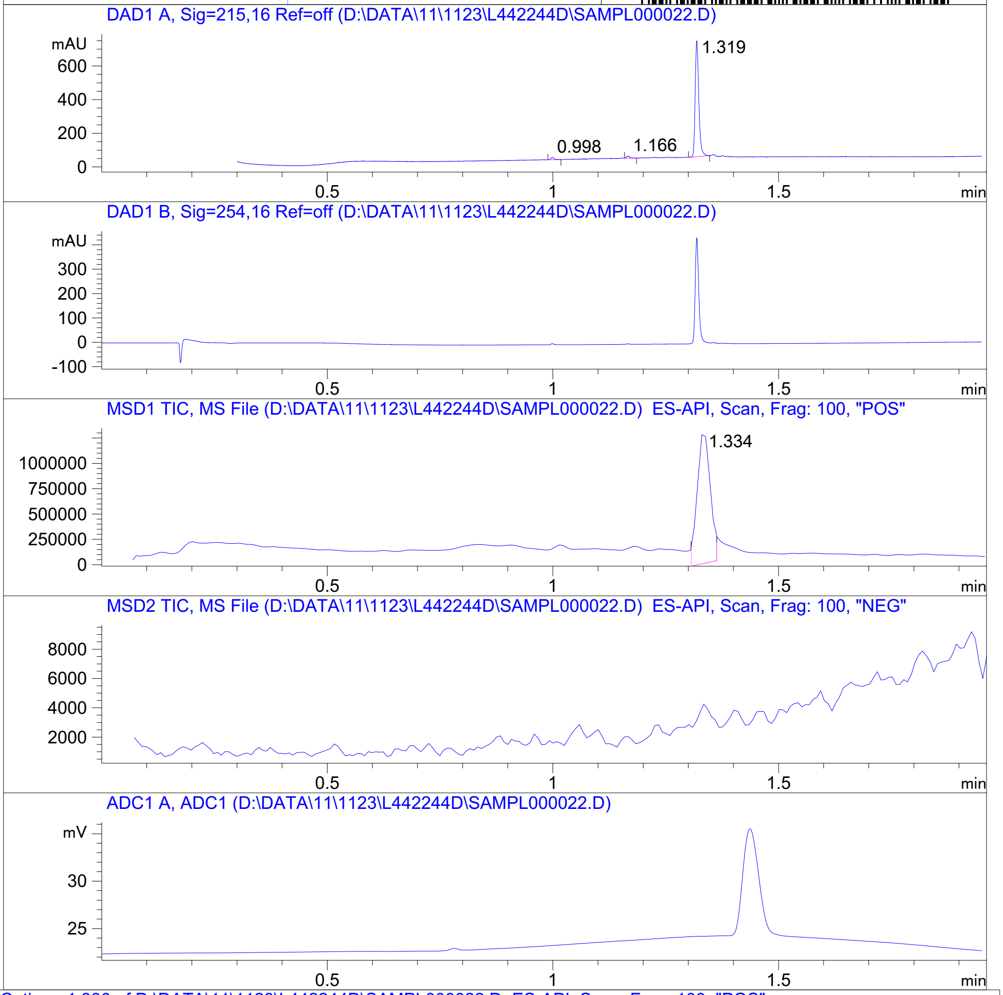


**Mol Wt** 304.35

**Exact Mass** 304.15

| # | Time | Area% |
|---|---|---|
| 1 | 0.998 | 1.94 |
| 2 | 1.166 | 1.79 |
| 3 | 1.319 | 96.28 |

RT 1.334

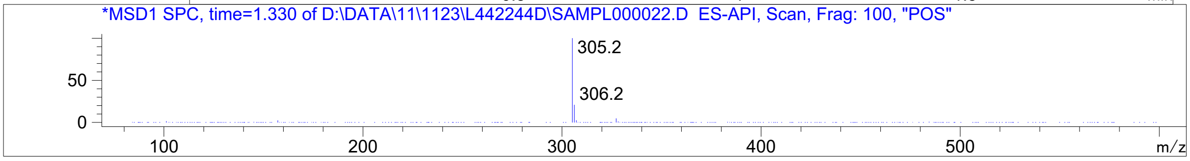


Inj.Date 11/22/2021 LT - 4 - Acq. Method C:\CHEM32\-> ->

Z3555684465_

MaxPeak: 100.00%
Ret_Time: 0.909 min

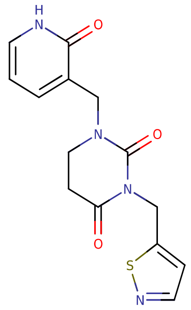


**Mol Wt 318.35**

**Exact Mass 318.08**

| # | Time | Area% |
|---|---|---|
| 1 | 0.909 | 100.00 |

W980660$2

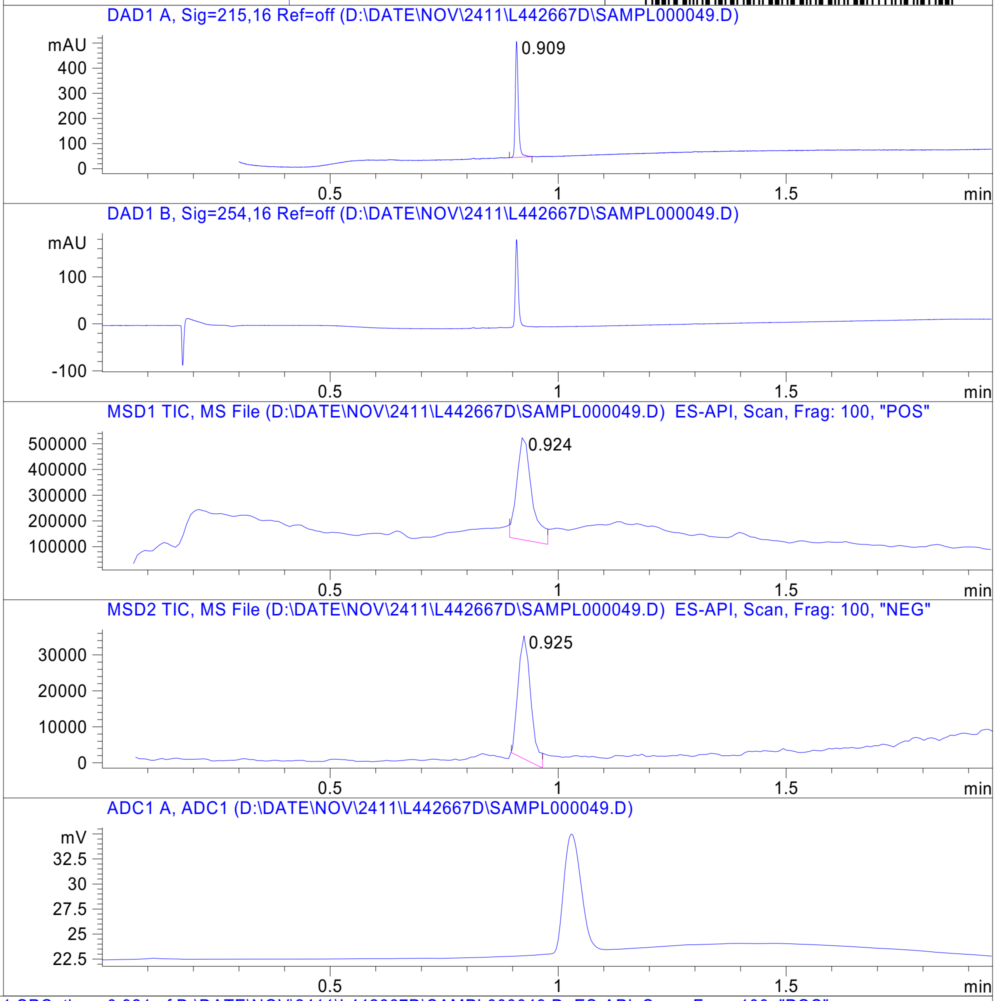


RT 0.924

RT 0.925

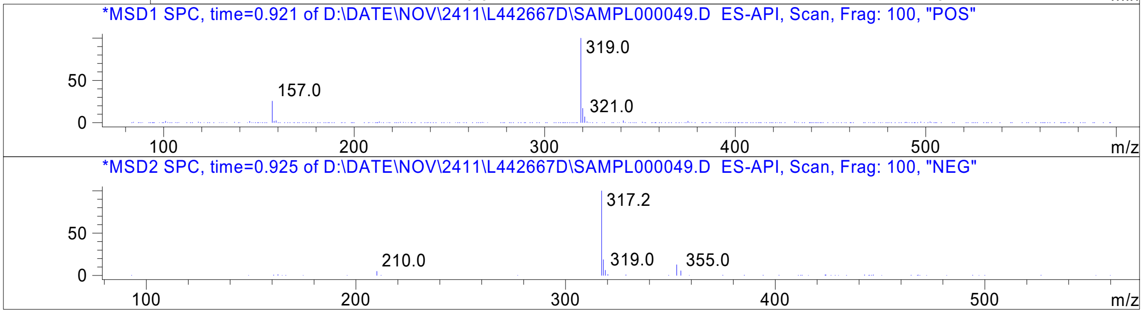


Inj.Date 11/23/2021 Y  Acq. Method C:\CHEM32\-> ->

Z3541227016_

MaxPeak: 98.00%
Ret_Time: 0.869 min

# X238577$2

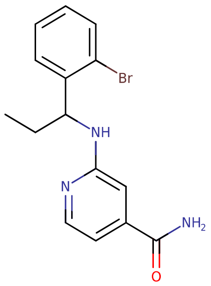

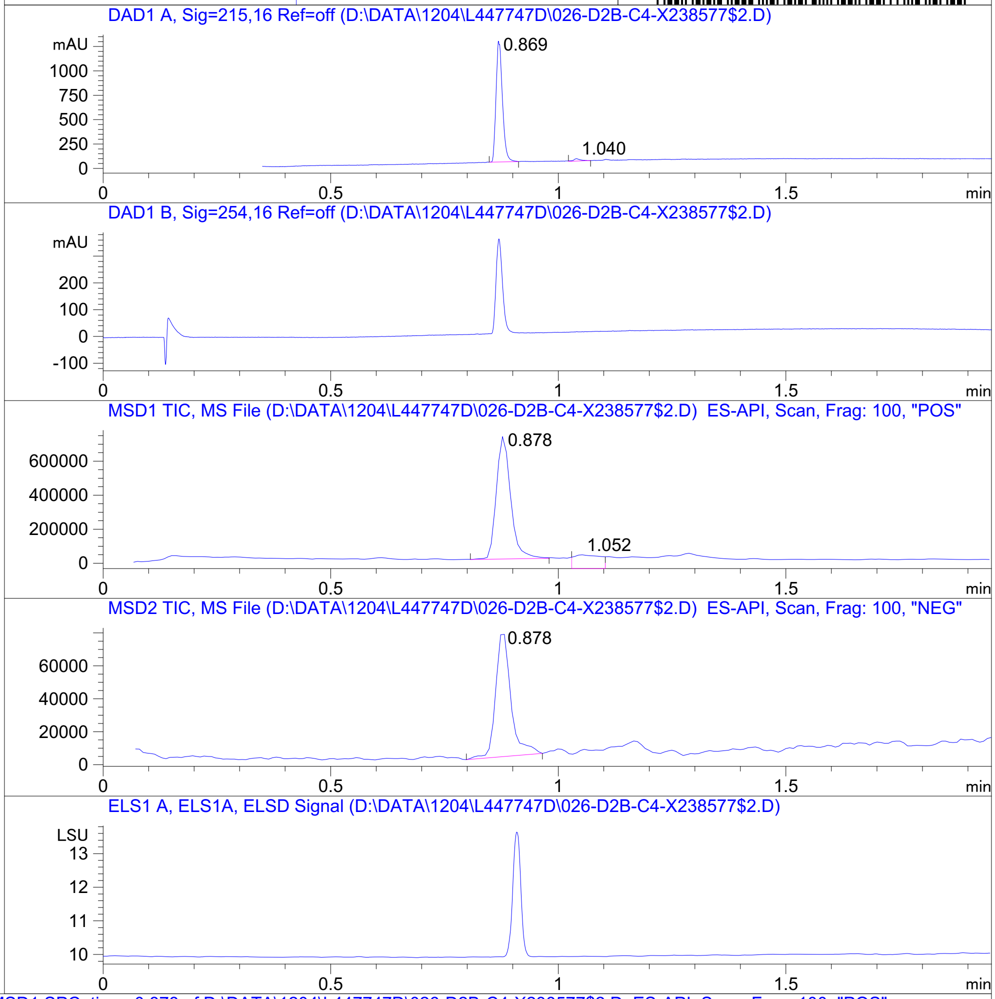


**Mol Wt** 334.21
**Exact Mass** 333.07

| # | Time | Area% |
|---|---|---|
| 1 | 0.869 | 98.00 |
| 2 | 1.040 | 2.00 |

RT 0.878

RT 1.052

RT 0.878

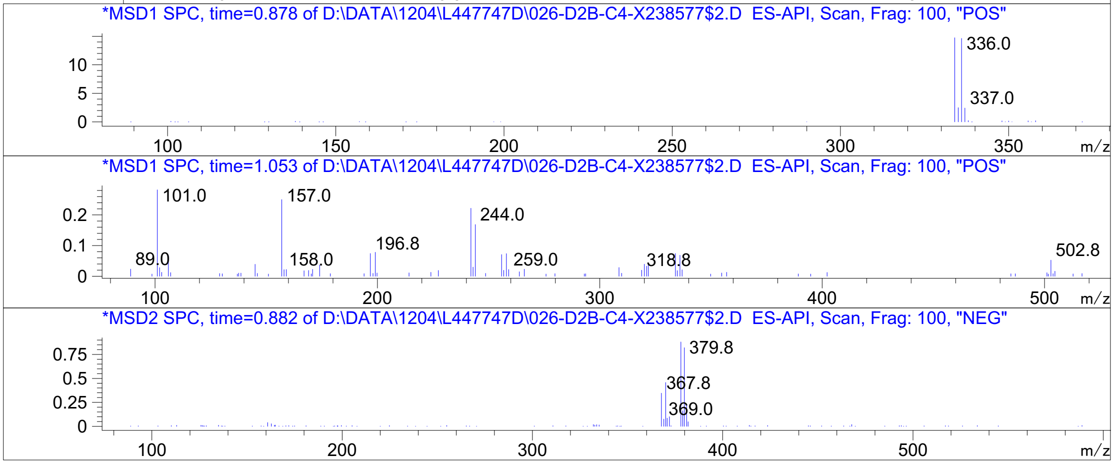




Z1425997900_

MaxPeak: 100.00%
Ret_Time: 0.589 min

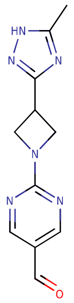

**Mol Wt** 244.25
**Exact Mass** 244.11

| # | Time | Area% |
|---|---|---|
| 1 | 0.589 | 100.00 |

S993158$1

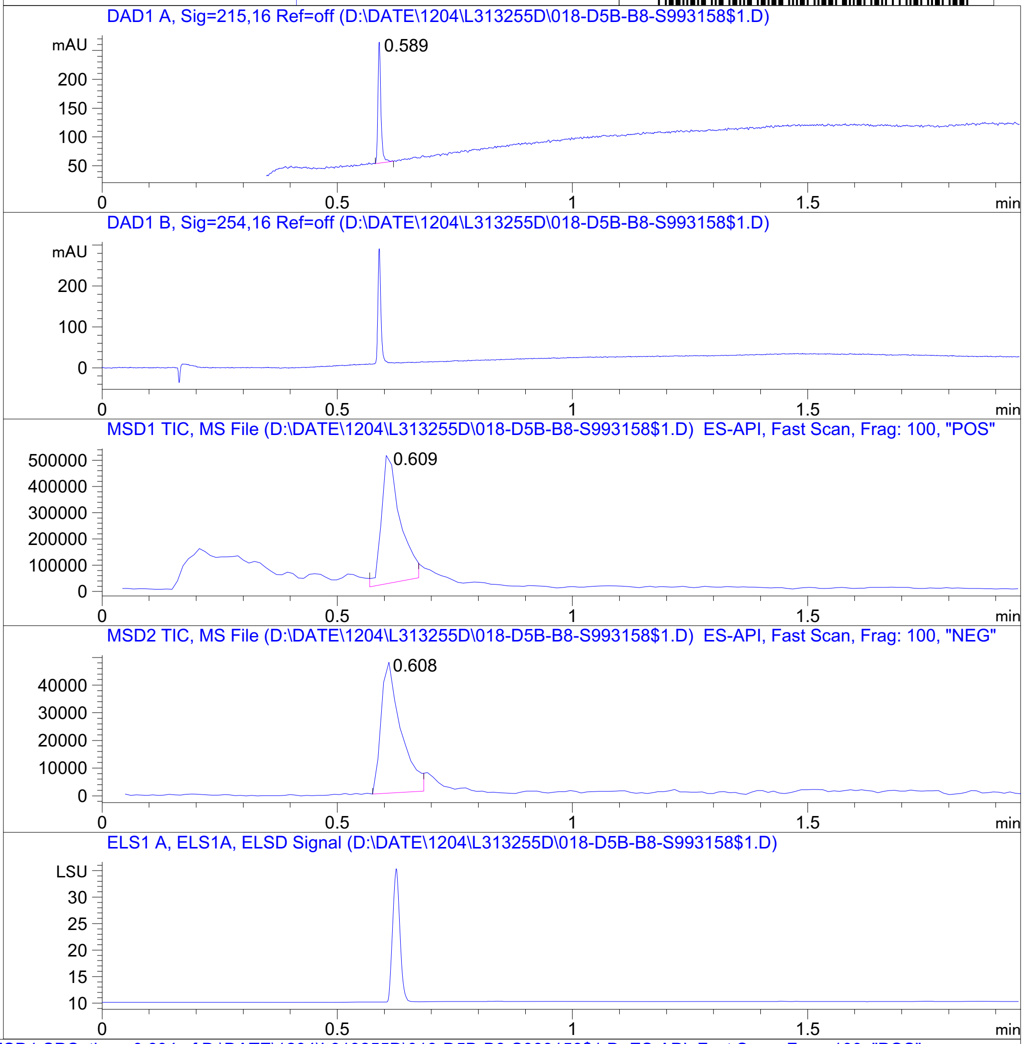


RT 0.609

RT 0.608

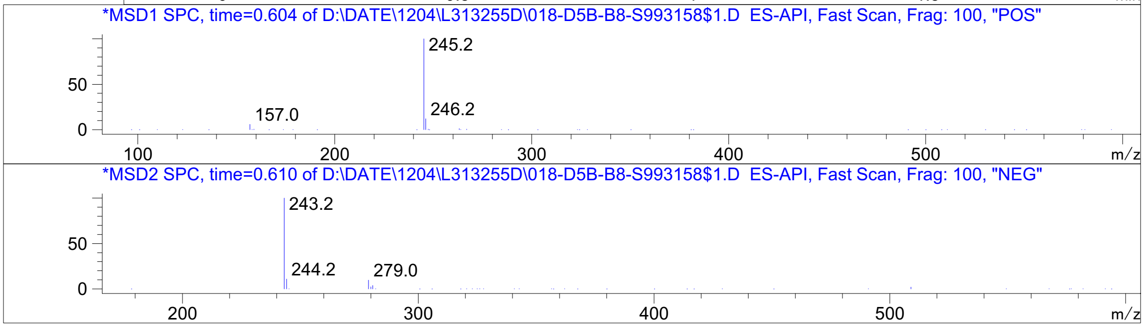


Inj.Date 12/3/2020 O Acq. Method C:\Users\ -> ->

Z2974471486_

MaxPeak: 100.00%
Ret_Time: 0.781 min

U211096$2

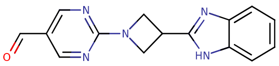


**Mol Wt** 279.3

**Exact Mass** 279.12

| # | Time | Area% |
|---|---|---|
| 1 | 0.781 | 100.00 |

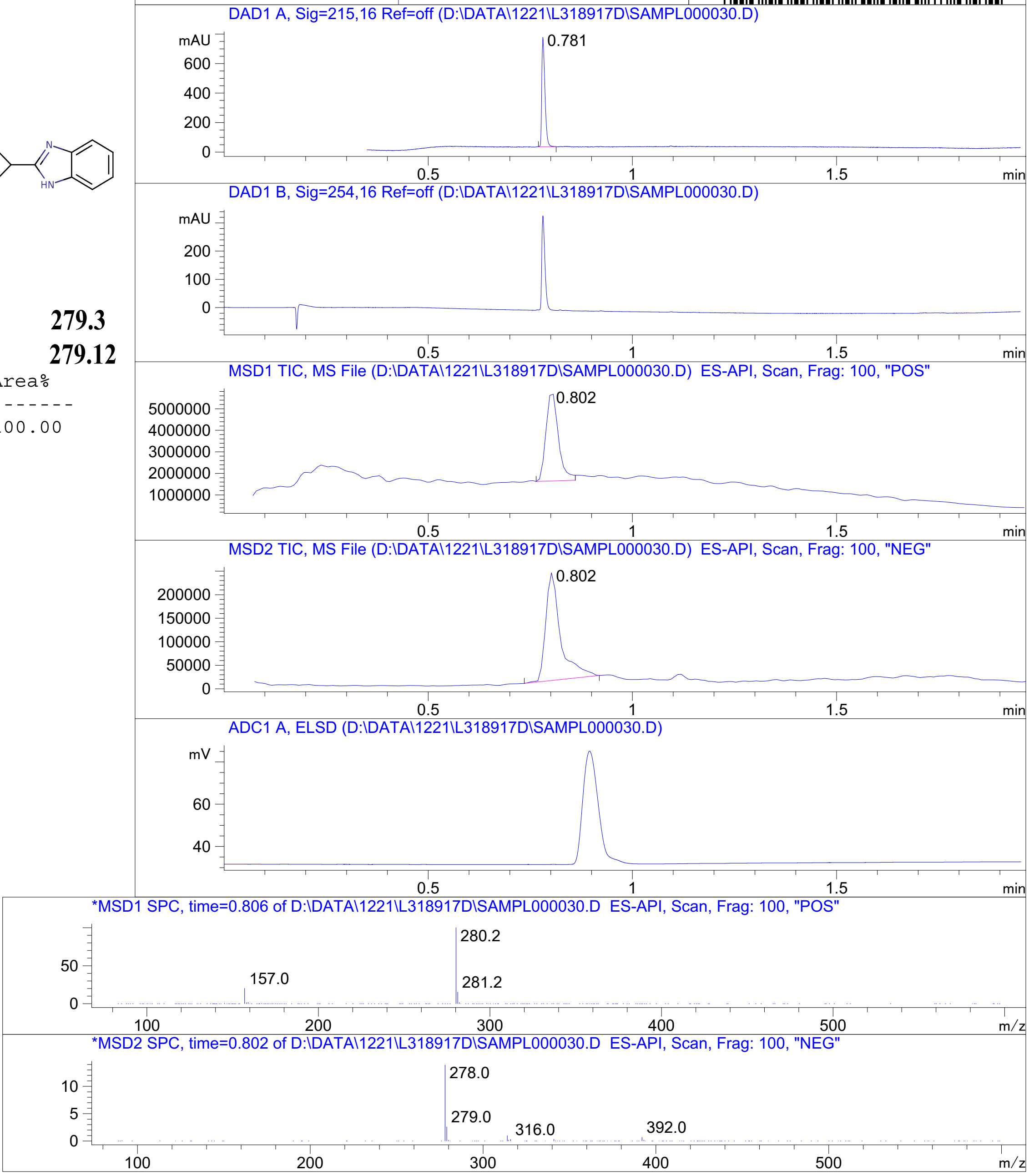




Z4497433835_

MaxPeak: 92.45%
Ret_Time: 1.228 min

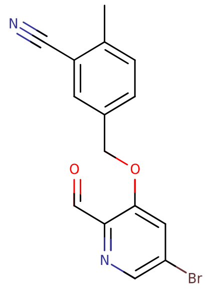

**Mol Wt** 331.16
**Exact Mass** 330.01

| # | Time | Area% |
|---|---|---|
| 1 | 1.228 | 92.45 |
| 2 | 1.378 | 5.40 |
| 3 | 1.787 | 2.15 |

U235907$1

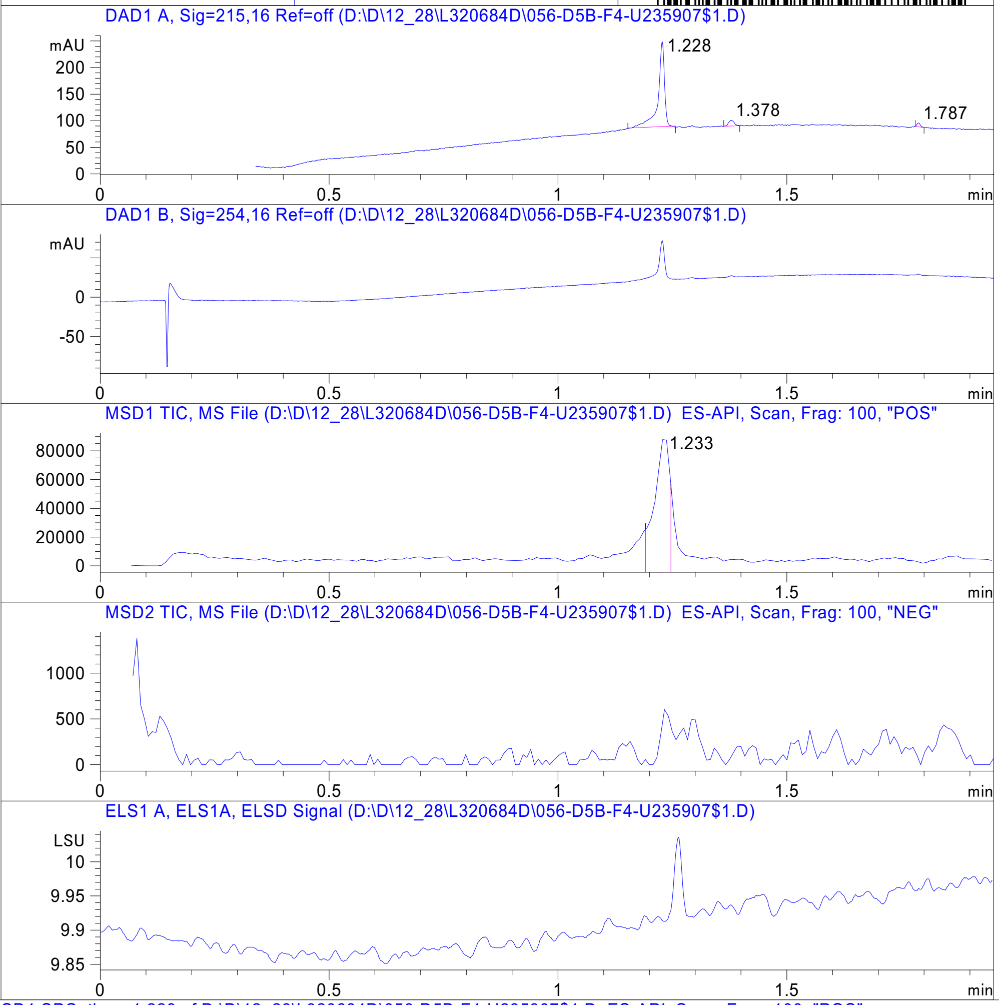


RT 1.233

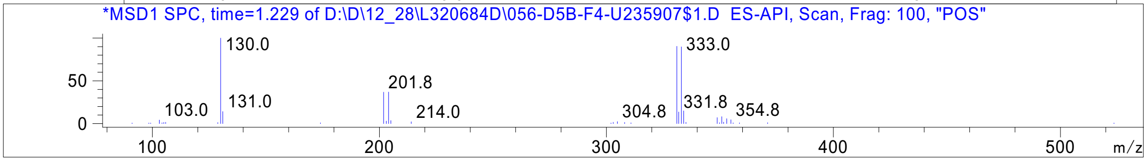




Z4767502624_

MaxPeak: 100.00%
Ret_Time: 1.316 min

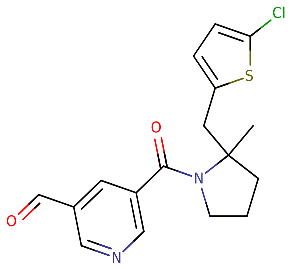


**Mol Wt** 348.85

**Exact Mass** 348.09

| # | Time | Area% |
|---|---|---|
| 1 | 1.316 | 100.00 |

U257602$1

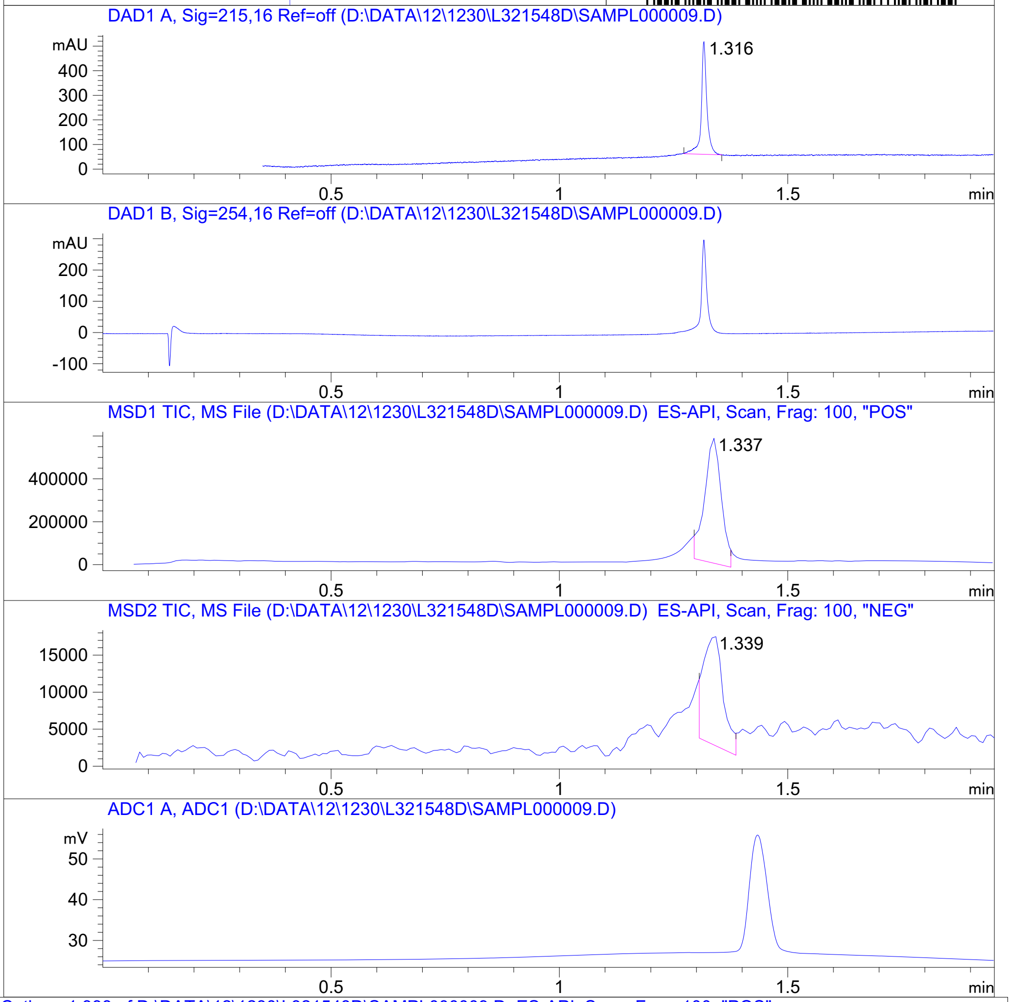


RT 1.337

RT 1.339

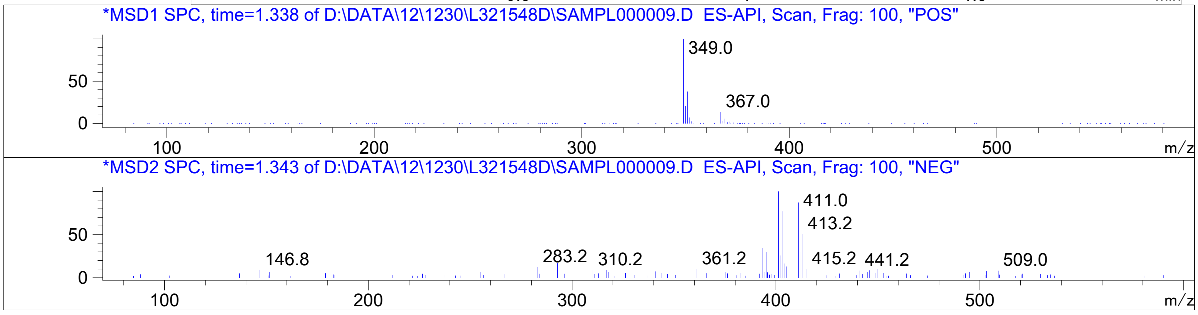




Z2962685976_

MaxPeak: 100.00%
Ret_Time: 1.340 min

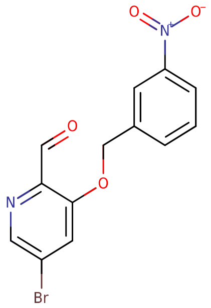


**Mol Wt** 337.13
**Exact Mass** 335.97

| # | Time | Area% |
|---|---|---|
| 1 | 1.340 | 100.00 |

U920097$5

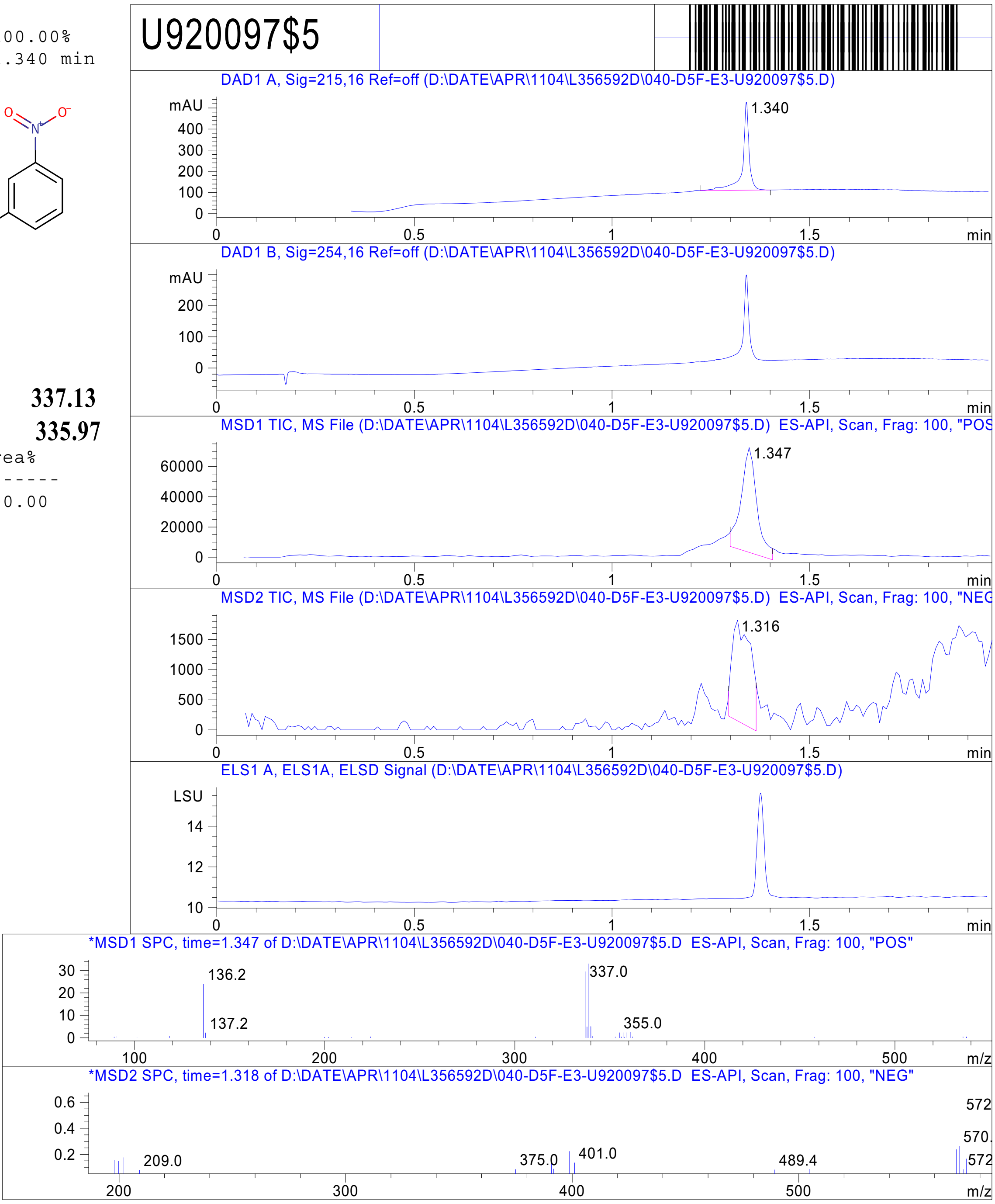


RT 1.347

RT 1.316

Inj.Date 4/11/2021 Y -14- Acq. Method C:\Chem32\-> ->

Z4954569112_

MaxPeak:  100.00%
Ret_Time: 1.421 min

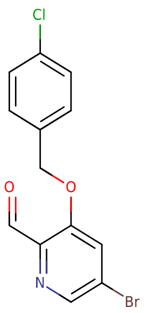


U920098$1

**Mol Wt** 326.57

**Exact Mass** 326.96

| # | Time | Area% |
|---|---|---|
| 1 | 1.421 | 100.00 |

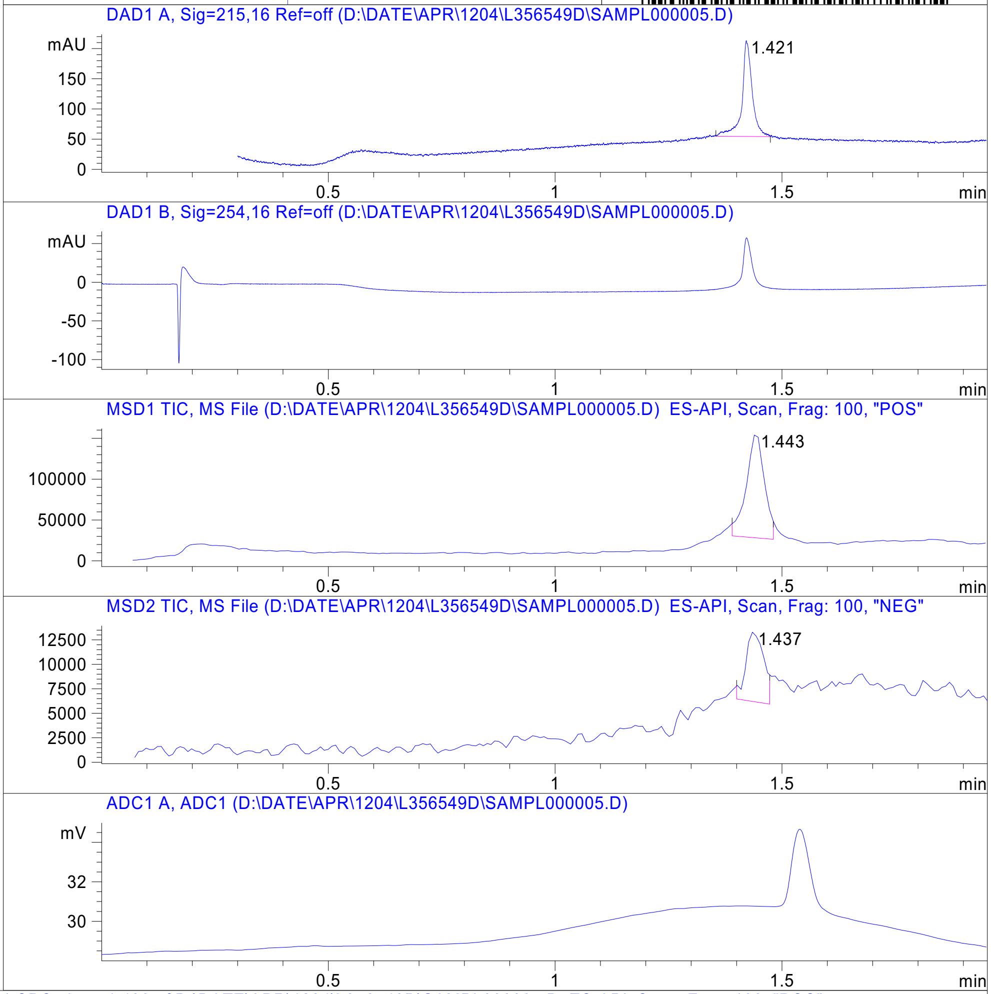


RT 1.443

RT 1.437

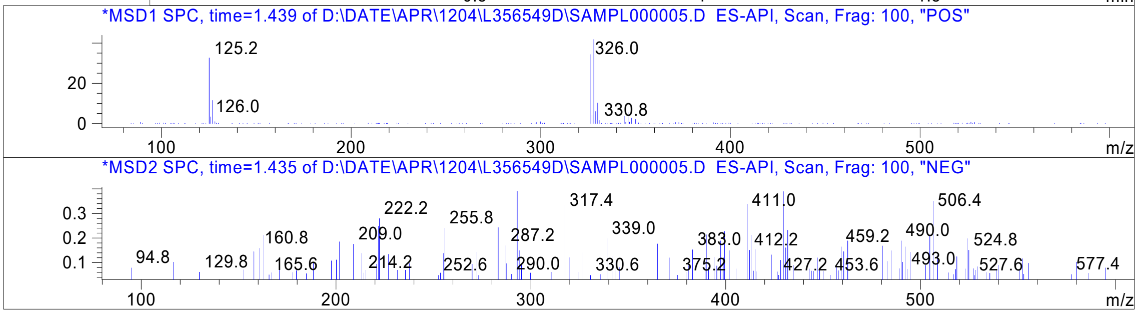




Z4954569113_

MaxPeak: 100.00%
Ret_Time: 1.402 min

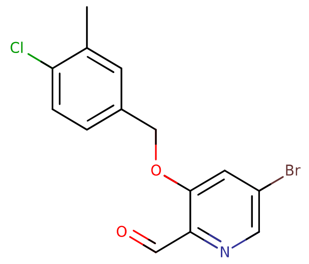


**Mol Wt** 340.6

**Exact Mass** 340.98

| # | Time | Area% |
|---|---|---|
| 1 | 1.402 | 100.00 |

# U920099$4

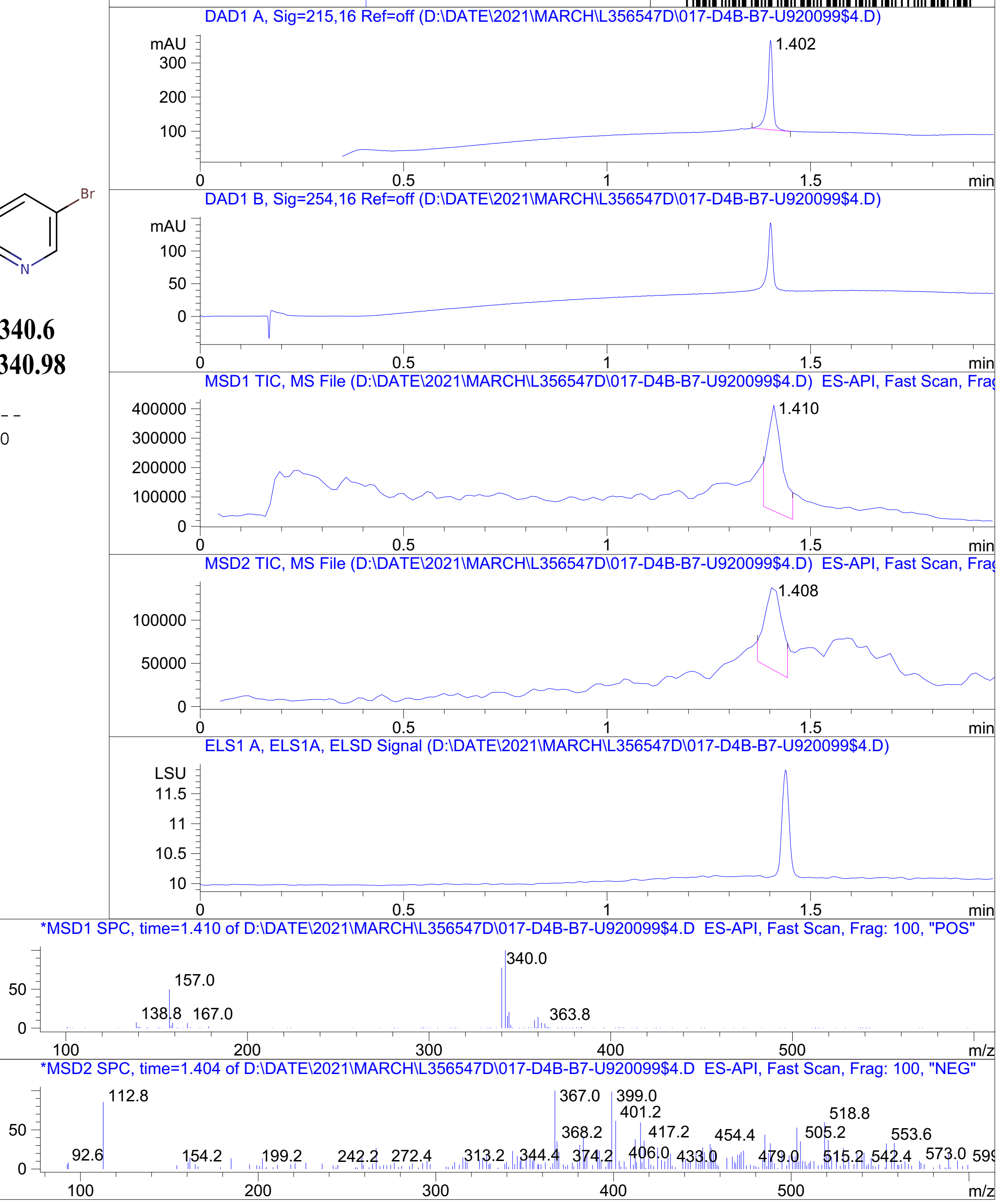


Inj.Date 4/11/2021 T <invalid> Acq. Method C:\Users\ -> ->

Z4954569120_

MaxPeak: 100.00%
Ret_Time: 1.446 min

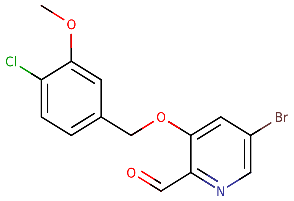


**Mol Wt** 356.6
**Exact Mass** 356.97

| # | Time | Area% |
|---|---|---|
| 1 | 1.446 | 100.00 |

U920102$3

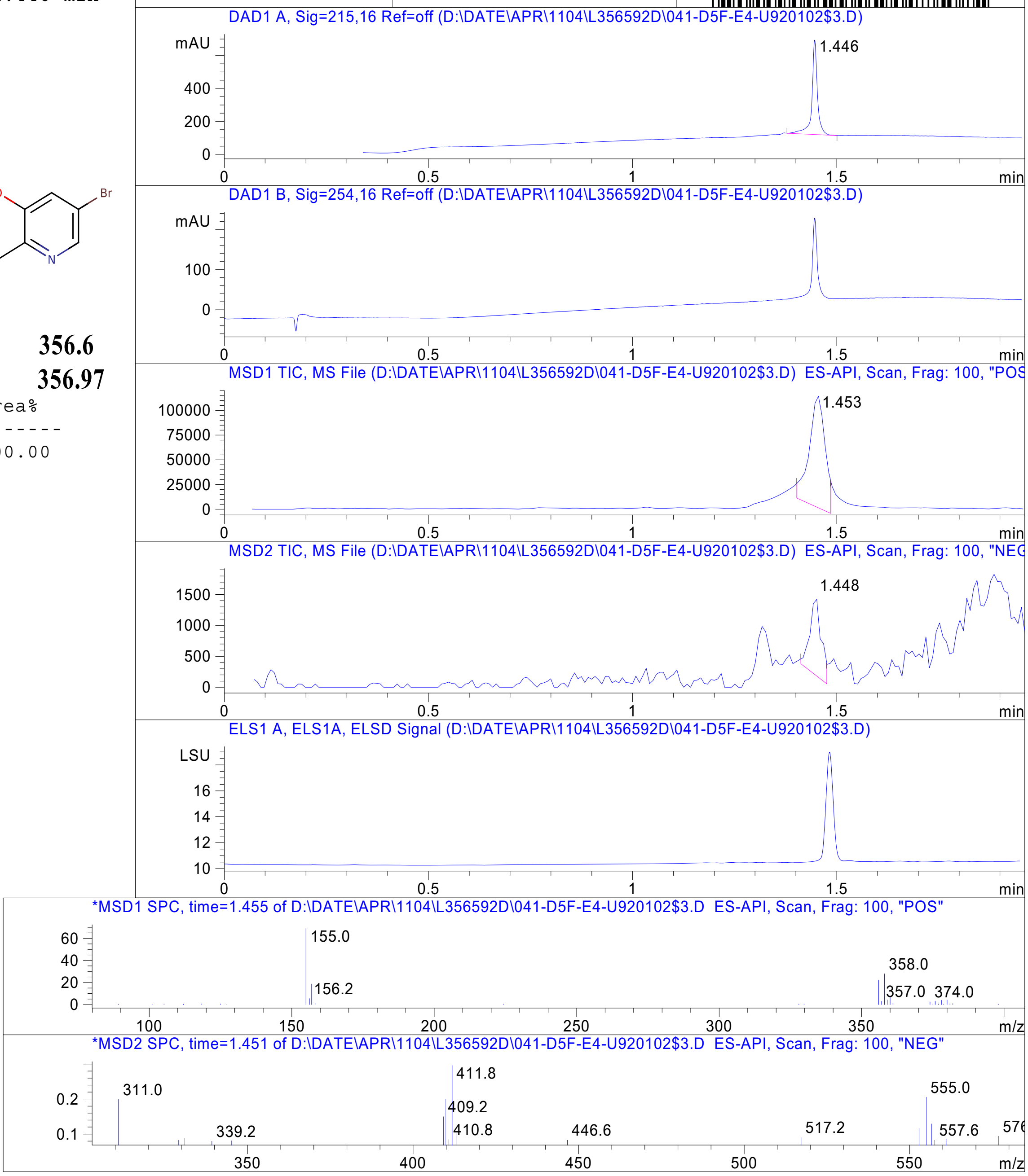


RT 1.453

RT 1.448

Inj.Date 4/11/2021 Y -14- Acq. Method C:\Chem32\-> ->

Z4954569125_

MaxPeak: 100.00%
Ret_Time: 1.336 min

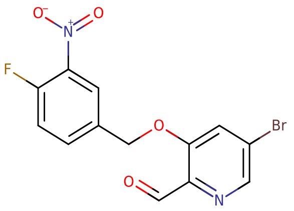


**Mol Wt** 355.12
**Exact Mass** 353.96

| # | Time | Area% |
|---|---|---|
| 1 | 1.336 | 100.00 |

U920111$1

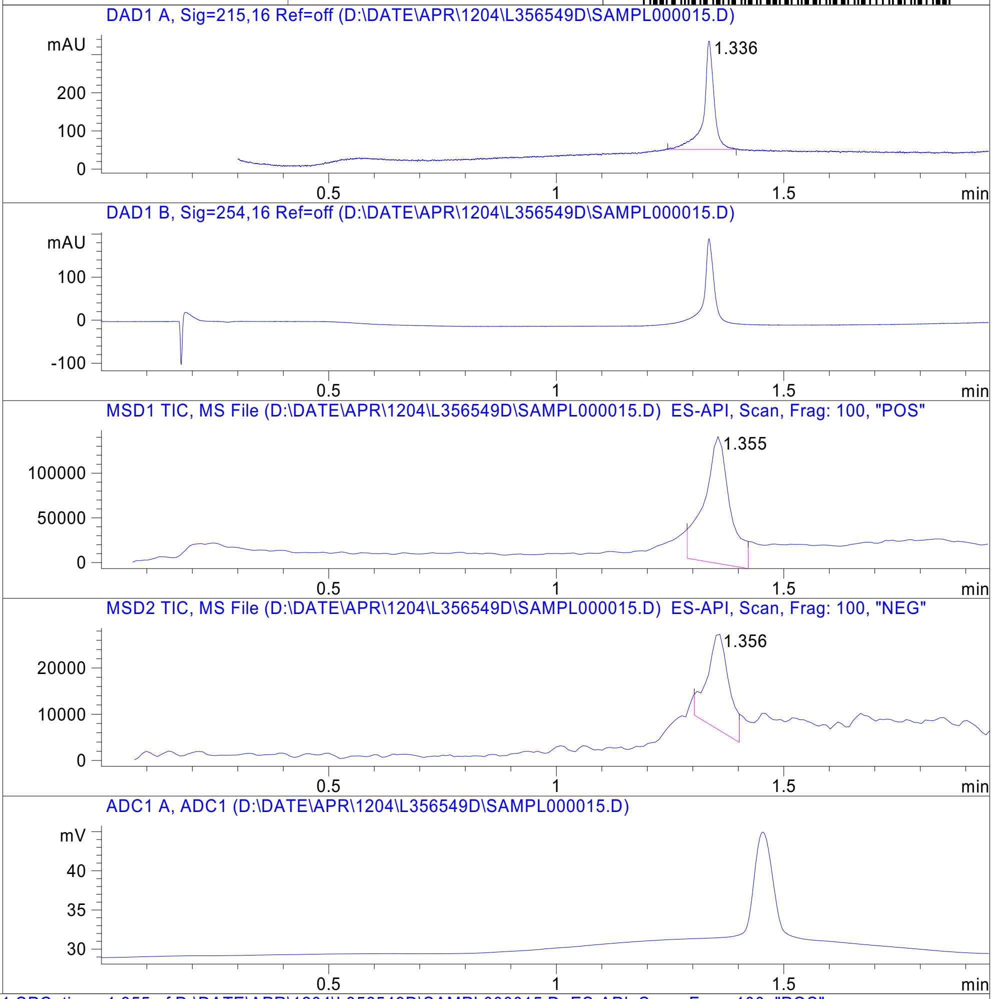


RT 1.355

RT 1.356

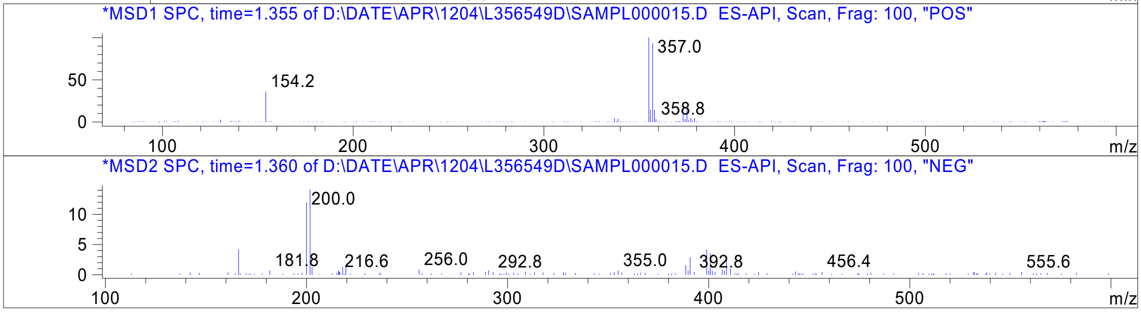




Z4954569219_

MaxPeak: 100.00%
Ret_Time: 1.528 min

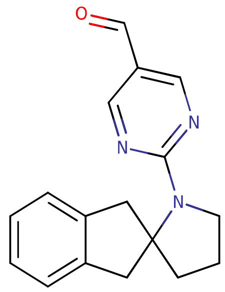


**Mol Wt** 279.34

**Exact Mass** 279.16

| # | Time | Area% |
|---|---|---|
| 1 | 1.528 | 100.00 |

# S993138$1

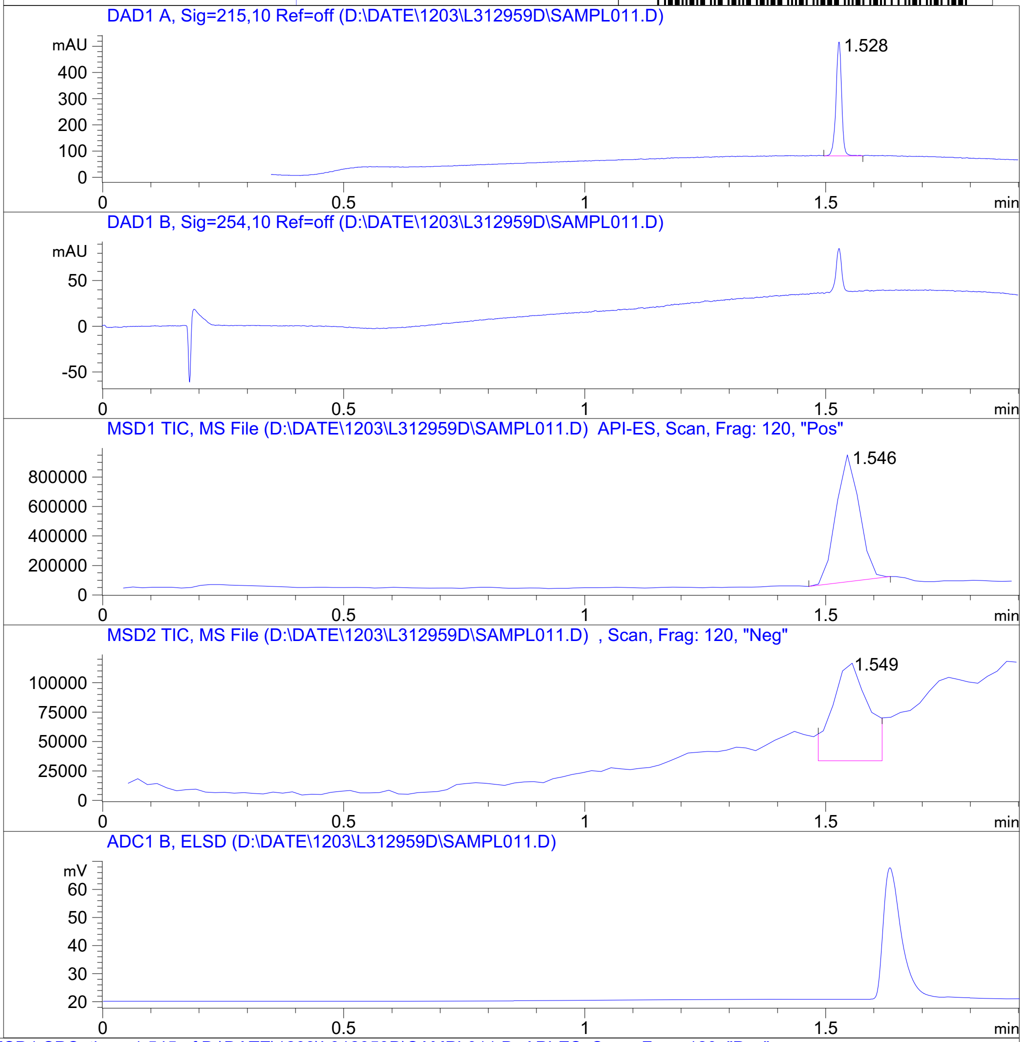


RT 1.546

RT 1.549

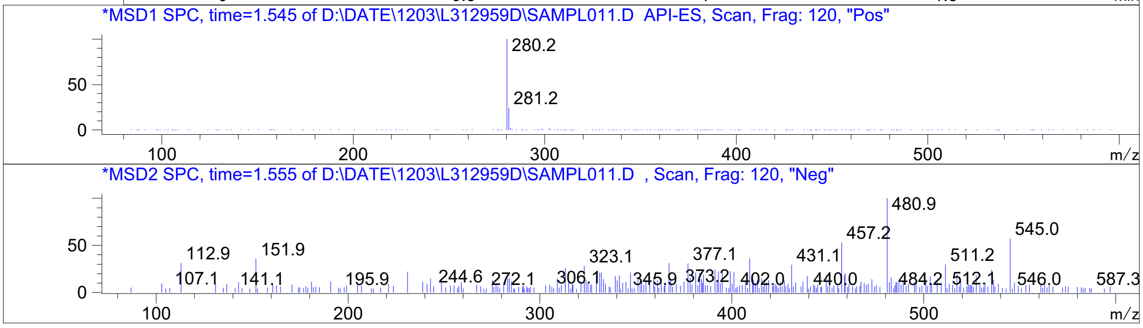


Inj.Date 12/3/2020 O -SL- Acq. Method C:\HPCHEM\-> ->

Z2851996828_

MaxPeak:  100.00%
Ret_Time: 1.648 min

# U339489$1

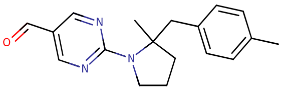

**Mol Wt** 295.38
**Exact Mass** 295.2

| # | Time | Area% |
|---|---|---|
| 1 | 1.648 | 100.00 |

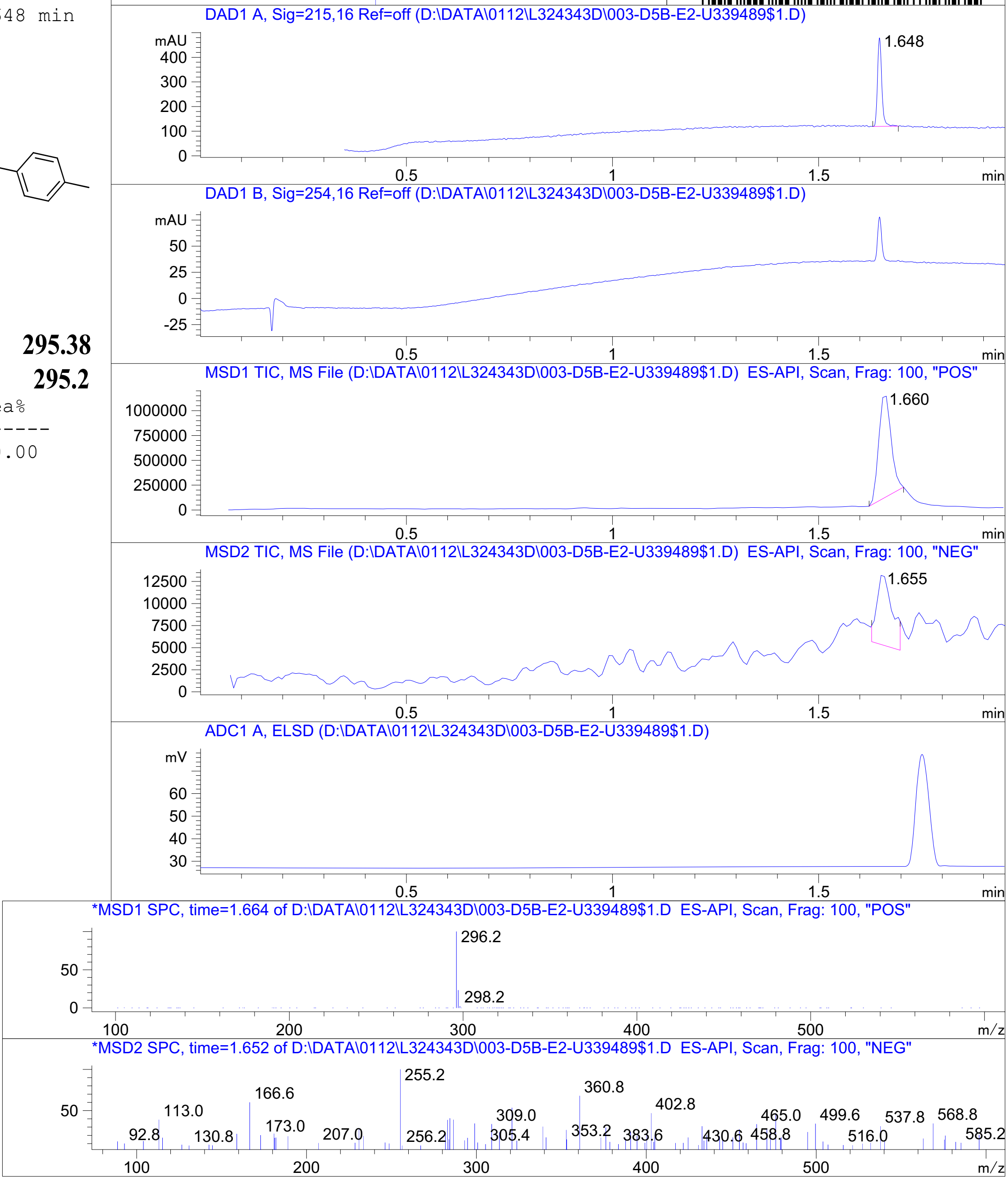


RT 1.660

RT 1.655

Inj.Date 1/11/2021 OA -12- Acq. Method C:\Chem32\-> ->

Z4658847374_

MaxPeak: 100.00%
Ret_Time: 1.508 min

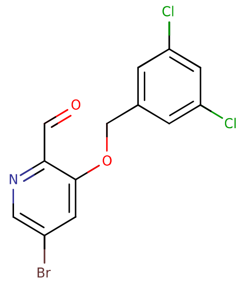


**Mol Wt** 361.02
**Exact Mass** 360.92

| # | Time | Area% |
|---|---|---|
| 1 | 1.508 | 100.00 |

W179754$4

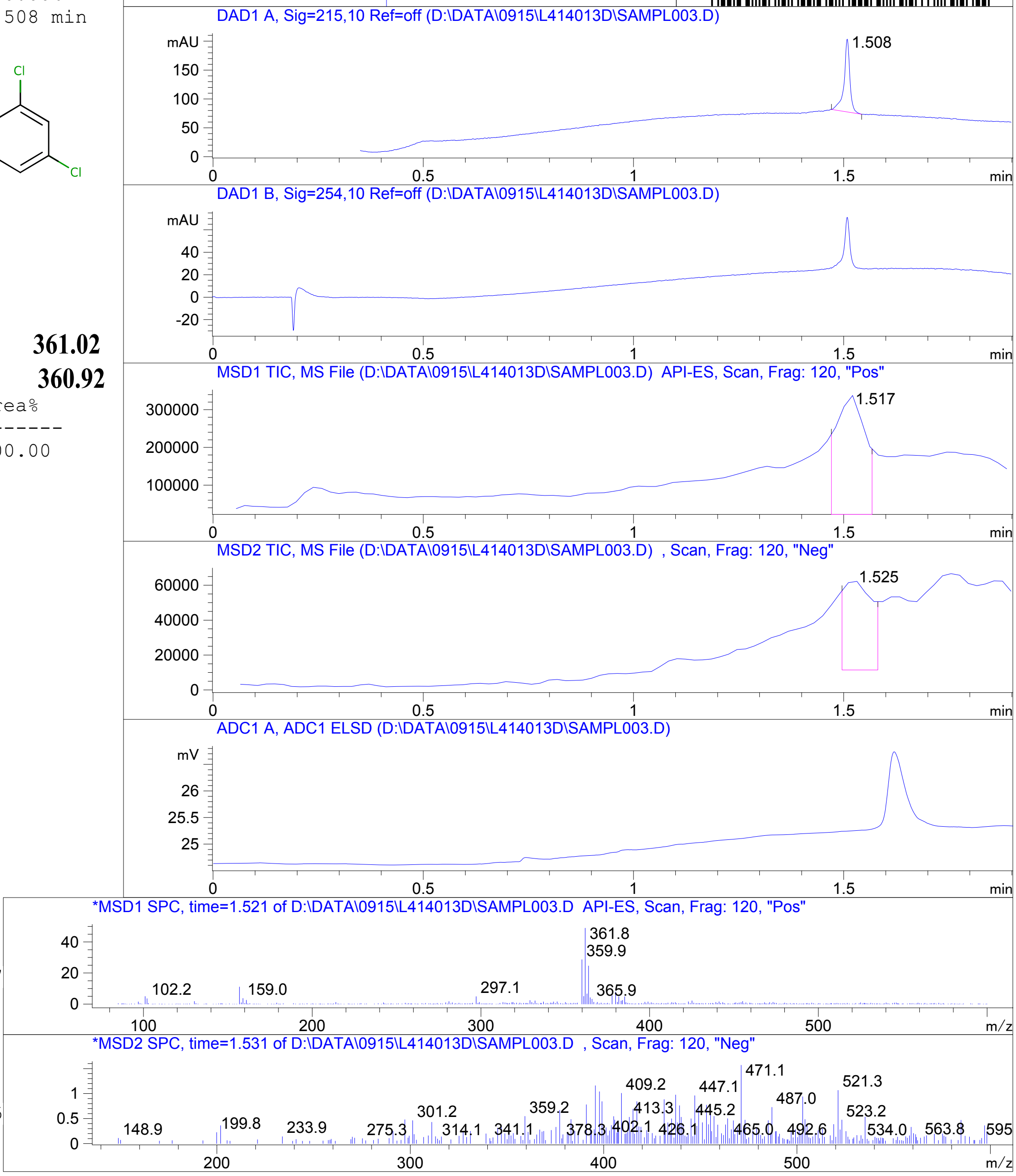


Inj.Date 9/16/2021 OA -VL- Acq. Method C:\HPCHEM\-> ->

Z5192779033_

MaxPeak: 100.00%
Ret_Time: 1.159 min

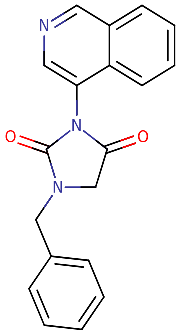

**Mol Wt 317.34**
**Exact Mass 317.13**

| # | Time | Area% |
|---|---|---|
| 1 | 1.159 | 100.00 |

# U887777

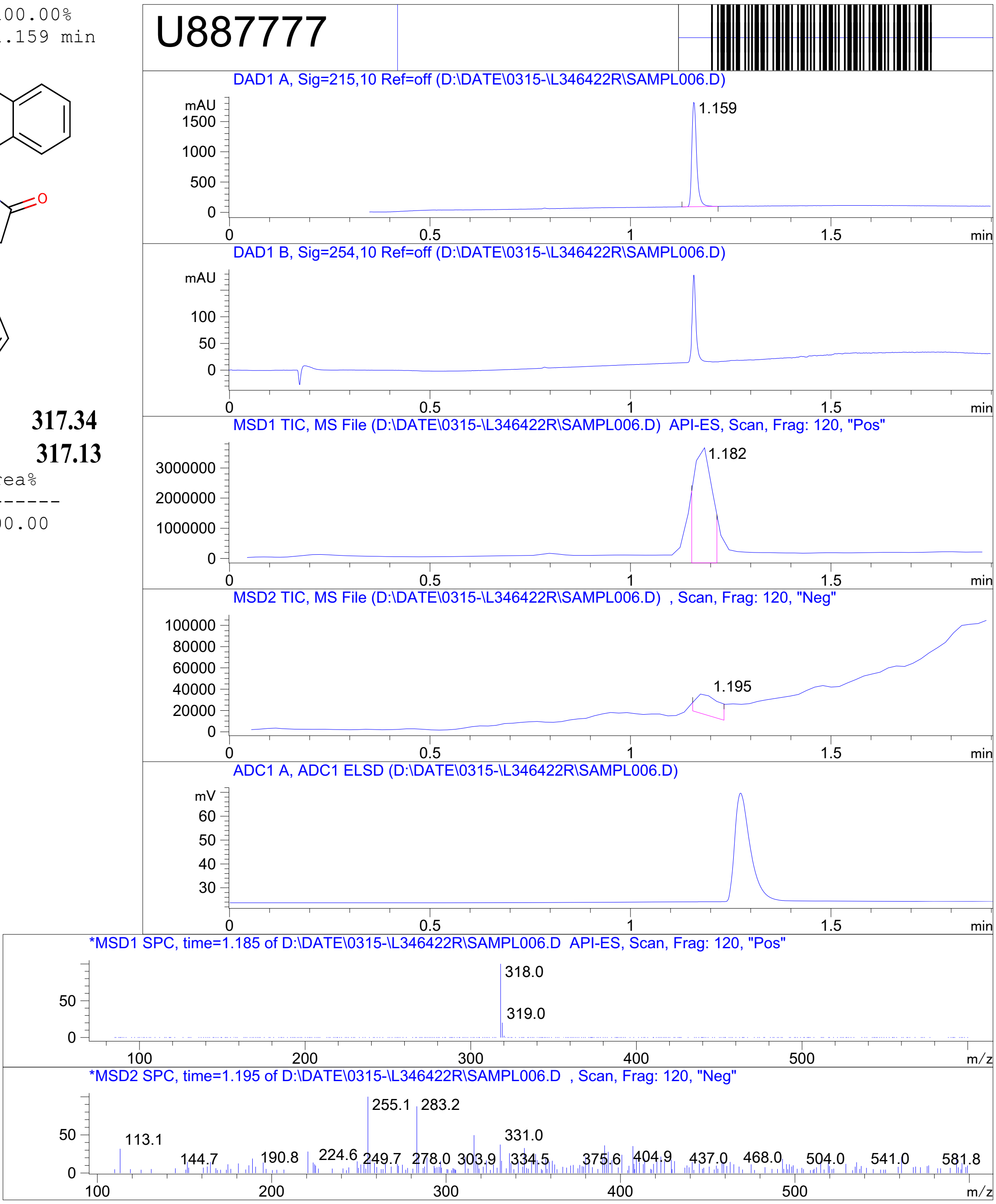


RT 1.182

RT 1.195

Inj.Date 3/16/2021 E -VL- Acq. Method C:\HPCHEM\-> ->

Z4929615903_

MaxPeak: 96.82%
Ret_Time: 1.258 min

# U887553

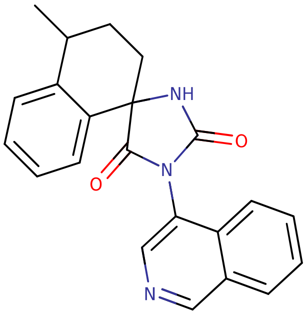

**Mol Wt** 357.4

**Exact Mass** 357.17

| # | Time | Area% |
|---|---|---|
| 1 | 0.555 | 0.19 |
| 2 | 0.688 | 1.05 |
| 3 | 0.851 | 1.12 |
| 4 | 0.998 | 0.81 |
| 5 | 1.258 | 96.82 |

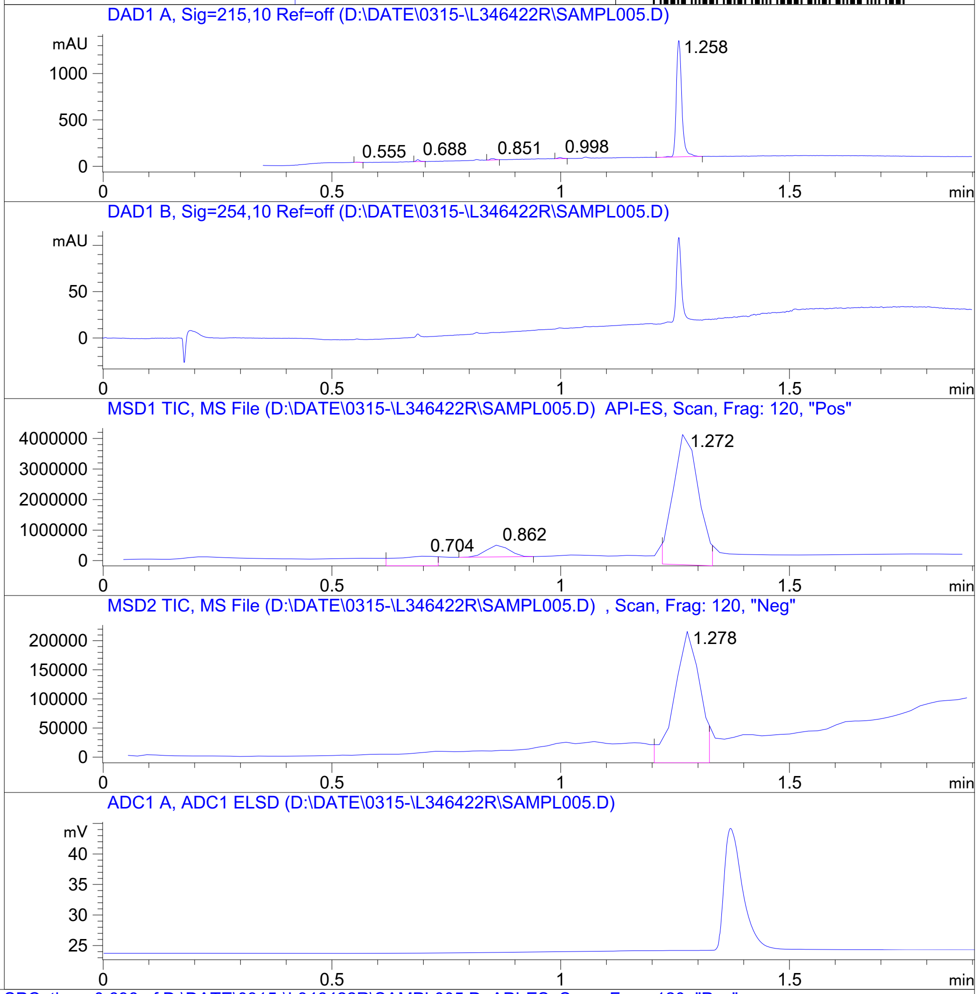


RT 0.704

RT 0.862

RT 1.272

RT 1.278

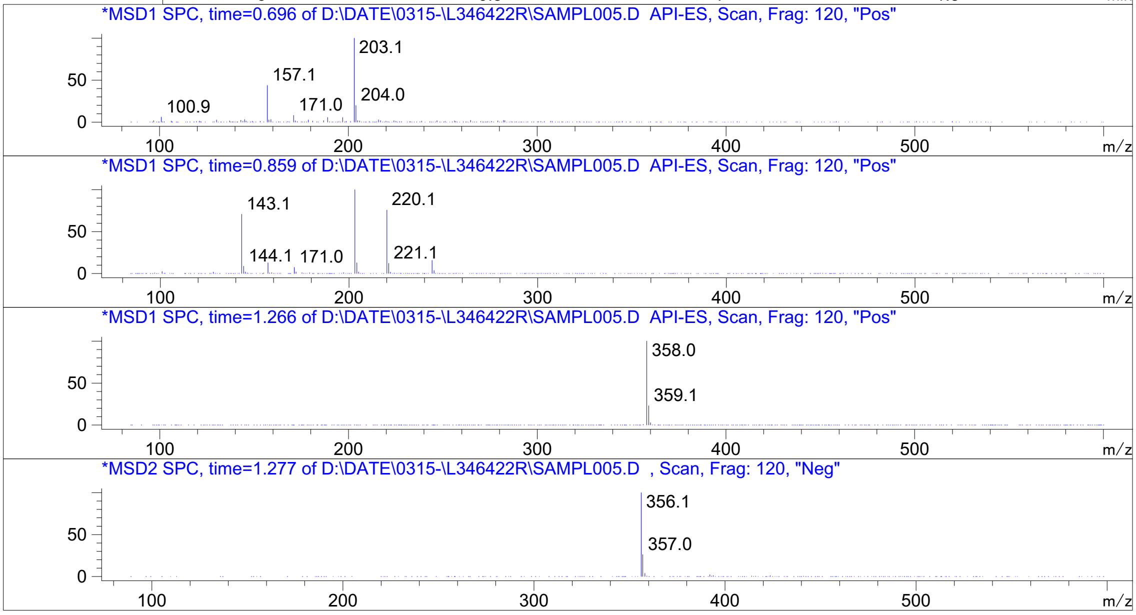


Inj.Date 3/16/2021 E -VL- Acq. Method C:\HPCHEM\-> ->

Z4929616137_

MaxPeak: 100.00%
Ret_Time: 1.029 min

# U887567

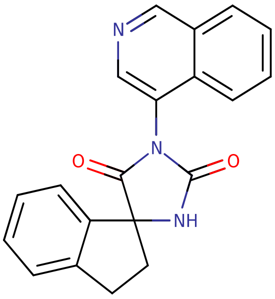


**Mol Wt** 329.35

**Exact Mass** 329.13

| # | Time | Area% |
|---|---|---|
| 1 | 1.029 | 100.00 |

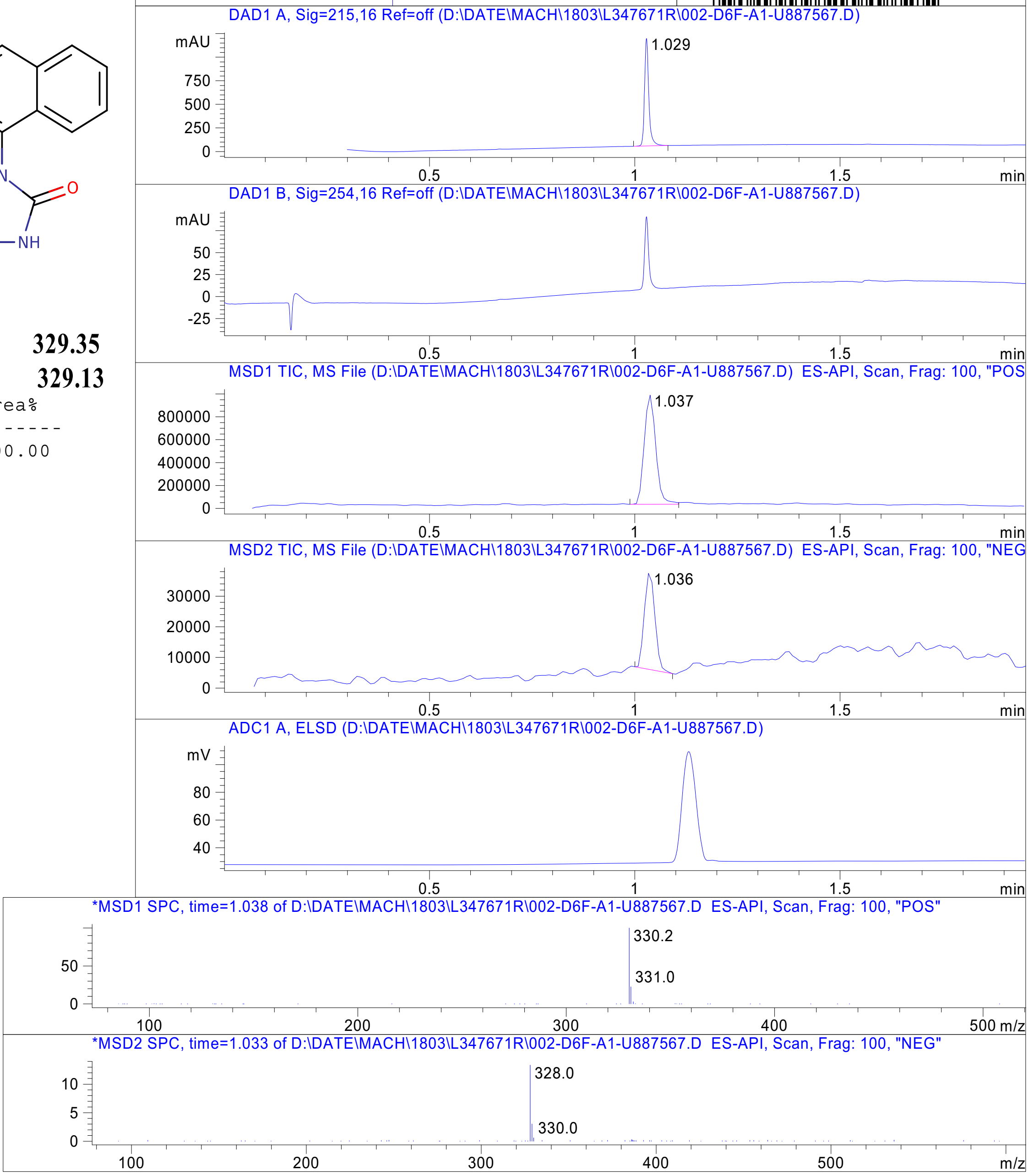


RT 1.037

RT 1.036

Inj.Date 3/17/2021 Y -8- Acq. Method C:\Chem32\-> ->

Z4929616111_

MaxPeak:  100.00%
Ret_Time: 0.940 min

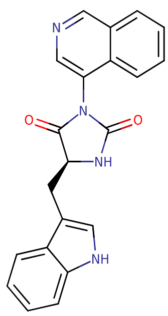


**Mol Wt 356.38**

**Exact Mass 356.14**

| # | Time | Area% |
|---|---|---|
| 1 | 0.940 | 100.00 |

U887560

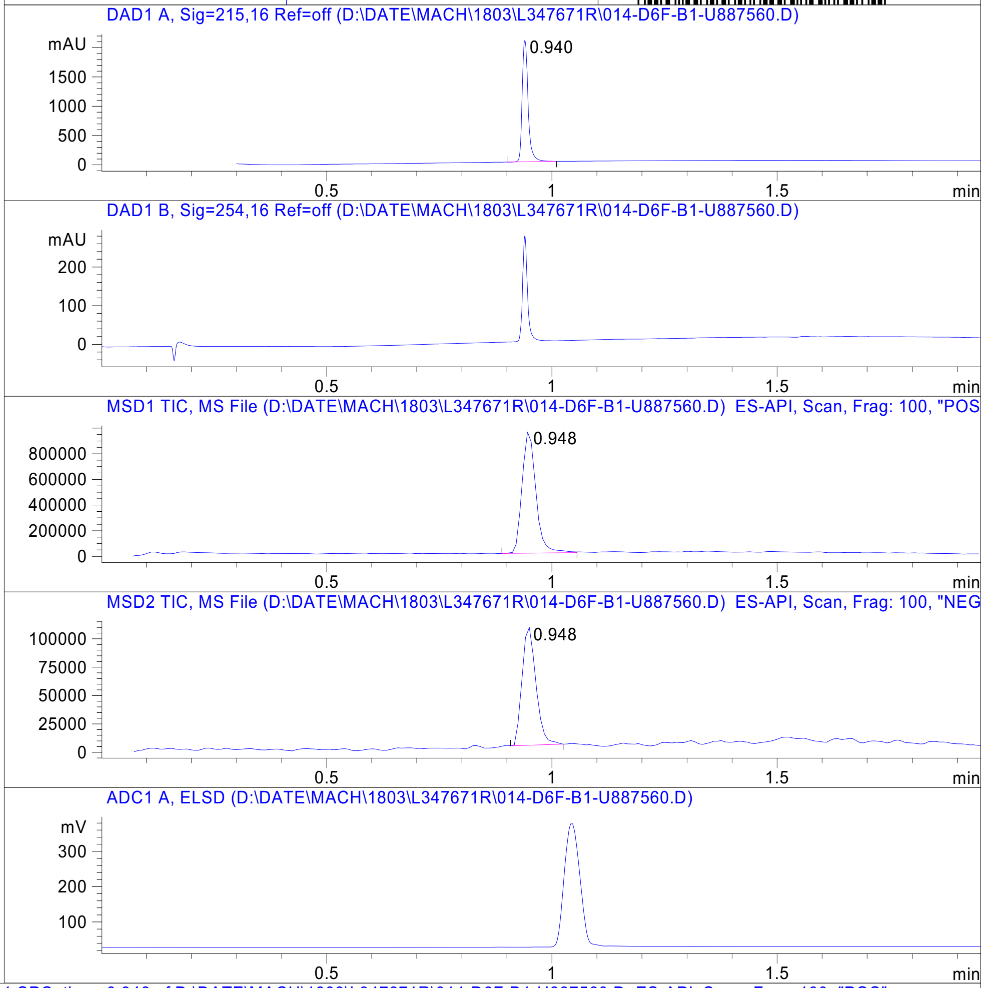


RT 0.948

RT 0.948

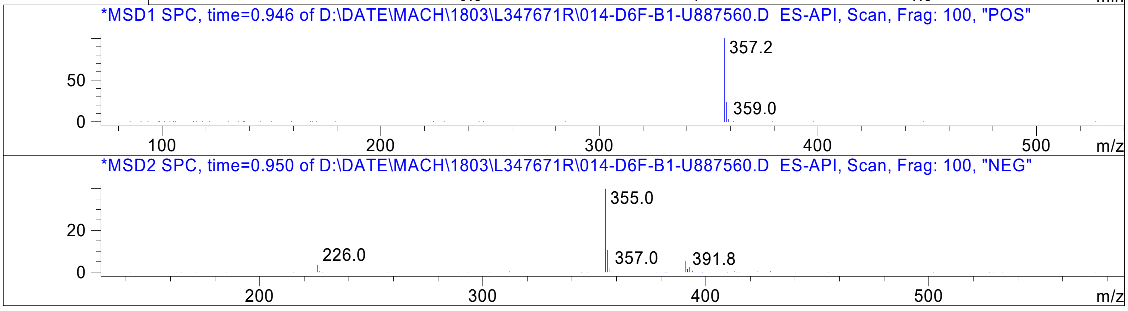




Z4929616023_

MaxPeak: 100.00%
Ret_Time: 1.050 min

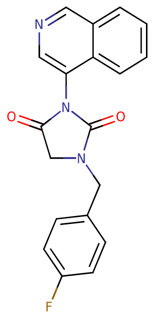

**Mol Wt** 335.33
**Exact Mass** 335.12

| # | Time | Area% |
|---|---|---|
| 1 | 1.050 | 100.00 |

U888029

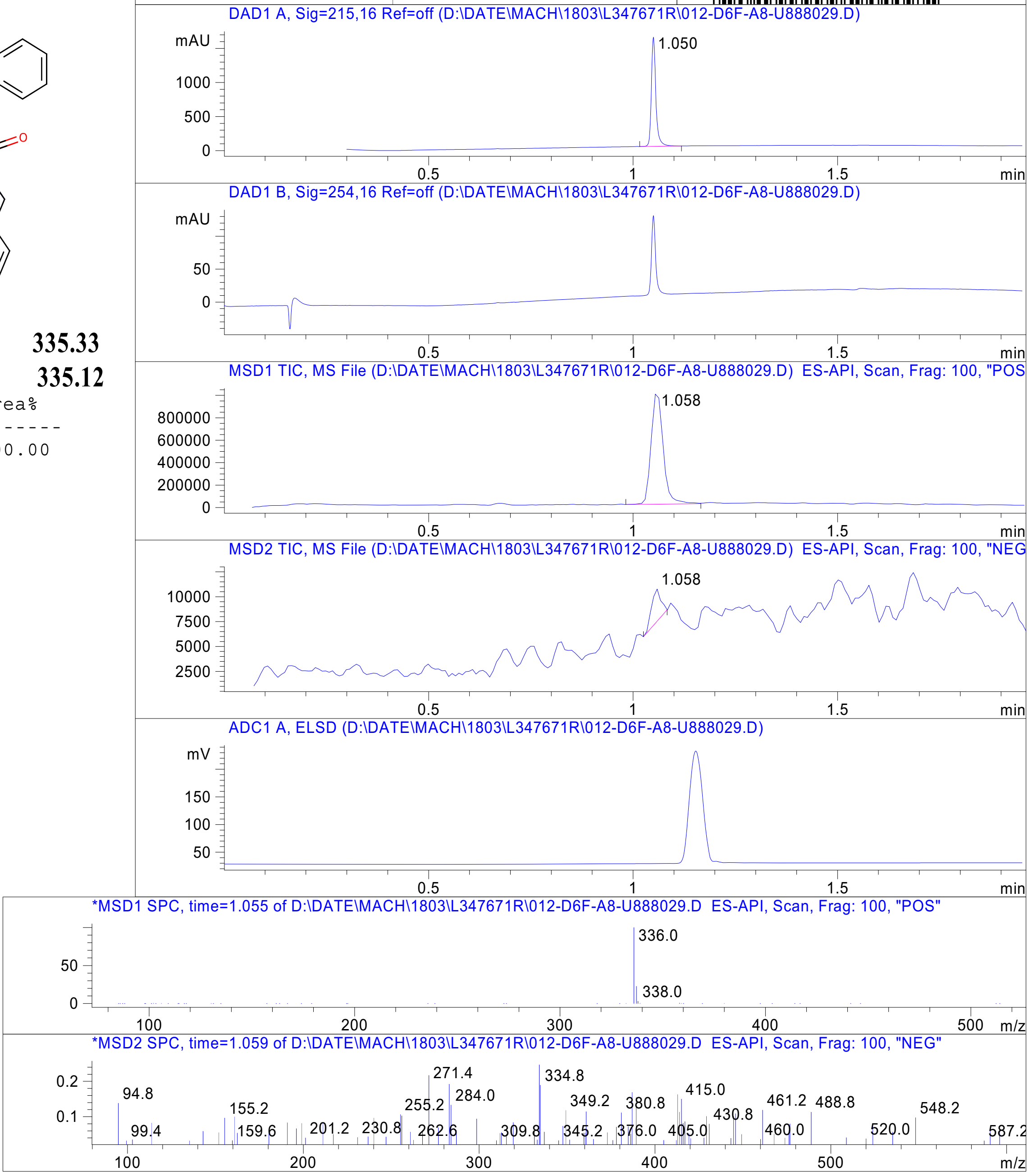


RT 1.058

RT 1.058



Z4929615548_

MaxPeak: 100.00%
Ret_Time: 1.005 min

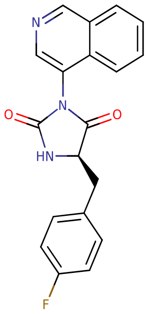


**Mol Wt** 335.33
**Exact Mass** 335.12

| # | Time | Area% |
|---|---|---|
| 1 | 1.005 | 100.00 |

U887628

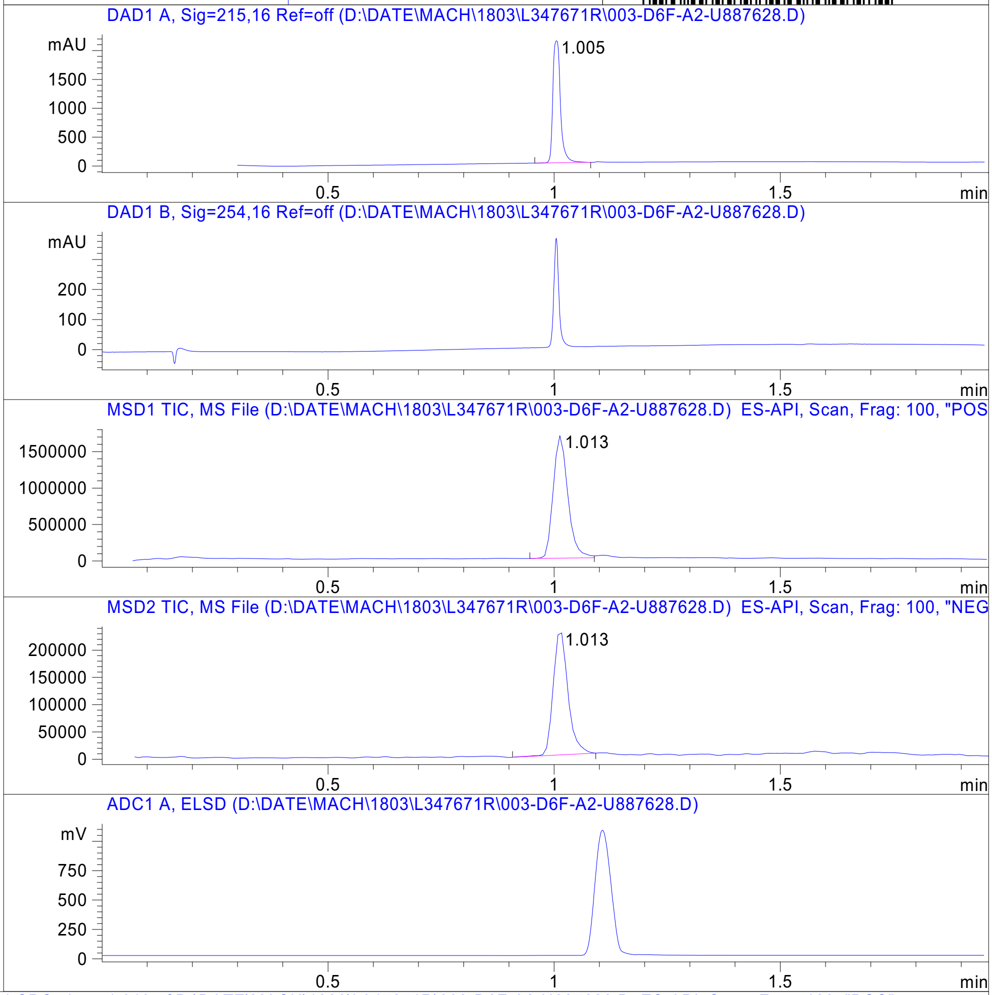


RT 1.013

RT 1.013

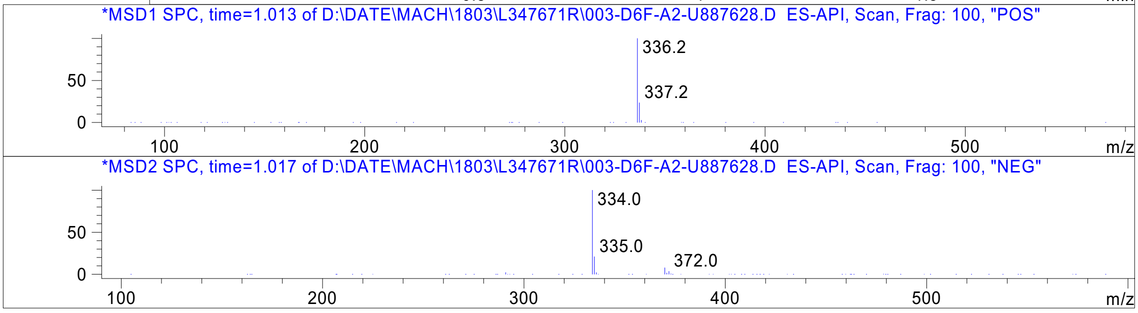




Z4929615788_

MaxPeak:  100.00%
Ret_Time: 1.004 min

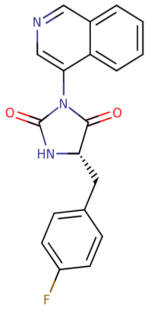


**Mol Wt** 335.33

**Exact Mass** 335.12

| # | Time | Area% |
|---|---|---|
| 1 | 1.004 | 100.00 |

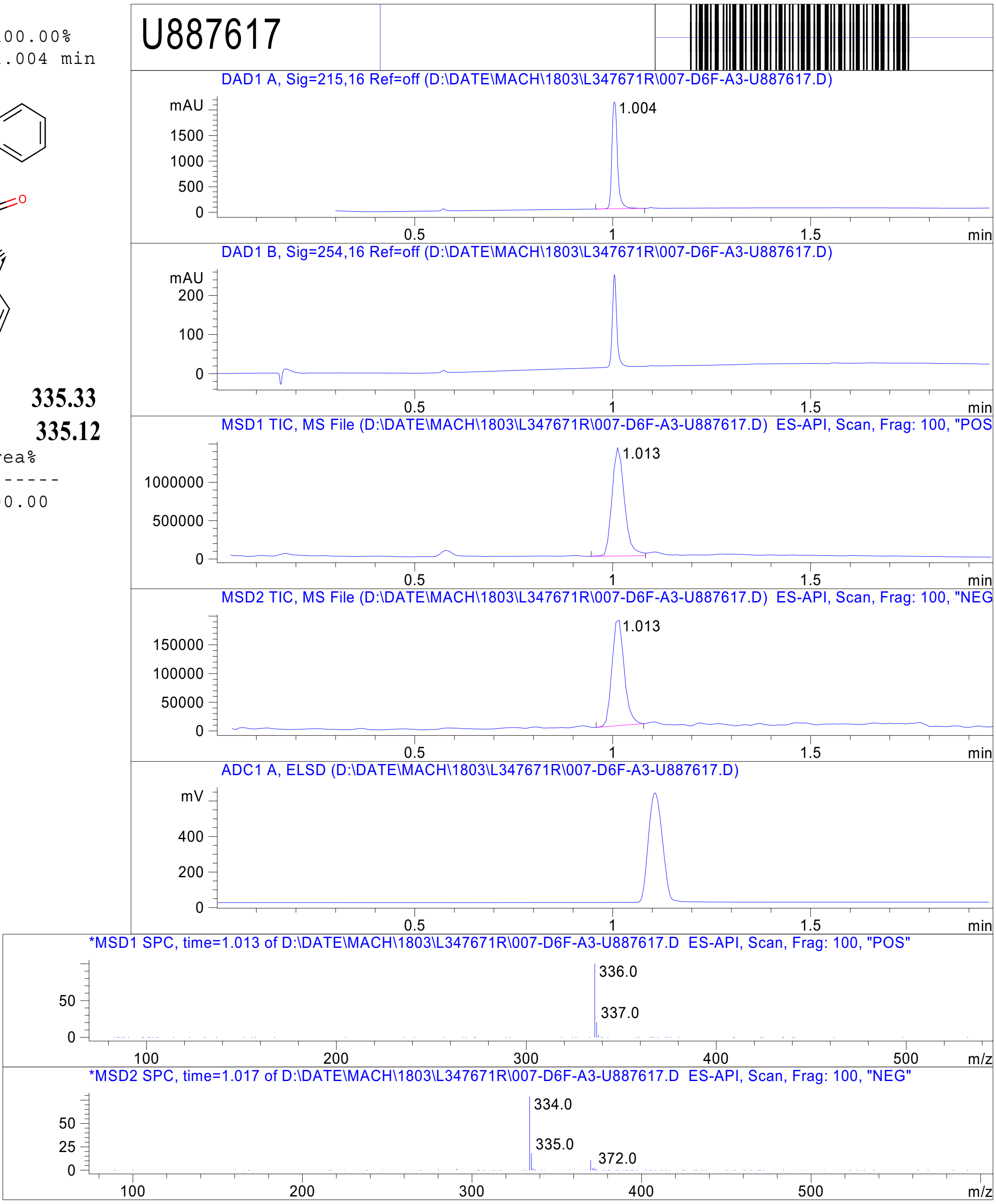


RT 1.013

RT 1.013



Z4929615650_

MaxPeak: 57.60%
Ret_Time: 1.022 min

U887779

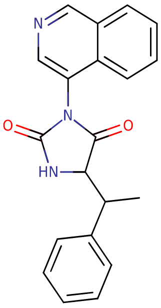


**Mol Wt** 331.37
**Exact Mass** 331.15

| # | Time | Area% |
|---|---|---|
| 1 | 1.022 | 57.60 |
| 2 | 1.044 | 42.40 |

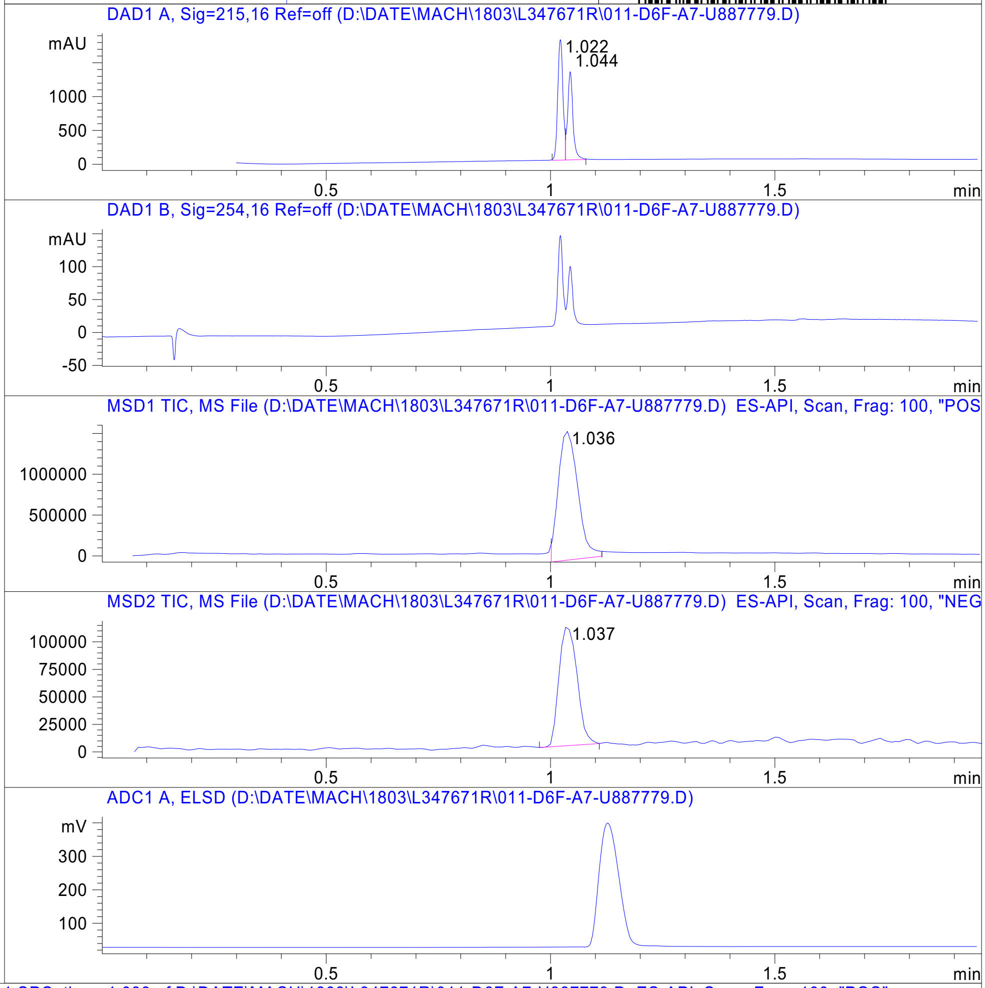


RT 1.036

RT 1.037

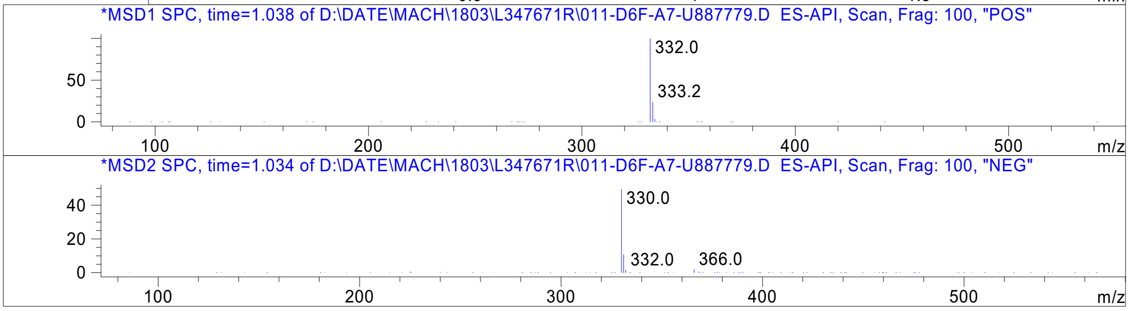




Z4929615582_

MaxPeak: 100.00%
Ret_Time: 1.093 min

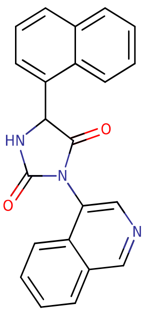


**Mol Wt** 353.37

**Exact Mass** 353.13

| # | Time | Area% |
|---|---|---|
| 1 | 1.093 | 100.00 |

U887831

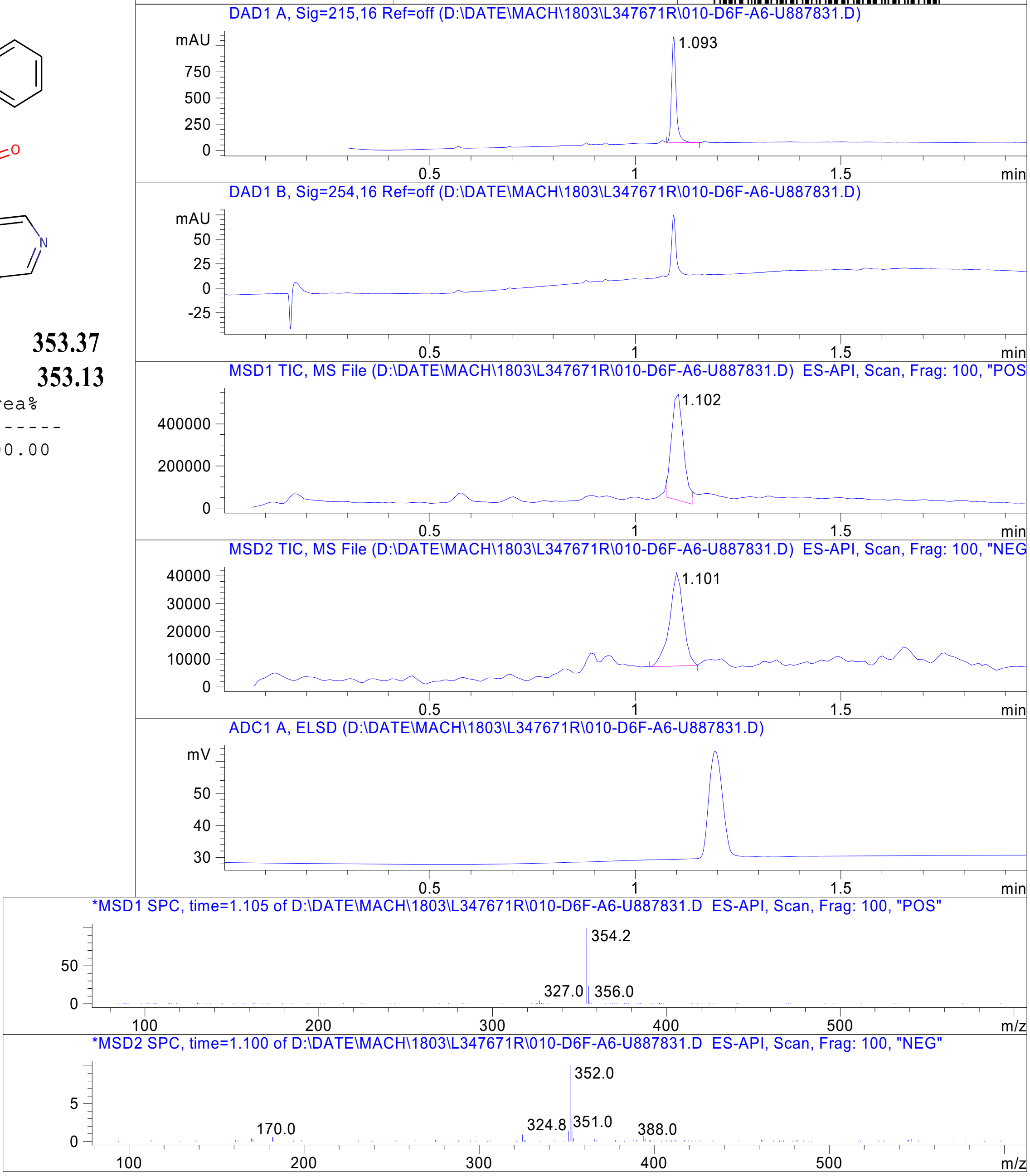


RT 1.102

RT 1.101

Inj.Date 3/17/2021 Y -8- Acq. Method C:\Chem32\-> ->

Z4929615577_

MaxPeak: 100.00%
Ret_Time: 1.073 min

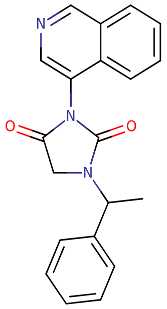

**Mol Wt 331.37**
**Exact Mass 331.15**

| # | Time | Area% |
| --- | --- | --- |
| 1 | 1.073 | 100.00 |

U888088

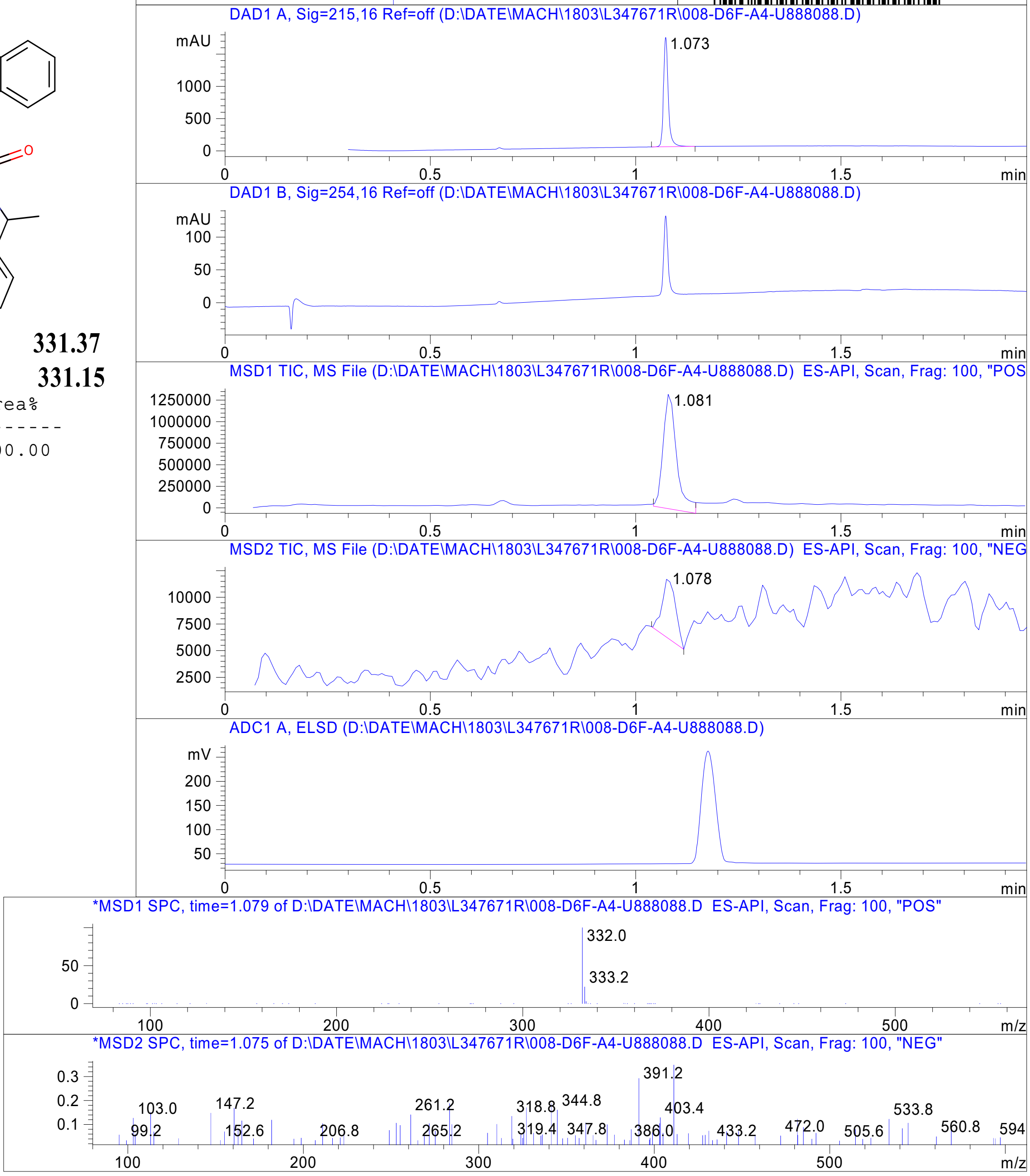


Inj.Date 3/17/2021 Y -8- Acq. Method C:\Chem32\-> ->

Z4925123498_

MaxPeak: 100.00%
Ret_Time: 1.177 min

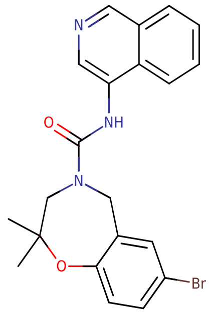


**Mol Wt** 426.31

**Exact Mass** 427.1

| # | Time | Area% |
|---|---|---|
| 1 | 1.177 | 100.00 |

V013590$2

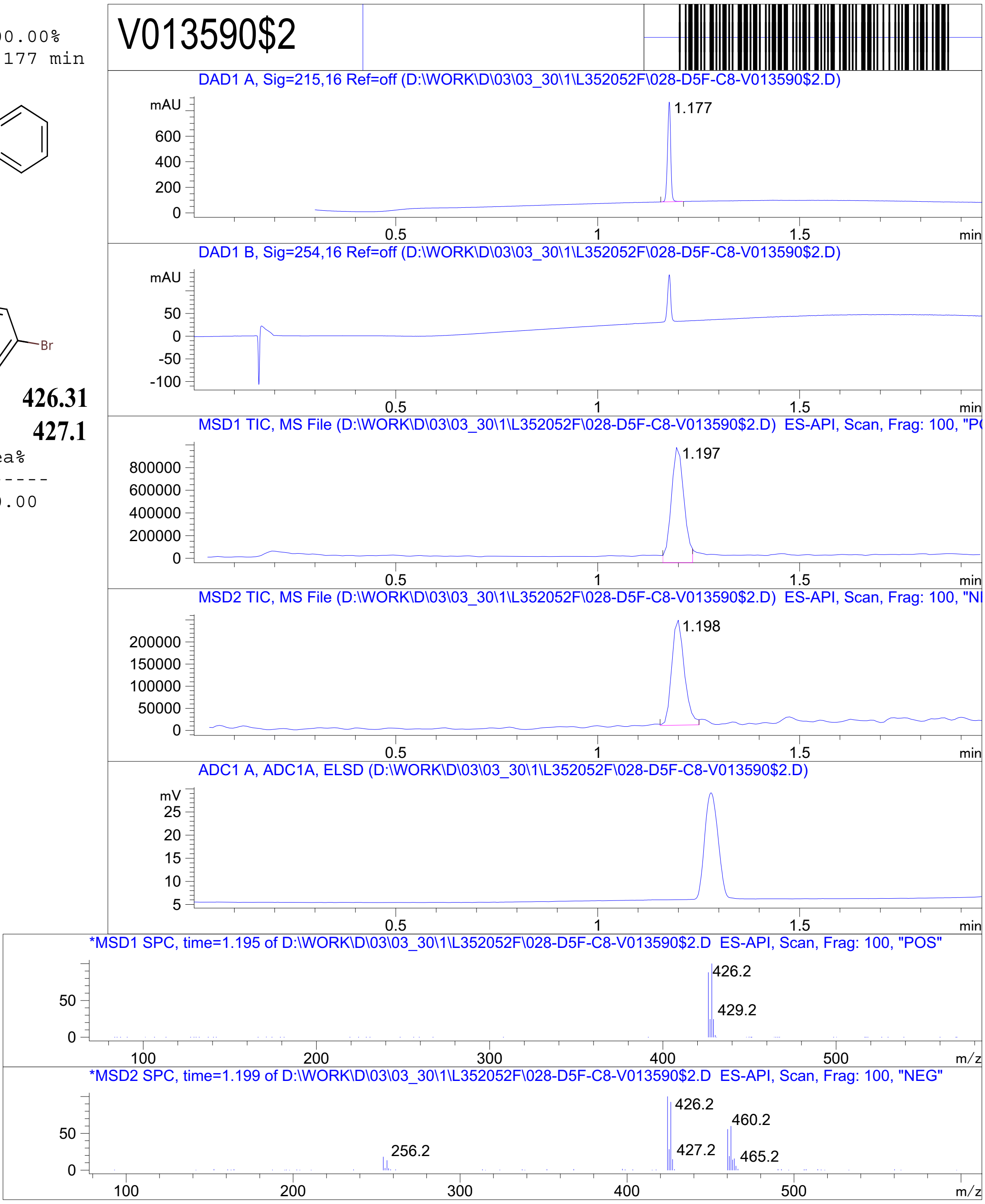


RT 1.197

RT 1.198



Z4924562413_

MaxPeak: 100.00%
Ret_Time: 0.828 min

V013589$1

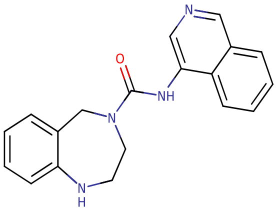

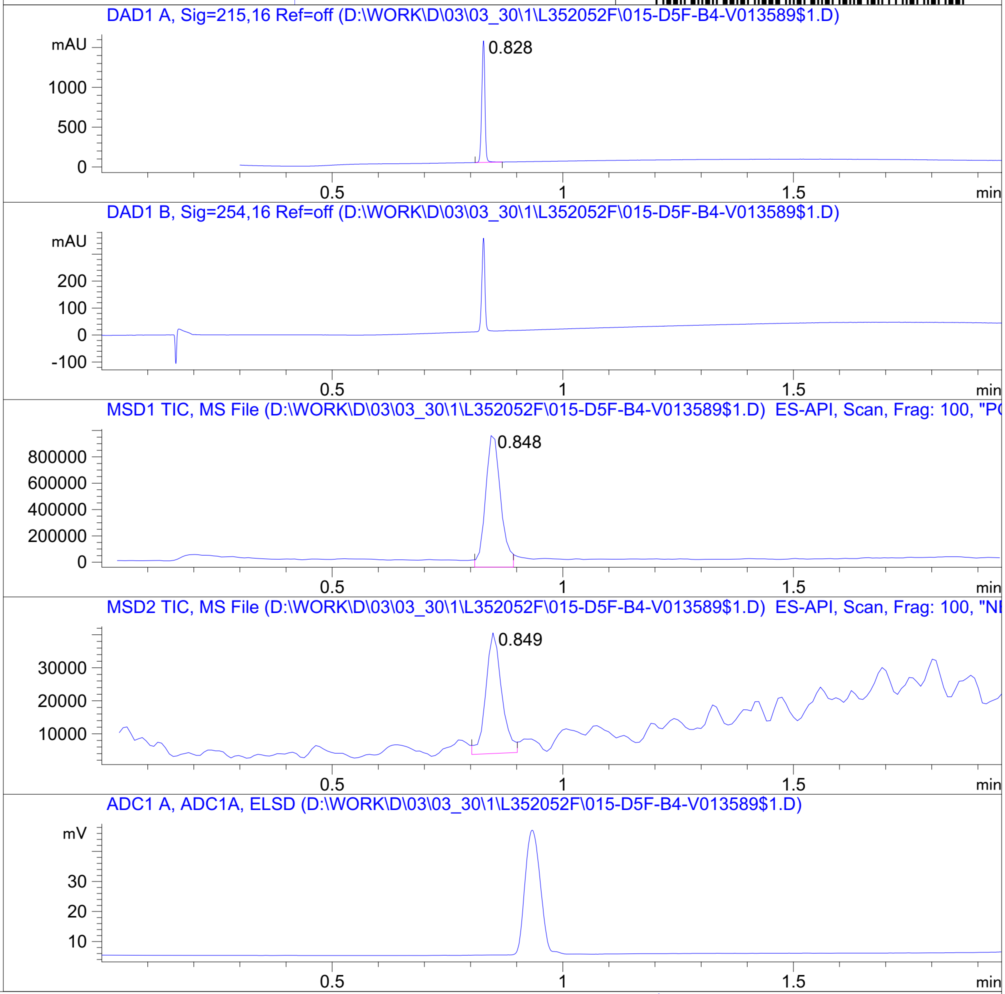


**Mol Wt** 318.37
**Exact Mass** 318.17

| # | Time | Area% |
|---|---|---|
| 1 | 0.828 | 100.00 |

RT 0.848

RT 0.849

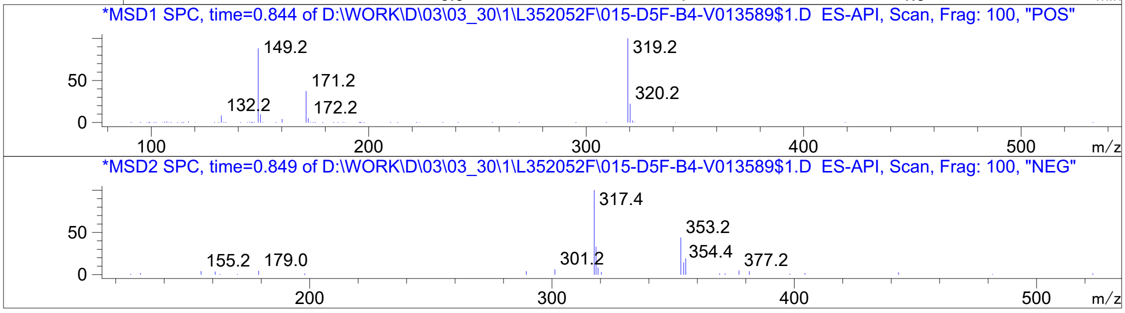


Inj.Date 3/30/2021 K -25- Acq. Method C:\Users\ -> ->

Z4924562415_

MaxPeak: 100.00%
Ret_Time: 2.443 min

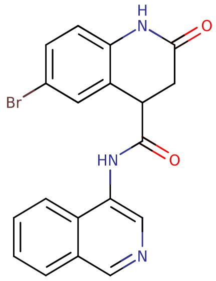


**Mol Wt** 396.24

**Exact Mass** 395.04

| # | Time | Area% |
|---|---|---|
| 1 | 2.443 | 100.00 |

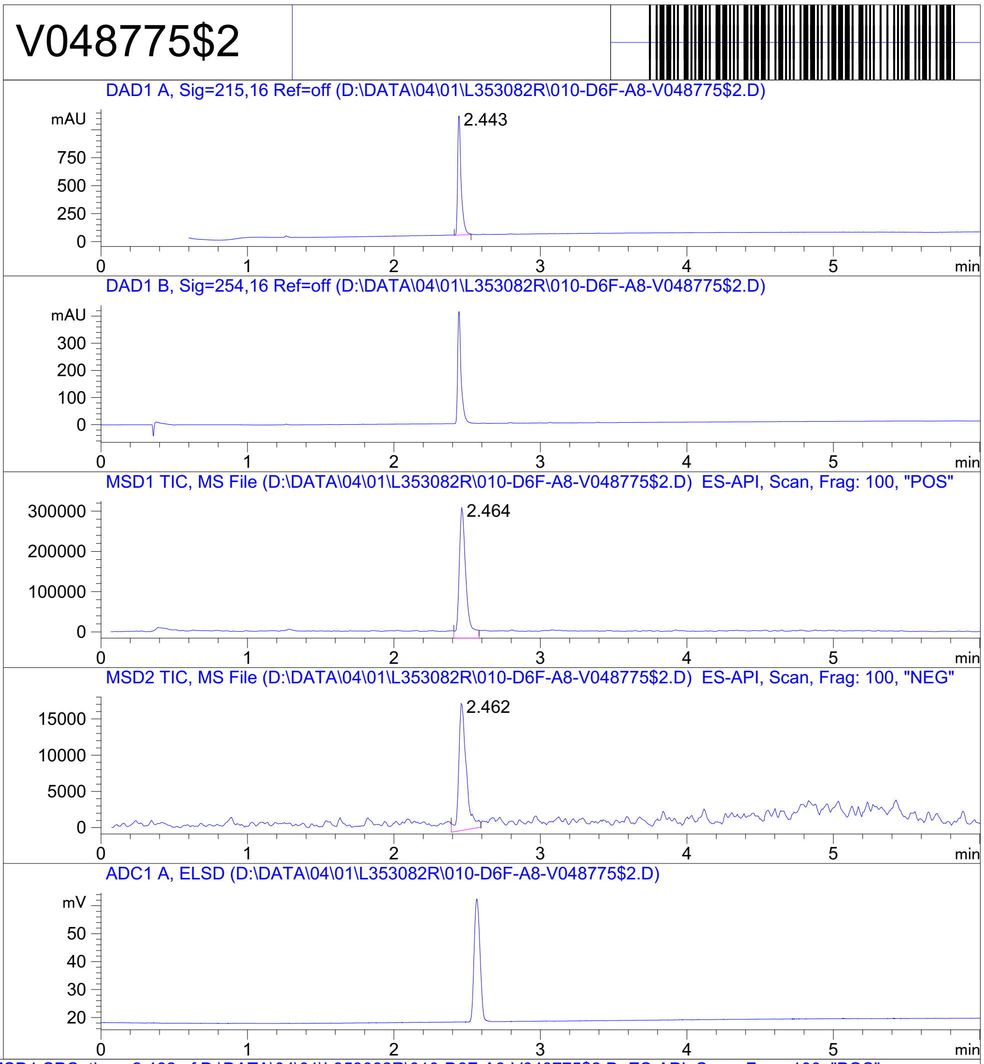


RT 2.464

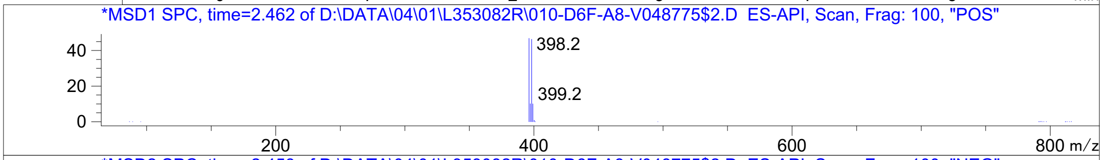


RT 2.462

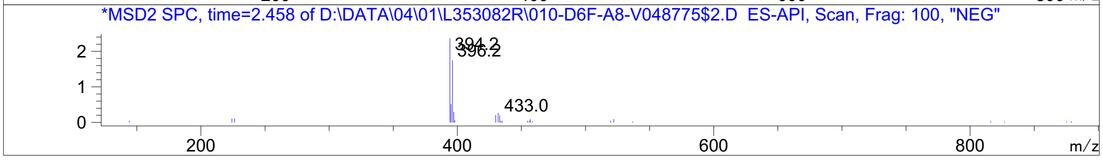


Inj.Date 4/1/2021 LB  Acq. Method C:\Chem32\-> ->

Z1892658668_

MaxPeak: 98.67%
Ret_Time: 2.380 min

# V053006$1

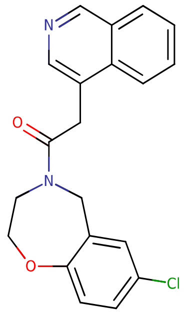

**Mol Wt** 352.81

**Exact Mass** 352.12

| # | Time | Area% |
|---|---|---|
| 1 | 2.380 | 98.67 |
| 2 | 3.345 | 1.33 |

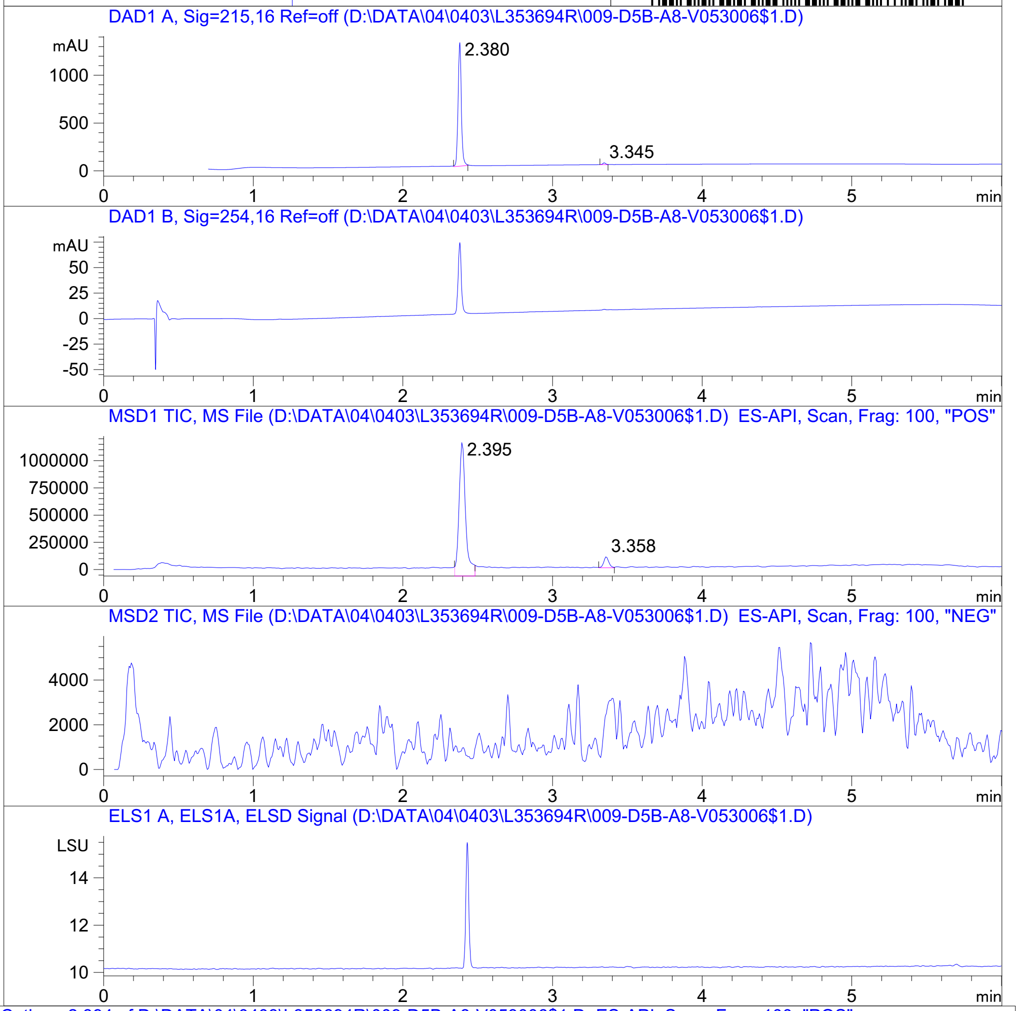


RT 2.395

RT 3.358

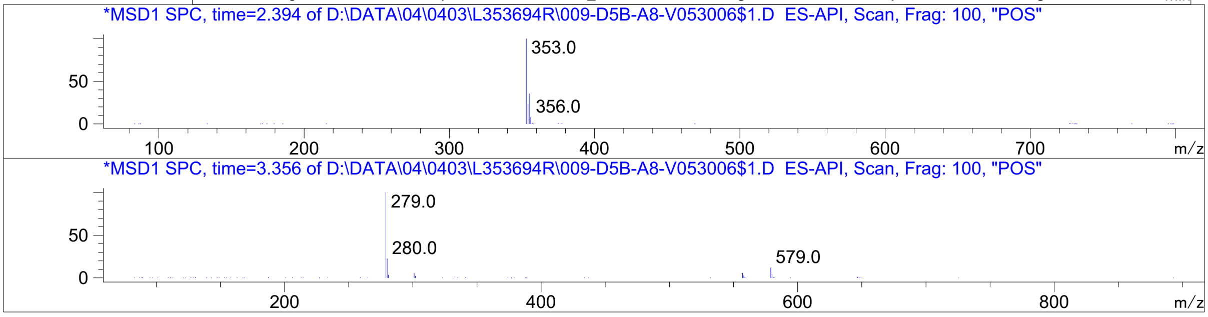




Z4946671001_

MaxPeak: 100.00%
Ret_Time: 1.393 min

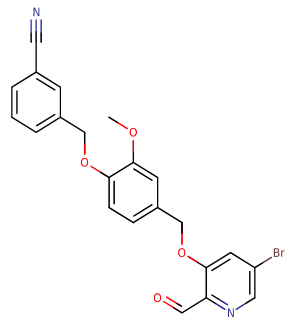

**Mol Wt** 453.28

**Exact Mass** 454.05

| # | Time | Area% |
|---|---|---|
| 1 | 1.393 | 100.00 |

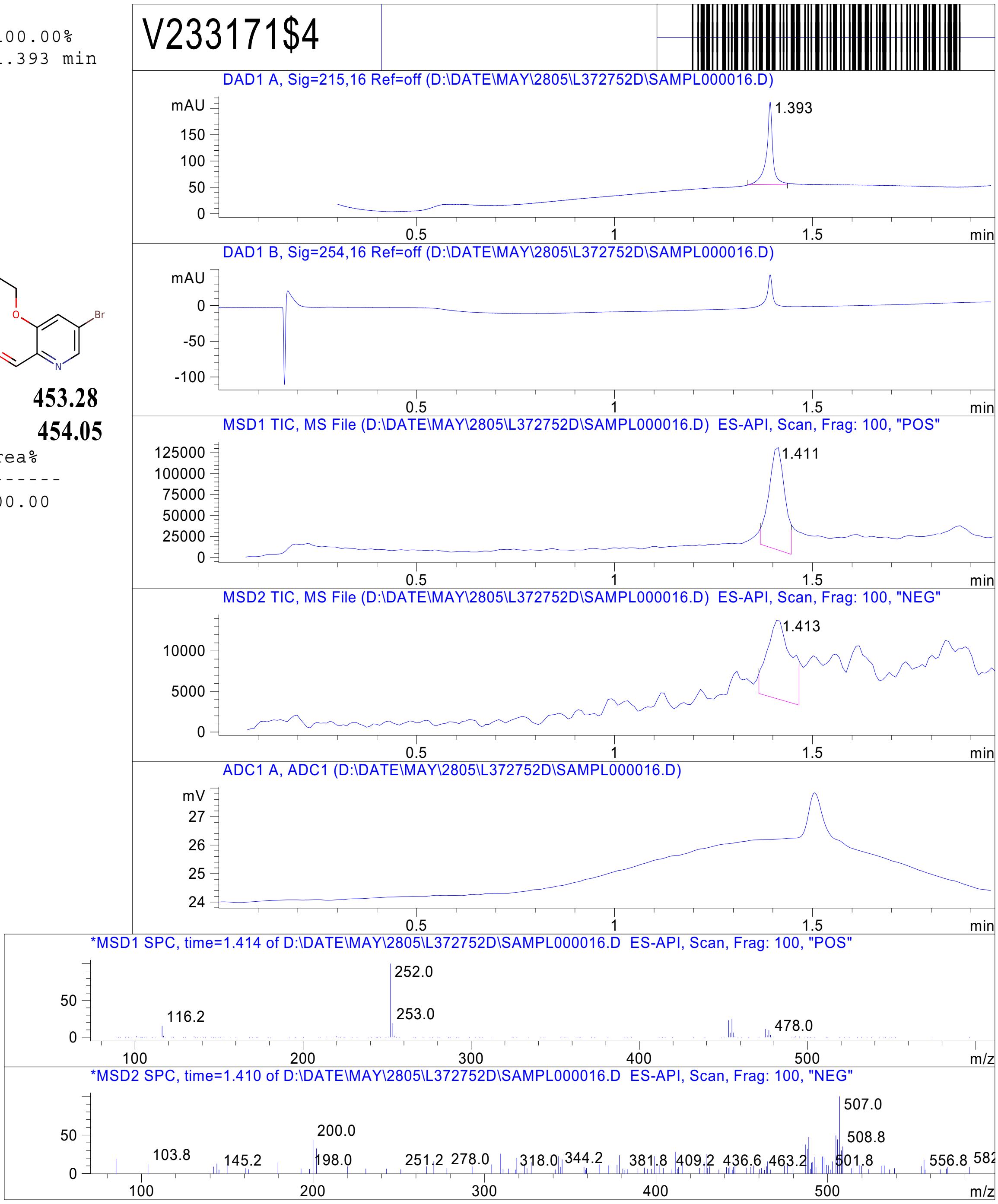


RT 1.411

RT 1.413

Inj.Date 5/28/2021 Y - 4 - Acq. Method C:\CHEM32\-> ->

Z5009000161_

MaxPeak: 100.00%
Ret_Time: 1.483 min

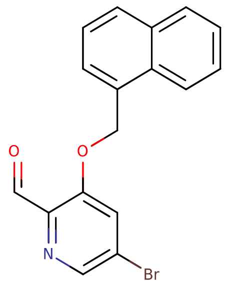


**Mol Wt** 342.19
**Exact Mass** 341.02

| # | Time | Area% |
|---|---|---|
| 1 | 1.483 | 100.00 |

# V608763$7

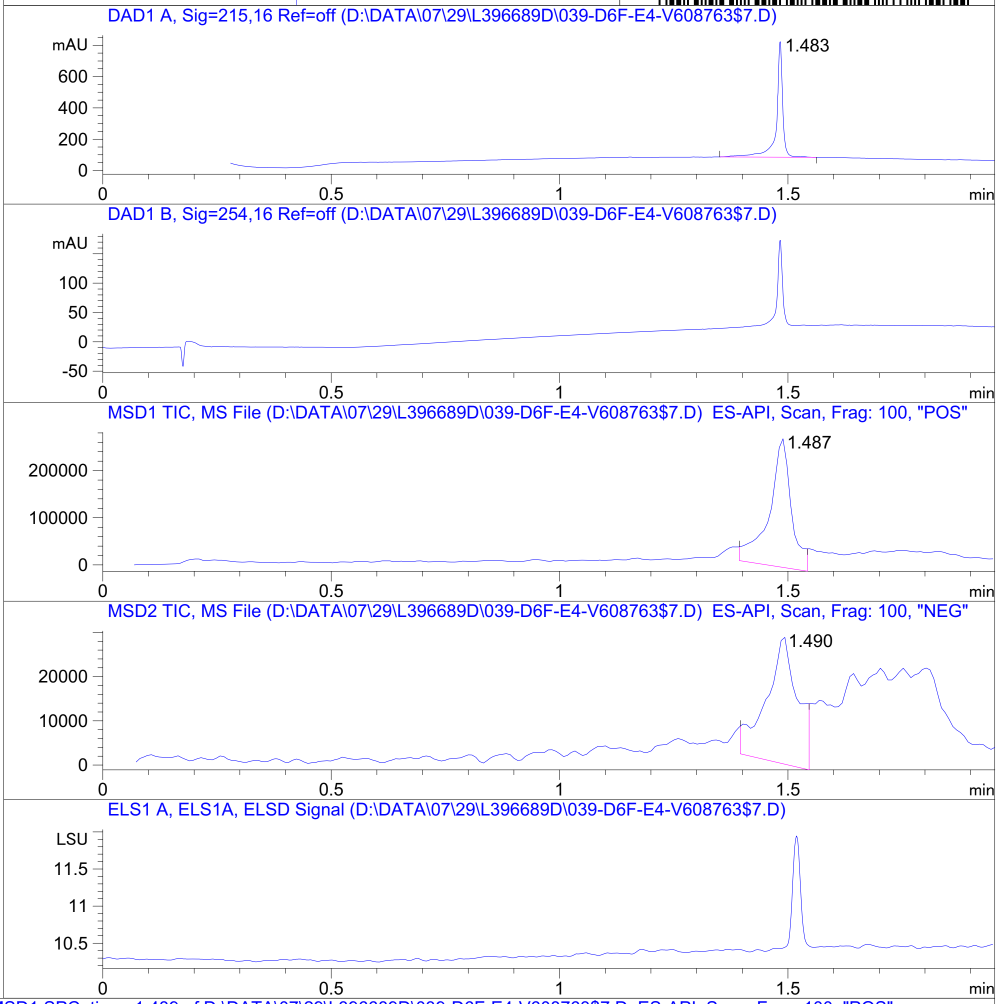


RT 1.487

RT 1.490

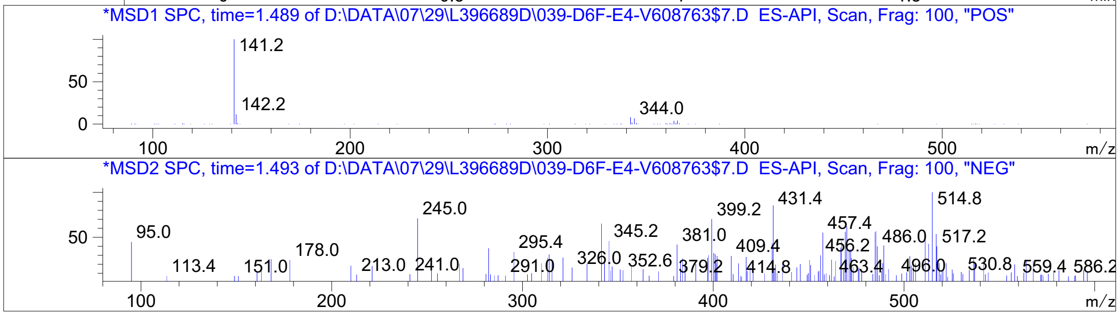




Z4989203251_

MaxPeak: 100.00%
Ret_Time: 1.488 min

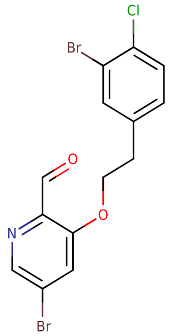


**Mol Wt** 419.5
**Exact Mass** 418.89

| # | Time | Area% |
|---|---|---|
| 1 | 1.488 | 100.00 |

W316824$12

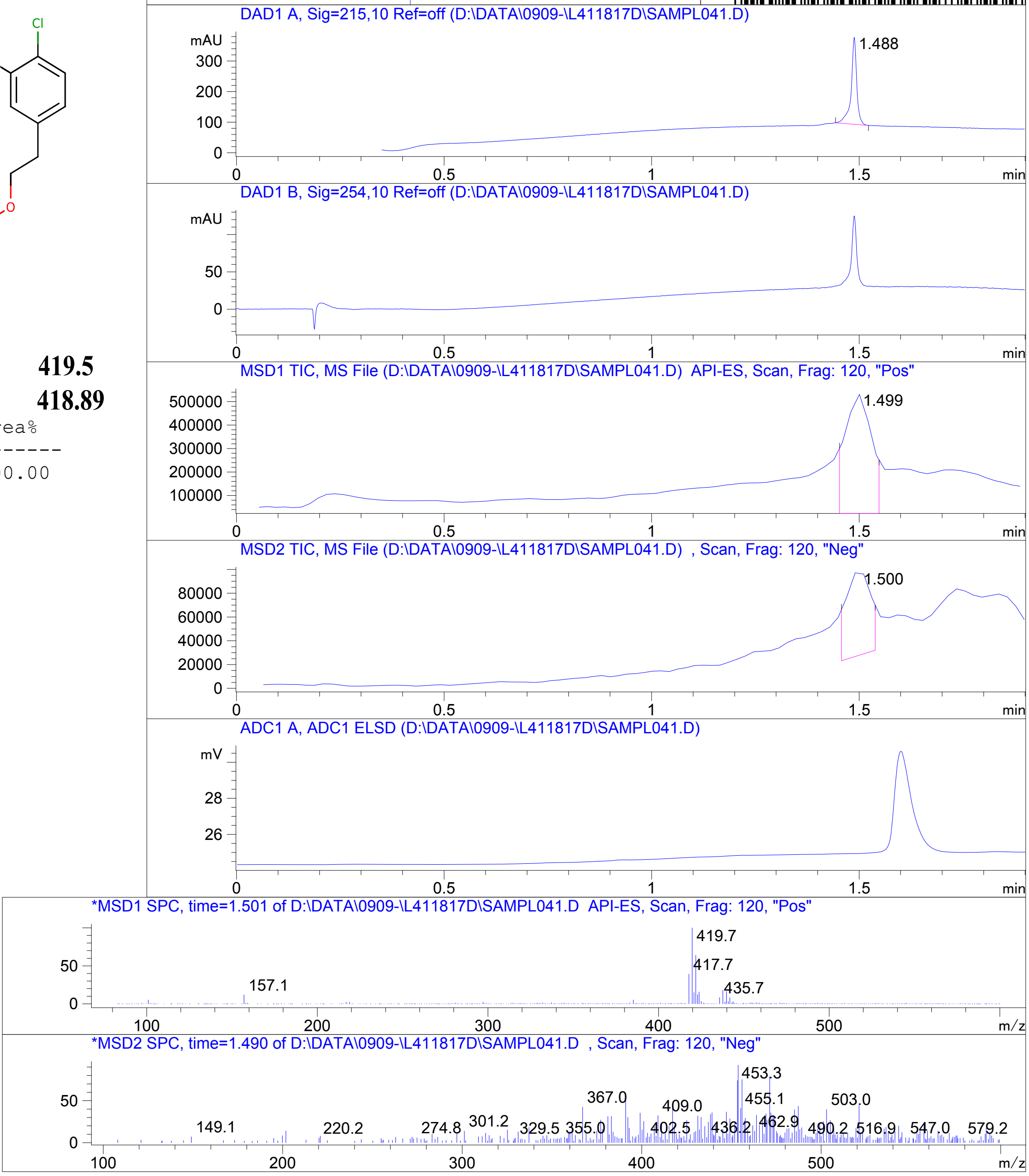


RT 1.499

RT 1.500

Inj.Date 9/9/2021 OA -VL- Acq. Method C:\HPCHEM\-> ->

Z5192779061_

MaxPeak: 100.00%
Ret_Time: 1.377 min

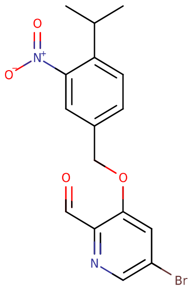


**Mol Wt** 379.2
**Exact Mass** 378.03

| # | Time | Area% |
|---|---|---|
| 1 | 1.377 | 100.00 |

# W495913$1

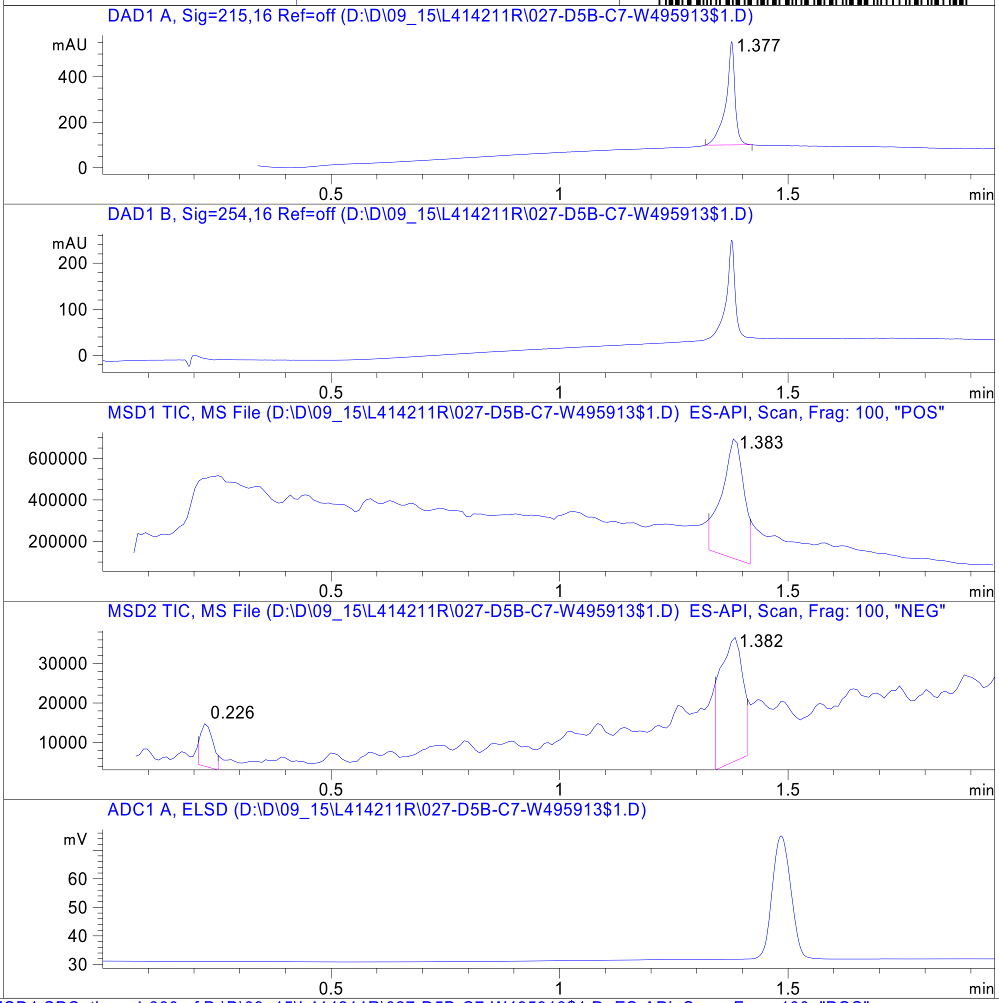


RT 1.383

RT 0.226

RT 1.382

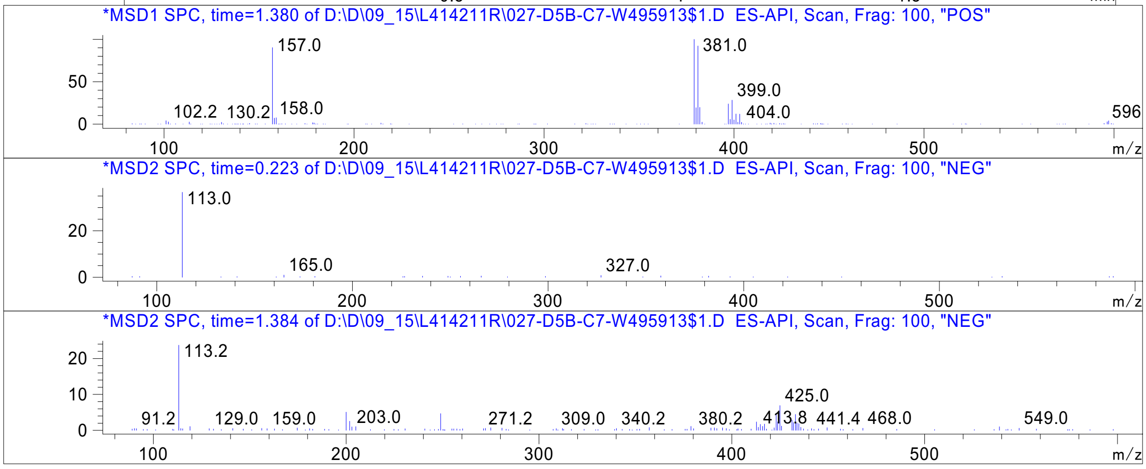


Inj.Date 9/15/2021 N  Acq. Method C:\Chem32\-> ->

Z5247643566_

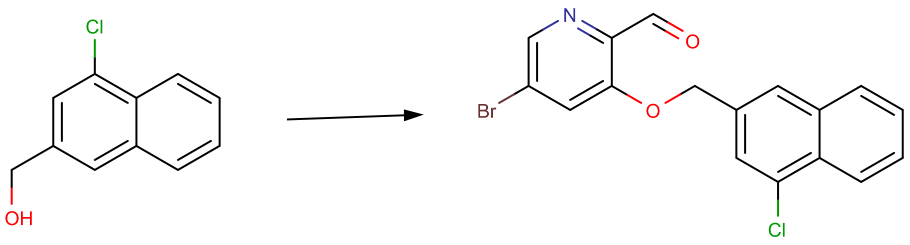


[192.03]

[374.97]

# V942078-1

L411799F
LCMS-16
SUPOR_30.M
09:49 09.09.2021
MaxPeak: 100.0%

| # | RT | DAD1A | DAD1B | MSD1 | MSD2 | ELSD | MSD1 ions | MSD1 rt | MSD2 ions | MSD2 rt | Info |
|---|---|---|---|---|---|---|---|---|---|---|---|
| 1 | 1.549 | 100.0% | 100.0% | 100.0% | --- | 100.0% | 175.0(51),377.8(26),375.8(16) | 1.554 | --- | --- | Reagent1 -OH,P +H+ |

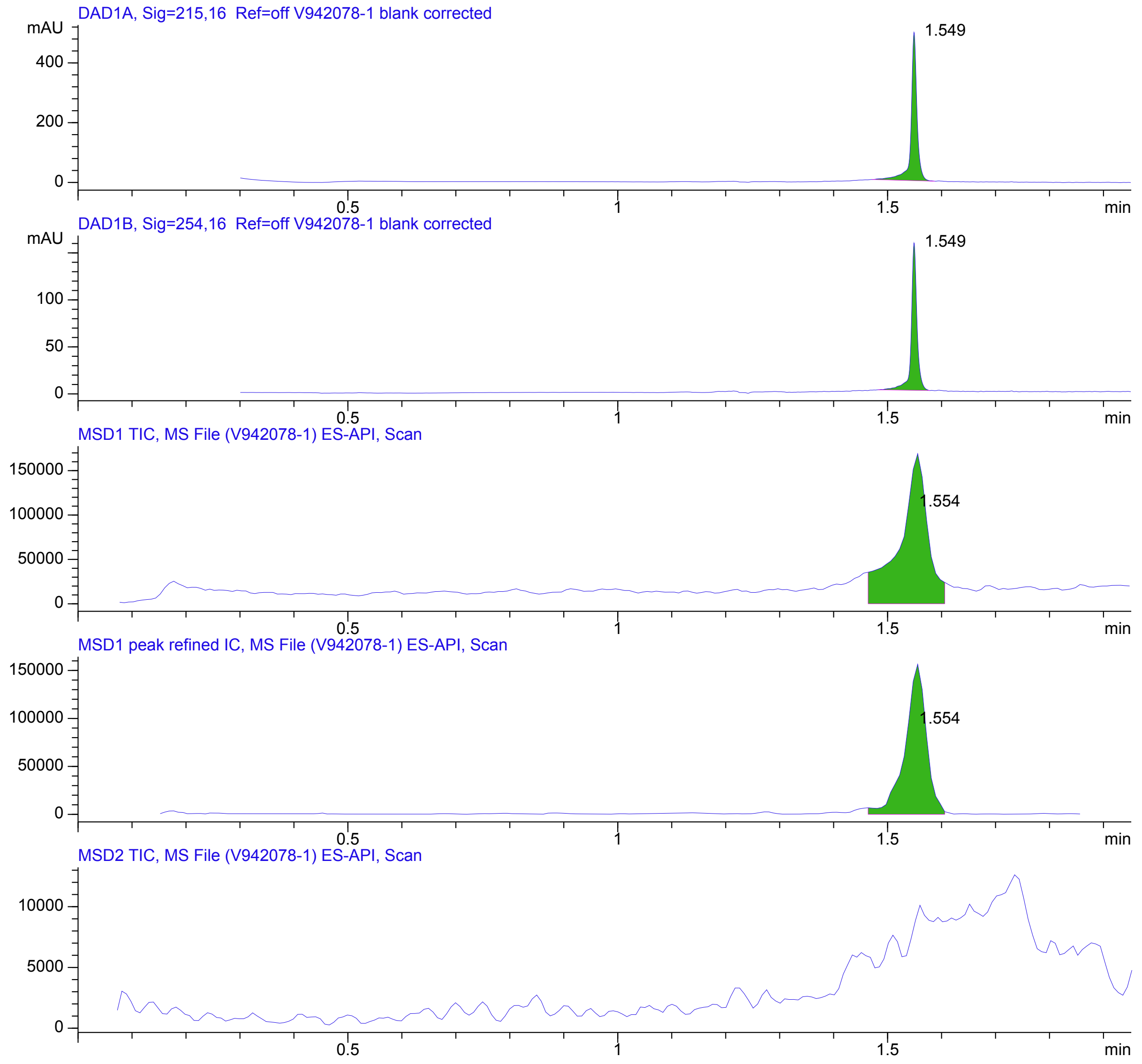


ICI: 19768
ELSD shift 2269 | +350 / -625
MSD shift 410 | +300 / -300
c:\ARCH\2021\16\09\09_09\L411799F\003-NV-B.D
c:\ARCH\2021\16\09\09_09\L411799F\005-D5F-A2-V942078-1.D
https://1-042-bot.work.local/alpha?instrument=LCMS-16&acquiredDate=1631170143000&barcode=V942078-1



Z5084954202_

MaxPeak: 100.00%
Ret_Time: 1.396 min

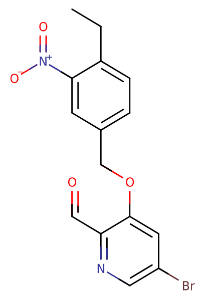

**Mol Wt** 365.18
**Exact Mass** 364.01

| # | Time | Area% |
|---|---|---|
| 1 | 1.396 | 100.00 |

W529955$2

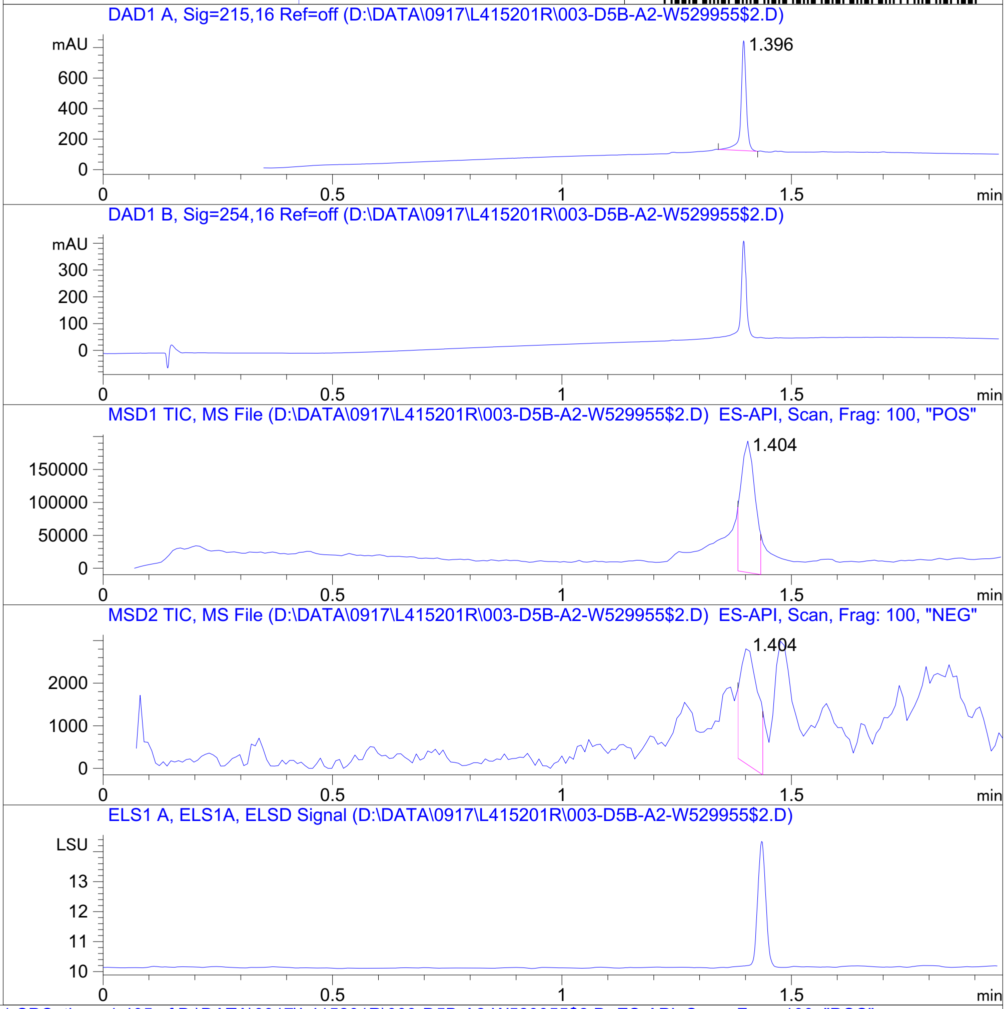


RT 1.404

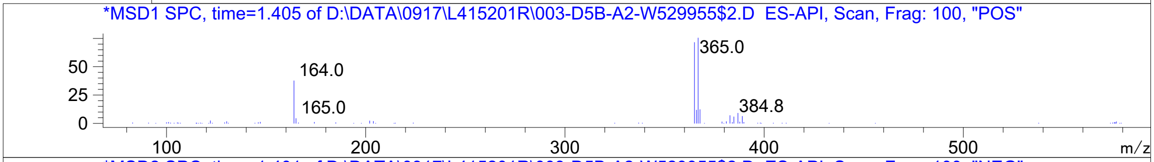


RT 1.404

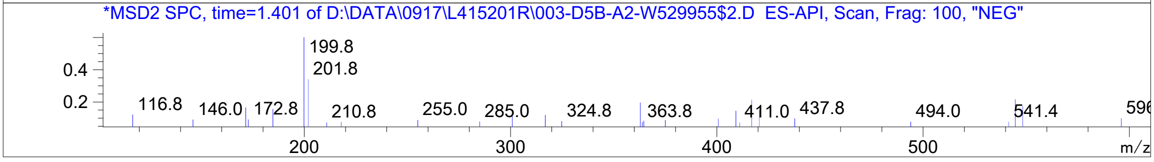


Inj.Date 9/17/2021 H 13 Acq. Method C:\Chem32\-> ->

Z5247643564_

MaxPeak: 100.00%
Ret_Time: 1.226 min

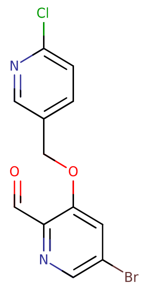


**Mol Wt** 327.56
**Exact Mass** 327.95

| # | Time | Area% |
|---|---|---|
| 1 | 1.226 | 100.00 |

U920104$26

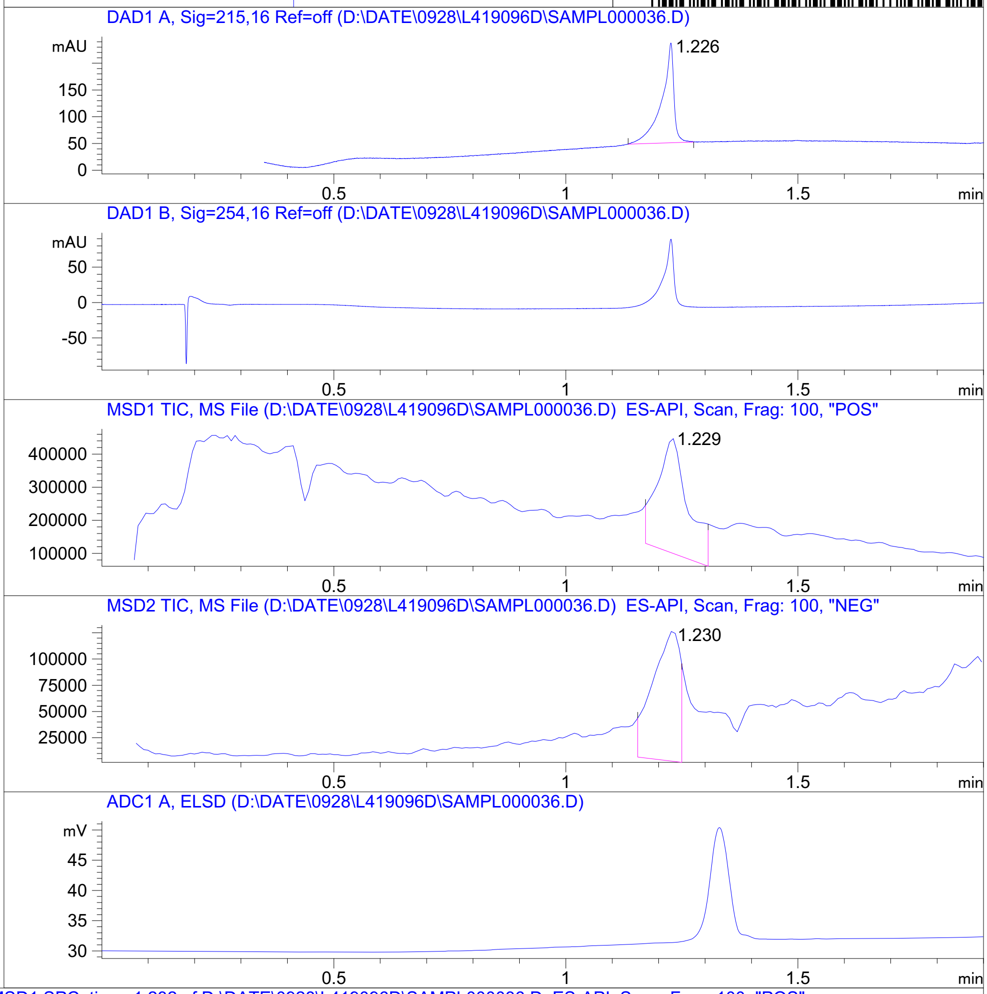


RT 1.229

RT 1.230

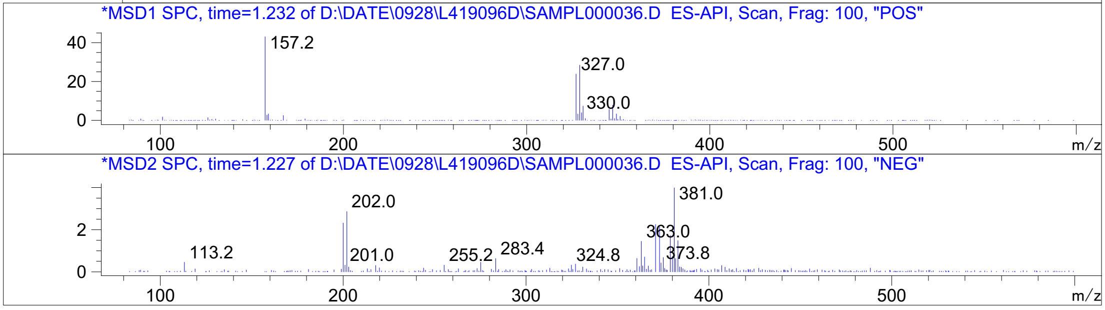




Z4954569140_

MaxPeak: 100.00%
Ret_Time: 1.300 min

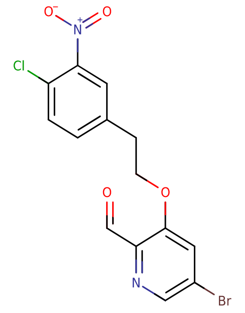

**Mol Wt** 385.6
**Exact Mass** 385.95

| # | Time | Area% |
| --- | --- | --- |
| 1 | 1.300 | 100.00 |

W828537$5

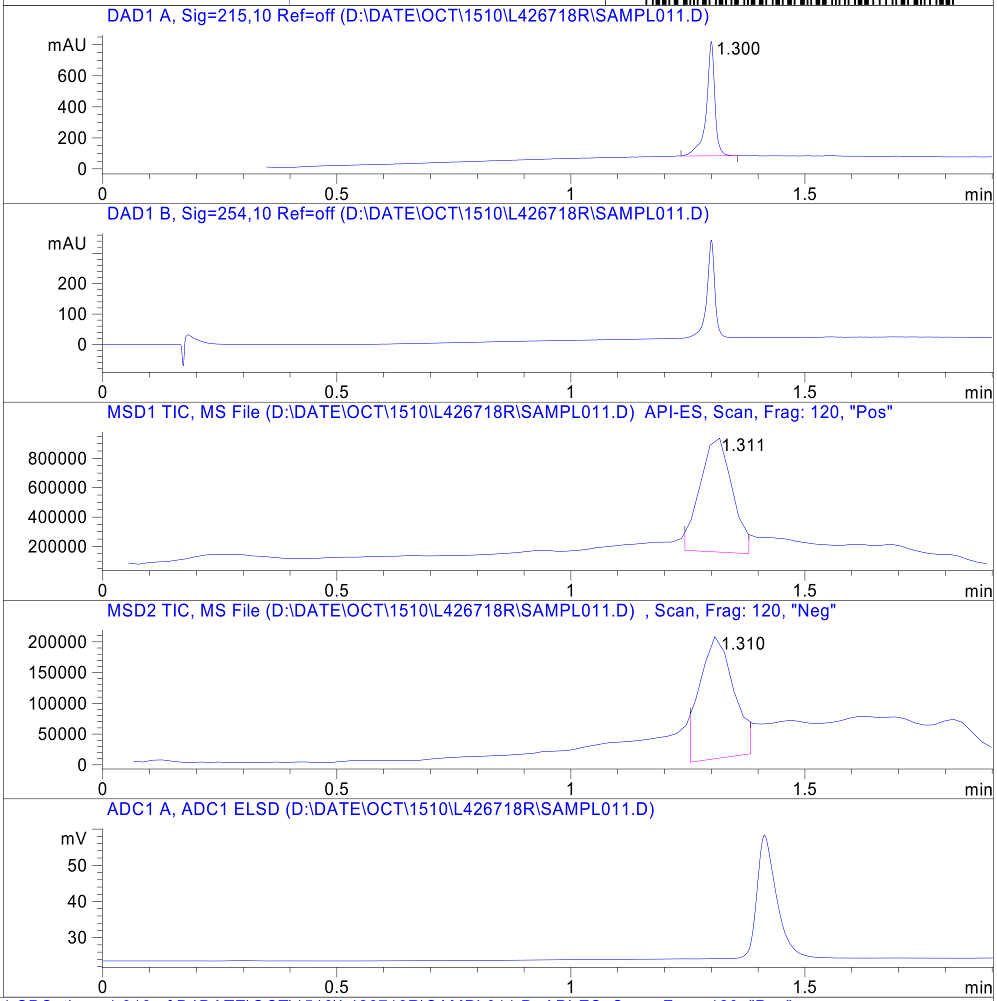


RT 1.311

RT 1.310

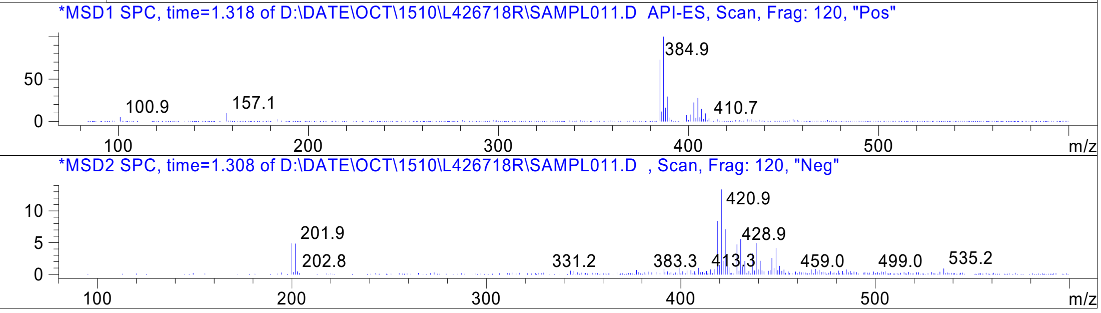


Inj.Date 10/15/2021 Y -VL- Acq. Method C:\HPCHEM\-> ->

Z5192779054_

MaxPeak: 100.00%
Ret_Time: 1.528 min

# W938418$2

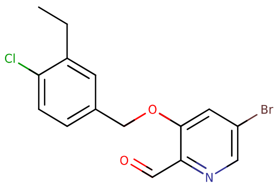


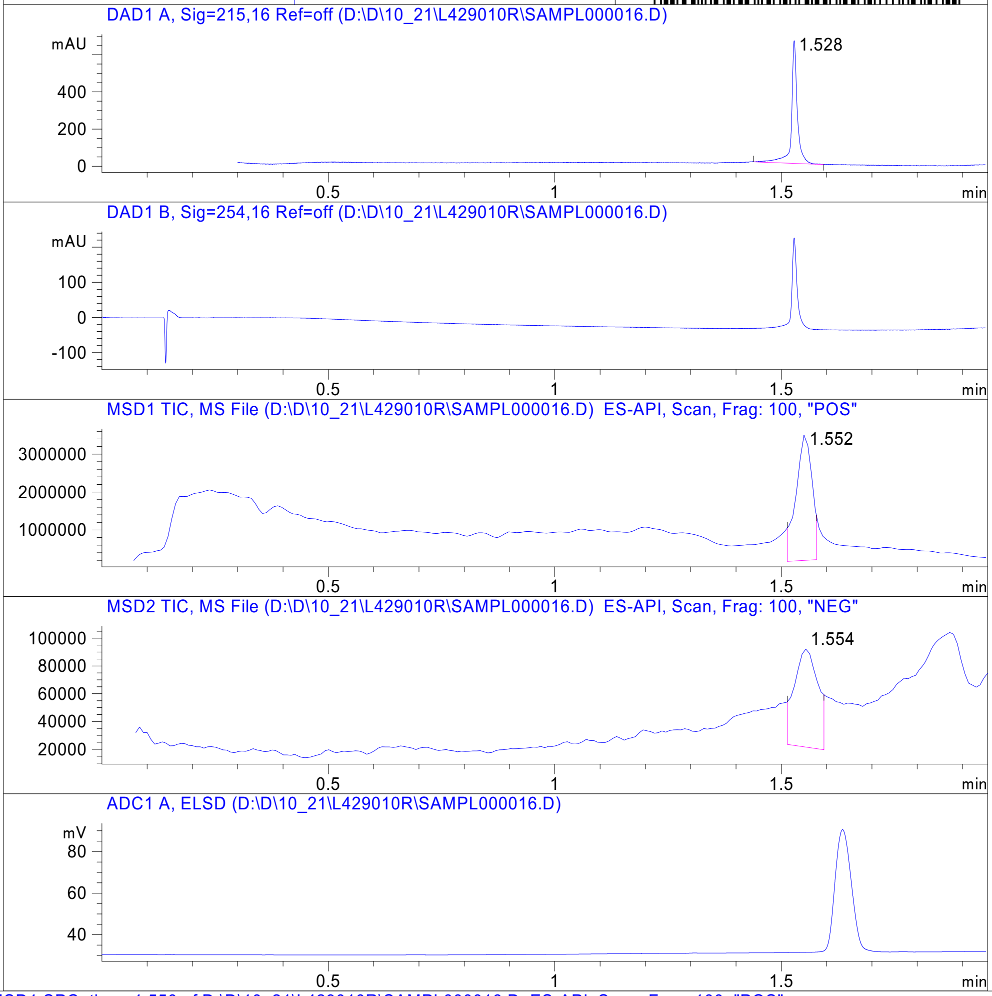


**Mol Wt** 354.63
**Exact Mass** 355

| # | Time | Area% |
|---|---|---|
| 1 | 1.528 | 100.00 |

RT 1.552

RT 1.554

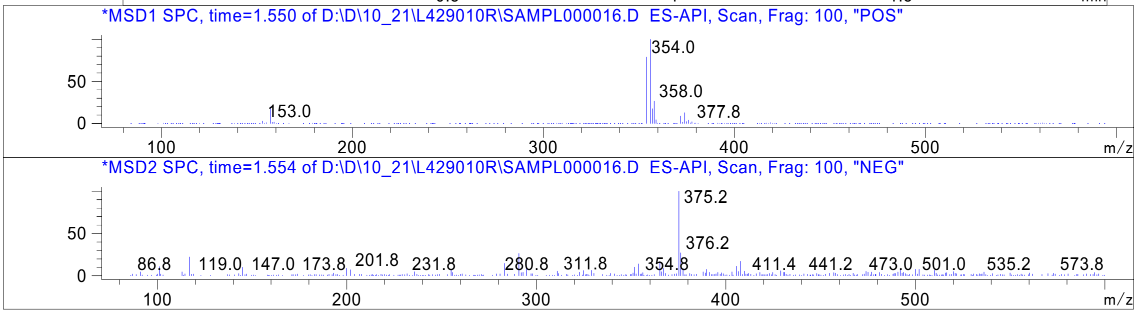




Z5192779014_

MaxPeak: 100.00%
Ret_Time: 1.483 min

# W976013$2

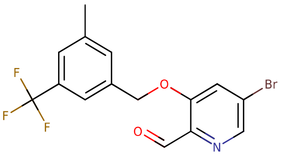


**Mol Wt** 374.15
**Exact Mass** 373.01

| # | Time | Area% |
|---|---|---|
| 1 | 1.483 | 100.00 |

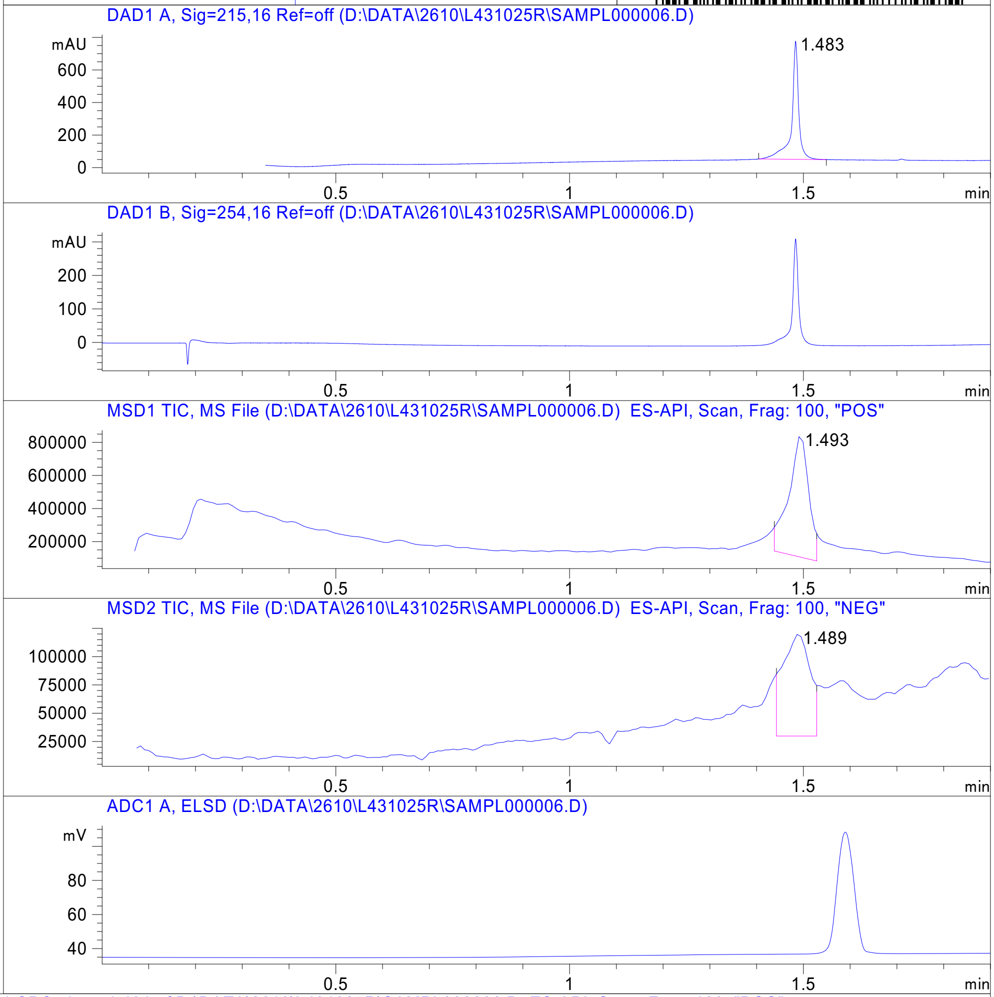


RT 1.493

RT 1.489

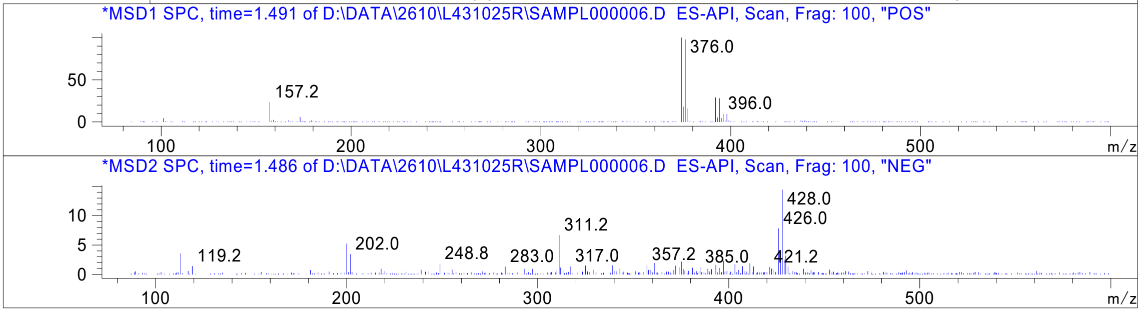




Z5336571669_

MaxPeak: 100.00%
Ret_Time: 1.325 min

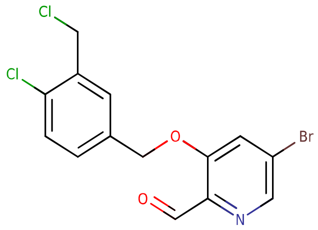

**Mol Wt** 375.04
**Exact Mass** 374.94

| # | Time | Area% |
|---|---|---|
| 1 | 1.325 | 100.00 |

# X349051$1

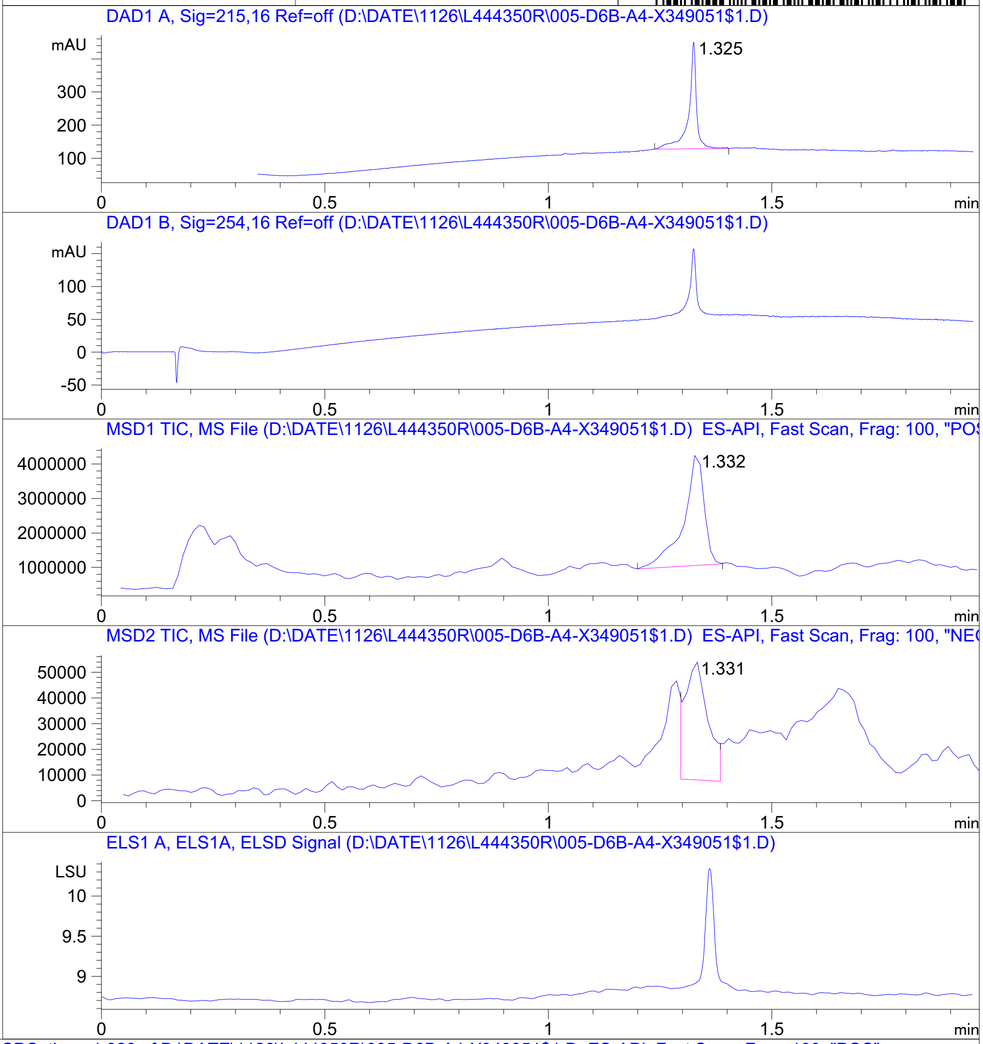


RT 1.332

RT 1.331

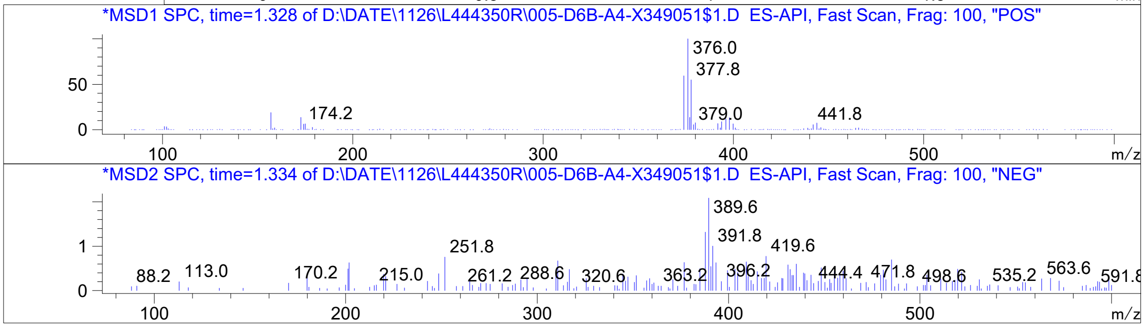


Inj.Date 11/26/2021 E Acq. Method C:\Users\ -> ->

Z5247643558_

### Analysis Report

#### Sample Information

| | | | |
|---|---|---|---|
| **Name** | R2571204 | **Data File Path** | D:\MassHunter\GCMS\1\data\04_20\R2571204.D |
| **Comment** | CH3CN | **Acq. Time (Local)** | 20-Apr-21 22:22:08 (UTC+03:00) |
| **Instrument** | GCMS-5 | **Method Path (Acq)** | D:\MassHunter\GCMS\1\methods\UNIV_ACN.M |
| **Position** | 33 | | |

#### Sample Chromatograms

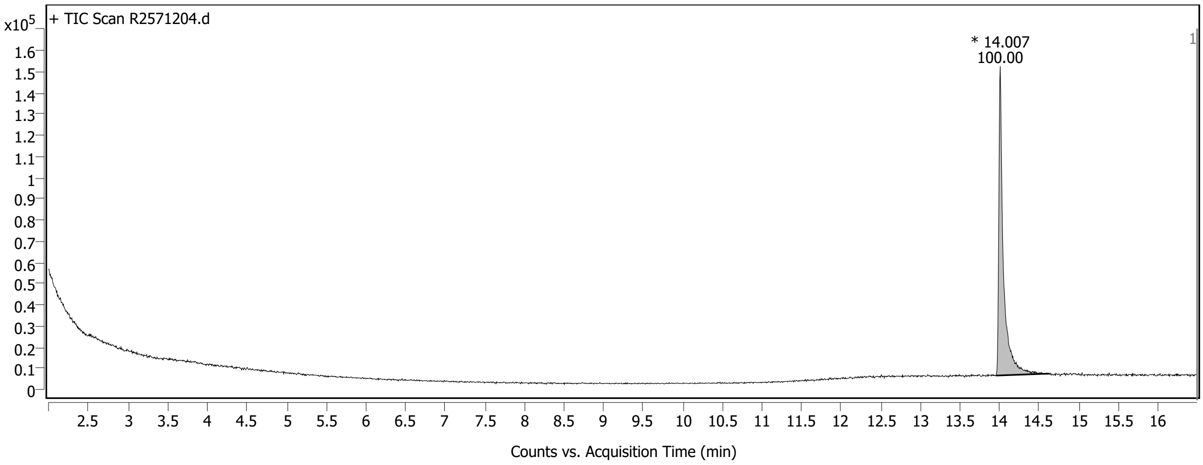


*Chromatogram Peaks*

| Peak | RT | Area | Area Sum % |
|---|---|---|---|
| 1 | 14.007 | 594625 | 100.00 |

**MassHunter Qual 10.0**
**(End of Report)**



Z4954569220_

MaxPeak: 100.00%
Ret_Time: 0.659 min

# LH103289174

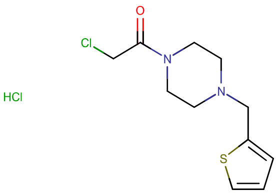


**Mol Wt** 295.23
**Exact Mass** 258.08

| # | Time | Area% |
|---|---|---|
| 1 | 0.659 | 100.00 |

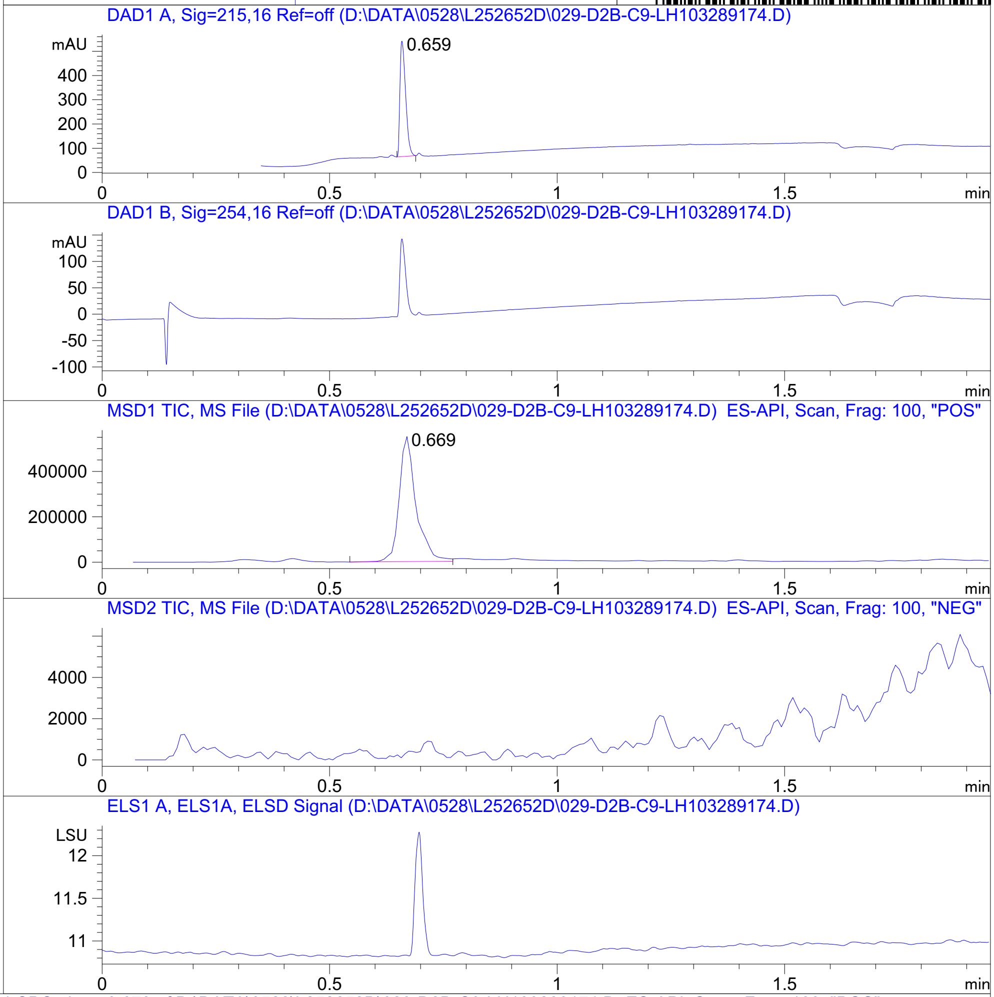


RT 0.669

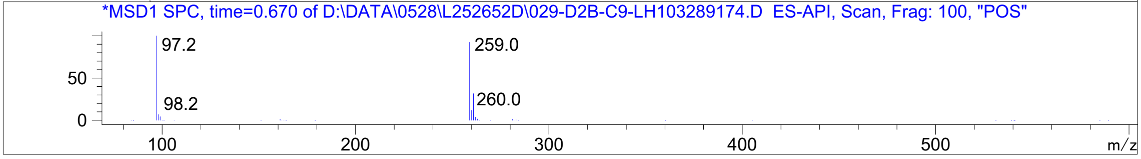




Z147647874_

MaxPeak: 98.15%
Ret_Time: 1.248 min

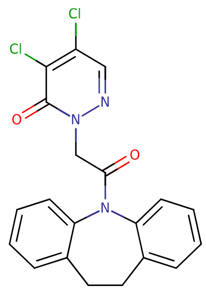

**Mol Wt** 400.26
**Exact Mass** 399.07

| # | Time | Area% |
|---|---|---|
| 1 | 1.248 | 98.15 |
| 2 | 1.292 | 1.85 |

T6201078

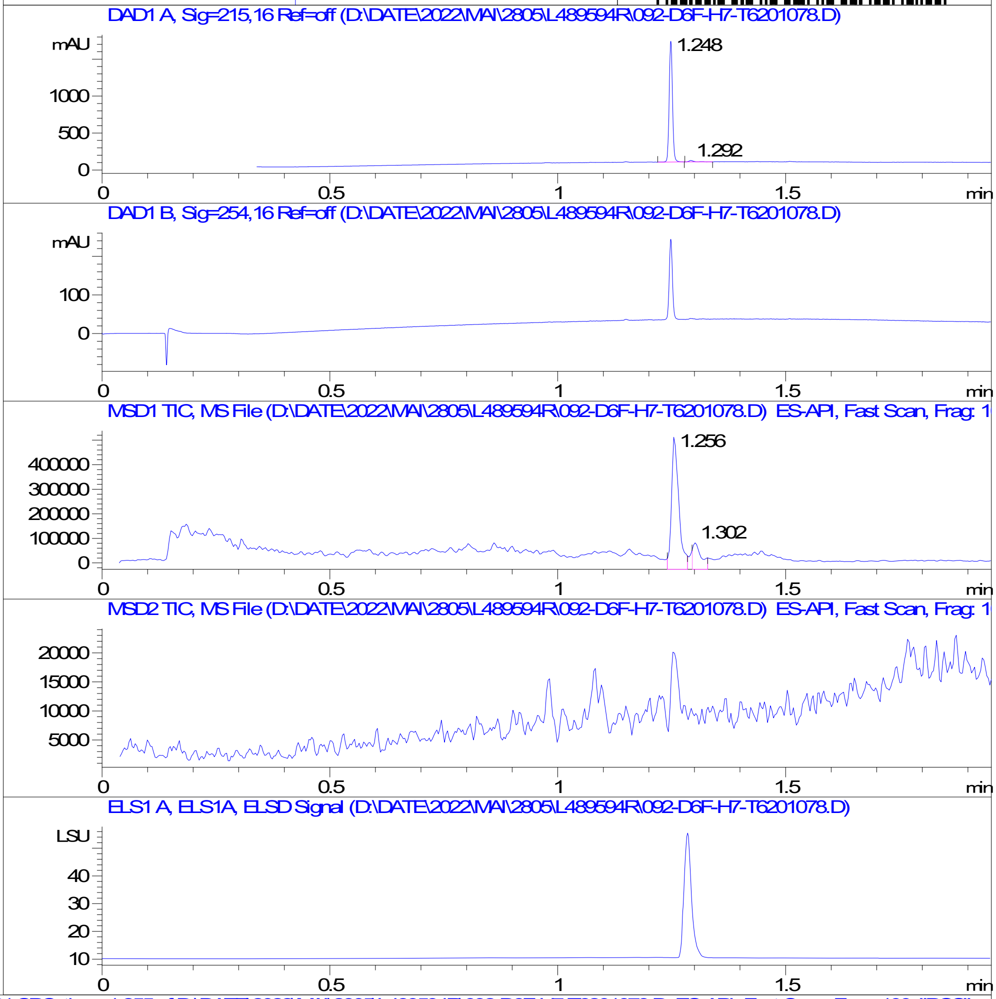


RT 1.256

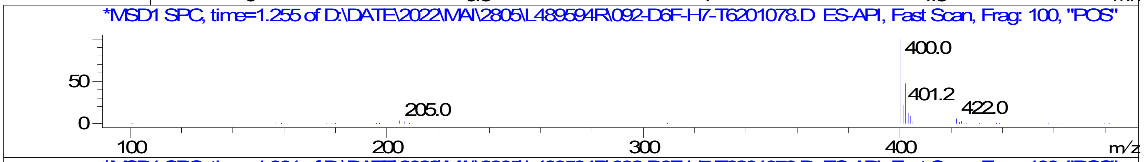


RT 1.302

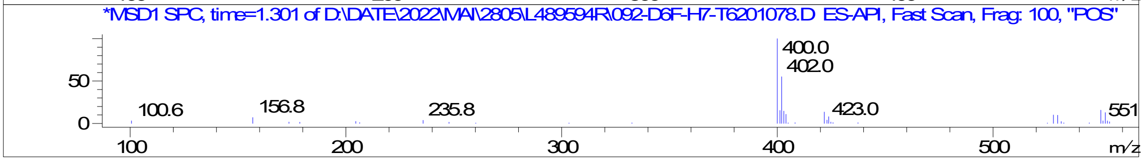


Inj.Date 28-May-22 Z <invalid> Acq. Method C:\Users\ -> ->

Z335721330_

MaxPeak: 100.00%
Ret_Time: 0.718 min

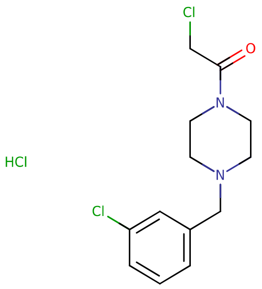


**Mol Wt** 323.65
**Exact Mass** 286.09

| # | Time | Area% |
|---|---|---|
| 1 | 0.718 | 100.00 |

# T6330170

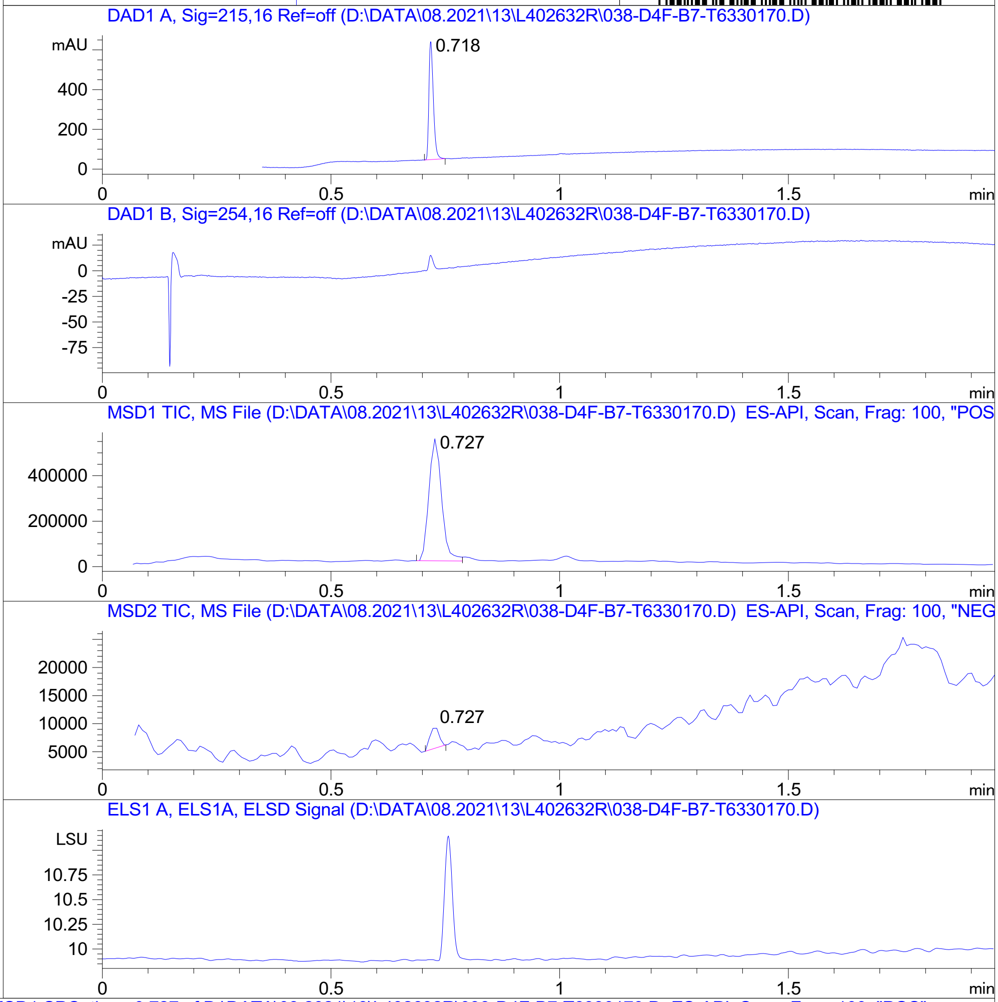


RT 0.727

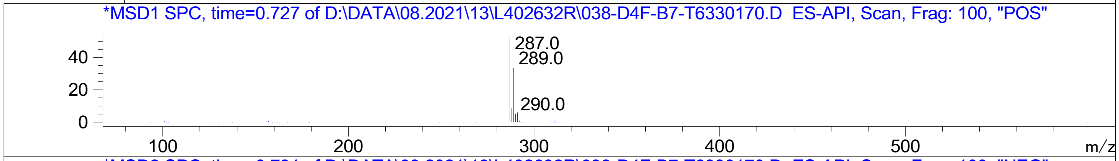


RT 0.727

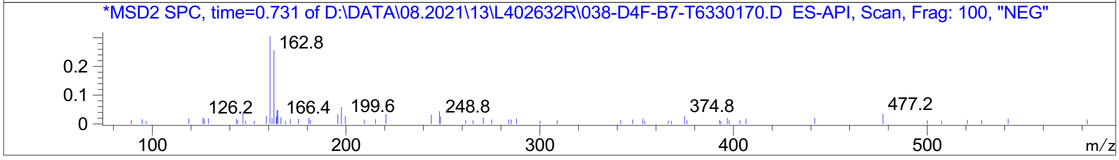


Inj.Date 8/13/2021 CH <invalid> -18- Acq. Method C:\Chem32\-> ->

Z147647726_

MaxPeak: 100.00%
Ret_Time: 1.440 min

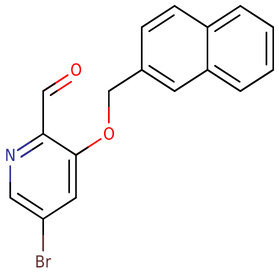


**V374147$4**

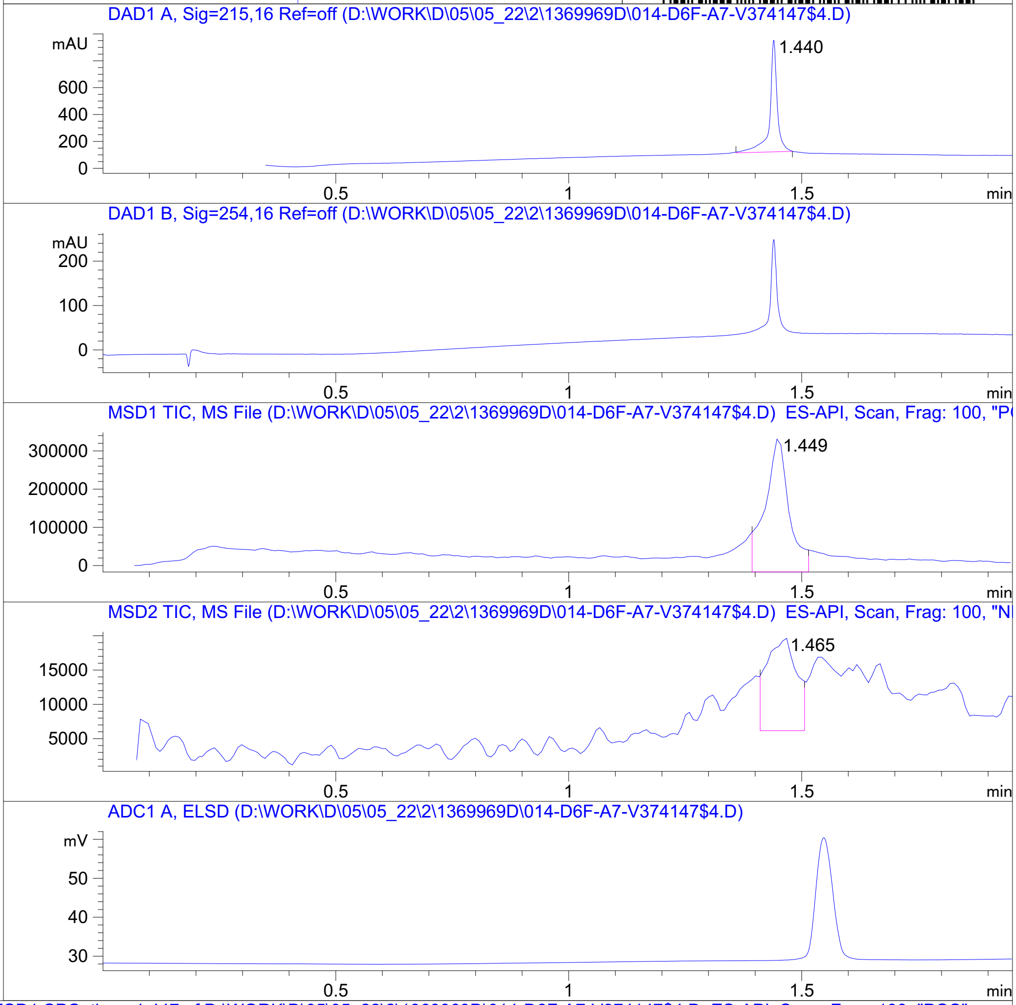


**Mol Wt** **342.19**

**Exact Mass** **341.02**

| # | Time | Area% |
|---|---|---|
| 1 | 1.440 | 100.00 |

RT 1.449

RT 1.465

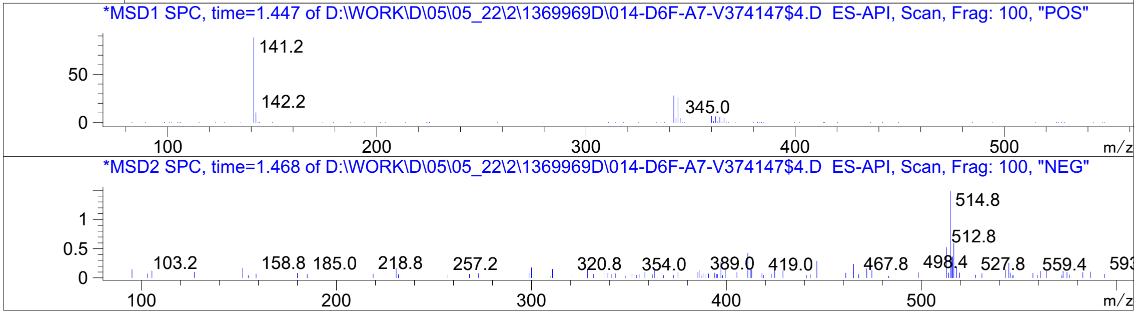




Z4658847356_

MaxPeak: 100.00%
Ret_Time: 1.550 min

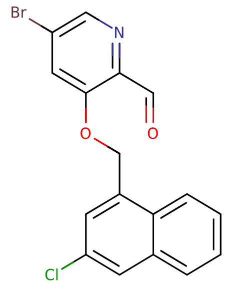


**Mol Wt** 376.63
**Exact Mass** 376.98

| # | Time | Area% |
|---|---|---|
| 1 | 1.550 | 100.00 |

# W557422$R

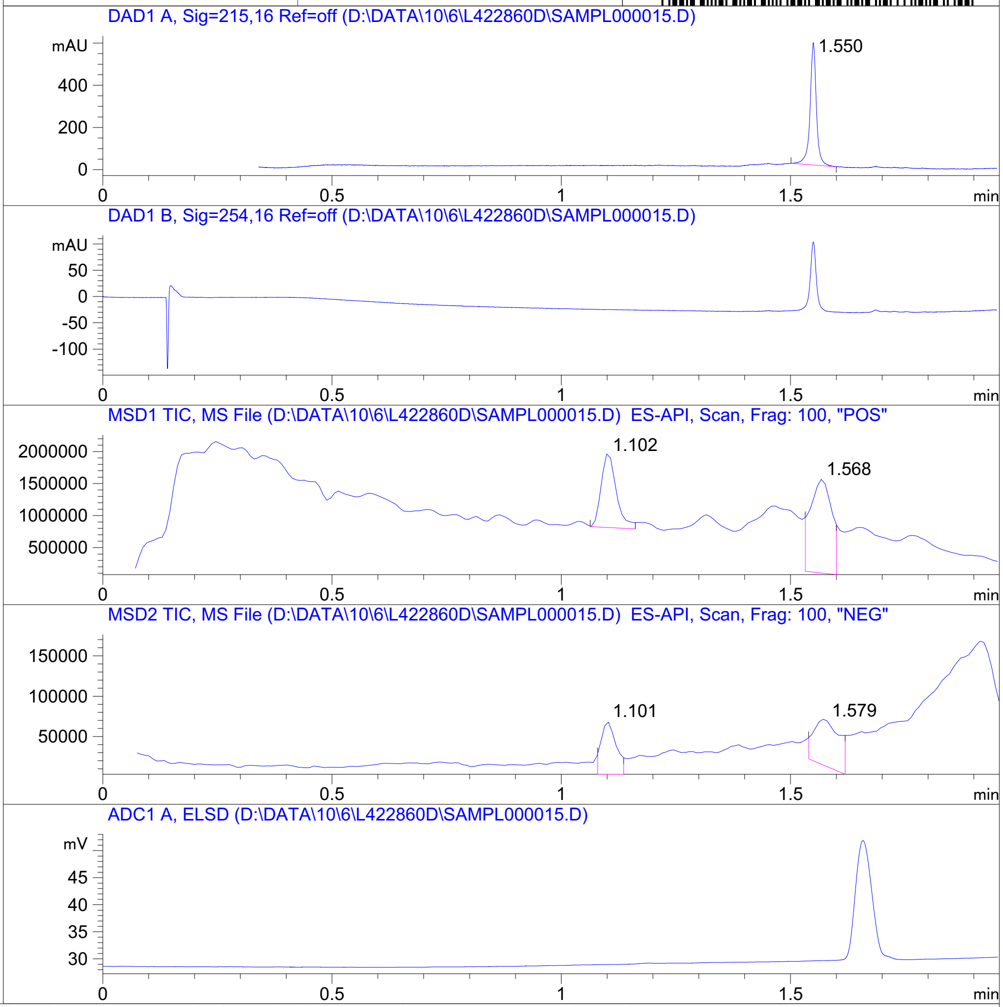


RT 1.102

RT 1.568

RT 1.101

RT 1.579

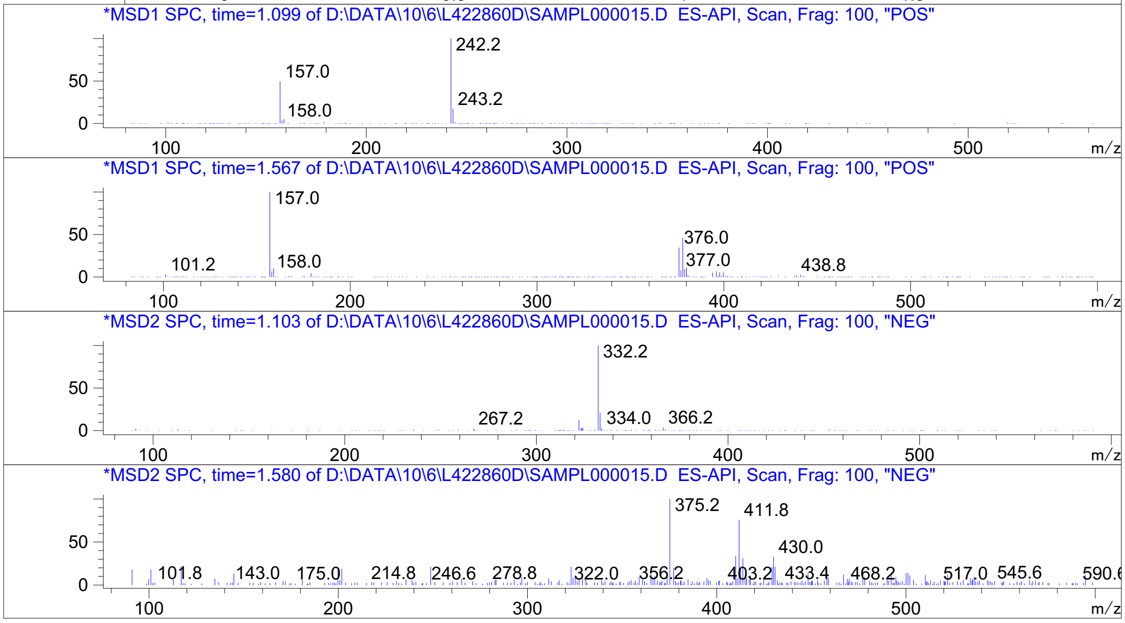


Inj.Date 10/6/2021 LB  Acq. Method C:\CHEM32\-> ->

Z5192779122_

MaxPeak: 80.83%
Ret_Time: 1.043 min

# H3599927

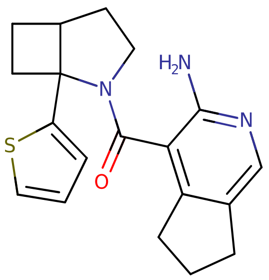


**Mol Wt** 339.46
**Exact Mass** 339.17

| # | Time | Area% |
|---|---|---|
| 1 | 0.632 | 19.17 |
| 2 | 1.043 | 80.83 |

RT 0.647

RT 1.061

RT 0.649

RT 1.060

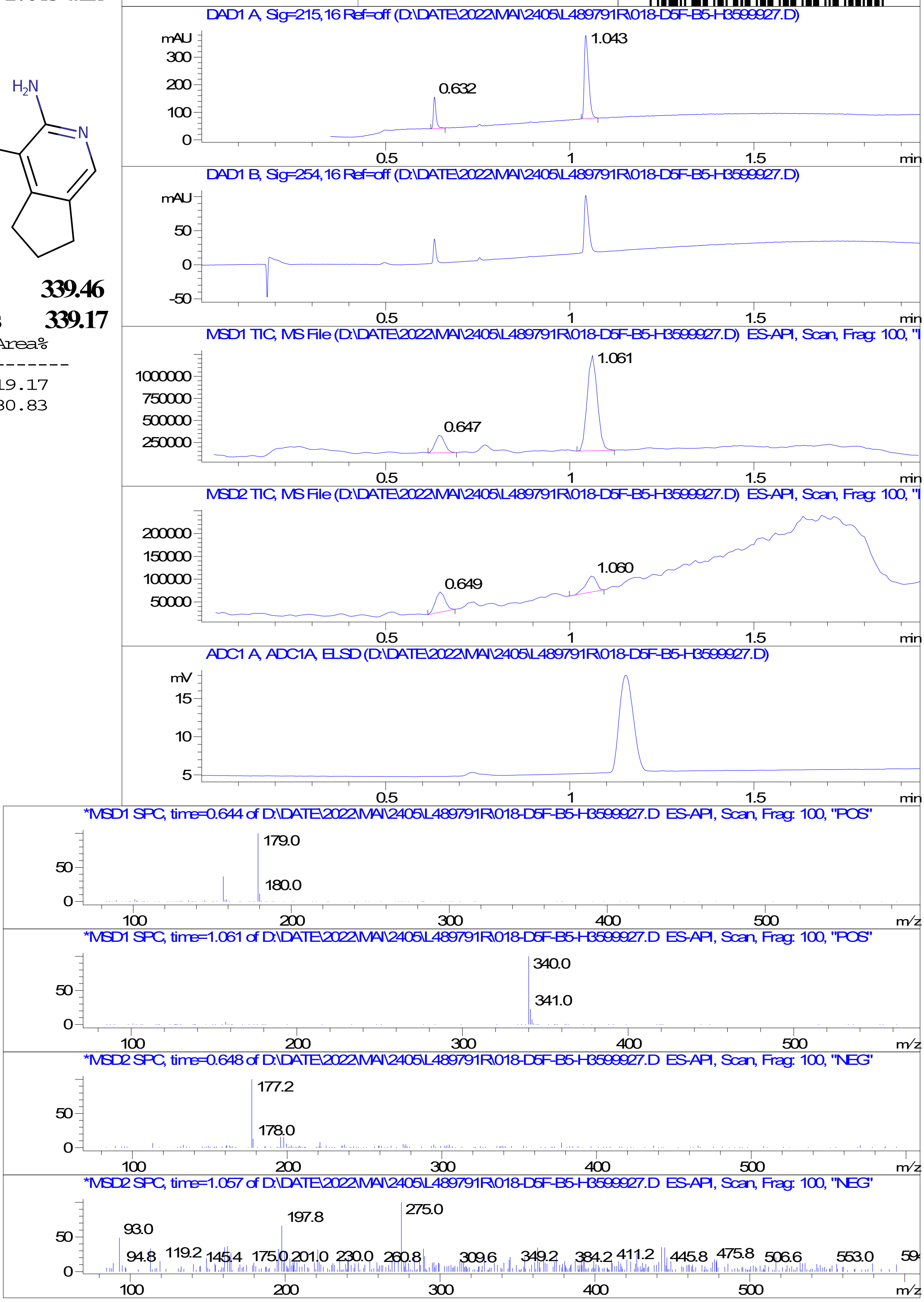


Inj.Date 24-May-22 Z <invalid> -25- Acq. Method C:\Users\ -> ->

Z4124468376_

MaxPeak: 100.00%
Ret_Time: 1.237 min

### GULO

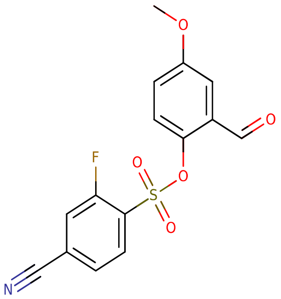

**Mol Wt** 335.31
**Exact Mass** 335.02

| # | Time | Area% |
|---|---|---|
| 1 | 1.237 | 100.00 |

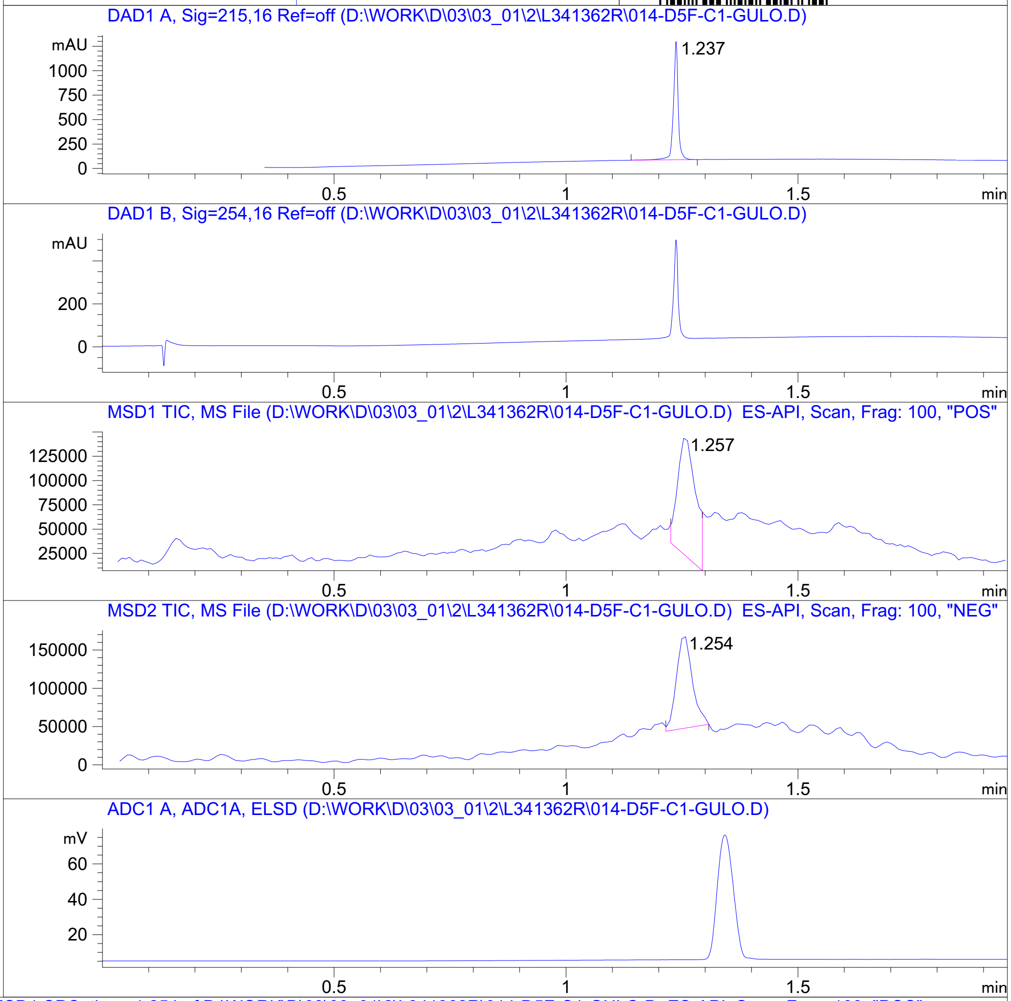


RT 1.257

RT 1.254

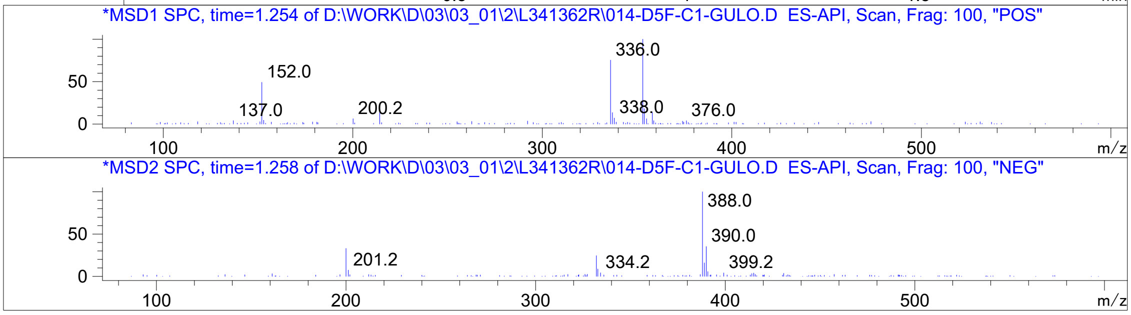


Inj.Date 3/1/2021 K  Acq. Method C:\Users\ -> ->

Z1247383311_

MaxPeak: 100.00%
Ret_Time: 1.468 min

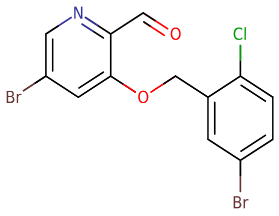


**Mol Wt** 405.47

**Exact Mass** 404.87

| # | Time | Area% |
|---|---|---|
| 1 | 1.468 | 100.00 |

V233166$4

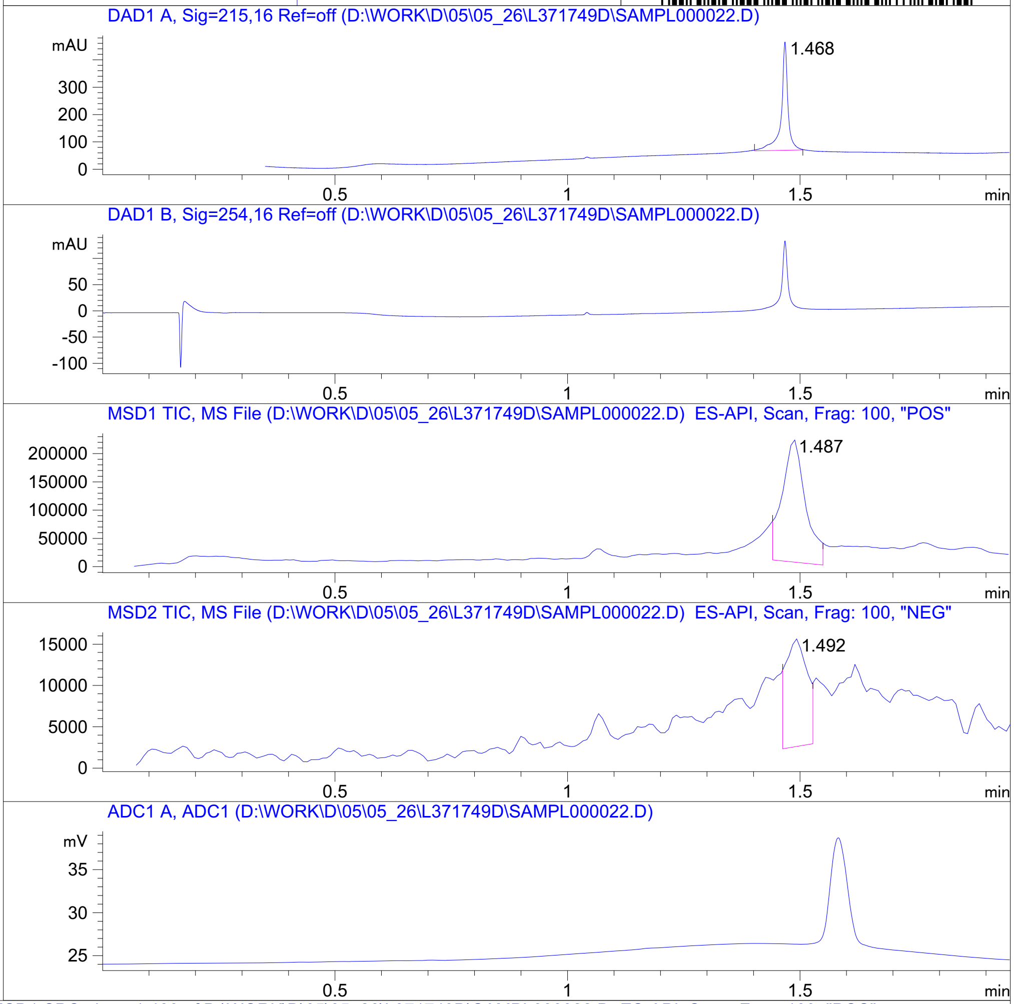


RT 1.487

RT 1.492

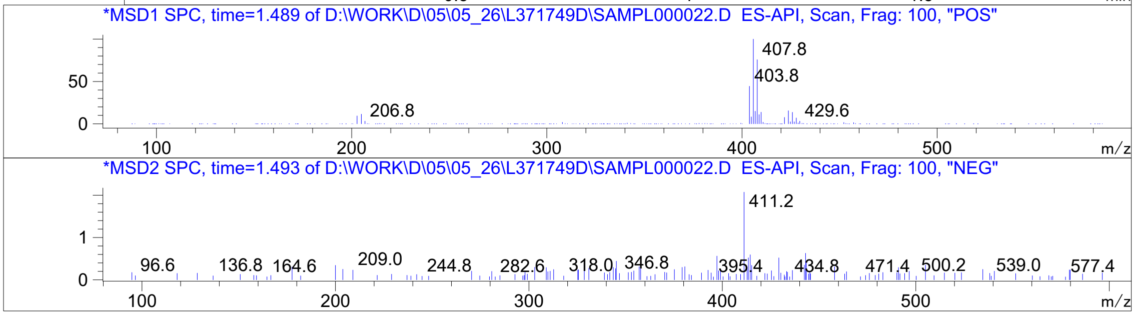




Z5009000155_

MaxPeak: 98.93%
Ret_Time: 1.247 min

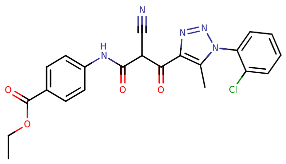


**Mol Wt** 451.86

**Exact Mass** 451.11

| # | Time | Area% |
|---|---|---|
| 1 | 1.030 | 1.07 |
| 2 | 1.247 | 98.93 |

S568083$4

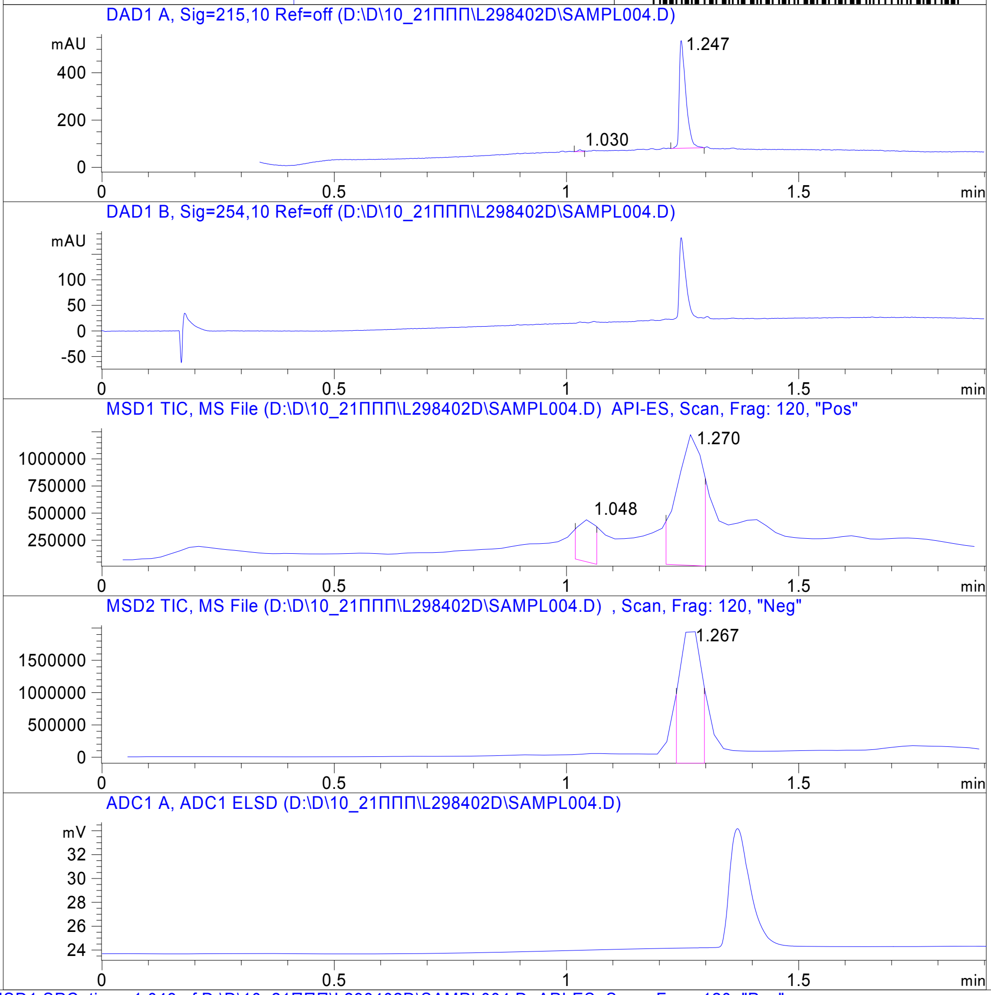


RT 1.048

RT 1.270

RT 1.267

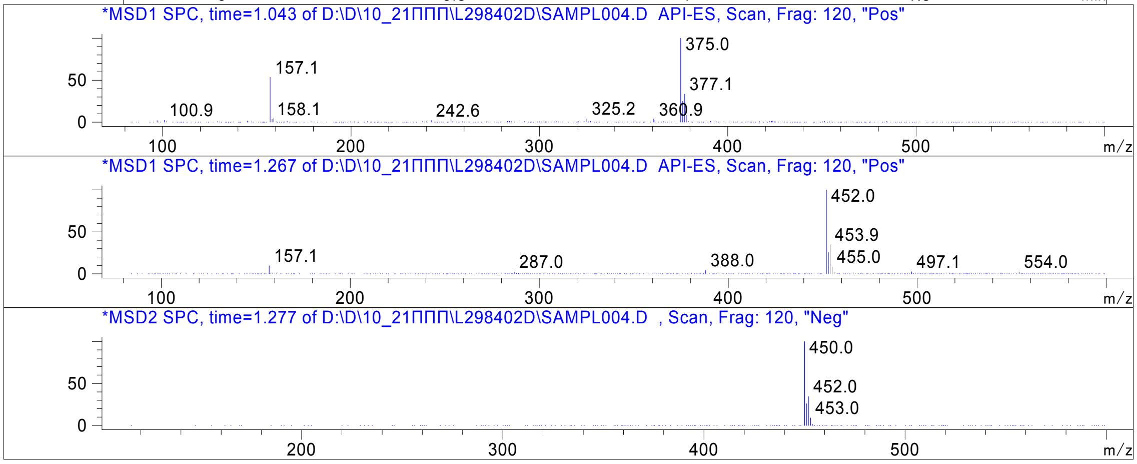


Inj.Date 10/22/2020 N -VL- Acq. Method C:\HPCHEM\-> ->

Z1123372000_

MaxPeak: 91.40%
Ret_Time: 1.332 min

S866357$B

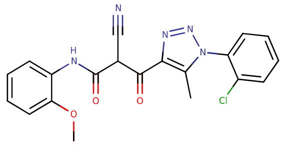

**Mol Wt** 409.83

**Exact Mass** 409.1

| # | Time | Area% |
|---|---|---|
| 1 | 0.913 | 6.77 |
| 2 | 1.221 | 1.83 |
| 3 | 1.332 | 91.40 |

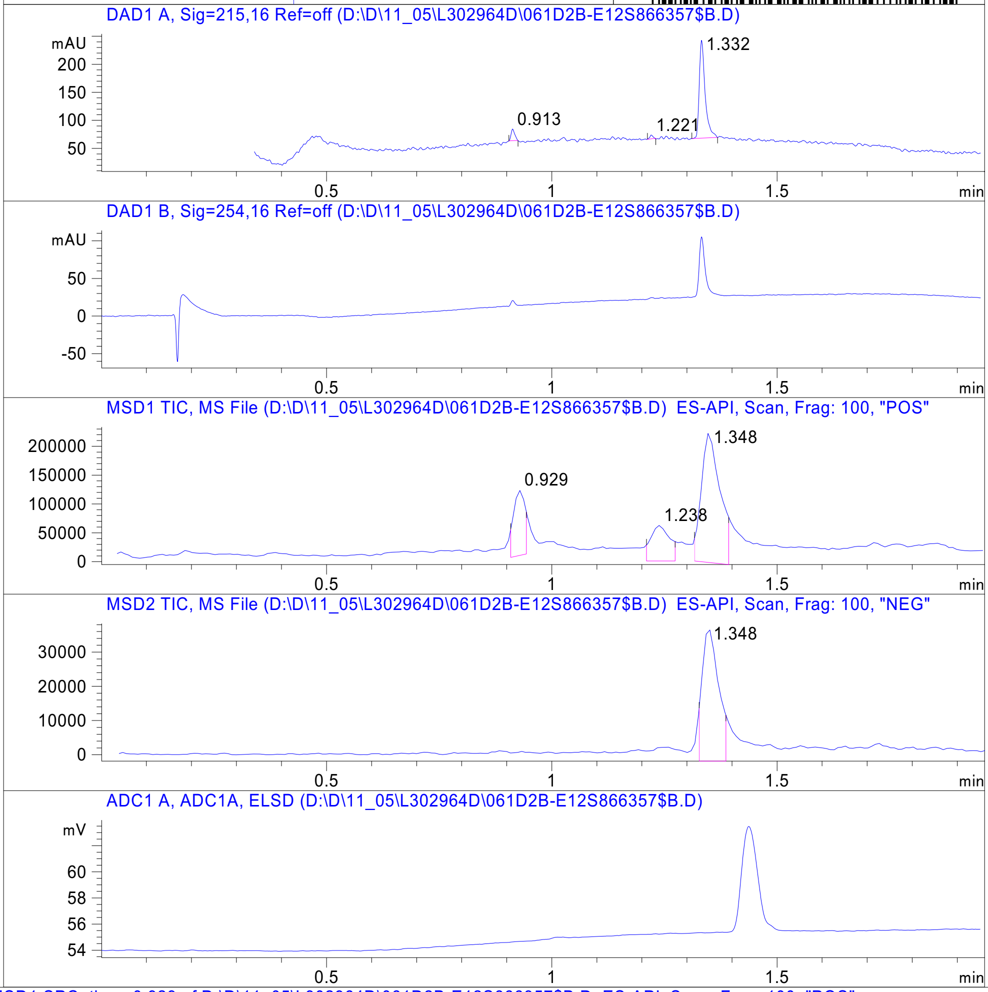


RT 0.929

RT 1.238

RT 1.348

RT 1.348

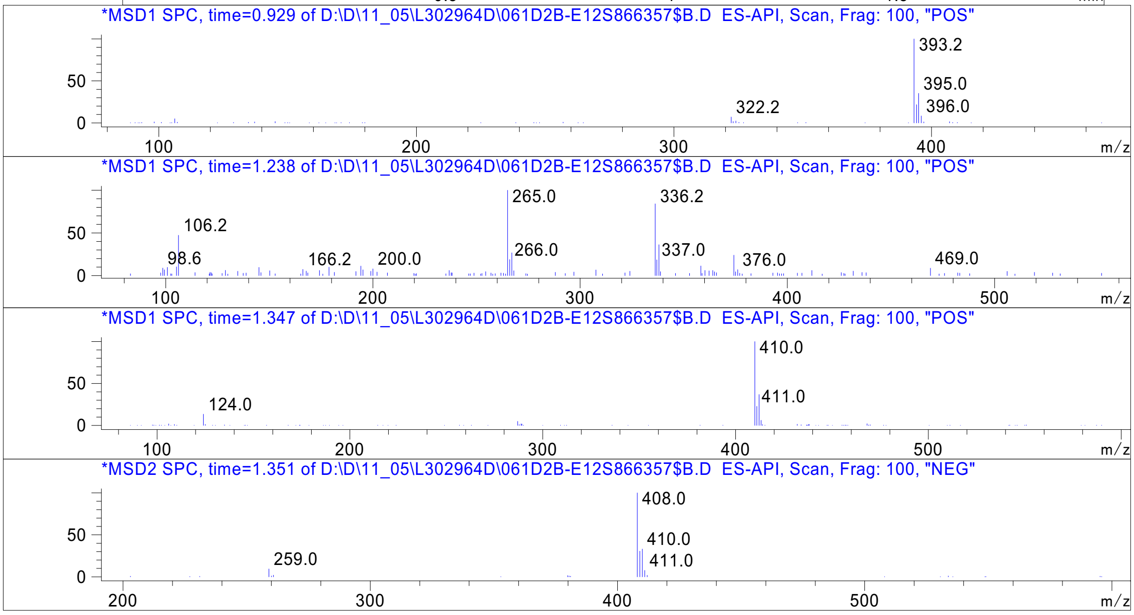




Z1123371668_

MaxPeak: 100.00%
Ret_Time: 1.795 min

# S568071$17

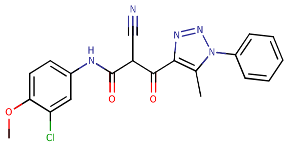


**Mol Wt** **409.83**

**Exact Mass** **409.1**

| # | Time | Area% |
|---|---|---|
| 1 | 1.795 | 100.00 |

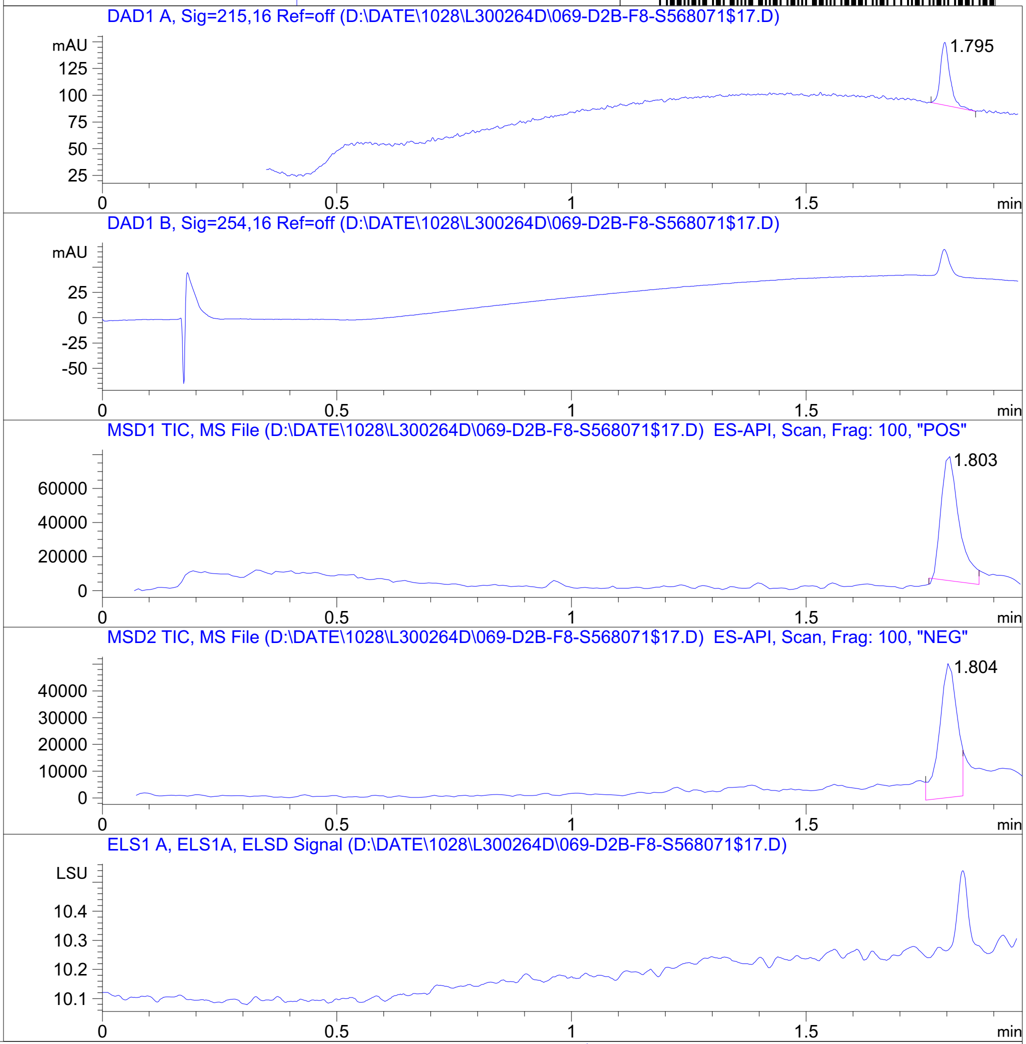


RT 1.803

RT 1.804

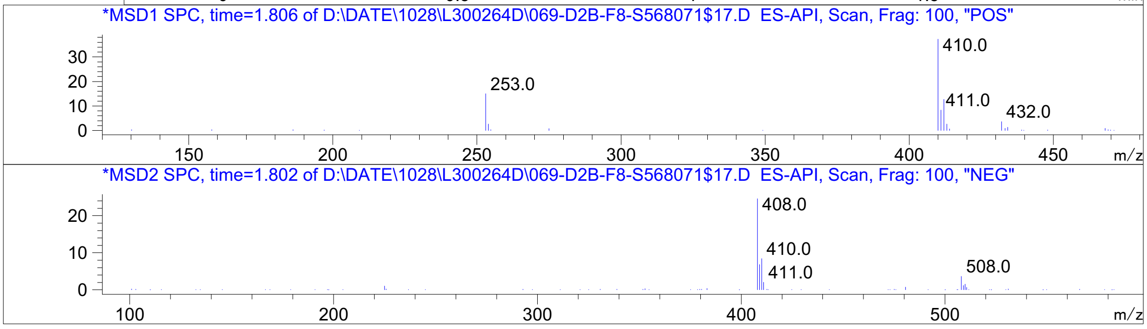


Inj.Date 10/28/2020 O -17- Acq. Method C:\Chem32\-> ->

Z762212060_

# S568078$C

MaxPeak: 95.28%
Ret_Time: 1.360 min

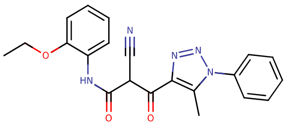

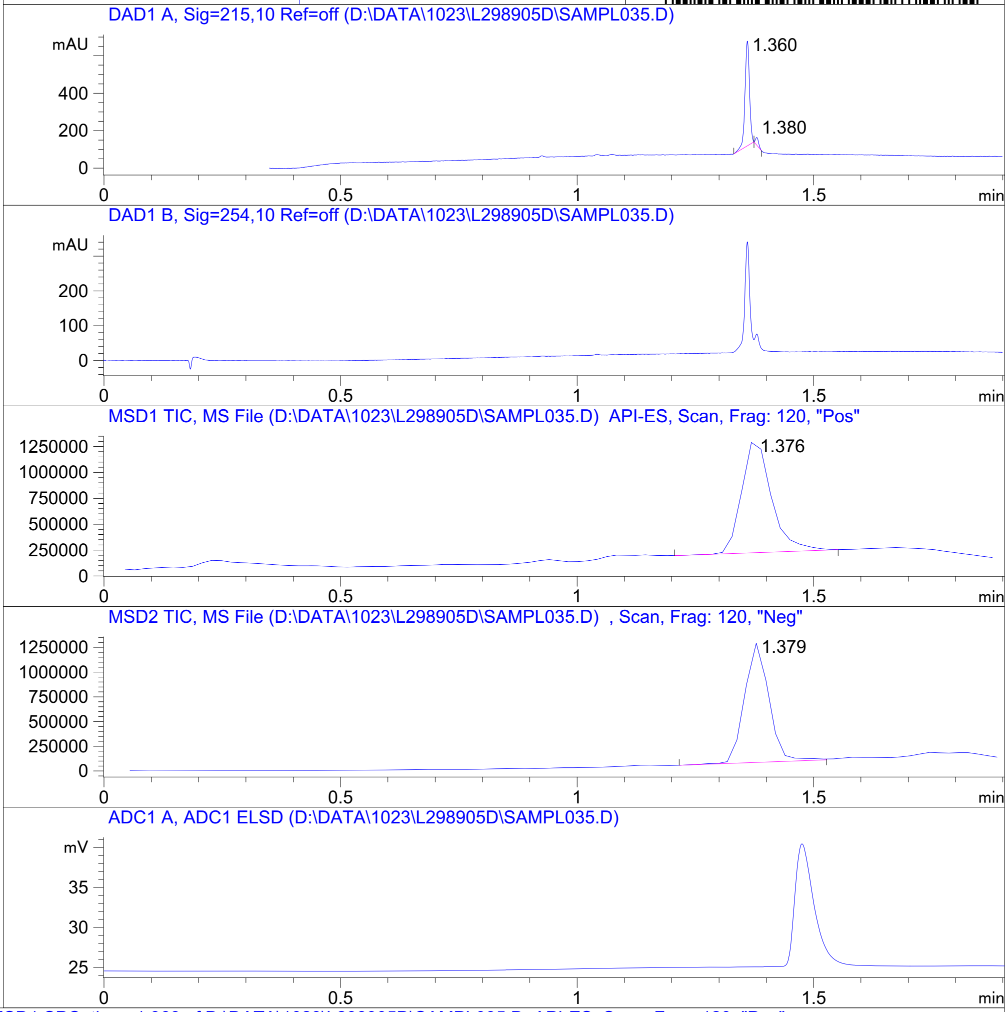


**Mol Wt** 389.41

**Exact Mass** 389.16

| # | Time | Area% |
|---|---|---|
| 1 | 1.360 | 95.28 |
| 2 | 1.380 | 4.72 |

RT 1.376

RT 1.379

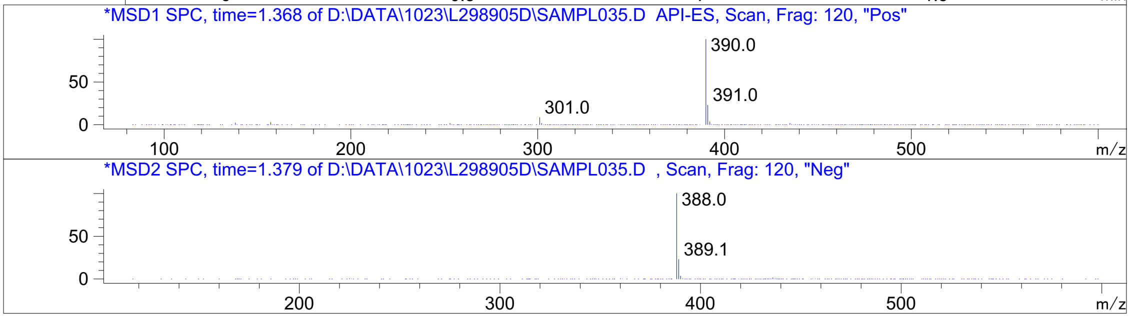


Inj.Date 10/23/2020 OA -VL- Acq. Method C:\HPCHEM\-> ->

Z762210282_

MaxPeak: 100.00%
Ret_Time: 1.205 min

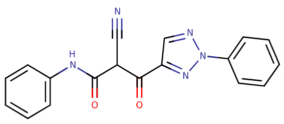

**Mol Wt** 331.33
**Exact Mass** 331.11

| # | Time | Area% |
|---|---|---|
| 1 | 1.205 | 100.00 |

# S993141$15

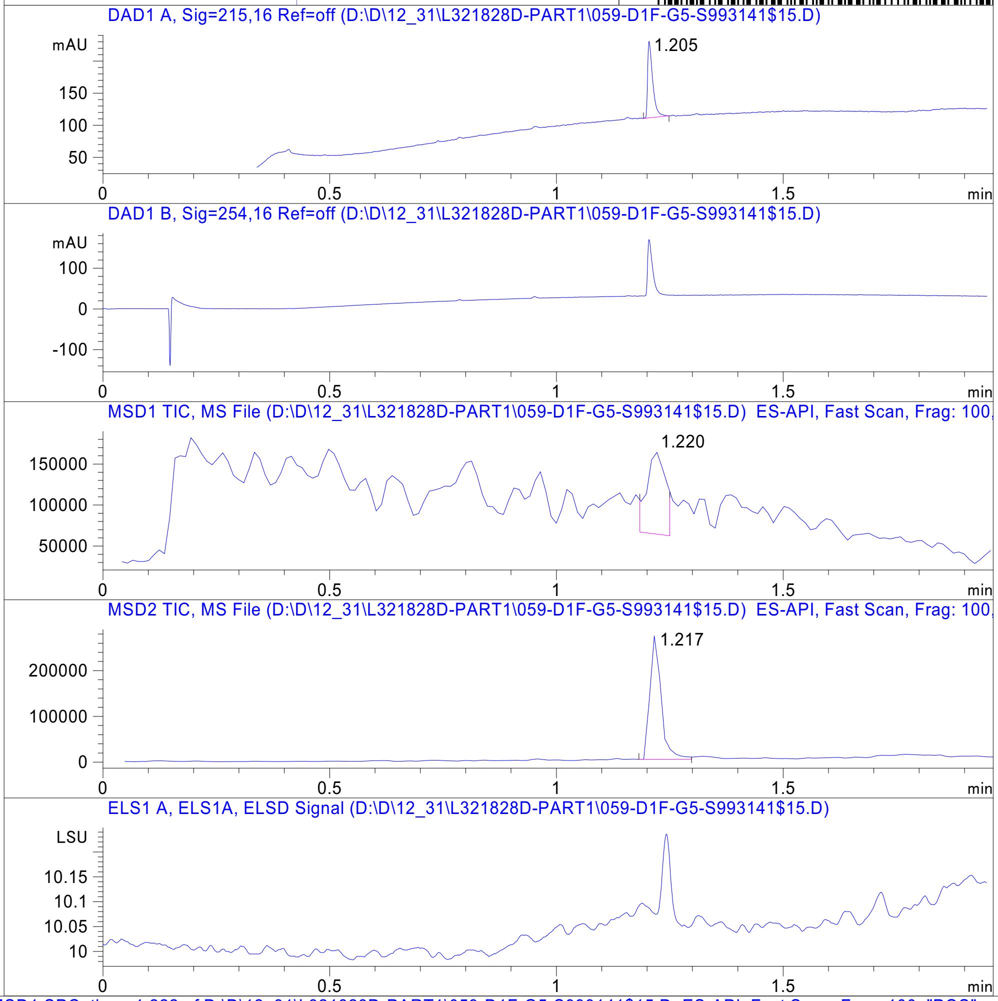


RT 1.220

RT 1.217

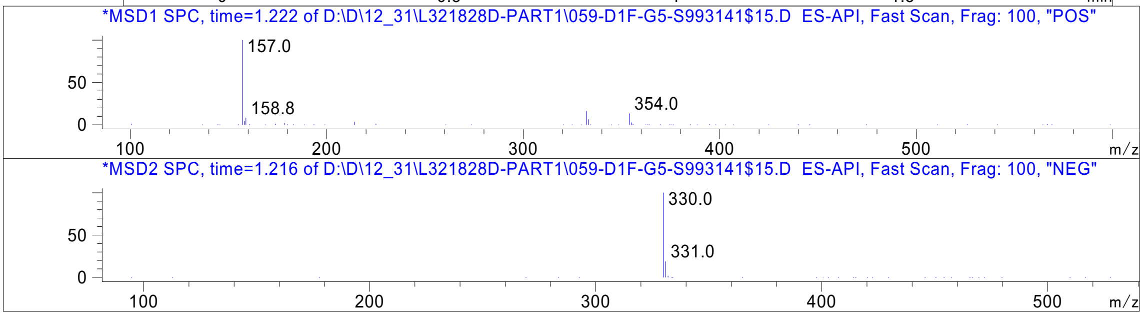


Inj.Date 12/30/2020 N Acq. Method C:\Users\ -> ->

Z638927484_

MaxPeak: 100.00%
Ret_Time: 2.073 min

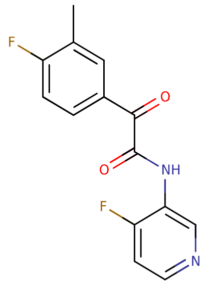


**Mol Wt** 276.24
**Exact Mass** 276.08

| # | Time | Area% |
|---|---|---|
| 1 | 2.073 | 100.00 |

# H3659984

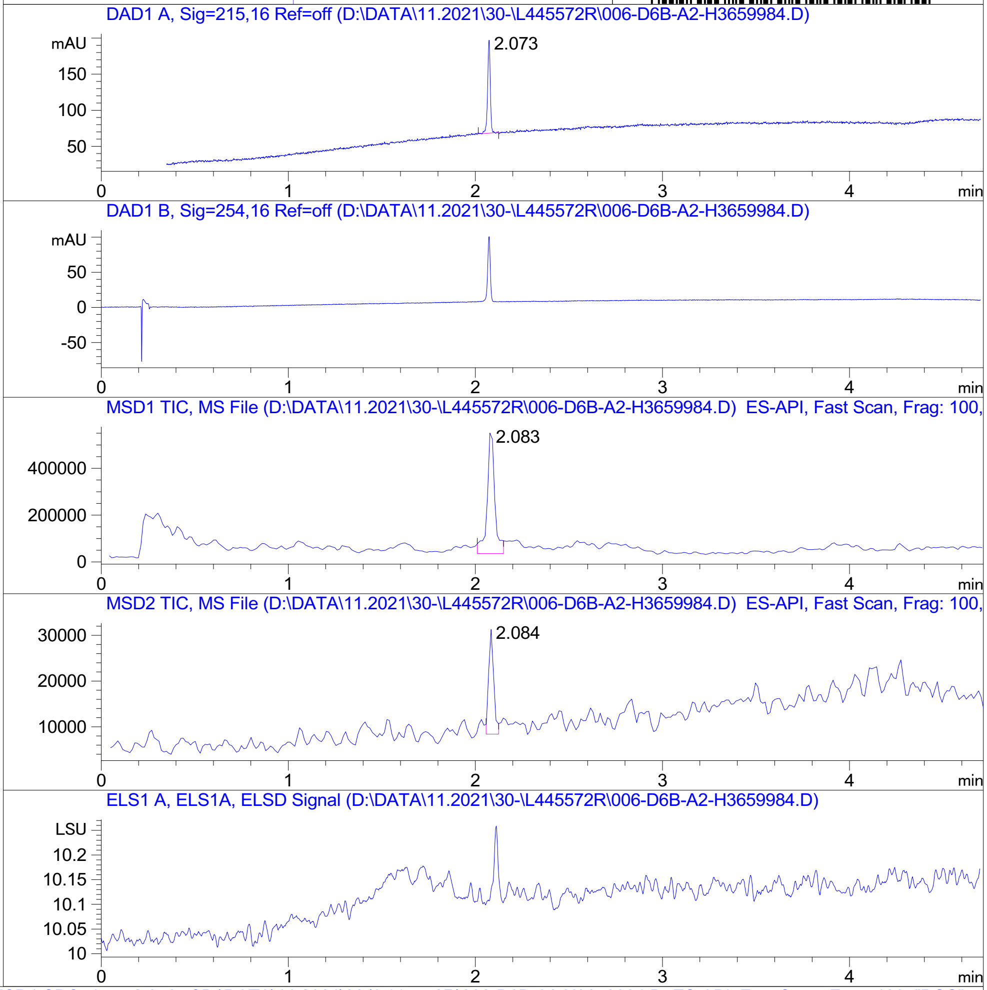


RT 2.083

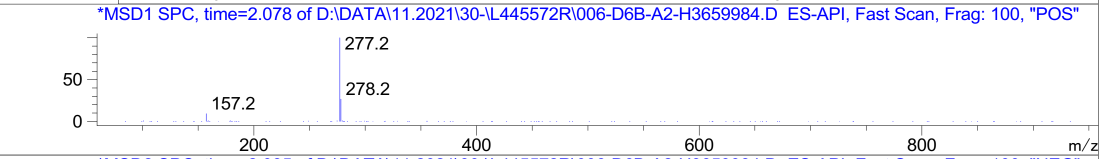


RT 2.084

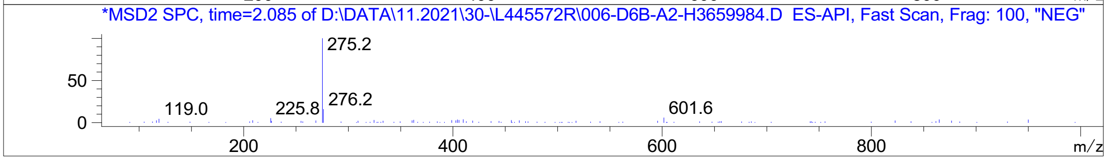


Inj.Date 11/30/2021 CH <invalid> Acq. Method C:\Users\ -> ->

Z5420738300_

# W179756$55

MaxPeak:  98.62%
Ret_Time: 1.333 min

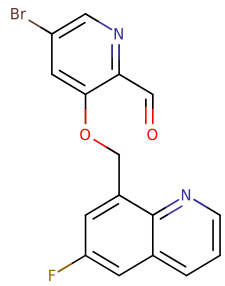


**Mol Wt** 361.16

**Exact Mass** 360

| # | Time | Area% |
|---|---|---|
| 1 | 1.197 | 1.38 |
| 2 | 1.333 | 98.62 |

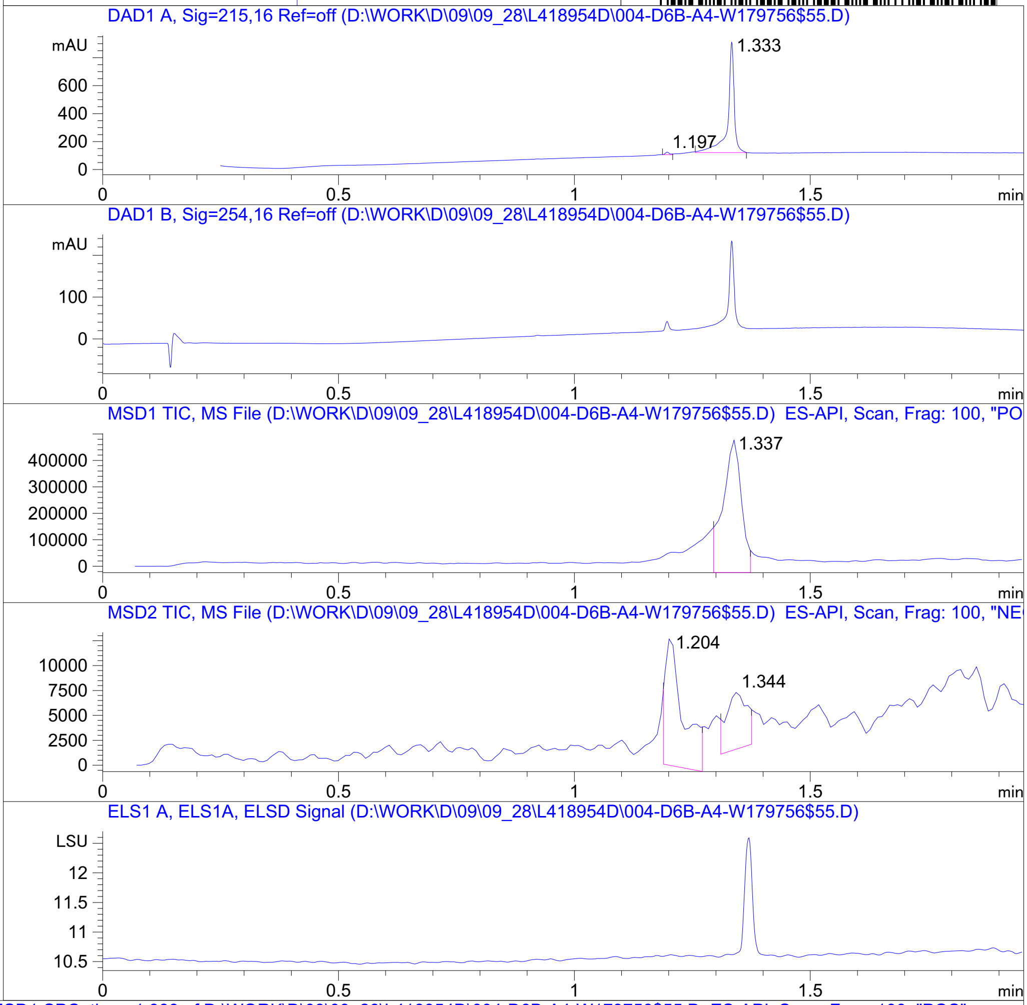


RT 1.337

RT 1.204

RT 1.344

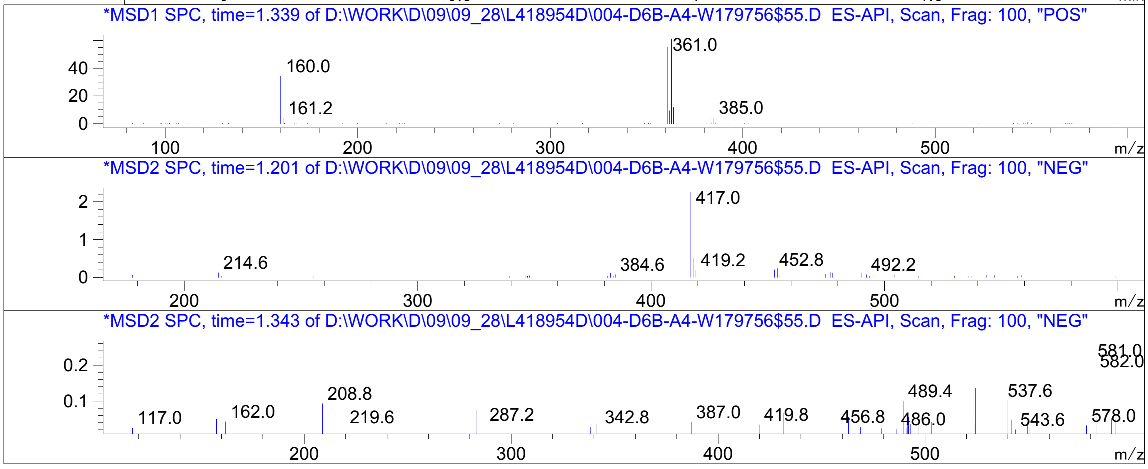


Inj.Date 9/28/2021 K <invalid> -15- Acq. Method C:\Chem32\-> ->

Z5192779144_

Z5192779028_

Library Search Report

```
Data Path : D:\MassHunter\GCMS\1\data\09_22\
Data File : R2726874.D
Acq On    : 22 Sep 2021  15:10
Operator  :
Sample    : R2726874
Misc      : CH3CN
ALS Vial  : 77   Sample Multiplier: 1

Search Libraries:   C:\Database\EMPTY.L               Minimum Quality:   0

Unknown Spectrum:   Apex
Integration Events: ChemStation Integrator - autoint1.e
```

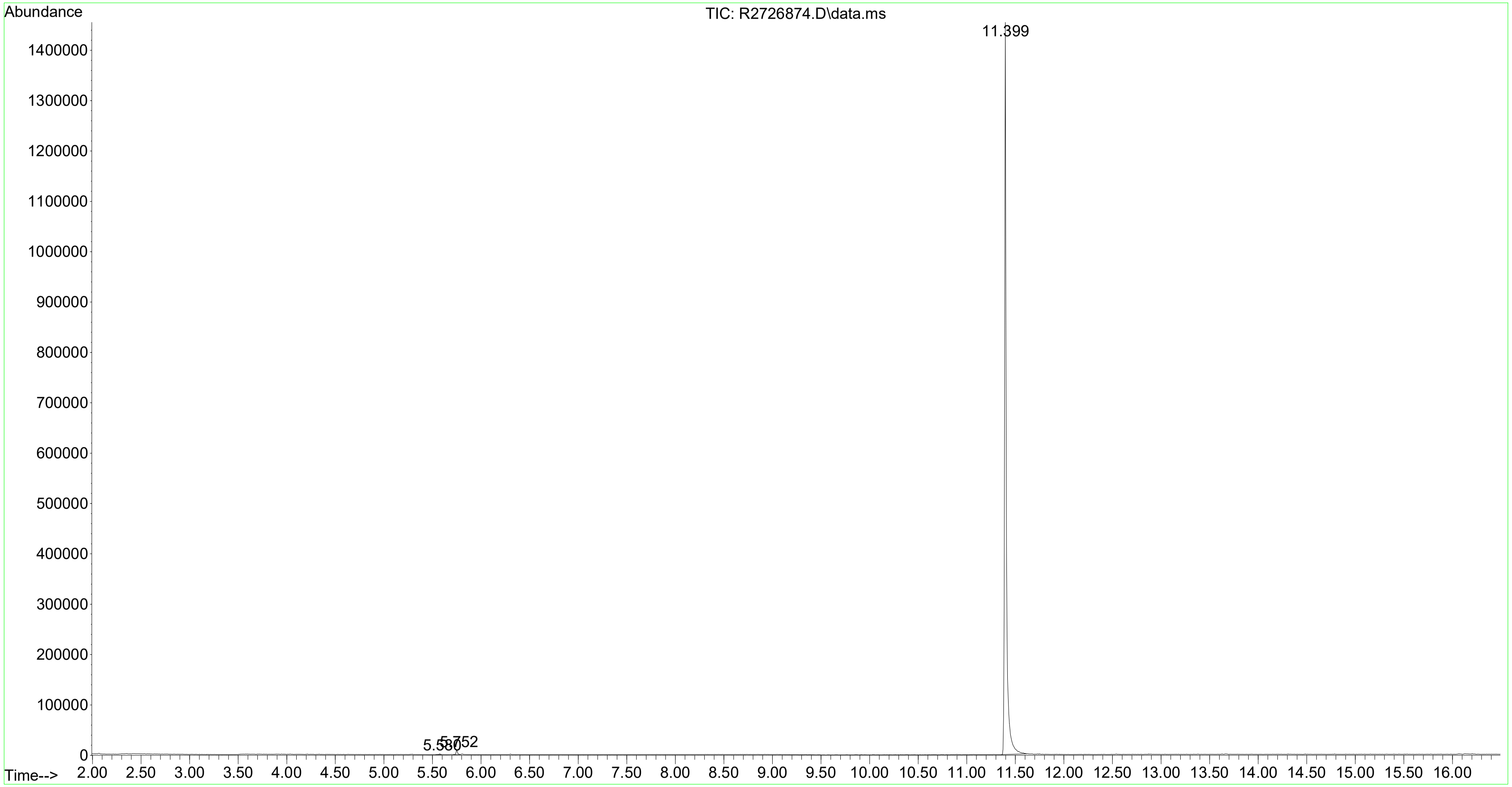

MaxPeak: 100.00%
Ret_Time: 1.568 min

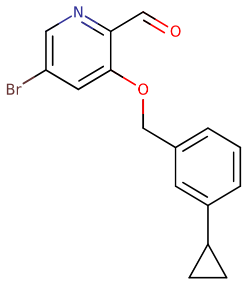

**Mol Wt** 332.19

**Exact Mass** 331.04

| # | Time | Area% |
|---|---|---|
| 1 | 1.568 | 100.00 |

V211811$A

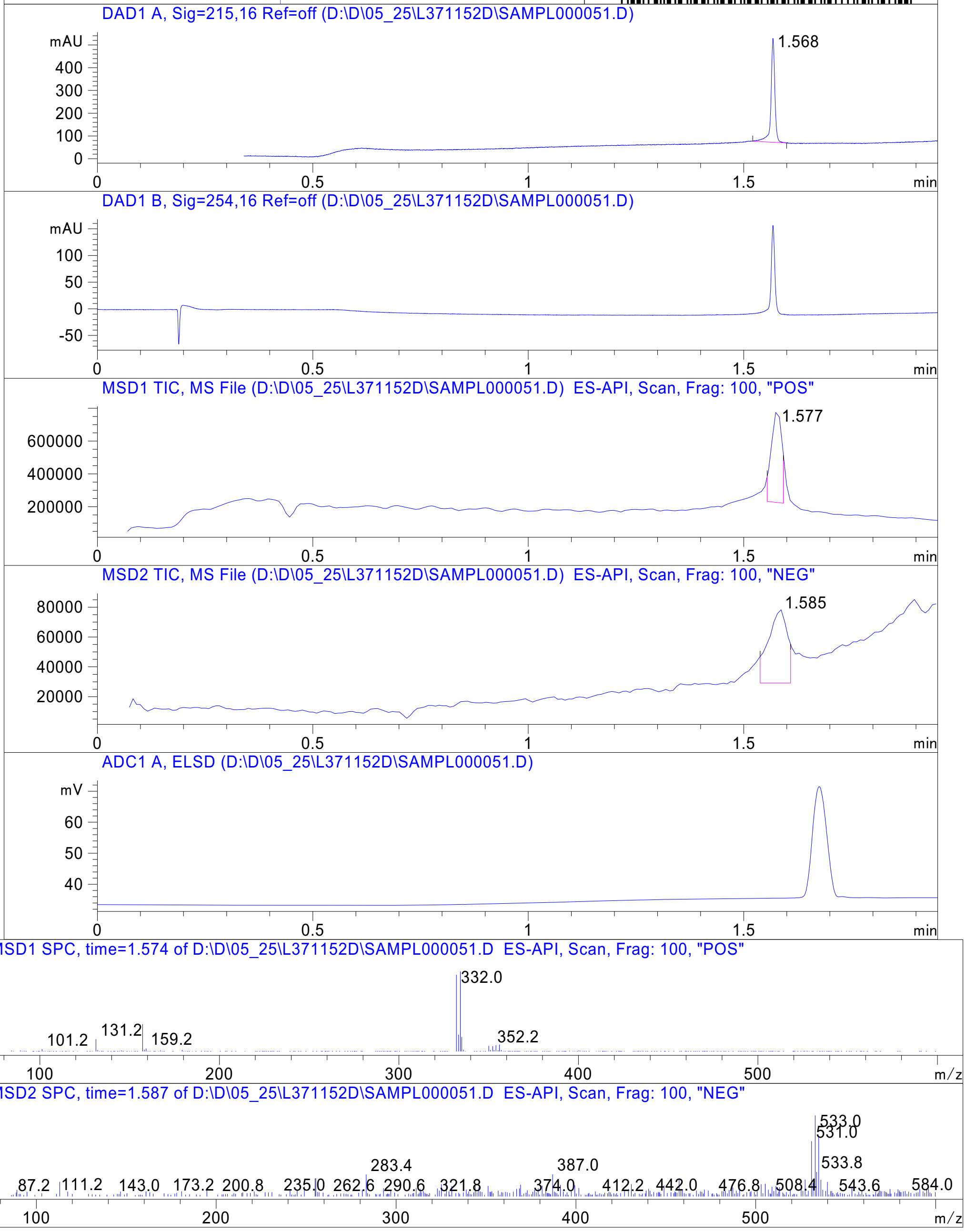




Z4989203264_

MaxPeak: 100.00%
Ret_Time: 1.615 min

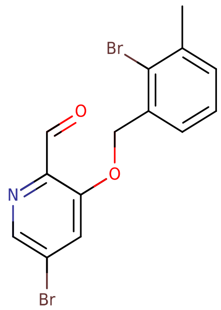


**Mol Wt** 385.05

**Exact Mass** 384.93

| # | Time | Area% |
|---|---|---|
| 1 | 1.615 | 100.00 |

V211809$3

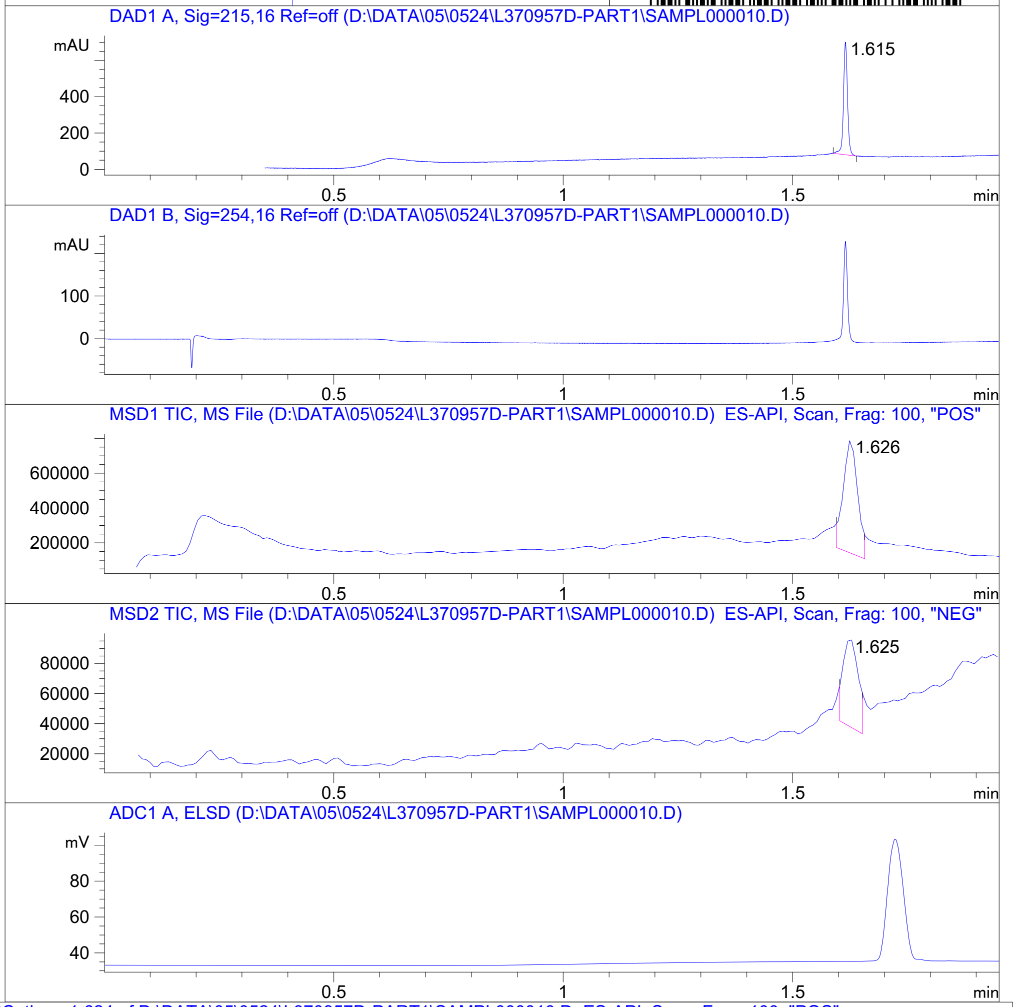


RT 1.626

RT 1.625

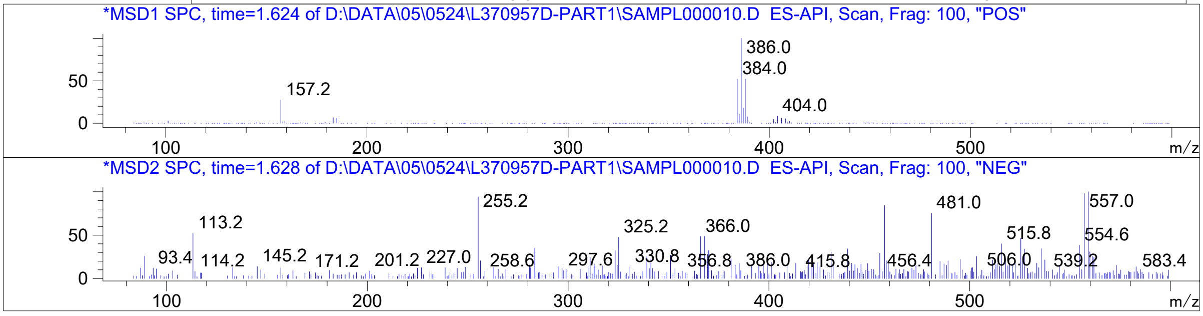




Z4989203259_

MaxPeak: 100.00%
Ret_Time: 1.295 min

V296342$4

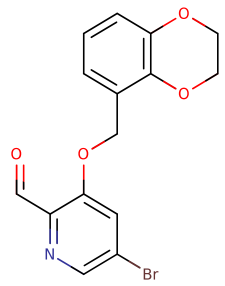

**Mol Wt** 350.16
**Exact Mass** 349

| # | Time | Area% |
|---|---|---|
| 1 | 1.295 | 100.00 |

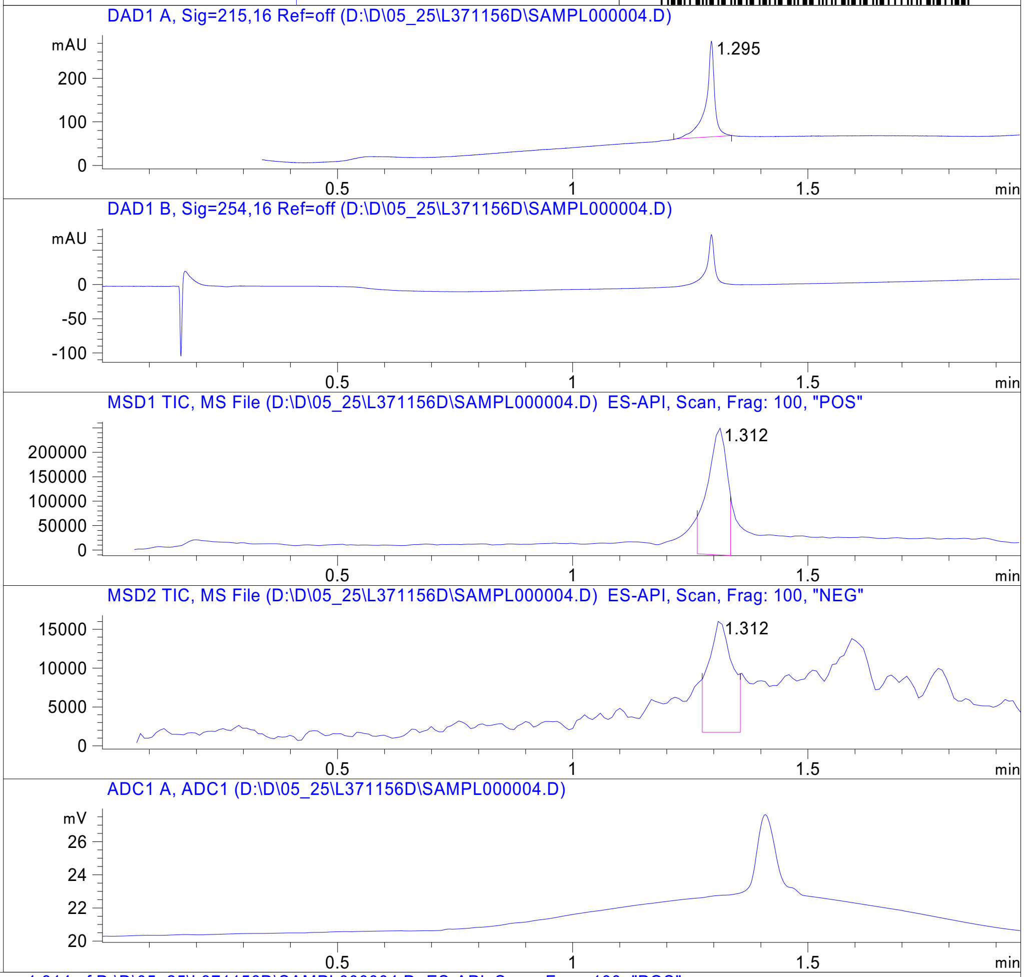


RT
1.312

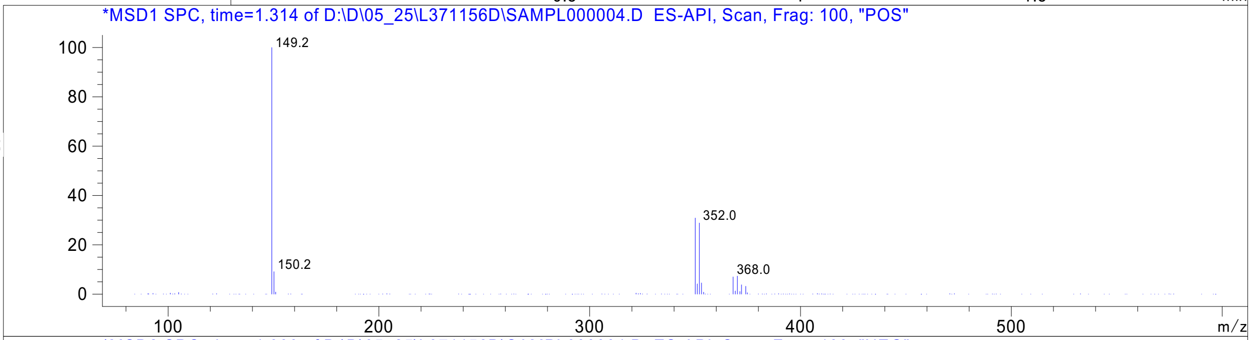


RT
1.312

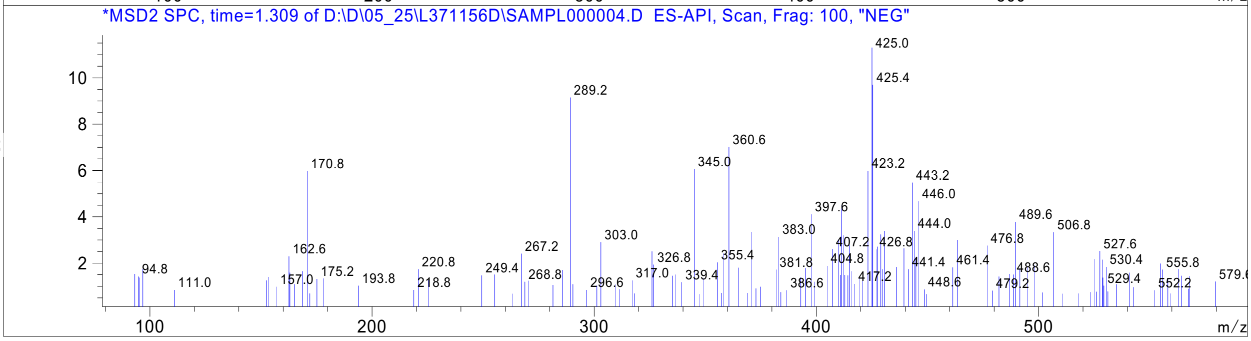


Inj.Date 5/23/2021 N - 4 - Acq. Method C:\CHEM32\->->

Z4989203255_

MaxPeak: 99.47%
Ret_Time: 2.993 min

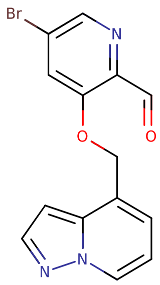


**V296341$6**

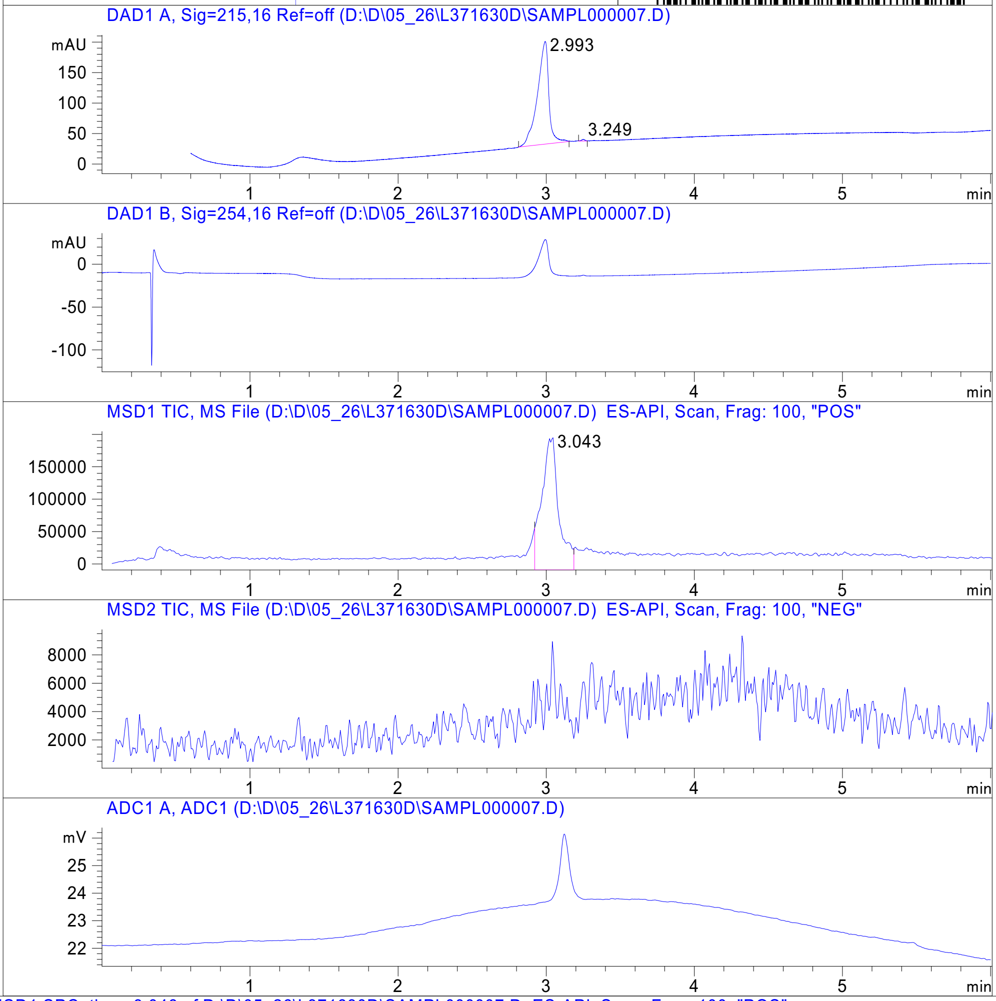


**Mol Wt** 332.15

**Exact Mass** 331

| # | Time | Area% |
|---|---|---|
| 1 | 2.993 | 99.47 |
| 2 | 3.249 | 0.53 |

RT 3.043

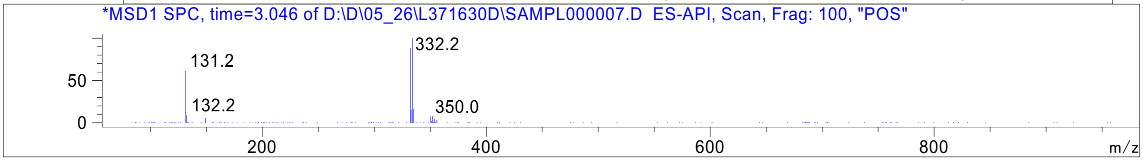




Z4989203254_

Z4954569231_

Library Search Report

```
Data Path : D:\MassHunter\GCMS\1\data\09_09\
Data File : R2714707.D
Acq On    : 09 Sep 2021  17:49
Operator  :
Sample    : R2714707
Misc      : CH3CN
ALS Vial  : 126   Sample Multiplier: 1

Search Libraries:   C:\Database\EMPTY.L              Minimum Quality:   0

Unknown Spectrum:   Apex
Integration Events: ChemStation Integrator - autoint1.e
```

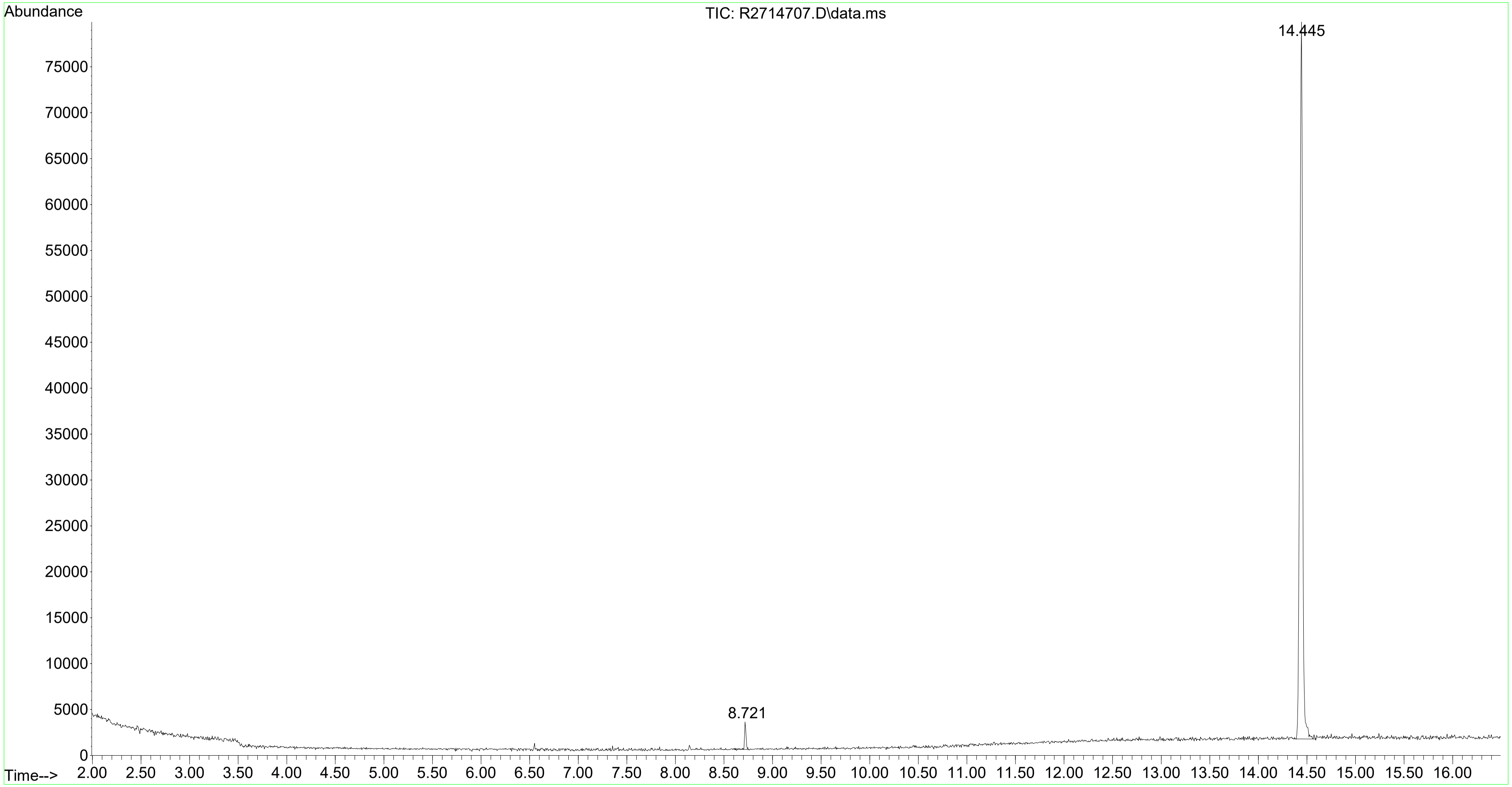

MaxPeak: 91.30%
Ret_Time: 0.860 min

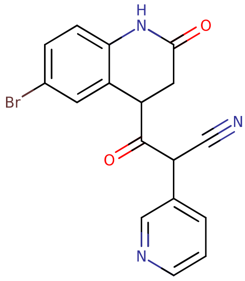


**Mol Wt** 370.2

**Exact Mass** 369.02

| # | Time | Area% |
|---|---|---|
| 1 | 0.765 | 3.46 |
| 2 | 0.834 | 5.24 |
| 3 | 0.860 | 91.30 |

# R2539922

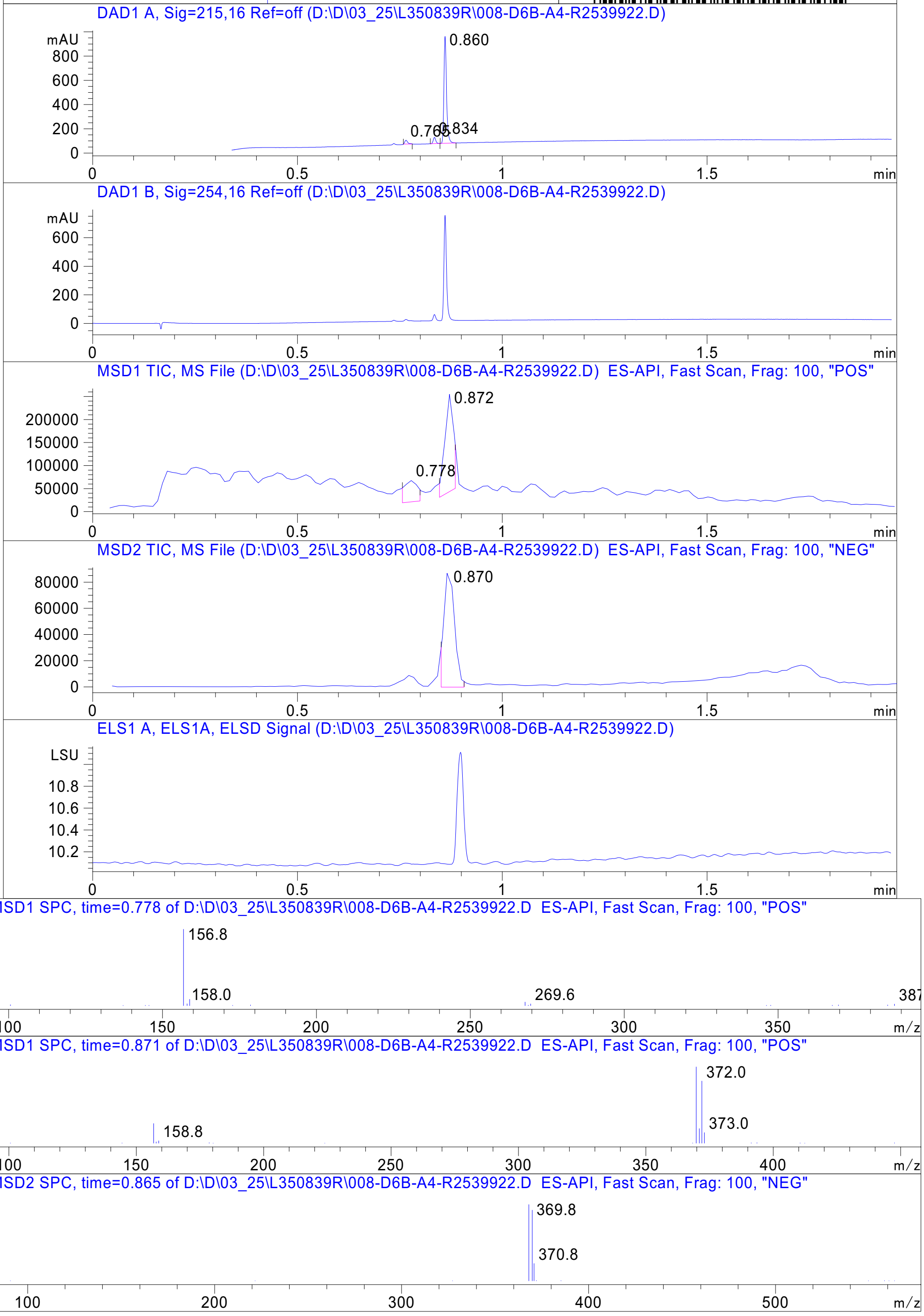


Inj.Date 3/25/2021 N Acq. Method C:\Users\ -> ->

Z4934489646_

MaxPeak: 100.00%
Ret_Time: 1.211 min

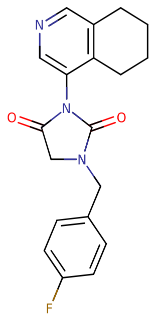


**Mol Wt** 339.36
**Exact Mass** 339.16

| # | Time | Area% |
|---|---|---|
| 1 | 1.211 | 100.00 |

U853195$2

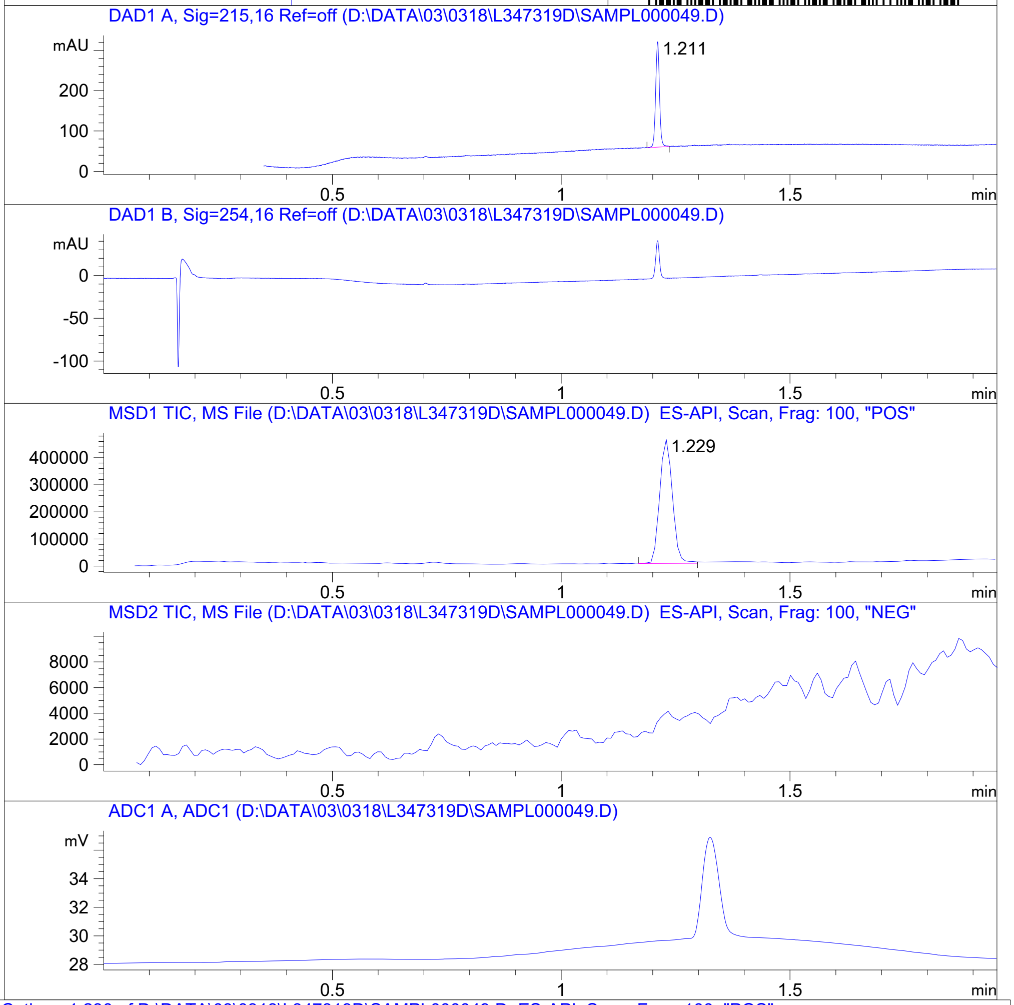


RT 1.229

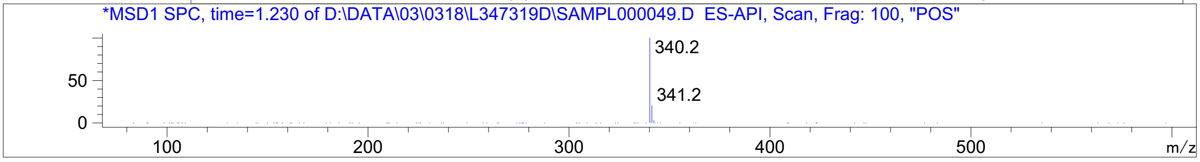




Z4929616328_

MaxPeak:  97.64%
Ret_Time: 1.055 min

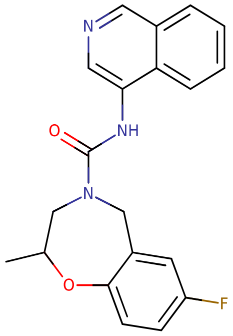


# V218888$1

**Mol Wt** 351.37

**Exact Mass** 351.16

| # | Time | Area% |
|---|---|---|
| 1 | 1.033 | 2.36 |
| 2 | 1.055 | 97.64 |

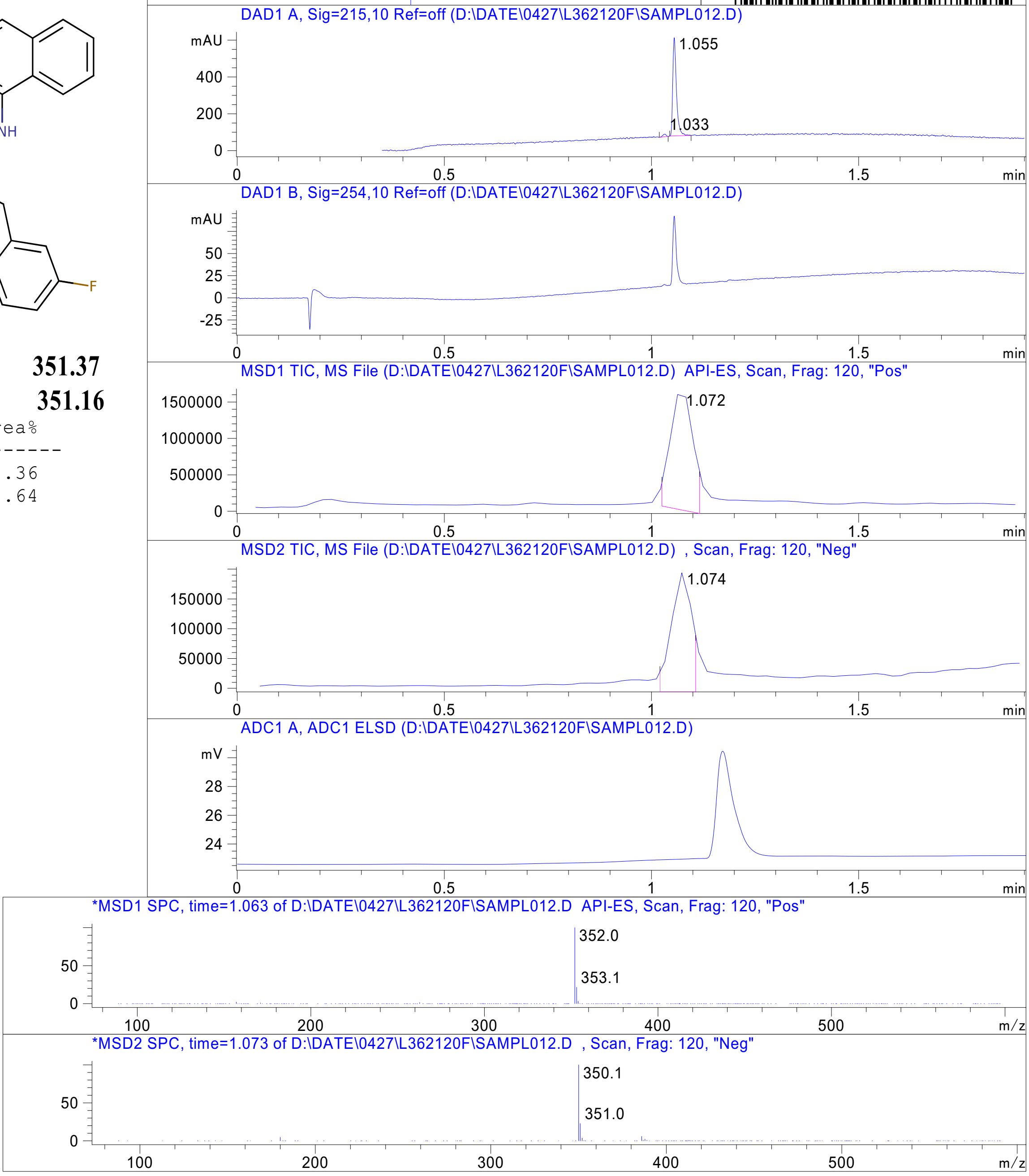


RT 1.072

RT 1.074

Inj.Date 4/27/2021    E    -VL-    Acq. Method C:\HPCHEM\->    ->

Z4924562400_

MaxPeak: 90.44%
Ret_Time: 1.579 min

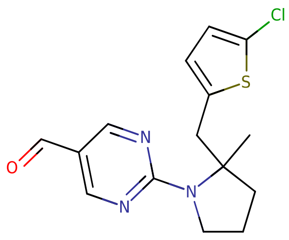

**Mol Wt 321.82**
**Exact Mass 321.09**

| # | Time | Area% |
|---|---|---|
| 1 | 1.311 | 9.56 |
| 2 | 1.579 | 90.44 |

Q763569$4

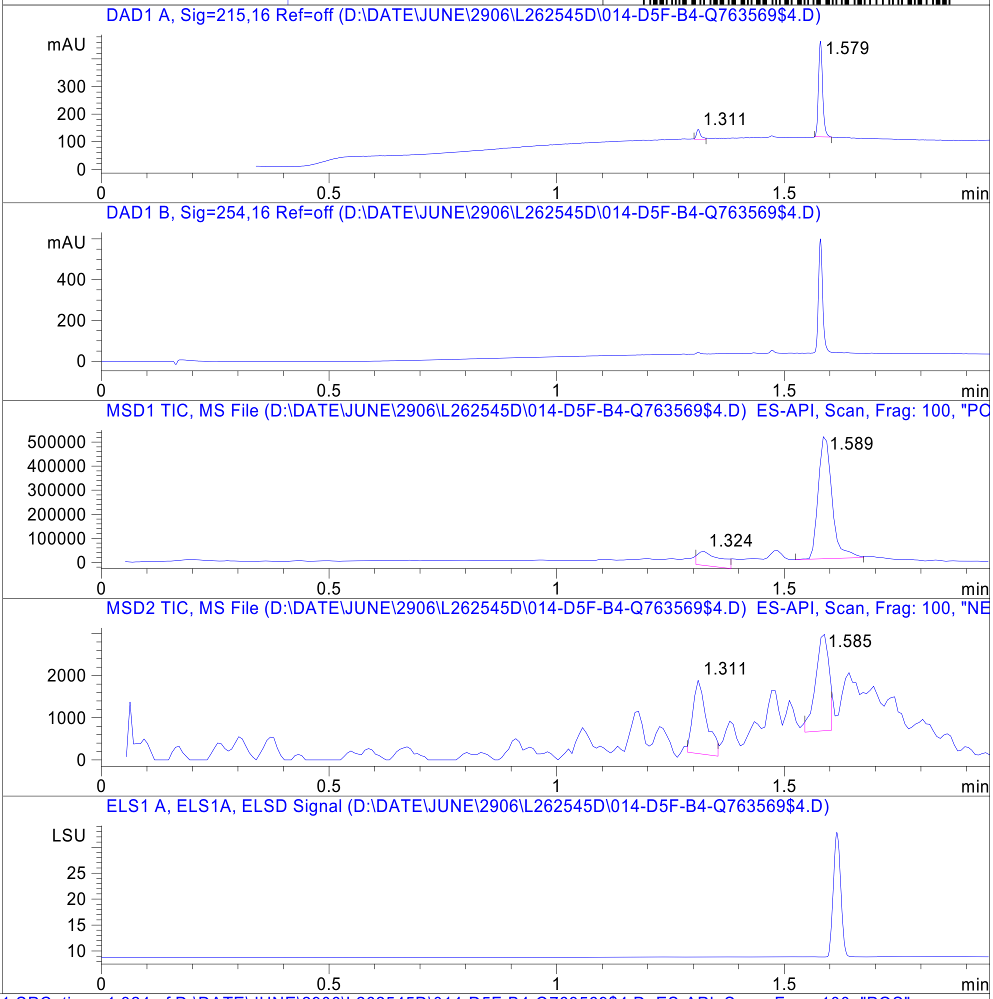


RT 1.324

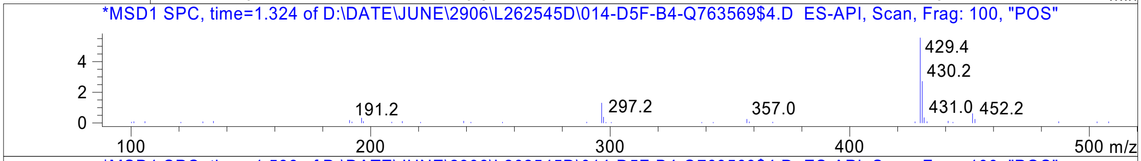


RT 1.589

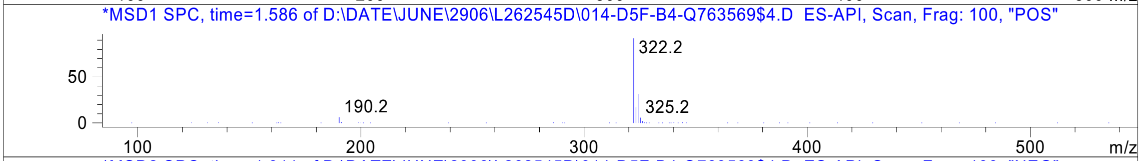


RT 1.311

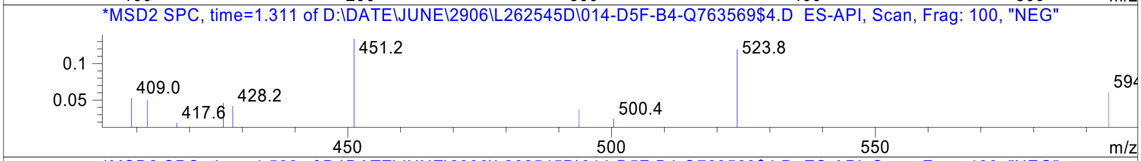


RT 1.585

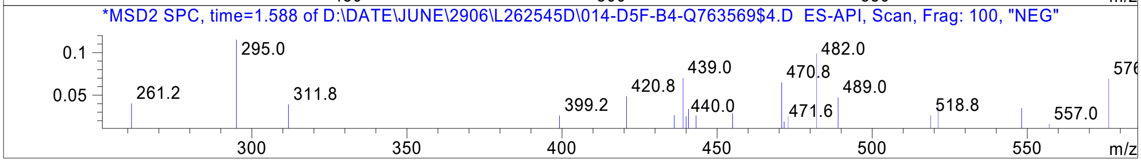




Z4497434898_

S993128$9

MaxPeak: 100.00%
Ret_Time: 1.350 min

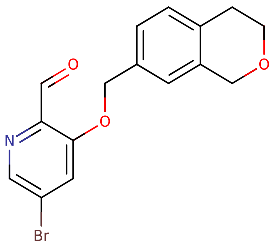


**Mol Wt** 348.19

**Exact Mass** 347.03

| # | Time | Area% |
|---|---|---|
| 1 | 1.350 | 100.00 |

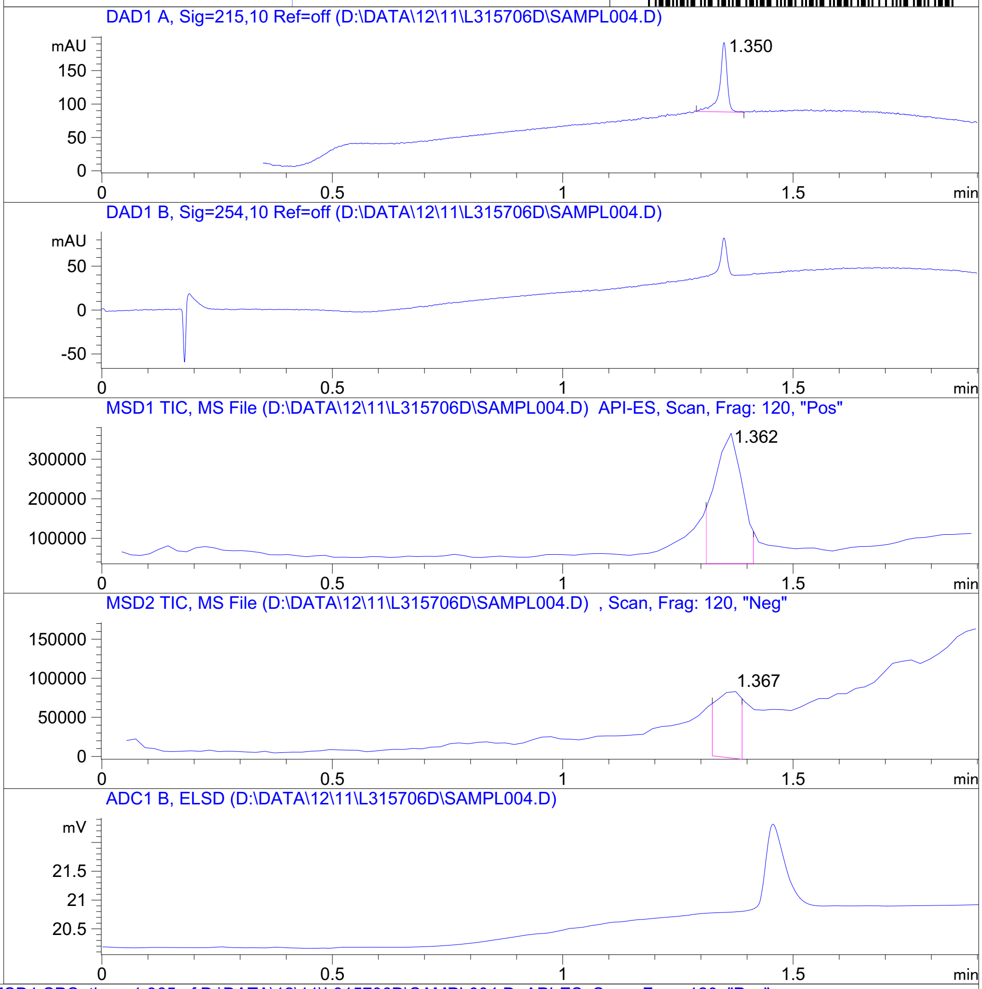


RT 1.362

RT 1.367

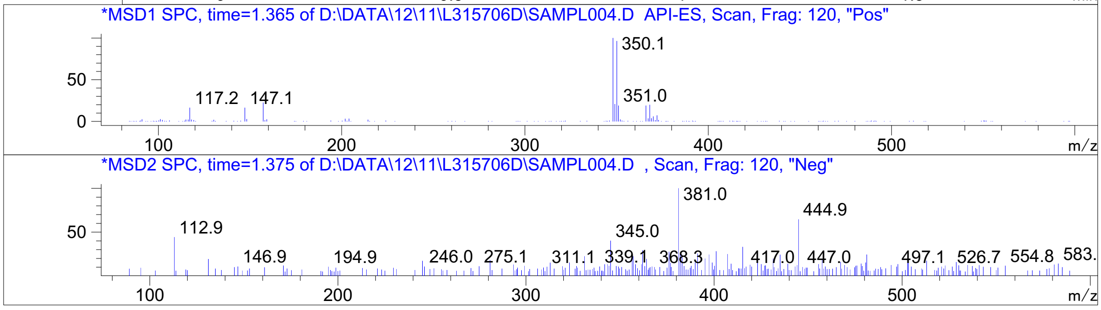


Inj.Date 12/11/2020 LB -SL- Acq. Method C:\HPCHEM\-> ->

Z3339728474_

MaxPeak: 100.00%
Ret_Time: 1.516 min

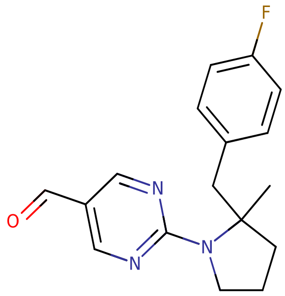


**Mol Wt** 299.34
**Exact Mass** 299.17

| # | Time | Area% |
|---|---|---|
| 1 | 1.516 | 100.00 |

Q763545$1

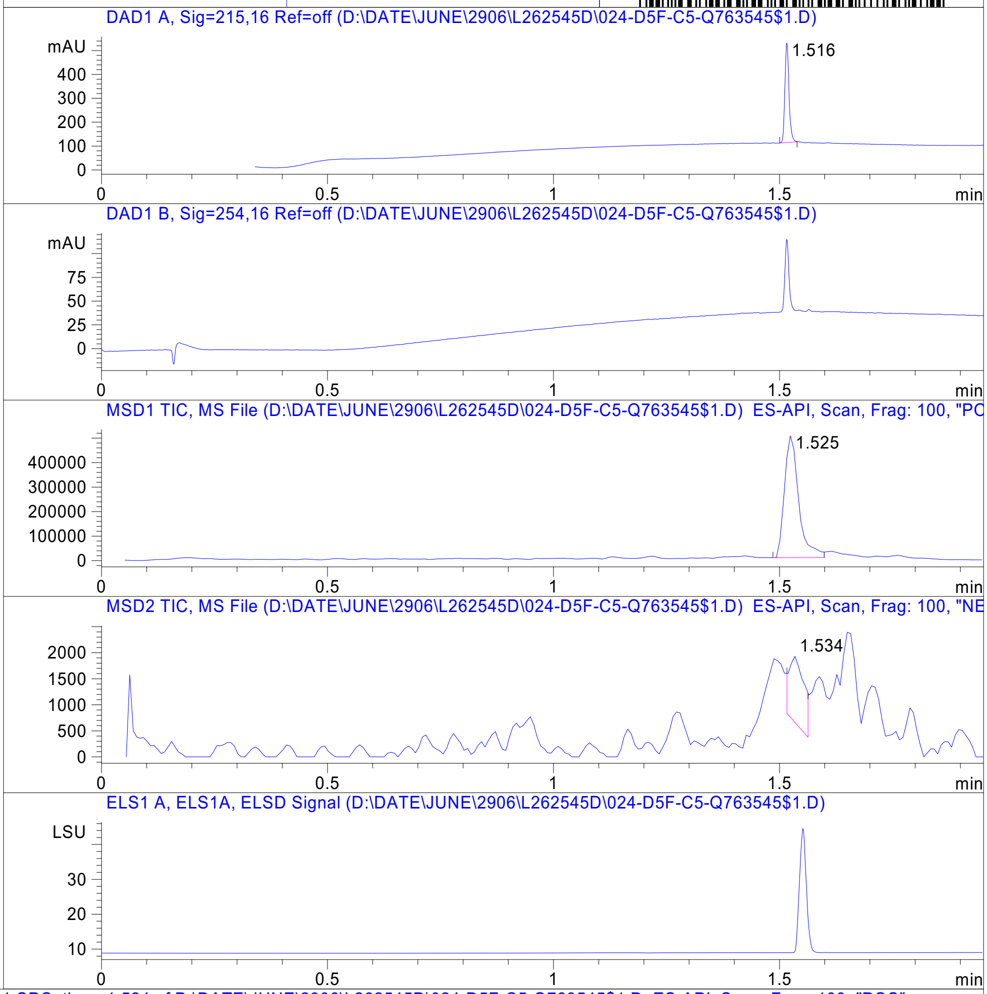


RT 1.525

RT 1.534

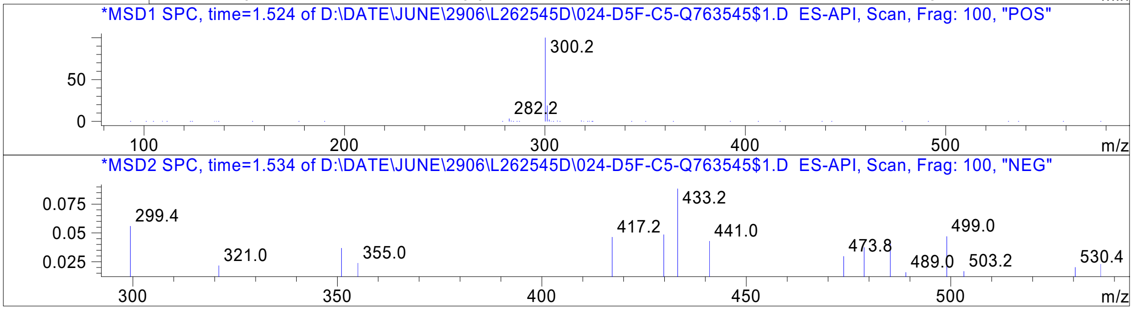




Z2851615440_

MaxPeak: 90.89%
Ret_Time: 1.145 min

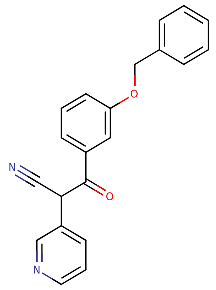

**Mol Wt** 328.36
**Exact Mass** 328.14

| # | Time | Area% |
|---|---|---|
| 1 | 1.145 | 90.89 |
| 2 | 1.276 | 9.11 |

R2541405

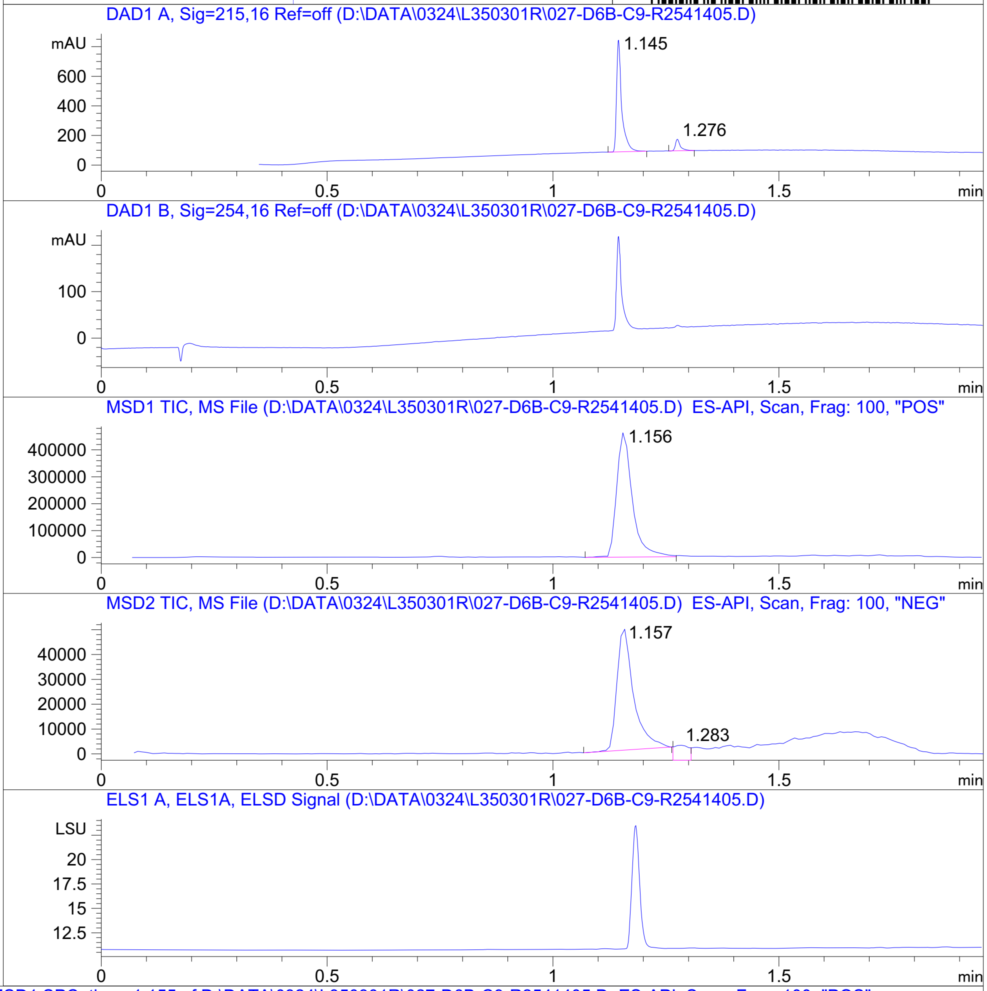


RT 1.156

RT 1.157

RT 1.283

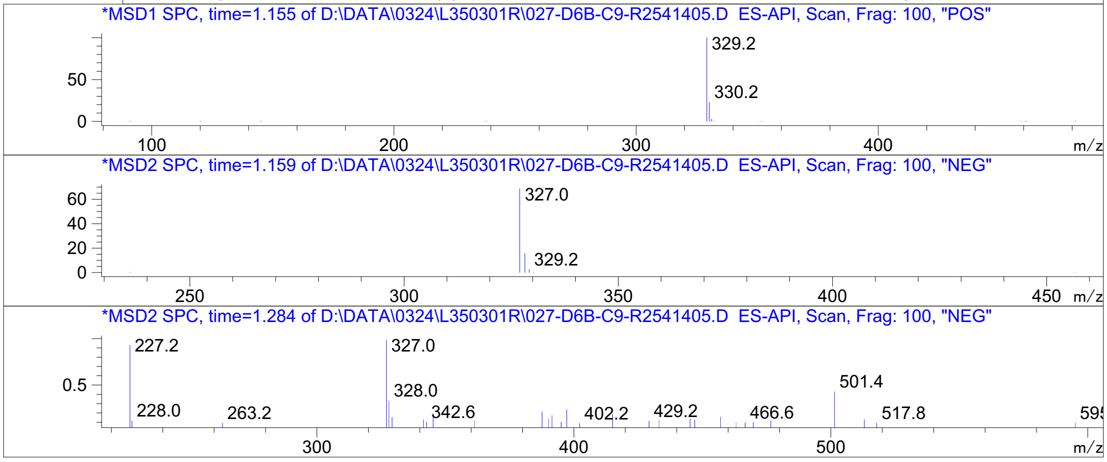


Inj.Date 3/24/2021 OA -14- Acq. Method C:\Chem32\-> ->

Z2168218606_

MaxPeak: 100.00%
Ret_Time: 0.876 min

R2538227

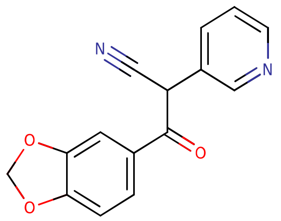


**Mol Wt 266.25**

**Exact Mass 266.07**

| # | Time | Area% |
|---|---|---|
| 1 | 0.876 | 100.00 |

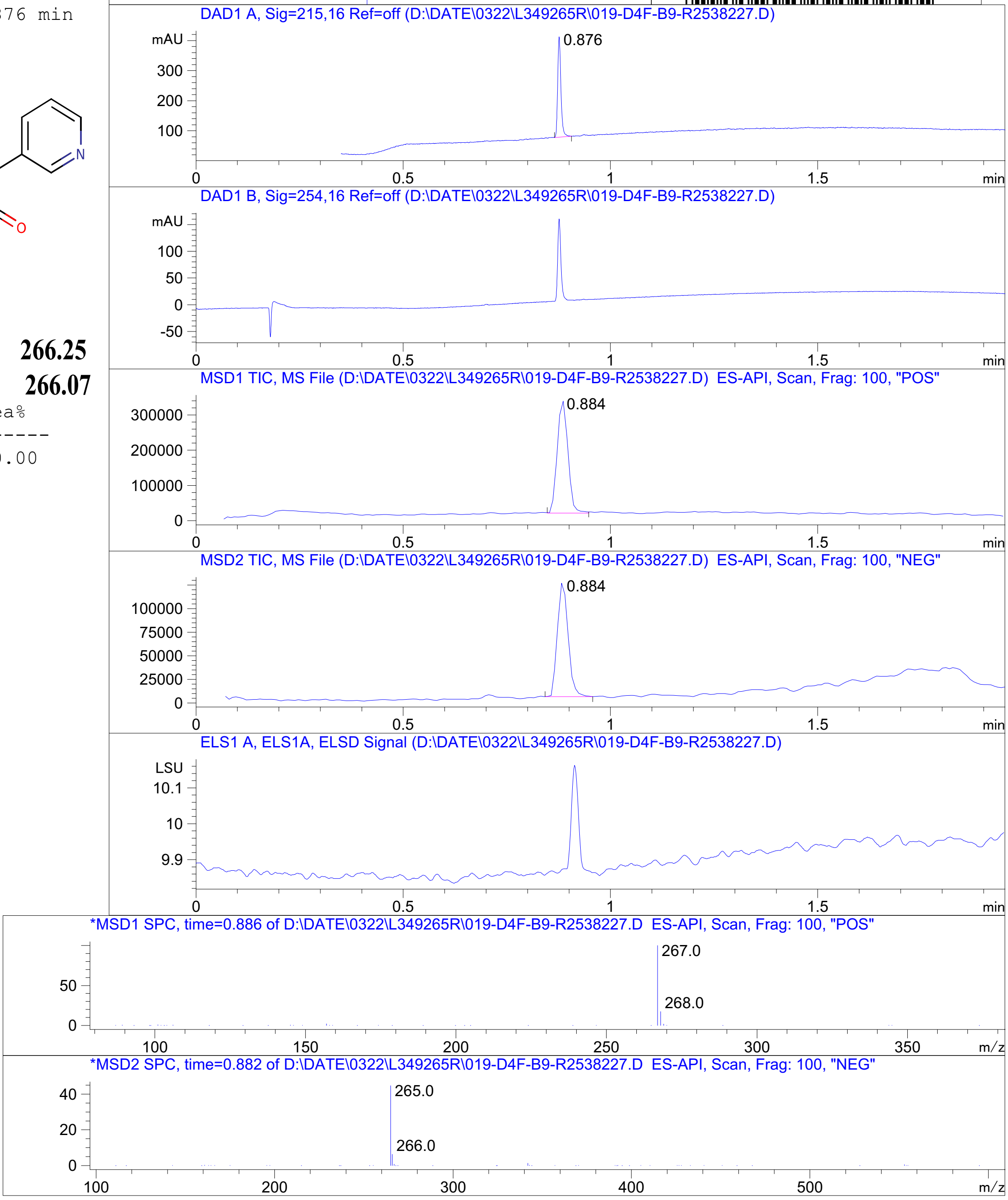


RT 0.884

RT 0.884

Inj.Date 3/22/2021 O -18- Acq. Method C:\Chem32\-> ->

Z1410957869_

MaxPeak: 100.00%
Ret_Time: 0.964 min

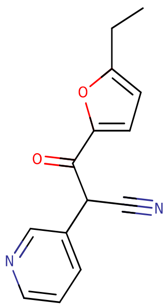

**Mol Wt 240.26**
**Exact Mass 240.1**

| # | Time | Area% |
|---|---|---|
| 1 | 0.964 | 100.00 |

R2532053

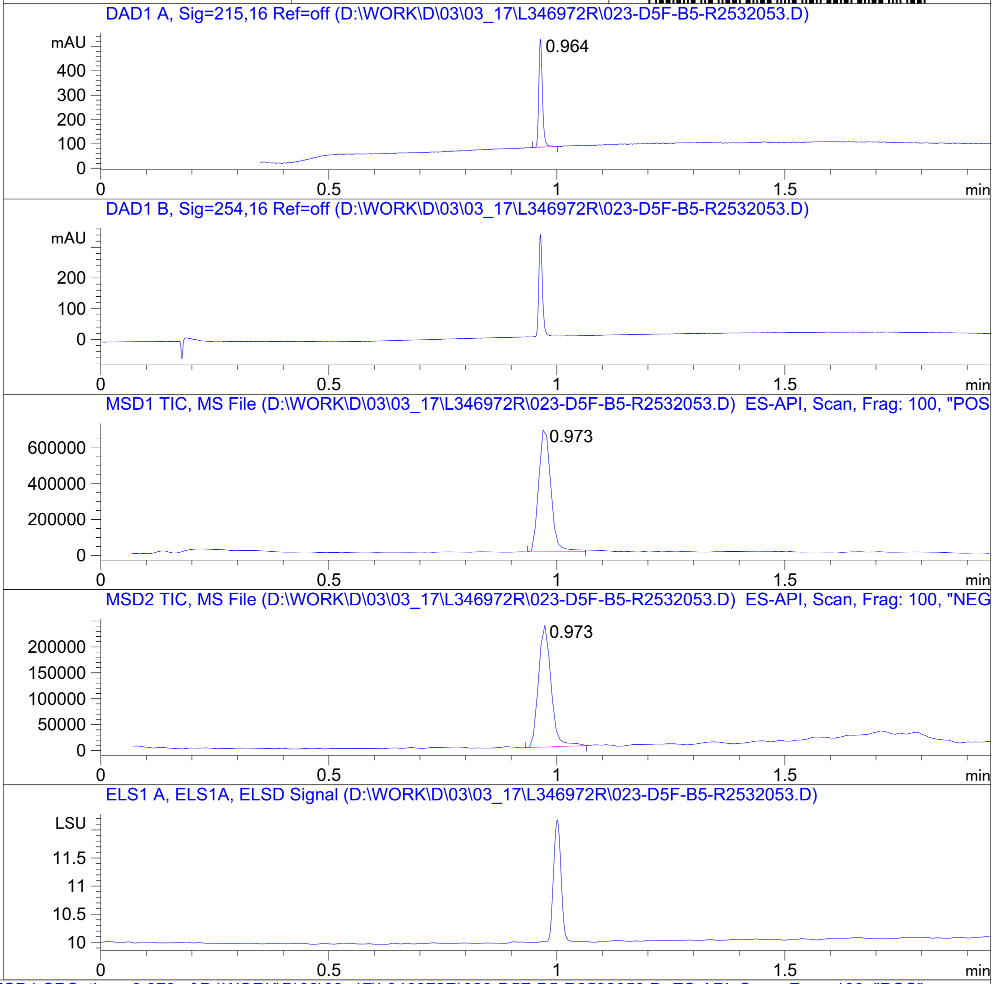


RT 0.973

RT 0.973

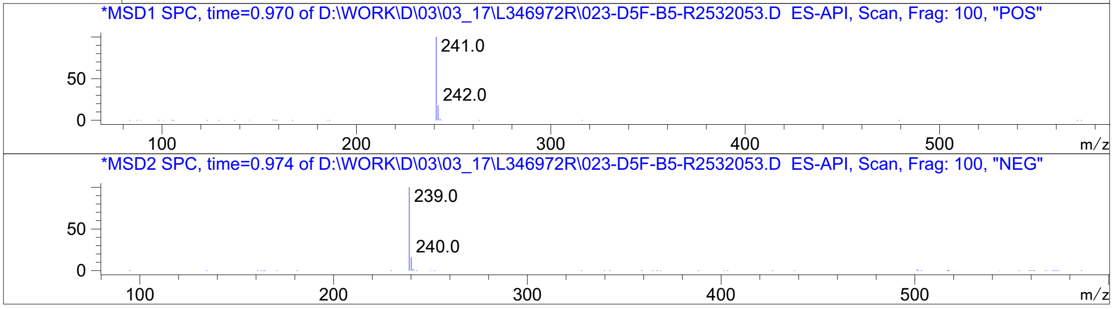


Inj.Date 3/16/2021 K -18- Acq. Method C:\Chem32\-> ->

Z1410957765_

Z1410957728_

Library Search Report

```
Data File : C:\MSDCHEM\1\DATA\03_23\R2539931.D          Vial: 16
Acq On    : 24 Mar 2021   4:20                      Operator:
Sample    : R2539931                                Inst    : GCMS2 #1
Misc      : CH3CN+CH2Cl2                            Multiplr: 1.00
                                               Sample Amount: 0.00
MS Integration Params: autoint1.e
Method   : C:\MSDCHEM\1\METHODS\UNIVERS.M (Chemstation Integrator)
Title    :
```

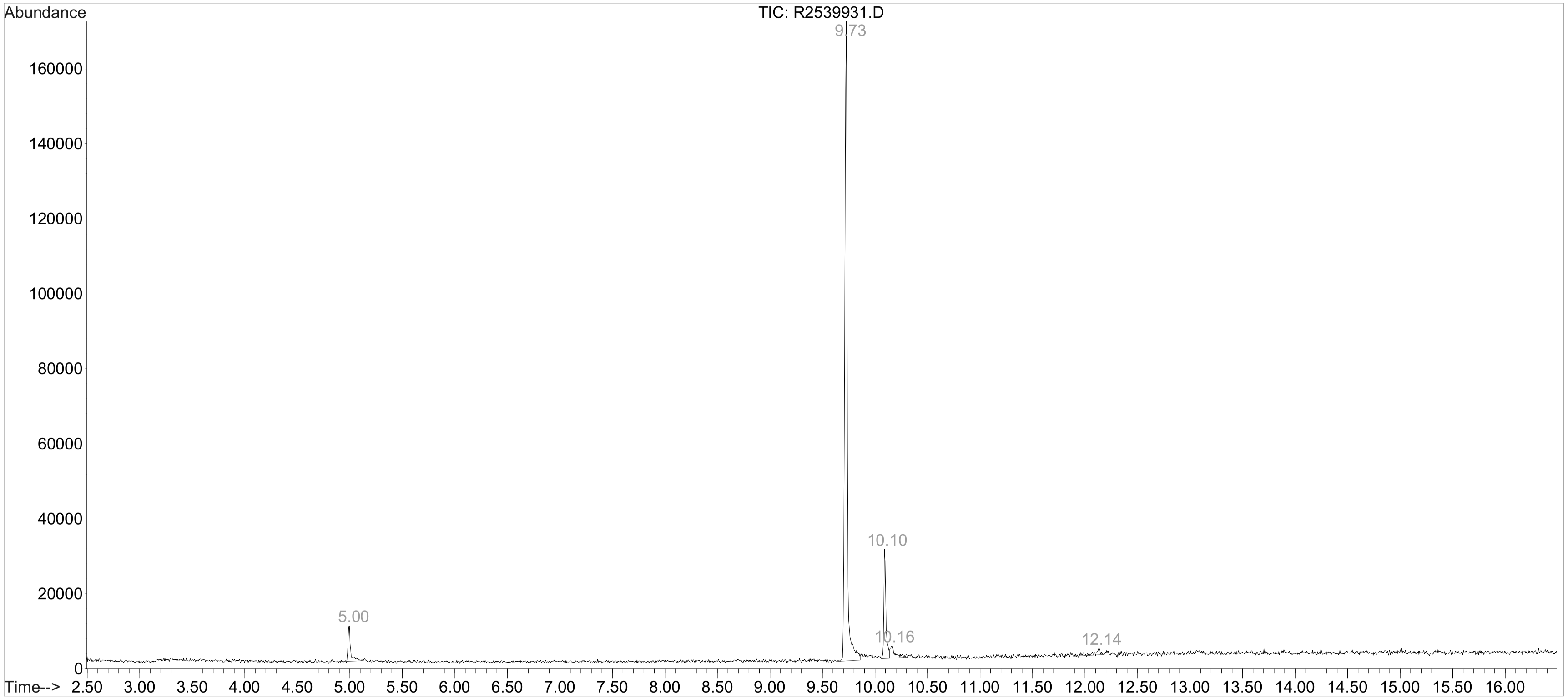

MaxPeak: 100.00%
Ret_Time: 0.926 min

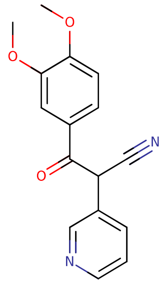

**Mol Wt** 282.29

**Exact Mass** 282.11

| # | Time | Area% |
|---|---|---|
| 1 | 0.926 | 100.00 |

R2535275

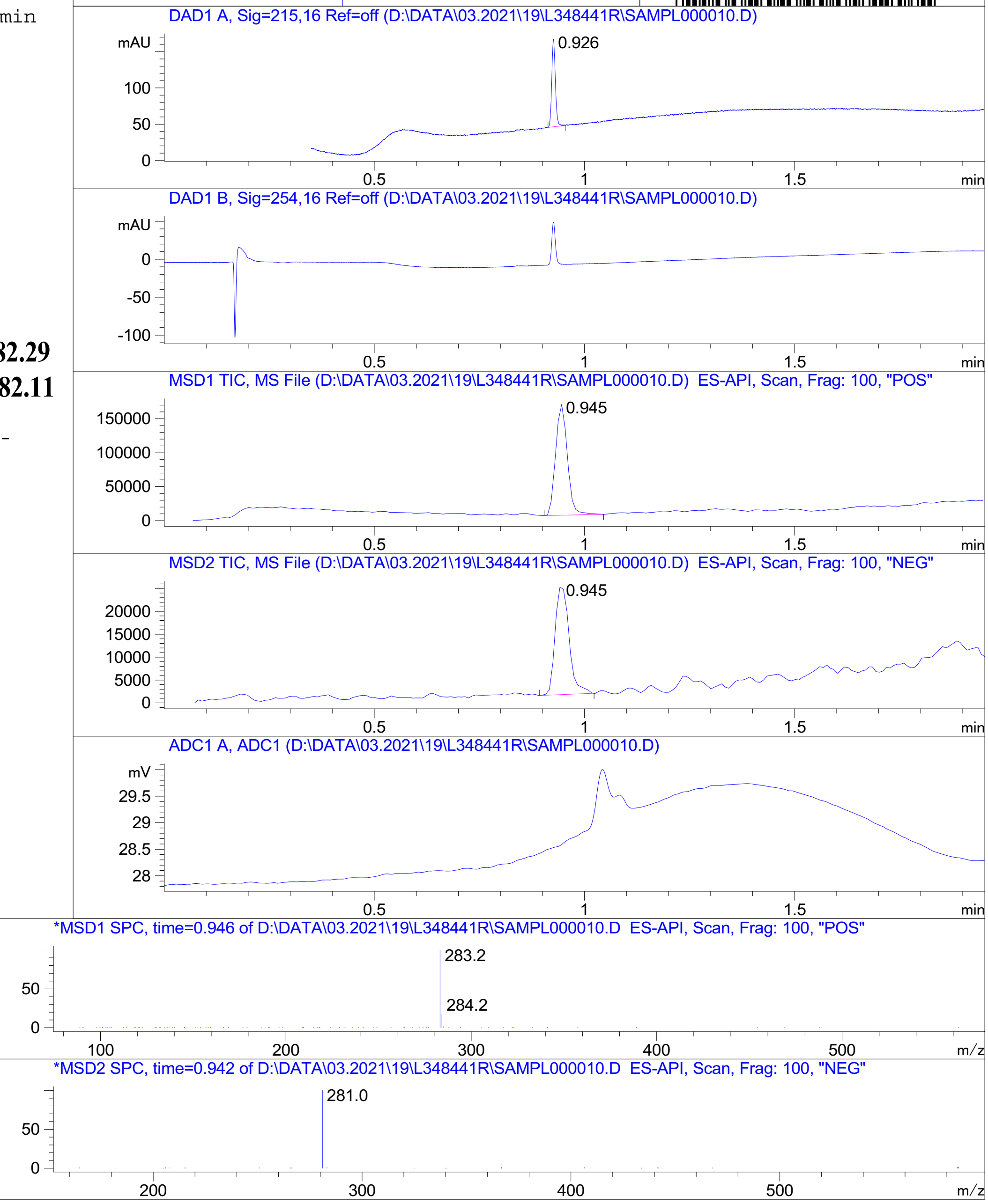


Inj.Date 3/19/2021 CH P2-B-02 - 4 - Acq. Method C:\CHEM32\-> ->

Z1410957725_

MaxPeak: 100.00%
Ret_Time: 1.542 min

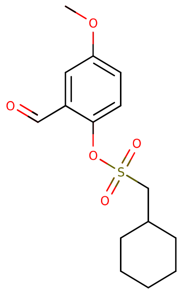


**Mol Wt** 312.38
**Exact Mass** 312.12

| # | Time | Area% |
|---|---|---|
| 1 | 1.542 | 100.00 |

Q786891$71

RT 1.555

RT 1.557

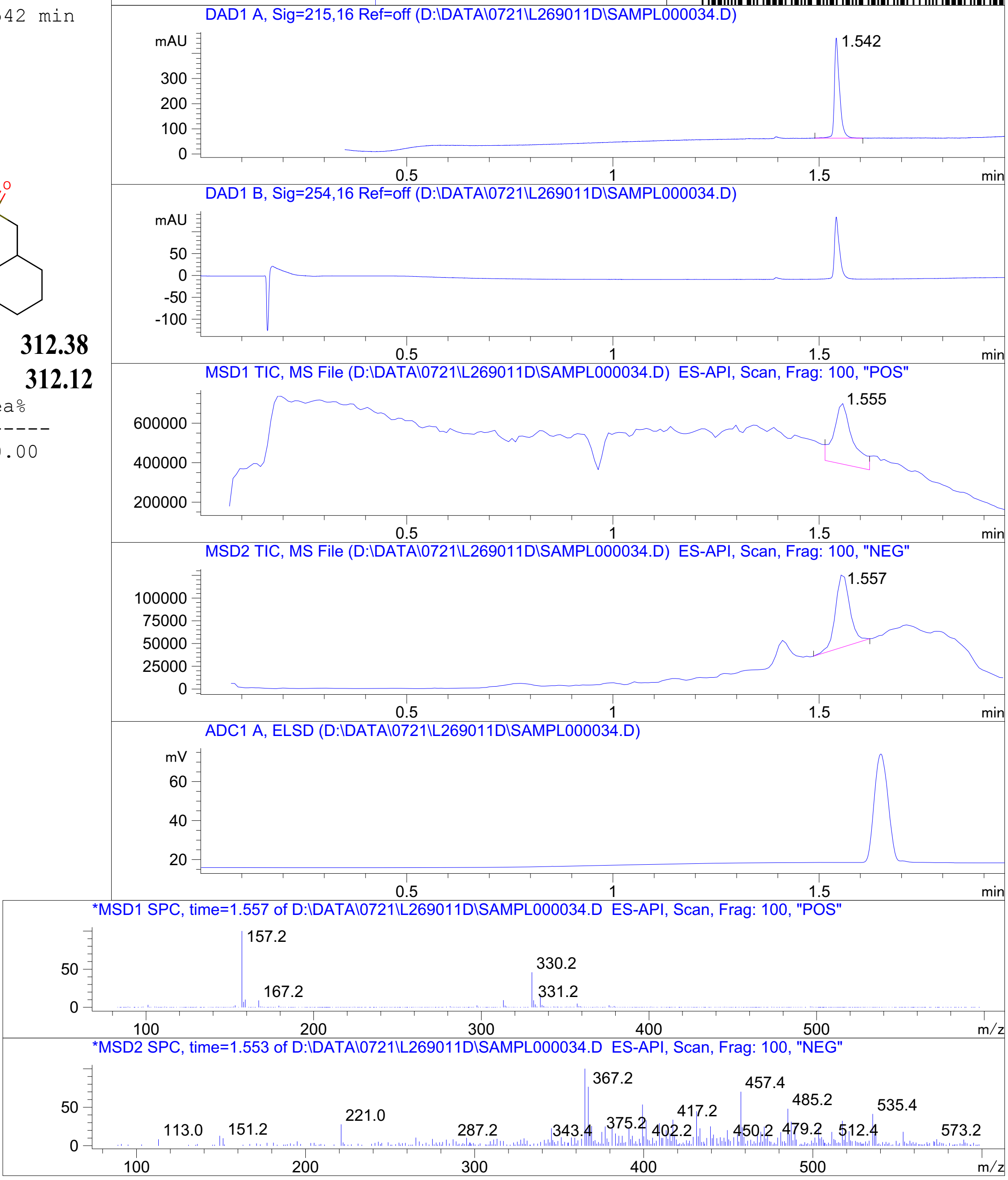




Z1218045697_

MaxPeak: 97.18%
Ret_Time: 1.417 min

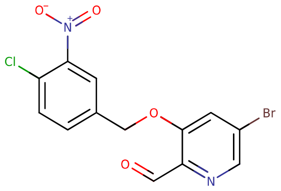


**Mol Wt** 371.57

**Exact Mass** 371.93

| # | Time | Area% |
|---|---|---|
| 1 | 1.417 | 97.18 |
| 2 | 1.463 | 2.82 |

V498758$I

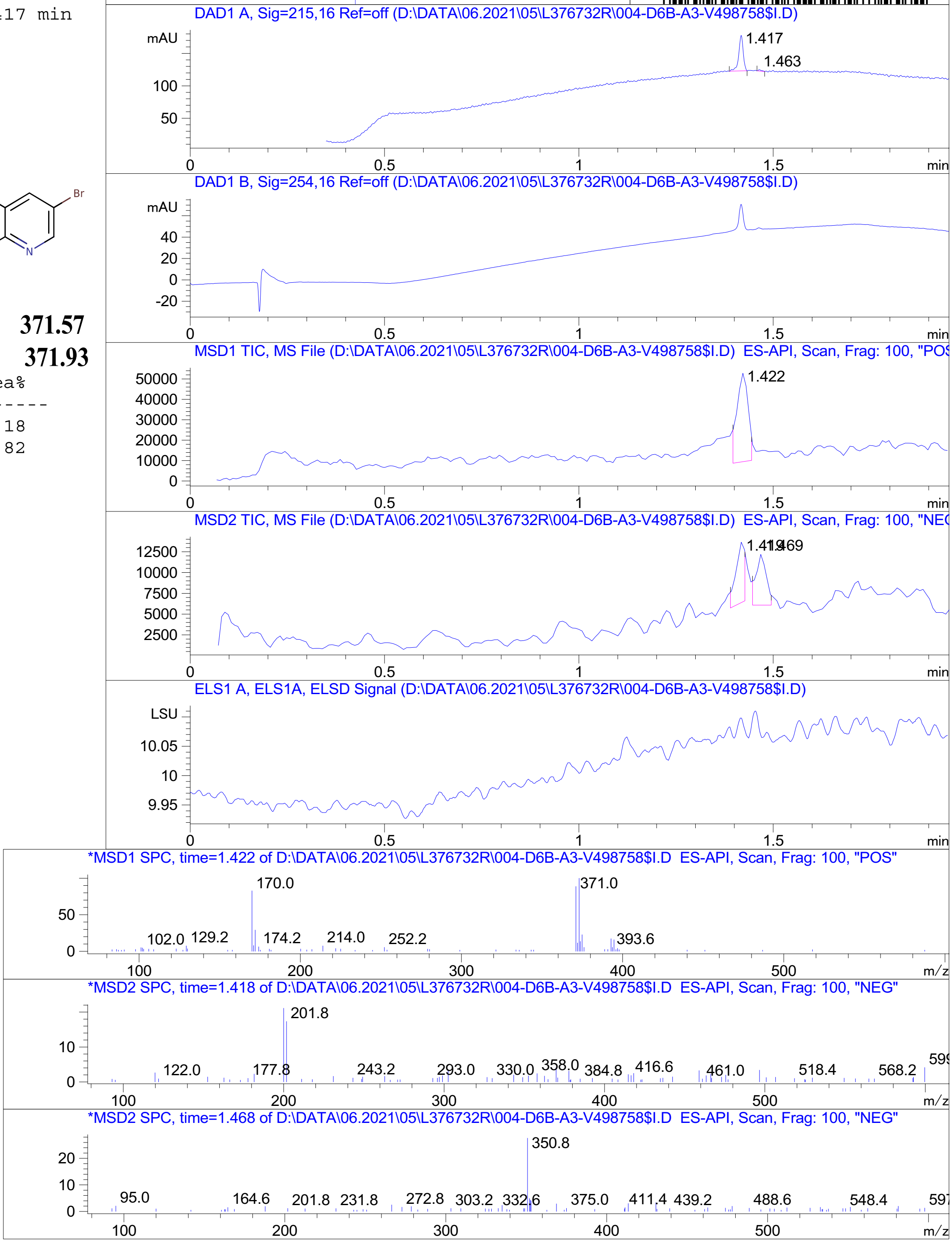


RT 1.422

RT 1.419

RT 1.469

Inj.Date 6/5/2021 CH <invalid> -17- Acq. Method C:\Chem32\-> ->

Z4767502618_

R2794156

MaxPeak: 100.00%
Ret_Time: 1.213 min

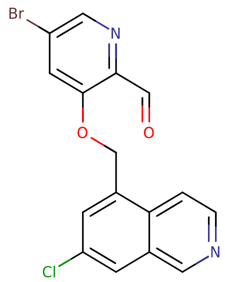


**Mol Wt** 377.62
**Exact Mass** 377.97

| # | Time | Area% |
| --- | --- | --- |
| 1 | 1.213 | 100.00 |

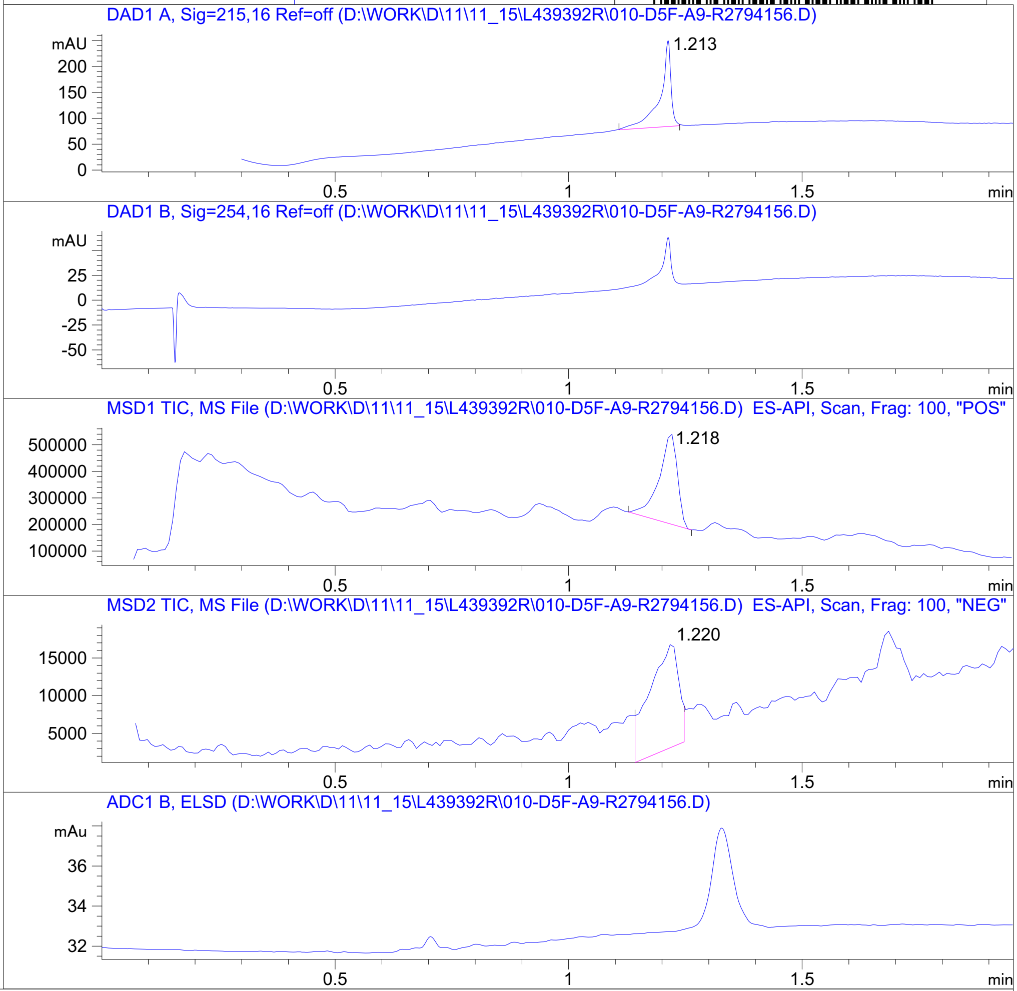


RT 1.218

RT 1.220

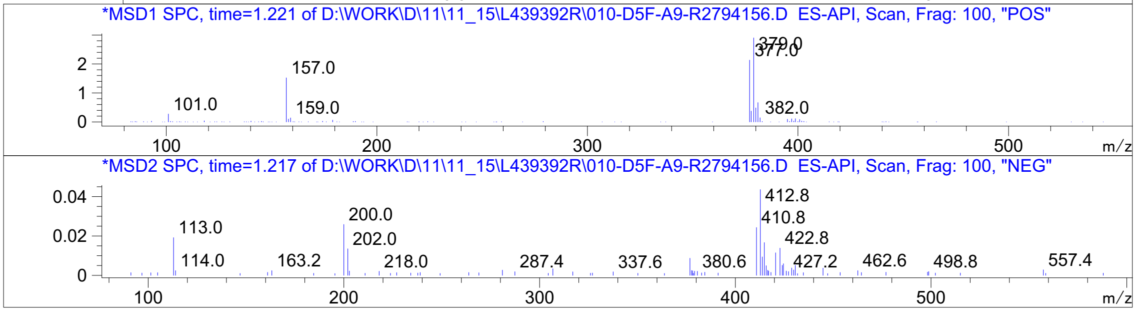


Inj.Date 11/15/2021 K <invalid> -10- Acq. Method C:\Chem32\-> ->

Z5192779138_

MaxPeak: 100.00%
Ret_Time: 1.035 min

# Q227942$1

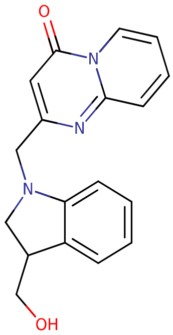

**Mol Wt** 307.35
**Exact Mass** 307.15

```
#   Time    Area%
------------------
1   1.035   100.00
```

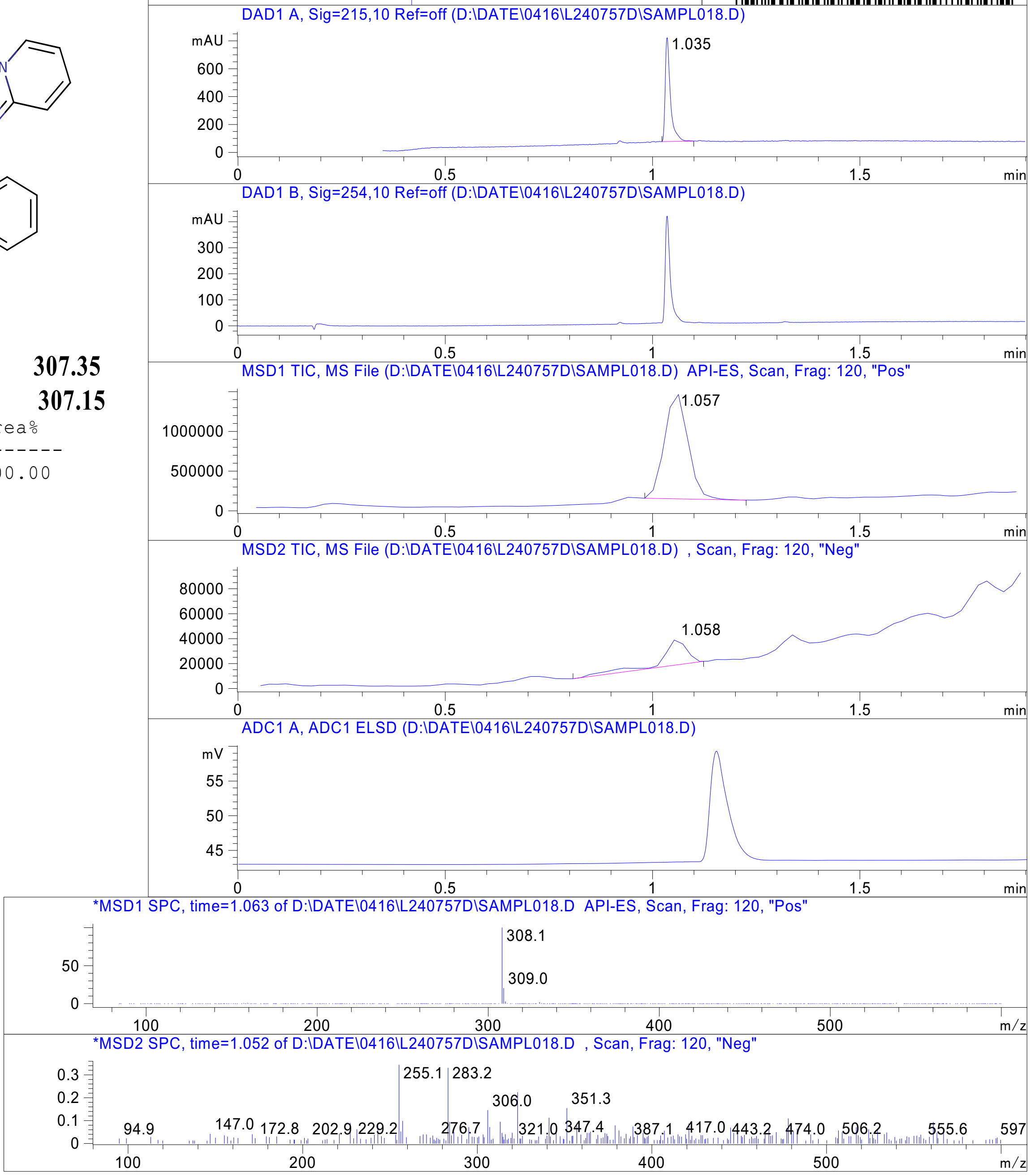


RT 1.057

RT 1.058

Inj.Date 4/16/2020 E -VL- Acq. Method C:\HPCHEM\-> ->

Z2215970681_

MaxPeak: 100.00%
Ret_Time: 1.375 min

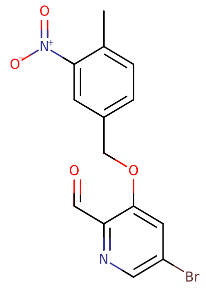


**Mol Wt** 351.15
**Exact Mass** 349.99

| # | Time | Area% |
|---|---|---|
| 1 | 1.375 | 100.00 |

# U920110$2

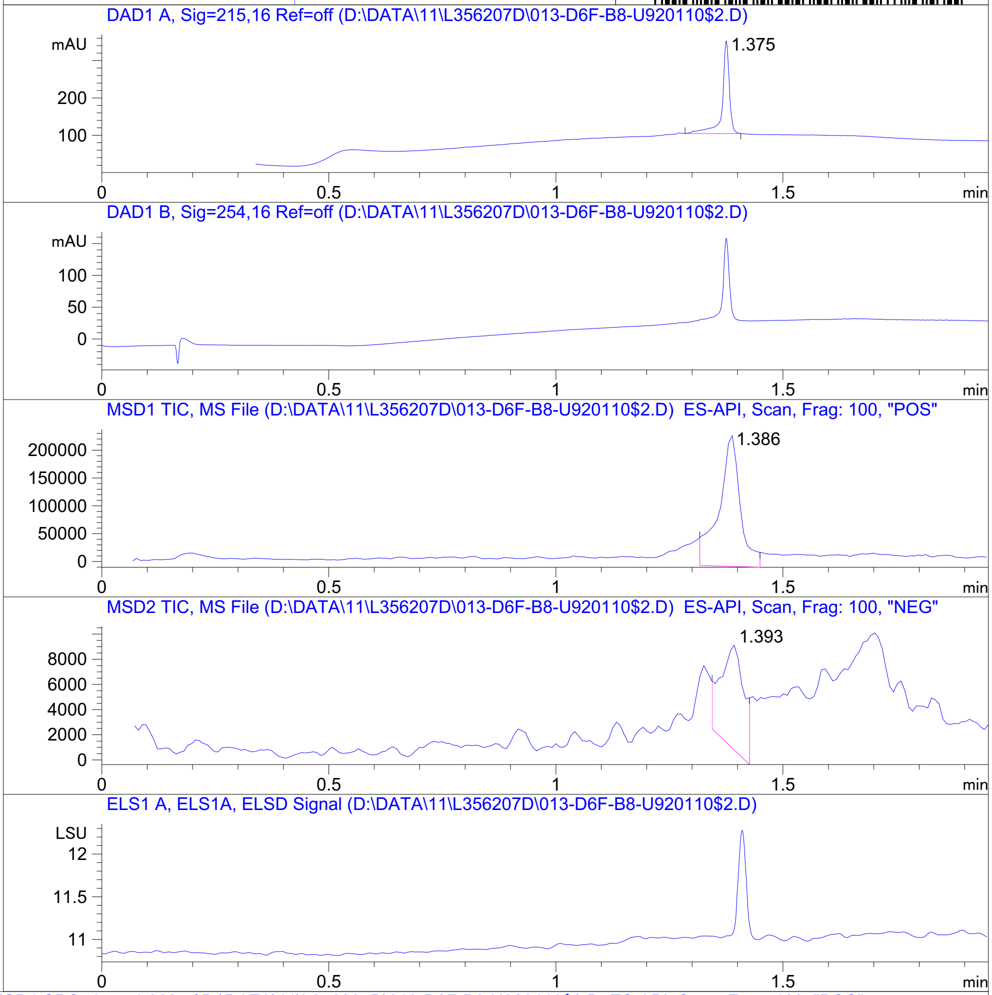


RT 1.386

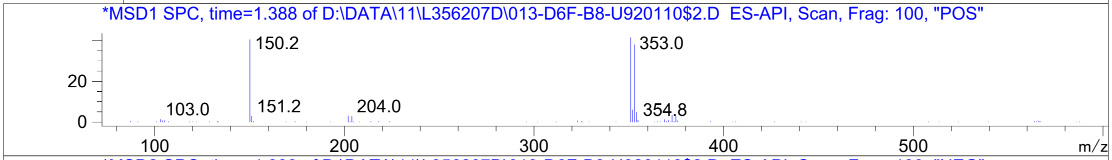


RT 1.393

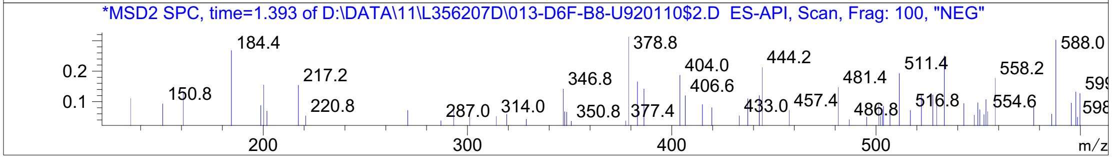


Inj.Date 4/10/2021 LB -16- Acq. Method C:\Chem32\-> ->

Z4954569218_

MaxPeak: 100.00%
Ret_Time: 1.494 min

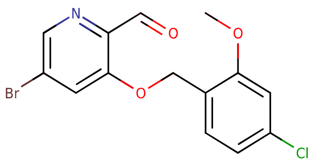

**Mol Wt** 356.6
**Exact Mass** 356.97

| # | Time | Area% |
|---|---|---|
| 1 | 1.494 | 100.00 |

# U920103$6

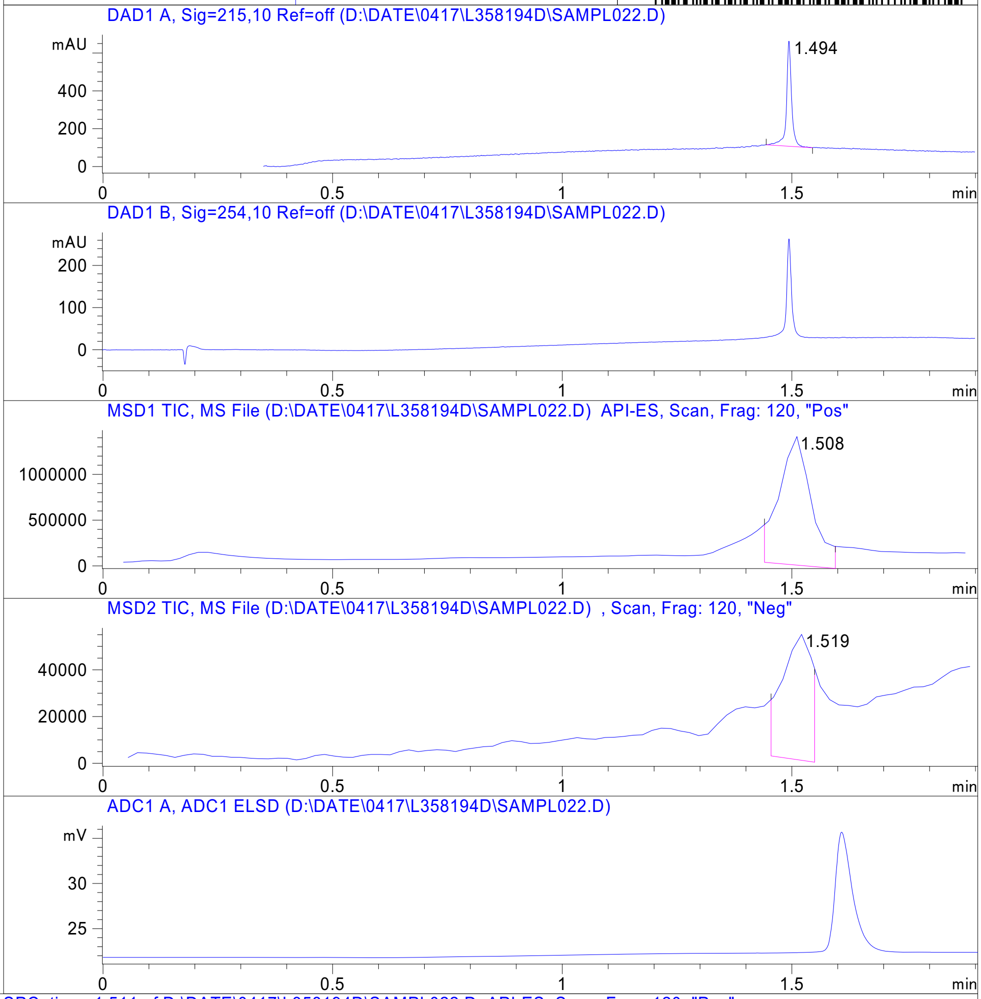


RT 1.508

RT 1.519

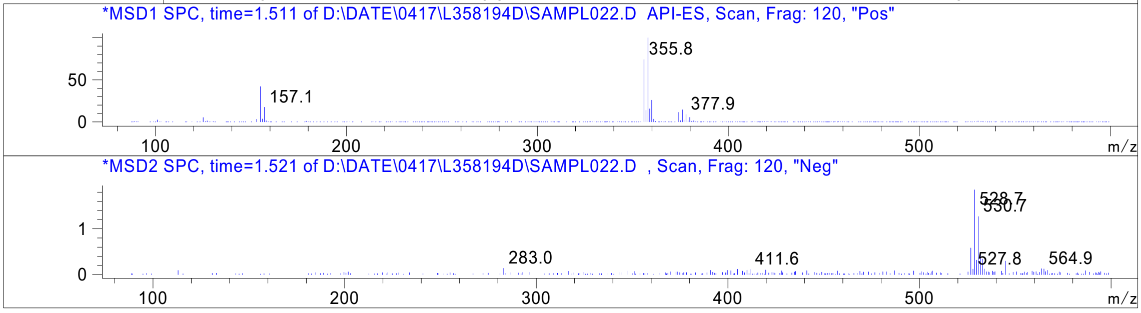


Inj.Date 4/17/2021 E -VL- Acq. Method C:\HPCHEM\-> ->

Z4954569127_

MaxPeak: 100.00%
Ret_Time: 1.415 min

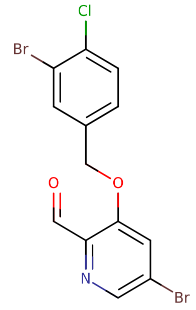


**Mol Wt** 405.47
**Exact Mass** 404.87

| # | Time | Area% |
|---|---|---|
| 1 | 1.415 | 100.00 |

# U920100$5

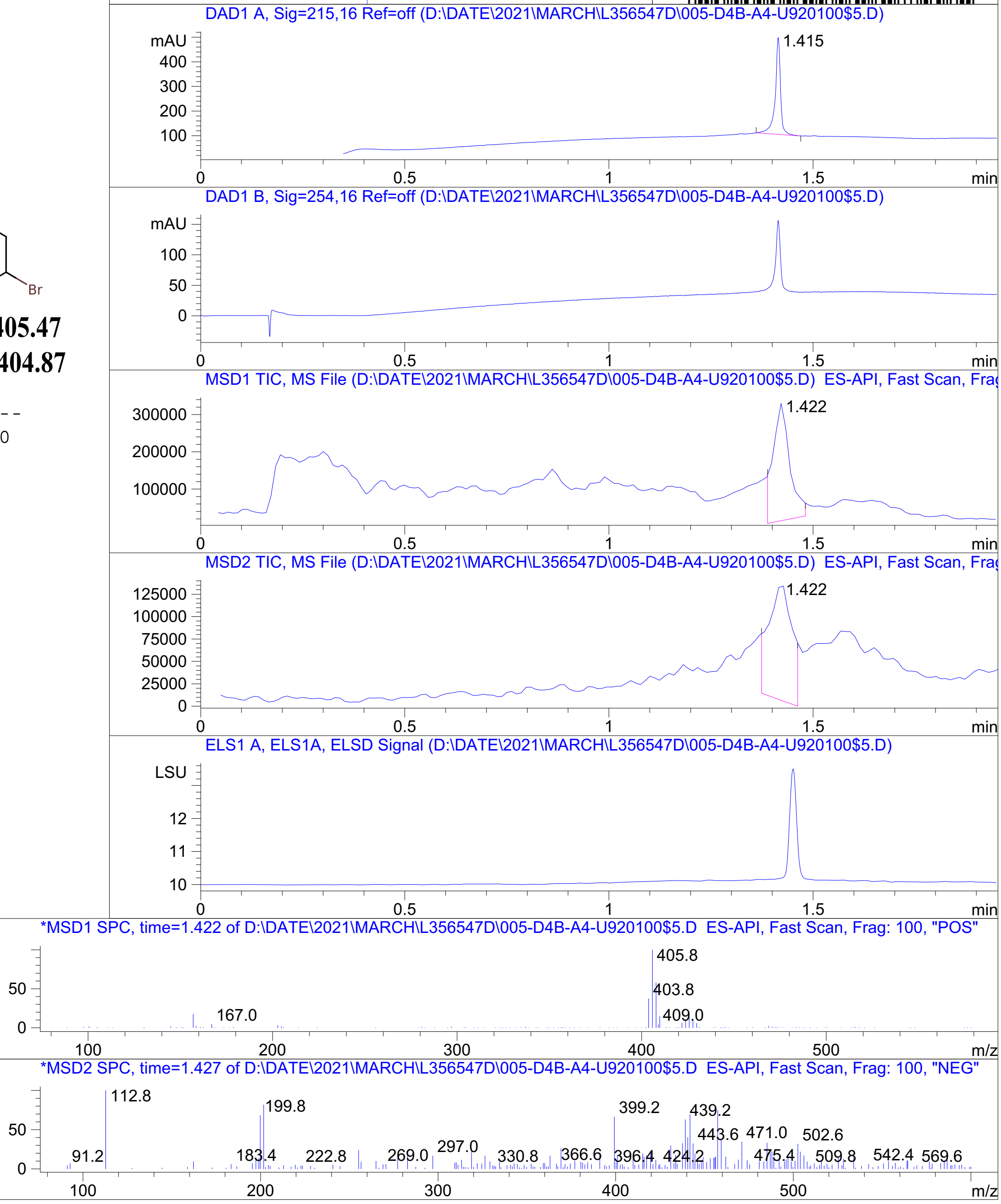


RT 1.422

RT 1.422

Inj.Date 4/11/2021 T <invalid> Acq. Method C:\Users\ -> ->

Z4954569121_

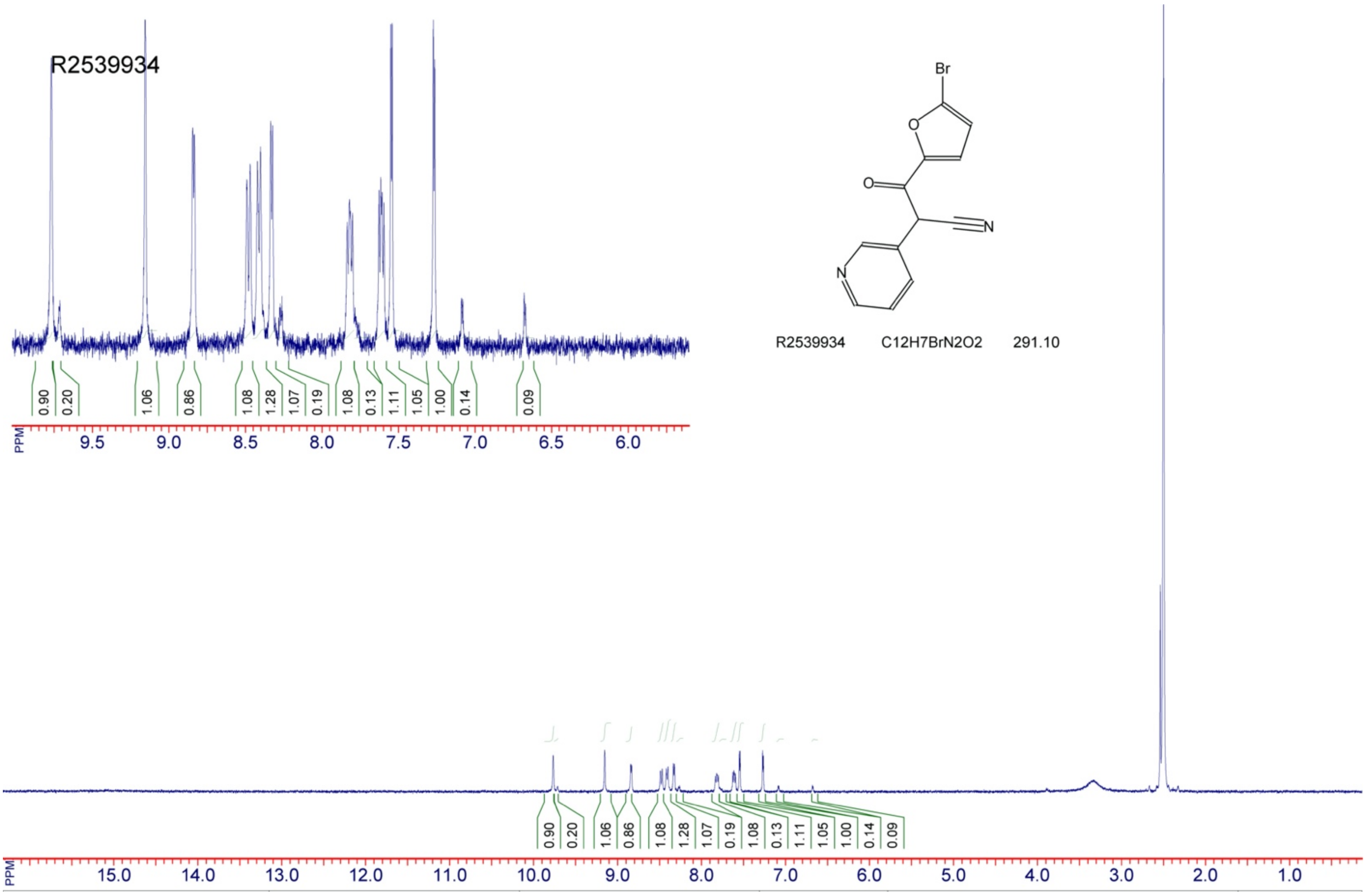


Z1410957817_

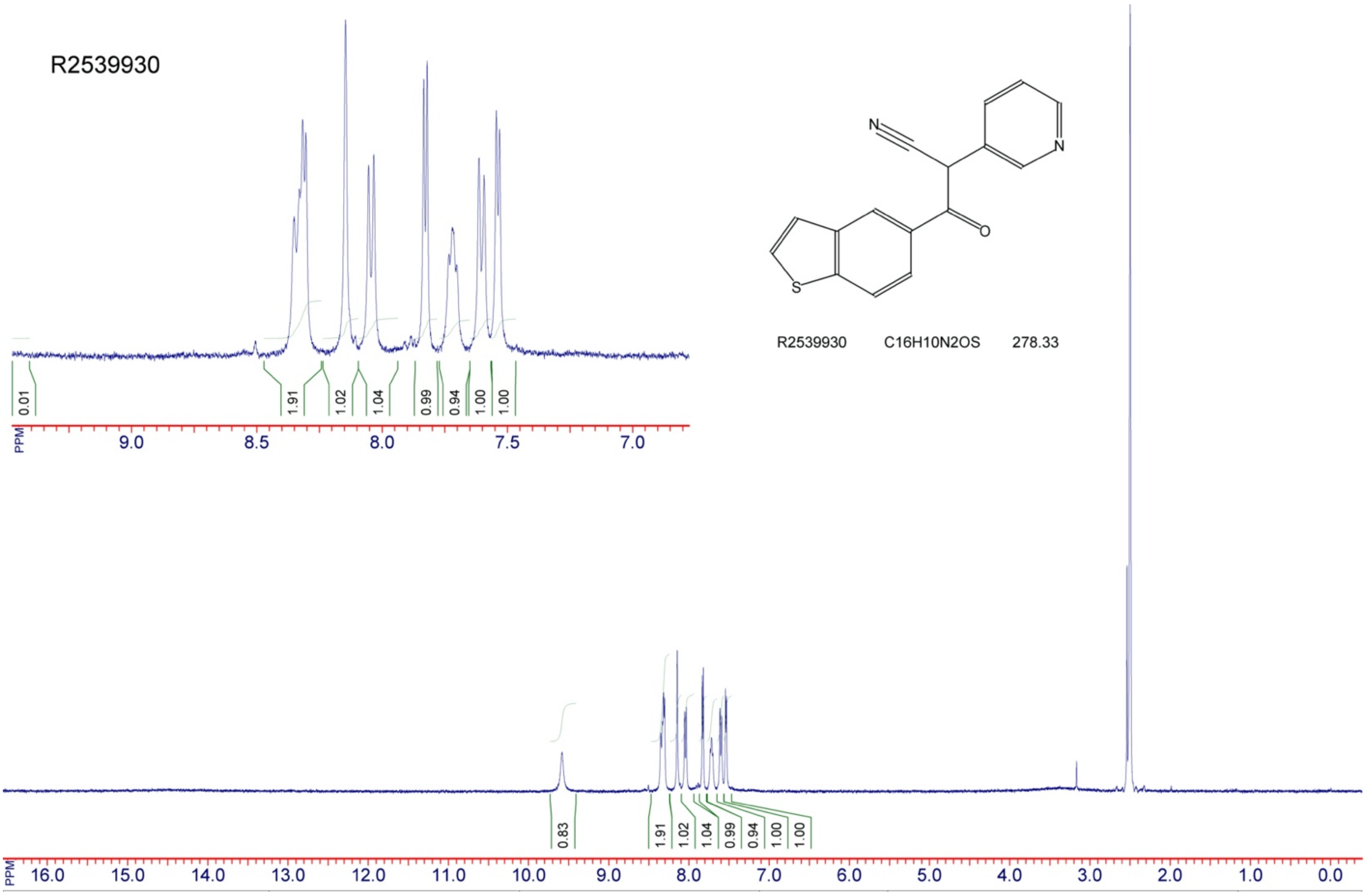


Z3302062818_
